# Extensive monolayer formation after transplantation depends on a RPE subpopulation derived from human iPSCs

**DOI:** 10.1101/2024.10.15.618147

**Authors:** Karen Tessmer, Sylvia J. Gasparini, Klara Schmidtke, Juliane Hammer, Trishla Adhikari, Lea Michalke, Anne Schön, Liliana R. Loureiro, Annika Last, Nadine Chelius, Annett Kurtz, Hannah Johannsen, Madalena Carido, Susanne Ferguson, Andreas Petzold, Ulrike A. Friedrich, Thomas Kurth, Andreas Dahl, Anja Feldmann, Seba Almedawar, Sebastian Knöbel, Marius Ader

## Abstract

Loss of the retinal pigment epithelium (RPE) in the eye leads to photoreceptor dysfunction and death causing eventually vision loss. Cell replacement strategies using RPE cells derived *in vitro* from pluripotent stem cells (PSCs) are currently evaluated as a potential therapeutic strategy. Generation of polarized monolayers represents an essential prerequisite for proper RPE function, however, monolayer formation following transplantation of RPE cell suspensions has not been systematically assessed. Using the sodium iodate mouse model of acute RPE depletion, significant increase in monolayer formation capacity of passage (P) 1 vs. P2 human iPSC-derived RPE cells was observed three weeks after transplantation. Transplant-derived monolayers showed characteristic apicobasal polarity, RPE marker expression, phagocytosis function, and preservation of the host outer nuclear layer. The cell surface marker panel CD54^+^/PSA-NCAM^-^ was identified to enrich for an RPE subpopulation with high potential for monolayer formation following transplantation. Results underline the importance of defining and isolating competent cell subpopulations for successful RPE transplantation.

## Introduction

Deterioration of the retinal pigment epithelium (RPE), an epithelial monolayer separating choroid and neuroretina in the eye, causes retinal degeneration including dysfunction and loss of the light sensitive photoreceptors. Diseases comprising RPE pathologies include subforms of inherited retinal degenerative diseases (IRD) like retinitis pigmentosa as well as age related macular degeneration (AMD), the main cause for vision impairment and blindness in industrialized societies ^1^. Loss of RPE cells is considered permanent given limitations of intrinsic regeneration in the mammalian retina, while cell transplantation approaches for the replacement of RPE are discussed as a potential therapeutic approach. Indeed, preclinical and clinical phase I/II studies using human pluripotent stem cell (PSC)- or RPE stem cell-derived RPE cells provide evidence for safety of the approach, long-term survival of donor cells and some signs of vision stabilization or even improvements^2,3^.

Two principal delivery approaches, both in most cases targeting the subretinal space, have mainly been used: injection of either an RPE cell suspension or a preformed RPE sheet or patch (i.e. RPE cultured on a scaffold). While sheets or patches have the advantage to resemble an already formed polarized monolayer, their delivery into the diseased eye is associated with major obstacles, requiring sophisticated surgery techniques with increasing danger of ruptures in significantly thinned and gliotic degenerative retinas ^3^. Cell suspensions, on the other hand, are more easily injected into the subretinal space, but still have to form a polarized monolayer on Bruch’s membrane in vivo. A recent direct comparison of suspension vs. patch transplantations provided evidence for improved survival and functionality of preformed patches in an acute laser-induced porcine-model of RPE damage ^4^. On the other hand, several clinical RPE replacement trials, with two moving forward to phase 2a, are based on suspension transplantation showing evidence for vision stabilization and improvement in AMD/geographic atrophy patients ^2,3^.

While PSC-derived RPE cells show significant changes in gene expression profiles and morphology over multiple passages leading eventually to de-differentiation and epithelial-to-mesenchymal transitions ^5^, recent single cell transcriptome analysis of stem cell-derived RPE in vitro provided evidence for a high heterogeneity within RPE cells also at low passages ^6,7^. Interestingly, a subpopulation of RPE cells derived from in vitro expanded RPE stem cells (RPESCs) showed improved transplantation efficacy in the pre-clinical RCS rat model when used from an early passage ^8^ and an RPE subpopulation expressing the lncRNA TREX displayed improved integration capacity in an existing RPE monolayer in vitro ^6^. However, a systematic and quantitative analysis for the competence of specific PSC-derived RPE subpopulations for monolayer formation in vivo has not been performed yet.

Using the sodium iodate (NaIO_3_) RPE depletion mouse model as recipient ^9,10^, we provide evidence that iPSC-RPE isolated from passage (P) 1 generate extensive polarized RPE monolayers within three weeks after suspension transplantation covering up to 35% of the host retinal surface with protection of the outer nuclear layer (ONL) structure. Contrarily, P2 RPE cells showed a reduced coverage in recipient mouse retinas, and their capacity to form polarized monolayers was significantly decreased, resulting in a disrupted ONL, choroid swelling and increased infiltration of immunogenic cells. Interestingly, enriching for a P1 RPE subpopulation that can be isolated by cell surface epitope sorting (CD54^+^/PSA-NCAM^-^) consistently results in higher monolayer percentages than usage of unsorted P1 RPE cells (up to 80% of transplant area). Our findings thus underline the importance of defining and isolating competent RPE subpopulations for successful RPE transplantation approaches.

## Results

### While morphologically similar, expression profiles of human iPSC-derived RPE differ after passaging

RPE cells were generated by differentiation of iPSC with initial epithelial cyst formation in Matrigel followed by polarized monolayer formation on cell culture inlets and repeated passaging (Fig. 1A) ^11–13^. Both, P1 and P2 RPE cells acquired pigmentation and a cobblestone-like morphology, with zonula occludens 1 (ZO1)-positive tight junctions and actin enrichment at the apical surface (Figs. 1B and C). Proper polarization was further evident in the cross-sectional view by basal and apical positioning of nuclei and pigment granules, respectively, as well as basal deposition of collagenIV (COL4), and apical enrichment of sodium/potassium-transporting ATPase (ATP1A1) and F-actin (Fig. 1C). Ultrastructural analysis confirmed the apicobasal polarization of the cells with basally located nucleus, basal labyrinth and mitochondria, apically located microvilli and melanosomes, and connections via tight and adherens junctions (Fig. 1D).

**Figure 1:**
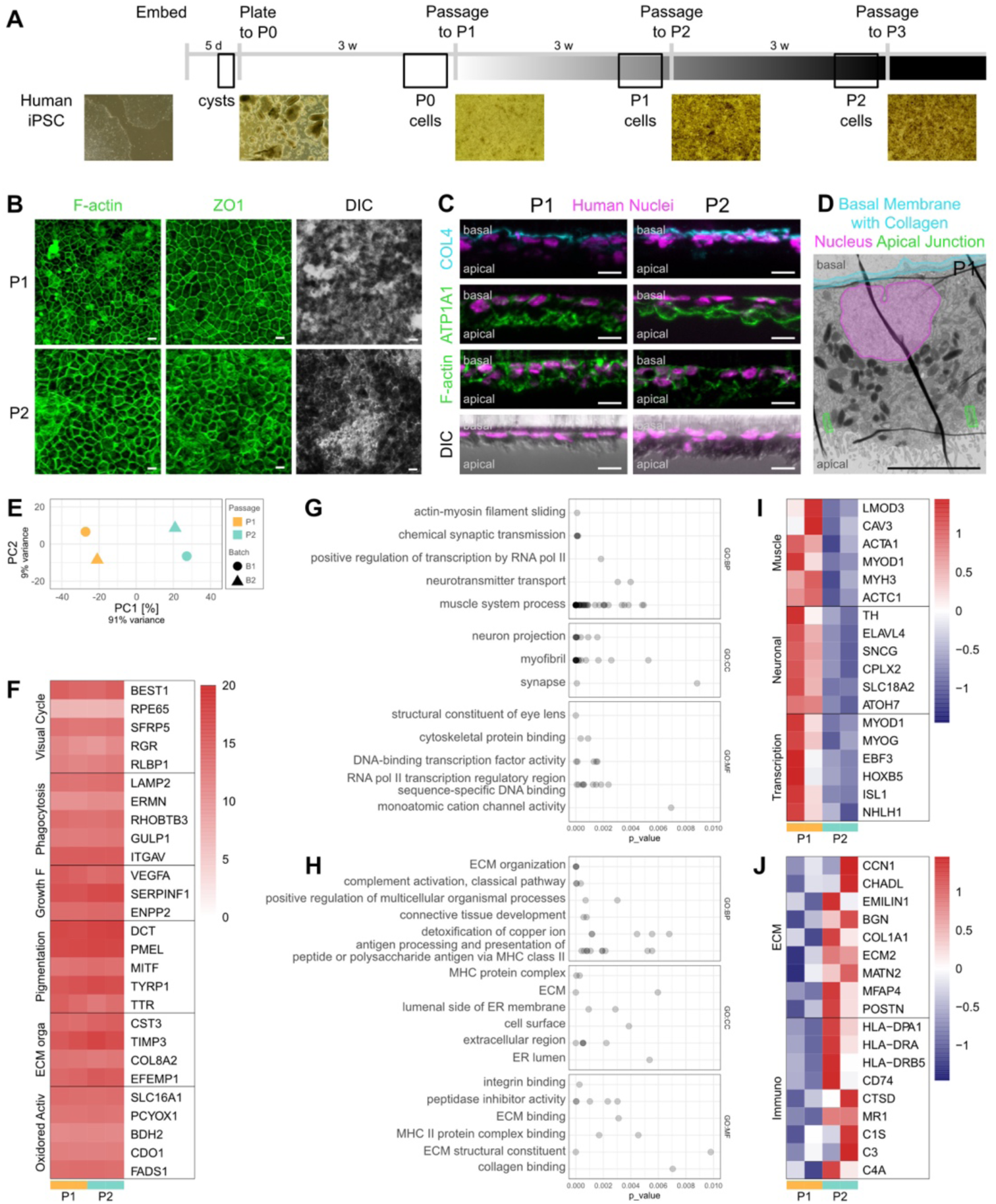
Characterization of P1 and P2 iPSC-derived RPE cells. (A) Scheme for the differentiation of iPSCs into RPE cells and representative images from different stages. In brief, iPSCs formed cysts in Matrigel within 5 days (d), followed by plating in RPE induction medium for 3 weeks (w). Passaging to cell culture inlets and culture for 3 w yielded passage (P) 1 cells and, after an additional passage and another 3 w, P2 cells. (B) P1 and P2 RPE cells form monolayers with cobblestone morphology visualized by immuno-fluorescence staining for F-actin or zonula occludens 1 (ZO1). P1 and P2 RPE cells are highly pigmented. n = 2 biological and 3 technical replicates. (C) Sections of P1 and P2 RPE monolayers show positive immunofluorescence staining for Human Nuclei, basally localized collagen IV (COL4), and apically localized ATPase (ATP1A1) and F-actin. Combined fluorescence staining (Human Nuclei) with DIC imaging reveals basal localization of RPE nuclei. n = 2 biological and 3 technical replicates. (D) Ultrastructure of differentiated P1 RPE cell showing features characteristic of polarized RPE monolayer features including a basal membrane with collagen (marked blue), a basally located nucleus (marked magenta), apically located pigmented melanosomes, apical microvilli, and tight/adherens junction (marked green). n = 2 biological and 3 technical replicates. (E) Principal component (PC) analysis after batch correction reveals close clustering of same passage number cells across cell culture batches (each symbol represents three averaged technical replicates of the same biological replicate). (F) Heatmap showing expression levels of RPE-related genes obtained by bulk RNAseq analysis from two batches each of P1 and P2 RPE cells. Values shown are log_2_ (normalized counts+1). (G, H) Gene ontology (GO) term analysis of differentially expressed genes shows GO term groups overrepresented in P1 (G) and P2 (H) cultures, respectively. Points represent single GO terms, grouped together according to the GO context view of gprofiler2, under the grouping label of the GO term with the lowest p-value. Overlapping points results in darker grey. Full GO term list and groupings are available in Doc. S2. GO:BP, GO terms concerning biological processes; GO:CC, GO terms concerning cellular components; GP:MF, GO terms concerning molecular functions. (I, J) Heatmaps showing z-scores of the differentially expressed genes contributing most often to the GO terms overrepresented in P1 (I) and P2 (J) cultures. Genes are aggregated according to their involvement in GO terms related to muscular, neuronal and transcriptional (I) or extracellular matrix (ECM) and immunologically relevant (J) terms. Scale bars: 10 µm (B, C), 5 µm (D).

Transcriptome analysis of P1 and P2 RPE populations by bulk RNA sequencing revealed close clustering of cells from the same passage across batches in principle component analysis (Fig. 1E). However, both P1 and P2 cells exhibited high expression levels of typical RPE genes characteristic for the visual cycle (e.g. BEST1, SFRP5, RLBP1), phagocytosis (e.g. LAMP2, ERMN, GULP1), or pigmentation (e.g. DCT, PMEL, MITF) and others (Fig. 1F and Doc. S2 sheet 1). Differential gene expression analysis returned a higher number of genes up-regulated in P1 cultures than in P2 (P1: 853, P2: 229, Doc. S2 sheets 2 and 5), possibly caused by a lower overall RPE purity or due to the developmentally younger P1 RPE not yet being as transcriptomically defined. Indeed, genes more highly expressed in P1 cells were overrepresented in GO terms related to neuronal and muscular cell identity and function as well as transcriptional processes (Figs. 1G and I, Doc. S2 sheets 3 and 4), suggesting the presence of non-RPE cells or young P1 RPE cells at a more pliable stage. In P2 cultures, the upregulated genes were overrepresented in GO terms related to the extracellular matrix and immune-related processes (Figs. 1H and J, Doc. S2 sheets 6 and 7).

In summary, while morphologically similar and showing RPE characteristics, P1 and P2 iPSC-RPE cultures also displayed significant differences in their expression profiles possibly affecting downstream applications.

### Single cell suspension transplantation of P1 but not P2 RPE yields large areas of polarized RPE monolayer

Using the NaIO_3_ mouse model of RPE degeneration as recipient ^10^, the potential of transplanted P1 and P2 iPSC-RPE cells for survival and monolayer formation was assessed. To that end, adult wild type mice (C57BL/6j) received a tail vein injection of 30 mg/kg NaIO_3_ which was sufficient to result in RPE depletion, disruption of the choroid-retina barrier as shown by retinal angiography, photoreceptor death as measured by TUNEL assay and function loss as measured by ERG within 3-7 days (Fig. S1). Seven days after NaIO_3_ injection, P1 or P2 RPE cell suspensions were transplanted into the subretinal space of hosts (Fig. 2A).

**Figure 2:**
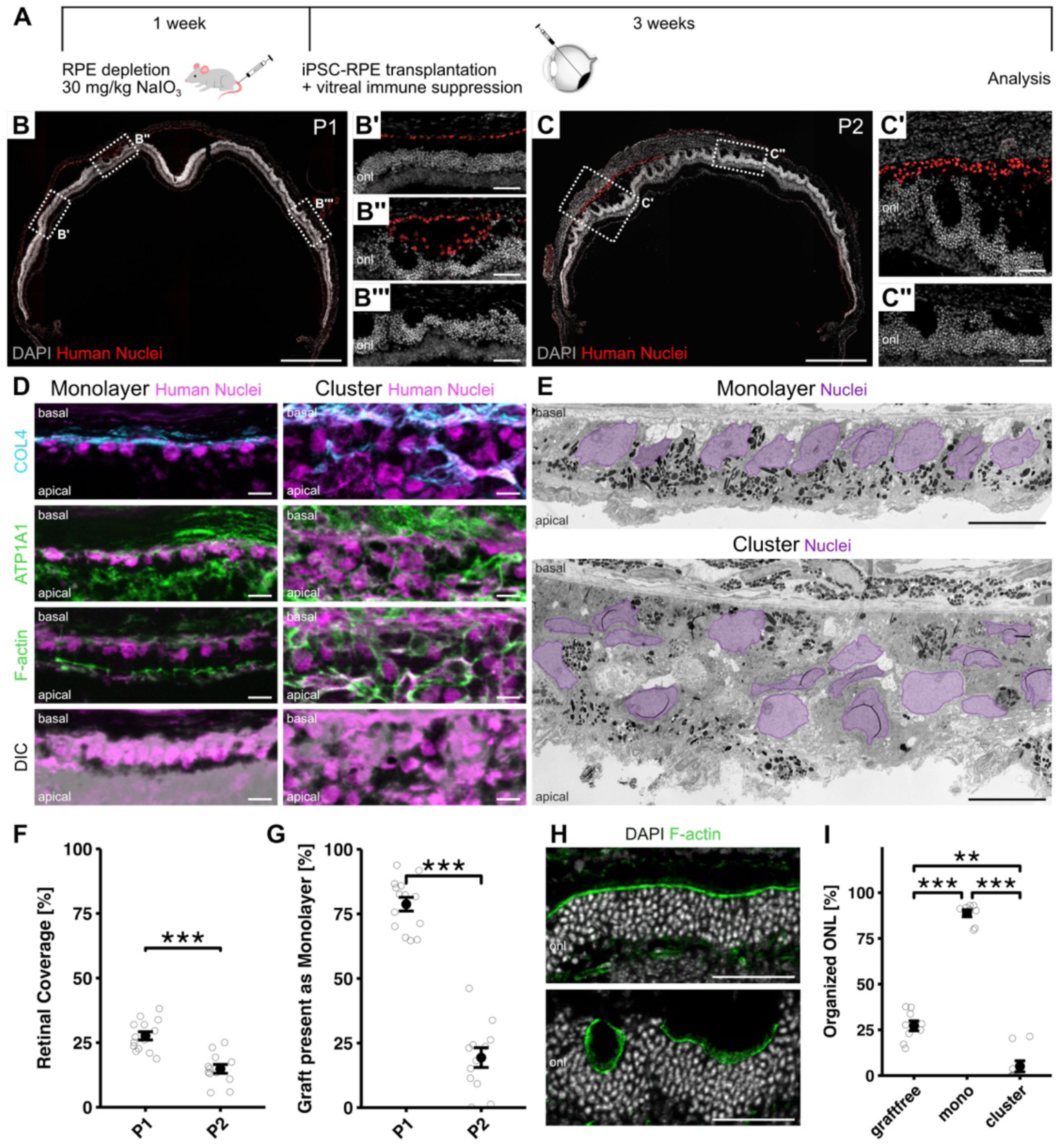
Single cell suspension transplantation of P1 but not P2 RPE yields large areas of polarized RPE monolayer. (A) Scheme of experimental design: Mice received a tail vein injection of 30 mg/kg bodyweight sodium iodate (NaIO_3_) causing depletion of endogenous RPE. One week later, a suspension of P1 or P2 RPE cells was injected into the subretinal space and the immune suppressant triamcinolone acetonide was supplied vitreally. Experimental eyes were analysed 3 weeks post-transplantation. (B) Representative section of a P1 RPE transplanted retina. Human-specific marker staining (Human Nuclei) revealed the survival of P1 RPE cells in large areas with many donor cells forming a monolayer-like structure between choroid and ONL (B, B’) and some cells forming multilayered cell clusters (B, B’’). While the ONL beneath donor cell-derived monolayers appears well organized (B, B’), the ONL beneath clusters or regions without RPE appears disorganized containing foldings and tubulations (B, B’’, B’’’). Cell nuclei counterstained with DAPI. n = 12 biological replicates. (C) Representative section of a P2 RPE transplanted retina. Human-specific marker staining (Human Nuclei) revealed the survival of P2 RPE cells in large areas with most donor cells forming multi-layered cell clusters and few monolayer-like structures (C, C’). The ONL beneath clusters or regions without RPE appears similarly disorganized containing foldings and tubulations (C, C’, C’’). Cell nuclei counterstained with DAPI. n = 12 biological replicates. (D) Immunofluorescence analysis of RPE-derived monolayers and clusters. RPE monolayers show positive immunofluorescence staining for human nuclei, that have a ‘pearls-on-a-string’ appearance, basally localized collagen IV (COL4), and apically localized ATPase (ATP1A1) and F-actin. Combined fluorescence staining with DIC imaging reveals basal localization of RPE nuclei and strong pigmentation apically. In clusters, human nuclei appear as multilayered, unorganized aggregates, that are positive for COL4, ATPase and F-actin, but lack a clear basal-apical polarity. n = 12 biological replicates. (E) Ultrastructural analysis of human RPE-derived monolayers reveals apicobasal polarity including basally localized nuclei, basal membrane/labyrinth and mitochondria, and apical melanosomes. Compare figure S2 for details. Donor-derived clusters show a disorganized appearance without cell polarization. n = 5 biological replicates. (F) Retinal coverage of transplanted P1 and P2 RPE cells. Three weeks after transplantation, P1 donor RPE cells covered almost double the host retinal surface as P2 RPE cells (P1: 27.6% ± 1.6%, n = 14 biological replicates; P2: 14.8% ± 1.7%, n = 12 biological replicates). (G) P1 RPE cells exhibited a significantly larger proportion of the graft as monolayers than P2 RPE transplants (P1: 78.8% ± 2.7%, n= 14 biological replicates; P2: 19.4% ± 3.8%, n = 12 biological replicates) (H) Experimental retinas show areas with organized (top) and disorganized (bottom) ONL, visualized by the OLM shown through staining for F-actin. In contrast to an organized ONL, a disorganized ONL shows foldings, rosettes and tubulations (compare also DAPI staining in figures 2B’ vs. 2B’’- 2C’’). n = 12 biological replicates. (I) Beneath donor RPE monolayers the majority of the ONL shows an organized structure, while the ONL beneath graft-free or cluster areas is mainly disorganized (graft-free: 27.2%±2.7; n = 9 biological replicates; monolayer: 88.8%±1.9, n = 8, cluster: 5.1%±3.0; n = 9 biological replicates). Scale bars: 500 µm (B, C), 50 µm (B’-B’’’, C’, C’’, H) and 10 µm (D, E). onl, outer nuclear layer. Data are represented as mean ± SEM. Data points represent single eyes (F,G,I). Statistics: p < 0.05, **p < 0.01, ***p<0.001 by two-sided Student’s t test (F,G) or by Kruskal-Wallis test with pairwise Wilcoxon test (I).

Three weeks after transplantation immunohistochemical analysis showed the presence of both P1 and P2 cells in the subretinal space (Figs. 2B and C). Retinal cryosections provided further insights into the distribution and organization of transplanted donor cells: While human P1 cells appeared to mainly form single-cell monolayers and only few cell clusters (Fig. 2B-B’’), P2 cells appeared to generate mainly cell clusters and only few monolayer-like structures (Figs. 2C and C’). Generally, RPE monolayers showed typical characteristics of apicobasal polarization including apical expression of ATPase and F-actin besides apical accumulation of pigment granules, as well as a basally located nucleus and basal expression of collagen IV (Fig. 2D). In contrast, clusters of donor cells showed few signs of polarized expression of these apical and basal markers, which were rather found distributed chaotically throughout the graft (Fig. 2D). The differences between cell monolayers and clusters in regard to cell organization and polarization were confirmed by ultrastructural analysis (Fig. 2E). Here, cells within monolayers showed a polarized distribution of RPE organelles, with melanosomes located apically, while the nucleus, basal labyrinth and mitochondria were located basally (Figs. 2E and S2A-A’’’). Clustered cells, in contrast, had a chaotic appearance without obvious signs of polarization and instead often contained large quantities of pigment granules (Fig. 2E). Additionally, laterally located adherens and tight junctions were regularly observed between cells organized in a monolayer, but missing in cell clusters (Figs. 2E, S2A’’ and S2A’’’).

Quantification of whole retina serial sections by fluorescence microscopy revealed that P1 cells occupied almost double the retinal surface as P2 cells (P1: 27.6%±1.6, P2: 14.8%±1.7, Fig. 2F). Additionally, while the majority of human P1 RPE occupied area presented as a monolayer (78.8%±2.7), only 19.4%±3.8 of the P2 RPE area adopted monolayer conformation, with the majority of cells instead forming clusters (Fig. 2G).

Figs. 2F and 2G both contain data from transplanting either 50,000 or 100,000 cells per eye, as aggregating the data by cell number transplanted revealed similar coverage and monolayer formation rates in both cases (Fig. S2B). While not statistically significant, transplanting the lower cell number even led to a slightly increased monolayer formation (Fig. S2B), thus for all further transplantations shown here, 50,000 cells/retina were used. Additionally, analysis at 1, 2, or 3 weeks after P1 RPE transplantation revealed a continuous increase in the proportion of monolayer area (week 1: 44.0%, week 2: 63.4%, week 3: 80.9%, Fig. S2C).

In summary, while human P1 and P2 RPE both survived for up to 3 weeks after transplantation in mouse RPE depleted recipients, P1 cells formed significantly larger areas of polarized, single cell monolayers than P2 cells, which instead mainly generated cell clusters.

### Human RPE monolayers preserve ONL organization and phagocytose outer segments

Proper RPE structure and function is required for photoreceptor homeostasis and light detection. Under homeostatic conditions, photoreceptor organization is characterized by tight stacking of cell bodies/nuclei in the ONL and the presence of a continuous, straight outer limiting membrane (OLM). Irregular stacking and an interrupted, strongly curved OLM indicate that the photoreceptors are stressed or damaged. Thus, as a proxy of photoreceptor support by RPE, we quantified the presence of ‘organized’ vs. ‘disorganized’ ONL areas, as shown by the OLM either being parallel or not parallel to the INL (Fig. 2H). Indeed, quantification revealed a strong photoreceptor support by RPE monolayers, as 88.8 ± 5.4 % of ONL beneath RPE monolayers retained an organized shape (Fig. 2I). In contrast, below cell clusters of either P1 or P2 RPE cells, only 5.7 ± 9.5 % of ONL appeared organized, with the majority of the ONL instead generating foldings, tubulations, rosettes, and exhibiting dislocated nuclei (Fig. 2I). This phenotype was even more pronounced than in areas without any RPE cells, where 27.2 ± 2.7 % of ONL area remained organized (Fig. 2I, compare also Figs. 2B-B’’’, C-C’’). These results highlight the notion that formation of a proper RPE monolayer is essential to support ONL preservation.

Another key RPE function is the continuous phagocytosis of shed outer segments. Phagosomes containing mouse outer segments can be found in the vicinity of human RPE monolayer nuclei in wild type NaIO_3_-treated animals (Fig. S3A). To further validate the capacity of transplanted human RPE to phagocytose mouse outer segments, P1 RPE cells were transplanted into NaIO_3_-treated rhoEGFP mice, which express a Rhodopsin-EGFP fusion protein that localizes to outer segments ^14^. Immunofluorescent staining (Figs. 3A and A’) and ultrastructural analysis by correlative light and electron microscopy (CLEM) at 3 weeks post transplantation confirmed the presence of GFP-positive structures within the human RPE cells, indicating their phagocytotic activity (Figs. 3B-B’’ and C-C’’).

**Figure 3:**
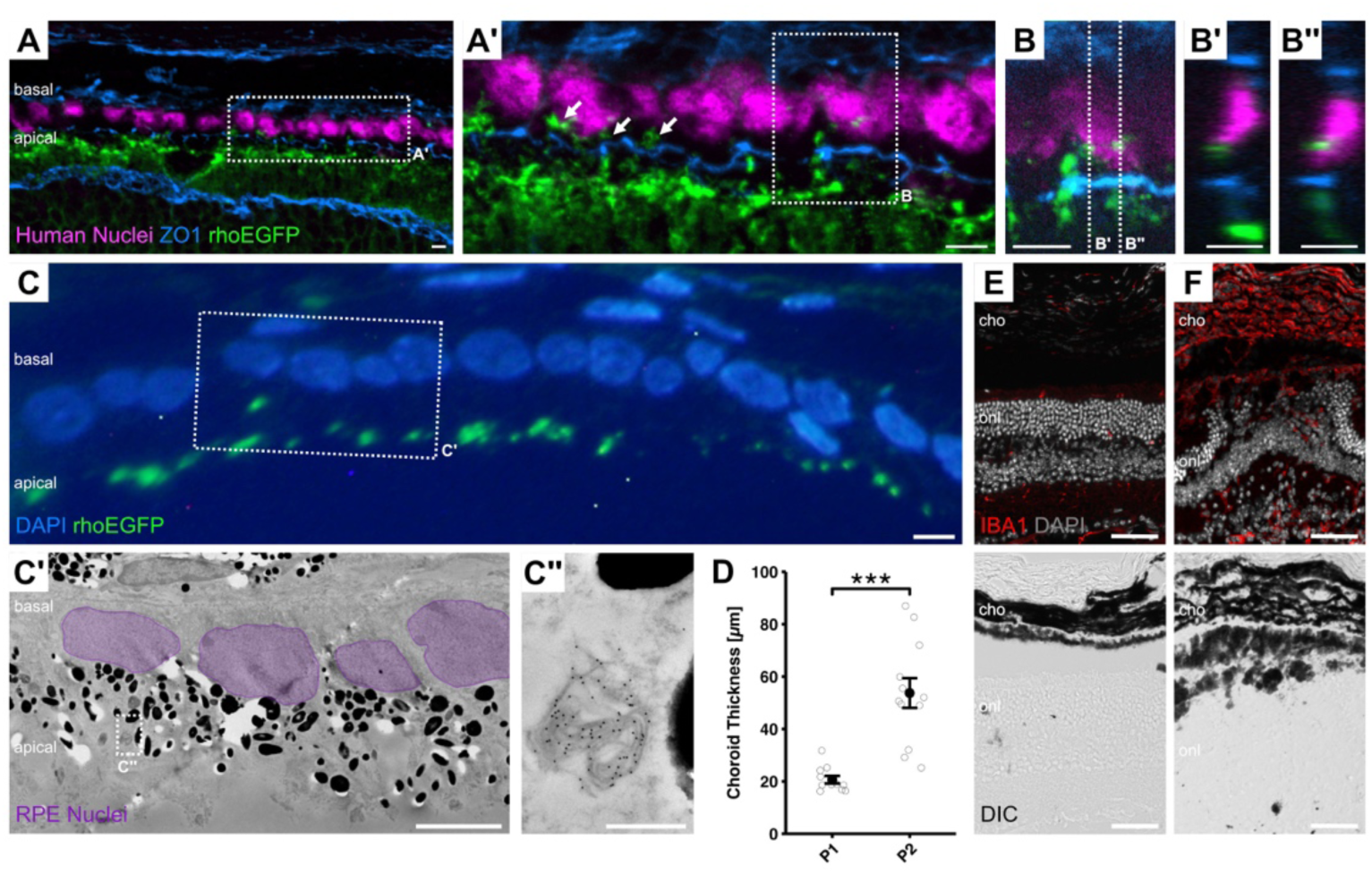
Donor RPE cells phagocytose photoreceptor outer segments and P2 cells induce strong immune reaction. (A) Phagocytosed photoreceptor outer segments can be detected in human RPE monolayer cells (arrows) three weeks after transplantation into NaIO_3_-treated rhoEGFP mice. n = 6 biological replicates. (B) Single plane of the z-stack area highlighted in A’. B shows the xy plane, B’ and B’’ show the zy planes at x positions marked in B. (C) Correlative light- and electron-microscopy (CLEM) reveals host outer segment particles within transplanted RPE cells, visualized by immunofluorescent (C) and immunogold (C’, C’’) labelling of rhoEGFP protein three weeks after transplantation into the retina of NaIO_3_ treated rhoEGFP mice. n = 3 biological replicates. (D) Retinas with P2 RPE donor cells show a significantly thicker choroid in comparison to P1 transplanted mice (C; P1: 20.7 µm ± 1.4, n = 11; P2: 53.7 μm ± 5.7, n = 12 biological replicates), pointing to a stronger immune response triggered by P2 transplants. (E, F) Retinas with P2 RPE donor cells (F) show increased numbers of ionized calcium-binding adapter molecule 1 (IBA1)^+^ cells both within the retina and the choroid than retinas that received P1 RPE donor cells (E). n = 12 biological replicates. Scale bars: 50 µm (E, F), 5 µm (A-C’) and 500 nm (C’’). cho, choroid; onl, outer nuclear layer. Data are represented as mean ± SEM. Statistics: p < 0.05, **p < 0.01, ***p<0.001 by two-sample Wilcoxon test (D).

### P2, but not P1 RPE transplants induce a strong immunological response

In healthy conditions, the retina represents an immune-privileged tissue, with resident microglia and Müller Glia (MG). Upon NaIO_3_-induced RPE and photoreceptor loss, MG become gliotic and GFAP upregulation was present in both P1- and P2- transplanted eyes (Fig. S3B). However, striking differences in choroid thickness and microglia activation were observed between P1 and P2 transplanted retinas. The choroid of eyes transplanted with P2 RPE cells was on average more than twice as thick compared to eyes transplanted with P1 RPE (53.7 ± 19.7 μm and 23.0 ± 9.3 μm respectively; Fig. 3D). Additionally, in P2 RPE-transplanted retinas a striking increase of cells positive for IBA1 (Ionized calcium-binding adapter molecule 1), a marker for microglia and macrophages, was observed both in the retina and the choroid (Fig. 3F). In P1 RPE transplanted eyes only few IBA1^+^ cells were located within the retina, mainly in the inner and outer plexiform layers, and almost no IBA1^+^ cells localized within the choroid (Fig. 3E).

In summary, transplantation of P2 RPE cell suspensions triggered a stronger immune reaction in comparison to P1 RPE cells.

### Different RPE P1 subpopulations can be identified using cell surface markers

Although in vitro differentiation of RPE is generally considered to yield a homogenous cell product, it has become more and more evident that differences between RPE subgroups exist both in vivo and in vitro, with the latter having important implications in cell performance after transplantation ^6,15^. Obvious phenotypic heterogeneity between cells, namely varying degrees of pigmentation between cells of the same passage P1 (Fig. 1B), inspired an empirical flow cytometry-based screening approach using a pre-formatted antibody panel covering 371 cell surface markers, with the aim to identify markers correlating with subpopulations of differentially pigmented cells. Side scatter signal intensity (SSC) was used as proxy for pigmentation and hits were selected based on either negative or positive correlation to identify depletion or enrichment of marker candidates, respectively (Fig. 4A, Doc. S3). Nine isotype controls and one empty well per screening plate served as controls.

**Figure 4:**
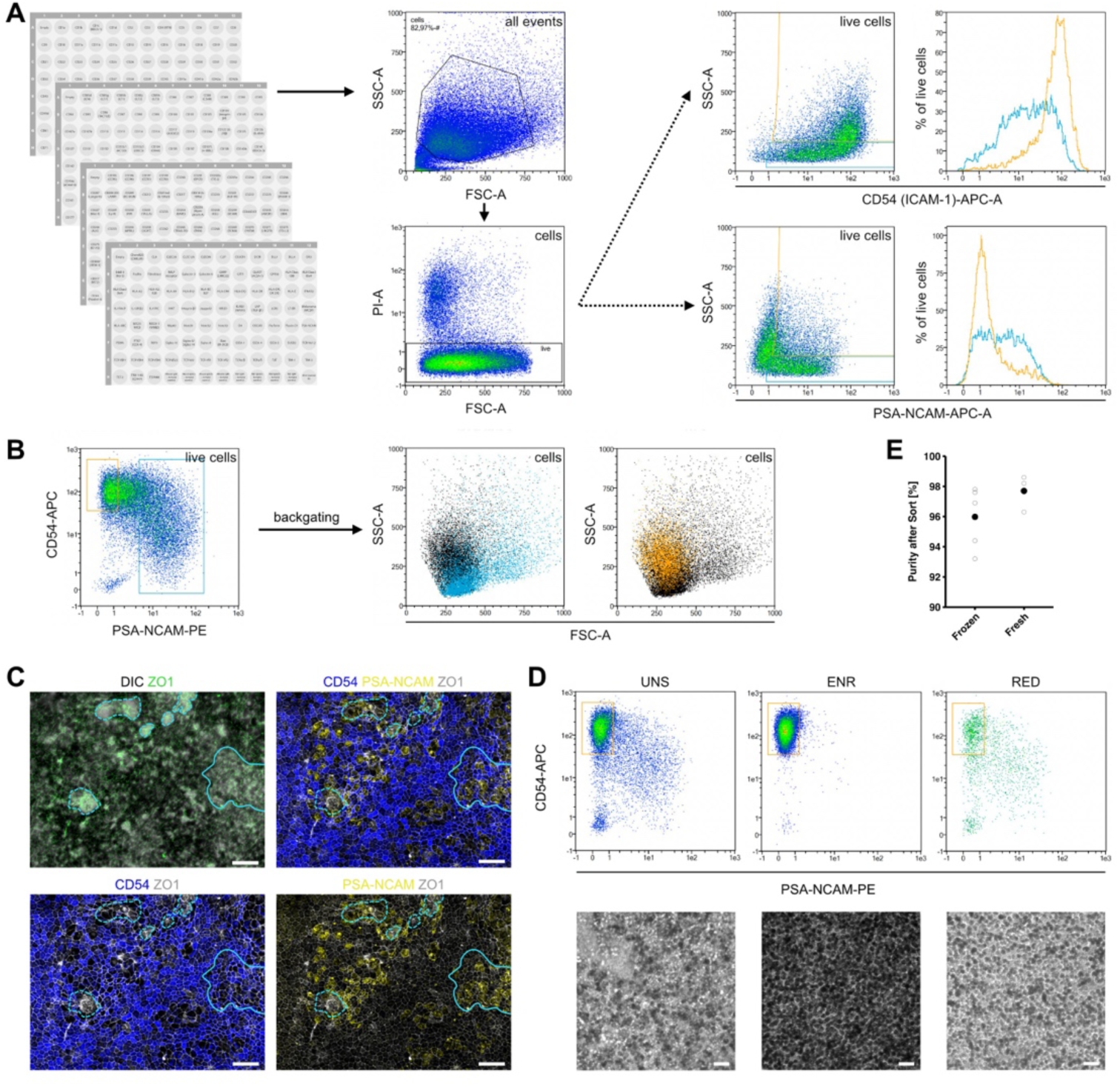
Identification and validation of cell surface sorting markers for iPSC-derived P1 RPE cells. (A) P1 RPE cells were screened for expression of 371 antibodies directed against cell surface markers using flow cytometry. Hits were identified based on correlation to side scatter (SSC) intensity as proxy for pigmentation, here shown for CD54 (ICAM-1) (positive correlation) and PSA-NCAM (negative correlation). (B) Combinatorial staining for CD54 and PSA-NCAM reliably discriminates SSC-low non-target (unpigmented, backgated in blue) and SSC-high target cells (pigmented, backgated in orange). (C) Immunofluorescent staining of RPE cells in culture shows cells expressing PSA-NCAM but little CD54 mostly in regions lacking pigmentation and proper cobblestone morphology (areas highlighted by dashed lines). Co-expression of CD54 and PSA-NCAM appears to occur primarily in regions of intermediate pigmentation levels and mixed cobblestone morphology (area highlighted by solid line). n = 2 biological and 3 technical replicates. (D) Flow scatter and corresponding transmitted light microscopy images for unsorted (UNS), CD54^+^/PSA-NCAM^-^ -enriched (ENR) and CD54^+^/PSA-NCAM^-^-reduced (RED) cell fractions. Cells were sorted using the MACSQuant Tyto cell sorter to maximize target cell purity. Replated fractions reveal that pigmented cells can be efficiently enriched with the dual marker strategy. All images were taken with the same exposure. n = 2 biological and 3 techical replicates. (E) CD54^+^/PSA-NCAM^-^ sorting of frozen and freshly harvested cells consistently yielded a target fraction of high purity. Circles show single biological replicates, solid points their average. Scale bars: 100 µm (D), 50 µm (C).

After validation of several initial hits the combination of CD54 (ICAM) positivity and PSA-NCAM negativity proved to be the most robust combination allowing identification (Figs. 4B and 4C) and reproducible isolation of highly pigmented target cells (Fig. 4D). Consequently, effective depletion of unpigmented cells and a high degree of pigmentation was observed for the target-enriched (ENR) fraction after replating in vitro (Fig. 4D, ENR), while the unsorted and target-reduced fractions showed a lower overall pigmentation level (Fig. 4D, UNS & RED). An average purity of 96.0% ICAM^+^/PSA-NCAM^-^ (Fig. 4E) was achieved in the ENR fraction from cell preparations used in the screening process, which were shipped frozen and stained immediately after thawing. Similarly, fresh cells collected from the transwell inserts directly before staining could be enriched to an average purity of 97.7%, ICAM^+^/PSA-NCAM^-^ (Fig. 4E), the latter being used for subsequent in vivo studies.

### Transplanted CD54^+^/PSA-NCAM^-^ P1 RPE cells show improved monolayer formation

To assess whether the highly pigmented cell surface marker-defined P1 RPE subpopulation has the capacity for monolayer formation after transplantation in vivo, P1 RPE cells were dissociated and sorted using the MACSQuant® Tyto® cell sorter (Miltenyi Biotec) to obtain two cell fractions: CD54^+^/PSA-NCAM^-^ enriched (ENR), and CD54^+^/PSA-NCAM^-^ reduced (RED) in addition to the unsorted (UNS) control. While the percentage of CD54^+^/PSA-NCAM^-^ cells fluctuated in the UNS and RED fraction depending on the differentiation round, sorting consistently yielded >96% pure CD54^+^/PSA-NCAM^-^ cells in the ENR fraction (Figs. 4E, S4A and S4B). All cell fractions were subretinally transplanted into the NaIO_3_ mouse model of RPE depletion and analyzed after three weeks.

Immunostaining of human nuclei and F-actin showed large grafts in flat-mounted experimental eyes (Fig. 5A (UNS), B (ENR), C (RED), dashed white outlines). Two main appearances could be distinguished: firstly, graft regions with an intense human nuclei signal but lacking cobblestone morphology, dotted with pigment-rich spheres (Figs. 5A and 5C, magnified in Fig. 5A’) likely representing macrophages (Fig. S5A); secondly graft regions exhibiting extensive cobblestone morphology but a seemingly weaker human nuclei signal (Figs. 5B, 5B’ and 5B’’). It is important to note that this comparatively weak human nuclei staining arises from concealment of the nuclei by the more apically positioned, light-absorbing pigment granules (Figs. S4C-F’) in the highly structured and polarized human RPE monolayer. Towards graft edges, the monolayer regions showed again higher human nuclei visibility (Fig. 5B’’ and Figs. S4C-F’), suggesting an absence of the characteristic pigment, while neither cobblestone morphology nor human nuclei staining was found outside of the graft areas (Fig. 5C’).

**Figure 5:**
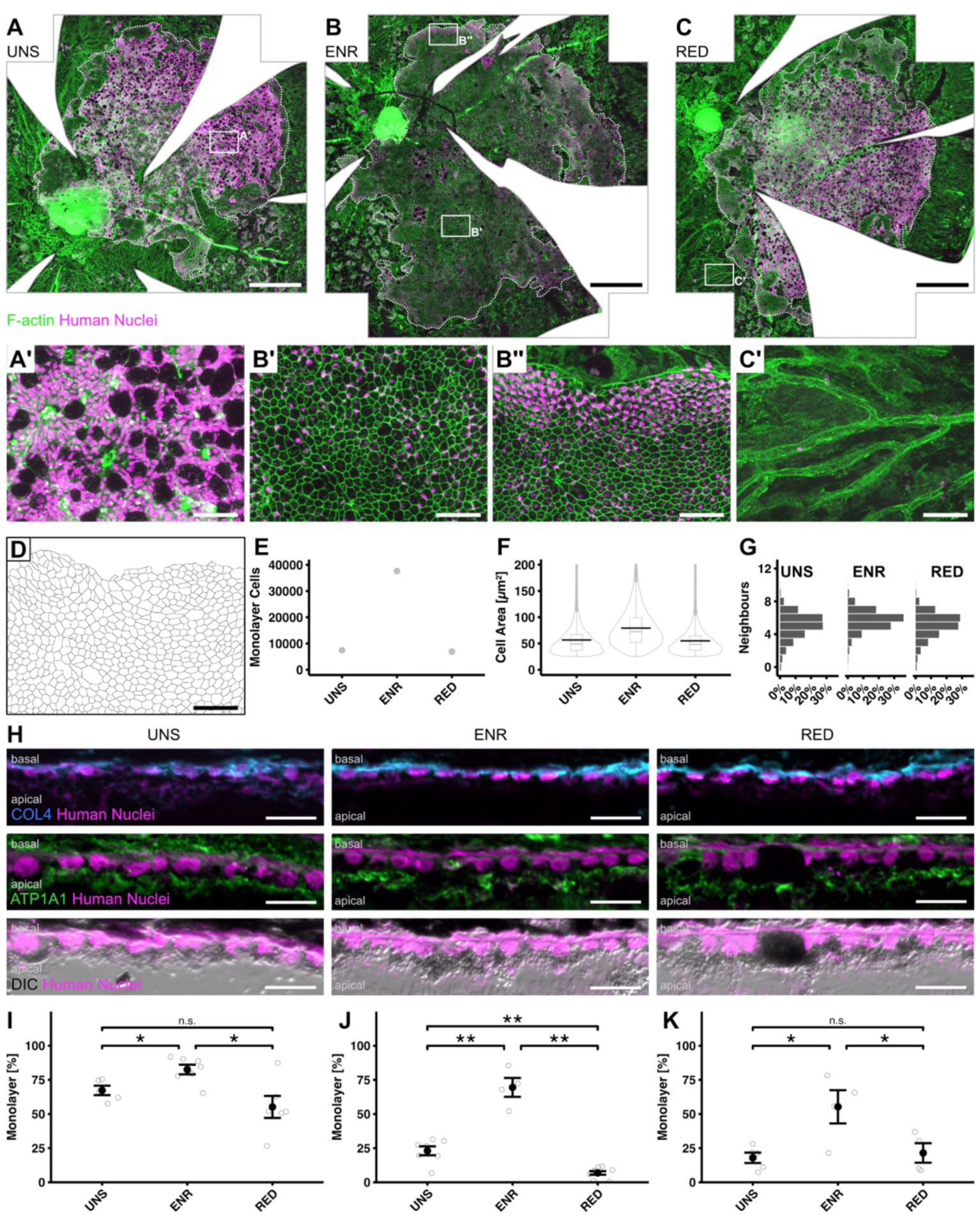
CD54^+^/PSA-NCAM^-^ RPE cells show increased monolayer formation after transplantation. (A – C) Flat-mounted eyecups that received P1 RPE single cell suspensions which were either unsorted (UNS; A), enriched (ENR, B), or reduced (RED, C) in CD54^+^/PSA-NCAM^-^ cells show large areas of donor cells identified by Human Nuclei staining (graft region outlined with dotted lines). While in UNS (A, A’) and RED (C) samples the proportion of area with cell clusters (visible through intense Human Nuclei staining) is high, in ENR transplanted retinas the proportion of monolayer with typical cobblestone morphology as seen by staining for F-actin is high (B, B’, B’’). Human nuclei visibility in properly structured monolayers is concealed due to high pigmentation of donor cells (B’’; for single z-planes see figures S4E and F), but better visible at monolayer edges (B’). Magnification of an RPE-depleted area without donor cells is shown in C’. n > 4 biological replicates. (D) Outlines of monolayer cells from B’’ as detected with the REShAPE tool^15^ show the possibility to analyse the grafts in flatmount view. (E-G) REShAPE analysis of the flatmounts A-C (corresponding REShAPE cell outline maps see figure S4G-I). (E) reveals a highly increased number of monolayer cells in the ENR graft in comparison to UNS or RED (UNS: 7453, ENR: 37607, RED: 6921). ENR monolayer cells on average appear larger than UNS or RED monolayer cells (F; mean areas: UNS 57 µm^2^, ENR 79 µm^2^, RED 55 µm^2^, violin plots of all cells detected with overlaid box-plots showing median and 25% quartiles, black lines showing the mean). In all three conditions human monolayer cells have mostly 5 or 6 neighboring cells, with the ENR graft having the highest ratio of cells with 5-7 neighbors (G; UNS: 67%, ENR: 82%, RED: 69%). (H) Monolayers formed in all conditions show a polarized morphology with basal collagen IV (COL4) and apical ATPase (ATP1A1) expression. Human nuclei are located basally, while pigmentation (dark in DIC images) appears mainly apically. Large, highly pigmented, round structures are frequently detected in clusters (see also A’) and sometimes in monolayer areas, likely representing pigment-filled macrophages (compare also figure S5A). n > 4 biological replicates. (I, J) Quantification of monolayer proportion after transplantation of RPE cells from two separate differentiation rounds. While in both rounds the CD54^+^/PSA-NCAM^-^ ENR donor cell fractions robustly generated similarly high proportions of monolayer (ENR - I : 82.5% ± 3.5%; n=7; ENR – J: 69.5% ± 6.9%; n=4), the amount of monolayer generated by UNS (UNS - I: 67.3% ± 3.5%; n=5; UNS - J: 23.1% ± 3.3%, n=7) and RED cell fractions (RED - I: 55.2% ± 8.1%; n=6; RED - J: 6.9% ± 1.3%, n=9) strongly varied, correlating with different amounts of CD54^+^/PSA-NCAM^-^ cell purity within the different fractions (see also figure S4A for purity data related to figure 5I and figure S4B for purity data related to figure 5J). All datapoints within one plot and fraction represent biological replicates, I and J represent technical replicates. (K) Quantification of monolayer proportion after transplantation of RPE cells from a third differentiation round, sorted with conventional flow cytometry (UNS: 17.9% ± 3.8%, n = 5; ENR: 55.3% ± 12.2%, n = 4; RED: 21.5% ± 7.2%, n = 4. All datapoints within one fraction represent biological replicates). Scale bars, 500 µm (A, B, C), 50 µm (A’, B’, B’’, C’, D) and 20 µm (H). Data are represented as mean ± SEM. Statistics: p < 0.05, **p < 0.01, ***p<0.001 by Kruskal-Wallis with pairwise Wilcoxon test (I,J,K).

Exploratory assessment with the image-analysis tool REShAPE ^15^ allowed the reconstruction and analysis of the graft monolayer regions of one flatmount per fraction (Figs. 5D and S4G-I). REShAPE analysis showed the ENR sample contained 5x and 5.4x as many monolayer cells as the UNS and RED samples, respectively (UNS: 7453, ENR: 37607, RED: 6921 cells; Fig. 5E), and overall generated the largest monolayer cells (mean areas: UNS 57 µm^2^, ENR 79 µm^2^, RED 55 µm^2^; Fig. 5F). The notion of proper RPE monolayer formation by the human cells was further underlined by high numbers of monolayer cells in all fractions neighboring 5 to 7 others, with ENR monolayer cells having the largest number of such cells (UNS: 67%, ENR: 82%, RED: 69%; Fig. 5G)

Analysis of retinal cryosections showed human cells again creating polarized single cell monolayers or unorganized clusters between the ONL and the choroid (Figs. 5H and S5B). Interestingly, P1 RPE fractions enriched for CD54^+^/PSA-NCAM^-^ cells showed a significantly increased monolayer formation when compared with unsorted cells or cells with a reduced proportion of CD54^+^/PSA-NCAM^-^ cells (Fig. 5I), despite similar overall retinal coverage (Fig. S5C). Importantly, enrichment of CD54^+^/PSA-NCAM^-^ cells yielded populations with similarly high monolayer formation capacity across experiments (Figs. 5I and 5J). The UNS and RED fractions on the other hand were found to differ strongly in that regard, with cell culture batches that contained a higher percentage of CD54^+^/PSA-NCAM^-^ cells yielding more monolayer (Fig. 5I: UNS 72.7% CD54^+^/PSA-NCAM^-^ cells) than batches with a lower percentage of CD54^+^/PSA-NCAM^-^ cells (Fig. 5J: 34.1% CD54^+^/PSA-NCAM^-^ cells).

To test whether enriching CD54^+^/PSA-NCAM^-^ cells robustly improves monolayer formation capacity across contexts, we varied sorting strategies. Similarly to the Tyto sorting, enrichment by conventional flow cytometry with a BD FACSAria™ III (BD Bioscience) yielded a purity of >97% in the ENR fraction (Fig. S5D), with the ENR fraction again showing an improved amount of monolayer formation after transplantation into the NaIO_3_ mouse model compared to cell fractions that were unsorted or reduced in CD54^+^/PSA-NCAM^-^ cells (Fig. 5K).

In summary, our combined results provide evidence that an RPE subpopulation sortable by the cell surface markers CD54^+^/PSA-NCAM^-^ at an early differentiation stage has the potential for extensive monolayer formation after subretinal transplantation into a mouse model of RPE depletion.

## Discussion

While not yet clinically established, transplantation of pluripotent stem cell-derived RPE cells for replacement approaches in the retina have continuously developed over the last two decades, with a number of clinical trials providing evidence for its safety and, in part, showing therapeutic effects like stabilization or even some improvement of visual functions ^3,16^. Advancing RPE cell generation and selection might be helpful to further improve transplantation outcomes. Here, we used the NaIO_3_ mouse model of RPE depletion to systematically assess the capacity of PSC-derived RPE cells to form polarized monolayers after transplantation in vivo. Our combined results provide evidence that human P1 RPE cells have a significantly increased potential in comparison to P2 RPE cells to generate extensive monolayers with rescue properties in the mouse retina. Additionally, cell surface marker sorting by CD54^+^/PSA-NCAM^-^ allowed isolation of a monolayer-forming subpopulation within the P1 RPE cell population, independent of the sorting method used. Thus, defining competent RPE cells appears as an important precondition to improve transplantation outcomes.

Polarized monolayer formation is an essential prerequisite for RPE cells to properly function. The RPE monolayer is part of the retina-blood-barrier in the eye and is involved e.g. in the phagocytosis of photoreceptor outer segments, ion buffering, epithelial transport of nutrients, ions and water, growth factor secretion, and light absorption ^17^. We saw striking differences in the potential of RPE cells at different developmental stages to generate monolayers after transplantation into a retinopathy mouse model depleted of endogenous RPE. While “young” RPE cells, i.e. P1 cells, generated extensive monolayers, P2 RPE cells instead mainly generated unorganized cell clusters. Immunohistochemical and ultrastructural analysis revealed a mature RPE phenotype within donor cell-derived monolayers including apicobasal polarized location of diverse organelles and proteins, which was absent in donor cells organized in clusters. The formation of RPE monolayers consequently had a significant influence on the underlying ONL: while underneath cell clusters the ONL showed massive pathological changes including foldings, tubulations and photoreceptor loss, as it was also observed in areas without RPE cells, the ONL in regions with newly formed RPE monolayers did not show signs of degeneration and had a highly organized appearance. Thus, the formation of monolayers appears to be essential for these therapeutic effects. Indeed, a recent study using RPE derived from RPESCs showed improved ONL and vision function rescue after transplantation at an early, defined developmental stage into the RCS model of RPE dysfunction ^8^. However, whether improved monolayer formation or other factors were the underlying mechanism for the described rescue effects was not assessed in that study ^8^.

In order for cell transplantations to be successful, the heterogeneity of RPE in vivo and in RPE cultures should be taken into account. Distinct RPE subpopulations were recently identified within concentric regions of the adult human retina, calling for further characterization and selection of RPE subpopulations in regard to function and potential therapeutic use in central versus peripheral retinal regions ^15^. Similarly, single cell RNAseq analysis of primary and stem cell-derived RPE populations revealed a higher heterogeneity, also in early passages in culture, than previously thought ^6,7,18–20^. Thus, also subpopulations of cultured RPE cells must be characterized more thoroughly with regards to their potential for integration and monolayer formation after transplantation in vivo. Interestingly, expression profiling in our study also showed significant differences in gene expression already at early passages, i.e. between P1 and P2 RPE cultures, though the morphological appearance including monolayer formation, pigmentation, and RPE-specific marker expression in vitro was still similar.

Given the high possible heterogeneity within and between cell culture batches, it is vital to understand the effect of cell subpopulations on transplantation outcomes when moving forward with clinical trials. Cell surface marker-based sorting therefore represents an important tool for high-throughput target cell isolation and quality control in a donor cell production pipeline. For stem cell-derived RPE cultures several studies identified RPE-specific cell surface markers, mainly with the goal to purify RPE cells from heterogenous cultures, particularly for protocols that were previously based on manual separation of pigmented cell populations ^18,21^. As such, removal of unwanted, non-RPE cells using CD56^+^ and CD24^+^ cell depletion has been introduced for the generation of RPE monolayers and patches in vitro for transplantation purposes ^21^. Reyes et al identified CD140b^+^/CD184^−^ sorting for improving RPE cultures that also have the capacity for monolayer formation after transplantation in rabbits ^18^, while the lncRNA TREX has been identified as a biomarker for RPESC-derived RPE with increased monolayer forming capacity in vitro and potential for functional improvement after transplantation into RCS rats ^6^. It would be of great interest to systematically and quantitatively assess these markers or their combinations in regard to RPE monolayer formation after transplantation in vivo.

Our human cell surface marker screen using 371 cell surface antibodies identified multiple cell surface markers capable of selecting highly pigmented RPE subpopulations. This screening data is a valuable resource for future refinement of sorting strategies confirming on the one hand the validity of already reported markers (CD24, CD56) and suggesting additional candidates (e.g. CD44, CD239; see also Doc. S3). In our hands, however, the marker combination CD54^+^/PSA-NCAM^-^ proved to be most robust during validation and indeed allowed to isolate a key RPE subpopulation contributing to monolayer formation after transplantation into the RPE-depleted NaIO_3_ mouse model. Direct quantitative comparison of CD54^+^/PSA-NCAM^-^ enriched (>95%) vs reduced (<50%) RPE fractions showed significant differences in monolayer formation in vivo. While the identification of potential cell surface markers for the enrichment of specific RPE subpopulations was based on pigmentation intensity, a recent study provided evidence that pigmentation intensity by itself does not correlate with specific RPE marker expression ^22^. Thus, further studies might be necessary to assess in more detail the direct (or indirect) context between pigmentation intensity and RPE maturation.

Indeed, the marker combination CD54^+^/PSA-NCAM^-^ might also mechanistically reflect its use for homogenous RPE monolayer formation as CD54 represents the intercellular adhesion molecule 1 (ICAM-1), an adhesion molecule of the immunoglobulin superfamily, classically involved in intercellular adhesion. The epitope PSA-NCAM, derived from the posttranslational modification of the neural cell adhesion molecule (NCAM) with polysialic acid, is considered a marker for immature neural precursors ^23^ and can be used to isolate primitive, proliferative neural progenitors derived from pluripotent stem cells ^24,25^. Thus, depletion of PSA-NCAM^+^ cells may reduce undesirable contaminants with proliferative potential and fate. The combinatorial approach with a positive marker confers a double layer of security to reduce heterogeneity of RPE cell compositions since an independent positive criterion may discriminate unpredictable off-target fates. Furthermore, the purification process allows for seamless implementation into GMP-manufacturing, e.g. based on MACS technology or GMP-compliant multiparametric cell sorting in a closed cartridge system using MACSQuant Tyto cell sorter.

When characterizing the human RPE grafts, the REShAPE artificial intelligence-based image analysis tool ^15^ allowed the assessment of entire human RPE transplant monolayers with single-cell resolution. Indeed, the donor cell-derived monolayers in the NaIO_3_ mouse model were highly similar in morphology, shape and number of neighboring cells to RPE monolayers in the human eye. While the average size of human RPE after transplantation in the mouse was significantly smaller than in adult human eyes (here: 60-80 µm^2, 15^ between 150 – 350 µm^2^;), the smaller size might be a result of the developmental stage of the iPSC-RPE used (P1) and time after transplantation (3 weeks), thus still representing fetal RPE cells rather than the far older RPE cells of donor ages 56-92 years described by Ortolan et al (2022). Further increasing the time of donor RPE in vivo would therefore be interesting to study. Additionally, specific signals or mechanic restrictions coming from the mouse environment might influence the growth of transplanted human RPE cells.

Currently, RPE replacement clinical trials either inject cell suspensions or preformed monolayers as sheets or on scaffolds as patches ^3,16^. An obvious advantage of sheets/patches is the already formed monolayer allowing for immediate activity and reduction of disorganized cell fractions. However, their placement underneath the retina, particularly in a degenerative environment, is associated with challenging surgery, while the subretinal application of a cell suspension is less traumatic. Interestingly, both delivery approaches appear to be safe and reproducible showing long-term survival with limited immune responses in some patients for more than three years ^3,16^. Sharma et al (2019) directly compared RPE suspension and patch injections in the RCS rat model of RPE dysfunction, showing similar ONL and functional rescue potential for both delivery methods, though the patch contained only 10,000 while the suspension contained 100,000 cells ^4^. However, the histological images in that study showed that the RPE suspensions formed only unorganized cell clusters and no monolayers in the recipients, suggesting that effects other than specific RPE function might be responsible for the observed histological and functional improvements: a well described phenomenon in RCS rats, where the injection of diverse cell populations including e.g. neural progenitors, fibroblasts, or Schwann cells resulted in rescue effects ^13,26,27^. In contrast, after injection into a laser-induced retinal degeneration porcine model, the RPE patches showed increased rescue potential in comparison to RPE suspensions at the tissue as well as functional level ^4^. Again, cell suspensions in that study did not generate single-cell monolayers but instead formed unorganized cell clusters that lost expression of the RPE maturity marker RPE65. Whether the poor performance of transplanted RPE suspensions in the porcine model was caused by not selecting for the proper RPE sub-population prior to injection or the animal model chosen would be interesting to assess. Indeed, specific environmental requirements, like a still remaining Bruch’s membrane, might be advantageous for proper monolayer formation after RPE suspension injections ^28^. This possibility not only calls for the systematic assessment of different RPE degeneration models to test RPE replacement strategies but also emphasizes the need to refine the selection of patients, disease forms and disease stages to identify suitable environments for successful transplantation approaches in clinical trials.

In conclusion, the developmental stage/differentiation time of in vitro generated RPE cells significantly influences their monolayer formation capacity after transplantation in vivo. Furthermore, the cell surface markers CD54^+^/PSA-NCAM^-^ allow for enrichment of a monolayer-forming RPE subpopulation, highlighting the need for the careful selection of appropriate donor cells for cell replacement approaches of RPE in the eye. These insights might be helpful for improving success of RPE suspension transplantation also in future clinical applications.

## STAR Methods

## KEY RESOURCES TABLE

See separate document.

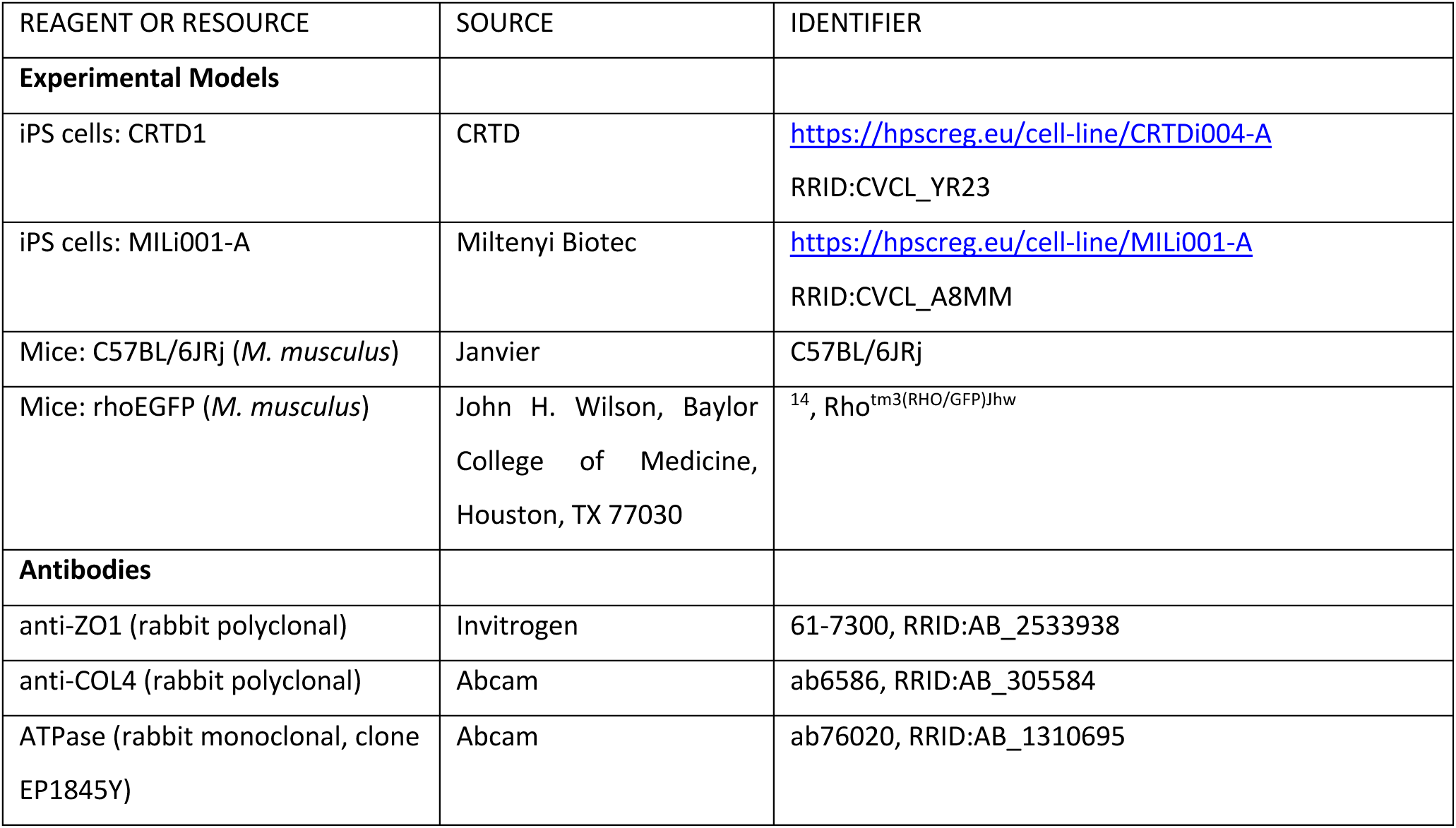
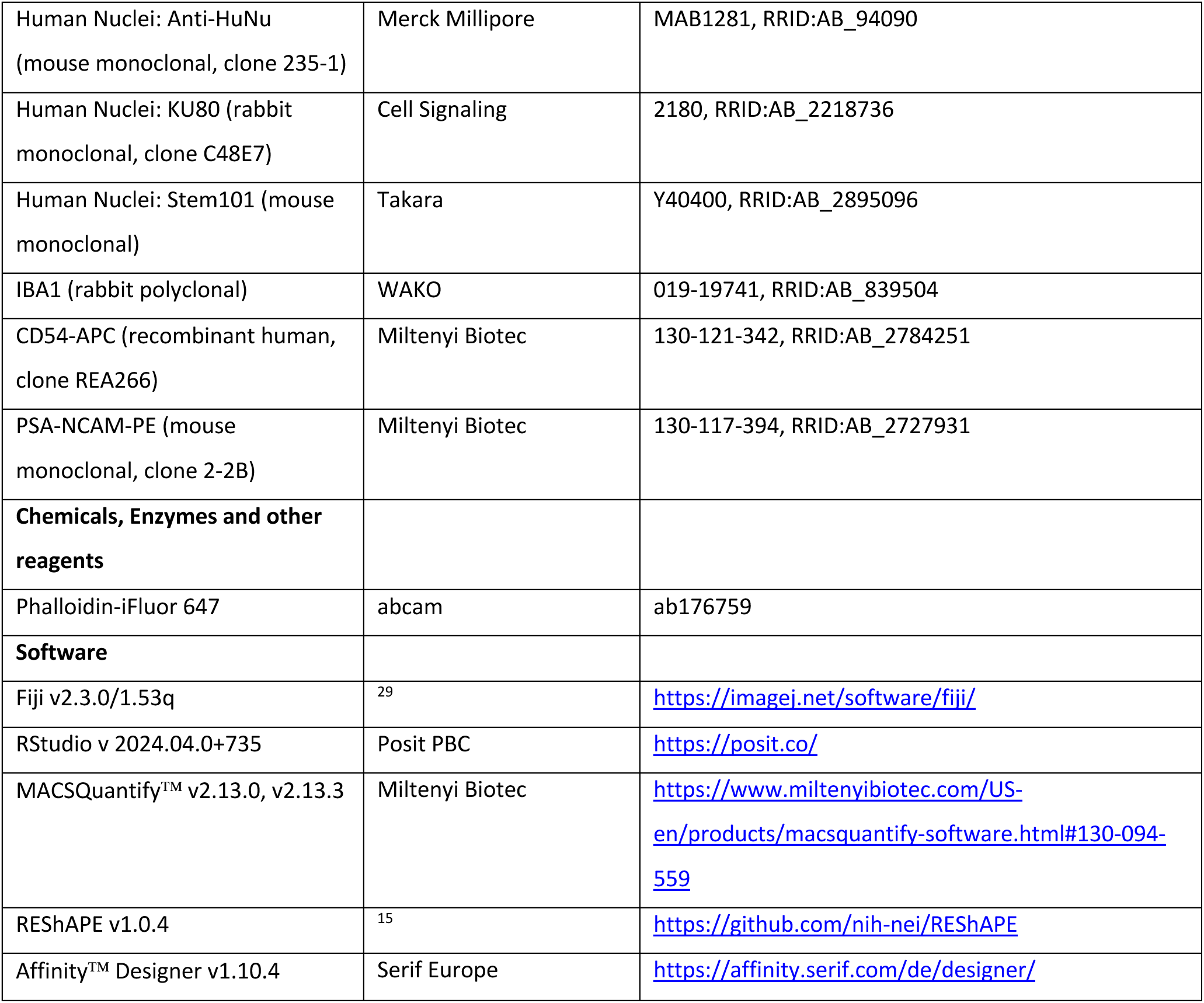

## RESOURCE AVAILABILITY

### Lead contact

- Further information and requests for resources and reagents should be directed to and will be fulfilled by the lead contact, Marius Ader.

### Materials availability

- This study did not generate new unique reagents.

### Data and code availability

- Bulk transcriptome data is available as raw count matrix in the supplementary files (Document S2). Raw sequencing data can be made available upon reasonable request.
- This paper does not report original code.
- Any additional information required to reanalyse the data reported in this paper is available from the lead contact upon request.

## EXPERIMENTAL MODEL AND STUDY PARTICIPANT DETAILS

### Animals

Details and sources of mouse lines can be found in the Key Resources table. Animals were maintained on a standard 12 hour light-dark cycle. Mice were provided with fresh bedding and nesting weekly and received food and water ad lib. One week before transplantation, endogenous RPE was ablated by a systemic injection of 30 mg/kg body weight sodium iodate (NaIO_3_) in isotonic NaCl solution into the tail vein. Animals were used for transplantation 7 days later, at 9-13 weeks of age, and kept for a further 3 weeks post-transplantation, under the conditions described above. Only females were used as recipients. Animals received treatment in both eyes, consisting of a subretinal injection of the cell suspension, followed by intravitreal injection of an immune suppressant (details below). No systemic immunosuppression was administered. Animals were sacrificed by cervical dislocation. All animal experiments adhered to the ARVO Statement for the Use of Animals in Ophthalmic and Vision Research and were approved by the animal ethics committee of the TU Dresden and the Landesdirektion Dresden (DD24.1-51531/449/69).

### Cell lines

Cell line details can be found in the Key Resources table. CRTD1 cells were cultured at 37°C, 5% CO_2_, in mTeSR PLUS (StemCell Technologies, 100-0276) on plates coated with Matrigel hESC-qualified matrix (Corning, 354277) with medium changes every 2-3 days. Colonies were passaged as aggregates using Dispase (StemCell Technologies, 07923) when reaching 40-60% confluency (every 3-7 days, depending on growth speed).

## METHOD DETAILS

### Animal model characterization

#### Quantification of RPE cell number and cell death

To characterize the animal model, RPE cell number was quantified by counting the number of RPE cells present within a distance of 200 μm around the optic nerve in cross sections where the optic nerve head (ONH) was present (7-24 sections per animal), including only cells with a wild type morphology and/or visible DAPI staining. To evaluate cell death, slides were submitted to the TUNEL assay (Roche, 11684795910), according to the manufacturer’s instructions, and the total of TUNEL-positive cells per section was counted in four sections per animal where the ONH was present.

#### Electroretinogram (ERG)

Mice were dark-adapted overnight and anesthetized under dim red light by an intraperitoneal injection of 30 µg/kg bodyweight (bw) Ketamine (medistar, 04-03-9264/23) and 1 µg/kg bw Medetomidine hydrocholoride (Orion Pharma, Domitor®) in sterile isotonic solution (Fresenius Kabi, B306175/03), followed by application of dilating eye drops containing 2.5% phenylephrine and 0.5% tropicamide (University Clinic Pharmacy Dresden). After 1-2 min, eye drops were replaced with clear moisturizing eye gel (Bausch+Lomb, Vidisic). Full-field ERGs were recorded using a Ganzfeld bowl (Roland Consult). Single-flash responses were obtained under dark-adapted (scotopic, with the following intensities: 0.0003, 0.001, 0.003, 0.01, 0.03. 0.1, 0.3, 1, 3 and 10 cd.s.m-2) and light-adapted (photopic, 10 cd.s.m-2) conditions. Light adaptation was achieved with a background illumination of 30 cd/m^2^ starting ten minutes before photopic recordings. Five responses were averaged with inter-stimuli interval of five (0.0003-0.3 cd.s.m-2) or 17 seconds (1-10 cd.s.m-2). For reversal of the anesthesia, animals were intraperitoneally injected with 10 mg/kg bw atipamezole hydrochloride (Orion Pharma, Antisedan®), placed in a warmed cage and monitored until fully awake.

#### Fundus imaging and Fluorescein angiography

Fundus images were taken with a Phoenix MICRON^®^ IV after animal anesthesia, pupil dilation and eye moisturizing as described above. For fluorescein angiography, animals were recorded 10-15 min after intraperitoneal injection of 10 µl/g bw 1% fluorescein (Alcon, H12588-0113, diluted in isotonic solution). Anesthesia was reversed as described above.

### Differentiation of RPE cells

#### Cyst generation

RPE cells were differentiated as described and patented in.^13,30–32^ Briefly, after removal of undifferentiated colonies, cells were washed twice with warm DMEM/F12 (Gibco, 31330-038) and incubated with Dispase for 7 min at 37°C. The Dispase was gently washed away twice with warm DMEM/F12, then warm N2B27 medium was added and the colonies scraped into a 15 ml Falcon tube, then centrifuged for 1 min at 92 g, room temperature (RT). The pellet containing roughly 50-100 µm large cell aggregates was resuspended in N2B27 medium (DMEM/F12+GlutaMAX (Gibco, 31331-028) and neurobasal medium (Gibco) at a 1:1 ratio, 0.5% GlutaMAX Supplement (Gibco, 35050061), 0.5x B-27 supplement (Gibco, 17504044), 0.5x N-2 supplement (Gibco, 17502048), 0.1 mM β-mercaptoethanol (Gibco, 11528926), 1x Penicillin-Streptomycin (Gibco, 15140-122)), placed on ice and gently mixed 2-3 times with cyst Matrigel (Corning, 354234; protein concentration 9-10 mg/ml) using P1000 wide bore tips. Per 1 mm pellet height, 20 µl N2B27 medium and 200 µl Matrigel were used. 100 µl cyst Matrigel with cells were then transferred to a glass bottom dish (Mattek, P35G-1.5-14-C), spreading the mix over the majority of the glass area and allowing the matrix to gel for 10 min at 37°C, the first 5 min upright, the second 5 min inverted. Then, N2B27 medium was added and exchanged every two days.

#### Plating

To release the cysts formed by day (D) 5 (embedding is counted as D0), 5 Matrigel drops were transferred to 1 ml cold Cell Recovery Solution (BD Biosciences, 354253) in a 15 ml Falcon, incubated on ice for 25 min with careful inversion every 2 min. After allowing the cysts to sink for 5 min, removal of the Cell Recovery Solution, a wash with PBS and 1 min centrifugation at 17 g, RT, the pellet was resuspended using a wide bore P1000 tip in TrypLE (GIBCO, 12563-011) diluted 1:6 in PBS. After incubation at 37°C with constant movement for 12 min or when cysts were broken up into single cells, dissociation was stopped by mixing with Soybean Trypsin Inhibitor solution (Sigma, T6522; stock at 10 mg/ml in distilled water) at 1 mg/ml and centrifugation for 2 min at 120 g, RT. The pellet was resuspended in N2B27 medium, filtered through a 40 µm cell strainer (BS Biosciences, 352340) and 300.000 cells per insert were plated onto Matrigel-coated Transwells (Corning, 3470; coating: incubation for 1 h at room temperature or 15 min at 37°C with Growth Factor-Reduced Matrigel (Corning, 356230) diluted 1:20 in DMEM/F12 (Gibco, 31330-038), using 105 µl per transwell). The lower chamber was filled RPE medium (DMEM+GlutaMAX (Gibco, 31966-047), 20% Knockout Serum Replacement (Gibco, β10828010), 0.5x L-Glutamine (Life Technologies, 25030-024), 1x MEM NEAA (Gibco, 11140-035), 0.1 mM ß-mercaptoethanol (Gibco, 11528926), 1x Penicillin-Streptomycin (Gibco, 15140-122), 10 ng/ml Activin A (R&D Systems, 338-AC; stock at 10 µg/ml in 4 mM HCl)) and 24 h after plating as well as every 2-3 days thereafter, medium in both upper and lower chamber was exchanged to fresh RPE medium. At this stage, cells were considered passage (P) 0.

#### Passaging

At 21 days after plating to P0 (protocol D26), cells were passaged to P1 and another 21 days later (protocol D47) to P2. For passaging, cells were washed and incubated with Trypsin in both upper and lower chamber for 10 min at 37°C, released from the transwells by pipetting up and down ∼12 times with a 100 µl tip and collected in warm RPE medium with 1 mg/ml Soybean Trypsin Inhibitor. The pellet of 3 min centrifugation at 800 rpm was resuspended in fresh RPE medium, filtered through a 40 µm cell strainer and plated onto fresh Matrigel-coated transwells at 600.000 cells per insert for P1 and 300.000 cells per insert for P2. Medium in both upper and lower chamber was changed every 2-3 days.

#### Cell release for sequencing and transplantation

For analysis and transplantation, P1 and P2 cells were retrieved from the inserts as for passaging. After centrifugation, cells were resuspended in RPE medium, filtered, counted and placed on ice until further use. For transplantation, cells were centrifuged again for 2 min at 200 g and resuspended at a concentration of 50,000 or 100,000 cells/µl in MACS buffer (PBS (Gibco, 10010-023), 2 mM EDTA, 0.5% w/v BSA) supplemented with 0.1 mg/ml DNaseI (Sigma, D5025-150KU).

### Sequencing

#### RNA preparation

To collect cells for sequencing, cells were detached from the transwell and processed as described above. 150,000 cells were then lysed in 200 µl RLT buffer (Qiagen, 79216), flash frozen and stored at -70°C until processing. RNA was isolated using the Qiagen RNeasy Mini RNA isolation kit (Qiagen, 74104) as instructed. Briefly, lysed samples were thawed on ice, the RNA precipitated with an equal volume of 70% ethanol and the sample transferred to an RNeasy spin column. After drying by centrifugation, a wash with Buffer RW1 and an on-column DNase digestion, the column was washed again with Buffer RW1, then Buffer RPE and finally RNA was eluted in water. All centrifugation steps were carried out for 30 s at 12,000 g. RNA integrity and concentration was measured using an Agilent 2100 Bioanalyzer System with the RNA 6000 Pico Kit (Agilent Technologies, 5067-1513).

#### Library preparation and Sequencing

mRNA was isolated from 100 or 500 ng total RNA by poly-dT enrichment using the NEBNext Poly(A) mRNA Magnetic Isolation Module (NEB) according to the manufacturer’s instructions. Samples were then directly subjected to the workflow for strand-specific RNA-Seq library preparation (Ultra II Directional RNA Library Prep, NEB). For ligation, the NEB Next Adapter for Illumina of the NEBNext Multiplex Oligos for Illumina Kit was used at a final concentration of 0.004 or 0.016 µM. After ligation, adapters were depleted by XP bead purification (Beckman Coulter), adding bead solution at a ratio of 0.9:1 to the samples (0.9X). Unique dual indexing was done during the following PCR enrichment (12 or 14 cycles) using amplification primers carrying the same sequence for i7 and i5 index (i5: AAT GAT ACG GCG ACC ACC GAG ATC TAC AC NNNNNNNN ACA TCT TTC CCT ACA CGA CGC TCT TCC GAT CT, i7: CAA GCA GAA GAC GGC ATA CGA GAT NNNNNNNN GTG ACT GGA GTT CAG ACG TGT GCT CTT CCG ATC T). After two more XP bead purifications (0.9X), libraries were quantified using the Fragment Analyzer (Agilent). Libraries were sequenced on an Illumina NovaSeq 6000 in 100 bp paired-end mode to an average depth of 40 million fragments per library.

#### Bioinformatics – adapted version

FastQC (http://www.bioinformatics.babraham.ac.uk/) was used to perform a basic quality control of the resulting sequencing data. Fragments were aligned to the human reference genome GRCm38 with support of the Ensembl 98 splice sites using the aligner gsnap (v2021-12-17).^33^ Counts per gene and sample were obtained based on the overlap of the uniquely mapped fragments with the same Ensembl annotation using featureCounts (v2.0.1).^34^ Raw counts of technical replicates were averaged and low count genes filtered out. For PCA analysis, data was transformed using the variance stabilizing transformation function of the DESeq2 package (v1.42.1)^35^ and batch-corrected using the limma package (v3.58.1).^36^ To identify differentially expressed genes (DEGs) between cell passage stages, counts were fitted to the negative binomial distribution and genes were tested between conditions using the Wald test of DESeq2 package including the batch as a covariate. Genes with a maximum of 5% false discovery rate (padj ≤ 0.05) were considered as significantly differentially expressed.

For GO term analysis DEGs were ordered by decreasing log2 fold change and used as input for an ordered query with gprofiler2 (package v0.2.3, version e111_eg58_p18_f463989d) with g:SCS multiple testing correction method applying significance threshold of 0.05.^37–39^ To limit the number of resulting terms, only DEGs upregulated in P1 cells with a log2 fold change > 3 were used, while no log2 fold change threshold was applied for DEGs upregulated in P2 cells. Using the gprofiler2 web application, resulting GO terms were grouped according to their connectivity in the GO context tab, with each context group bearing the name of the contributing term with the lowest p-value. Context groups were again thematically aggregated and z-scores of the regularized log-transformed data of the DEGs contributing most to each aggregate were plotted as heatmaps.

### Cell Surface Marker Screening

P1 RPE cells derived from CRTD1 iPSC were screened for expression of 371 antibodies directed against cell surface markers using the flow cytometry-based MACS® Marker Screen, anti-human, version 01 (Miltenyi Biotec, 130-110-055) according to the manufacturer’s instructions. Undifferentiated CRTD1 and MIL001-A pre-labelled with the CellTrace Violet Cell Proliferation Kit (Life Technologies, C34557) were pooled together with P1 RPE and used as internal control and reference. Data acquisition was done on MACSQuant Analyzer 10 (Miltenyi Biotec, 130-096-343). MACS Quantify Software (Miltenyi Biotec) was used for analysis. Debris was excluded based on scatter properties. Dead cells were excluded based on propidium iodide staining. RPE cells were gated based on negative CellTrace Violet signal. Sidescatter (SSC) signal, as proxy for pigmentation, was used to gate target (pigmented) and non-target cells (unpigmented). Full gating strategy and raw data is available as supplementary data (Document S4). Combinations of candidate markers for pigmented and unpigmented cells were validated on independent P1 RPE specimen.

For cell surface marker screening at Miltenyi Biotec Co., iPSC-derived P1 RPE differentiated by the Ader lab using the standard protocol as described above were sent as frozen vials and used for screening directly after thawing in cultivation medium (DMEM/F12, GlutaMAX (Gibco, 10565018), 20% Knockout Serum Replacement (Gibco, 10828028), 0.8 mM L-Glutamine (Fagron, 700846), 1x MEM NEAA (Lonza, 13-114E), 0.1 mM ß-mercaptoethanol (Sigma, M7522), 1x Penicillin/Streptomycin (Lonza, 17-602E), 10 ng/ml Activin A (Miltenyi, 130-115-010)) freshly supplemented with 10 µM StemMACS Y-27632 (Miltenyi, 130-106-538) and 1.5% Enzyme Mix 2 from the Neural Tissue Dissociation Kit – Postnatal Neurons (Miltenyi, 130-094-802). Cell count was determined using the flow cytometer MACSQuant Analyzer 10 (Miltenyi, 130-096-343) and viability assessed with Propidium iodide (Miltenyi, 130-093-233).

### Cell Surface Marker Sorting

#### Live cell staining

1×10^6^ cells were stained in 100 µl cultivation medium or RPE medium (see RPE differentiation protocol) supplemented with 10 µM StemMACS Y-27632 and 1.5% Enzyme Mix 2. P1 cells were stained with CD54-APC (Miltenyi, 130-121-342) and PSA-NCAM-PE (Miltenyi, 130-117-394) according to data sheet. For washing steps, supplemented medium was used and cells were centrifuged 5 min at 200 g.

#### Sorting using the Tyto platform

Cells in supplemented medium were sorted at a maximum concentration of 1×10^6^ cells/ml in the MACSQuant Tyto cell sorter (Miltenyi, 130-103-931) using a Tyto HS cartridge (Miltenyi, 130-121-549). Before sorting, the cartridge was primed using the priming fixture according to protocol and 200-400 µl supplemented medium. Cells were filtered over a 20 µm pre-separation filter (Miltenyi, 130-101-812) into the Tyto cartridge via a 10 ml syringe with luer lock tip (Braun, 4617100V-02). For later analysis, an unsorted sample was taken from the input chamber of the cartridge using a 200 µl gel loading tip (VWR, 732-2671). The sort was performed in the MACSQuant Tyto cell sorter using software 1.0.0.

After the sort, fractions were retrieved from sort (target cell-enriched fraction, ENR) and non-sort (target cell-reduced fraction, RED) chambers using 200 µl gel loading tips and both collection chambers were washed with supplemented medium. Samples from all fractions were analyzed with a MACSQuant Analyzer 10 (Miltenyi, 130-096-343). Viability was determined using propidium iodide (Miltenyi, 130-093-233) or DAPI. MACS Quantify Software (Miltenyi) was used to analyze the sort performance.

#### Sorting using the Aria III platform

For sorting of cells with the BD FACSAria III (BD Biosciences), stained cells were resuspended in MACS/DNaseI buffer (PBS (Gibco, 10010-023), 2 mM EDTA, 0.5% w/v BSA, 0.1 mg/ml DNaseI (Sigma, D5025-150KU)) and an UNS fraction was set aside. The gating strategy included gating out of debris (FCS-A v. SSC-A), doublets (FSC-A v. FSC-H, SSC-H v. SSC-W) and dead cells based on DAPI uptake. Sorting was performed at 4°C, using a 100 µm nozzle.

#### Cell preparation for transplantation

For transplantation, UNS, ENR and RED fractions obtained from both Tyto or Aria III were centrifuged for 2 min at 200 g and resuspended at a concentration of 50,000 cells/µl in MACS/DNaseI buffer.

#### Plating of sorted cells

24-well transwell plates (Merck, CLS3470) were coated with Matrigel (Corning, 354234), diluted 1:20 in D-MEM/F-12 (Gibco, 11320-074) on the previous day, using 120 µl of coating solution per well. Coating solution was removed before plating 300.000 cells per 24-Transwell. Duplicates from unsorted cells (UNS), positive sorted (ENR) and negative sorted (RED) fractions were plated. Cells were plated in upper part of the transwell in 200 µl cultivation medium containing 10 µM StemMACS Y-27632. The lower part was filled with 600 µl cultivation medium. Medium was changed every 3-4 days (cultivation medium without Y-Compound).

### Transplantation

Subretinal transplantation was performed as described in detail in.^40^ Briefly, mice were anesthetized by intraperitoneal injection of 30 µg/kg bodyweight (bw) Ketamine (medistar, 04-03-9264/23) and 1 µg/kg bw Medetomidine hydrocholoride (Orion Pharma, Domitor®) in sterile isotonic solution (Fresenius Kabi, B306175/03), followed by application of dilating eye drops containing 2.5% phenylephrine and 0.5% tropicamide (University Clinic Pharmacy Dresden). After 1-2 min, eye drops were replaced with clear moisturizing eye gel (Bausch+Lomb, Vidisic) and a coverslip placed on top for visibility through the operating microscope. The mouse head was fixed with a stereotactic head holder (World Precision Instruments, 505242). Using a sterile 30G needle, a small incision was made below the ora serrata opposite to the transplantation target region. 1 µl cell suspension and 0.2 µl air were loaded into a 5 µl Hamilton syringe (Hamilton, 065-7634-01) with a blunt 34-gauge needle (Hamilton, special request; 12 mm length). The Hamilton syringe was mounted onto a micromanipulator (H. Saur Laborbedarf, M1) and the needle steered through the pre-made incision, until it reached the retina at the target region. There, the air was used to generate a pre-bleb, followed by the subretinal release of the cell suspension and slow needle retraction for self-sealing of the injection site. Transplantation success and possible bleeding was noted and only eyes with an estimated ≥80% of cell suspension successfully placed subretinally were considered in the experiments.

Immediately after transplantation, 80 µg filtered triamcinolone acetonide (University Clinic Pharmacy Dresden; 80 µg/µl in NaCl, preservative-free, filtered through 35 µm mesh) were injected vitreally through the same incision using a handheld 10 µl Hamilton syringe (Hamilton, 065-7635-01). Then, animals were released from the head holder, intraperitoneally injected with 10 mg/kg bw atipamezole hydrochloride (Orion Pharma, Antisedan®), placed in a warmed cage and monitored until fully awake.

### Immunohistochemistry

#### Sample collection and preparation

Cultured cells were briefly washed in PBS and fixed by 10 min incubation in 4% paraformaldehyde (PFA) in PBS at room temperature (RT). Cell-carrying membranes were cut out and either stained directly for the en face view or embedded in NEG50 (Richard-Allan Scientific, 6502), frozen and cryosectioned with the Epredia^TM^ CryoStar^TM^ NX70 (Leika Mikrosysteme, Wetzlar, Germany) in 12 µm sections. Sections were air-dried at 37°C for 20 min and stained directly.

Experimental animals were anesthetized with isoflurane and decapitated. Eyes were enucleated immediately and placed in 4% PFA in PBS for fixation over the course of 1 hour at 4°C. Samples for cryosectioning were then dissected, removing lens and cornea, and cryoprotected overnight at 4°C in 30% w/v sucrose solution in PBS. After embedding in NEG50 and freezing, eyes were serially sectioned at 12 µm thickness with the CryoStar^TM^. Sections were air-dried at 37°C for 20 min, stored at -70°C and allowed to come to RT before staining.

For flatmounts, fixed eyes were dissected removing lense, cornea and ocular muscle tissue. Then, the retina was detached by cutting right below the ora serrata and incisions were made, cutting the eyball into petals to allow later flattening. Flatmounts were stained immediately after collection.

#### Staining

All immunohistochemistry samples were blocked in blocking solution (0.3% Triton X-100, 1% bovine serum albumin, 5% normal goat serum) for 1 h at RT and incubated overnight at 4°C with primary antibodies idluted in blocking solution. After washing with PBS, samples were incubated for 1-2h in PBS containing secondary antibodies and 0.2 µg/ml 4ʹ,6-diamidino-2-phenylindole (DAPI). Samples were washed again with PBS and then distilled water before mounting using AquaPolymount (Polysciences, 18606).

### Imaging and Image analysis

Stained cryosections were imaged with an Axio Imager.Z1 microscope with the ApoTome.2 enabled (Zeiss). Unless noted otherwise, single images shown are maximum intensity projections of z-stacks with 5-6 µm thickness. Flatmounts were imaged using the tiling setup of the Observer.Z1 microscope with the ApoTome.2 enabled (Zeiss), with z-stack depths of ca. 35 µm to ensure proper graft visualization despite a wavy surface.

For graft size and monolayer coverage quantification, every fourth serial section containing human graft cells stained with an antibody against human nuclei was imaged with an AxioScan.Z1 (Zeiss). The graft length per section was measured using the active curve tool, differentiating between monolayer and cluster graft regions. Monolayer percentage was calculated dividing the sum of monolayer length measurements by the sum of total graft measurements. The retinal coverage was calculated by diving total graft length by total retina length. For total retina length, a standardized value was used, averaged from three eyes in which retina length was measured after imaging serial sections of the entire eye.

Flatmount analysis with REShAPE was carried out using the code published on github (see key resources table), adapted to run on the local server infrastructure.

### Electron Microscopy (EM)

Eyes to be analyzed through TEM were collected from the animals as described for immunohistochemistry. After fixation in 4% PFA in 100 mM phosphate buffer for 20-24 h, cornea, iris, lens and muscle tissue were cut away and the globe was halved along the vertical axis (temporally located scar tissue from the incision below the ora serrata during transplantation allowed for proper orientation). The nasal portion containing the area of interest was further processed for EM as described in.^41,42^

#### Transmission Electron Microscopy (TEM)

For TEM, the samples were further dissected to small pieces (0.5-1 mm) and postfixed at least overnight in Modified Karnovsky’s Fixative (2% glutaraldehyde, 2% paraformaldehyde in 100 mM phosphate buffer) at 4°C. Samples were further postfixed and contrasted following the OTO-procedure by incubating them in a 2% OsO_4_ solution (2% OsO4, 1.5% potassium ferrocyanide, 2 mM CaCl2 in water), then in 1% aqueous thiocarbohydrazide, followed by another osmium step in 2% aqueous OsO_4_ with washes in water between all incubation steps. Finally, samples were *en-bloc* contrasted with 1% uranyl acetate in water, and washed several times in water. Contrasted samples were then dehydrated in a graded ethanol series from 30% to 100% on a molecular sieve (30%, 50%, 20 min each, 70%, 90%, 95%, 3x 100%, 30 min each), before embedding in EMBed 812 through stepwise infiltration (25%, 50%, 75% resin 1h each, pure resin overnight, pure resin 5 h) and curing at 65°C overnight. Semithin sections were made and stained with toluidine blue/borax to find the graft region. For TEM, 70 nm ultrathin sections were cut with a Leica UC6 ultramicrotome using a diamond knife (Diatome), collected on formvar-coated slot grids and contrasted once more with lead citrate^43^ and uranyl acetate. Samples were then imaged with a Jeol JEM1400 Plus transmission electron microscope (camera: Ruby, Jeol) running at 80 kV acceleration voltage.

#### Correlative light and electron microscopy (CLEM)

CLEM samples were stored in 1% PFA in PBS, 4°C until selection and dissection of the ROIs under a fluorescence stereo microscope (Leica MZ10F). Selected ROIs were subjected to progressive lowering of temperature embedding in the methacrylate resin Lowicryl K4M.^44^ Samples were dehydrated in a graded ethanol series (30%, 50% ethanol/water 30 min each at 4°C, 70%, 90% at -20°C, 96% 1h at -35°C, 2x 100% on molecular sieve at -35°C, 1h each. The last dehydration steps at -35°C and the following infiltration steps in Lowicryl K4M (1:2 resin:ethanol, 1:1, and 2:1 for 1-2 h each, pure resin overnight, pure resin 5-6 h, all at -35°C) were conducted in a Leica AFS2 freeze substitution unit. Samples were placed and oriented in flat embedding forms and cured by UV-irradiation at -35°C for 48 h. 70 nm ultrathin sections were cut using a Leica UC6 ultramicrotome with a diamond knife, mounted onto formvar-coated mesh grids and immunolabeled as described previously ^45–47^. Briefly, sections were blocked with 1% BSA in PBS, sequentially incubated with the primary antibody (rabbit anti-GFP, TP401 from Torrey Pines, 1:100) followed by protein A 10 nm gold (Utrecht CMC, 1:50), a fluorophore-conjugated secondary antibody (goat-anti-rabbit Alexa488, 1:100) for immunofluorescence, and DAPI to counterstain the nuclei. In between incubations, sections were washed with PBS, and labeled grids were then mounted in glycerol-water (1:1) before fluorescent imaging with a Keyence Biozero 8000 fluorescence microscope. Grids chosen for TEM were demounted, washed thoroughly in water, contrasted with uranyl acetate, dried and imaged as described above for TEM samples.

## QUANTIFICATION AND STATISTICAL ANALYSIS

All values are represented as mean ± SEM (standard error of the mean) unless otherwise stated, with n = number of retinas examined, where appropriate. Statistical significance was assessed using RStudio and denoted as p < 0.05 = *; p < 0.01 = **; p < 0.001 = ***. Samples were tested for outliers using Grubbs or Dixon test, and outliers were removed before further analysis. Sample distributions were tested for normality using the Shapiro-Wilk’s test, and for homoscedacity using Bartlett’s and Levene’s test. Appropriate statistical tests were then applied including: unpaired t-test, and Kruskal-Wallis with pairwise Wilcoxon test. Figures were generated in Affinity Designer.

## Ressource Availability

### Lead contact

Requests for further information and resources should be directed to and will be fulfilled by the lead contact, Prof. Marius Ader.

### Materials availability

This study did not generate new unique reagents.

### Data and code availability

Single-cell RNA-seq data have been deposited at GEO at GEO: accession number and are publicly available as of the date of publication. Any additional information required to reanalyze the data reported in this paper is available from the lead contact upon request.

## Acknowledgements

The authors would like to thank the Stem Cell Engineering, Flow Cytometry, Animal facility, Light Microscopy and Genome Center at the CRTD/CMCB, Technische Universität Dresden for assistance in iPSC cell culture, cell sorting, mouse maintenance, microscopy and sequencing/bioinformatic analysis, respectively. Additional technical and laboratory assistance was provided by Jochen Hentschel and Patrick Schäfer and IT assistance by Alexandre Mestiashvili. This work was supported by the Bundesministerium für Bildung und Forschung (BMBF): ReSight - 01EK1613A to M.A., 01EK1613C to S.K.), and Foundation Fighting Blindness (FFB Award Number: TA-RM-0522-0824-TUD-TRAP to M.A.).

## Author contributions

K.T. & S.J.G.: Conceptualization, Methodology, Validation, Formal Analysis, Investigation, Writing – Original Draft, Writing – Review and Editing, Visualization, Supervision, and Project Administration. K.S.: Conceptualization, Methodology, Validation, Formal Analysis, Investigation, Visualization, and Project Administration. T.A. & L.M.: Validation and Investigation. J.H.: Validation, Formal Analysis, Investigation, Visualization, Supervision, and Project Administration. L.R.L, A.L., N.C.: Investigation and Resources. M.C.: Formal Analysis and Investigation. S.A.: Methodology and Resources. S.F., A.D., A.F.: Resources. T.K., A.P., U.A.F.: Formal Analysis. A.K.: Formal Analysis, Investigation, Resources and Supervision. H.J.: Formal Analysis, Resources, Writing – Original Draft, Writing – Review & Editing, and Supervision. S.K.: Formal Analysis, Resources, Writing – Original Draft, Writing – Review & Editing, Supervision, Project Administration, and Funding Acquisition. M.A.: Conceptualization, Methodology, Writing – Original Draft, Writing – Review and Editing, Supervision, Project Administration, and Funding Acquisition.

## Supplemental information

Document S1. Figures S1-S5.

Document S2. Excel file containing raw counts, differentially expressed genes, GO terms and GO term groups obtained from bulk sequencing of P1 versus P2 cells, related to Figure 1.

Document S3. PDF containing screening results of all cell surface antibodies tested with the MACS Marker Screen, related to Figure 4.

## Supplemental figures and legends

**Figure S1:**
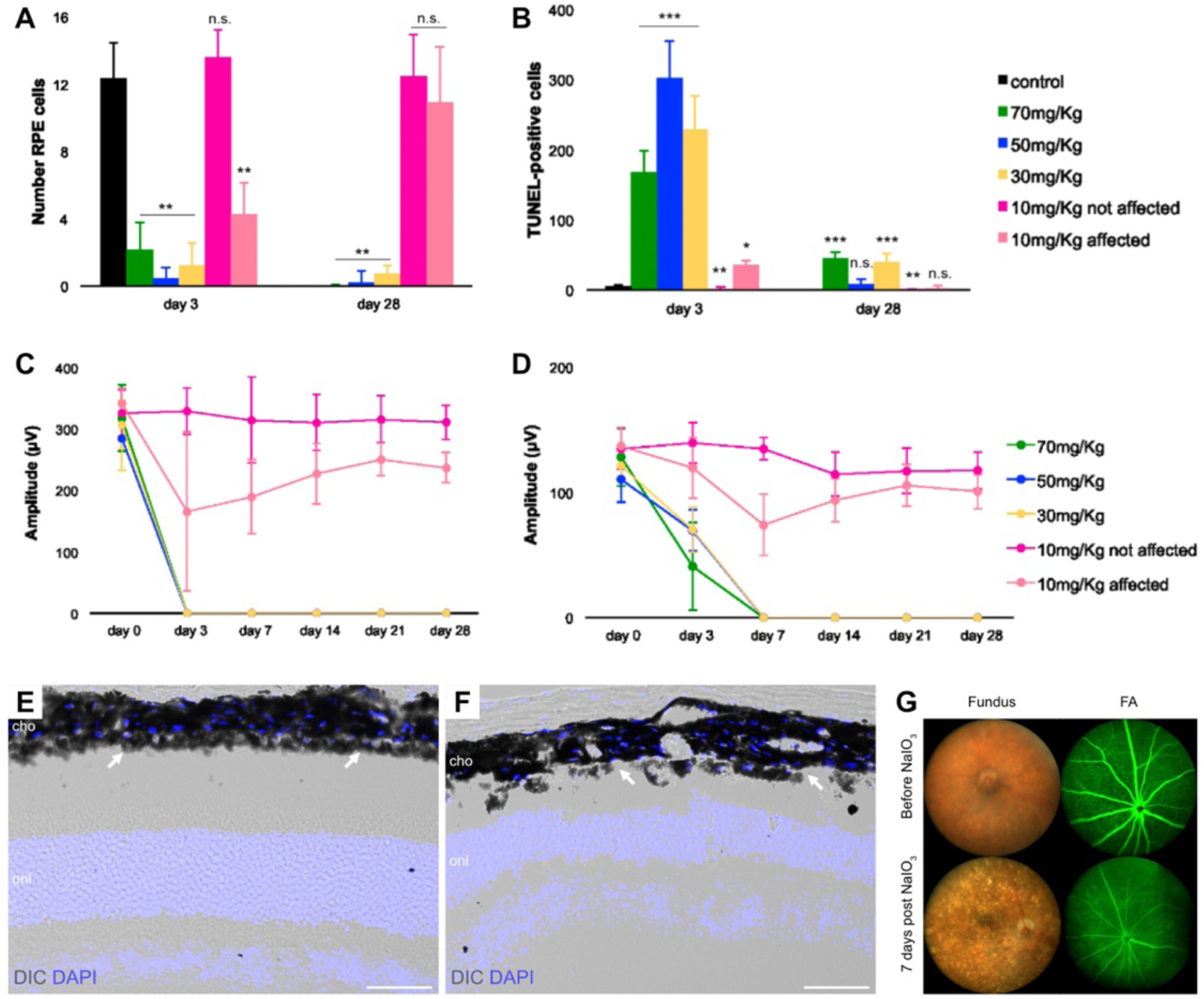
Injection of 30 mg/kg bodyweight NaIO_3_ suffices to cause RPE degeneration and loss of visual function. (A) While a low dosage (10 mg/kg bodyweight (bw) NaIO_3_) had variable effects on RPE cell number, an injection of the intermediate dosage of 30 mg/kg bw NaIO_3_ was sufficient to cause RPE degeneration to a similar degree as high dosages (50 and 70 mg/kg bw NaIO_3_) at day 3 and day 28. Control: n=4; 70mg/Kg - day 3: n=14, day 28: n=7; 50mg/Kg - day 3: n=5, day 28: n=5; 30mg/Kg - day 3: n=6, day 28: n=7; 10mg/Kg not affected - day 3: n=4, day 28: n=5; 10mg/Kg affected - day 3: n=1, day 28: n=4. n: individual mice. (B) In intermediate and high dosage groups, RPE cell death occurred quickly after NaIO_3_ injection (day 3), and was strongly reduced at 4 weeks after injection (day 28). n: see A. (C, D) Visual function of both, rods and cones, was compromised after injection of intermediate and high NaIO_3_ dosages, as shown by the amplitudes of the scotopic (C, 0.003 cd*s*m-2) and photopic (D, 10 cd*s*m-2) b-waves over time. 70mg/Kg - day 0: n=14, day 3: n=14, day 7: n=14, day 14: n=14, day 21: n=11, day 28: n=7; 50mg/Kg - day 0: n=29, day 3: n=5, day 7: n=5, day 14: n=5, day 21: n=5, day 28: n=5; 30mg/Kg - day 0: n=7, day 3: n=11, day 7: n=8, day 14: n=3, day 21: n=8, day 28: n=7; 10mg/Kg - day 0: n=14, day 3: n=14, day 7: n=9, day 14: n=9, day 21: n=9, day 28: n=9. n: individual mice. (E, F) DAPI staining and DIC imaging at 7 days after injection shows a normal RPE (arrows) and neuroretinal phenotype in the control group (E, saline injection). The RPE in animals injected with 30 mg/kg bw NaIO_3_ (F) is no longer a continuous monolayer, but patchy and unorganized (arrows). n = 4 biological replicates. (G) Injection of 30 mg/kg bw NaIO_3_-injection causes the development of highly reflective speckles in the previously homogenous fundus image. Fluorescein angiography (FA) further shows the change in blood-retina barrier permeability as a result of the NaIO_3_-injection, with the fluorescein-bright inner plexus vessels clearly standing out from the dark retina initially. At 7 days post NaIO_3_ injection, the contrast is strongly reduced due to fluorescein leakage through the destroyed RPE layer. n = 4 biological replicates. Scale bars: 50 µm (E,F). cho, choroid; onl, outer nuclear layer; FA, fluorescein angiography. Data are represented as mean ± SEM. Statistics: *p < 0.05, **p < 0.01, ***p<0.001 by one-way ANOVA (A-D).

**Figure S2:**
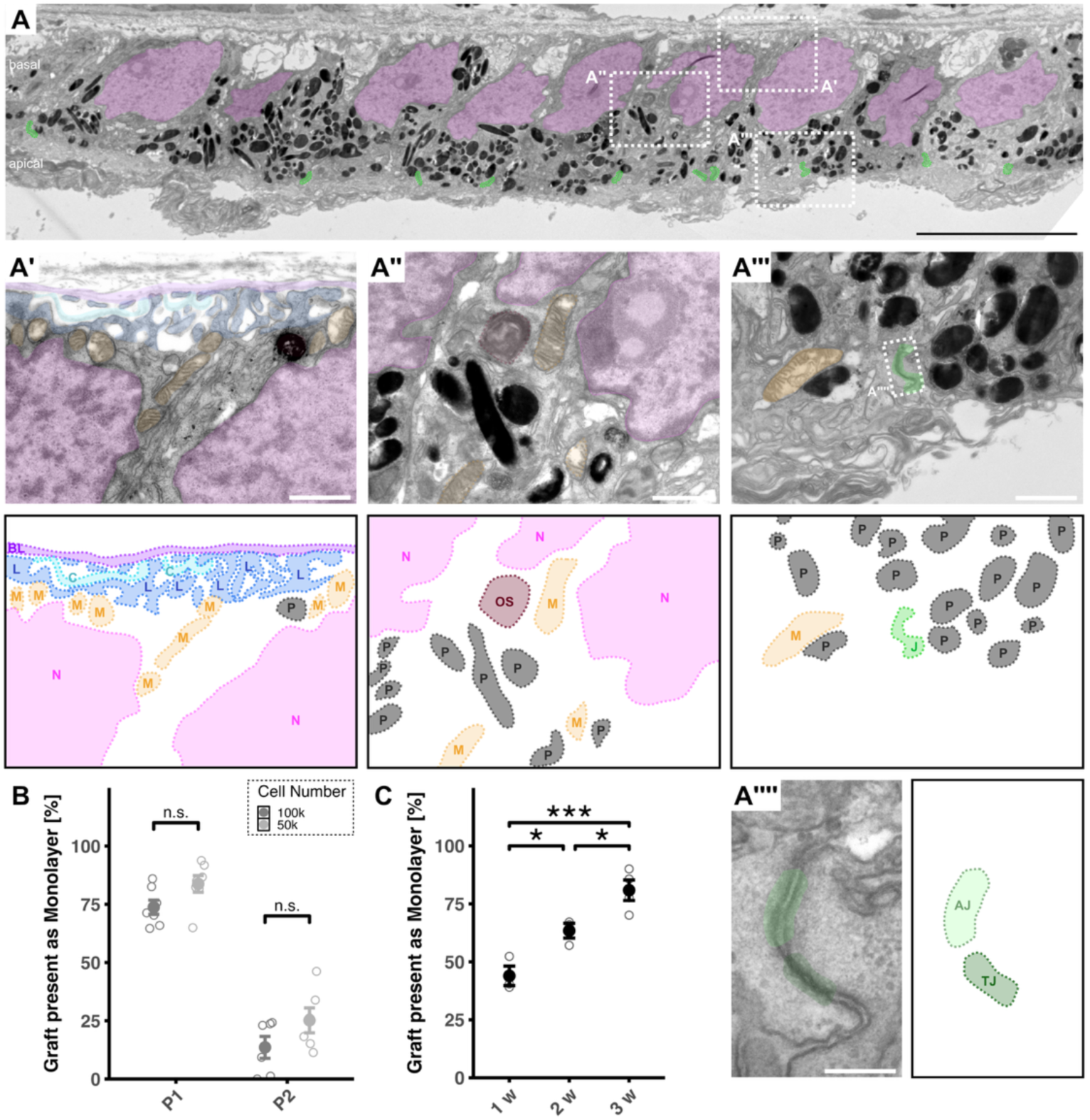
RPE-derived monolayer formation and characterisation (A) Transmission electron microcopy (TEM) image presented in figure 2E with higher magnifications showing a basal (A’), an intermediate (A’’) and an apical (A’’’) portion of the cells with RPE-characteristic features. Apical junctions are further highlighted per green overlay in A and one junctional complex from A’’’ is magnified in A’’’’. TEM images A’-A’’’’ are depicted with coloured overlays, with the labelled overlay additionally positioned next to the respective image for better visibility. n = 3 biological replicates. (B) Monolayer percentage data presented in figure 2G, replotted to show differences between eyes having obtained 50,000 or 100,000 cells. While the difference is not statistically significant, transplantation of 50,000 cells tends to generate a higher monolayer percentage than transplantation of 100,000 cells. P1 100k: n=7; P1 50k: n=7; P2 100k: n=6; P2 50k: n=6. All datapoints within one group represent biological replicates. (C) Monolayer percentage increases over time. Here, empty points represent graft quantification of a single central section per eye, filled points their average. n=3, 3, and 4 biological replicates in groups 1w, 2,w, and 3w, respectively. Scale bars: 10 µm (A), 1 µm (A’-A’’’), 200 nm (A’’’’). Data are represented as mean ± SEM. AJ, adherens junctions; BL, basal lamina; C, collagen; J, junctional complexes; L, basal labyrinth; M, mitochondrium; N, nucleus; OS, phagocytosed outer segment; P, pigment granule; TJ, tight junctions; w, weeks. Statistics: p < 0.05, **p < 0.01, ***p<0.001 by two-sided Student’s t-test (B) and one-way ANOVA with Tukey’s post hoc test (C).

**Figure S3:**
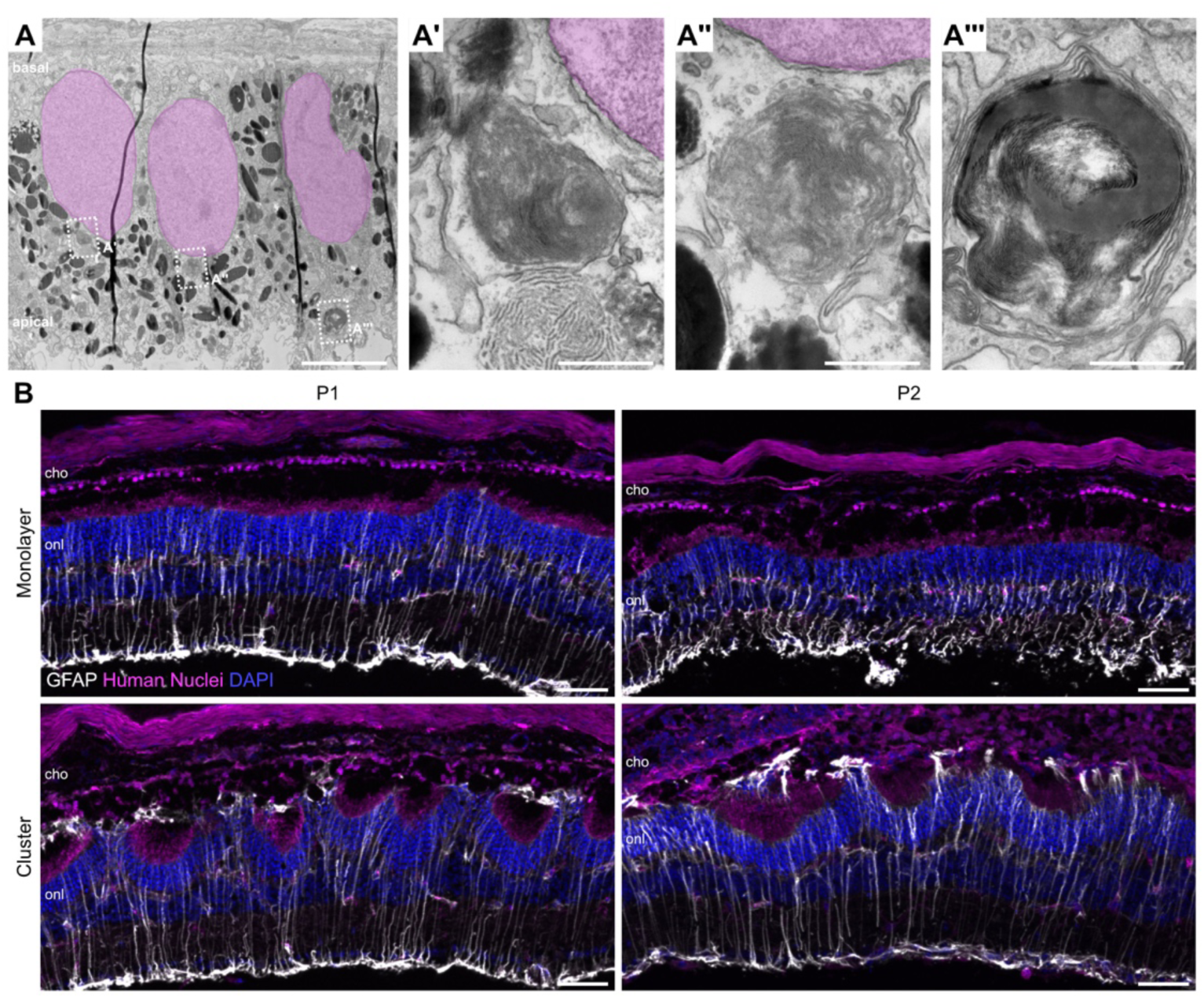
Supplementary data for Figure 3. (A) Transmission electron microcopy images of RPE-derived monolayers show outer segment particles within transplanted cells. A’-A’’’ show boxed areas in A magnified. n = 3 biological replicates. (B) GFAP staining found throughout all neuroretinal layers shows reactive Müller Glia irrespective of RPE passage transplanted or graft cell conformation as monolayer or cluster. n = 5 biological replicates. Scale bars: 50 µm (B), 5 µm (A) and 500 nm (A’-A’’’). cho, choroid; onl, outer nuclear layer.

**Figure S4:**
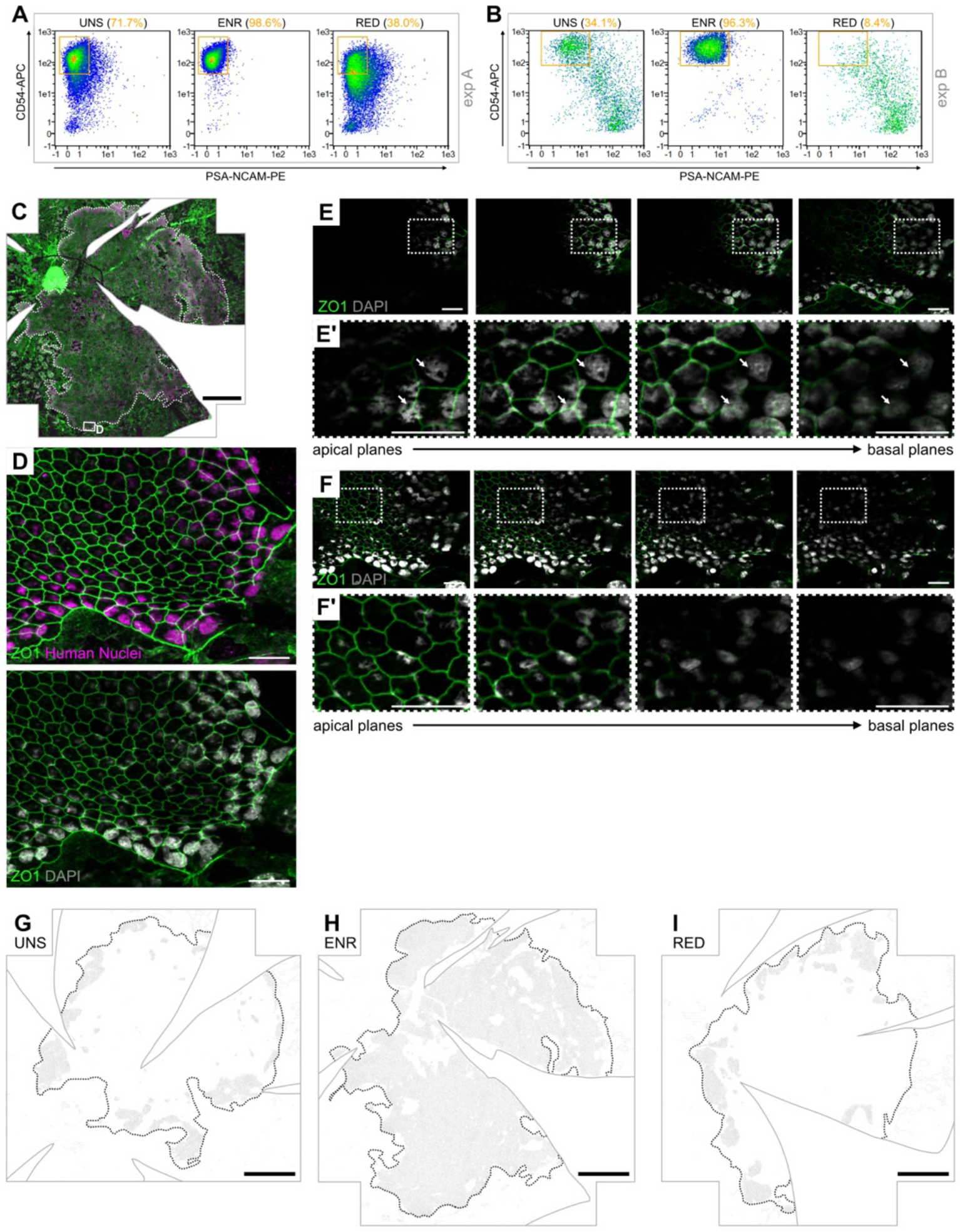
Supplementary data for Figure 5. (A, B) Flow cytometry re-analysis of unsorted (UNS), CD54^+^/PSA-NCAM^-^-enriched (ENR) and -reduced (RED) fractions before transplantation in experiment A (exp A, see also figure 5I) and experiment B (exp B, see also figure 5J). Orange numbers show purity of the target cell population in each fraction, with a high percentage of CD54^+^/PSA-NCAM^-^ cells in exp A (71.7%) and a lower percentage of CD54^+^/PSA-NCAM^-^ cells in exp B (34.1%). (C) Flatmount of the ENR graft as presented in figure 5B, here with an additional region framed and magnified in D-F’. (D) Immunohistochemistry shows higher visibility of the Human Nuclei and DAPI staining towards the edge of the human graft. (E, F) Single z-planes of the maximum intensity projection in D, with E’ and F’ showing the boxed regions in E and F enlarged, highlighting the obstruction of nucleus visibility through the apically located pigment (arrows). DAPI brightness is increased in F and F’ for better visibility. (G-I) REShAPE cell outlines of the flatmounted eyecups shown in figures 5A-C, with human graft areas denoted by dotted lines. Eyes received transplantations of cell suspensions that were either unsorted (G), enriched (H), or reduced (I) in CD54^+^/PSA-NCAM^-^ cells. Scale bars: 500 µm (C, G-I), 20 µm (D-F’).

**Figure S5:**
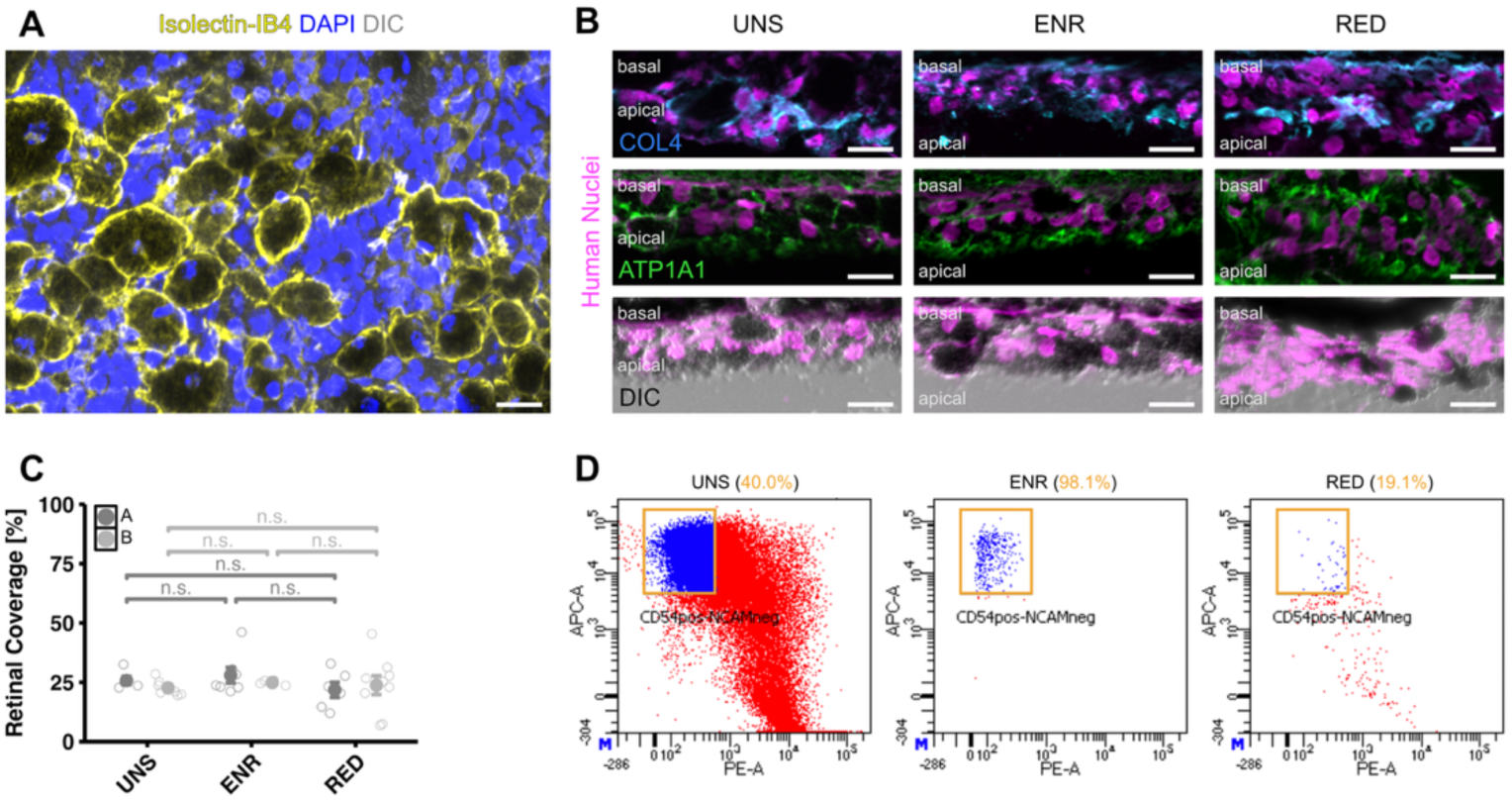
Supplementary data for Figure 5, continued. (A) Pigmented, spherical clumps likely represent macrophages as they are larger than the average RPE cells, contain a single nucleus and stain positive for isolectin IB_4_. Here, pigmented clumps are shown stained on top of the outer nuclear layer of a retinal flatmount opposing an unsorted human RPE graft. n = 3 biological replicates. (B) Clusters formed by all graft fractions lack polarization, as basal marker COL4 and the apical marker ATP1A1 are expressed chaotically throughout the cluster. Similarly, the pigment (dark in DIC images) is not confined to the apical cytosol but mostly situated throughout the cluster. n > 4 biological replicates. (C) Quantification of retinal coverage after transplantation of RPE cells from experiments A and B (exp A and exp B, representing the experiments depicted in figures 5I and 5J, respectively) shows a similar coverage in all fractions, irrespective of CD54^+^/PSA-NCAM^-^ cell purity (exp A: n=5, 7, and 6 biological replicates in groups UNS, ENR and RED, respectively. exp B: n=7, 4, and 9 biological replicates in groups UNS, ENR and RED, respectively). (D) Reanalysis of the unsorted (UNS), CD54^+^/PSA-NCAM^-^-enriched (ENR) and -reduced (RED) fractions of RPE sorted with an Aria III flow cytometer. Orange boxes show the target cell sorting gate and orange numbers show purity of the target cell population per fraction. Scale bars: 20 µm (A, B). Statistics: p < 0.05, **p < 0.01, ***p<0.001 by Kruskal-Wallis test (exp. A) and one-way ANOVA (exp. B) (C).

