## Supplementary material for "Extensive monolayer formation after transplantation depends on a RPE subpopulation derived from human iPSCs": Document S3

Date:2024-06-14

Operator: Sebastian Knoebel

Version:2.13.0+54.g95abf0f4

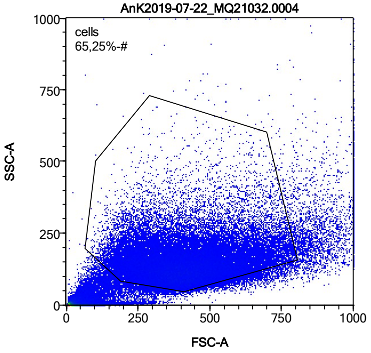


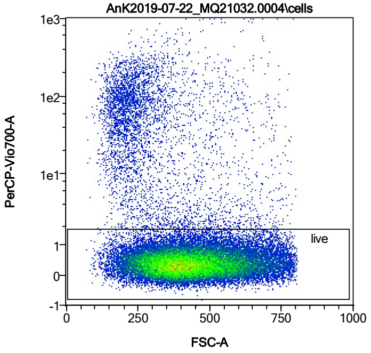


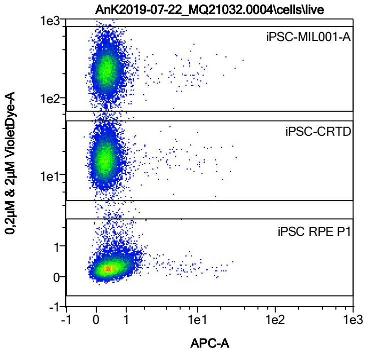


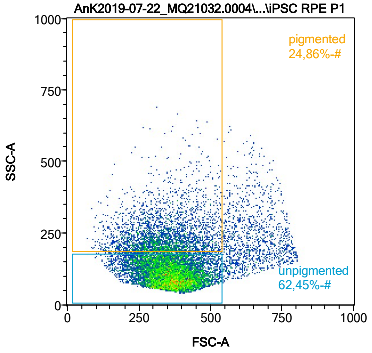


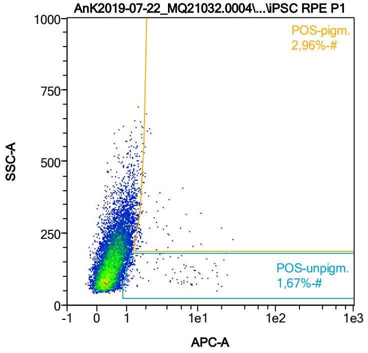


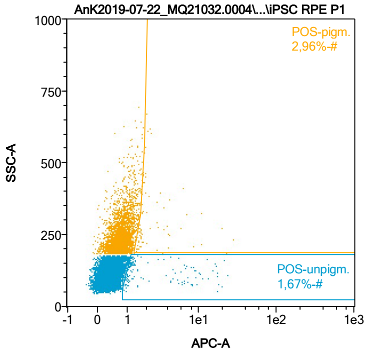


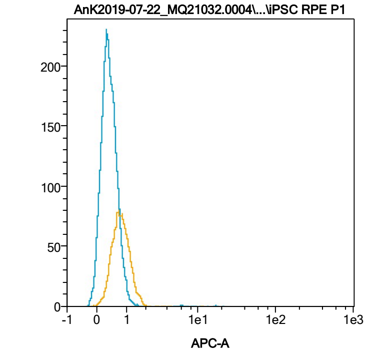


| Name | %-# | APC-A Mean | APC-A Median |
|---|---|---|---|
| POS-unpigm. | 1,67 | 4,39 | 1,34 |
| POS-pigm. | 2,96 | 3,20 | 1,78 |

Date:2024-06-14

Operator: Sebastian Knoebel

Version:2.13.0+54.g95abf0f4

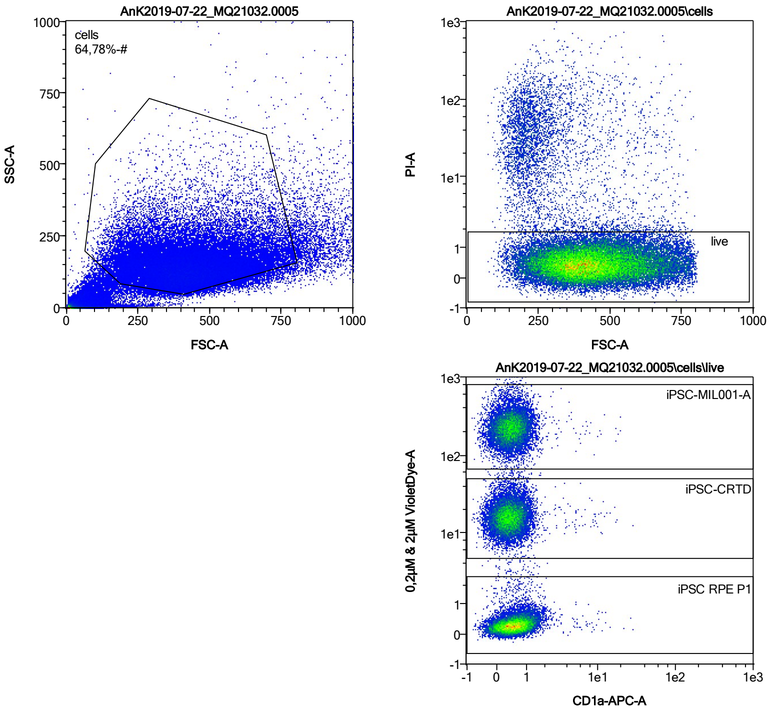


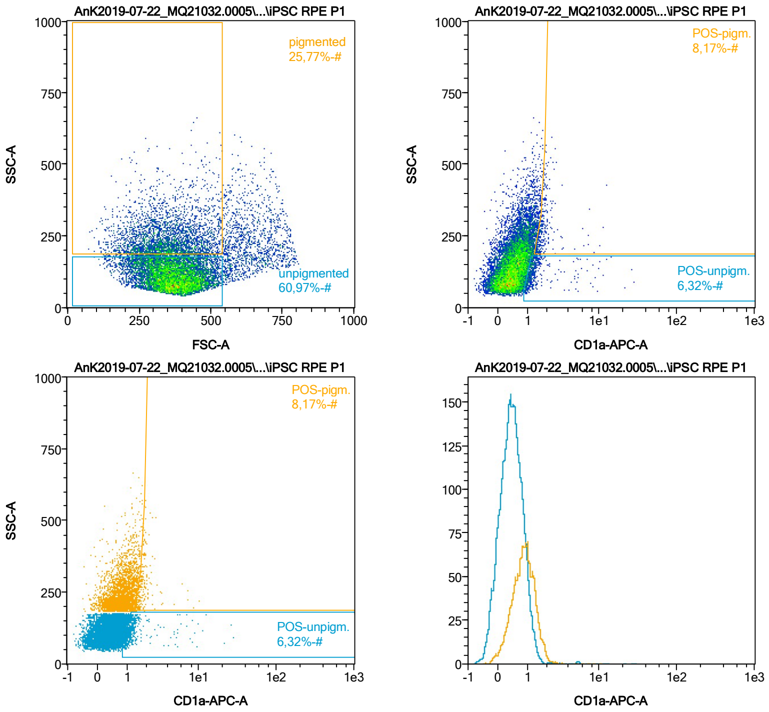


| Name | %-# | CD1a-APC-A Mean | CD1a-APC-A Median |
|---|---|---|---|
| POS-unpigm. | 6,32 | 1,64 | 1,17 |
| POS-pigm. | 8,17 | 1,87 | 1,68 |

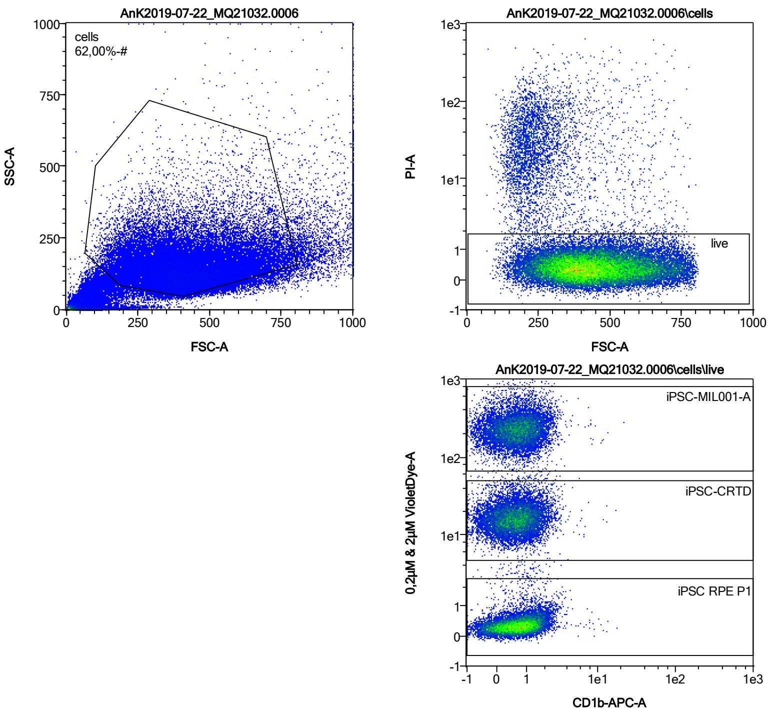


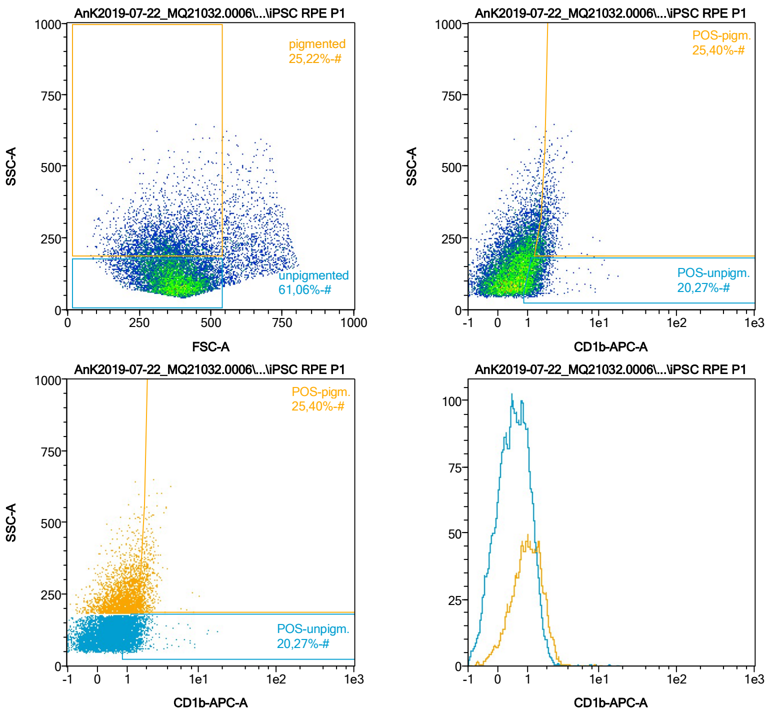


| Name | %-# | CD1b-APC-A Mean | CD1b-APC-A Median |
|---|---|---|---|
| POS-unpigm. | 20,27 | 1,40 | 1,25 |
| POS-pigm. | 25,40 | 1,92 | 1,78 |

Date:2024-06-14

Operator: Sebastian Knoebel

Version:2.13.0+54.g95abf0f4

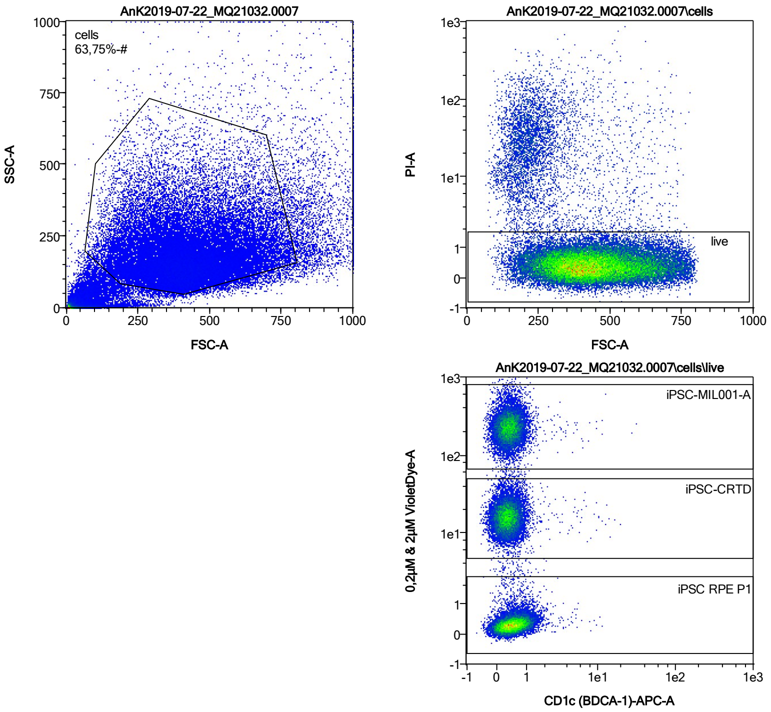


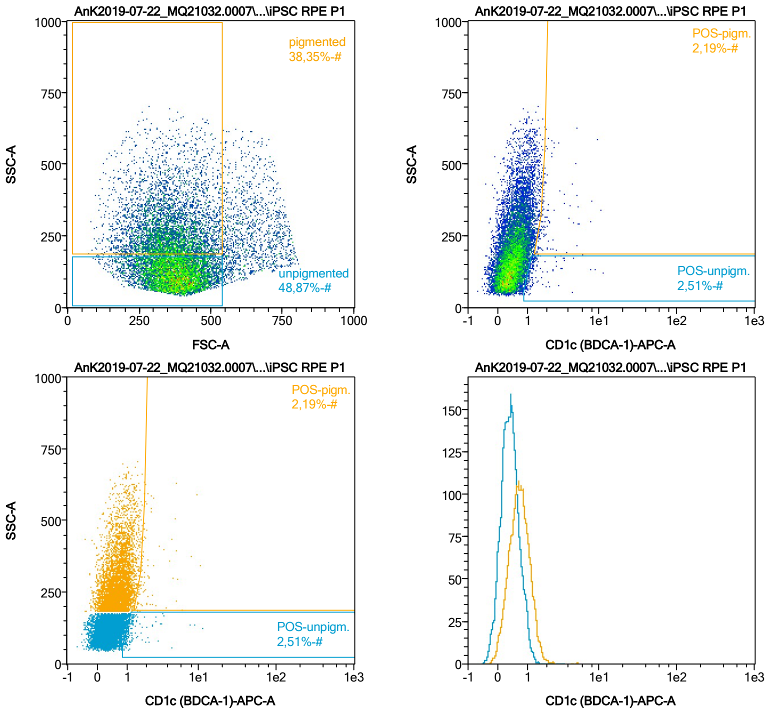


| Name | %-# | CD1c (BDCA-1)-APC-A Mean | CD1c (BDCA-1)-APC-A Median |
|---|---|---|---|
| POS-unpigm. | 2,51 | 1,46 | 1,12 |
| POS-pigm. | 2,19 | 2,37 | 1,83 |

Date:2024-06-14

Operator: Sebastian Knoebel

Version:2.13.0+54.g95abf0f4

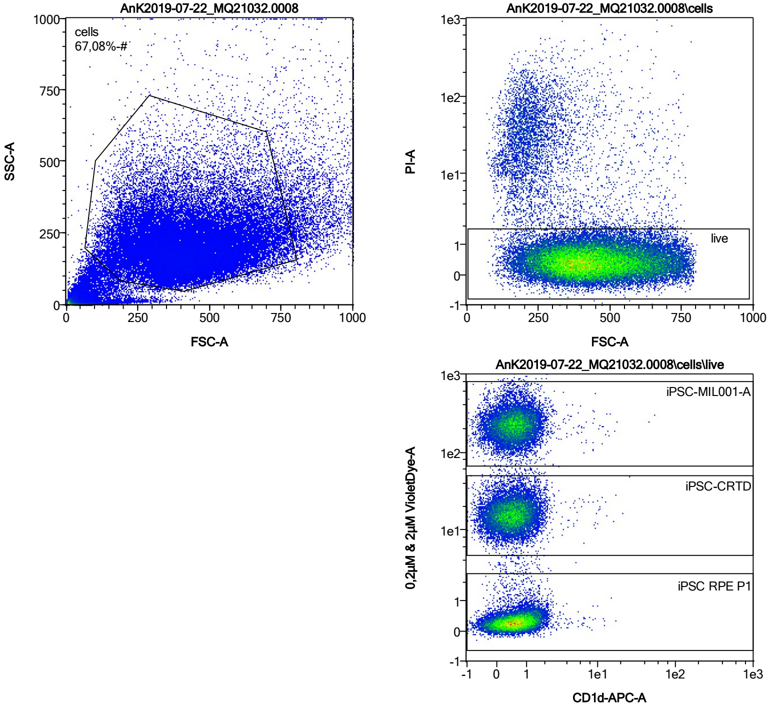


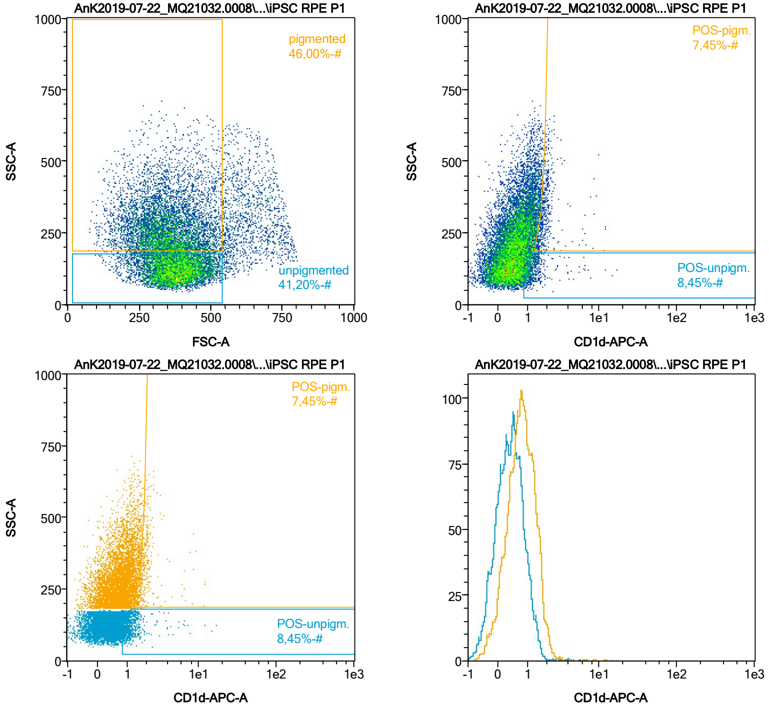


| Name | %-# | CD1d-APC-A Mean | CD1d-APC-A Median |
|---|---|---|---|
| POS-unpigm. | 8,45 | 1,52 | 1,21 |
| POS-pigm. | 7,45 | 2,06 | 1,79 |

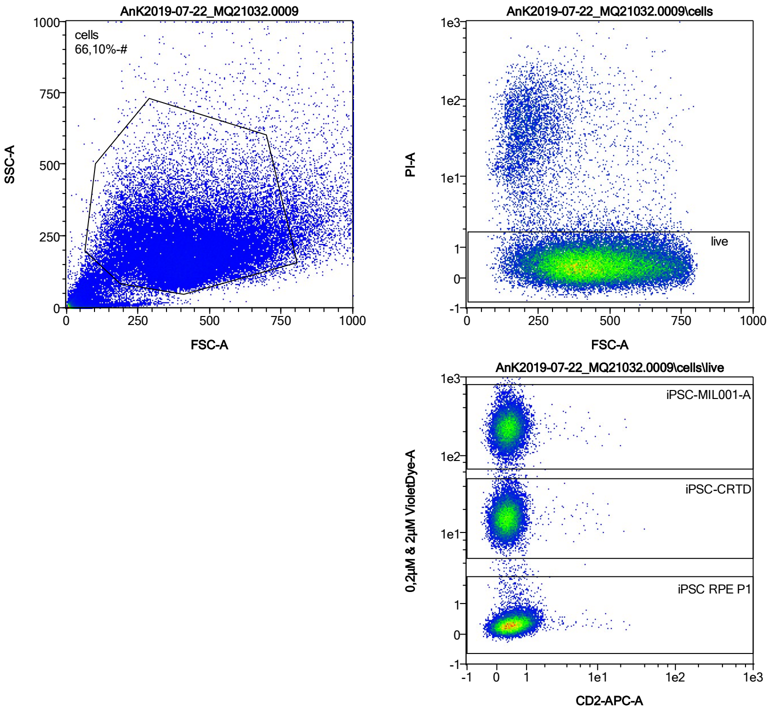


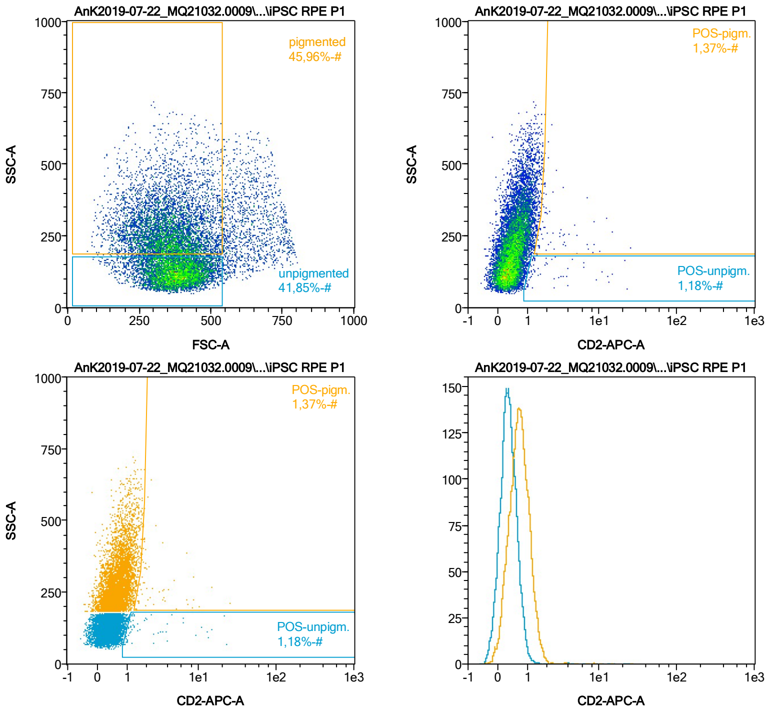


| Name | %-# | CD2-APC-A Mean | CD2-APC-A Median |
|---|---|---|---|
| POS-unpigm. | 1,18 | 3,33 | 1,28 |
| POS-pigm. | 1,37 | 3,10 | 1,91 |

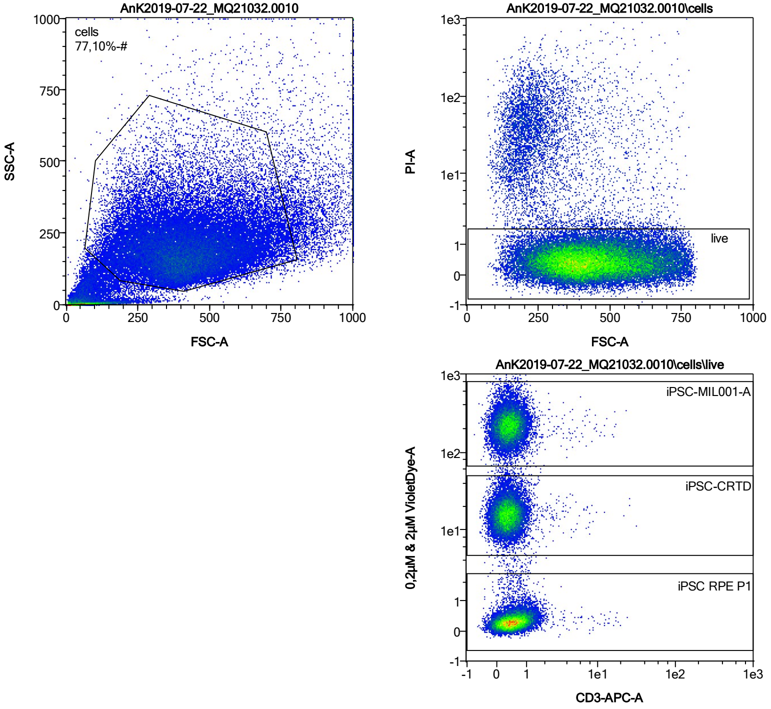


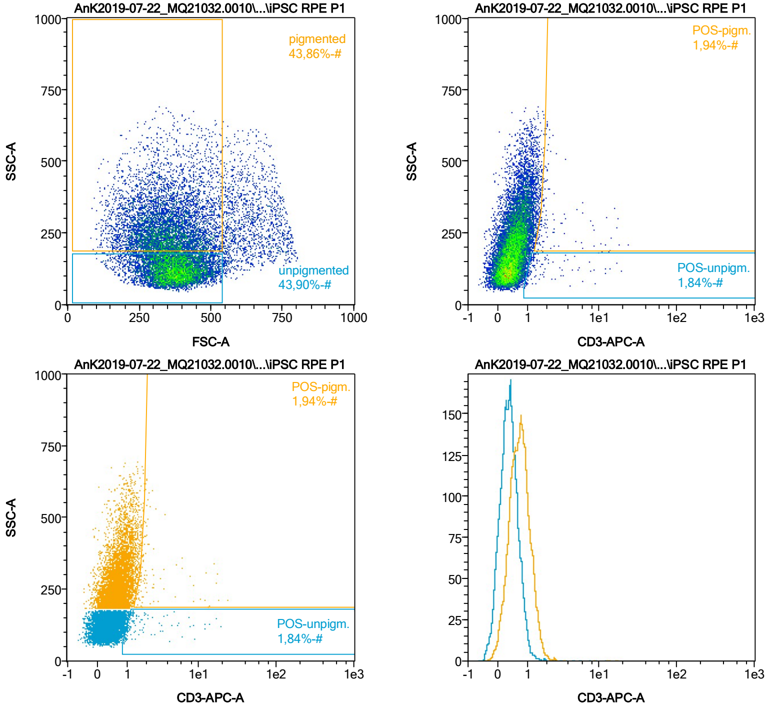


| Name | %-# | CD3-APC-A Mean | CD3-APC-A Median |
|---|---|---|---|
| POS-unpigm. | 1,84 | 2,44 | 1,27 |
| POS-pigm. | 1,94 | 3,33 | 1,88 |

Date:2024-06-14

Operator: Sebastian Knoebel

Version:2.13.0+54.g95abf0f4

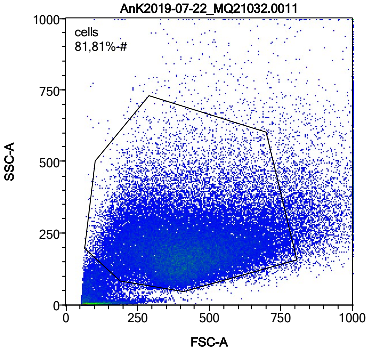


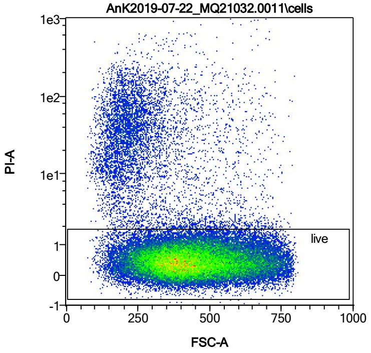


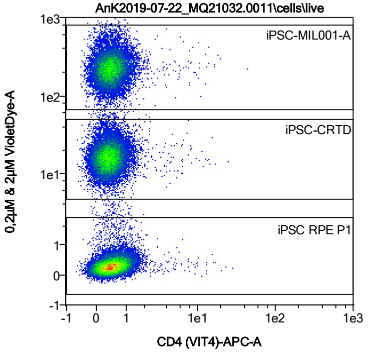


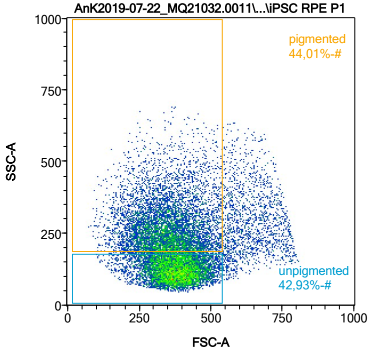


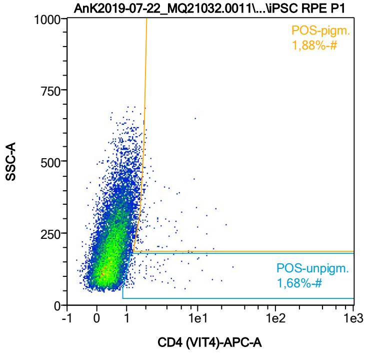


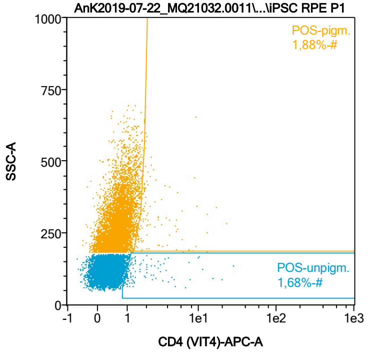


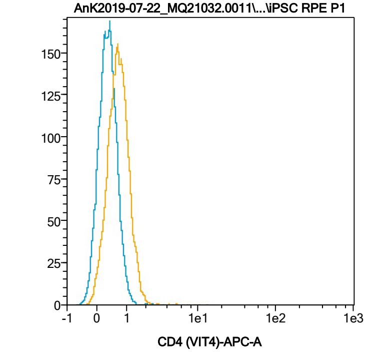


| Name | %-# | CD4 (VIT4)-APC-A Mean | CD4 (VIT4)-APC-A Median |
|---|---|---|---|
| POS-unpigm. | 1,68 | 2,59 | 1,25 |
| POS-pigm. | 1,88 | 3,25 | 1,97 |

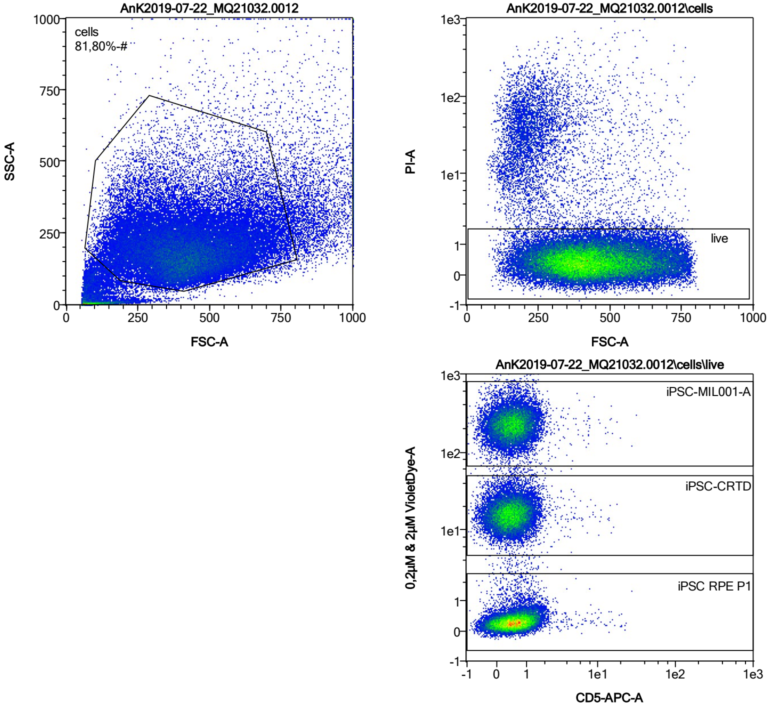


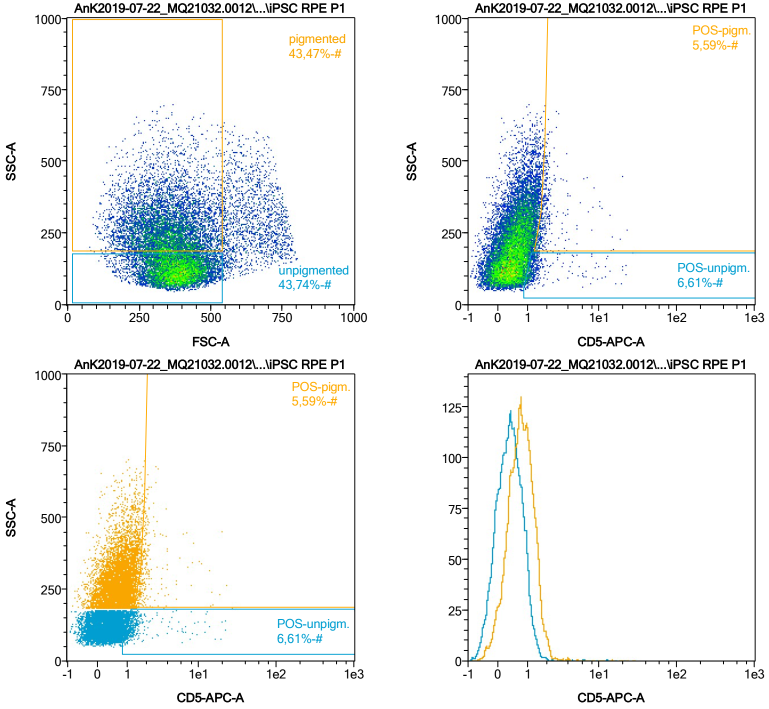


| Name | %-# | CD5-APC-A Mean | CD5-APC-A Median |
|---|---|---|---|
| POS-unpigm. | 6,61 | 1,84 | 1,19 |
| POS-pigm. | 5,59 | 2,42 | 1,78 |

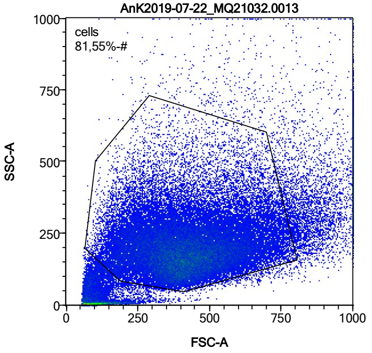


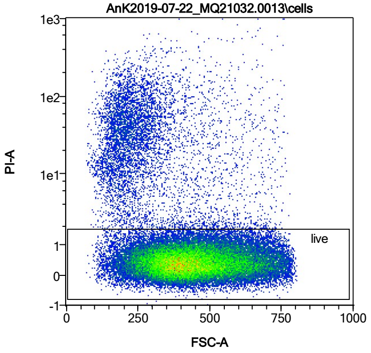


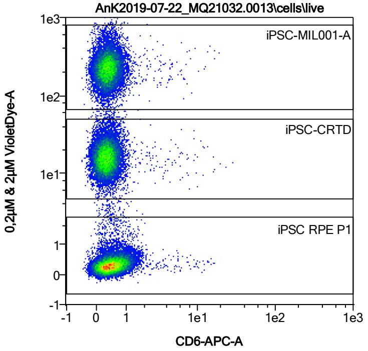


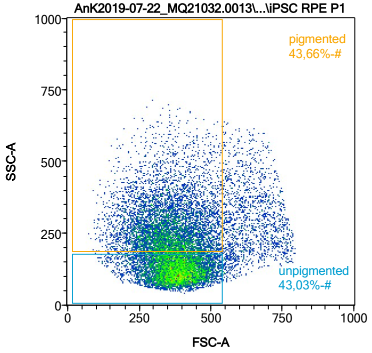


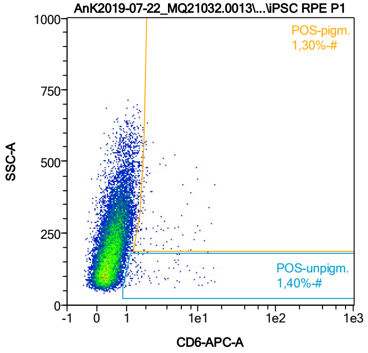


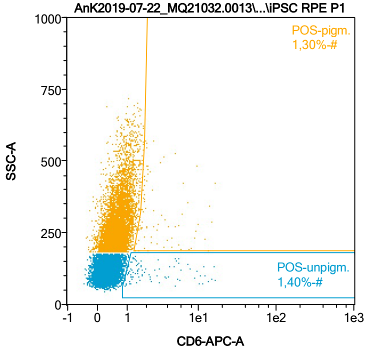


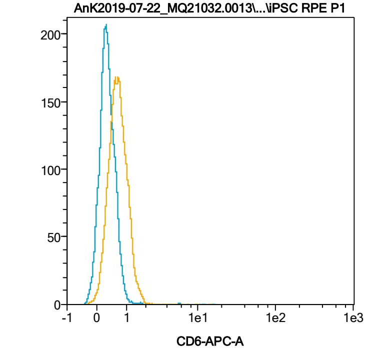


| Name | %-# | CD6-APC-A Mean | CD6-APC-A Median |
|---|---|---|---|
| POS-unpigm. | 1,40 | 3,46 | 1,59 |
| POS-pigm. | 1,30 | 3,70 | 2,08 |

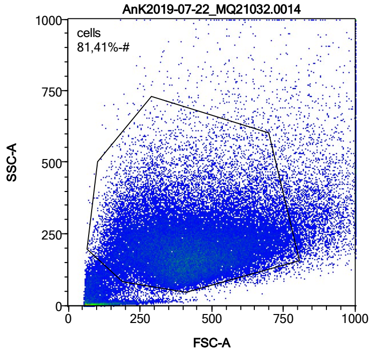


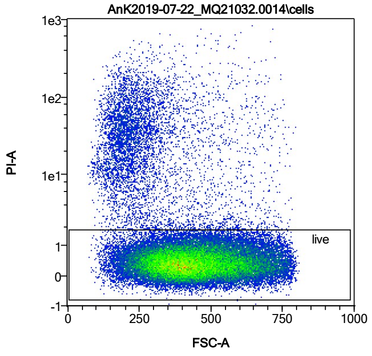


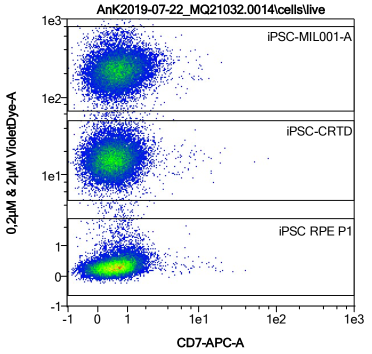


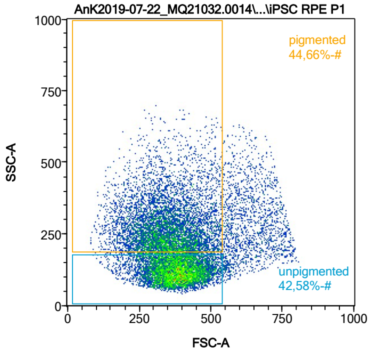


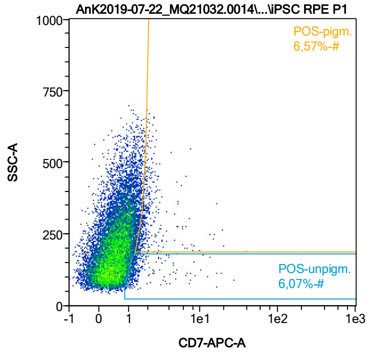


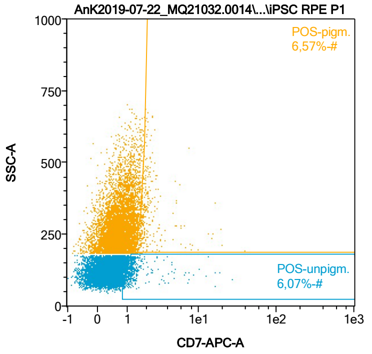


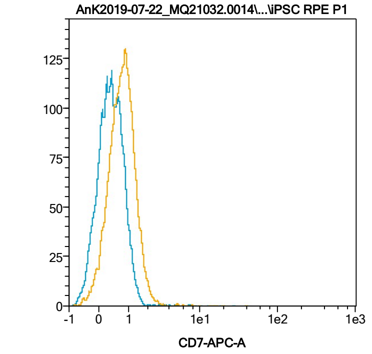


| Name | %-# | CD7-APC-A Mean | CD7-APC-A Median |
|---|---|---|---|
| POS-unpigm. | 6,07 | 1,74 | 1,20 |
| POS-pigm. | 6,57 | 2,34 | 1,79 |

Date:2024-06-14

Operator: Sebastian Knoebel

Version:2.13.0+54.g95abf0f4

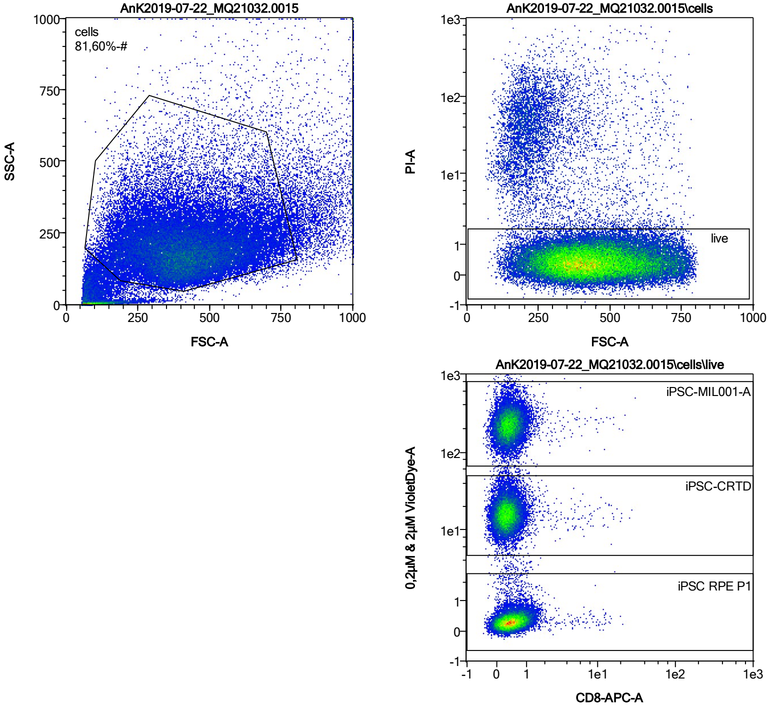


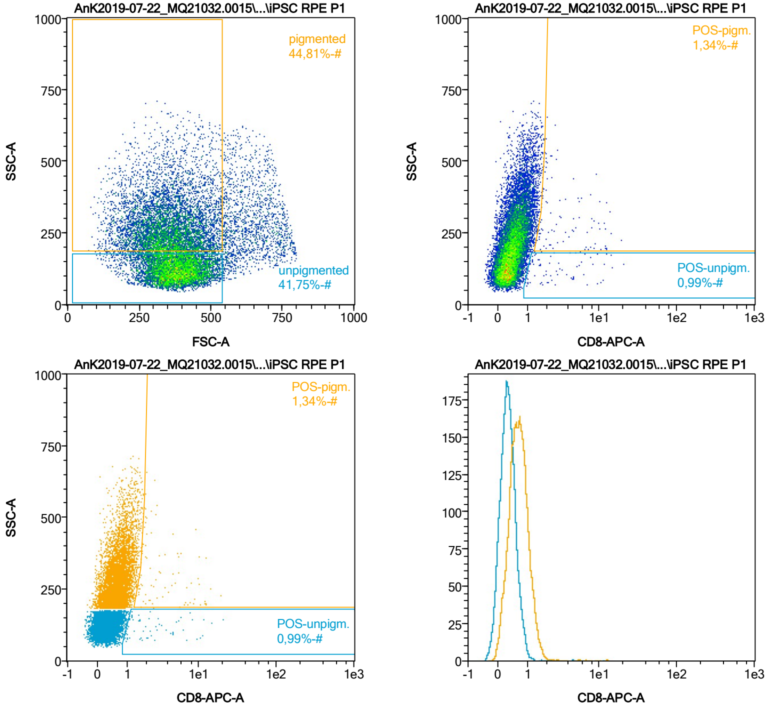


| Name | %-# | CD8-APC-A Mean | CD8-APC-A Median |
|---|---|---|---|
| POS-unpigm. | 0,99 | 3,52 | 1,96 |
| POS-pigm. | 1,34 | 4,32 | 2,43 |

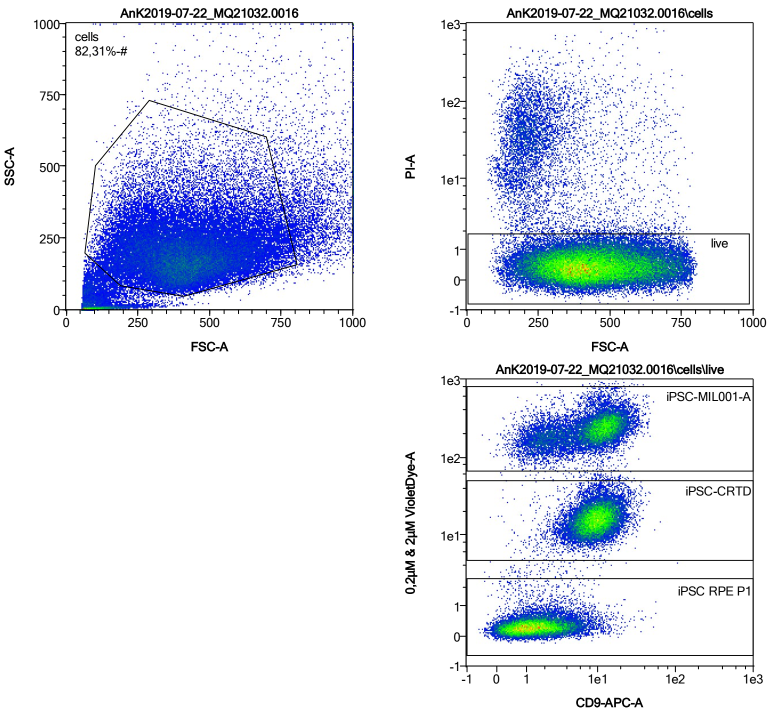


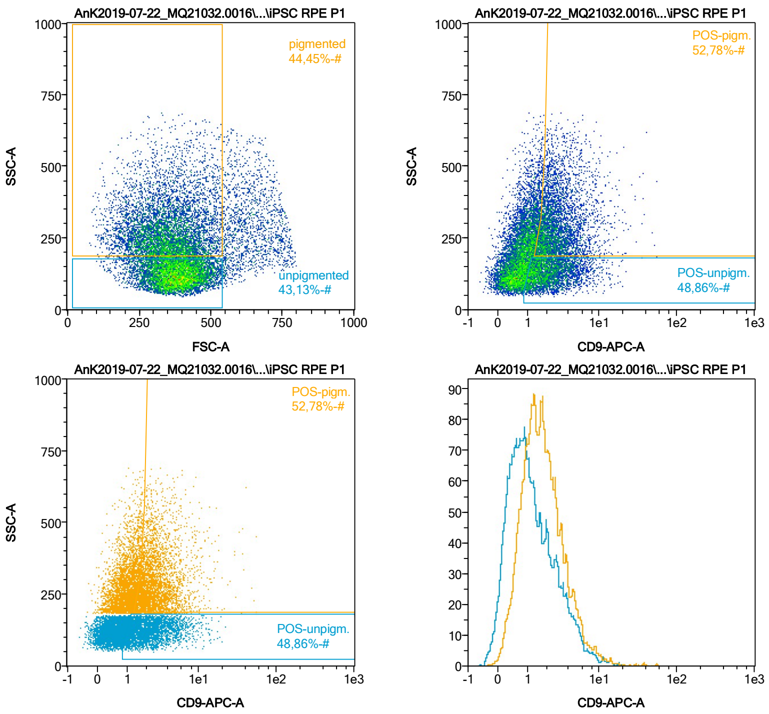


| Name | %-# | CD9-APC-A Mean | CD9-APC-A Median |
|---|---|---|---|
| POS-unpigm. | 48,86 | 2,60 | 1,98 |
| POS-pigm. | 52,78 | 3,28 | 2,50 |

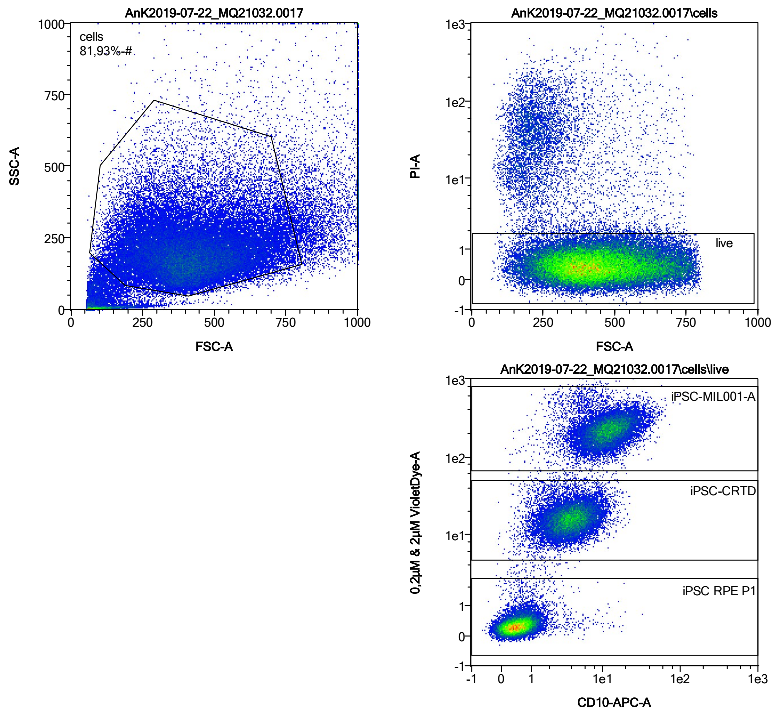


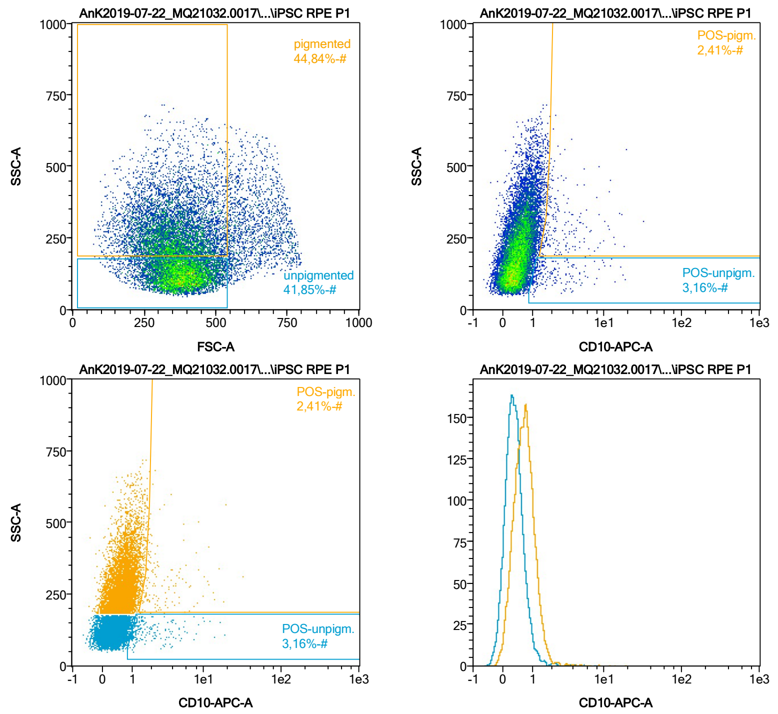


| Name | %-# | CD10-APC-A Mean | CD10-APC-A Median |
|---|---|---|---|
| POS-unpigm. | 3,16 | 2,40 | 1,41 |
| POS-pigm. | 2,41 | 3,58 | 2,14 |

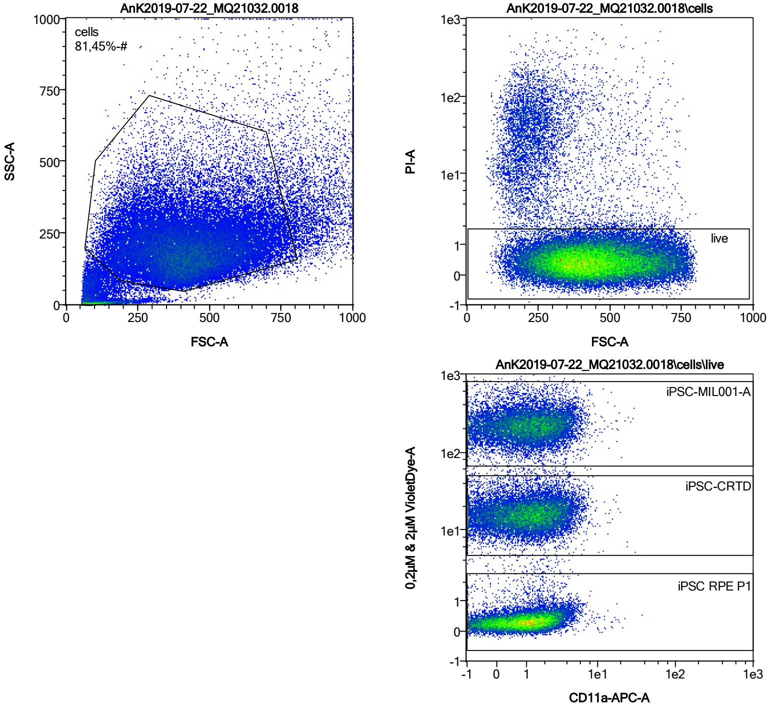


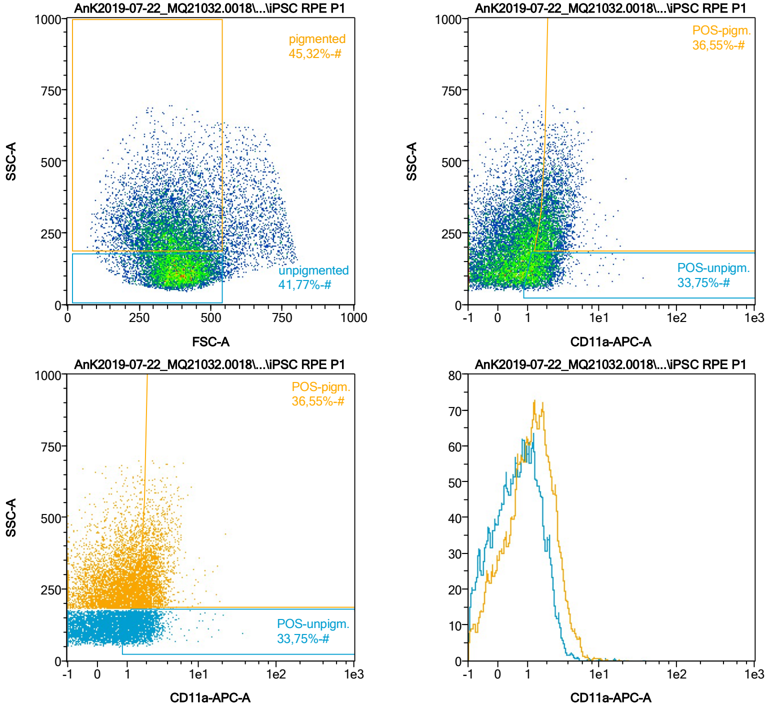


| Name | %-# | CD11a-APC-A Mean | CD11a-APC-A Median |
|---|---|---|---|
| POS-unpigm. | 33,75 | 1,81 | 1,55 |
| POS-pigm. | 36,55 | 2,53 | 2,23 |

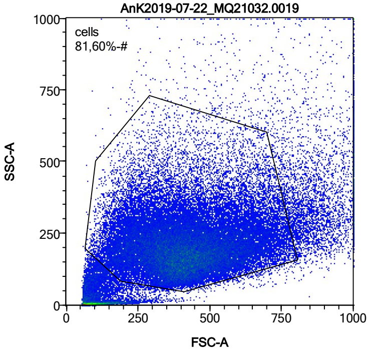


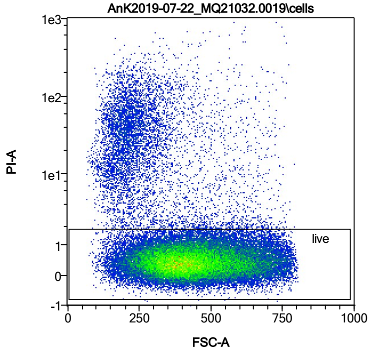


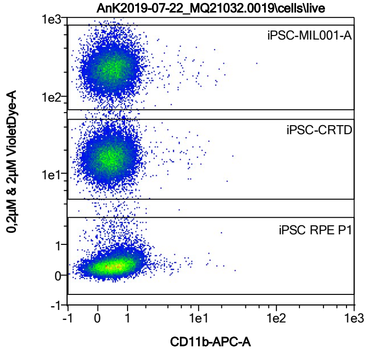


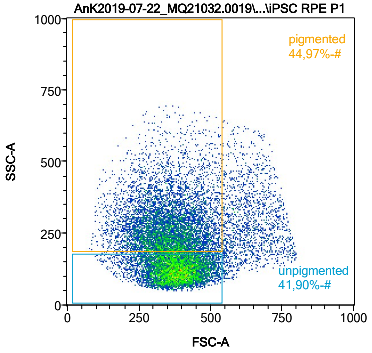


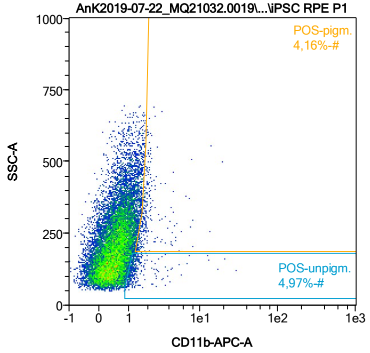


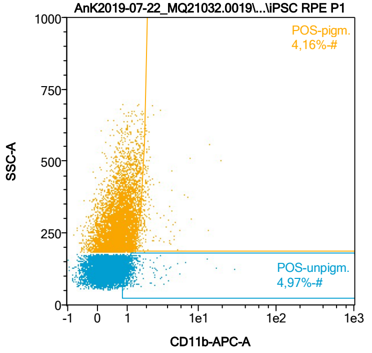


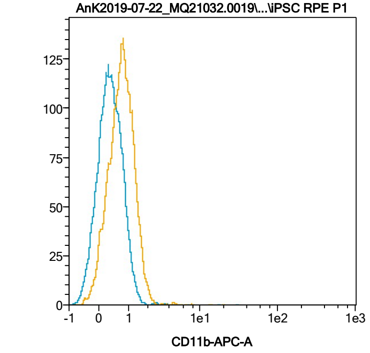


| Name | %-# | CD11b-APC-A Mean | CD11b-APC-A Median |
|---|---|---|---|
| POS-unpigm. | 4,97 | 1,58 | 1,20 |
| POS-pigm. | 4,16 | 2,23 | 1,82 |

Date:2024-06-14

Operator: Sebastian Knoebel

Version:2.13.0+54.g95abf0f4

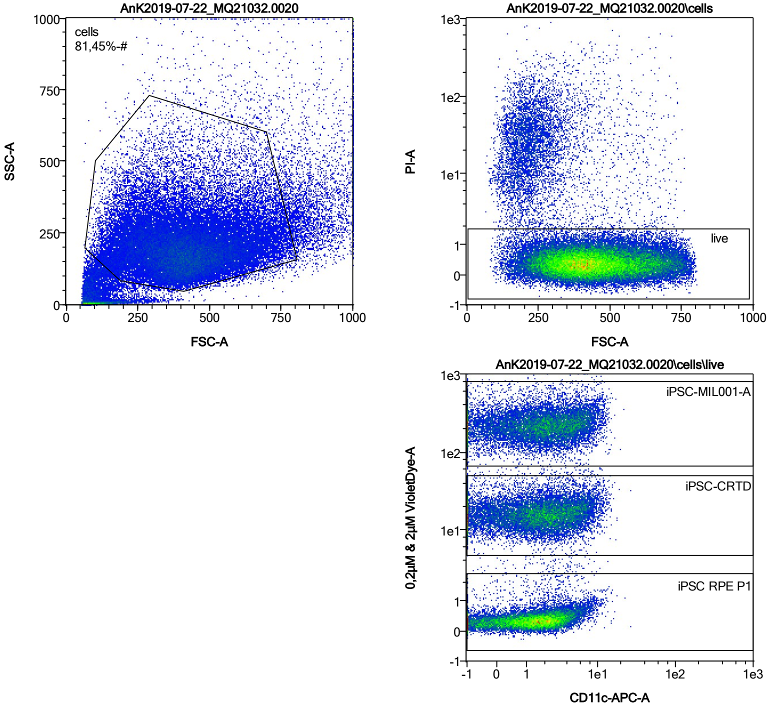


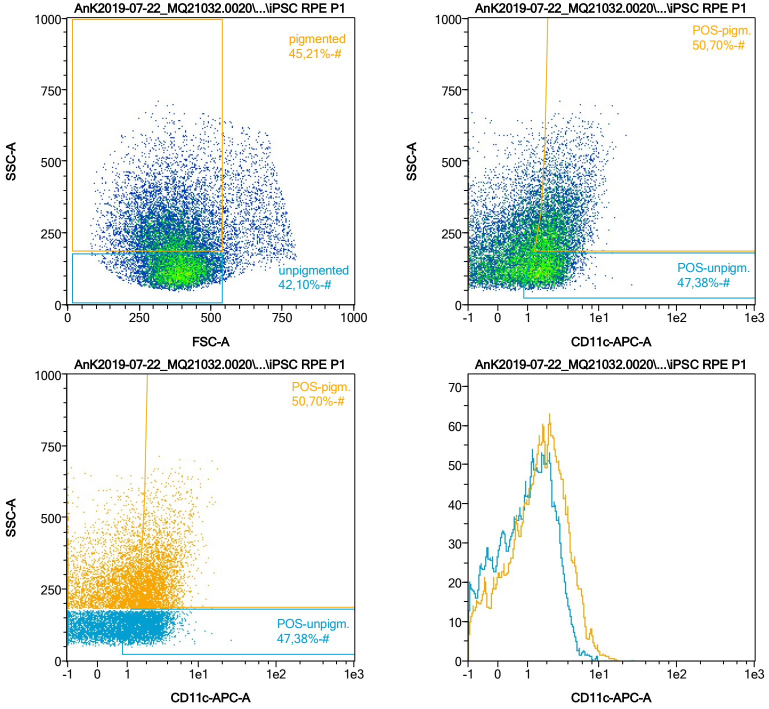


| Name | %-# | CD11c-APC-A Mean | CD11c-APC-A Median |
|---|---|---|---|
| POS-unpigm. | 47,38 | 2,20 | 1,94 |
| POS-pigm. | 50,70 | 3,21 | 2,70 |

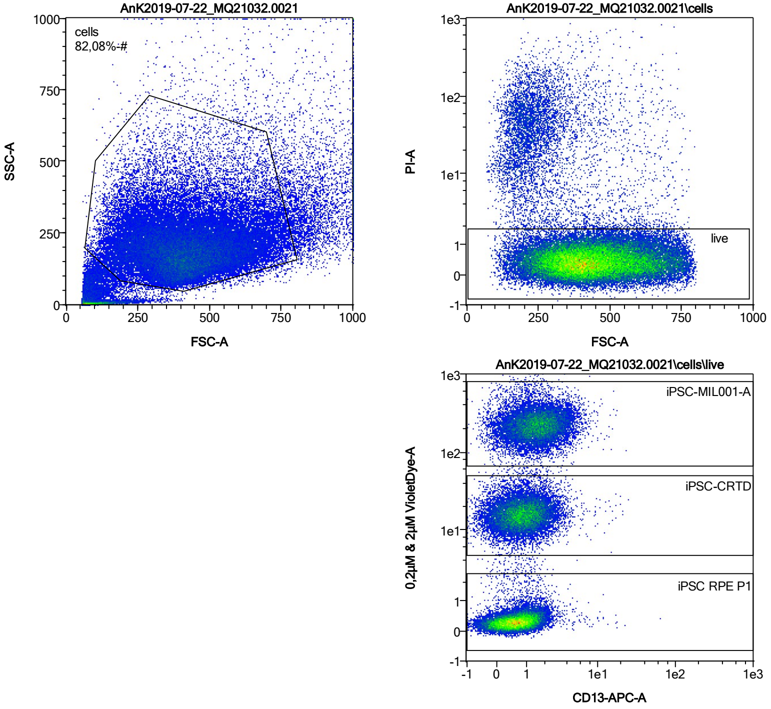


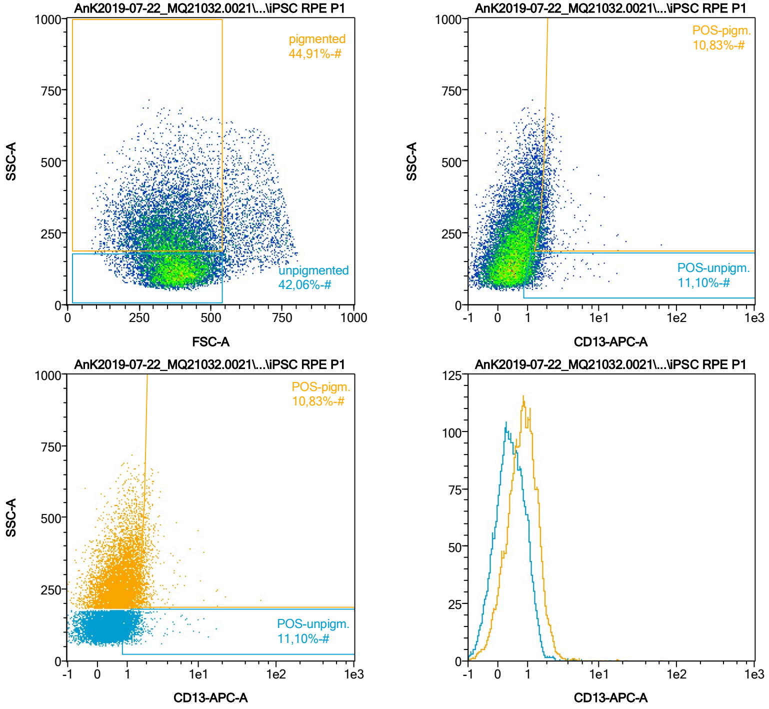


| Name | %-# | CD13-APC-A Mean | CD13-APC-A Median |
|---|---|---|---|
| POS-unpigm. | 11,10 | 1,47 | 1,23 |
| POS-pigm. | 10,83 | 2,20 | 1,83 |

Date:2024-06-14

Operator: Sebastian Knoebel

Version:2.13.0+54.g95abf0f4

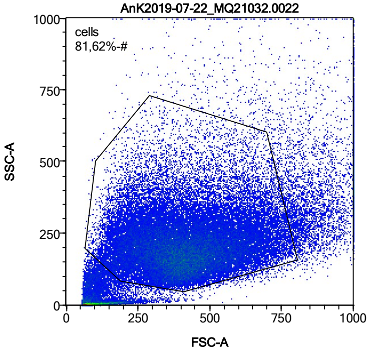


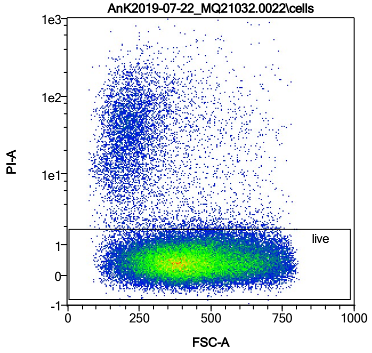


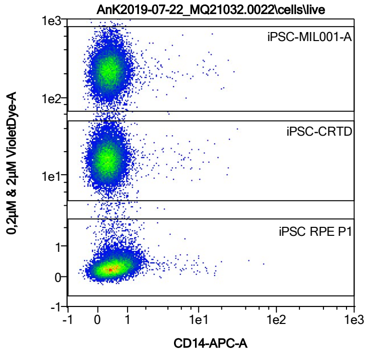


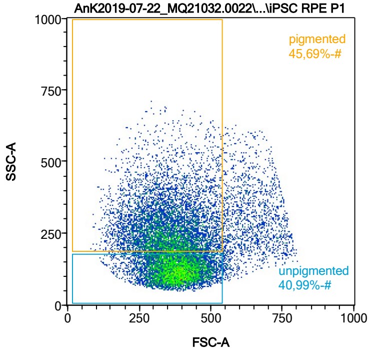


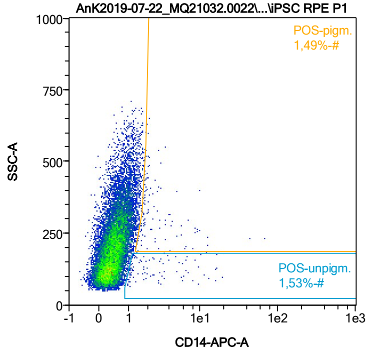


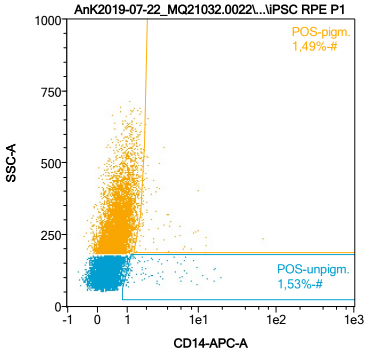


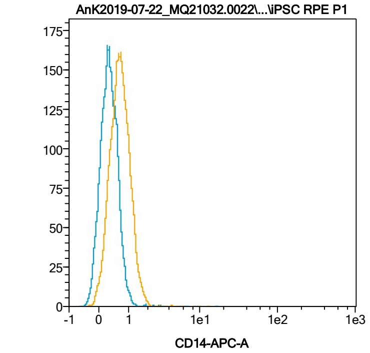


| Name | %-# | CD14-APC-A Mean | CD14-APC-A Median |
|---|---|---|---|
| POS-unpigm. | 1,53 | 3,41 | 1,79 |
| POS-pigm. | 1,49 | 3,64 | 1,93 |

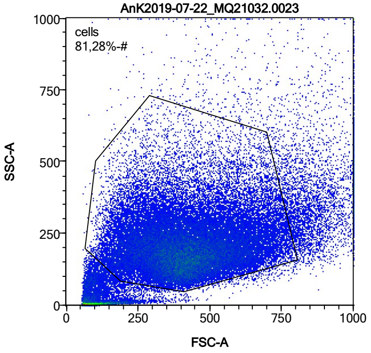


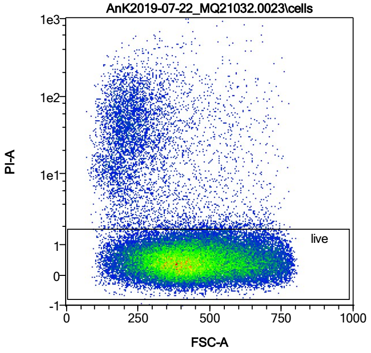


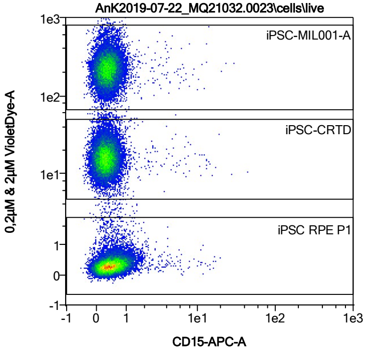


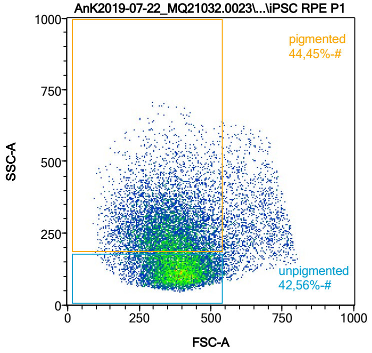


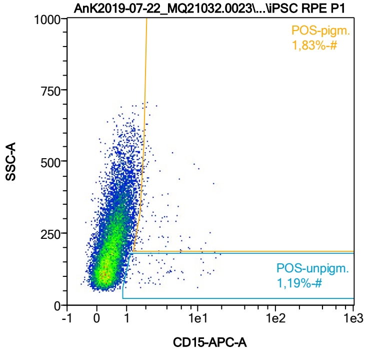


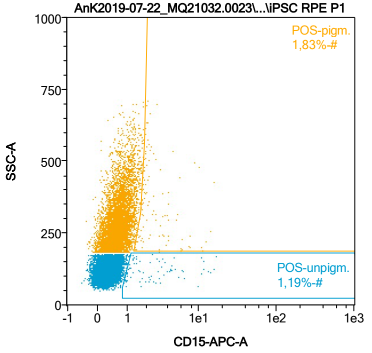


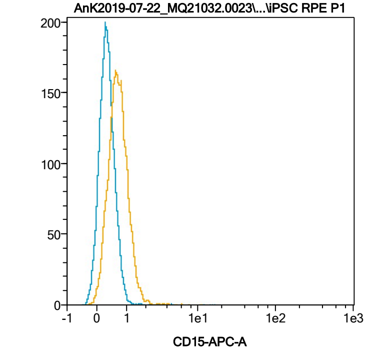


| Name | %-# | CD15-APC-A Mean | CD15-APC-A Median |
|---|---|---|---|
| POS-unpigm. | 1,19 | 3,62 | 2,18 |
| POS-pigm. | 1,83 | 3,15 | 2,08 |

Date:2024-06-14
Operator: Sebastian Knoebel
Version:2.13.0+54.g95abf0f4

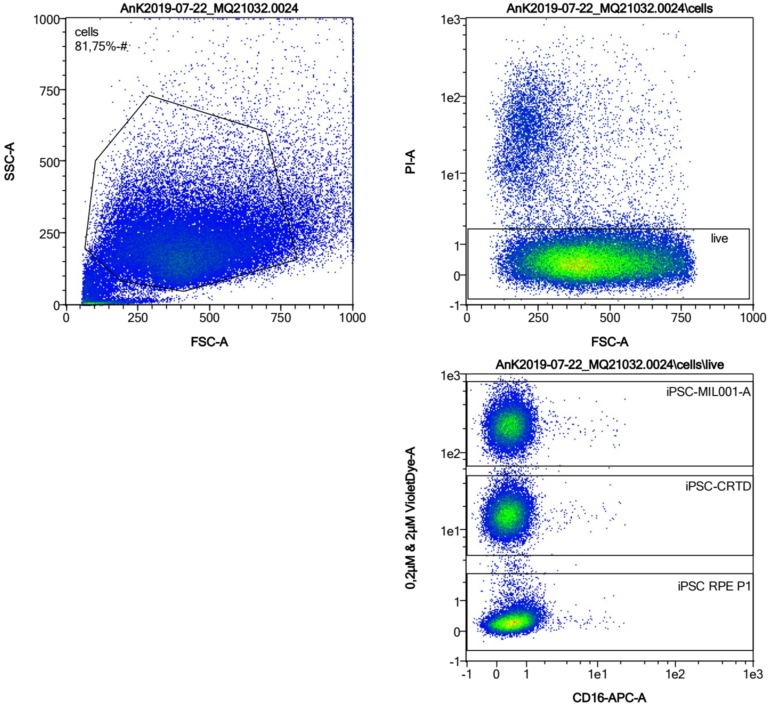


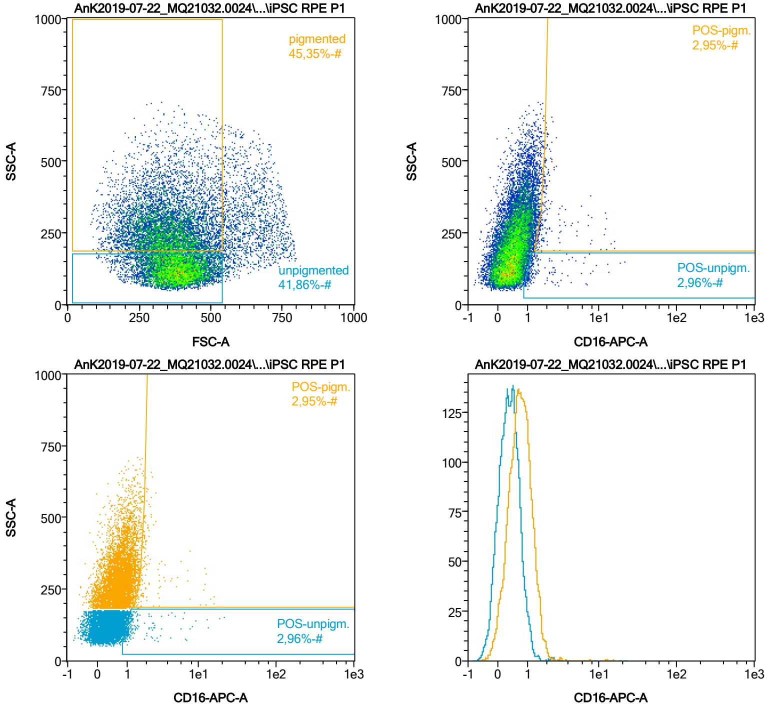


| Name | %-# | CD16-APC-A Mean | CD16-APC-A Median |
|---|---|---|---|
| POS-unpigm. | 2,96 | 2,00 | 1,17 |
| POS-pigm. | 2,95 | 2,59 | 1,82 |

Date:2024-06-14
Operator: Sebastian Knoebel
Version:2.13.0+54.g95abf0f4

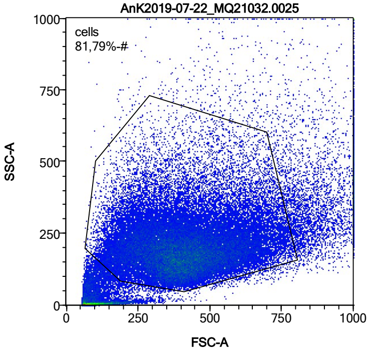


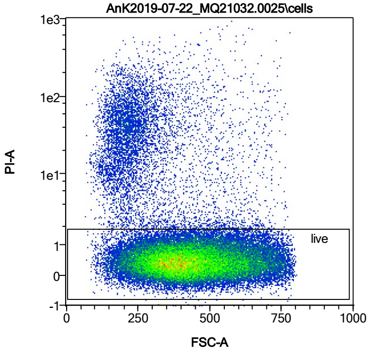


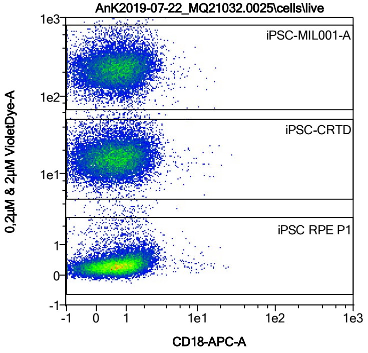


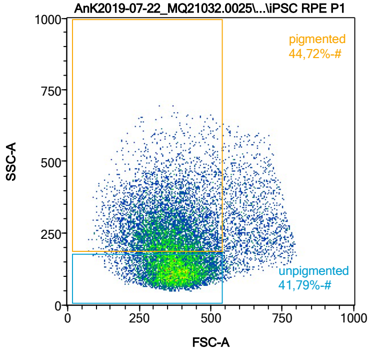


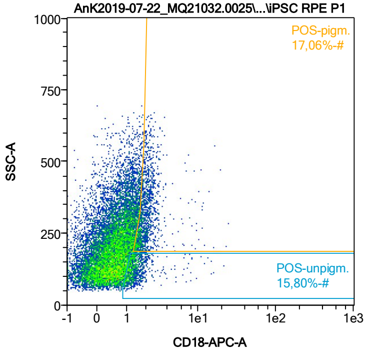


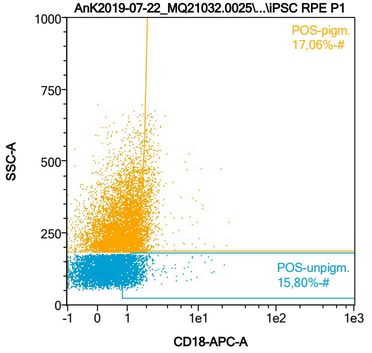


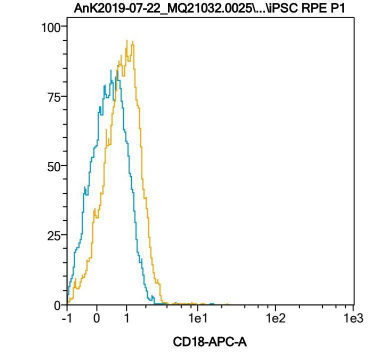


| Name | %-# | CD18-APC-A Mean | CD18-APC-A Median |
|---|---|---|---|
| POS-unpigm. | 15,80 | 1,55 | 1,28 |
| POS-pigm. | 17,06 | 2,16 | 1,89 |

Date:2024-06-14

Operator: Sebastian Knoebel

Version:2.13.0+54.g95abf0f4

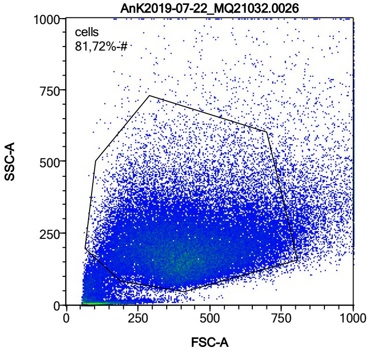


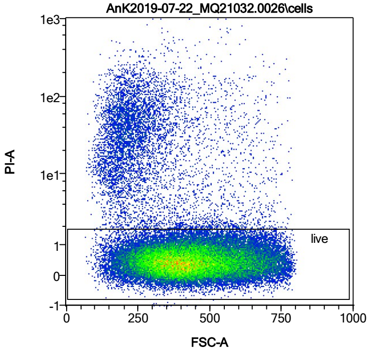


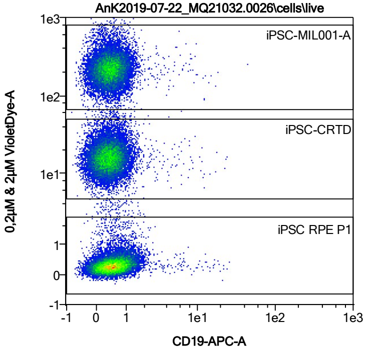


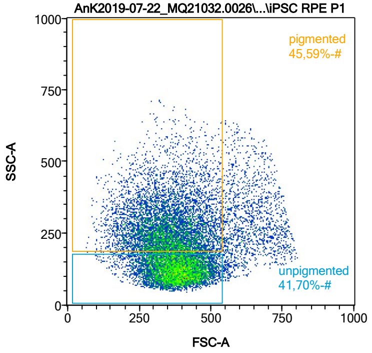


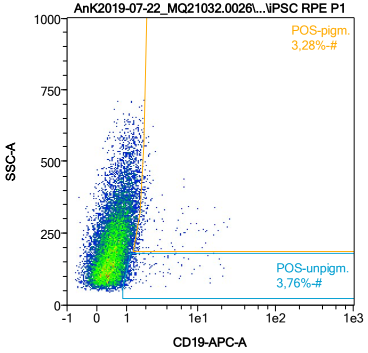


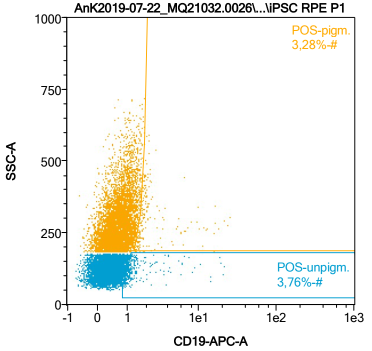


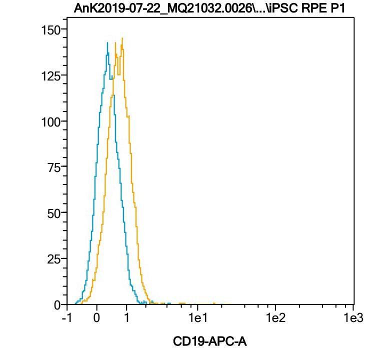


| Name | %-# | CD19-APC-A Mean | CD19-APC-A Median |
|---|---|---|---|
| POS-unpigm. | 3,76 | 2,06 | 1,20 |
| POS-pigm. | 3,28 | 3,01 | 1,81 |

Date:2024-06-14

Operator: Sebastian Knoebel

Version:2.13.0+54.g95abf0f4

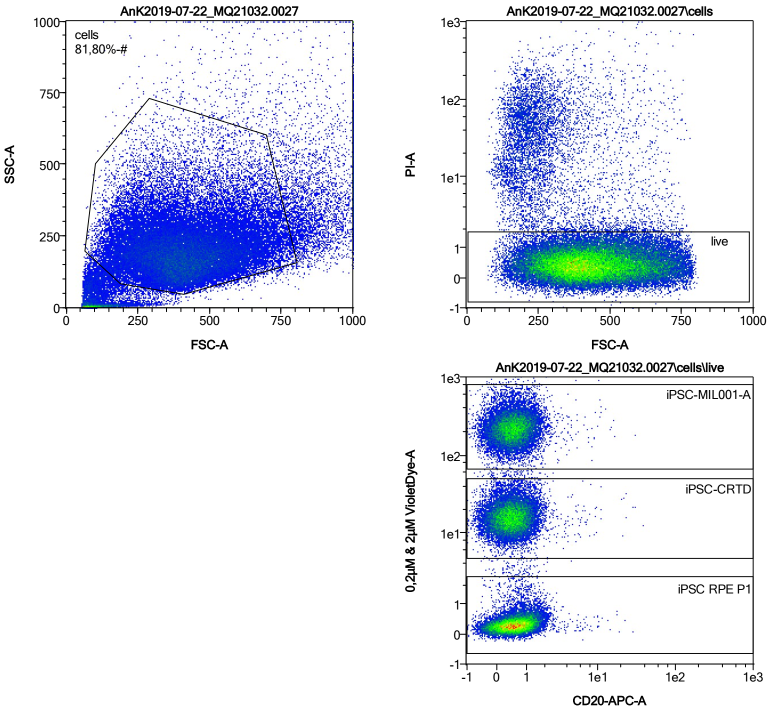


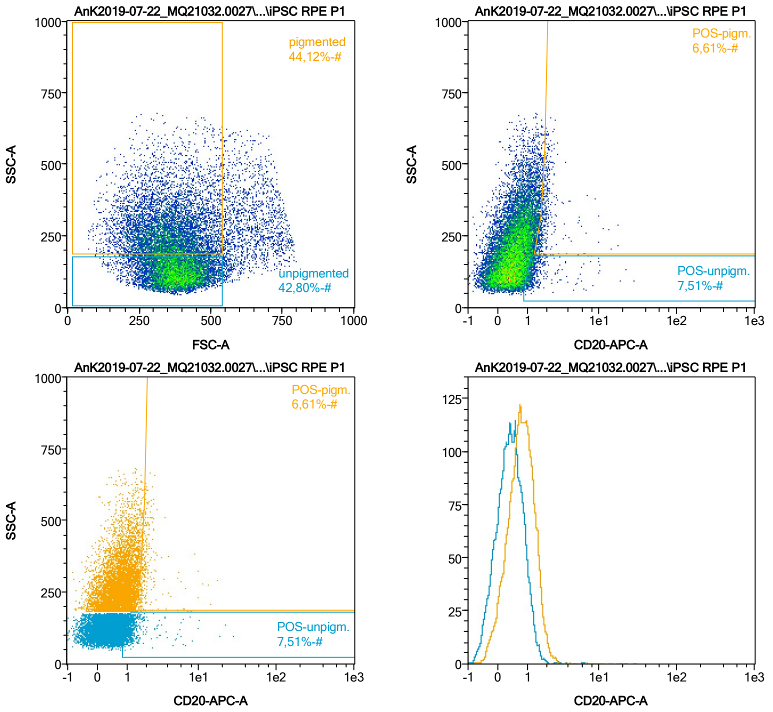


| Name | %-# | CD20-APC-A Mean | CD20-APC-A Median |
|---|---|---|---|
| POS-unpigm. | 7,51 | 1,59 | 1,20 |
| POS-pigm. | 6,61 | 2,11 | 1,78 |

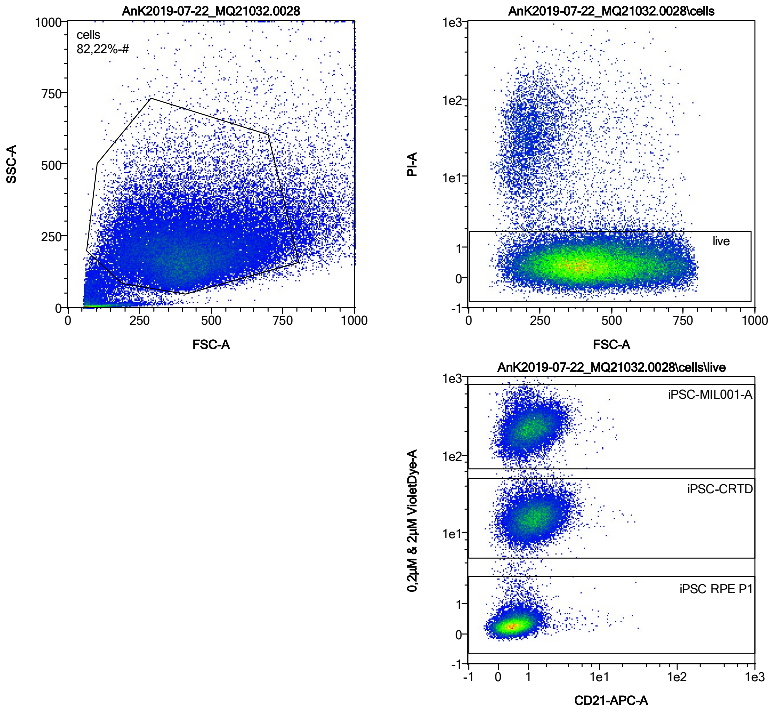


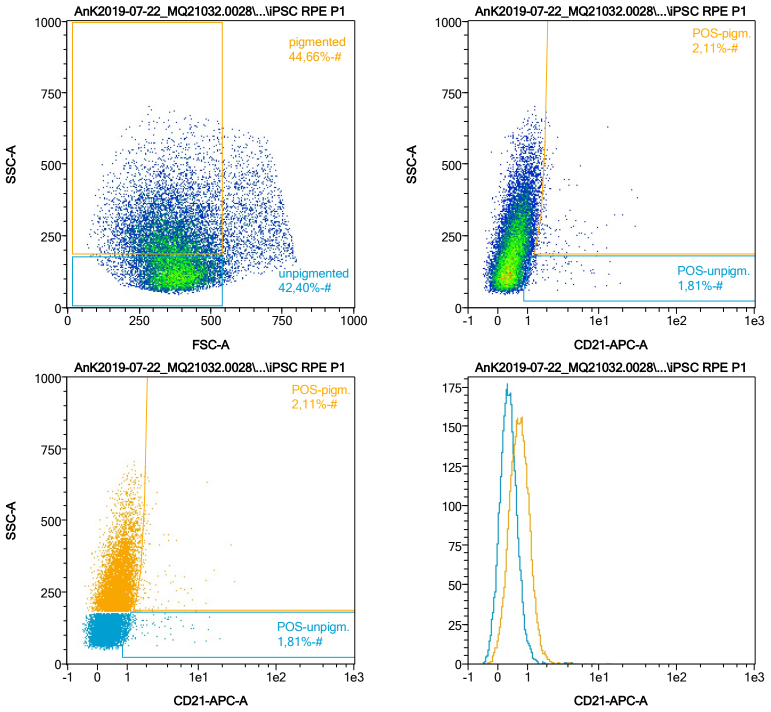


| Name | %-# | CD21-APC-A Mean | CD21-APC-A Median |
|---|---|---|---|
| POS-unpigm. | 1,81 | 2,42 | 1,25 |
| POS-pigm. | 2,11 | 3,13 | 1,94 |

Date:2024-06-14
Operator: Sebastian Knoebel
Version:2.13.0+54.g95abf0f4

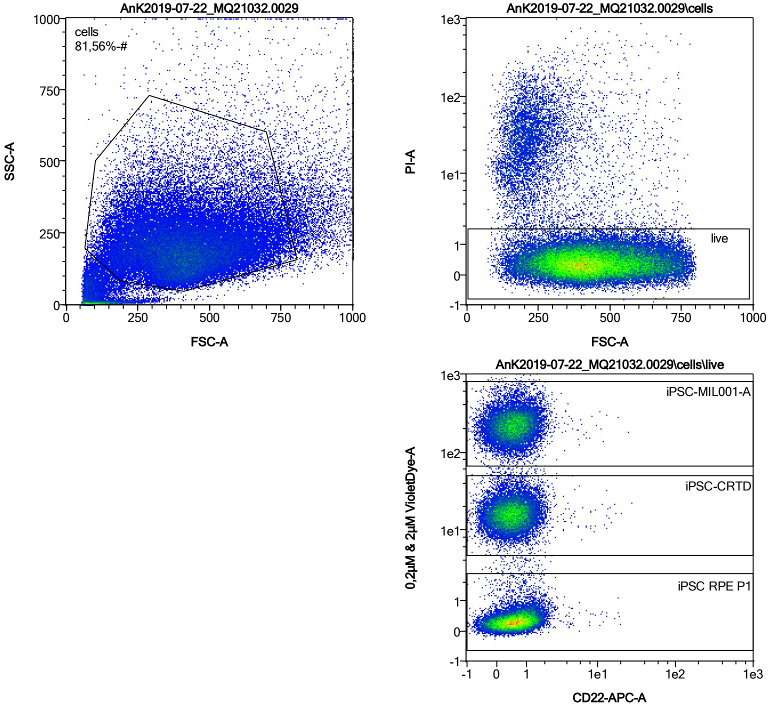


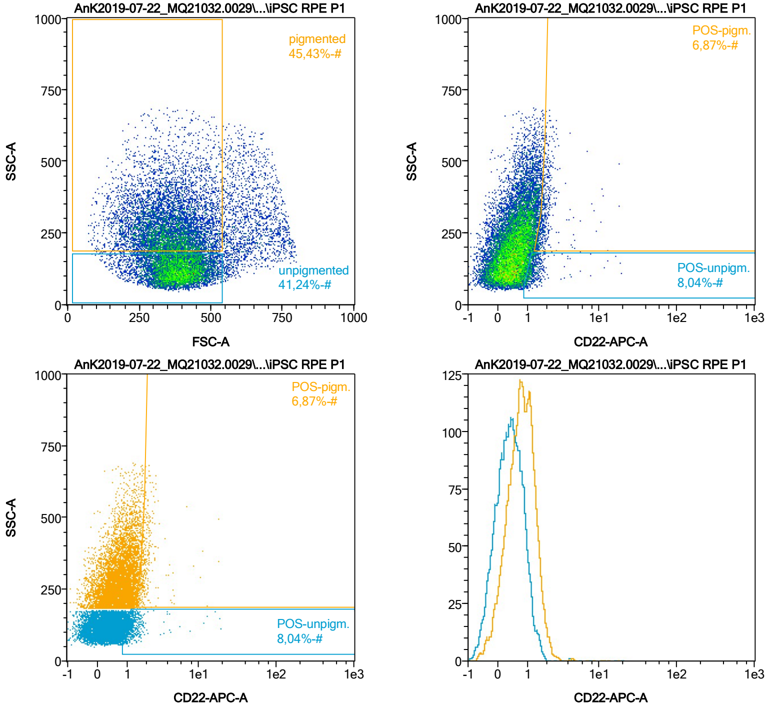


| Name | %-# | CD22-APC-A Mean | CD22-APC-A Median |
|---|---|---|---|
| POS-unpigm. | 8,04 | 1,45 | 1,18 |
| POS-pigm. | 6,87 | 2,04 | 1,80 |

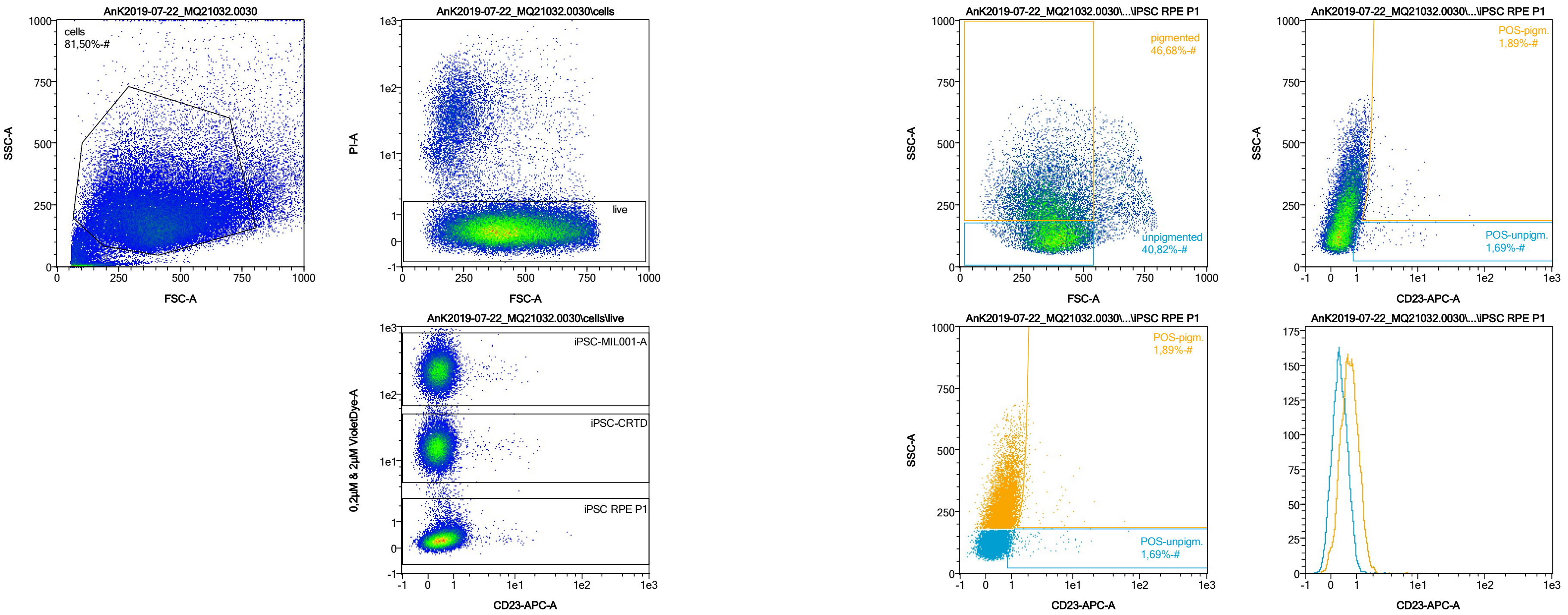


| Name | %-# | CD23-APC-A Mean | CD23-APC-A Median |
|---|---|---|---|
| POS-unpigm. | 1,69 | 3,41 | 1,34 |
| POS-pigm. | 1,89 | 3,44 | 1,92 |

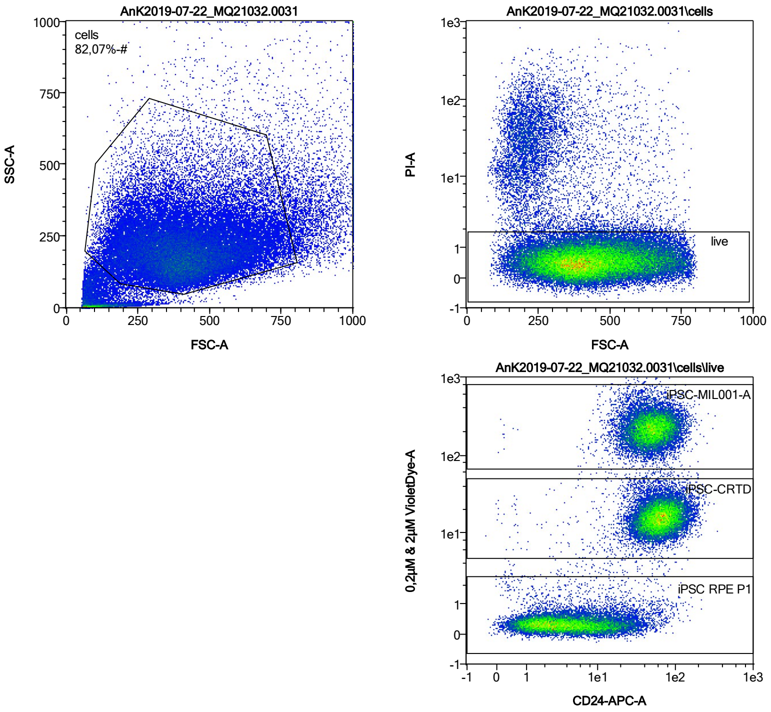


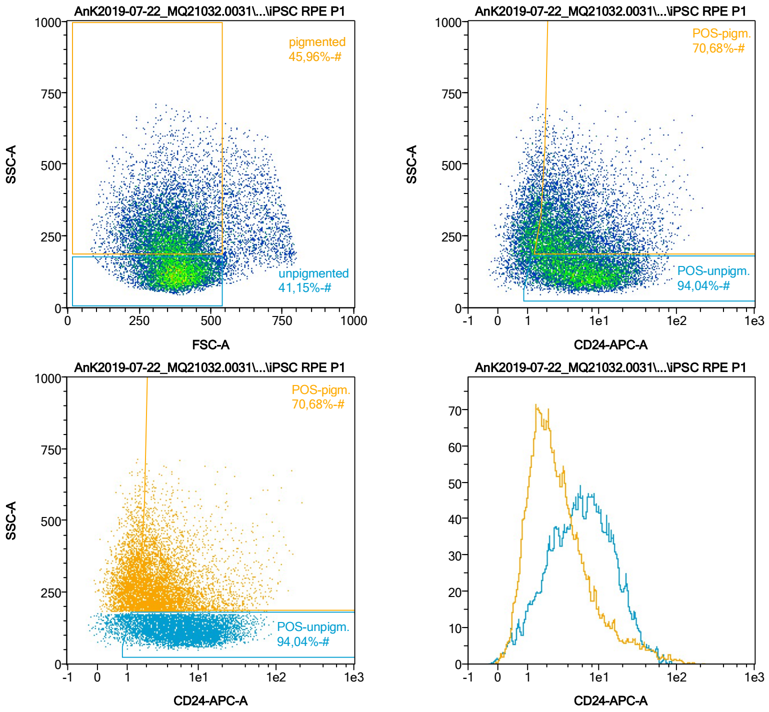


| Name | %-# | CD24-APC-A Mean | CD24-APC-A Median |
|---|---|---|---|
| POS-unpigm. | 94,04 | 9,18 | 6,36 |
| POS-pigm. | 70,68 | 7,13 | 3,40 |

Date:2024-06-14
Operator: Sebastian Knoebel
Version:2.13.0+54.g95abf0f4

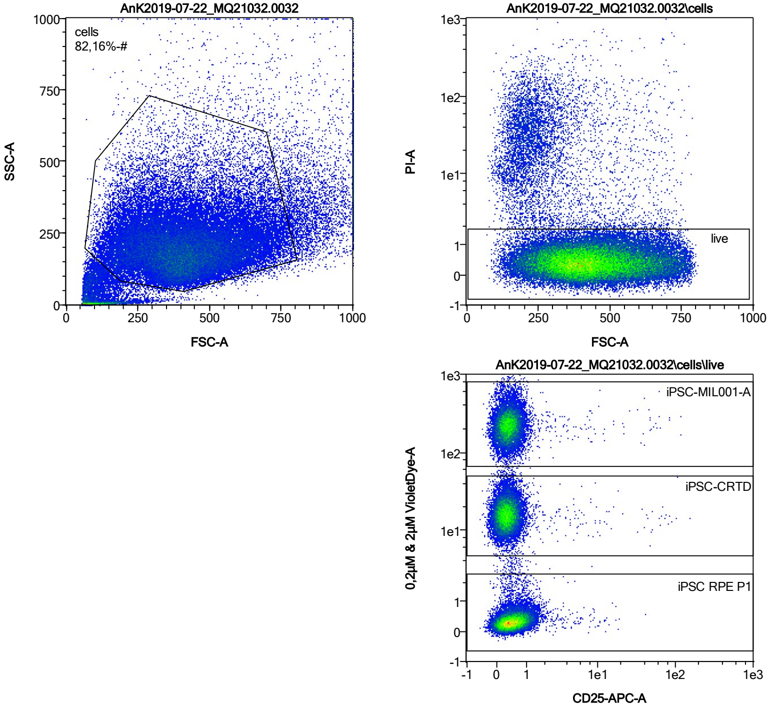


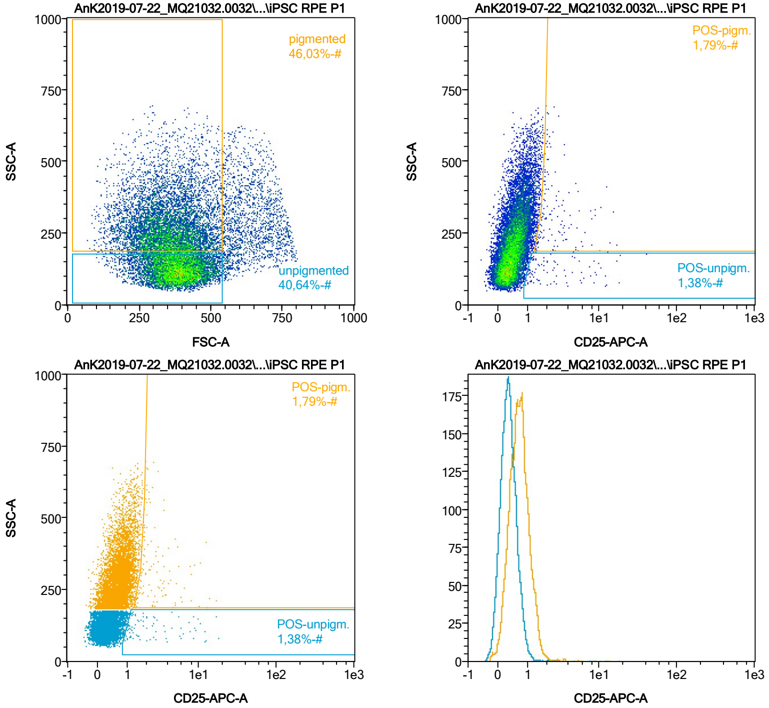


| Name | %-# | CD25-APC-A Mean | CD25-APC-A Median |
|---|---|---|---|
| POS-unpigm. | 1,38 | 3,38 | 1,66 |
| POS-pigm. | 1,79 | 3,18 | 2,17 |

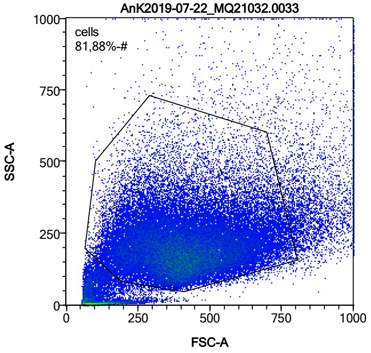


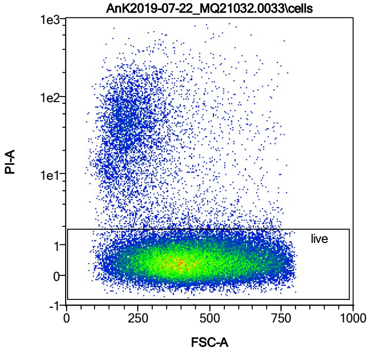


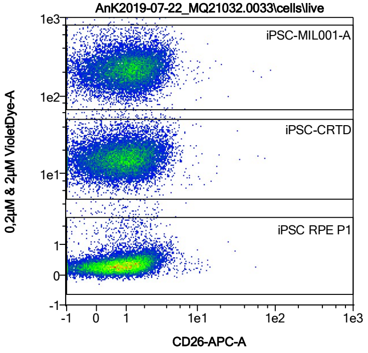


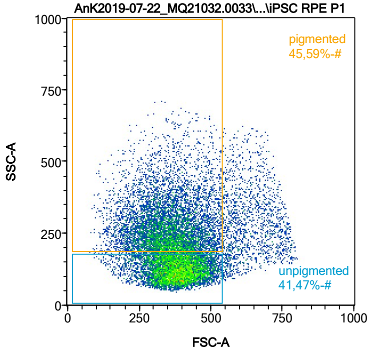


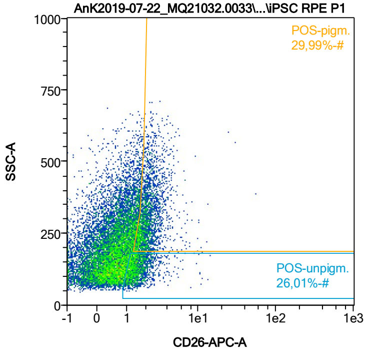


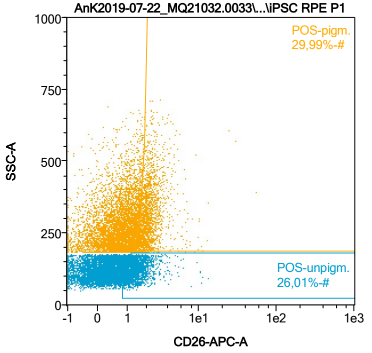


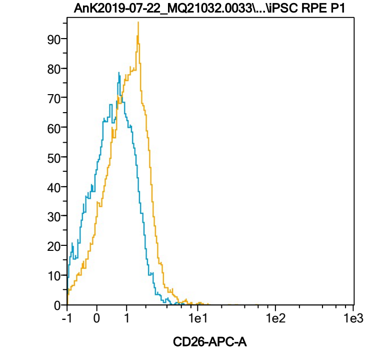


| Name | %-# | CD26-APC-A Mean | CD26-APC-A Median |
|---|---|---|---|
| POS-unpigm. | 26,01 | 1,60 | 1,41 |
| POS-pigm. | 29,99 | 2,32 | 2,03 |

Date:2024-06-14

Operator: Sebastian Knoebel

Version:2.13.0+54.g95abf0f4

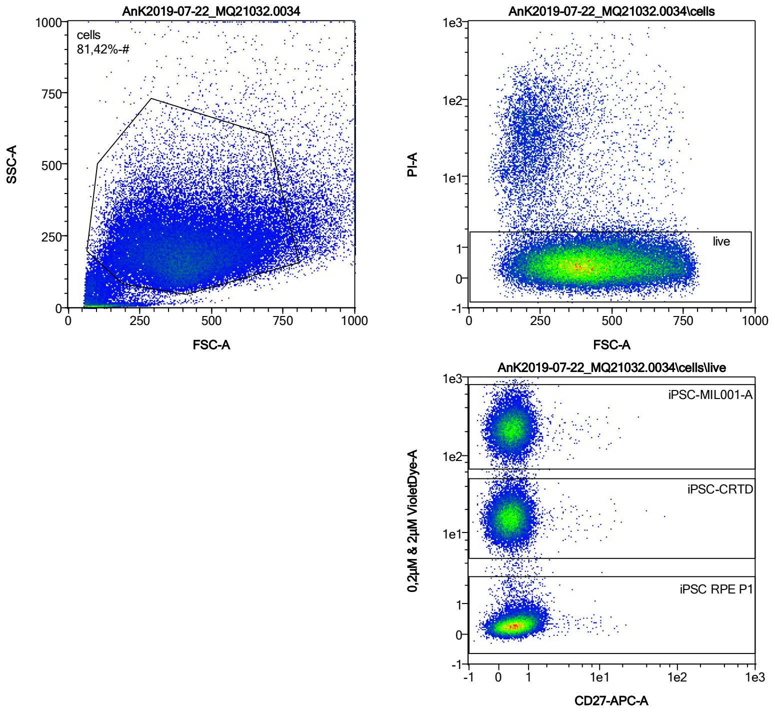


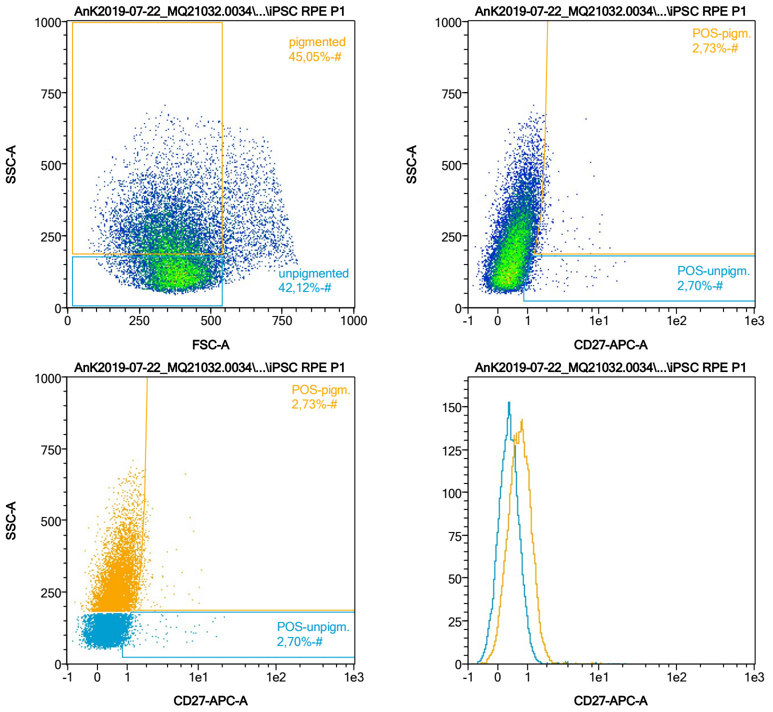


| Name | %-# | CD27-APC-A Mean | CD27-APC-A Median |
|---|---|---|---|
| POS-unpigm. | 2,70 | 2,22 | 1,18 |
| POS-pigm. | 2,73 | 2,27 | 1,85 |

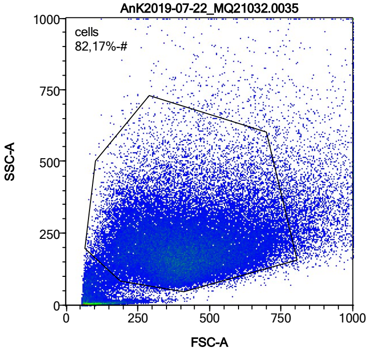


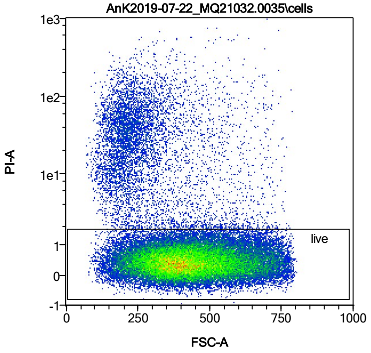


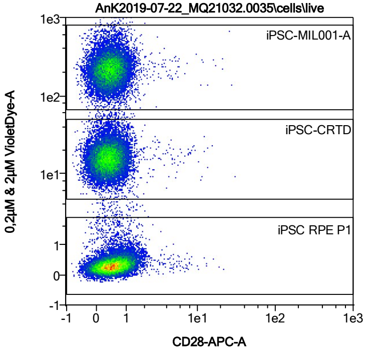


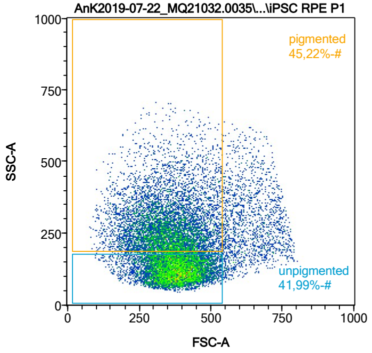


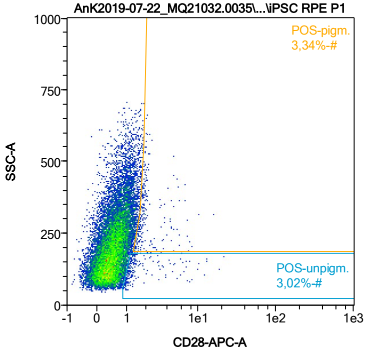


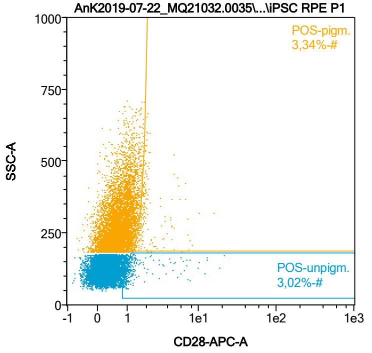


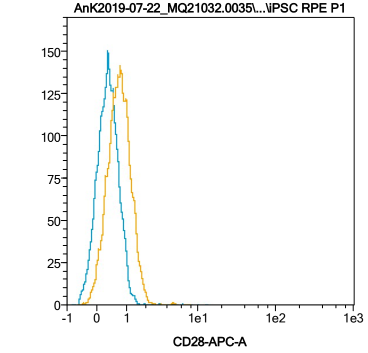


| Name | %-# | CD28-APC-A Mean | CD28-APC-A Median |
|---|---|---|---|
| POS-unpigm. | 3,02 | 2,14 | 1,19 |
| POS-pigm. | 3,34 | 2,37 | 1,85 |

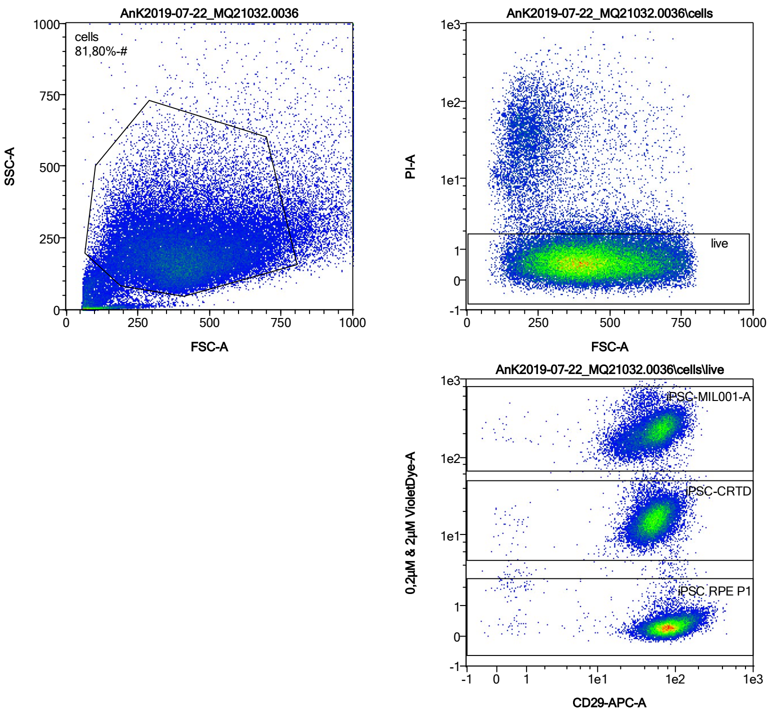


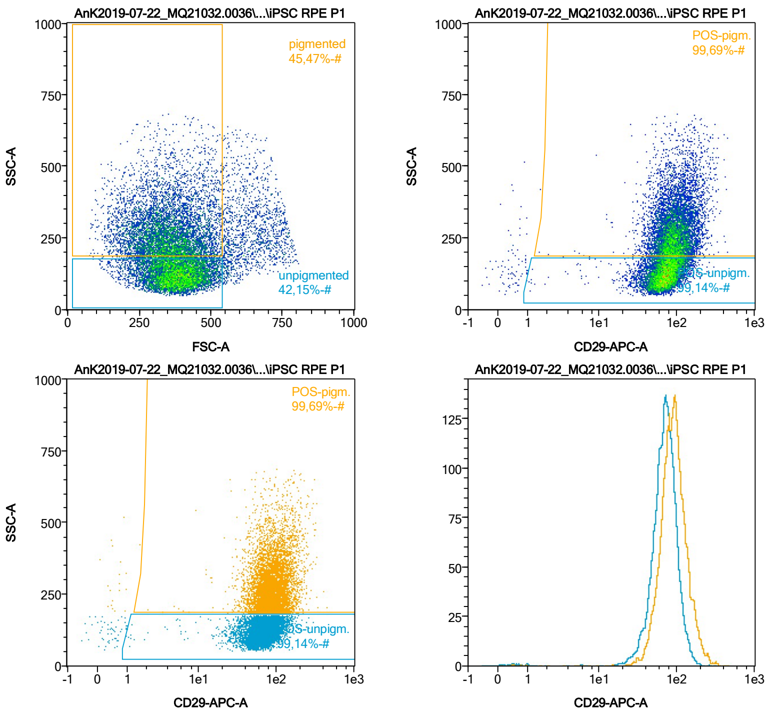


| Name | %-# | CD29-APC-A Mean | CD29-APC-A Median |
|---|---|---|---|
| POS-unpigm. | 99,14 | 72,40 | 69,94 |
| POS-pigm. | 99,69 | 96,14 | 89,02 |

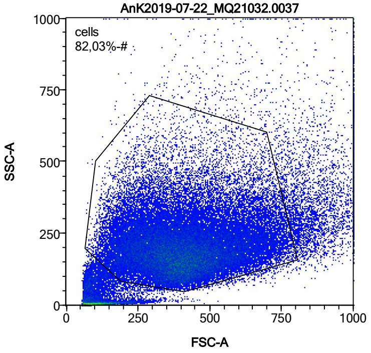


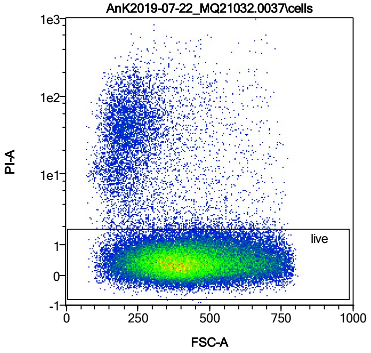


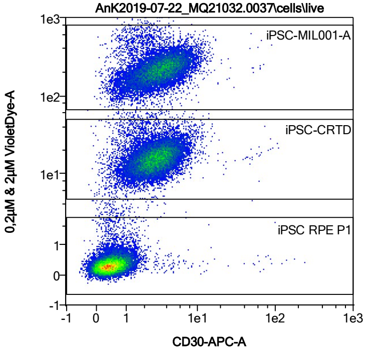


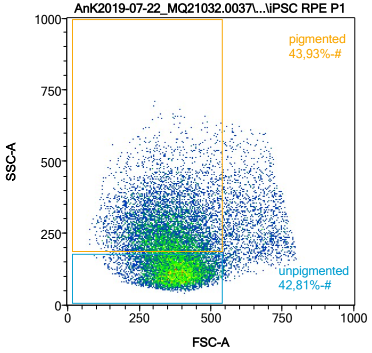


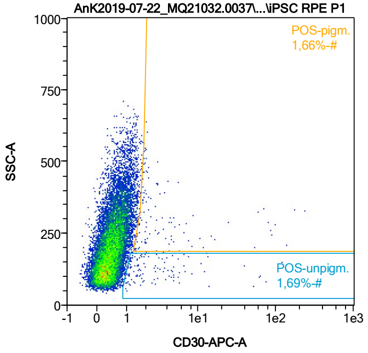


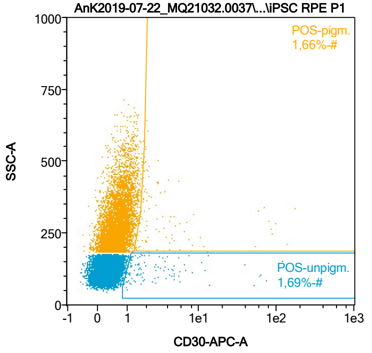


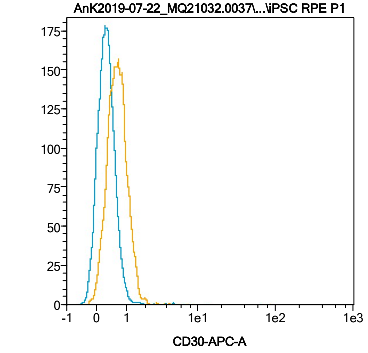


| Name | %-# | CD30-APC-A Mean | CD30-APC-A Median |
|---|---|---|---|
| POS-unpigm. | 1,69 | 6,30 | 1,69 |
| POS-pigm. | 1,66 | 10,29 | 2,04 |

Date:2024-06-14

Operator: Sebastian Knoebel

Version:2.13.0+54.g95abf0f4

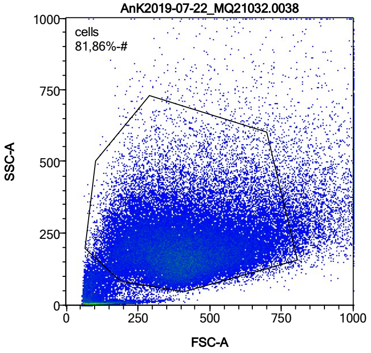


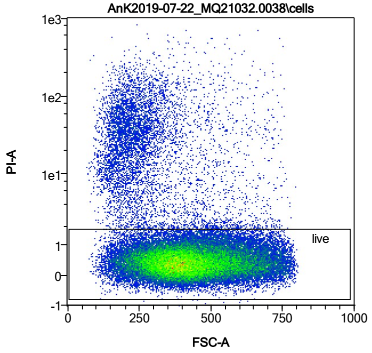


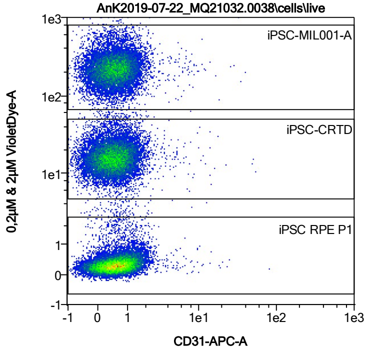


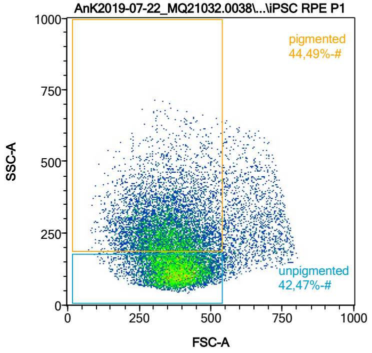


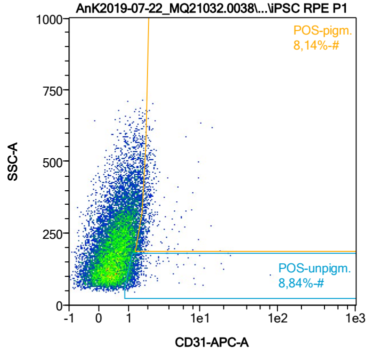


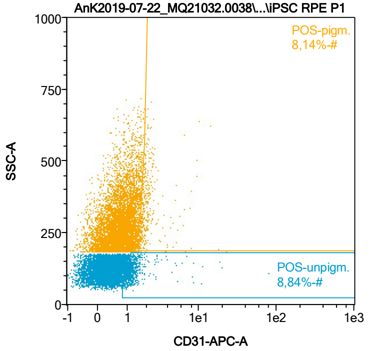


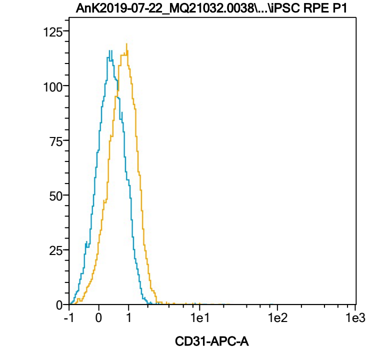


| Name | %-# | CD31-APC-A Mean | CD31-APC-A Median |
|---|---|---|---|
| POS-unpigm. | 8,84 | 1,59 | 1,18 |
| POS-pigm. | 8,14 | 2,09 | 1,79 |

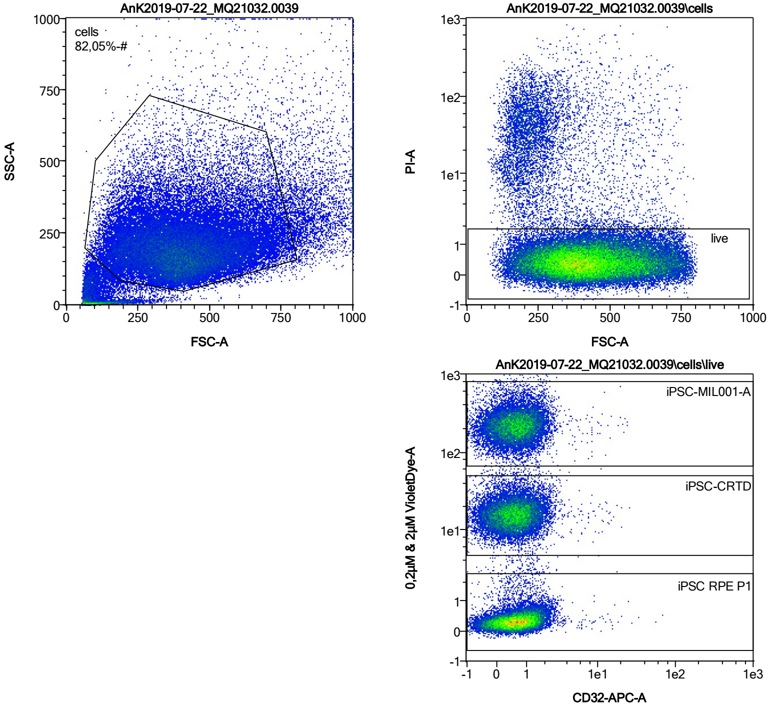


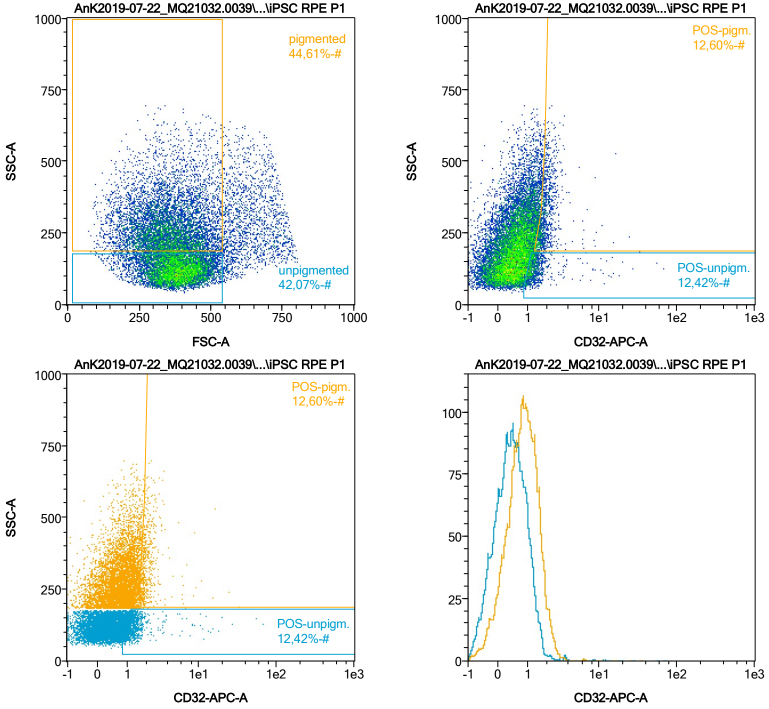


| Name | %-# | CD32-APC-A Mean | CD32-APC-A Median |
|---|---|---|---|
| POS-unpigm. | 12,42 | 1,77 | 1,25 |
| POS-pigm. | 12,60 | 2,08 | 1,82 |

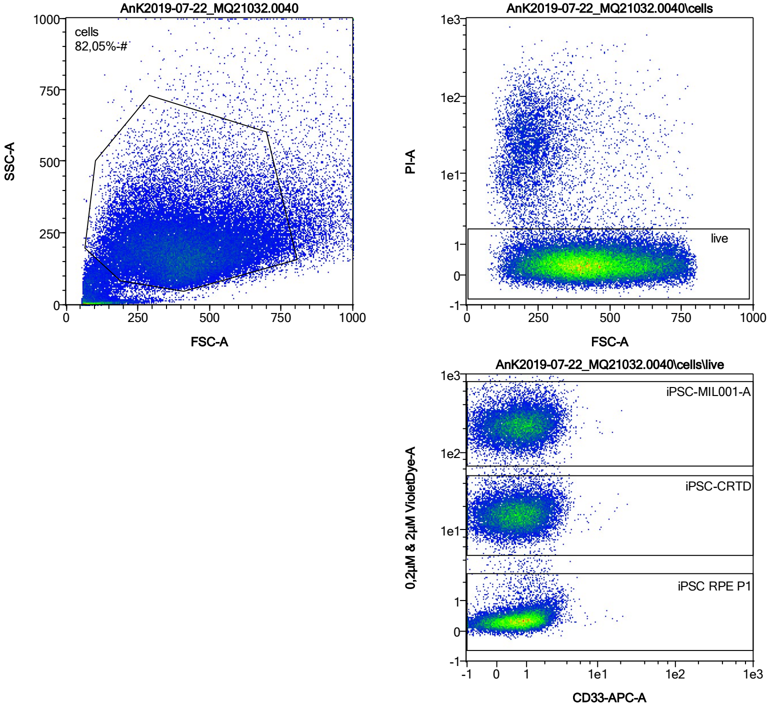


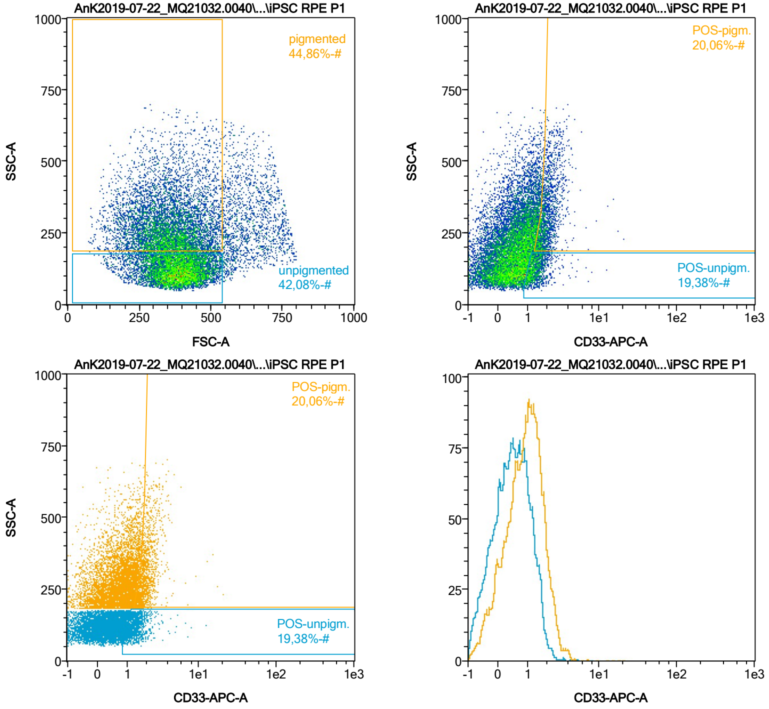


| Name | %-# | CD33-APC-A Mean | CD33-APC-A Median |
|---|---|---|---|
| POS-unpigm. | 19,38 | 1,43 | 1,30 |
| POS-pigm. | 20,06 | 2,10 | 1,90 |

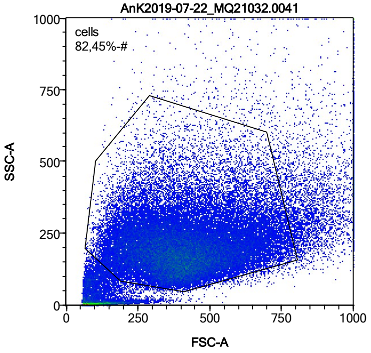


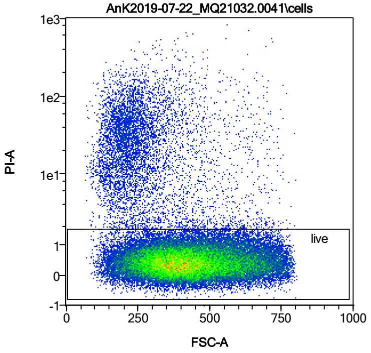


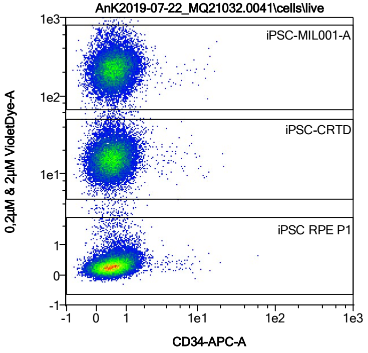


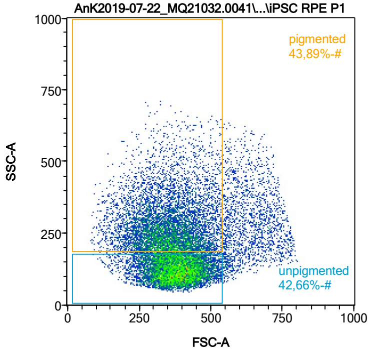


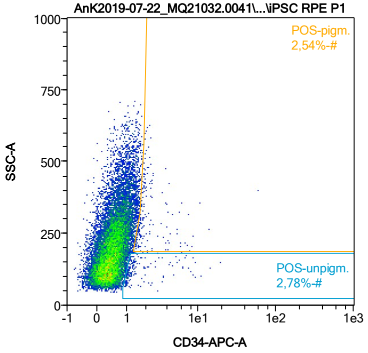


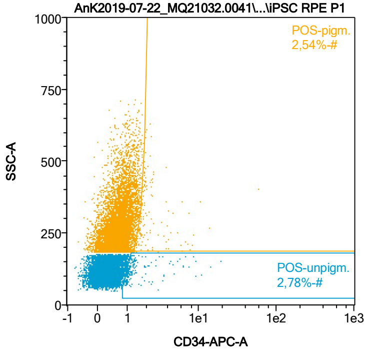


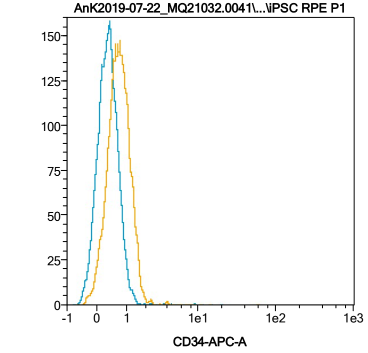


| Name | %-# | CD34-APC-A Mean | CD34-APC-A Median |
|---|---|---|---|
| POS-unpigm. | 2,78 | 1,91 | 1,18 |
| POS-pigm. | 2,54 | 2,69 | 1,80 |

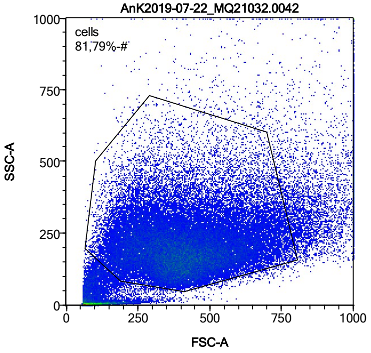


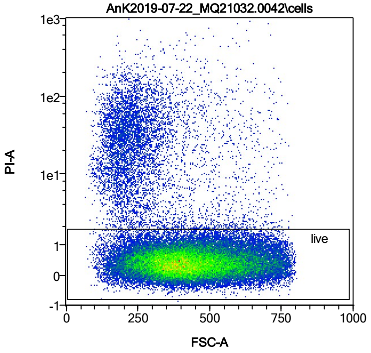


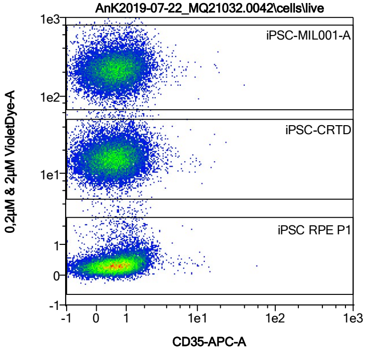


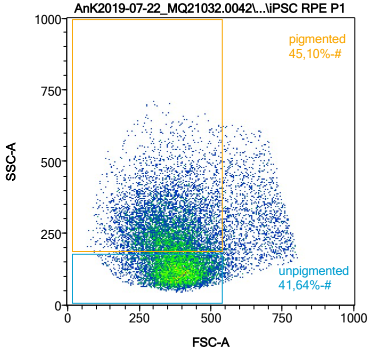


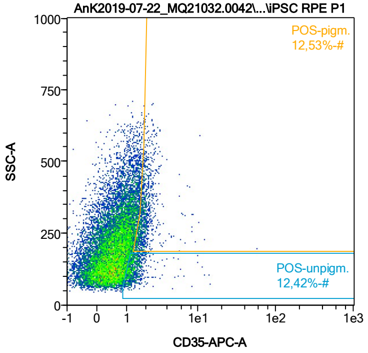


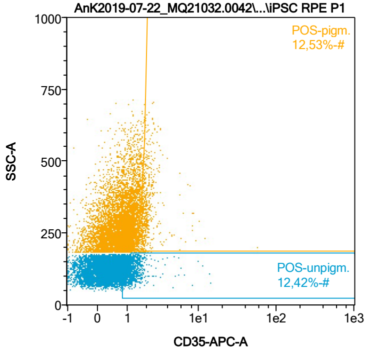


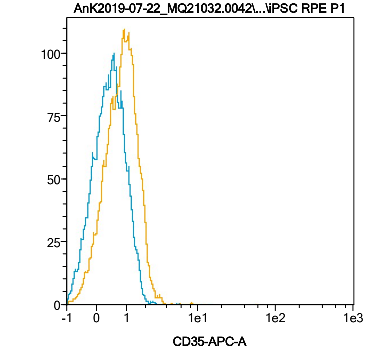


| Name | %-# | CD35-APC-A Mean | CD35-APC-A Median |
|---|---|---|---|
| POS-unpigm. | 12,42 | 1,42 | 1,25 |
| POS-pigm. | 12,53 | 2,09 | 1,85 |

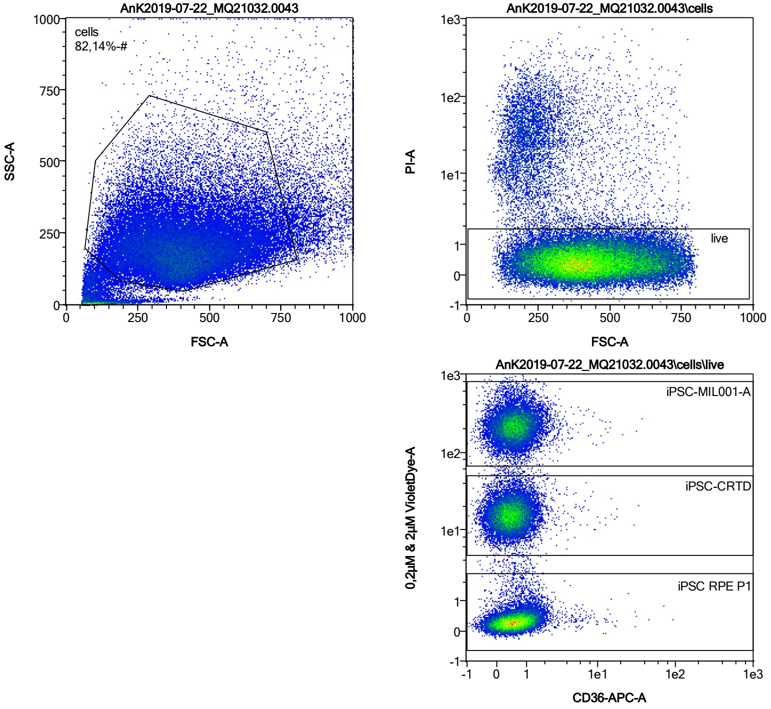


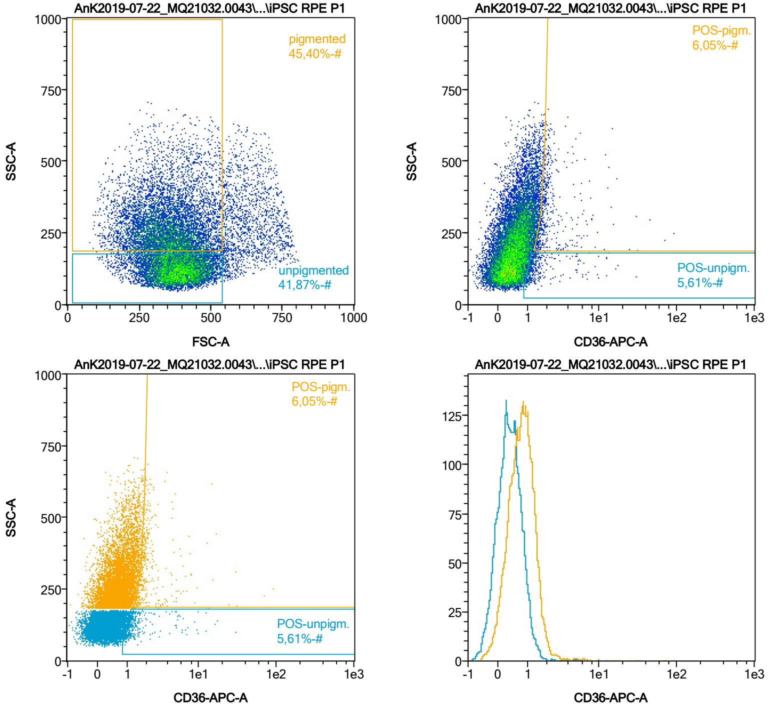


| Name | %-# | CD36-APC-A Mean | CD36-APC-A Median |
|---|---|---|---|
| POS-unpigm. | 5,61 | 1,81 | 1,20 |
| POS-pigm. | 6,05 | 2,93 | 1,86 |

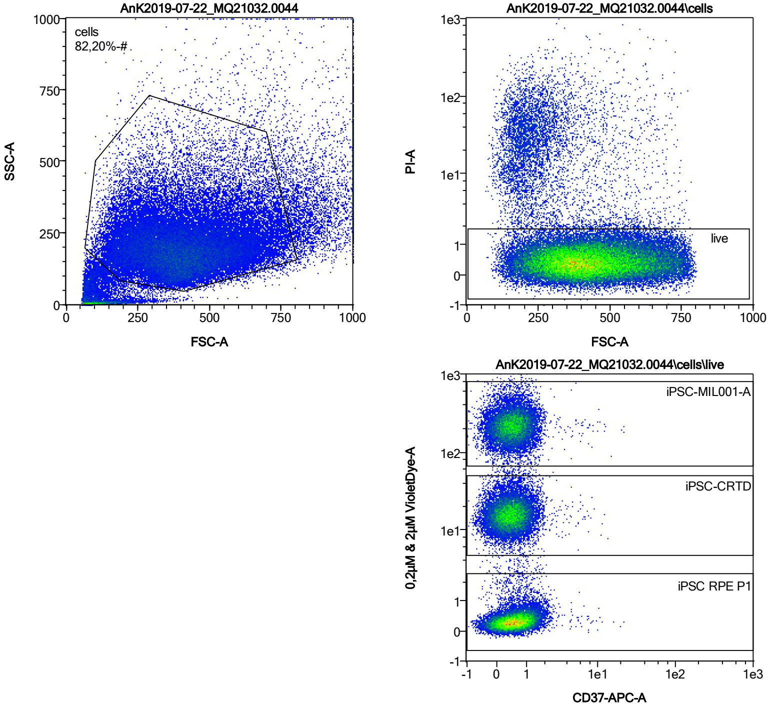


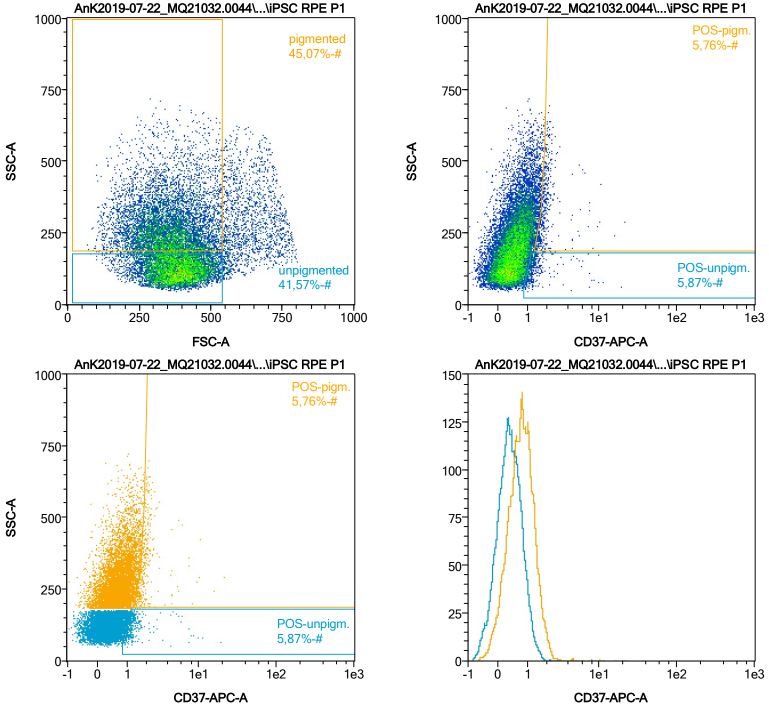


| Name | %-# | CD37-APC-A Mean | CD37-APC-A Median |
|---|---|---|---|
| POS-unpigm. | 5,87 | 1,48 | 1,17 |
| POS-pigm. | 5,76 | 2,12 | 1,84 |

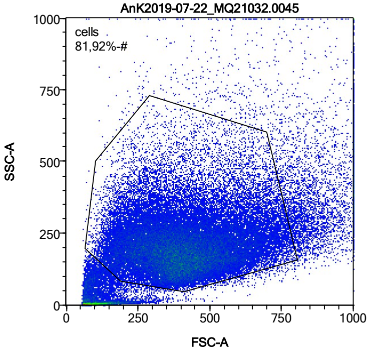


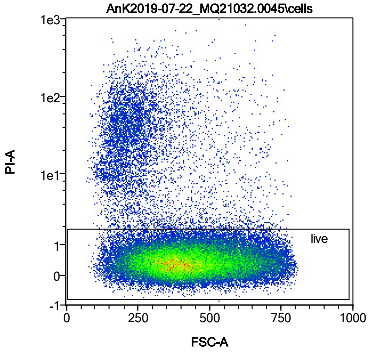


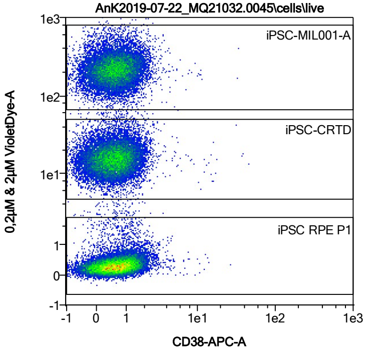


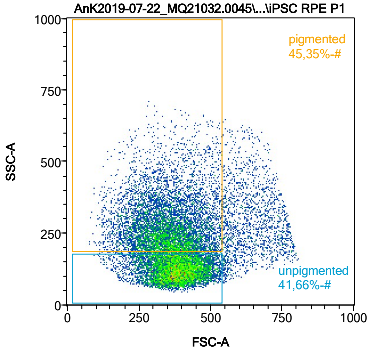


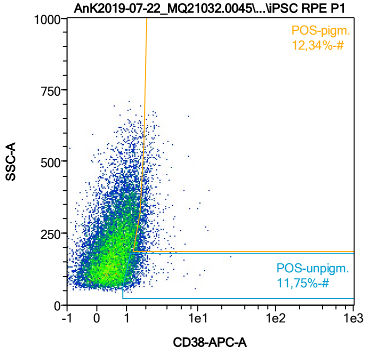


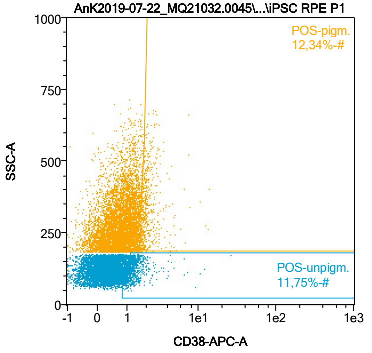


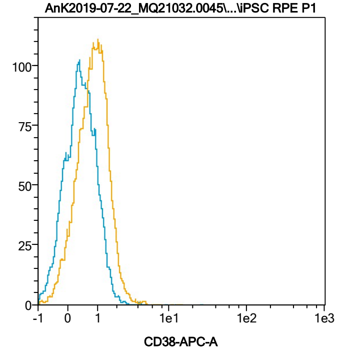


| Name | %-# | CD38-APC-A Mean | CD38-APC-A Median |
|---|---|---|---|
| POS-unpigm. | 11,75 | 1,45 | 1,24 |
| POS-pigm. | 12,34 | 2,06 | 1,83 |

Date:2024-06-14
Operator: Sebastian Knoebel
Version:2.13.0+54.g95abf0f4

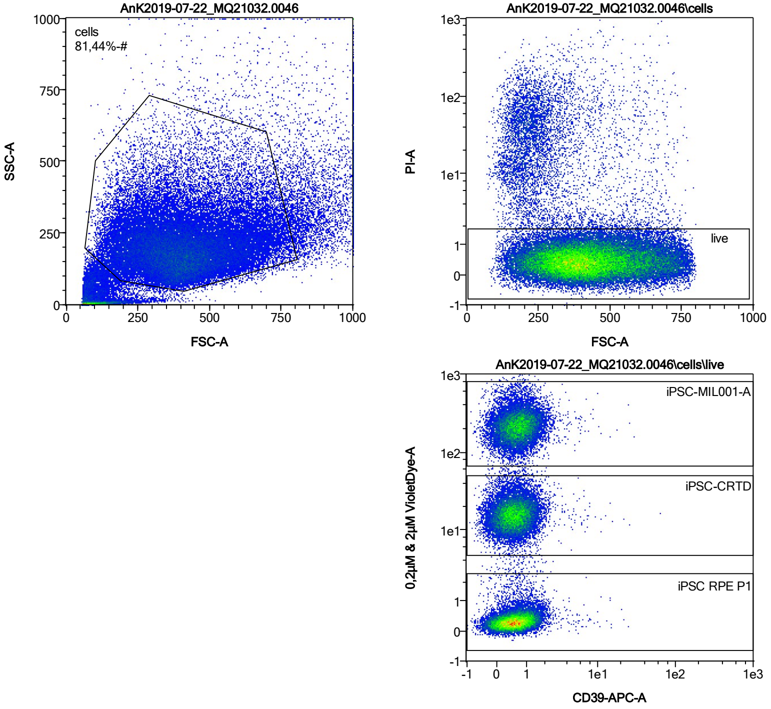


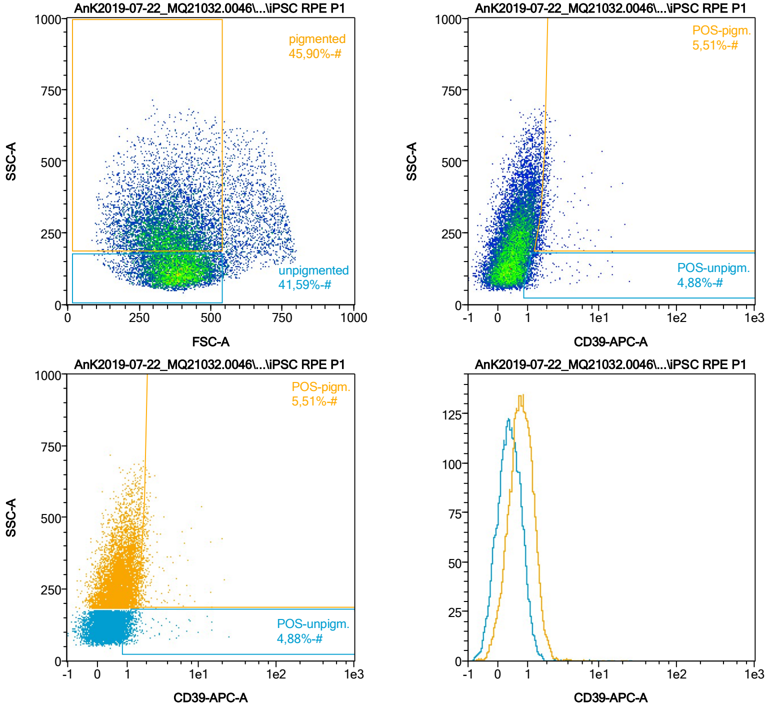


| Name | %-# | CD39-APC-A Mean | CD39-APC-A Median |
|---|---|---|---|
| POS-unpigm. | 4,88 | 1,62 | 1,17 |
| POS-pigm. | 5,51 | 2,26 | 1,80 |

Date:2024-06-14
Operator: Sebastian Knoebel
Version:2.13.0+54.g95abf0f4

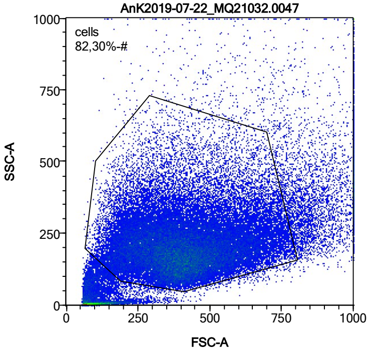


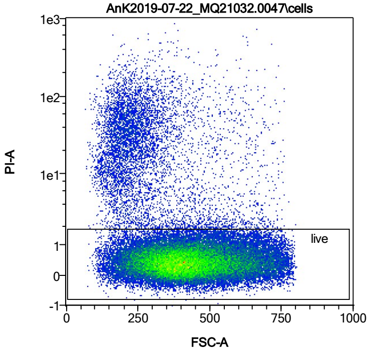


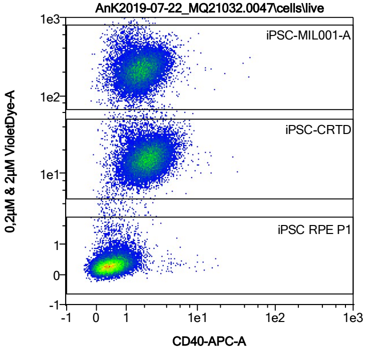


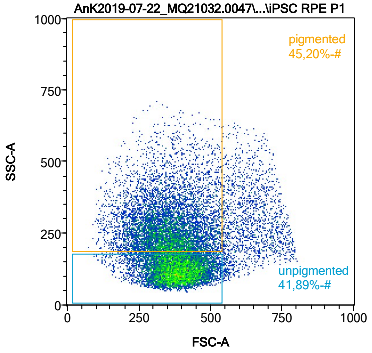


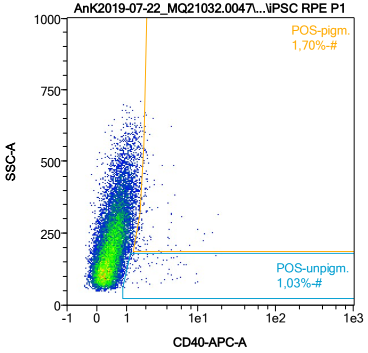


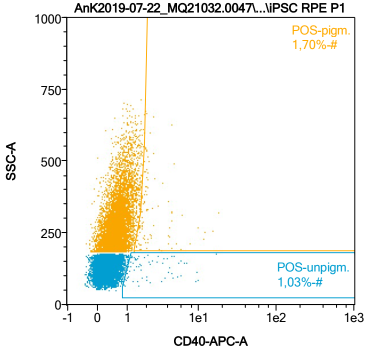


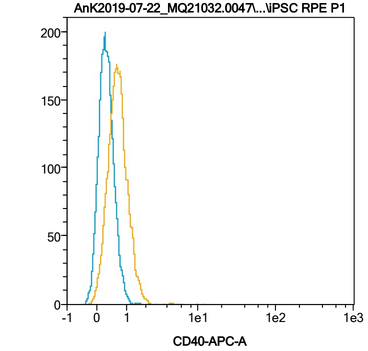


| Name | %-# | CD40-APC-A Mean | CD40-APC-A Median |
|---|---|---|---|
| POS-unpigm. | 1,03 | 2,92 | 1,71 |
| POS-pigm. | 1,70 | 2,77 | 1,93 |

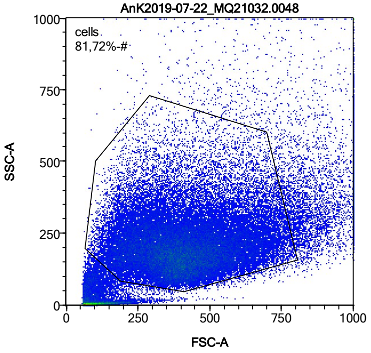


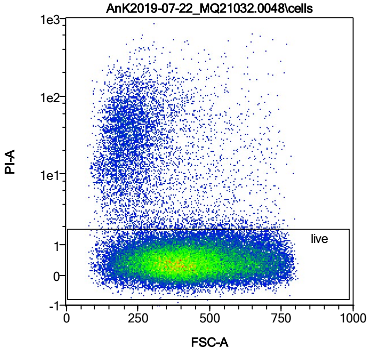


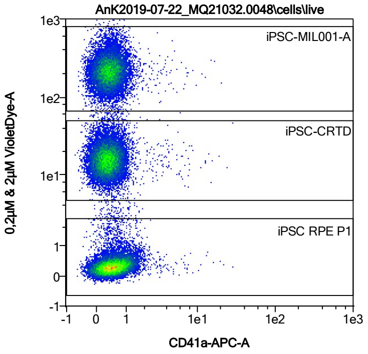


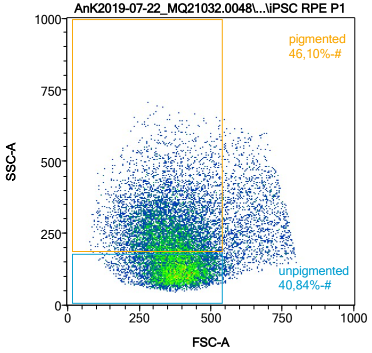


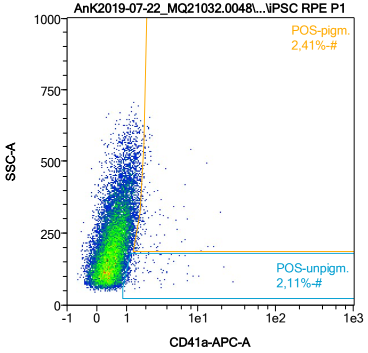


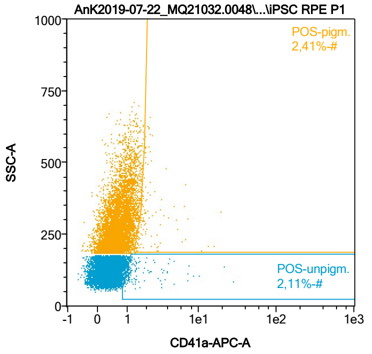


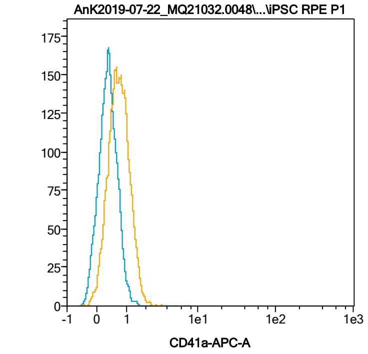


| Name | %-# | CD41a-APC-A Mean | CD41a-APC-A Median |
|---|---|---|---|
| POS-unpigm. | 2,11 | 2,16 | 1,21 |
| POS-pigm. | 2,41 | 2,40 | 1,80 |

Date:2024-06-14

Operator: Sebastian Knoebel

Version:2.13.0+54.g95abf0f4

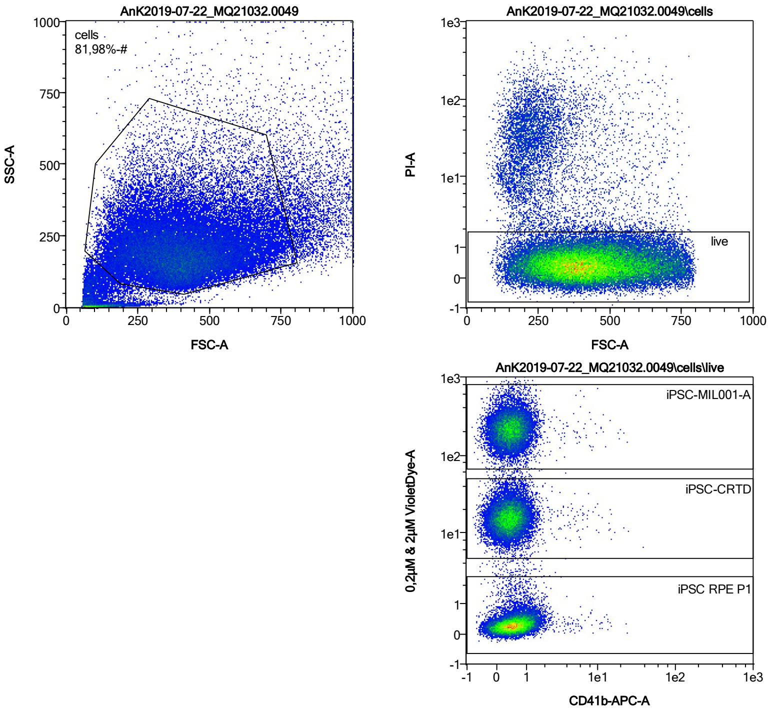


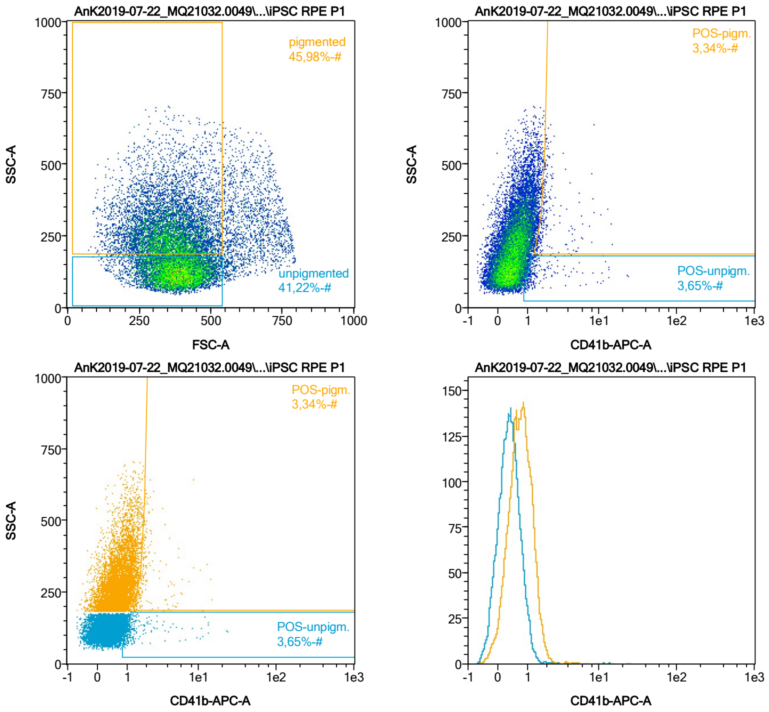


| Name | %-# | CD41b-APC-A Mean | CD41b-APC-A Median |
|---|---|---|---|
| POS-unpigm. | 3,65 | 2,03 | 1,19 |
| POS-pigm. | 3,34 | 2,43 | 1,85 |

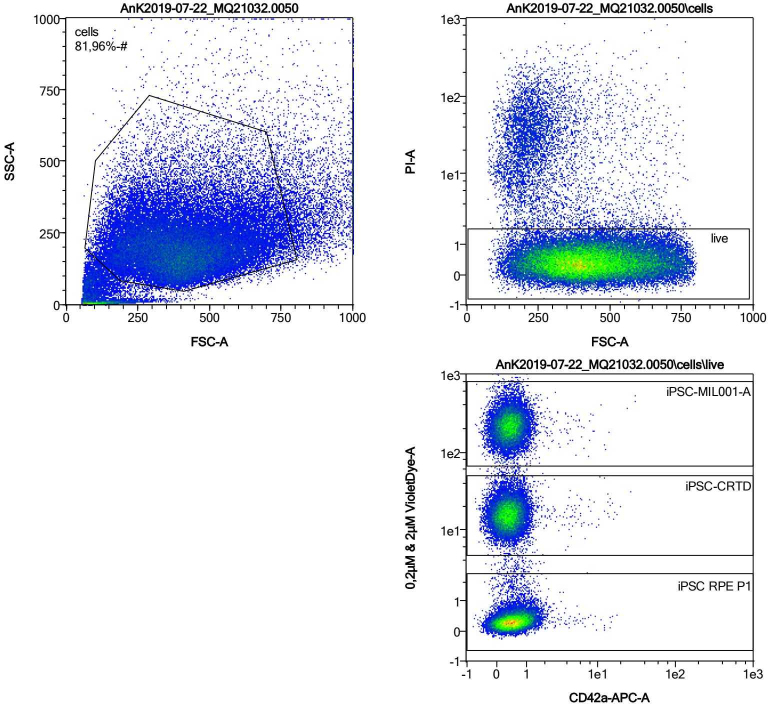


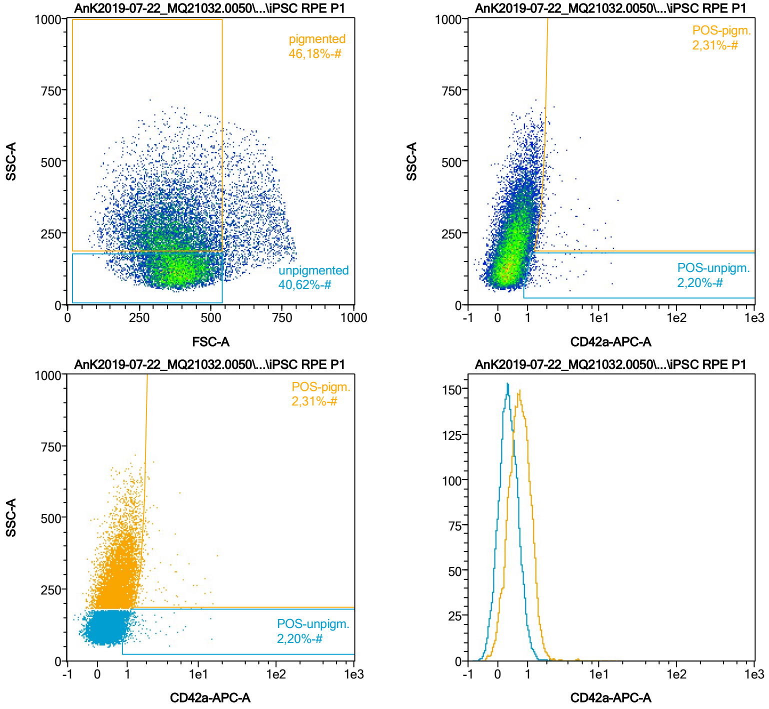


| Name | %-# | CD42a-APC-A Mean | CD42a-APC-A Median |
|---|---|---|---|
| POS-unpigm. | 2,20 | 1,70 | 1,21 |
| POS-pigm. | 2,31 | 2,85 | 1,87 |

Date:2024-06-14

Operator: Sebastian Knoebel

Version:2.13.0+54.g95abf0f4

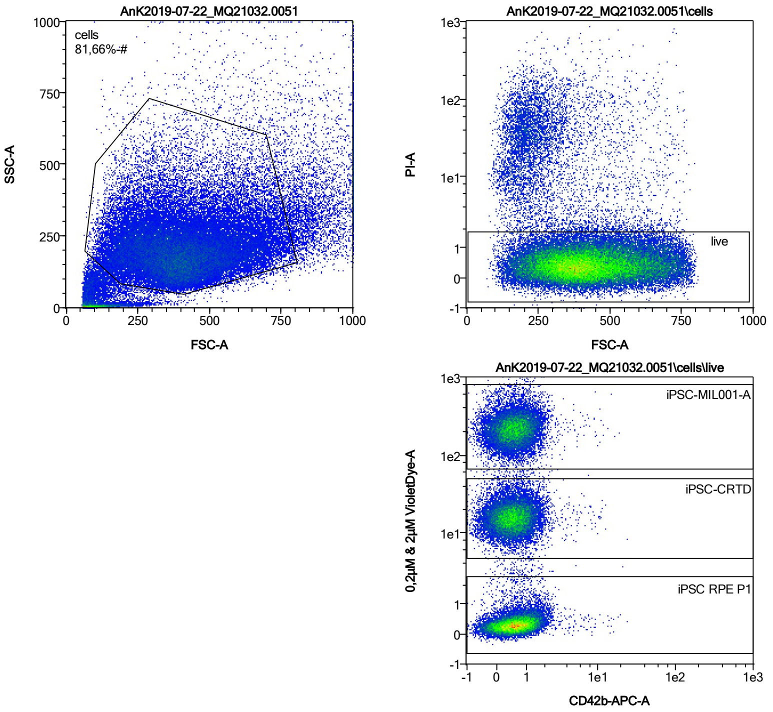


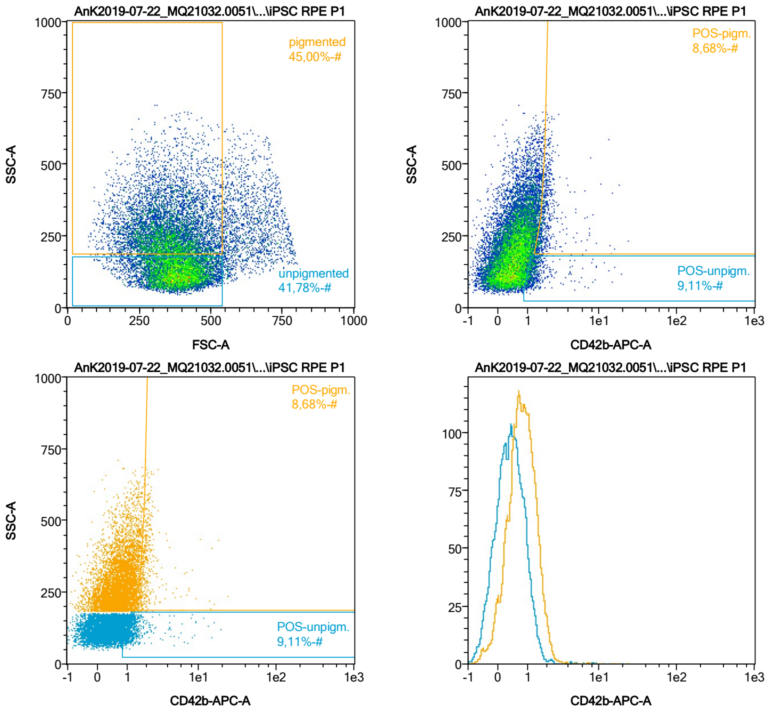


| Name | %-# | CD42b-APC-A Mean | CD42b-APC-A Median |
|---|---|---|---|
| POS-unpigm. | 9,11 | 1,59 | 1,22 |
| POS-pigm. | 8,68 | 2,17 | 1,82 |

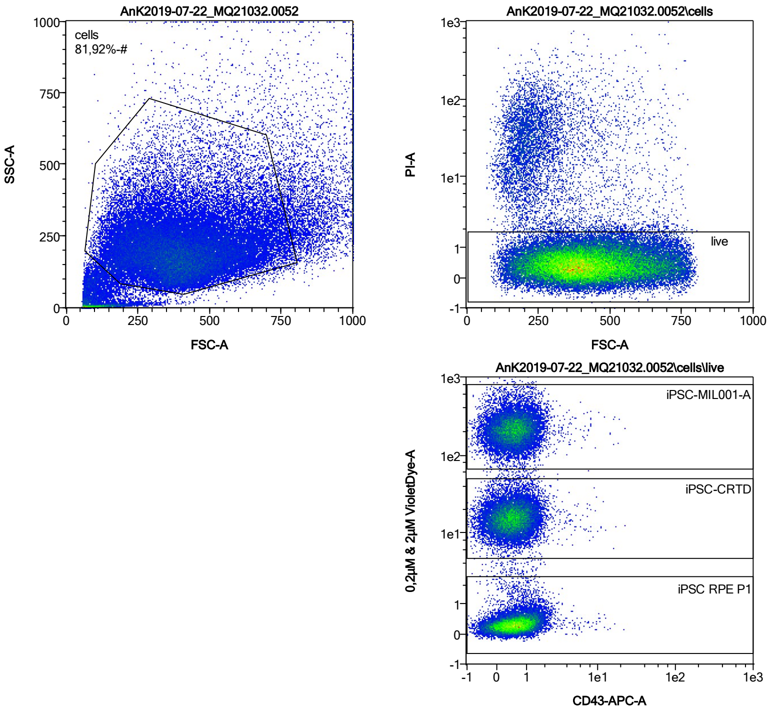


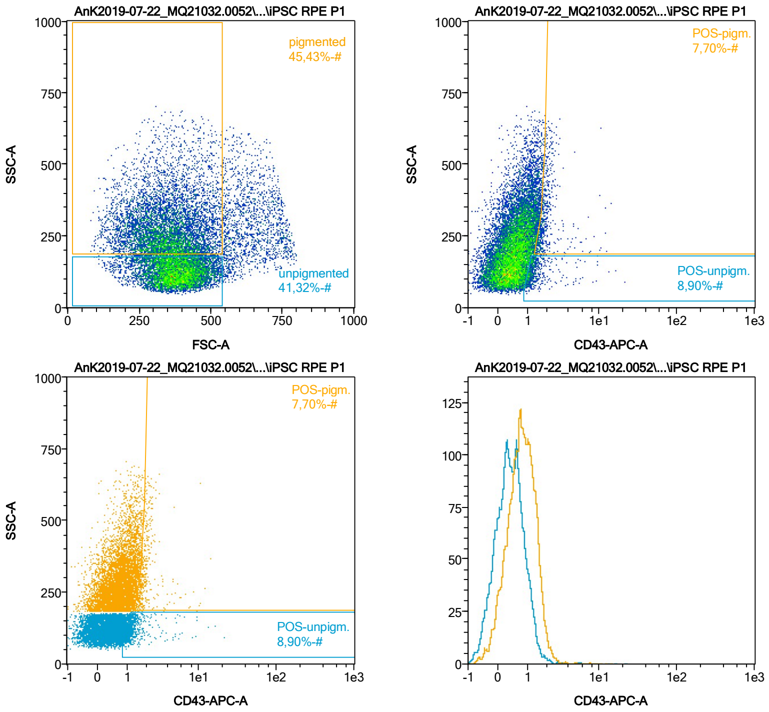


| Name | %-# | CD43-APC-A Mean | CD43-APC-A Median |
|---|---|---|---|
| POS-unpigm. | 8,90 | 1,56 | 1,22 |
| POS-pigm. | 7,70 | 2,08 | 1,78 |

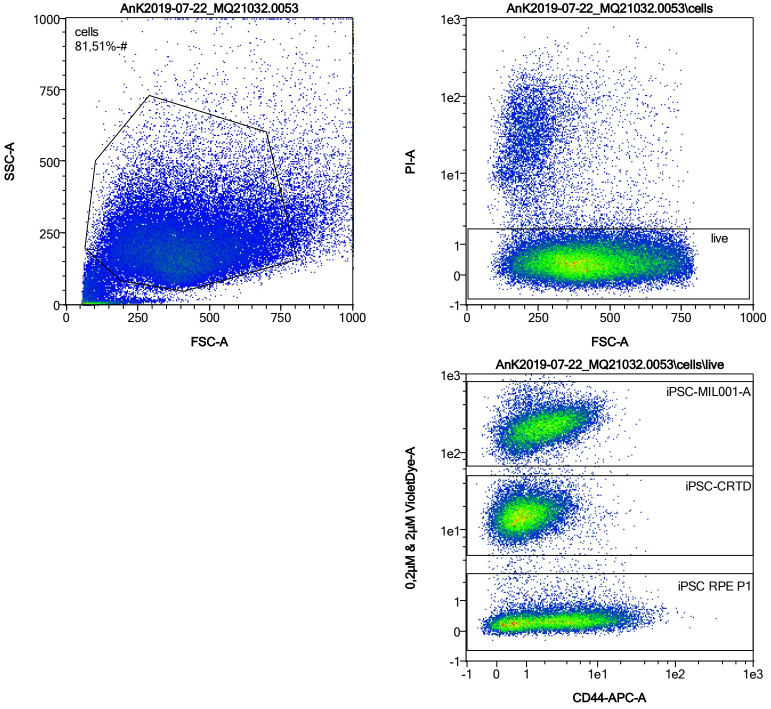


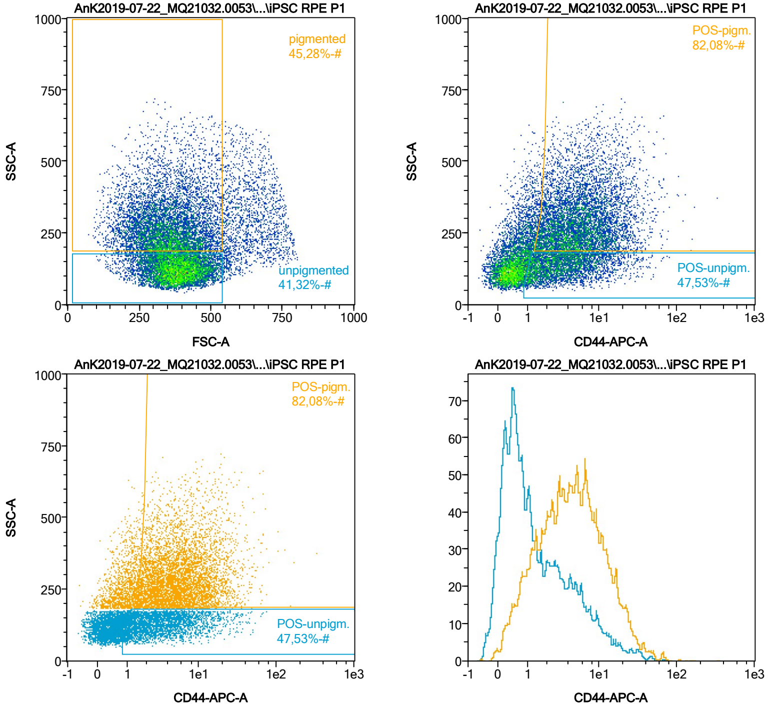


| Name | %-# | CD44-APC-A Mean | CD44-APC-A Median |
|---|---|---|---|
| POS-unpigm. | 47,53 | 4,98 | 2,94 |
| POS-pigm. | 82,08 | 8,01 | 5,31 |

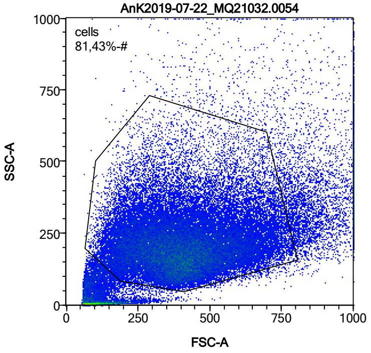


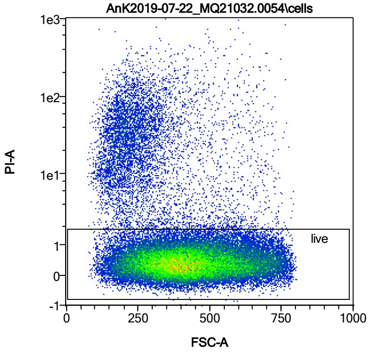


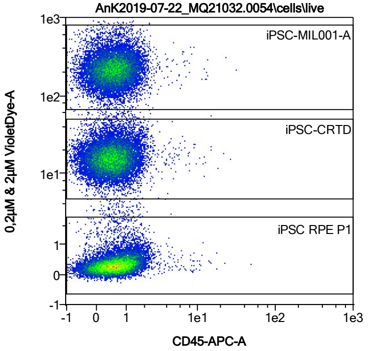


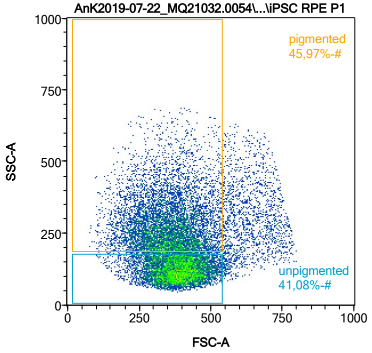


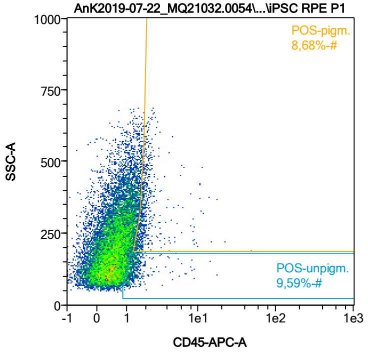


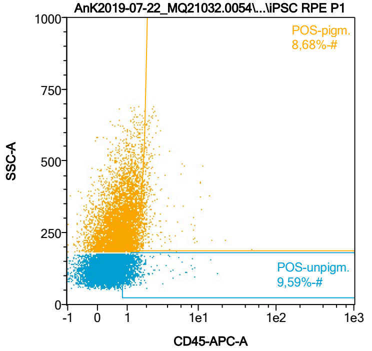


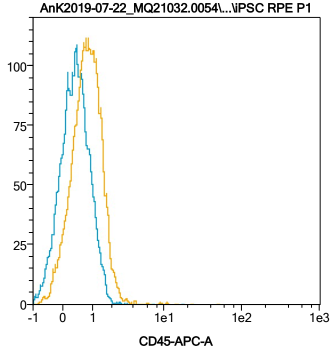


| Name | %-# | CD45-APC-A Mean | CD45-APC-A Median |
|---|---|---|---|
| POS-unpigm. | 9,59 | 1,48 | 1,23 |
| POS-pigm. | 8,68 | 2,31 | 1,84 |

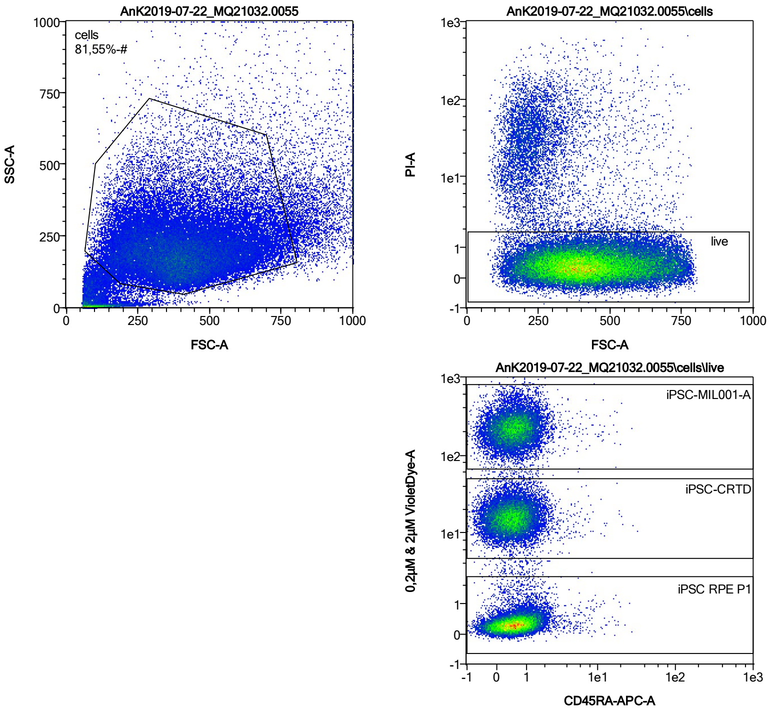


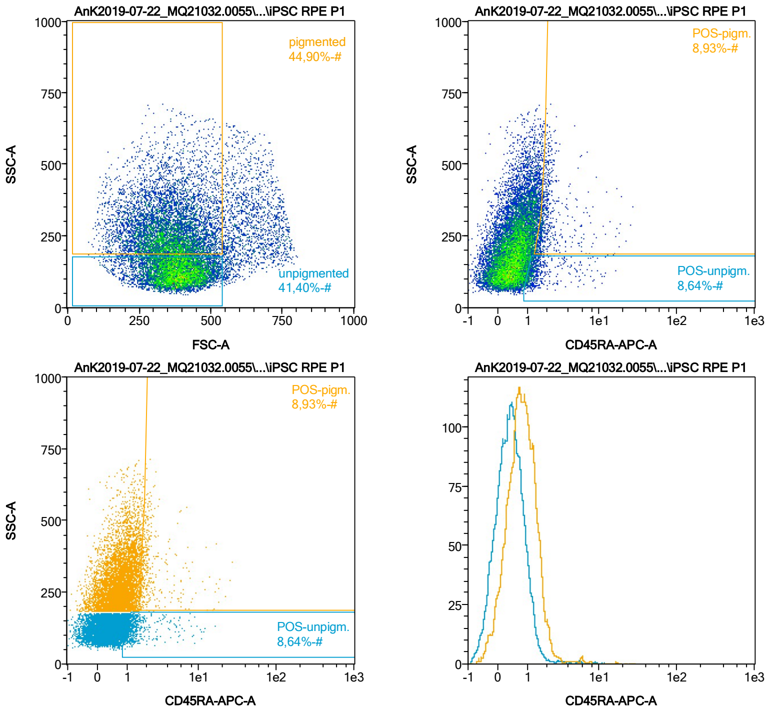


| Name | %-# | CD45RA-APC-A Mean | CD45RA-APC-A Median |
|---|---|---|---|
| POS-unpigm. | 8,64 | 1,77 | 1,25 |
| POS-pigm. | 8,93 | 2,61 | 1,85 |

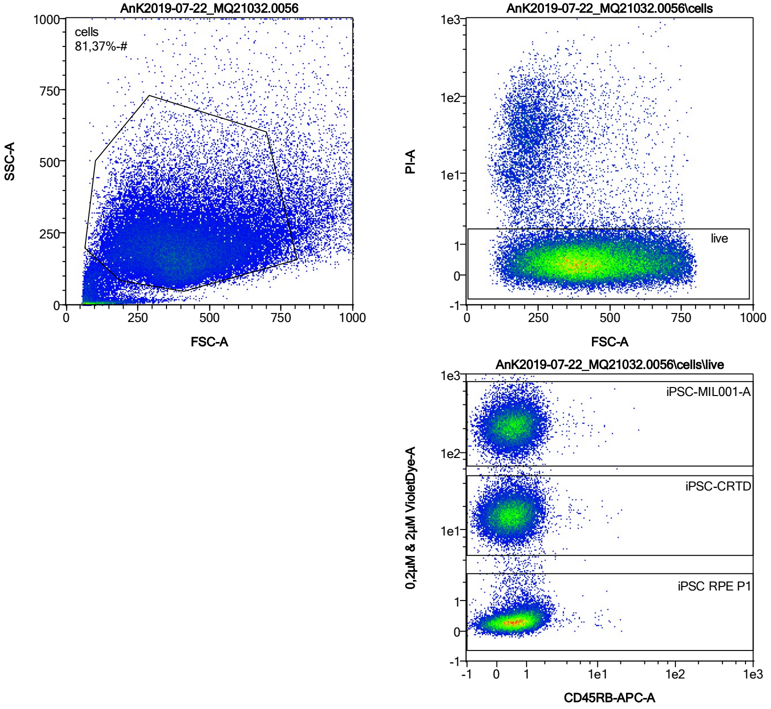


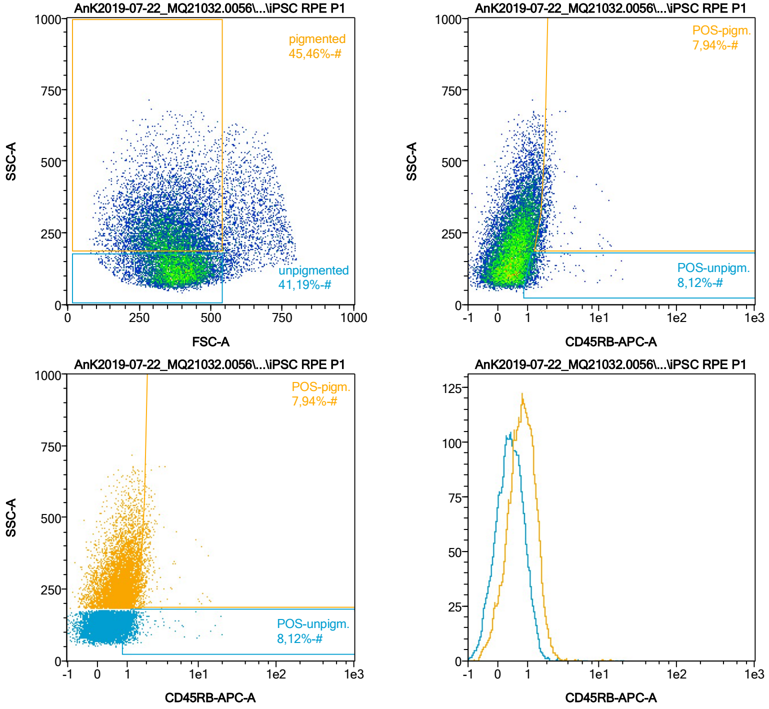


| Name | %-# | CD45RB-APC-A Mean | CD45RB-APC-A Median |
|---|---|---|---|
| POS-unpigm. | 8,12 | 1,54 | 1,23 |
| POS-pigm. | 7,94 | 2,08 | 1,80 |

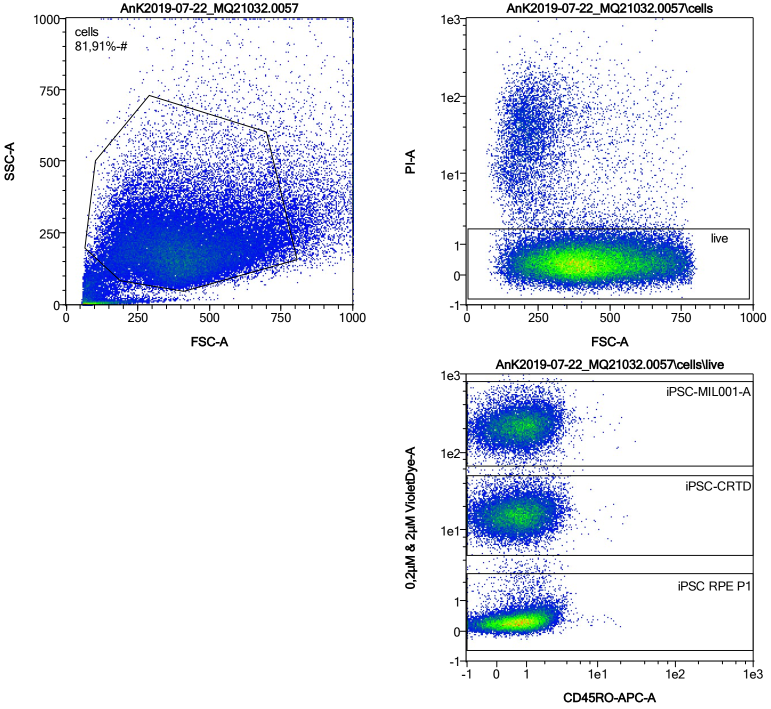


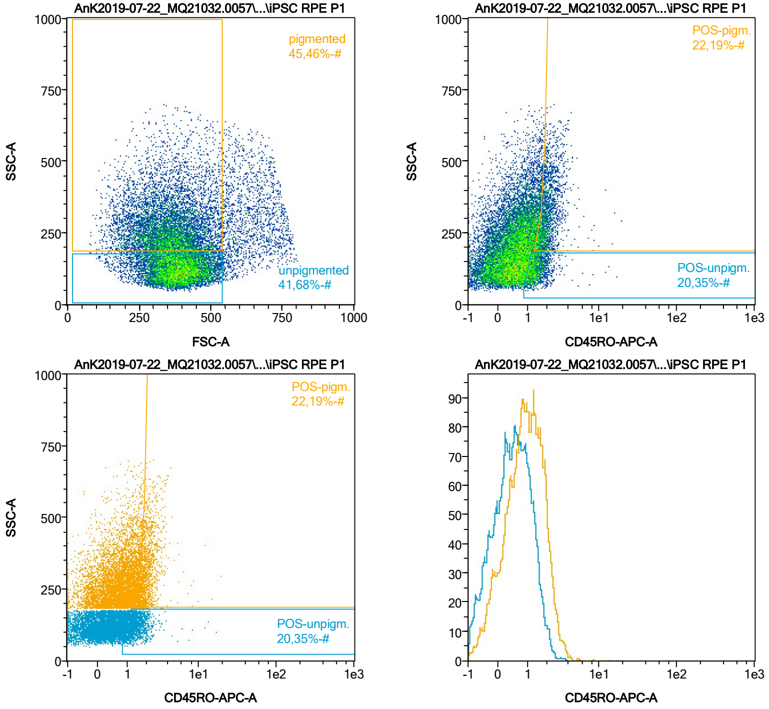


| Name | %-# | CD45RO-APC-A Mean | CD45RO-APC-A Median |
|---|---|---|---|
| POS-unpigm. | 20,35 | 1,47 | 1,32 |
| POS-pigm. | 22,19 | 2,10 | 1,93 |

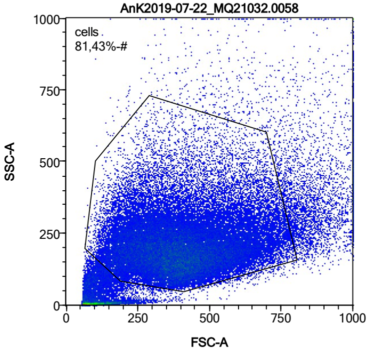


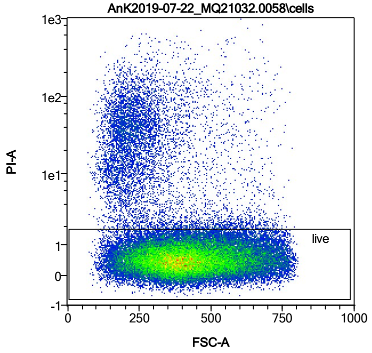


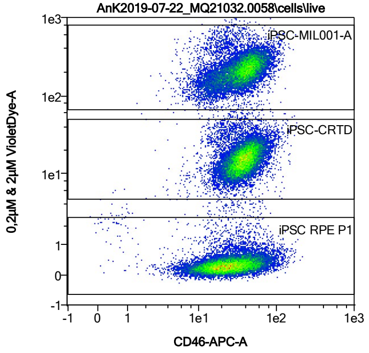


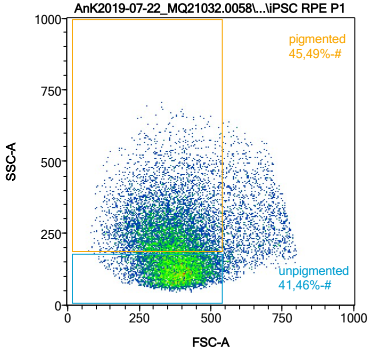


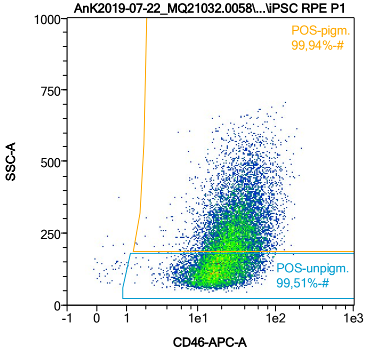


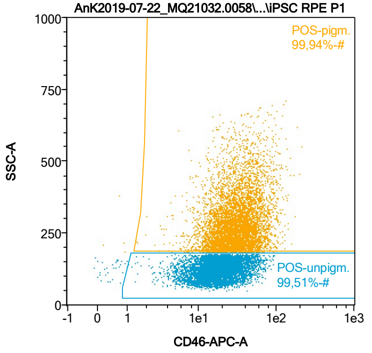


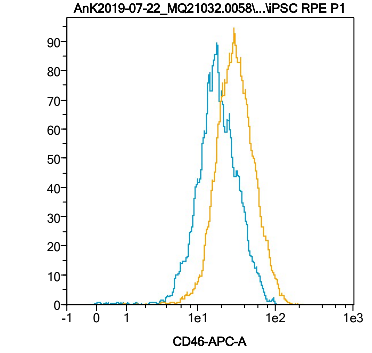


| Name | %-# | CD46-APC-A Mean | CD46-APC-A Median |
|---|---|---|---|
| POS-unpigm. | 99,51 | 20,96 | 17,66 |
| POS-pigm. | 99,94 | 34,77 | 29,71 |

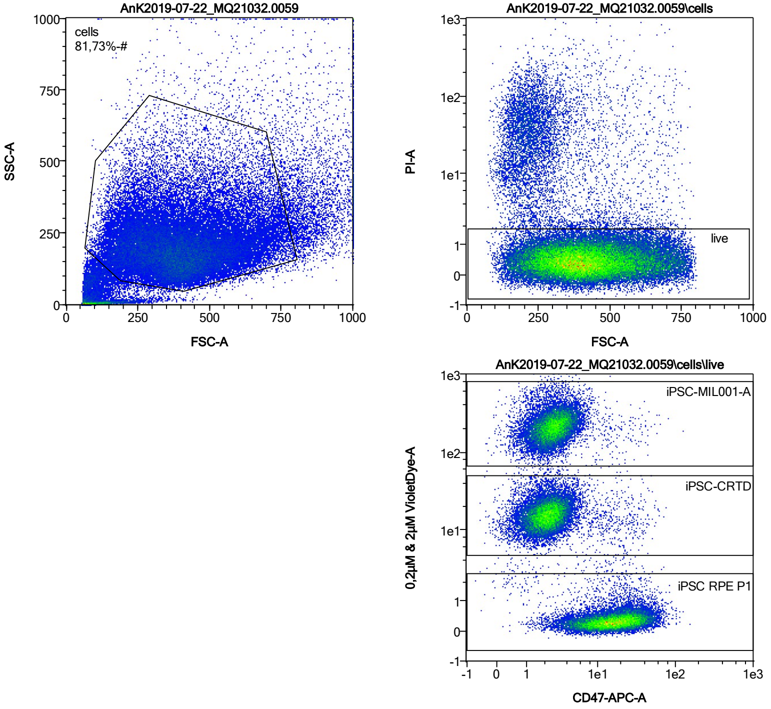


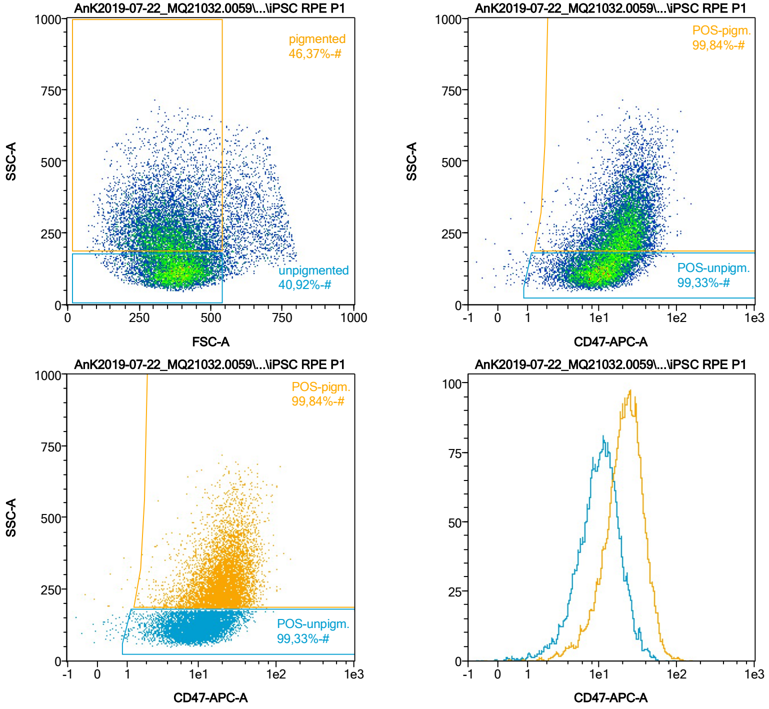


| Name | %-# | CD47-APC-A Mean | CD47-APC-A Median |
|---|---|---|---|
| POS-unpigm. | 99,33 | 11,73 | 10,30 |
| POS-pigm. | 99,84 | 23,80 | 21,78 |

Date:2024-06-14

Operator: Sebastian Knoebel

Version:2.13.0+54.g95abf0f4

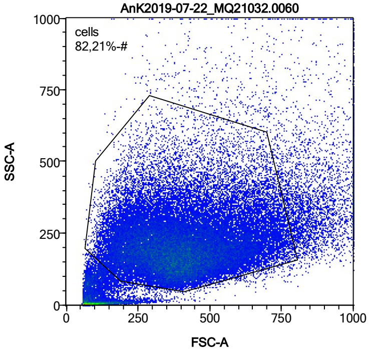


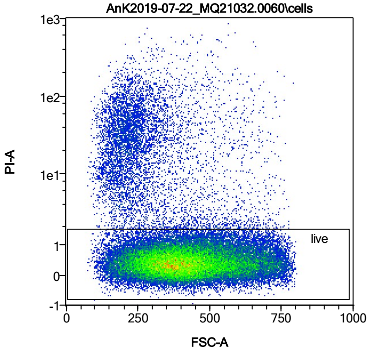


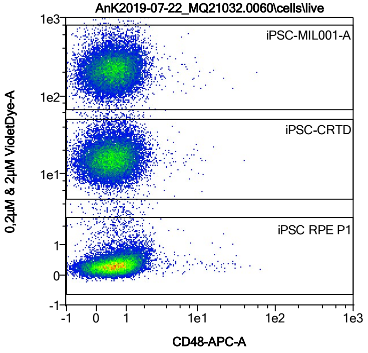


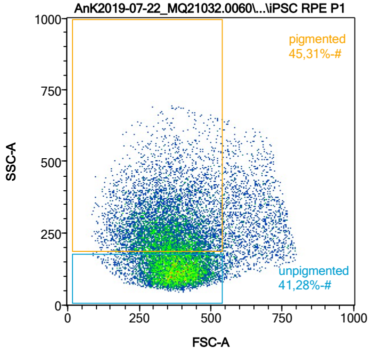


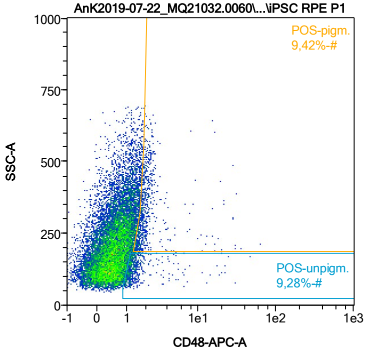


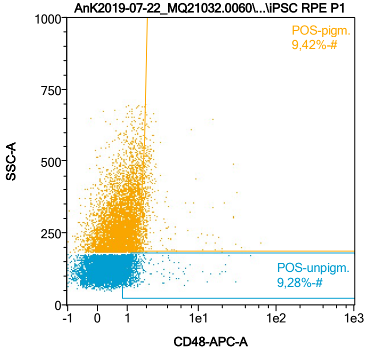


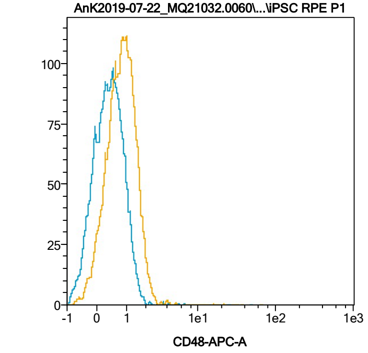


| Name | %-# | CD48-APC-A Mean | CD48-APC-A Median |
|---|---|---|---|
| POS-unpigm. | 9,28 | 1,79 | 1,22 |
| POS-pigm. | 9,42 | 2,65 | 1,85 |

Date:2024-06-14

Operator: Sebastian Knoebel

Version:2.13.0+54.g95abf0f4

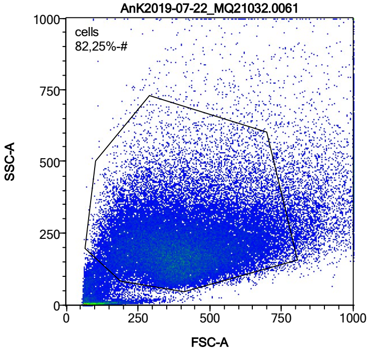


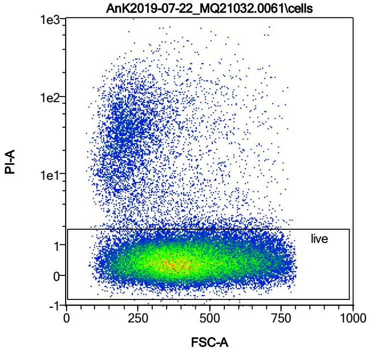


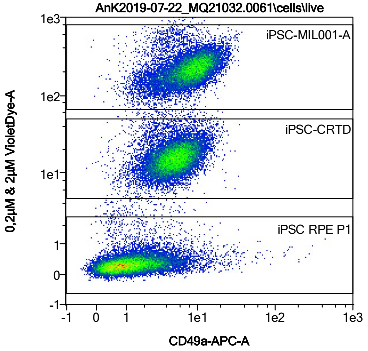


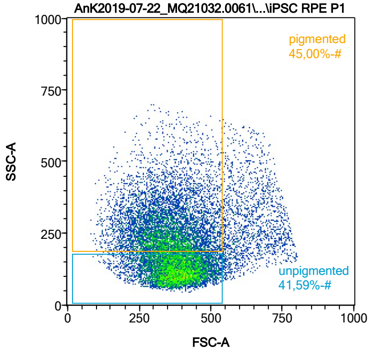


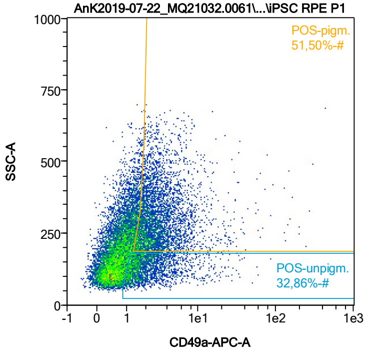


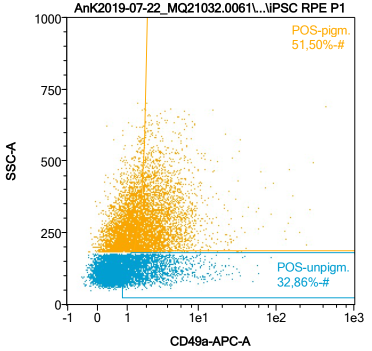


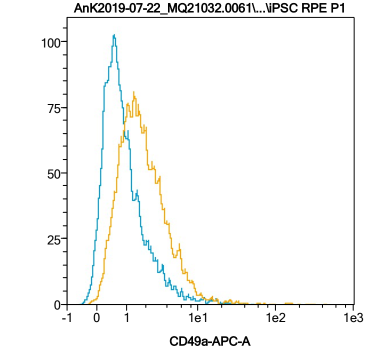


| Name | %-# | CD49a-APC-A Mean | CD49a-APC-A Median |
|---|---|---|---|
| POS-unpigm. | 32,86 | 2,81 | 1,67 |
| POS-pigm. | 51,50 | 4,41 | 2,65 |

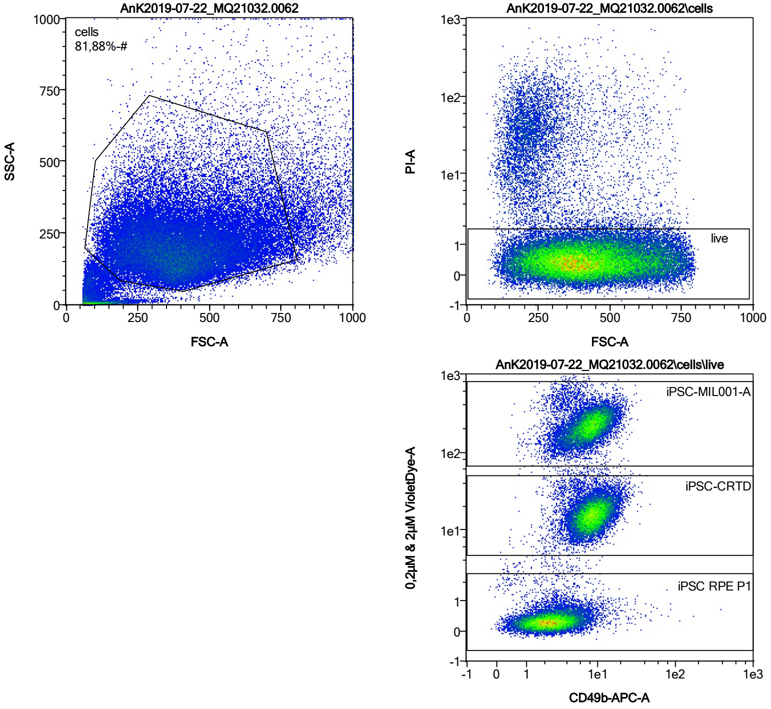


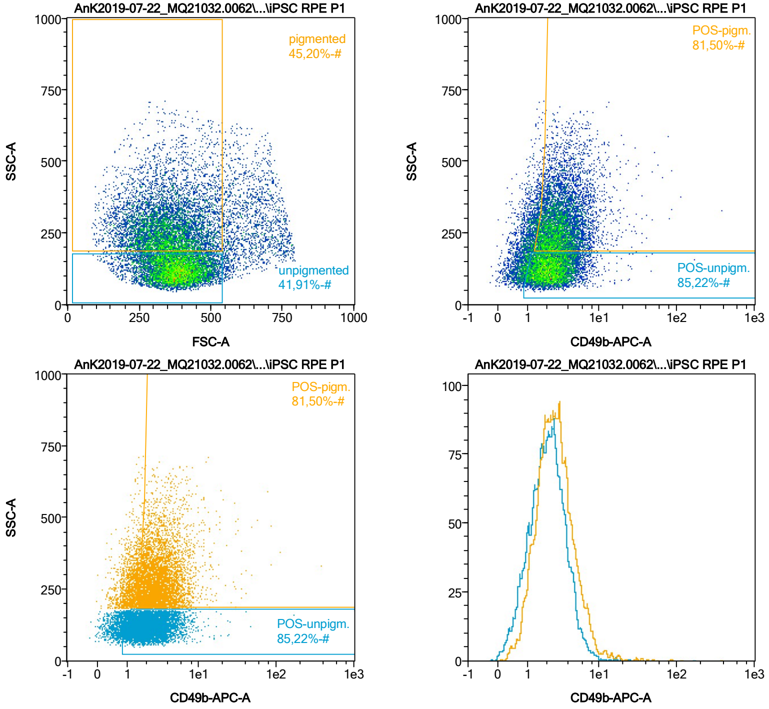


| Name | %-# | CD49b-APC-A Mean | CD49b-APC-A Median |
|---|---|---|---|
| POS-unpigm. | 85,22 | 2,66 | 2,26 |
| POS-pigm. | 81,50 | 3,62 | 2,84 |

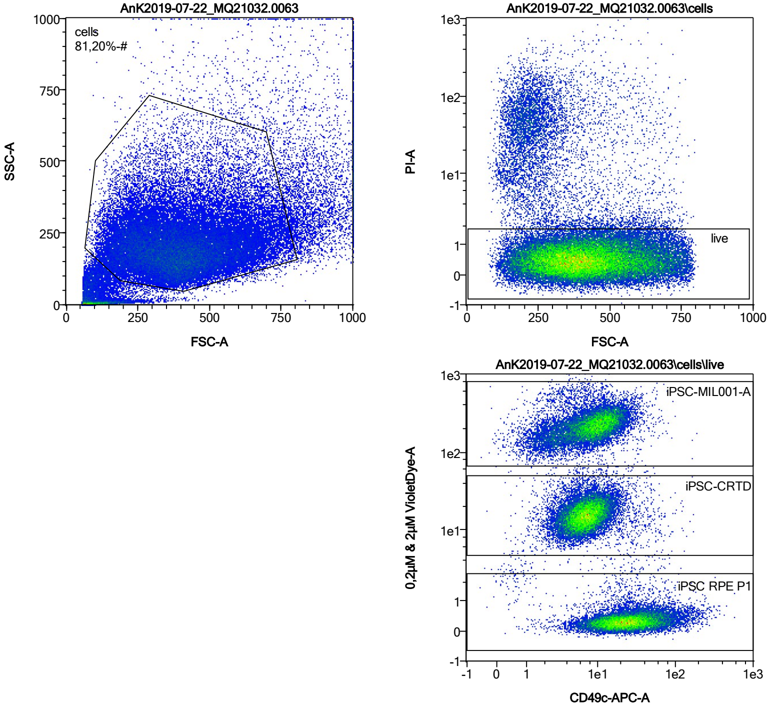


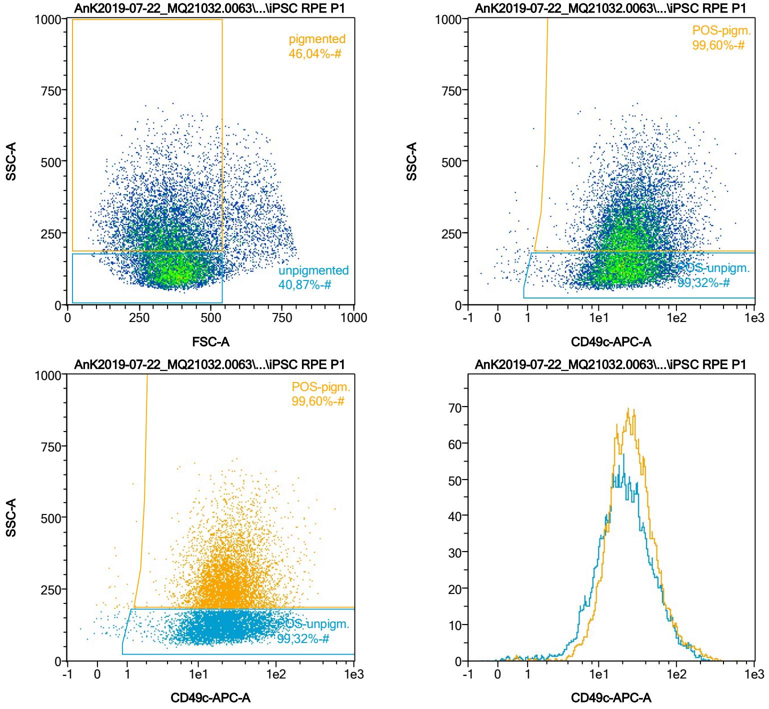


| Name | %-# | CD49c-APC-A Mean | CD49c-APC-A Median |
|---|---|---|---|
| POS-unpigm. | 99,32 | 29,08 | 21,12 |
| POS-pigm. | 99,60 | 32,83 | 24,43 |

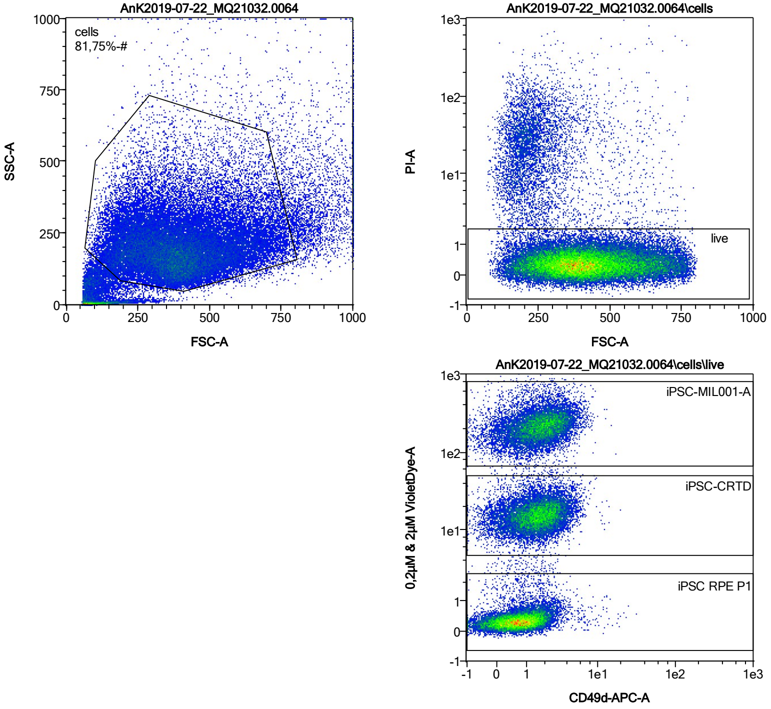


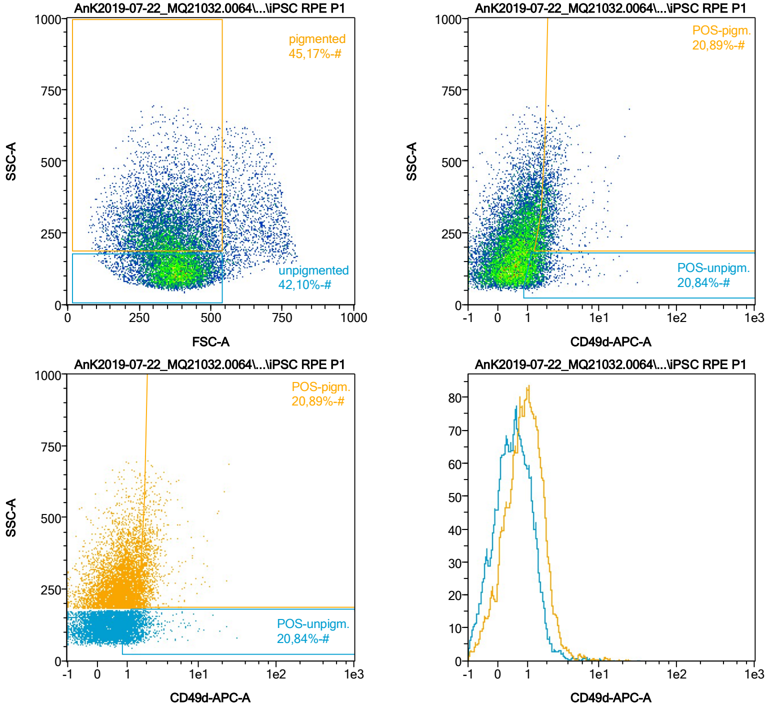


| Name | %-# | CD49d-APC-A Mean | CD49d-APC-A Median |
|---|---|---|---|
| POS-unpigm. | 20,84 | 1,63 | 1,33 |
| POS-pigm. | 20,89 | 2,29 | 1,92 |

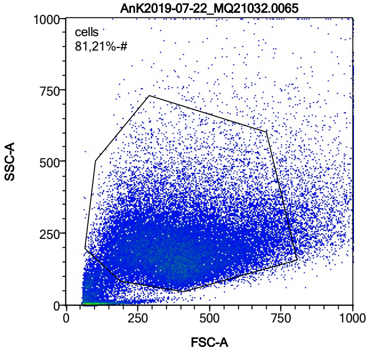


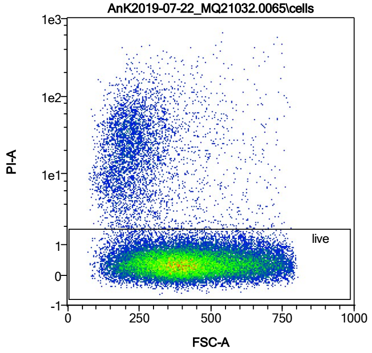


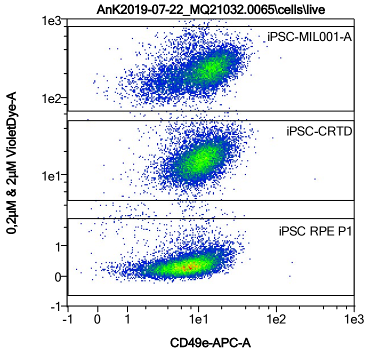


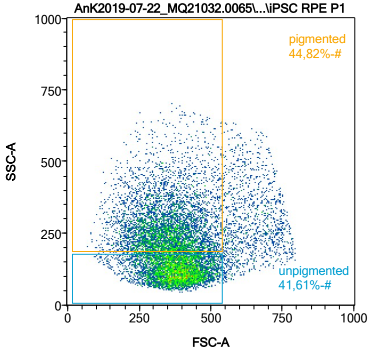


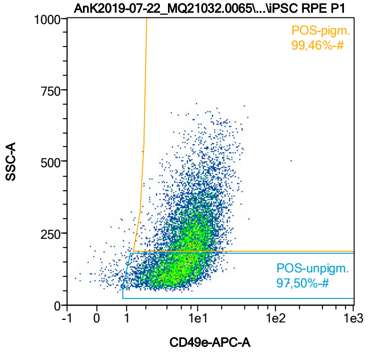


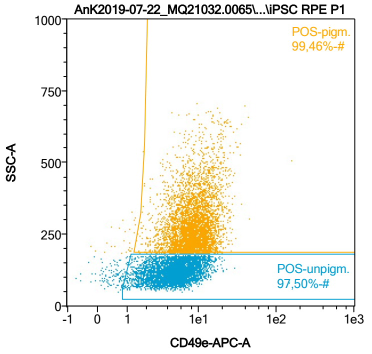


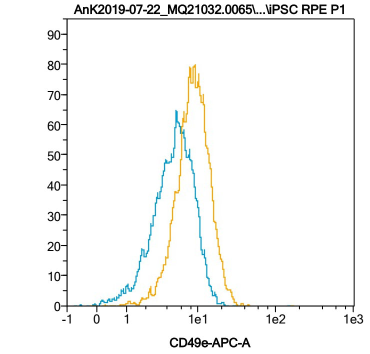


| Name | %-# | CD49e-APC-A Mean | CD49e-APC-A Median |
|---|---|---|---|
| POS-unpigm. | 97,50 | 5,64 | 5,08 |
| POS-pigm. | 99,46 | 9,50 | 8,67 |

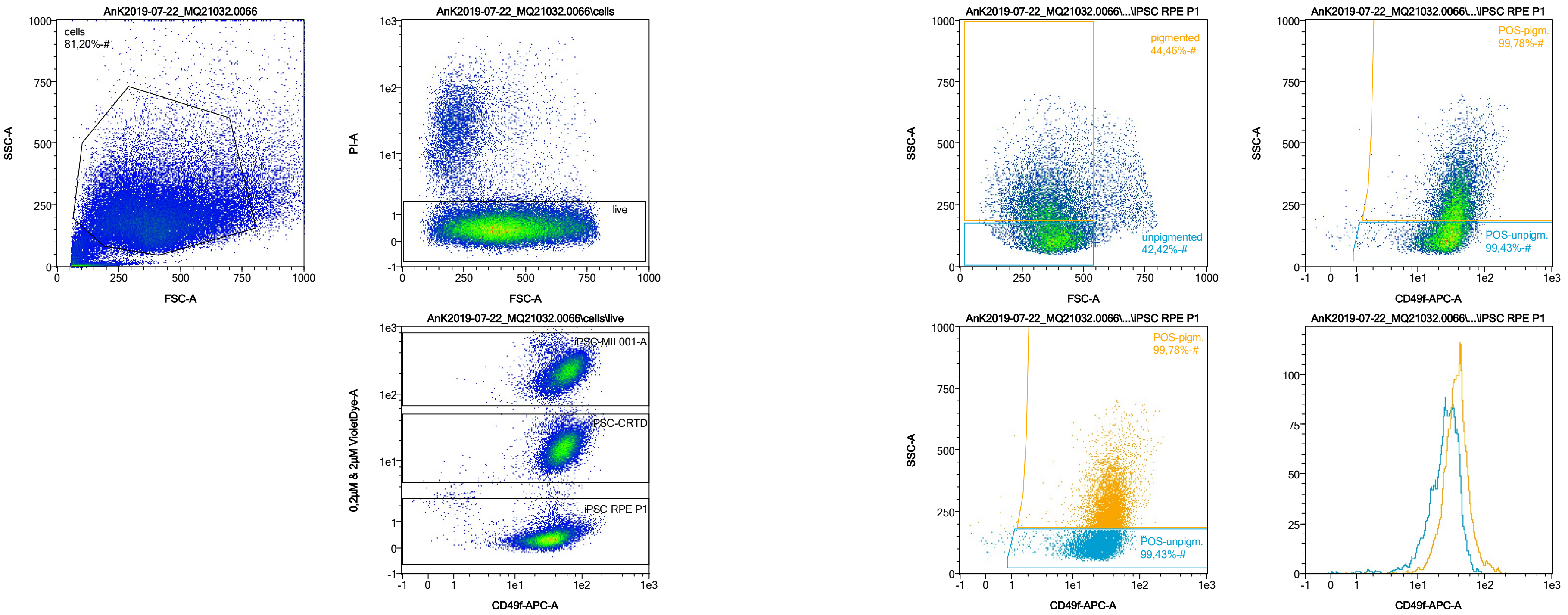


| Name | %-# | CD49f-APC-A Mean | CD49f-APC-A Median |
|---|---|---|---|
| POS-unpigm. | 99,43 | 27,05 | 26,22 |
| POS-pigm. | 99,78 | 39,64 | 36,92 |

Date:2024-06-14

Operator: Sebastian Knoebel

Version:2.13.0+54.g95abf0f4

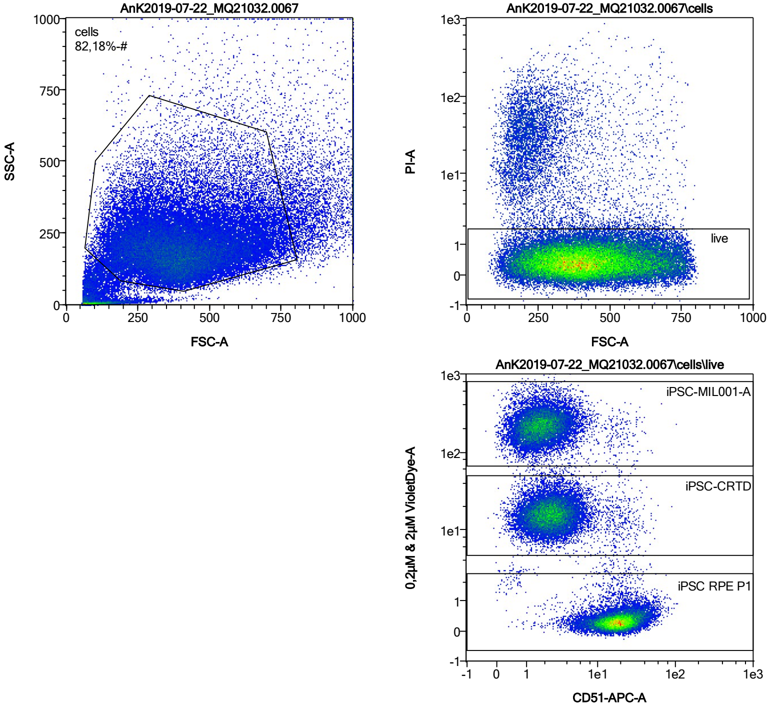


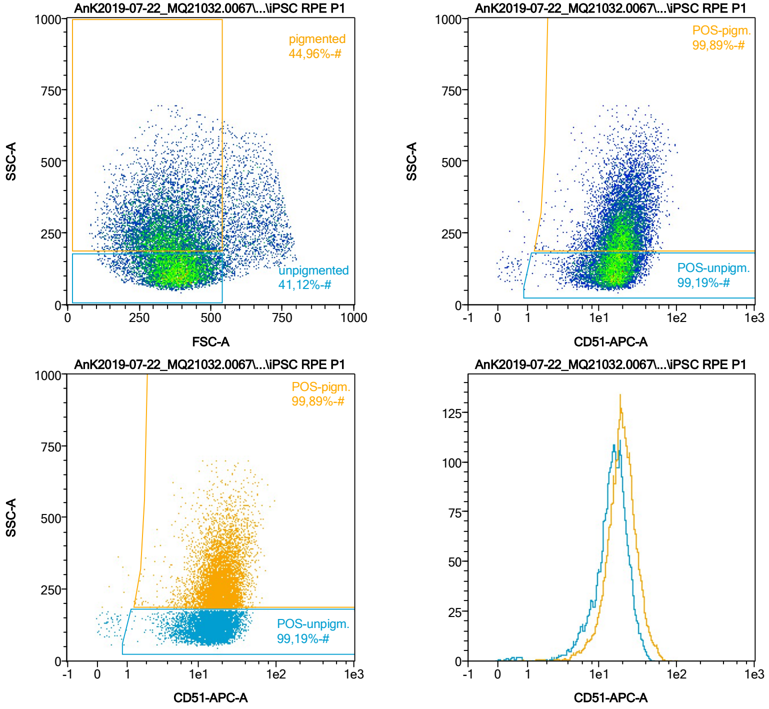


| Name | %-# | CD51-APC-A Mean | CD51-APC-A Median |
|---|---|---|---|
| POS-unpigm. | 99,19 | 15,99 | 15,32 |
| POS-pigm. | 99,89 | 20,87 | 19,43 |

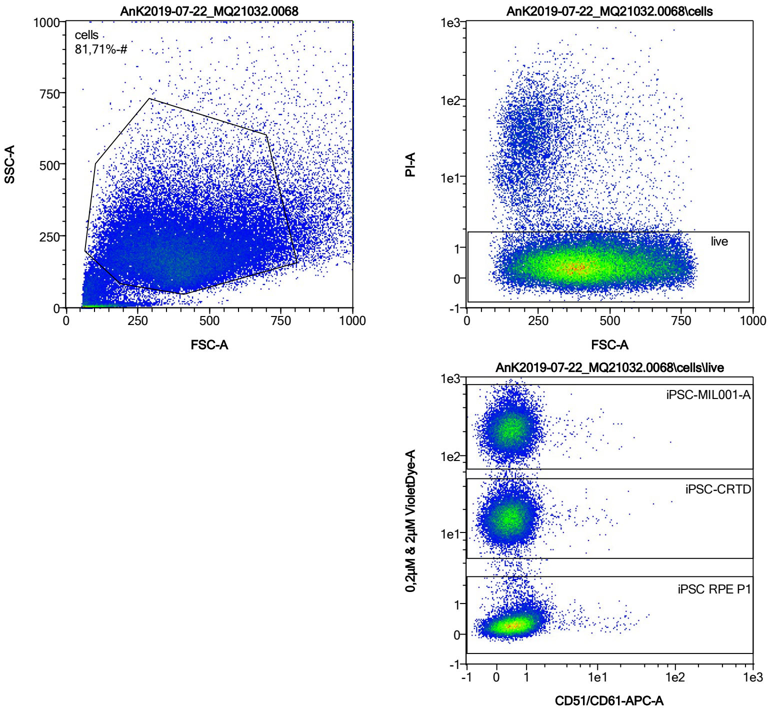


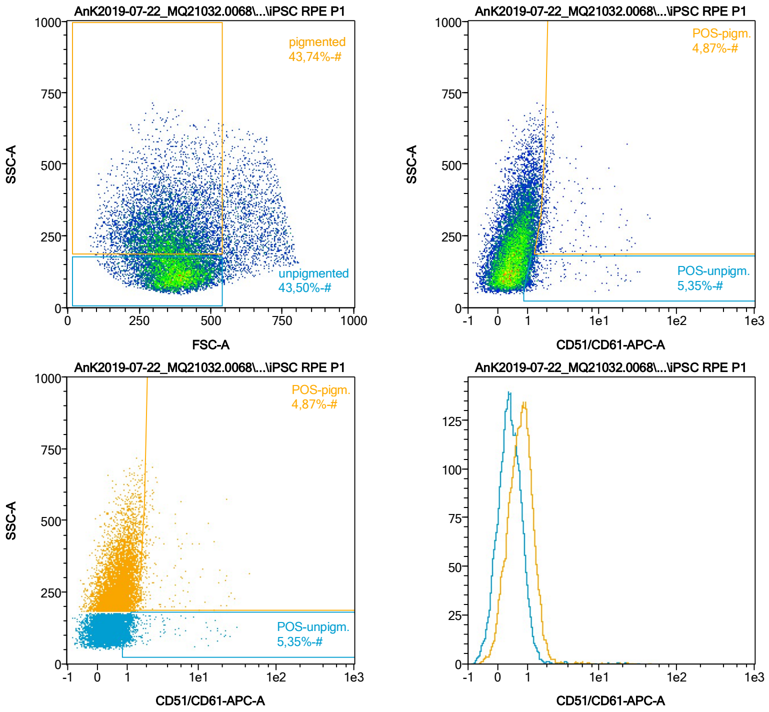


| Name | %-# | CD51/CD61-APC-A Mean | CD51/CD61-APC-A Median |
|---|---|---|---|
| POS-unpigm. | 5,35 | 2,25 | 1,18 |
| POS-pigm. | 4,87 | 3,21 | 1,85 |

Date:2024-06-14

Operator: Sebastian Knoebel

Version:2.13.0+54.g95abf0f4

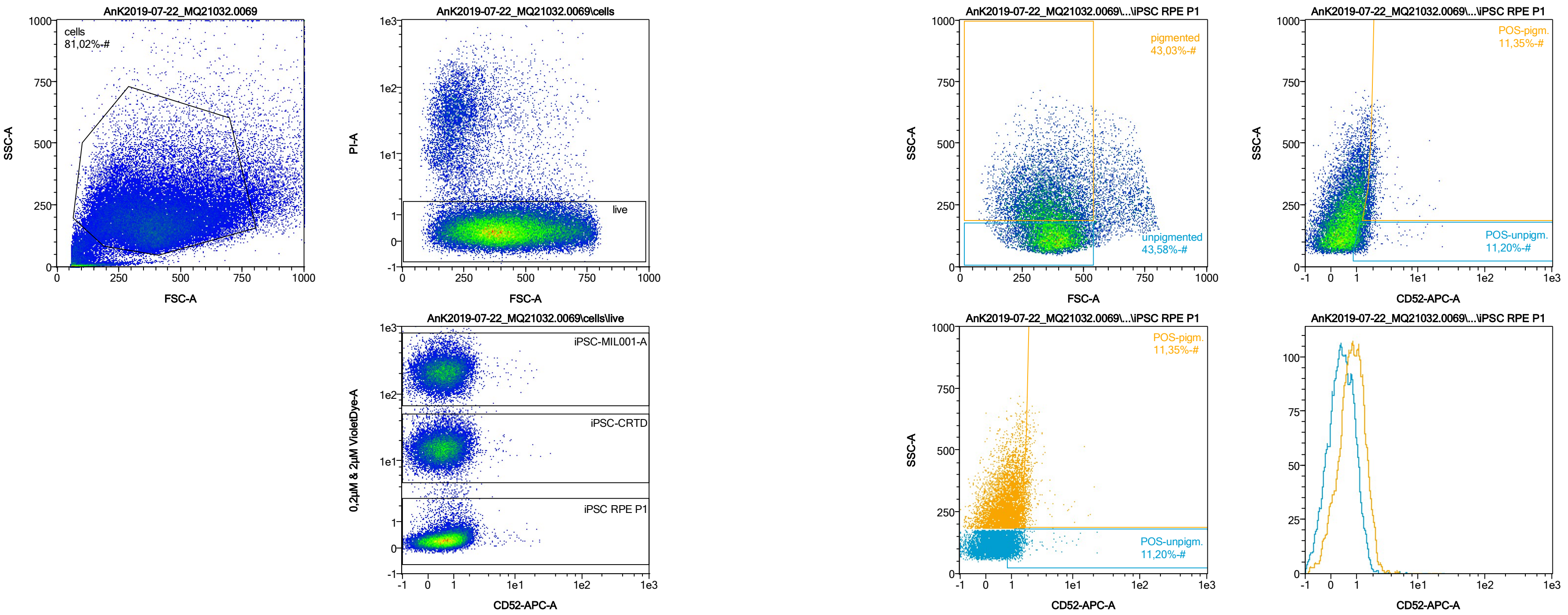


| Name | %-# | CD52-APC-A Mean | CD52-APC-A Median |
|---|---|---|---|
| POS-unpigm. | 11,20 | 1,47 | 1,23 |
| POS-pigm. | 11,35 | 2,03 | 1,82 |

Date:2024-06-14
Operator: Sebastian Knoebel
Version:2.13.0+54.g95abf0f4

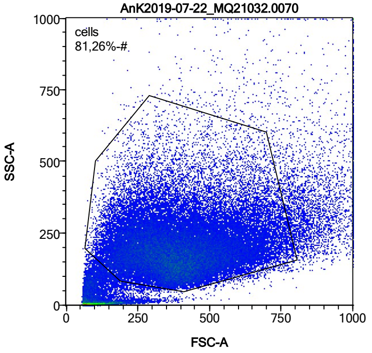


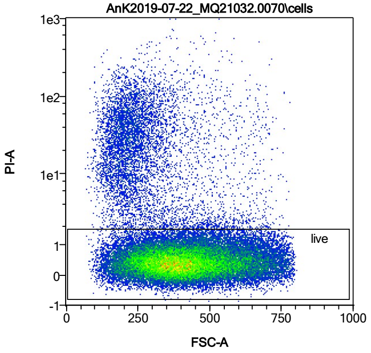


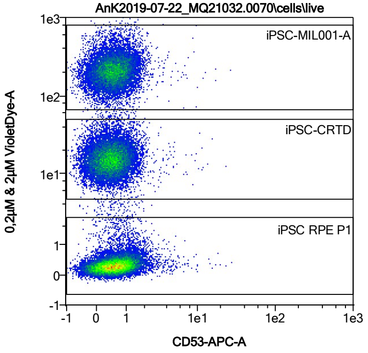


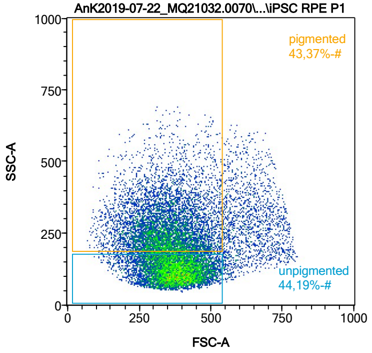


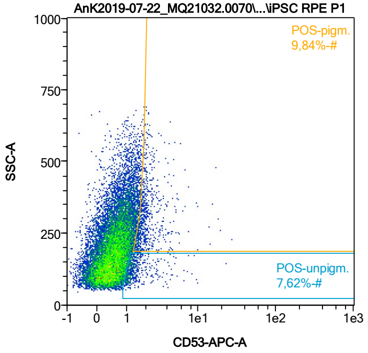


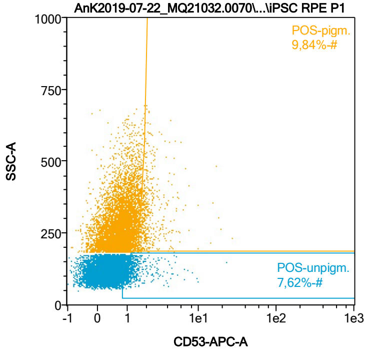


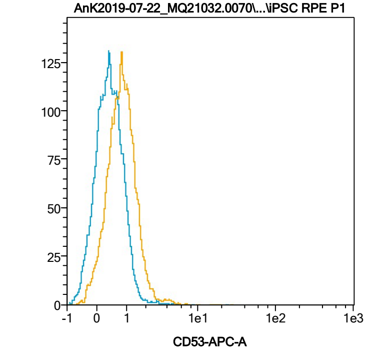


| Name | %-# | CD53-APC-A Mean | CD53-APC-A Median |
|---|---|---|---|
| POS-unpigm. | 7,62 | 1,56 | 1,22 |
| POS-pigm. | 9,84 | 2,32 | 1,88 |

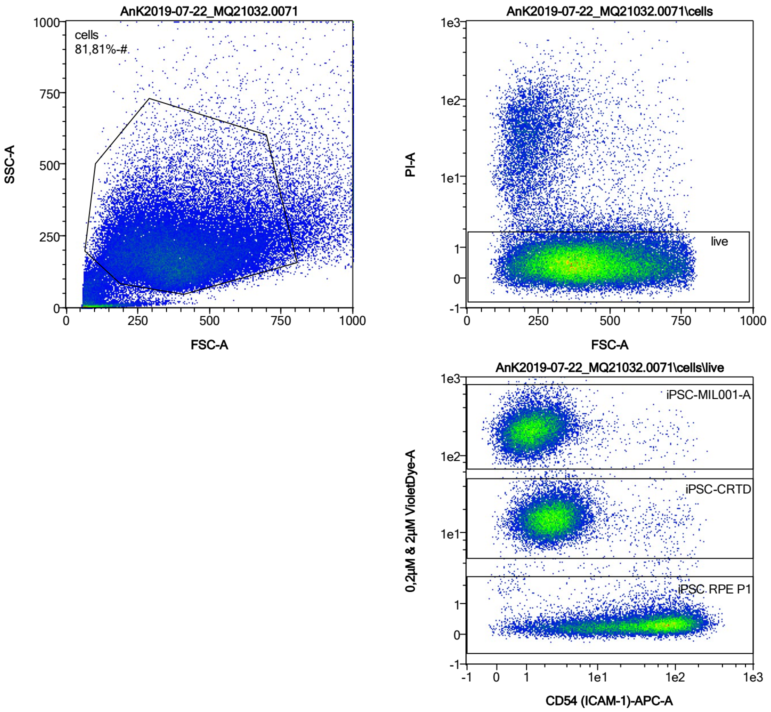


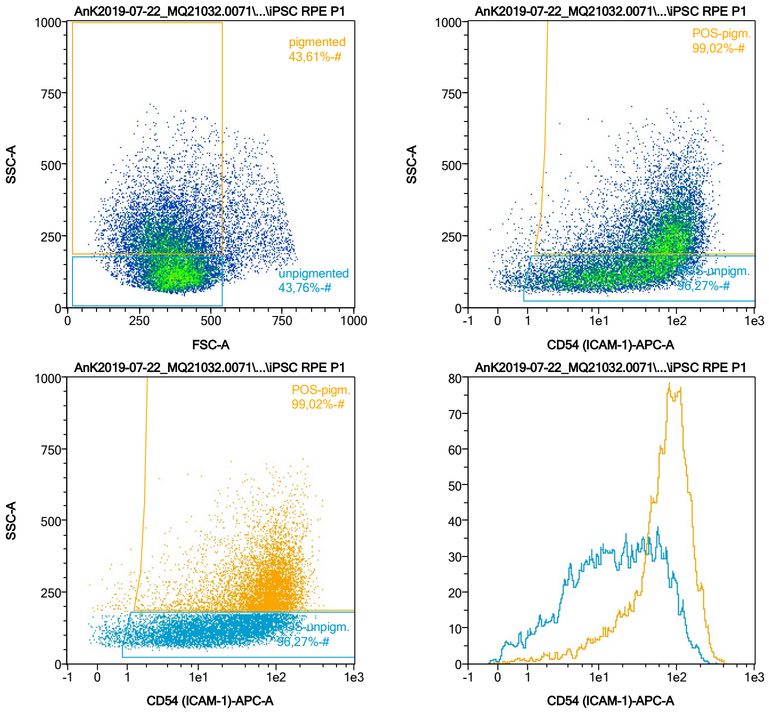


| Name | %-# | CD54 (ICAM-1)-APC-A Mean | CD54 (ICAM-1)-APC-A Median |
|---|---|---|---|
| POS-unpigm. | 96,27 | 32,02 | 18,14 |
| POS-pigm. | 99,02 | 84,34 | 76,68 |

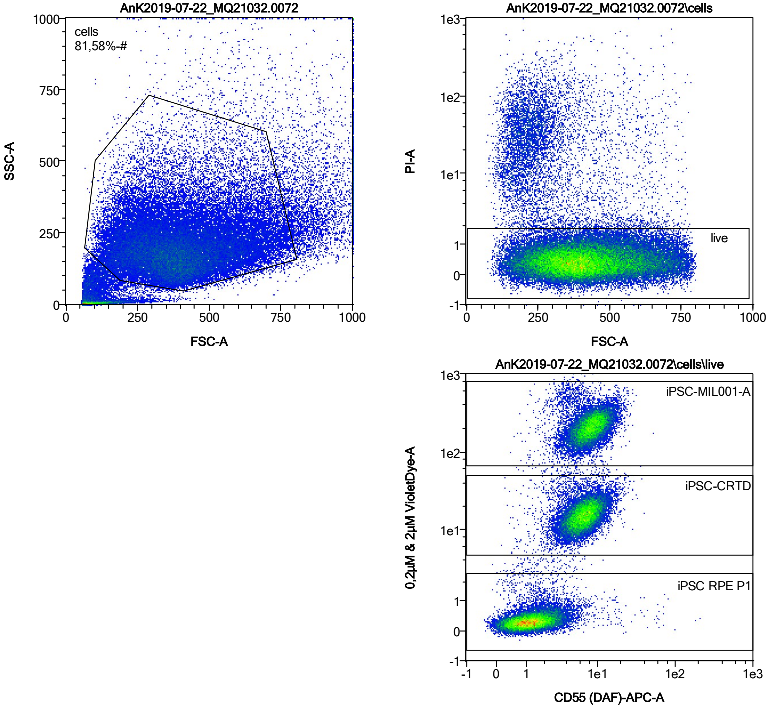


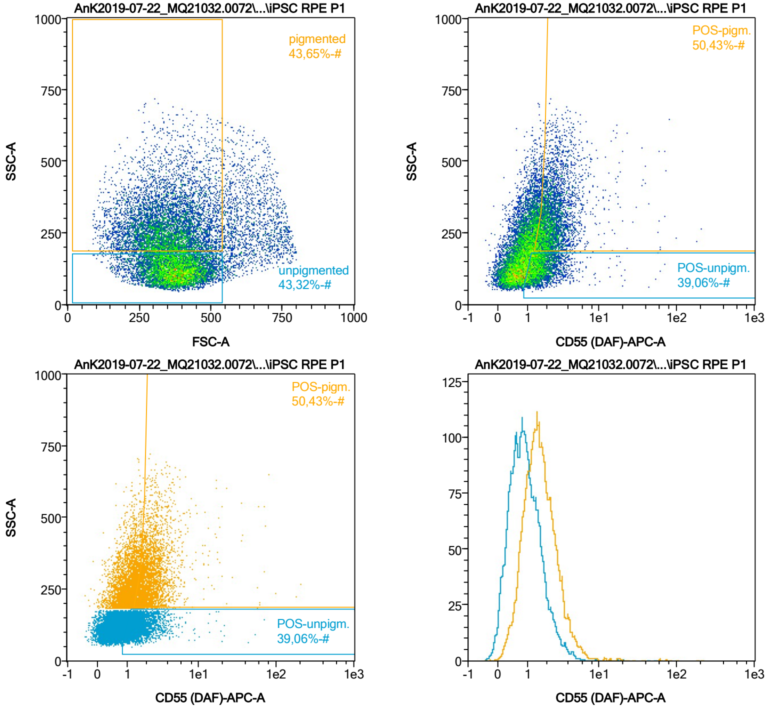


| Name | %-# | CD55 (DAF)-APC-A Mean | CD55 (DAF)-APC-A Median |
|---|---|---|---|
| POS-unpigm. | 39,06 | 1,85 | 1,49 |
| POS-pigm. | 50,43 | 2,92 | 2,18 |

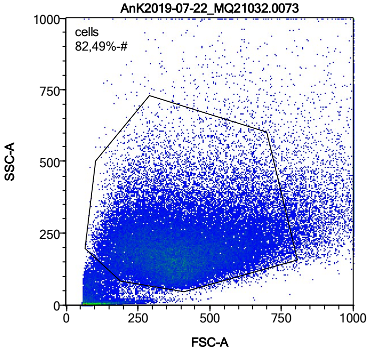


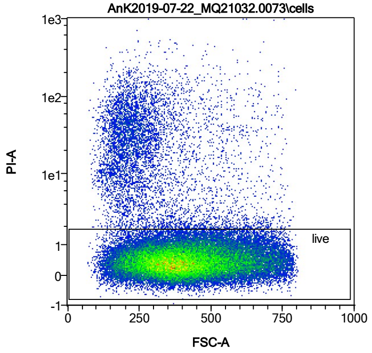


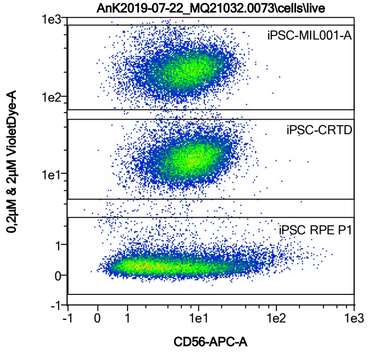


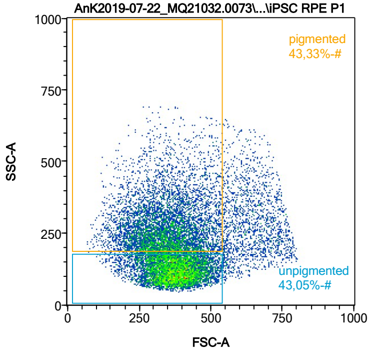


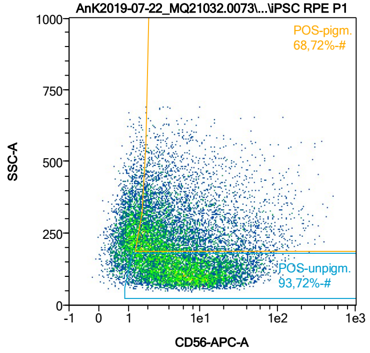


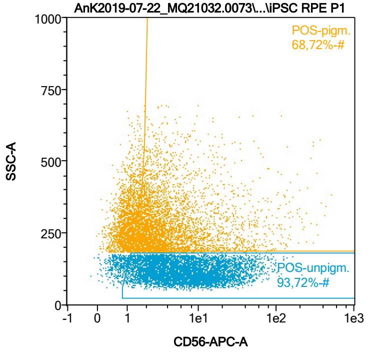


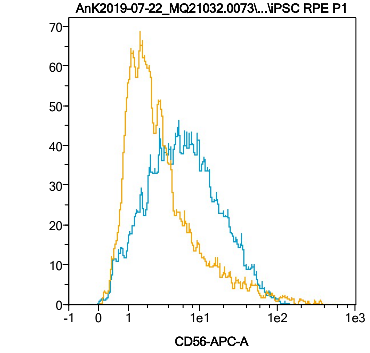


| Name | %-# | CD56-APC-A Mean | CD56-APC-A Median |
|---|---|---|---|
| POS-unpigm. | 93,72 | 11,08 | 6,44 |
| POS-pigm. | 68,72 | 10,32 | 3,34 |

Date:2024-06-14

Operator: Sebastian Knoebel

Version:2.13.0+54.g95abf0f4

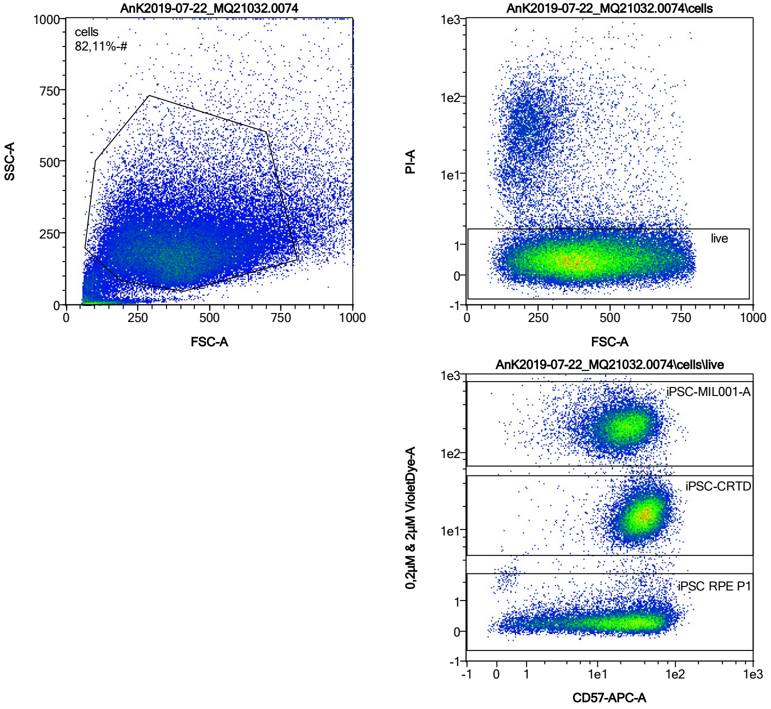


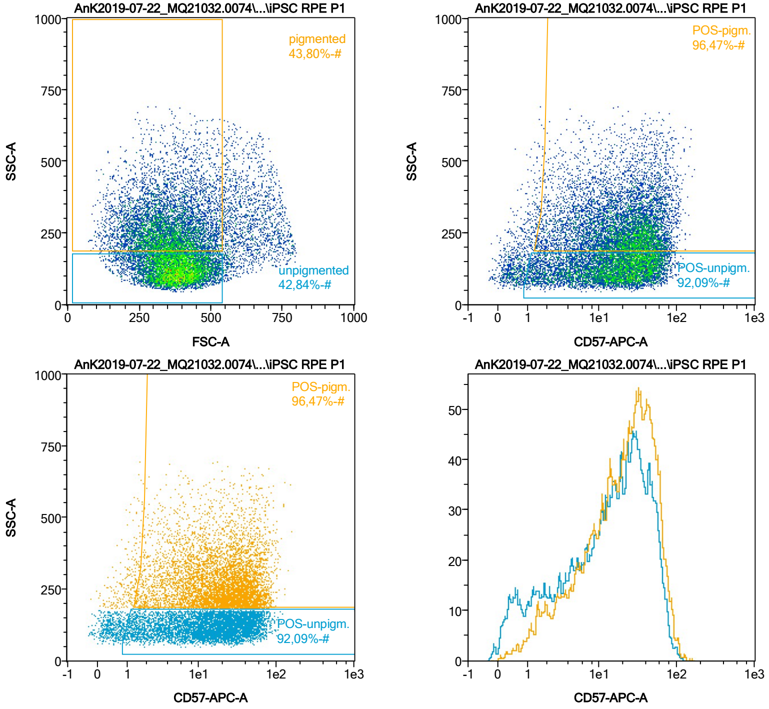


| Name | %-# | CD57-APC-A Mean | CD57-APC-A Median |
|---|---|---|---|
| POS-unpigm. | 92,09 | 22,09 | 17,03 |
| POS-pigm. | 96,47 | 26,70 | 21,99 |

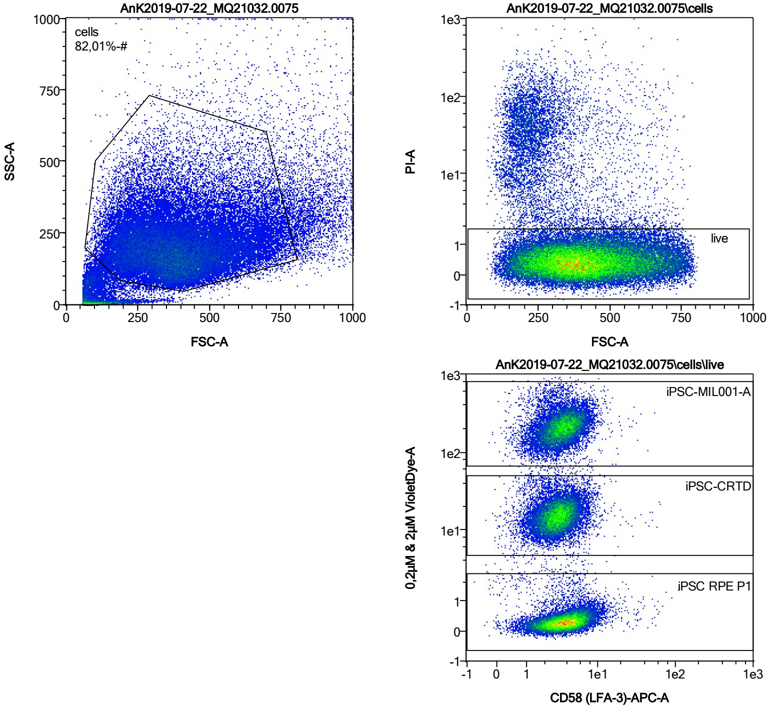


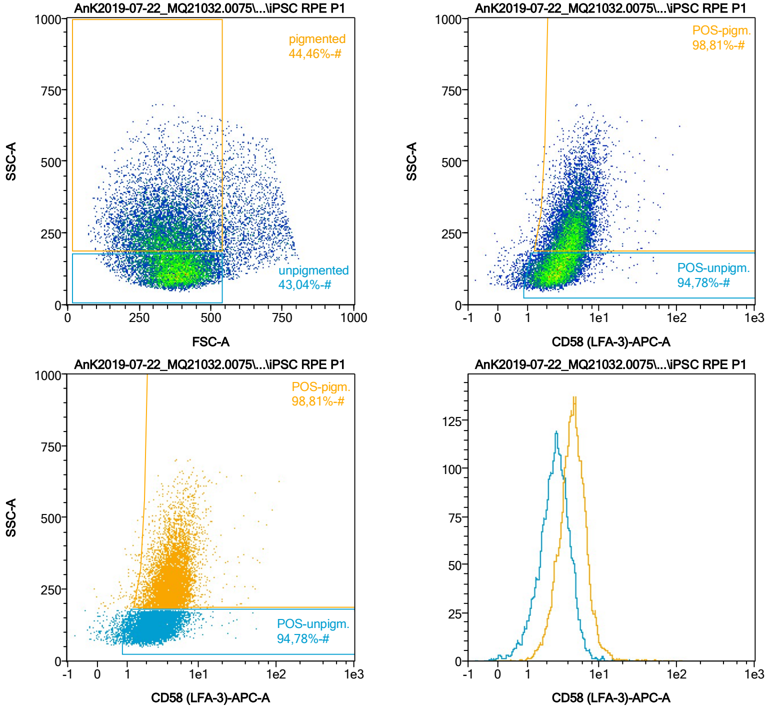


| Name | %-# | CD58 (LFA-3)-APC-A Mean | CD58 (LFA-3)-APC-A Median |
|---|---|---|---|
| POS-unpigm. | 94,78 | 2,99 | 2,72 |
| POS-pigm. | 98,81 | 4,98 | 4,53 |

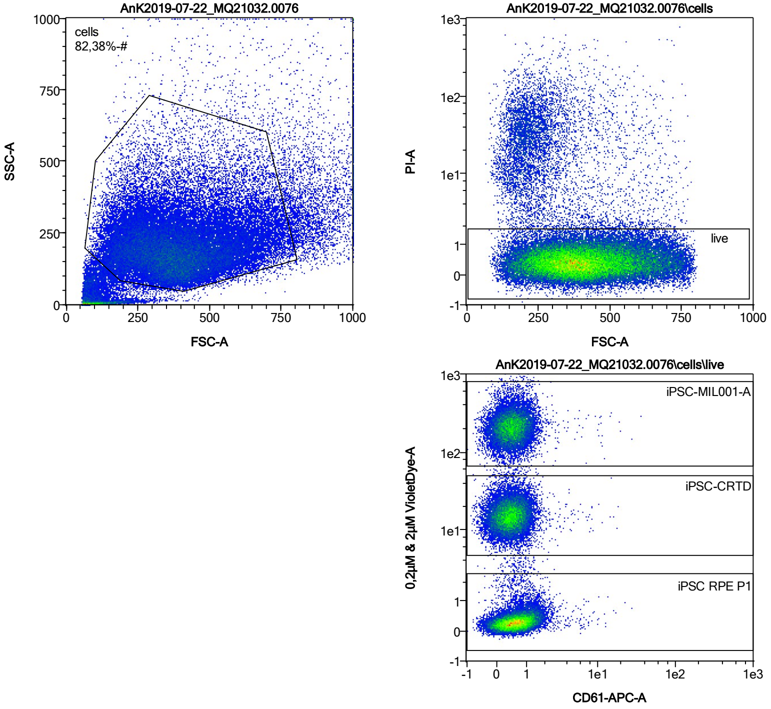


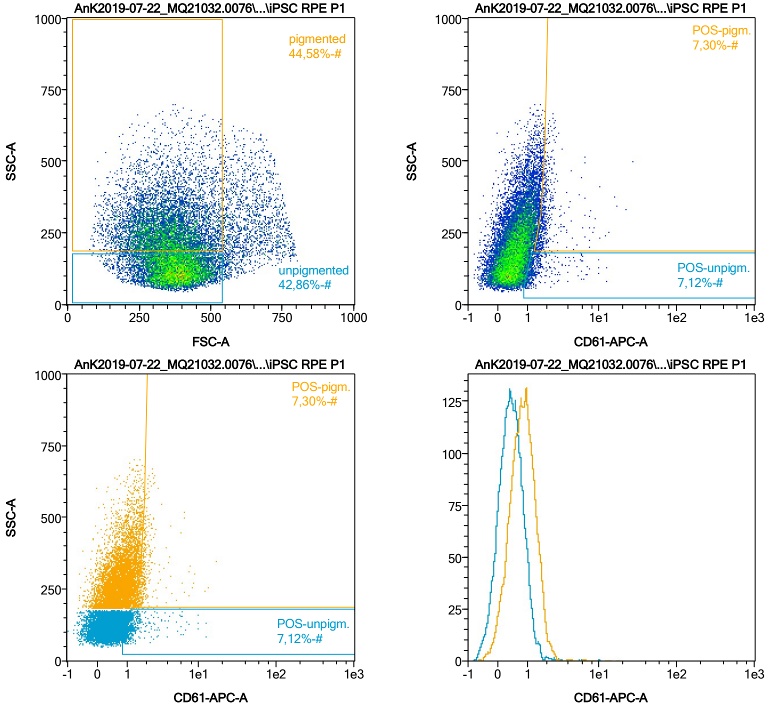


| Name | %-# | CD61-APC-A Mean | CD61-APC-A Median |
|---|---|---|---|
| POS-unpigm. | 7,12 | 1,56 | 1,19 |
| POS-pigm. | 7,30 | 2,08 | 1,78 |

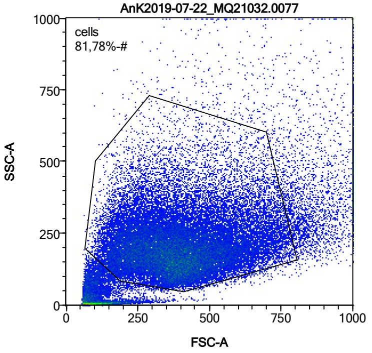


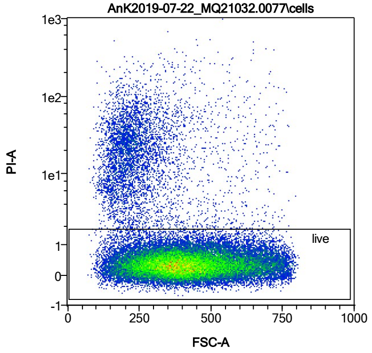


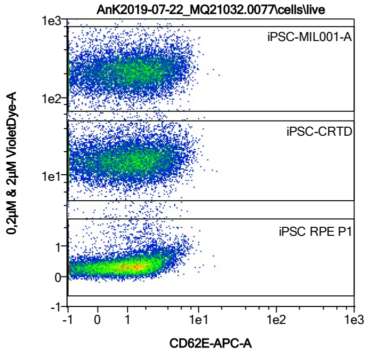


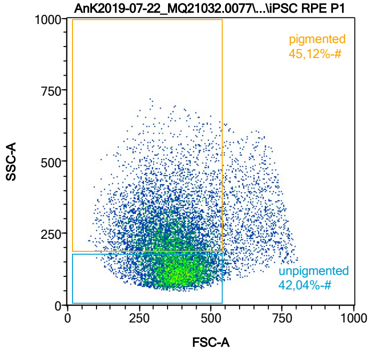


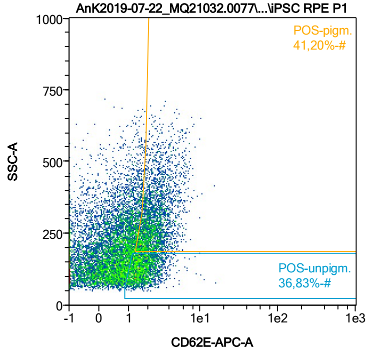


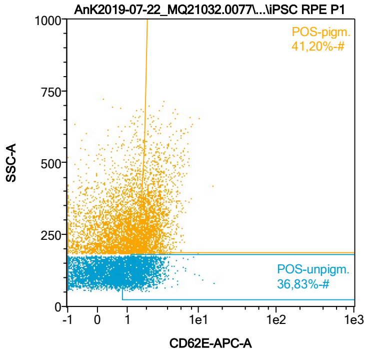


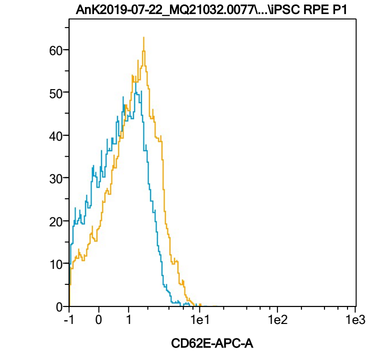


| Name | %-# | CD62E-APC-A Mean | CD62E-APC-A Median |
|---|---|---|---|
| POS-unpigm. | 36,83 | 1,82 | 1,62 |
| POS-pigm. | 41,20 | 2,64 | 2,36 |

Date:2024-06-14

Operator: Sebastian Knoebel

Version:2.13.0+54.g95abf0f4

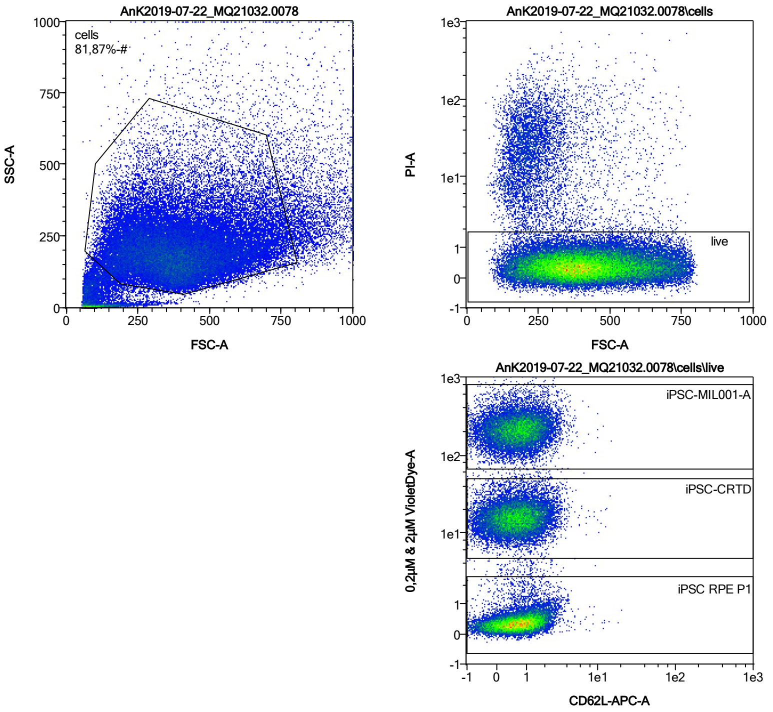


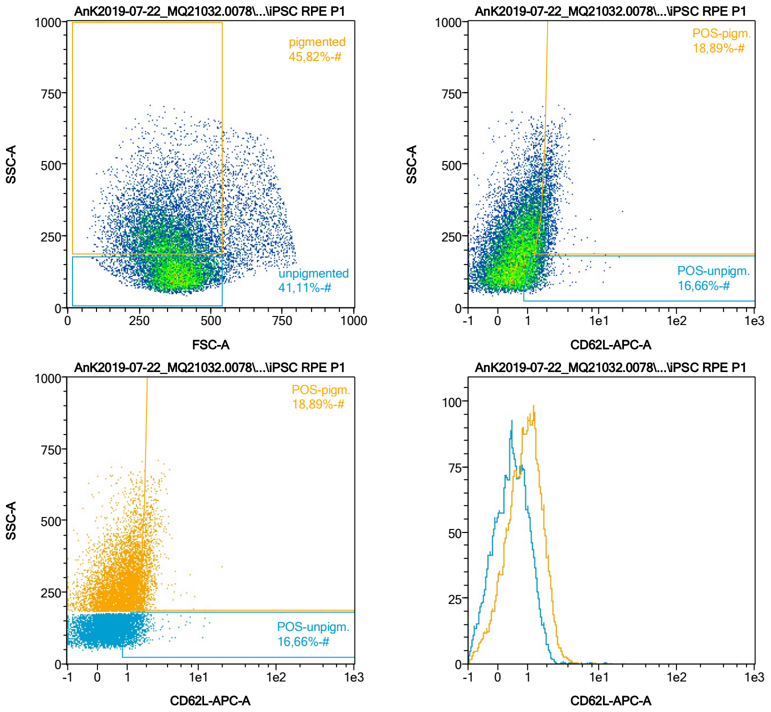


| Name | %-# | CD62L-APC-A Mean | CD62L-APC-A Median |
|---|---|---|---|
| POS-unpigm. | 16,66 | 1,47 | 1,29 |
| POS-pigm. | 18,89 | 2,07 | 1,90 |

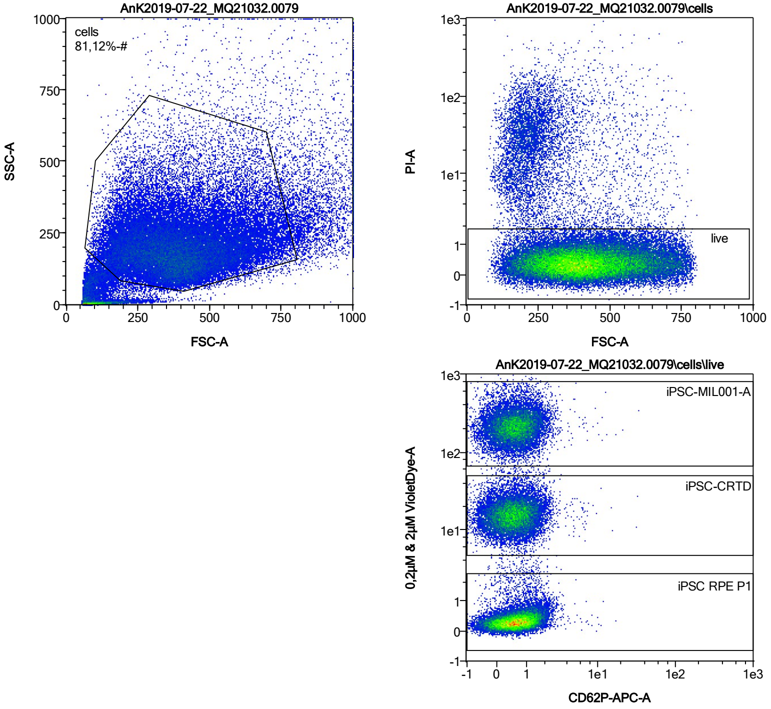


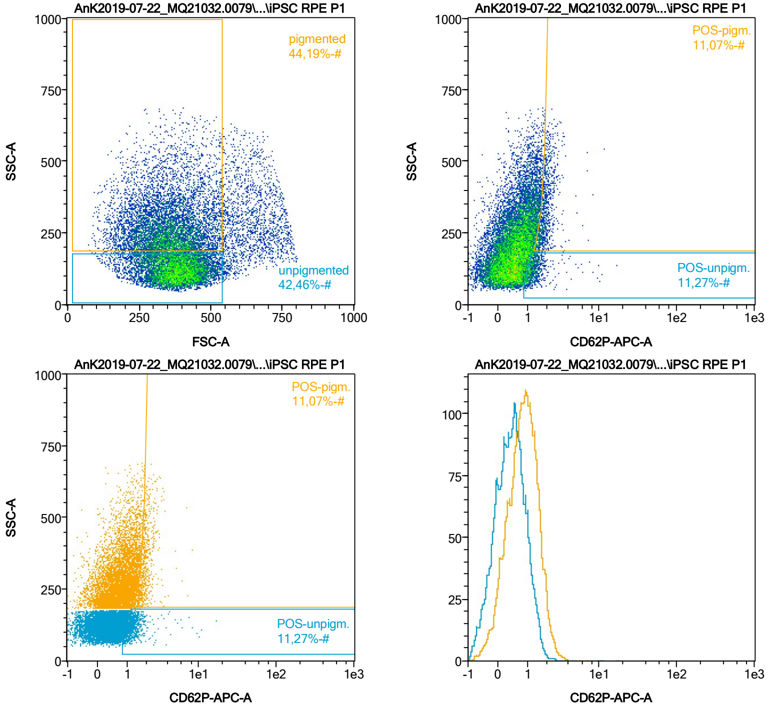


| Name | %-# | CD62P-APC-A Mean | CD62P-APC-A Median |
|---|---|---|---|
| POS-unpigm. | 11,27 | 1,41 | 1,22 |
| POS-pigm. | 11,07 | 1,95 | 1,86 |

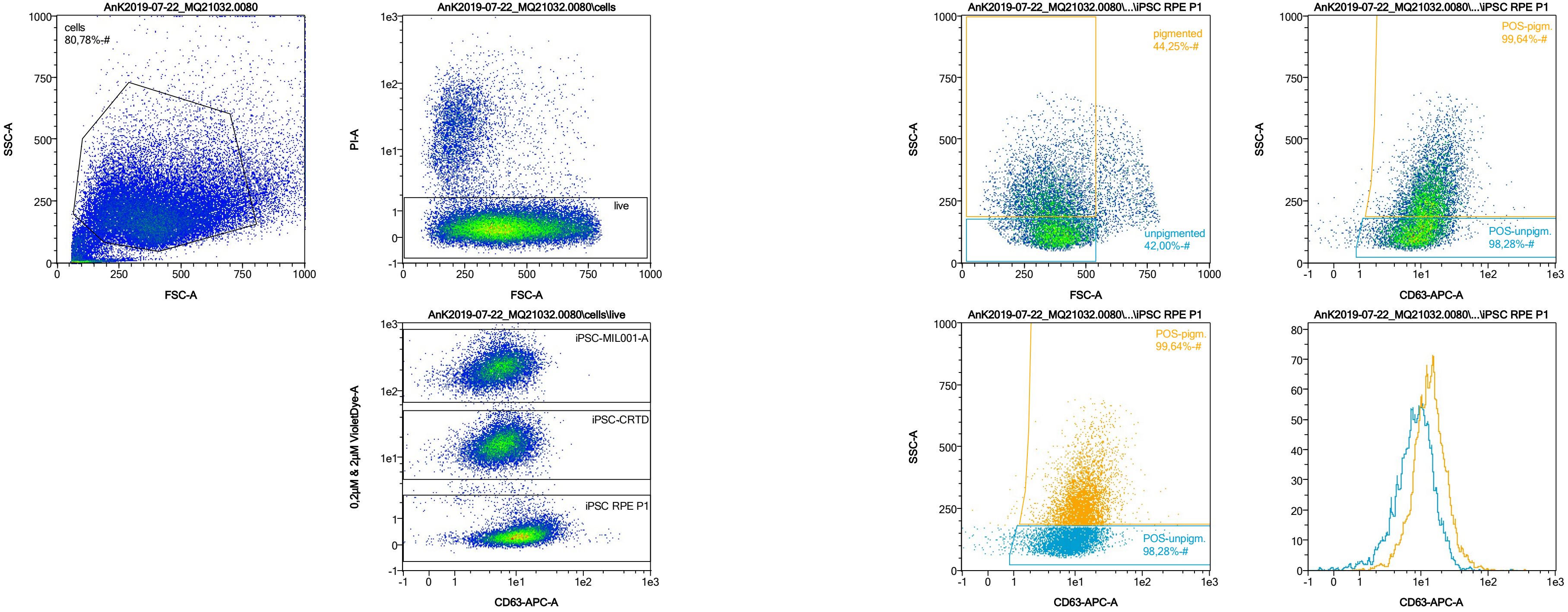


| Name | %-# | CD63-APC-A Mean | CD63-APC-A Median |
|---|---|---|---|
| POS-unpigm. | 98,28 | 10,02 | 8,48 |
| POS-pigm. | 99,64 | 15,58 | 13,04 |

Date:2024-06-14

Operator: Sebastian Knoebel

Version:2.13.0+54.g95abf0f4

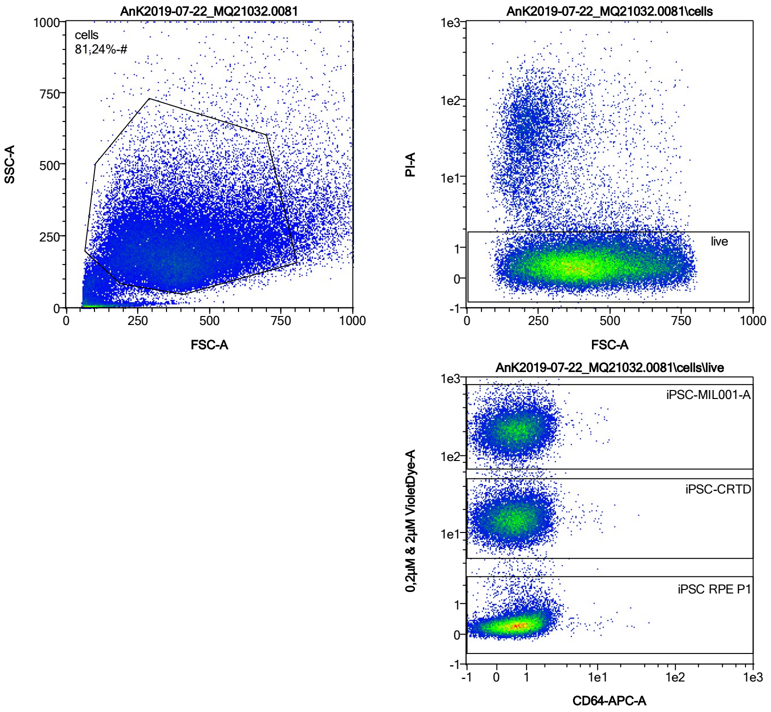


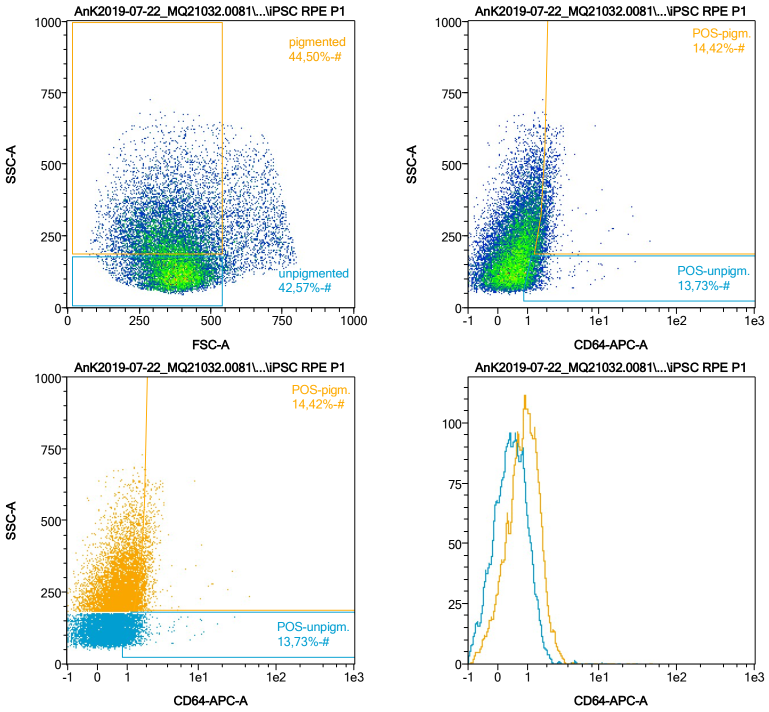


| Name | %-# | CD64-APC-A Mean | CD64-APC-A Median |
|---|---|---|---|
| POS-unpigm. | 13,73 | 1,49 | 1,25 |
| POS-pigm. | 14,42 | 2,09 | 1,83 |

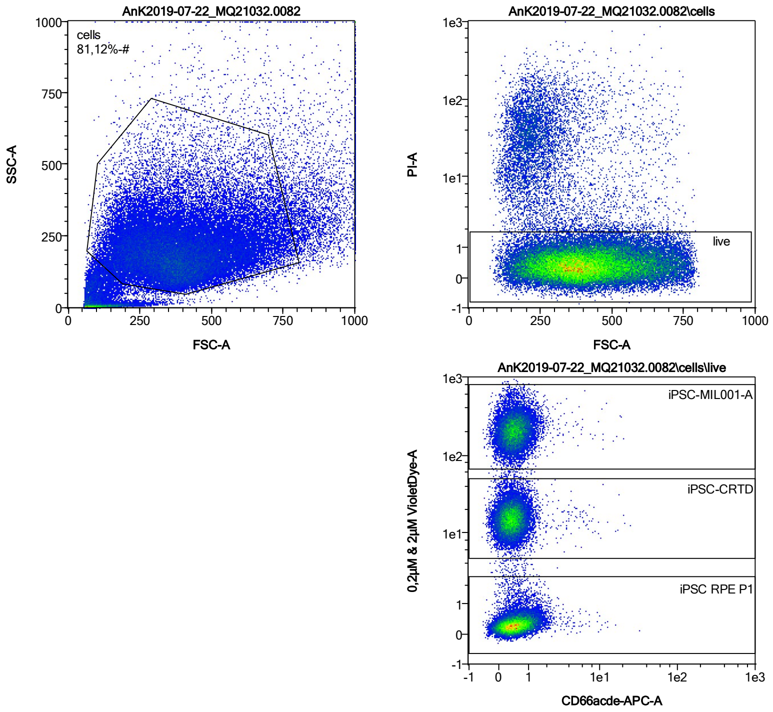


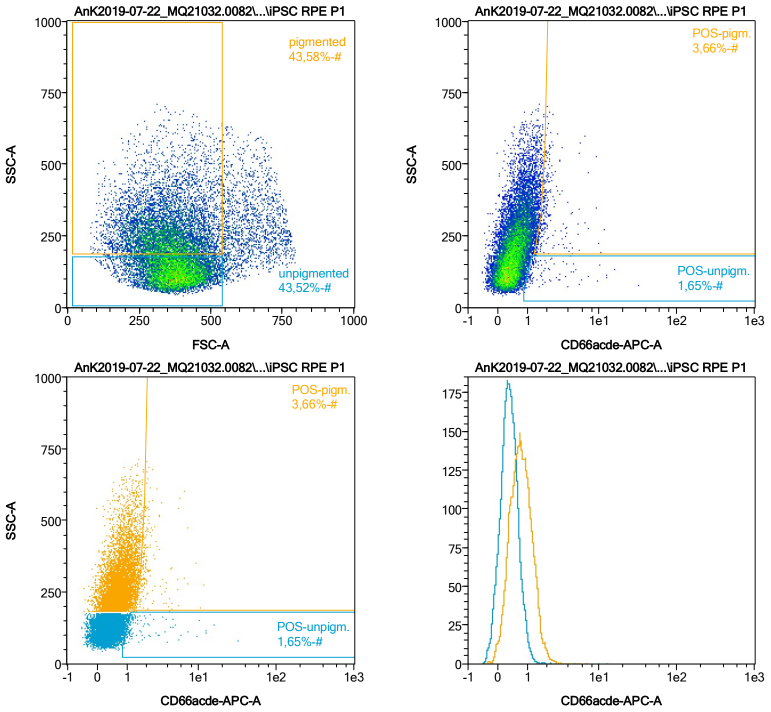


| Name | %-# | CD66acde-APC-A Mean | CD66acde-APC-A Median |
|---|---|---|---|
| POS-unpigm. | 1,65 | 2,42 | 1,24 |
| POS-pigm. | 3,66 | 2,24 | 1,86 |

Date:2024-06-14
Operator: Sebastian Knoebel
Version:2.13.0+54.g95abf0f4

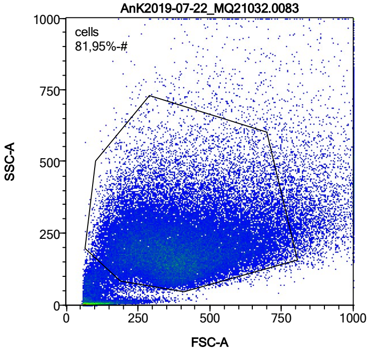


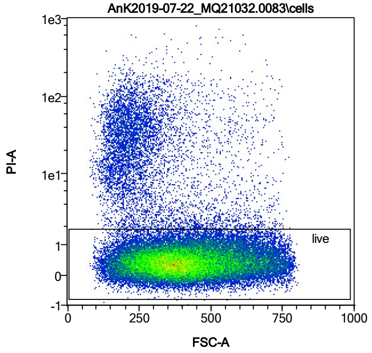


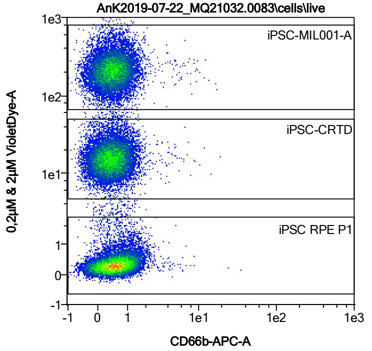


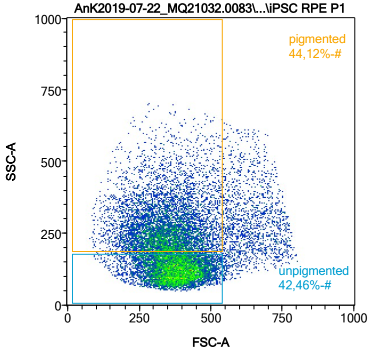


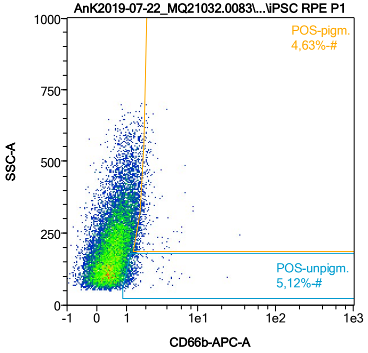


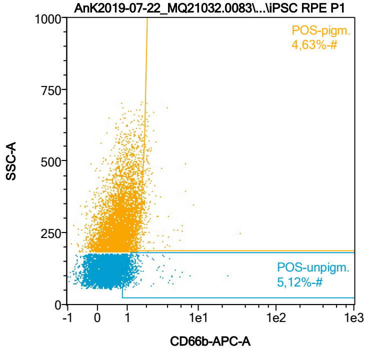


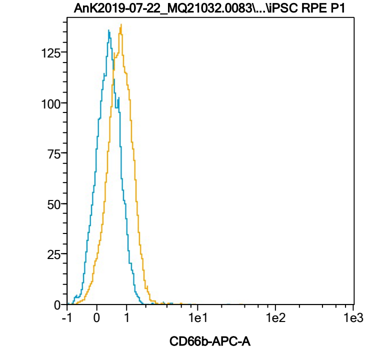


| Name | %-# | CD66b-APC-A Mean | CD66b-APC-A Median |
|---|---|---|---|
| POS-unpigm. | 5,12 | 1,45 | 1,17 |
| POS-pigm. | 4,63 | 2,10 | 1,80 |

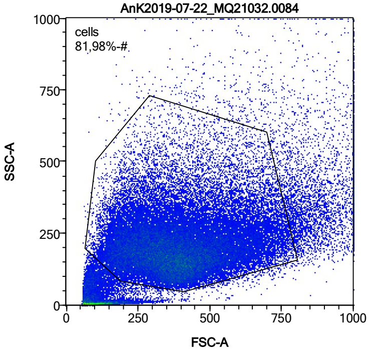


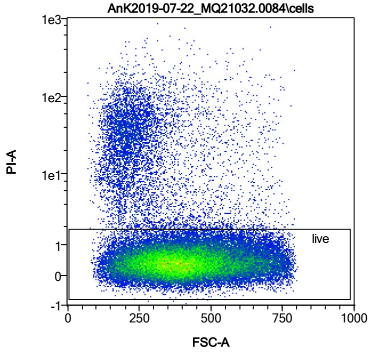


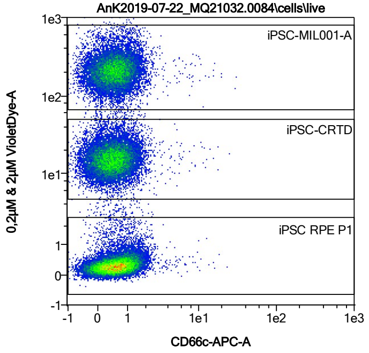


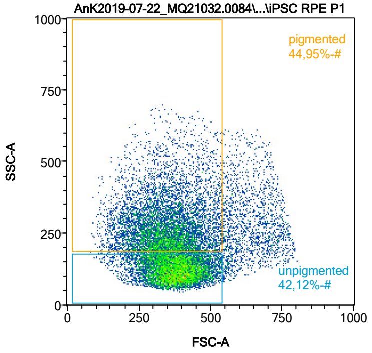


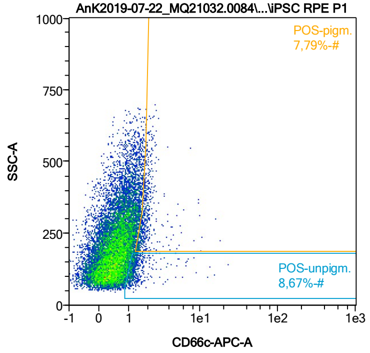


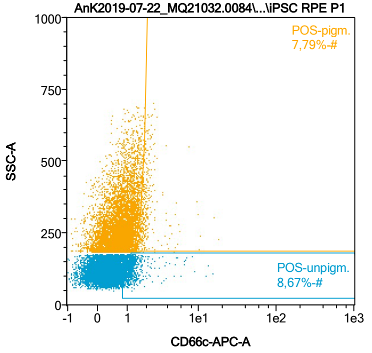


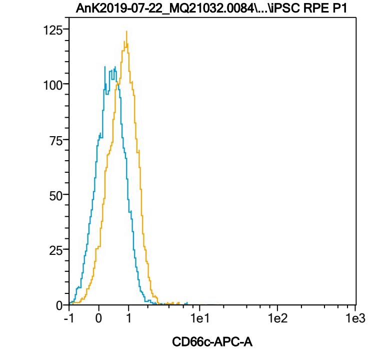


| Name | %-# | CD66c-APC-A Mean | CD66c-APC-A Median |
|---|---|---|---|
| POS-unpigm. | 8,67 | 1,47 | 1,20 |
| POS-pigm. | 7,79 | 2,04 | 1,80 |

Date:2024-06-14

Operator: Sebastian Knoebel

Version:2.13.0+54.g95abf0f4

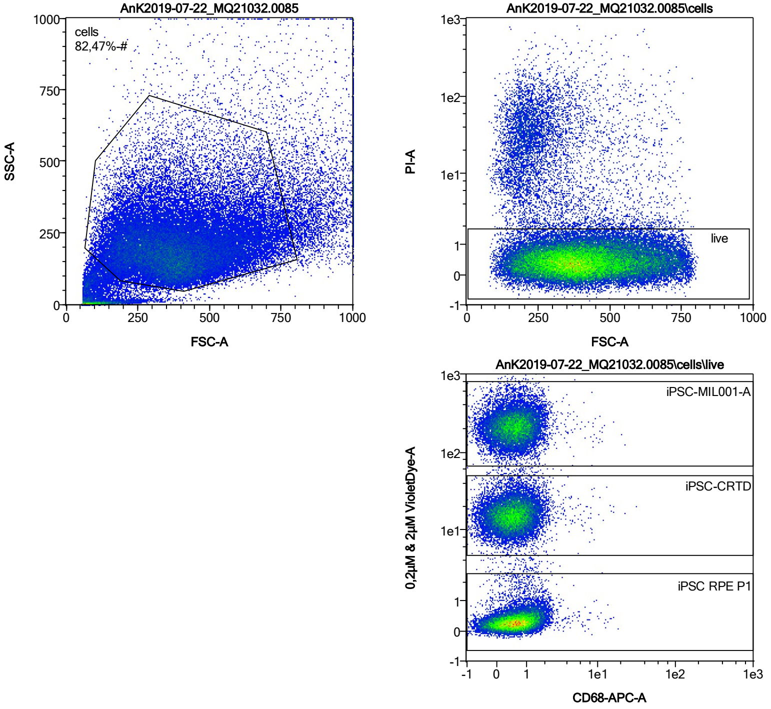


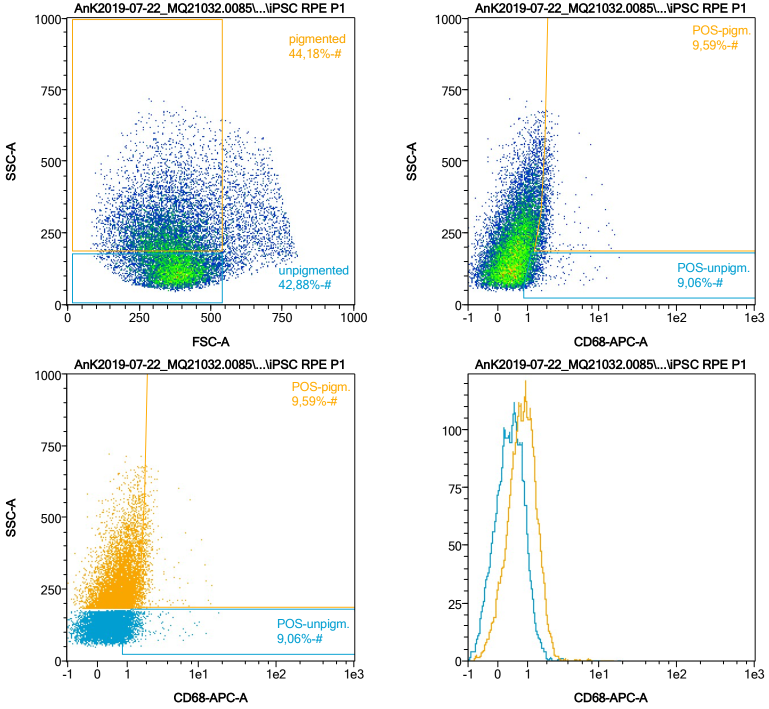


| Name | %-# | CD68-APC-A Mean | CD68-APC-A Median |
|---|---|---|---|
| POS-unpigm. | 9,06 | 1,50 | 1,22 |
| POS-pigm. | 9,59 | 2,09 | 1,83 |

Date:2024-06-14

Operator: Sebastian Knoebel

Version:2.13.0+54.g95abf0f4

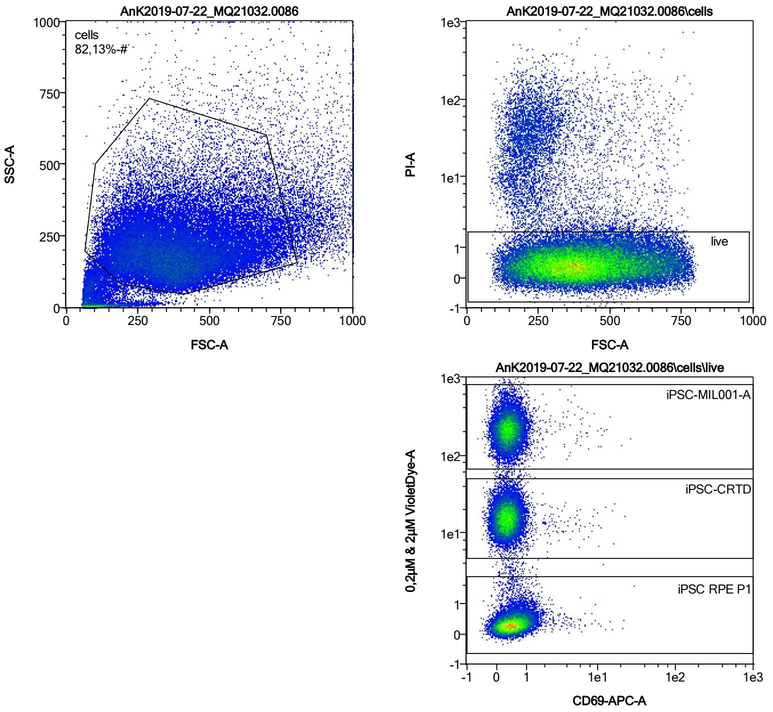


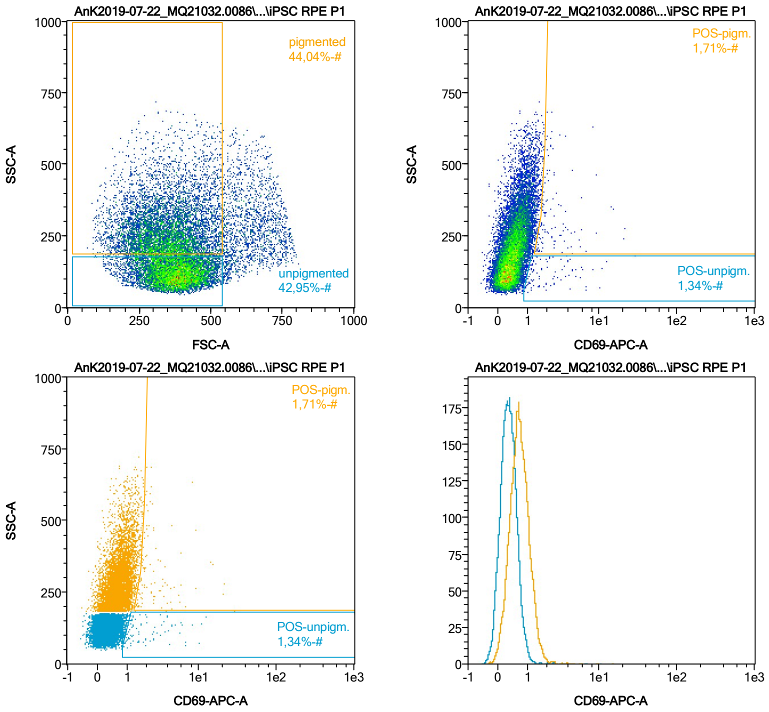


| Name | %-# | CD69-APC-A Mean | CD69-APC-A Median |
|---|---|---|---|
| POS-unpigm. | 1,34 | 2,45 | 1,50 |
| POS-pigm. | 1,71 | 3,36 | 2,03 |

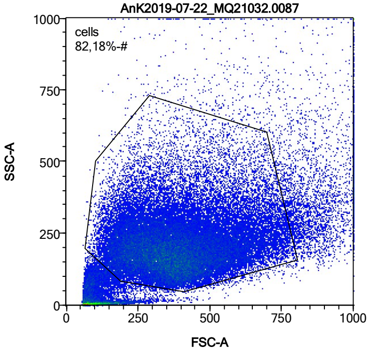


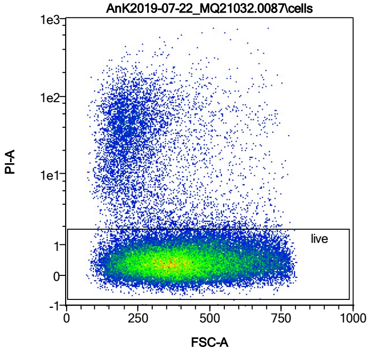


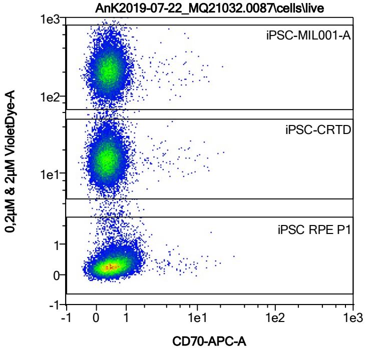


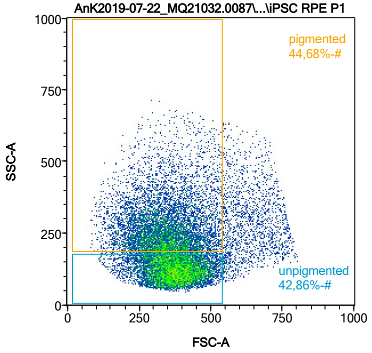


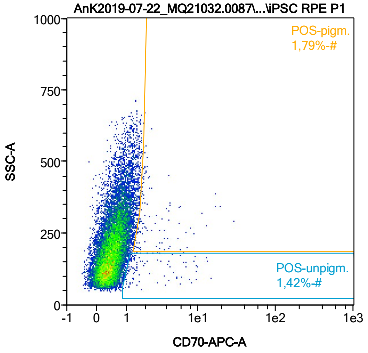


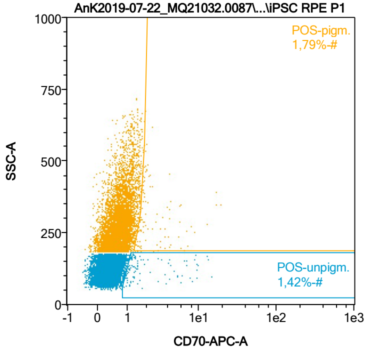


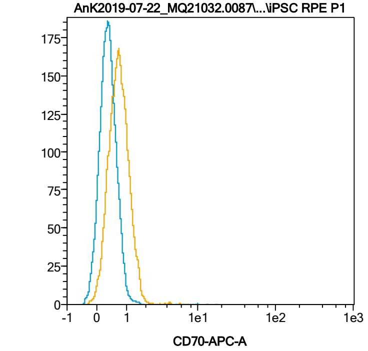


| Name | %-# | CD70-APC-A Mean | CD70-APC-A Median |
|---|---|---|---|
| POS-unpigm. | 1,42 | 2,97 | 1,37 |
| POS-pigm. | 1,79 | 2,95 | 1,89 |

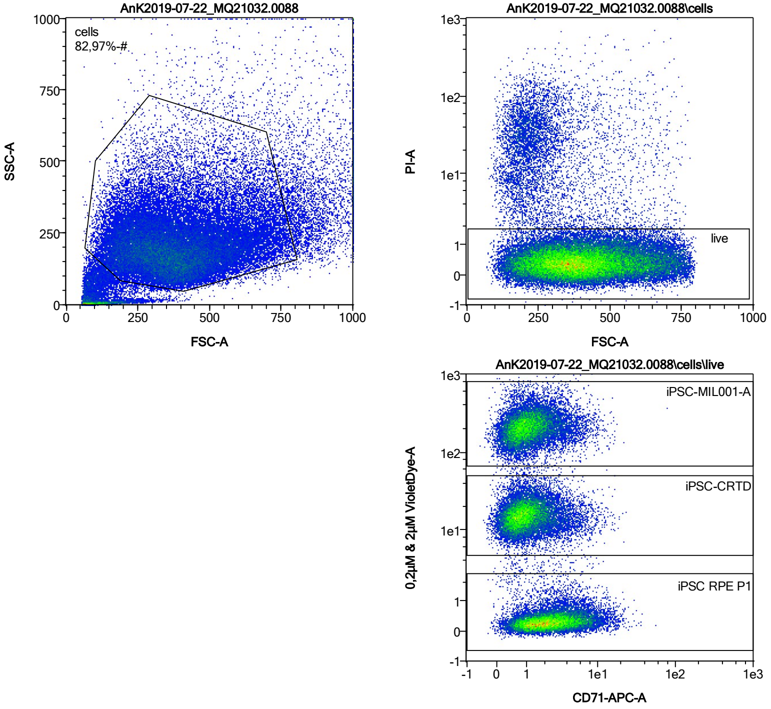


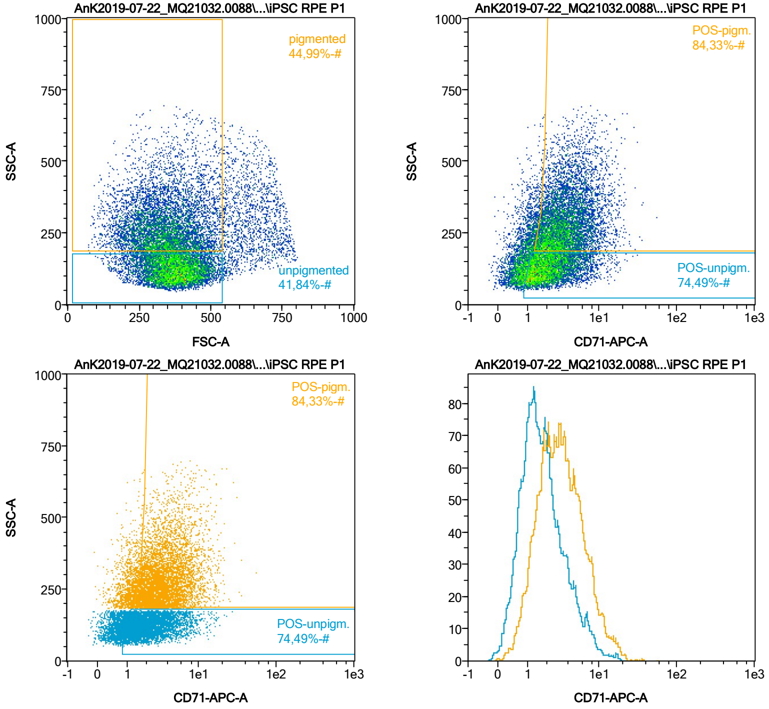


| Name | %-# | CD71-APC-A Mean | CD71-APC-A Median |
|---|---|---|---|
| POS-unpigm. | 74,49 | 2,66 | 1,99 |
| POS-pigm. | 84,33 | 4,34 | 3,40 |

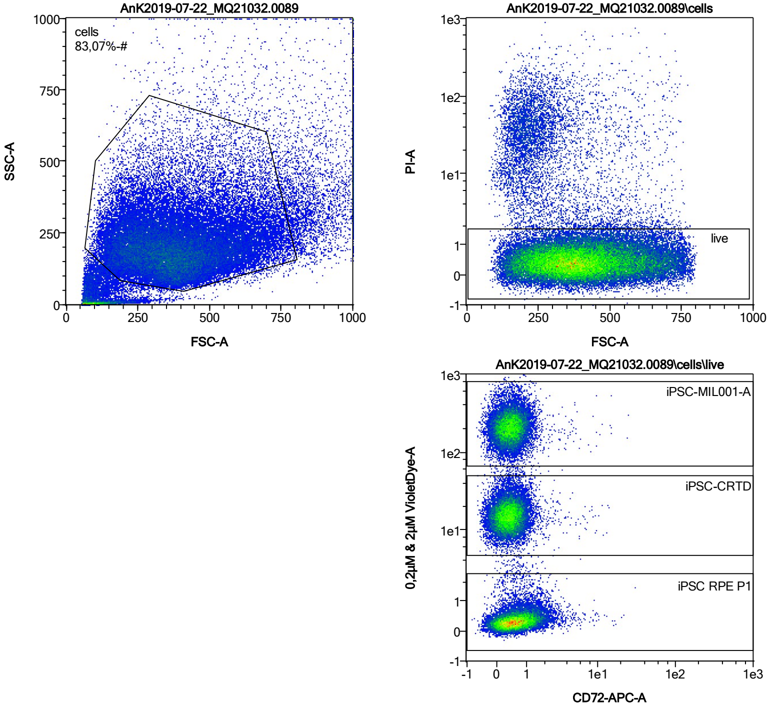


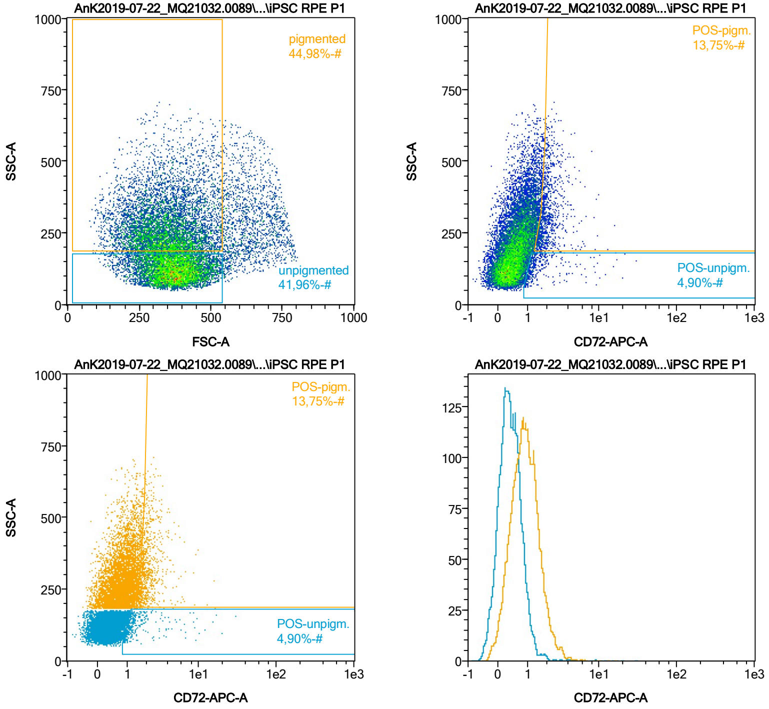


| Name | %-# | CD72-APC-A Mean | CD72-APC-A Median |
|---|---|---|---|
| POS-unpigm. | 4,90 | 1,99 | 1,24 |
| POS-pigm. | 13,75 | 2,20 | 1,93 |

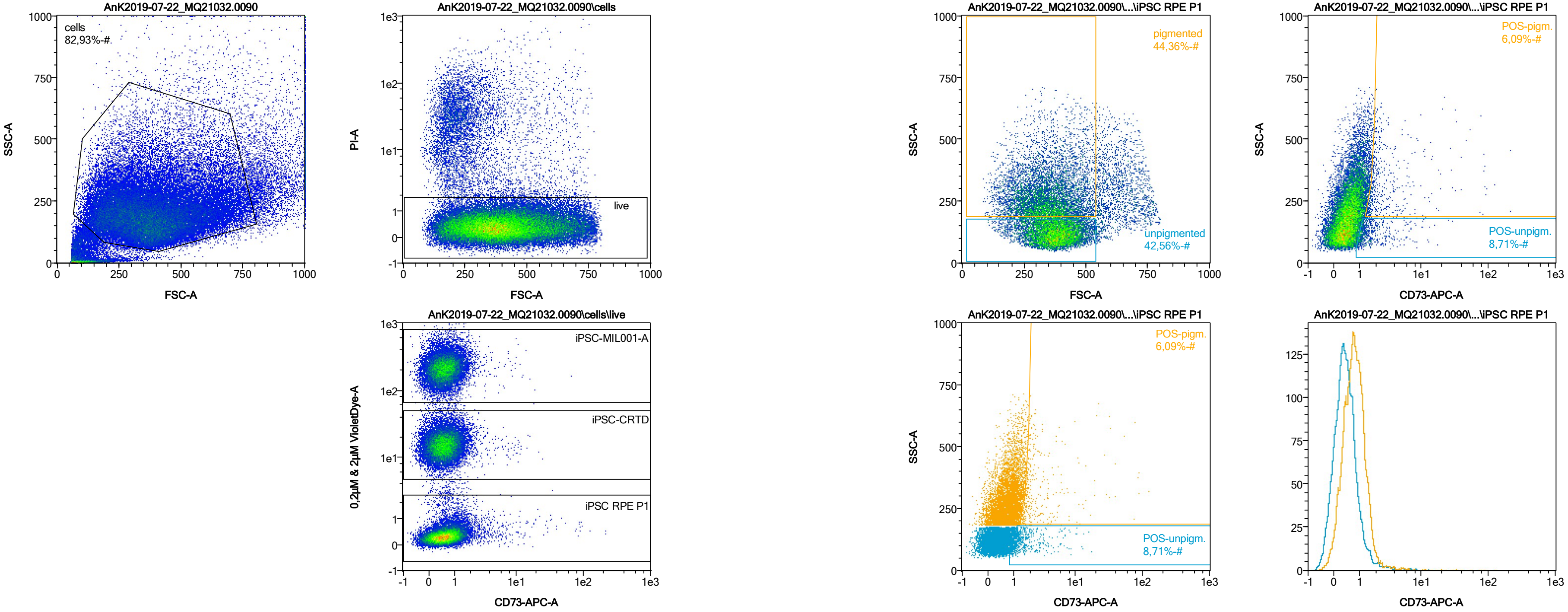


| Name | %-# | CD73-APC-A Mean | CD73-APC-A Median |
|---|---|---|---|
| POS-unpigm. | 8,71 | 1,90 | 1,31 |
| POS-pigm. | 6,09 | 4,74 | 1,95 |

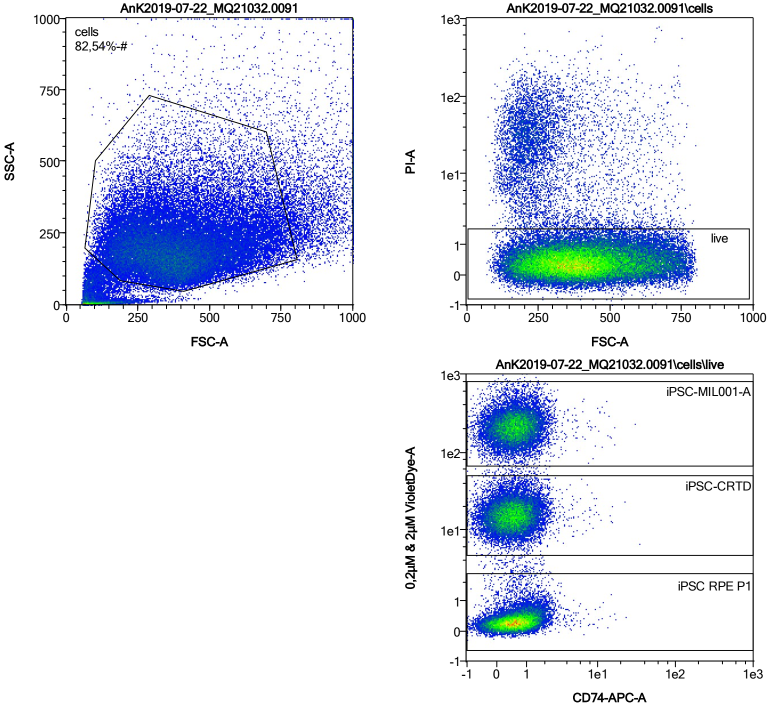


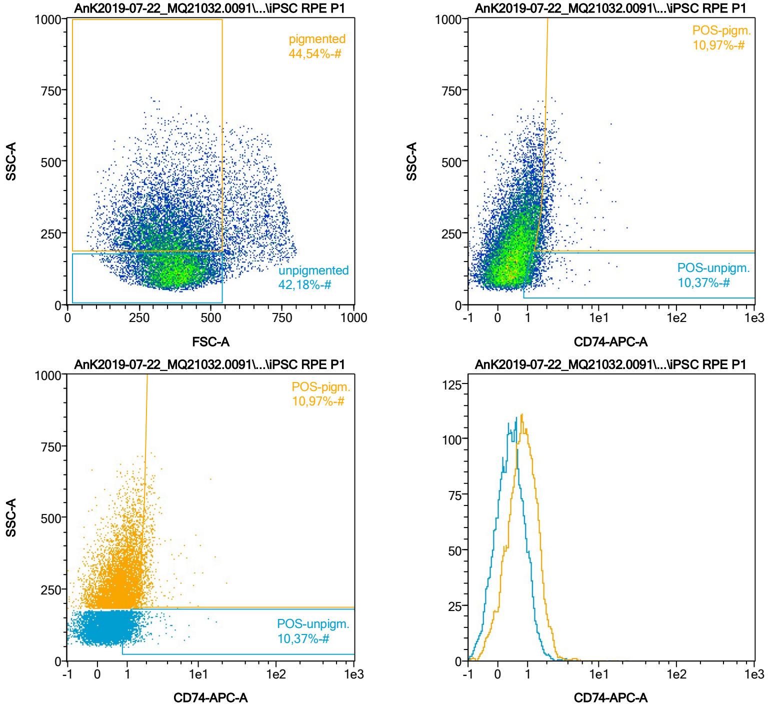


| Name | %-# | CD74-APC-A Mean | CD74-APC-A Median |
|---|---|---|---|
| POS-unpigm. | 10,37 | 1,43 | 1,21 |
| POS-pigm. | 10,97 | 2,12 | 1,84 |

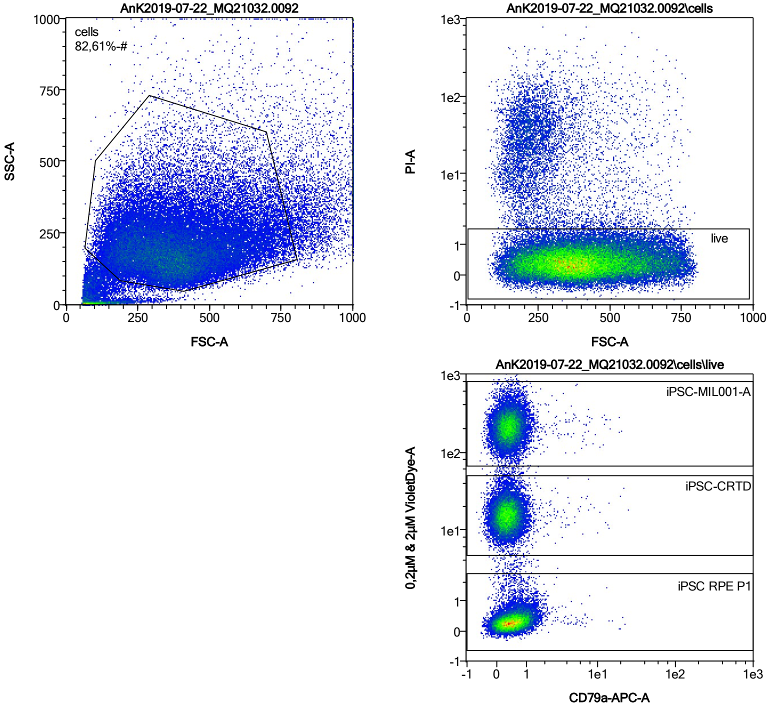


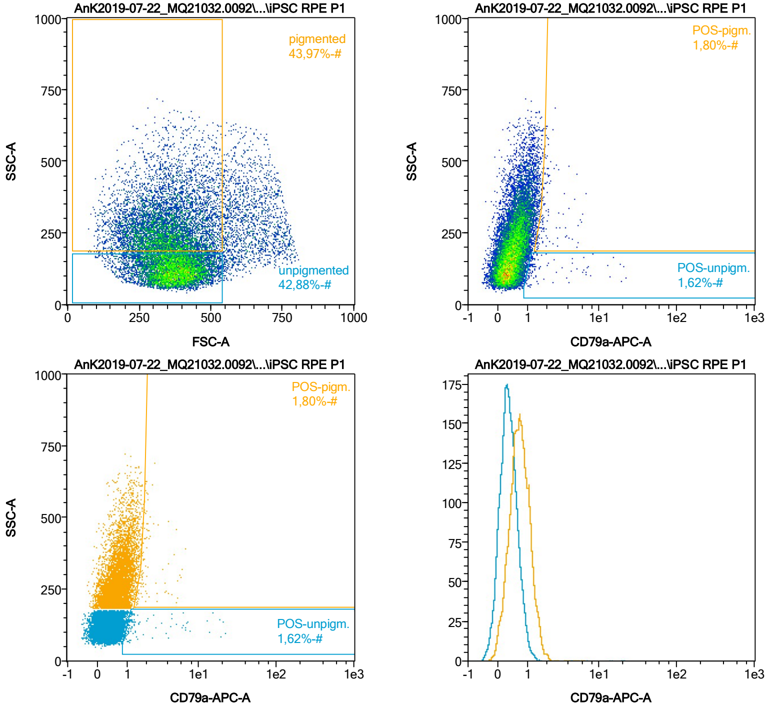


| Name | %-# | CD79a-APC-A Mean | CD79a-APC-A Median |
|---|---|---|---|
| POS-unpigm. | 1,62 | 2,97 | 1,22 |
| POS-pigm. | 1,80 | 2,28 | 1,87 |

Date:2024-06-14

Operator: Sebastian Knoebel

Version:2.13.0+54.g95abf0f4

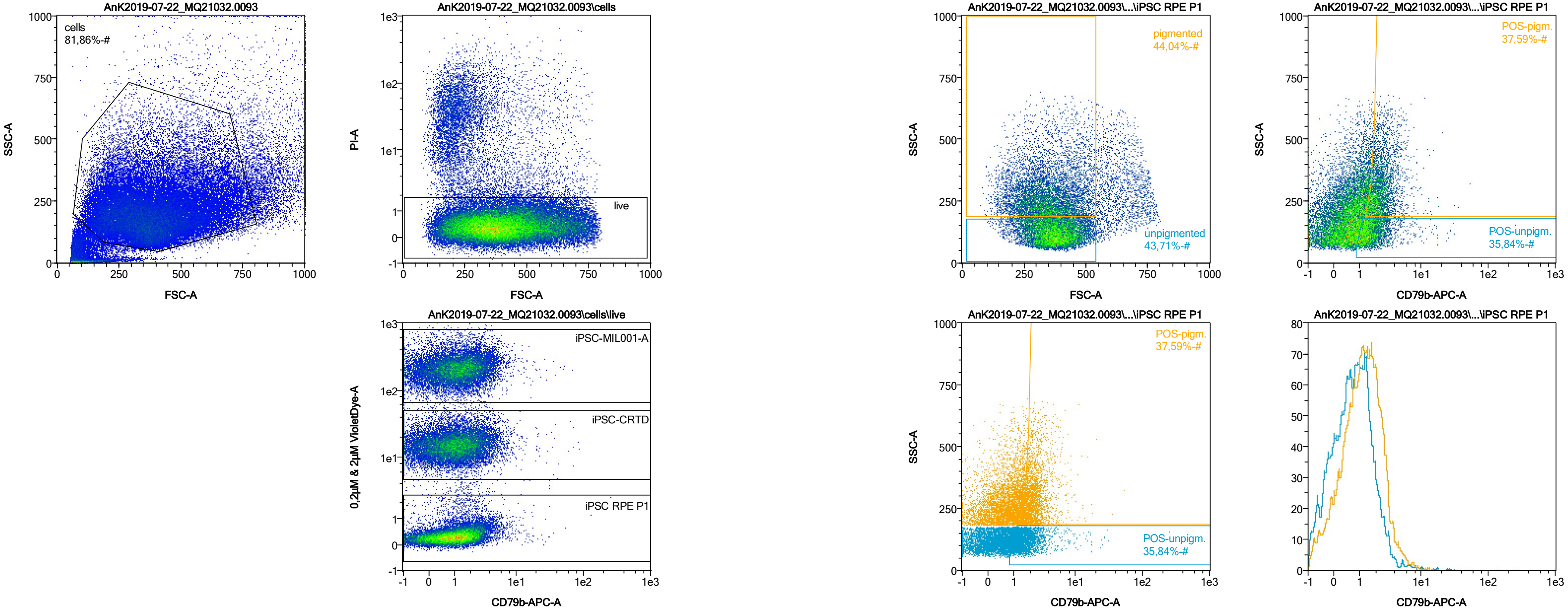


| Name | %-# | CD79b-APC-A Mean | CD79b-APC-A Median |
|---|---|---|---|
| POS-unpigm. | 35,84 | 2,13 | 1,52 |
| POS-pigm. | 37,59 | 2,56 | 2,17 |

Date:2024-06-14
Operator: Sebastian Knoebel
Version:2.13.0+54.g95abf0f4

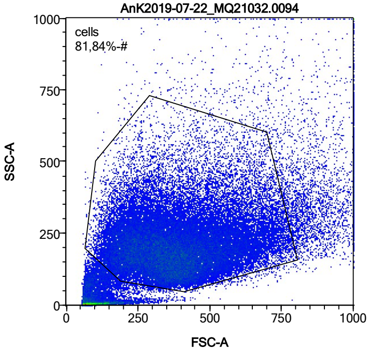


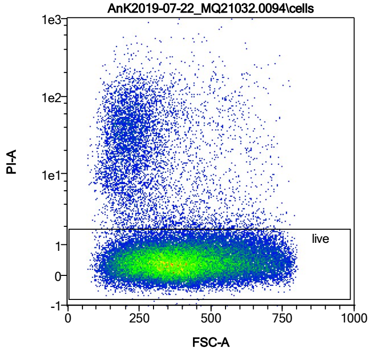


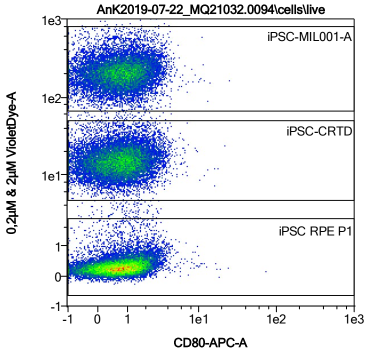


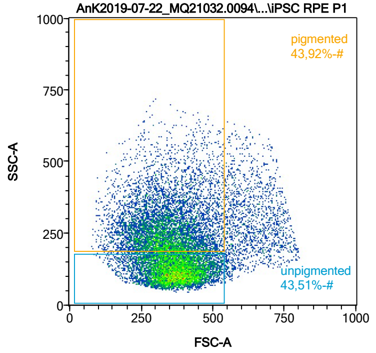


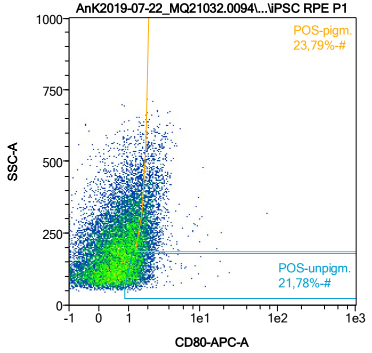


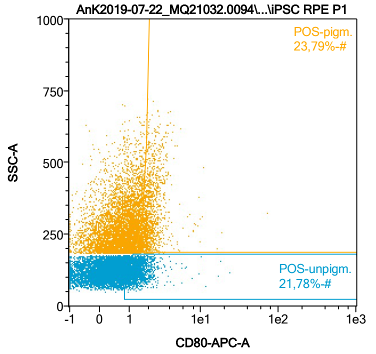


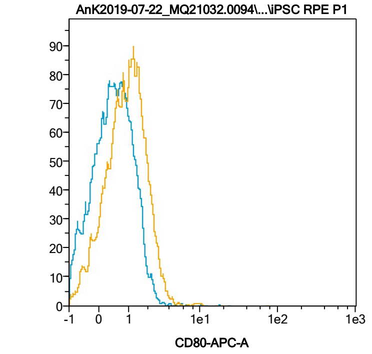


| Name | %-# | CD80-APC-A Mean | CD80-APC-A Median |
|---|---|---|---|
| POS-unpigm. | 21,78 | 1,54 | 1,36 |
| POS-pigm. | 23,79 | 2,23 | 1,97 |

Date:2024-06-14
Operator: Sebastian Knoebel
Version:2.13.0+54.g95abf0f4

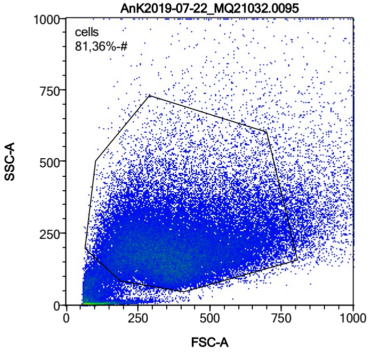


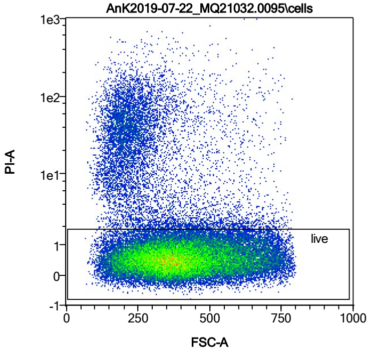


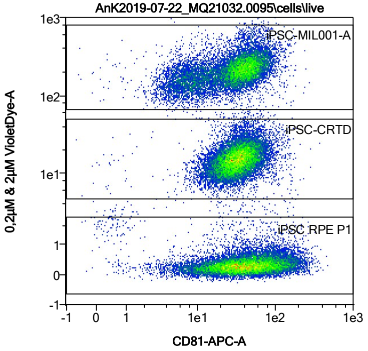


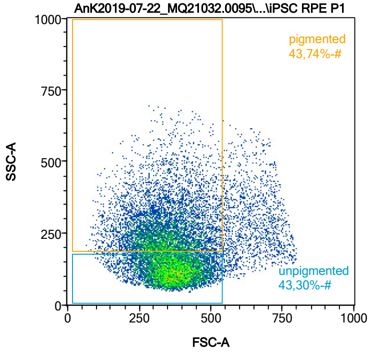


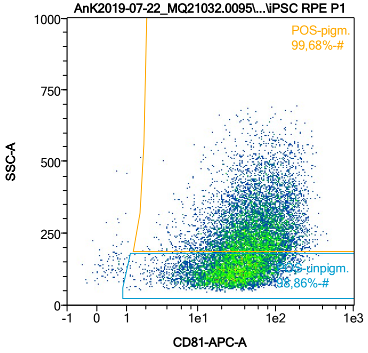


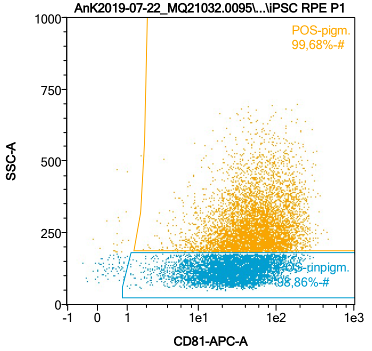


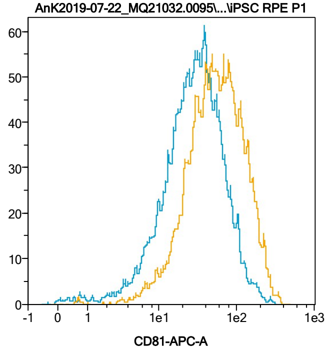


| Name | %-# | CD81-APC-A Mean | CD81-APC-A Median |
|---|---|---|---|
| POS-unpigm. | 98,86 | 37,87 | 28,98 |
| POS-pigm. | 99,68 | 70,51 | 54,59 |

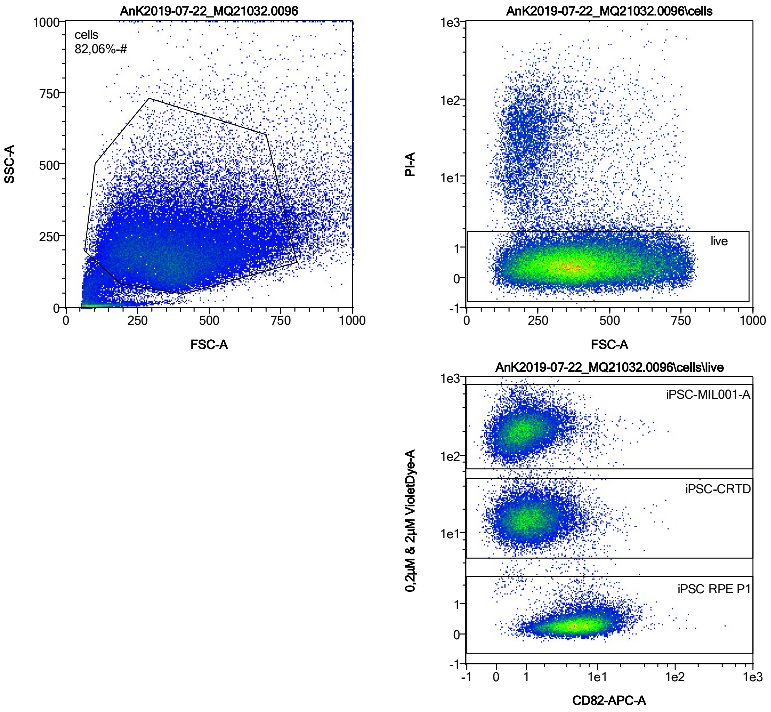


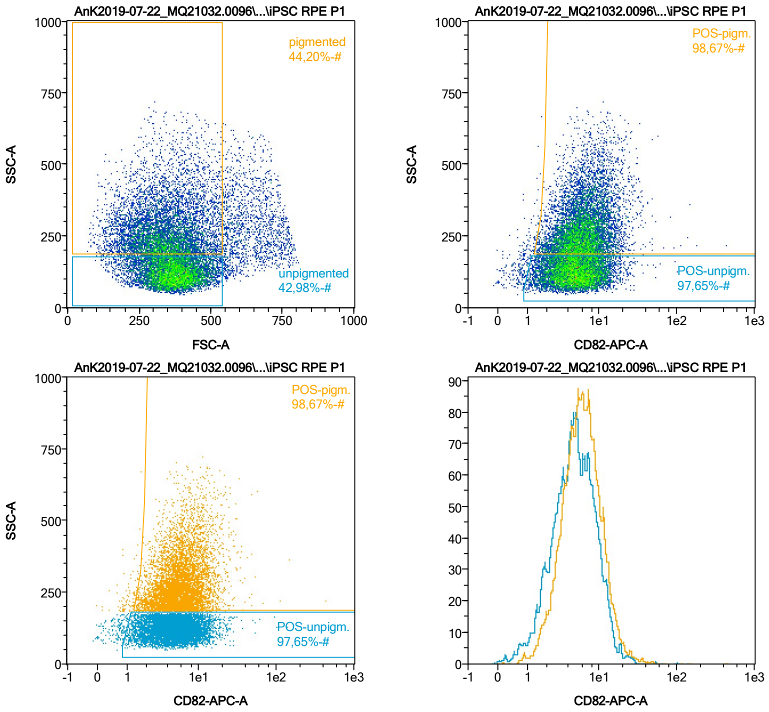


| Name | %-# | CD82-APC-A Mean | CD82-APC-A Median |
|---|---|---|---|
| POS-unpigm. | 97,65 | 5,75 | 4,85 |
| POS-pigm. | 98,67 | 6,98 | 5,89 |

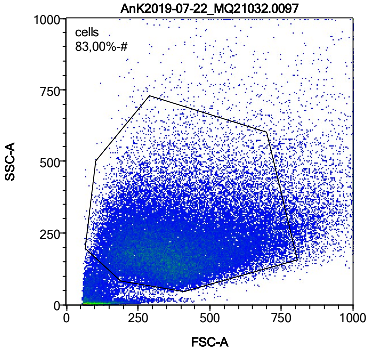


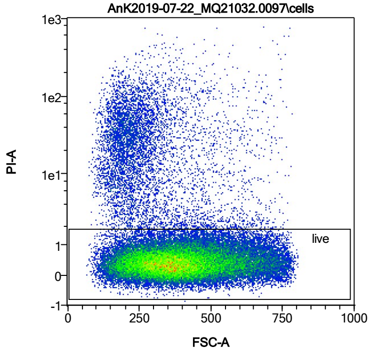


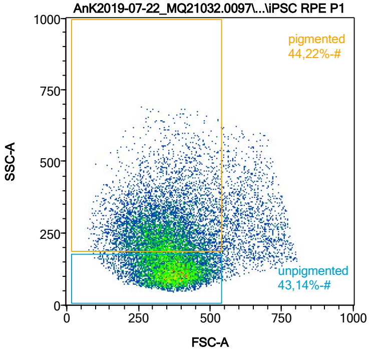


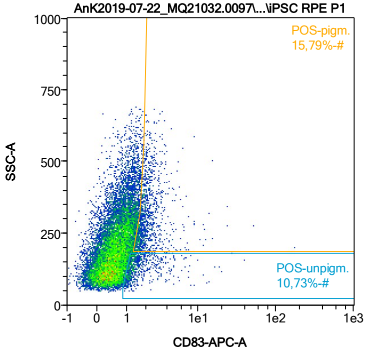


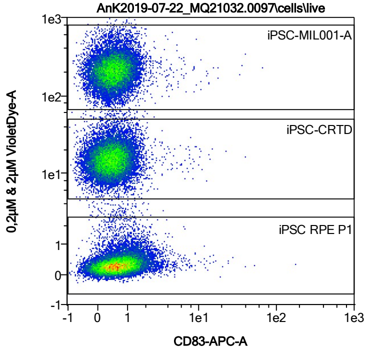


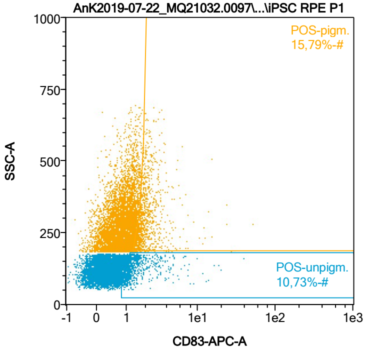


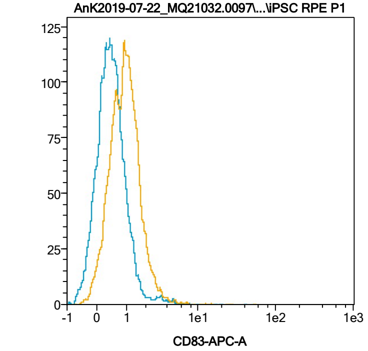


| Name | %-# | CD83-APC-A Mean | CD83-APC-A Median |
|---|---|---|---|
| POS-unpigm. | 10,73 | 1,85 | 1,30 |
| POS-pigm. | 15,79 | 2,33 | 1,91 |

Date:2024-06-14

Operator: Sebastian Knoebel

Version:2.13.0+54.g95abf0f4

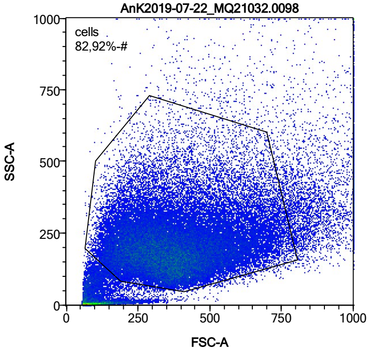


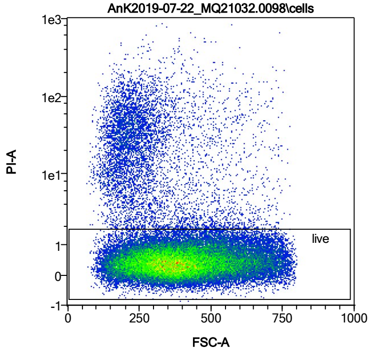


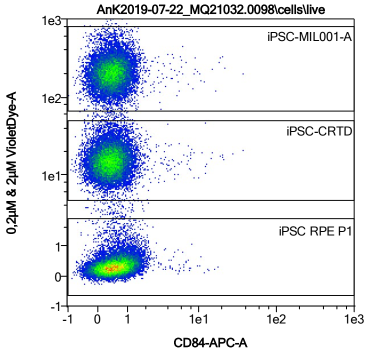


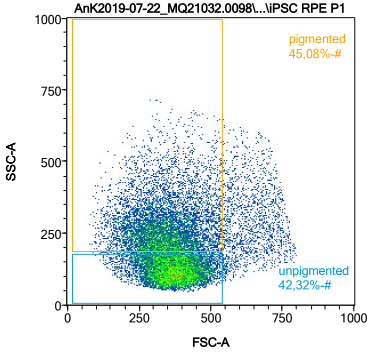


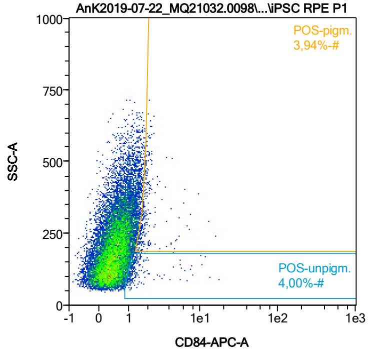


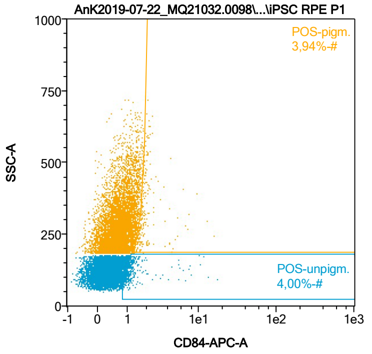


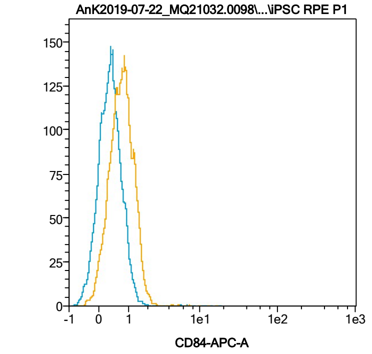


| Name | %-# | CD84-APC-A Mean | CD84-APC-A Median |
|---|---|---|---|
| POS-unpigm. | 4,00 | 1,69 | 1,18 |
| POS-pigm. | 3,94 | 2,11 | 1,79 |

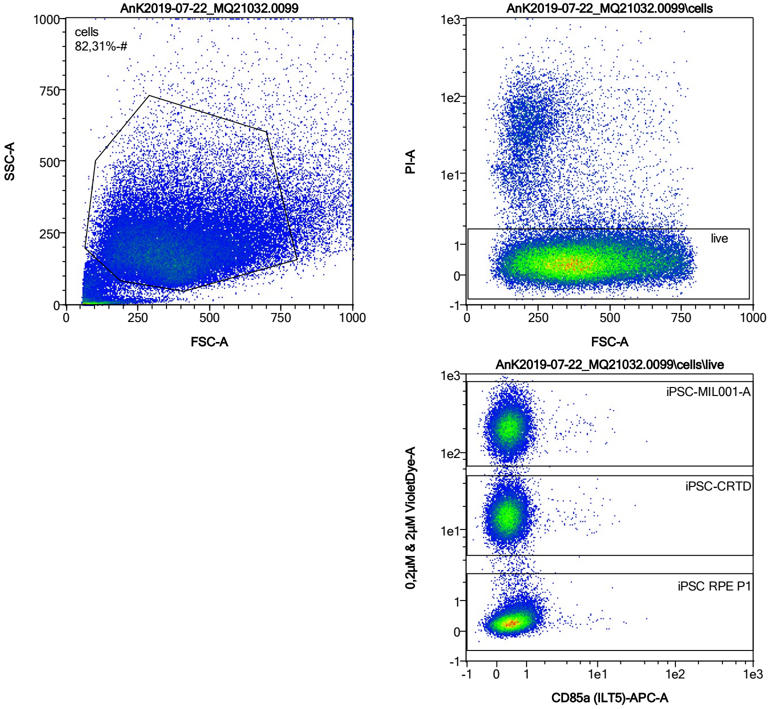


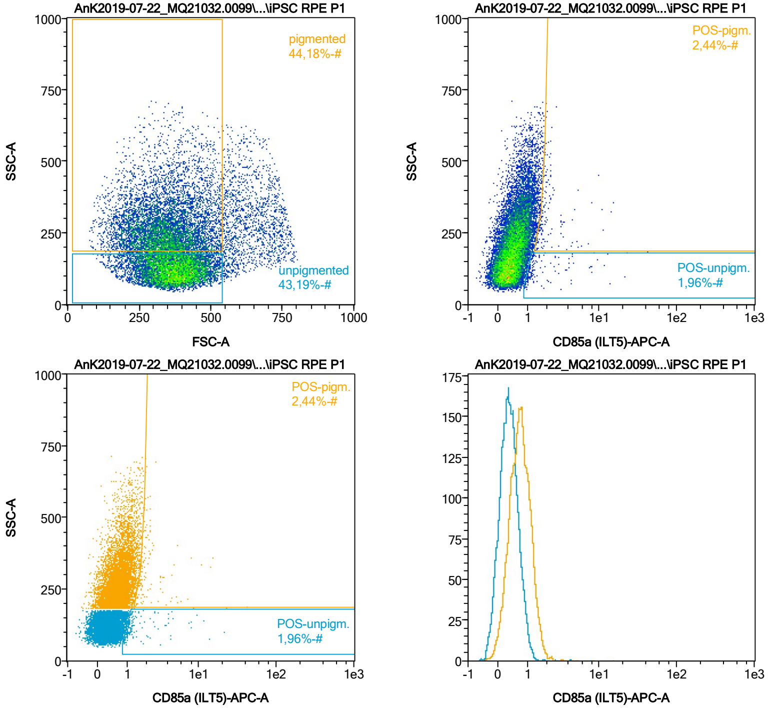


| Name | %-# | CD85a (ILT5)-APC-A Mean | CD85a (ILT5)-APC-A Median |
|---|---|---|---|
| POS-unpigm. | 1,96 | 2,05 | 1,19 |
| POS-pigm. | 2,44 | 2,75 | 1,83 |

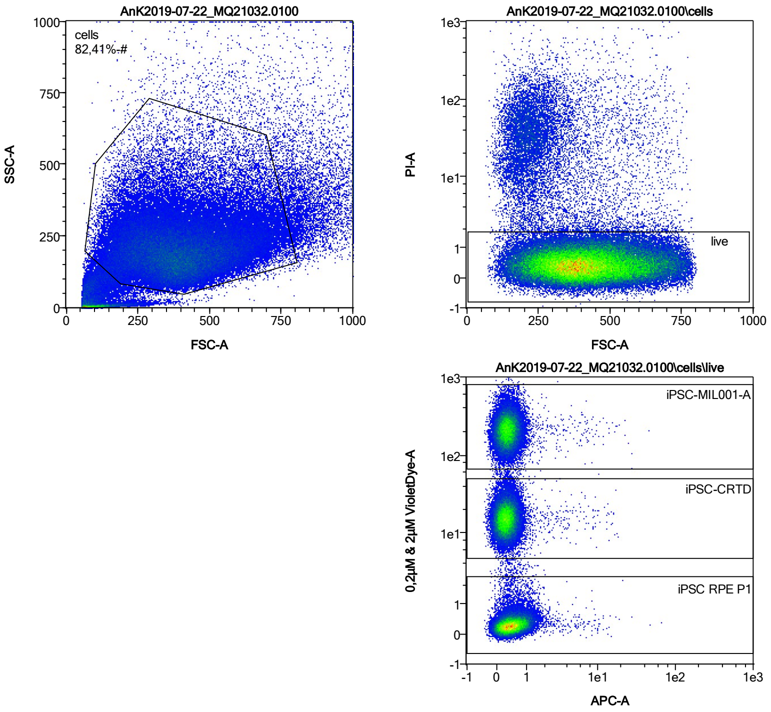


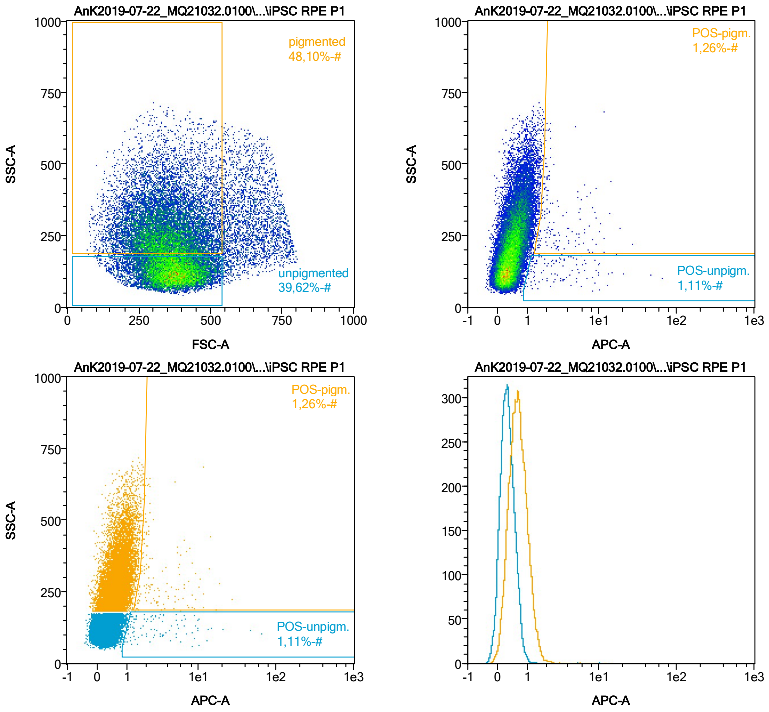


| Name | %-# | APC-A Mean | APC-A Median |
|---|---|---|---|
| POS-unpigm. | 1,11 | 4,61 | 2,37 |
| POS-pigm. | 1,26 | 3,83 | 2,24 |

Date:2024-06-14

Operator: Sebastian Knoebel

Version:2.13.0+54.g95abf0f4

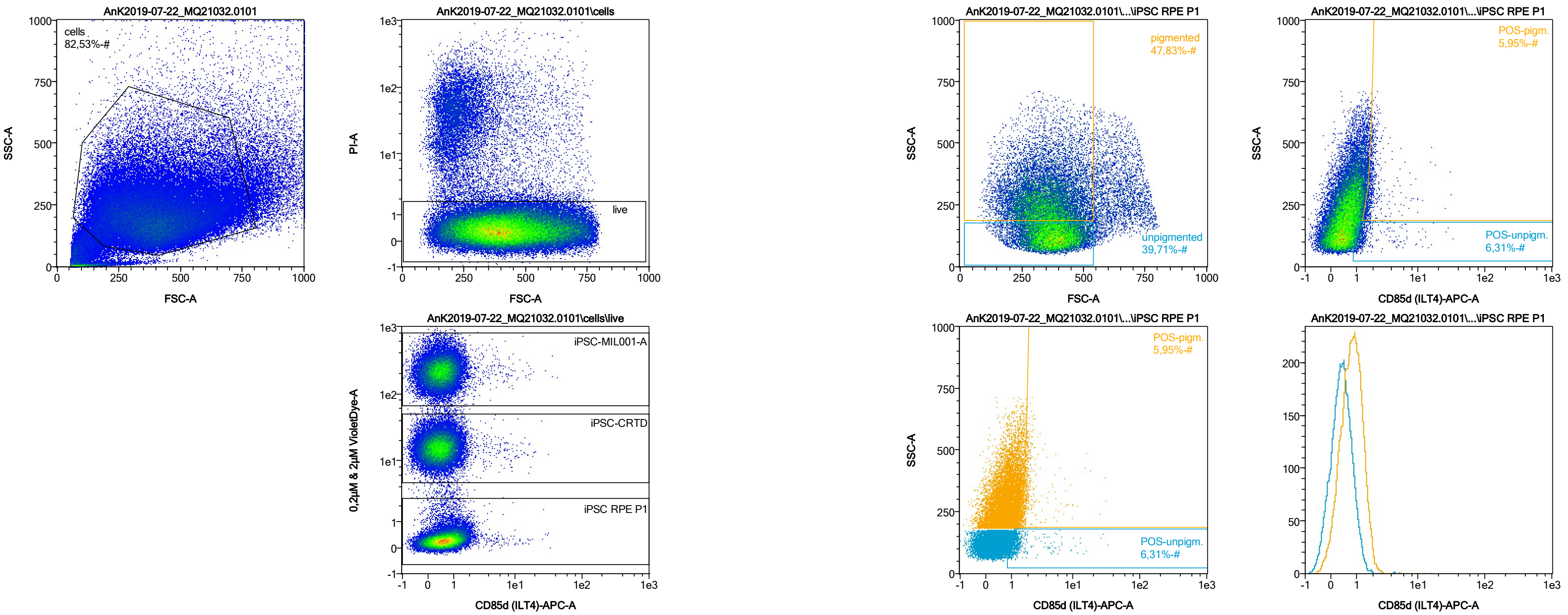


| Name | %-# | CD85d (ILT4)-APC-A Mean | CD85d (ILT4)-APC-A Median |
|---|---|---|---|
| POS-unpigm. | 6,31 | 1,70 | 1,20 |
| POS-pigm. | 5,95 | 2,18 | 1,81 |

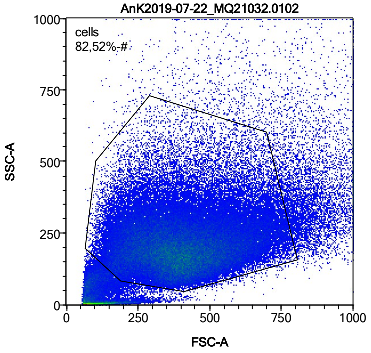


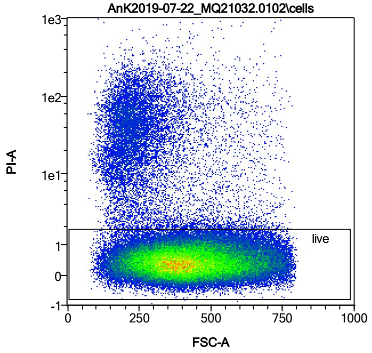


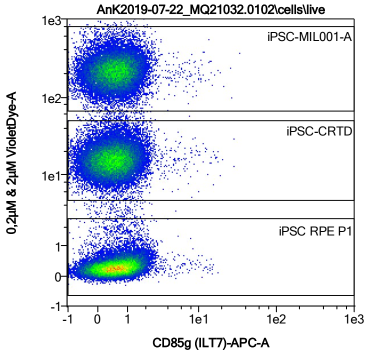


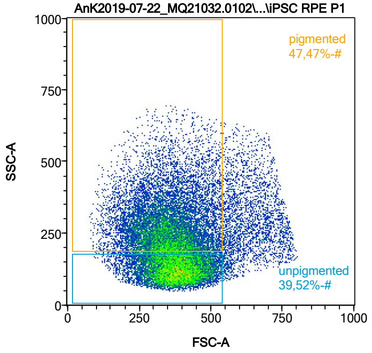


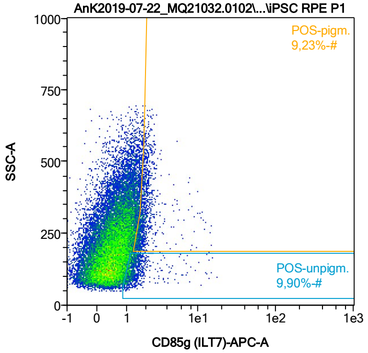


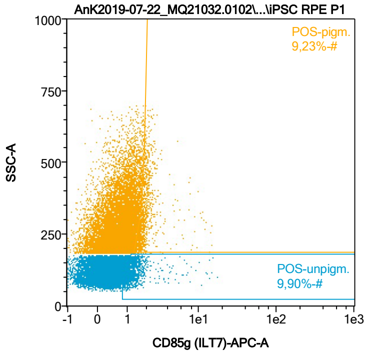


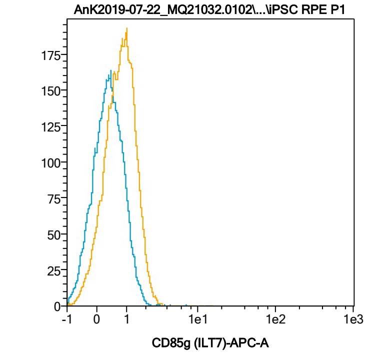


| Name | %-# | CD85g (ILT7)-APC-A Mean | CD85g (ILT7)-APC-A Median |
|---|---|---|---|
| POS-unpigm. | 9,90 | 1,52 | 1,23 |
| POS-pigm. | 9,23 | 2,08 | 1,83 |

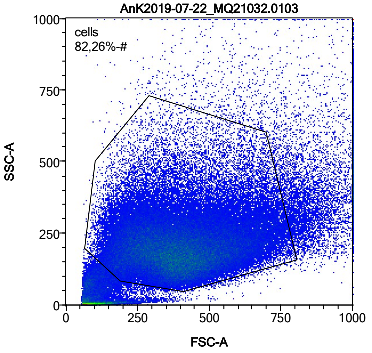


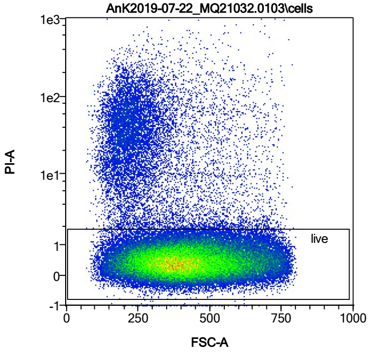


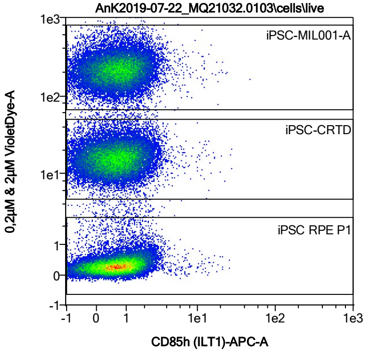


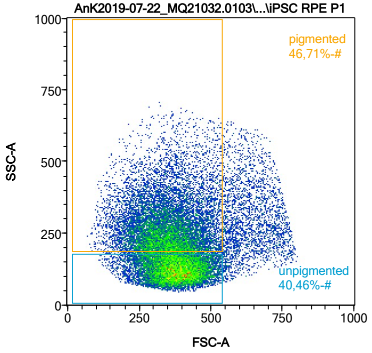


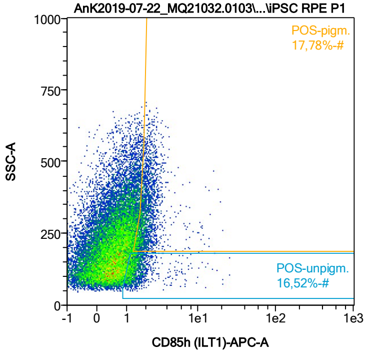


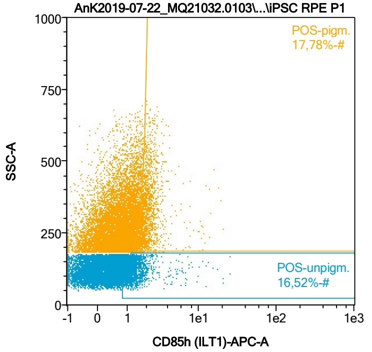


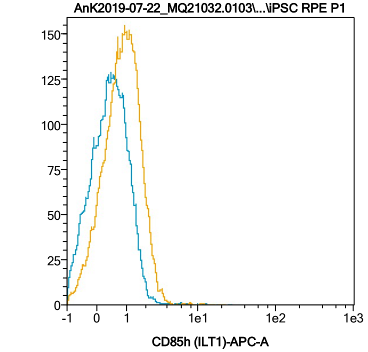


| Name | %-# | CD85h (ILT1)-APC-A Mean | CD85h (ILT1)-APC-A Median |
|---|---|---|---|
| POS-unpigm. | 16,52 | 1,60 | 1,30 |
| POS-pigm. | 17,78 | 2,15 | 1,90 |

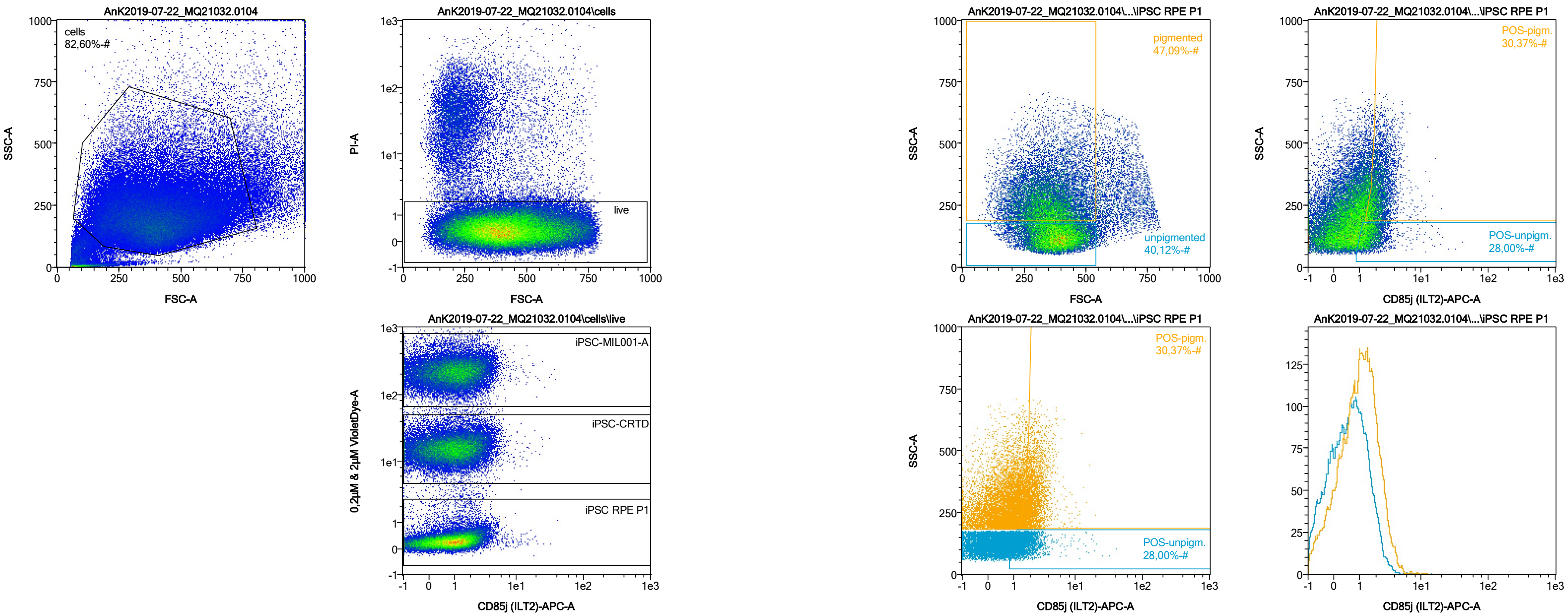


| Name | %-# | CD85j (ILT2)-APC-A Mean | CD85j (ILT2)-APC-A Median |
|---|---|---|---|
| POS-unpigm. | 28,00 | 1,76 | 1,47 |
| POS-pigm. | 30,37 | 2,34 | 2,07 |

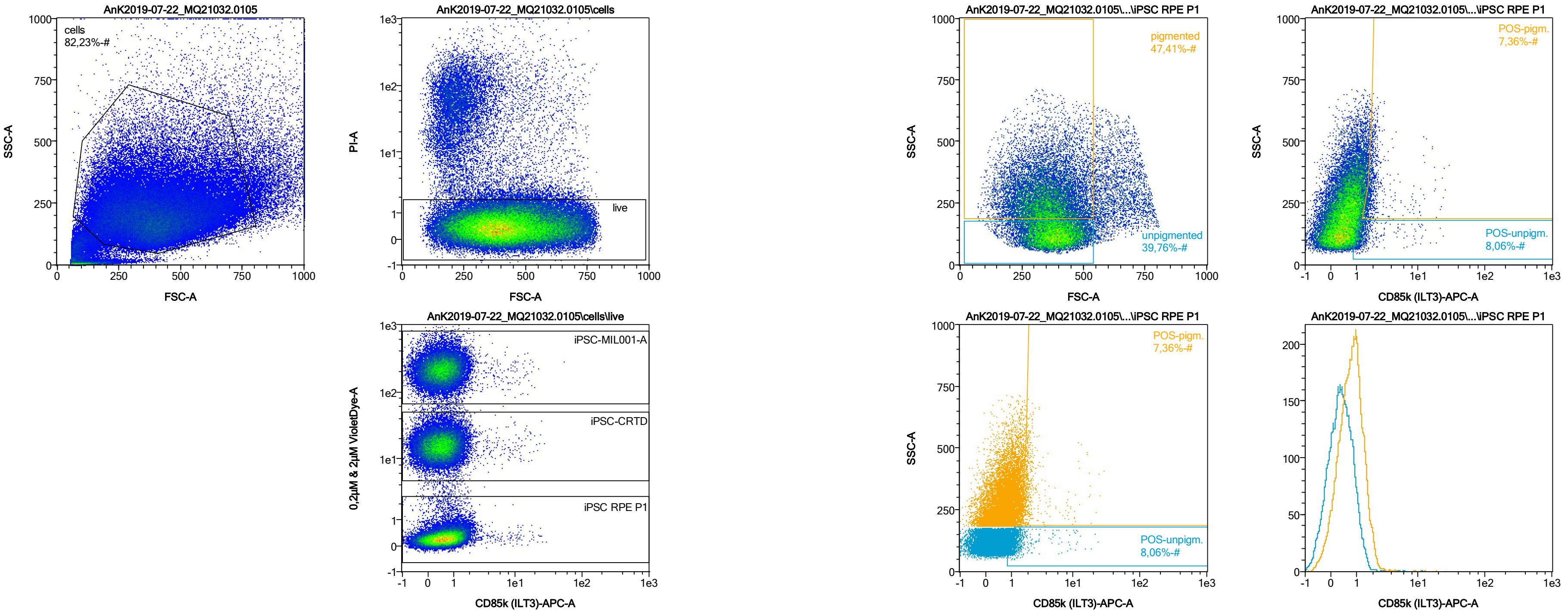


| Name | %-# | CD85k (ILT3)-APC-A Mean | CD85k (ILT3)-APC-A Median |
|---|---|---|---|
| POS-unpigm. | 8,06 | 1,63 | 1,21 |
| POS-pigm. | 7,36 | 2,37 | 1,85 |

Date:2024-06-14

Operator: Sebastian Knoebel

Version:2.13.0+54.g95abf0f4

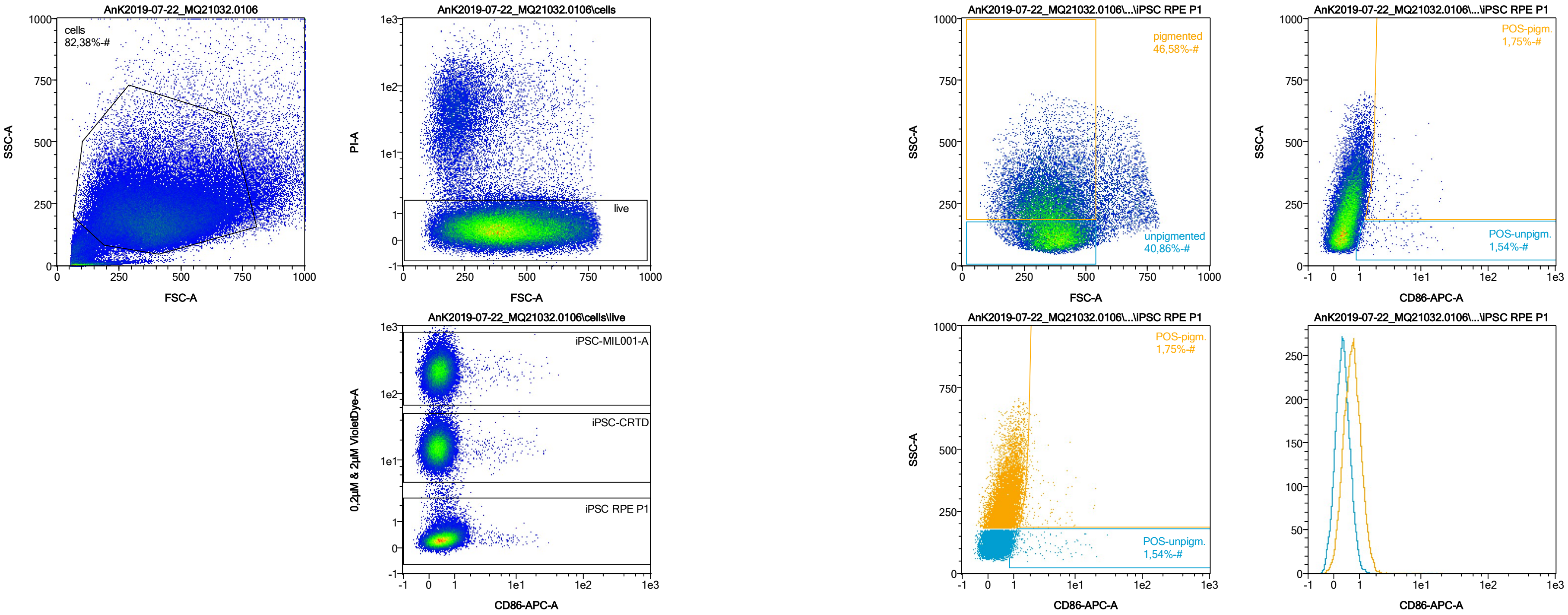


| Name | %-# | CD86-APC-A Mean | CD86-APC-A Median |
|---|---|---|---|
| POS-unpigm. | 1,54 | 3,28 | 1,36 |
| POS-pigm. | 1,75 | 2,88 | 1,87 |

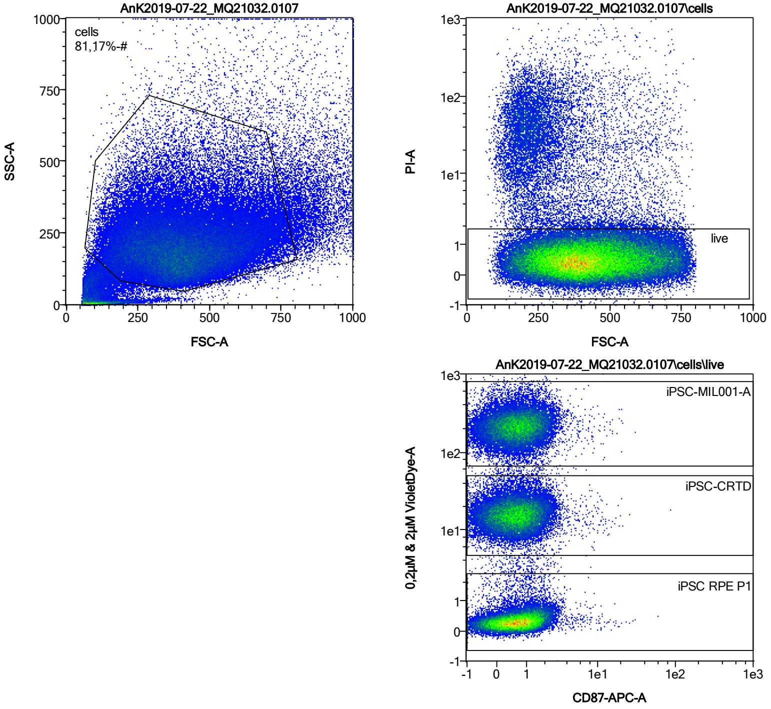


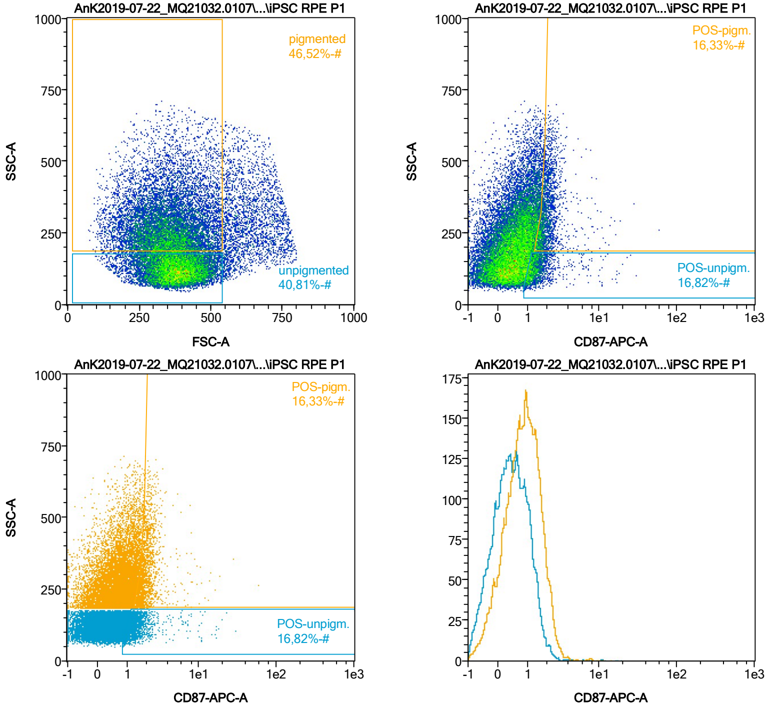


| Name | %-# | CD87-APC-A Mean | CD87-APC-A Median |
|---|---|---|---|
| POS-unpigm. | 16,82 | 1,62 | 1,30 |
| POS-pigm. | 16,33 | 2,16 | 1,88 |

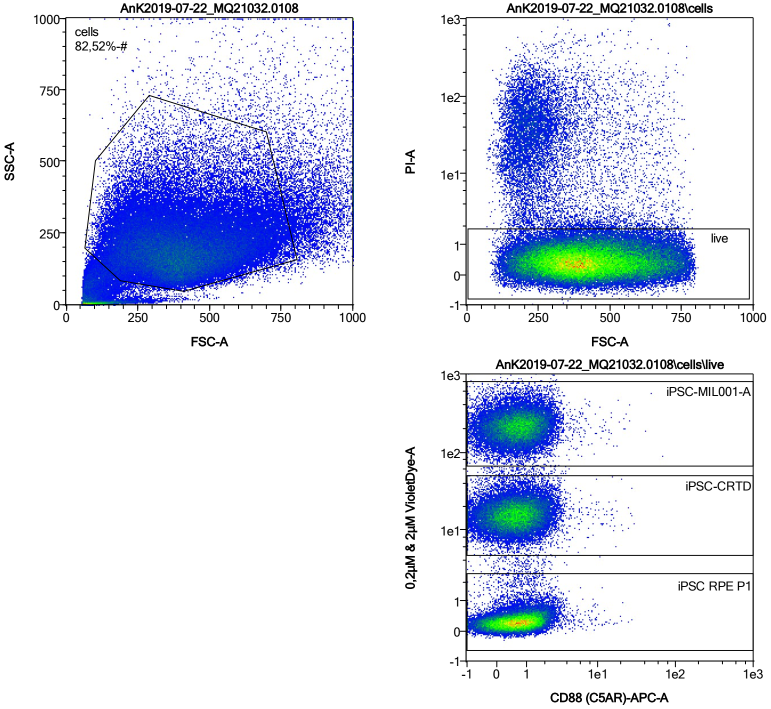


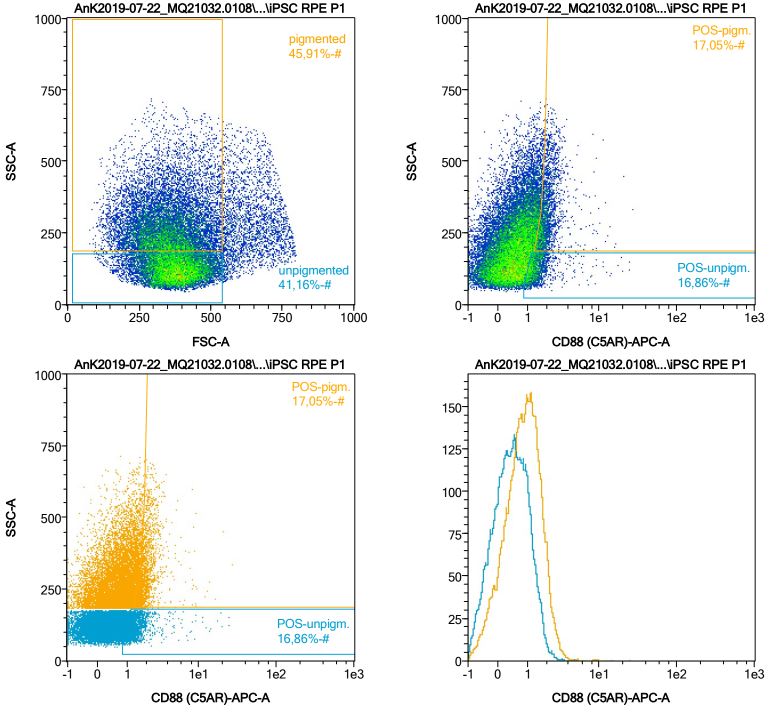


| Name | %-# | CD88 (C5AR)-APC-A Mean | CD88 (C5AR)-APC-A Median |
|---|---|---|---|
| POS-unpigm. | 16,86 | 1,54 | 1,29 |
| POS-pigm. | 17,05 | 2,11 | 1,88 |

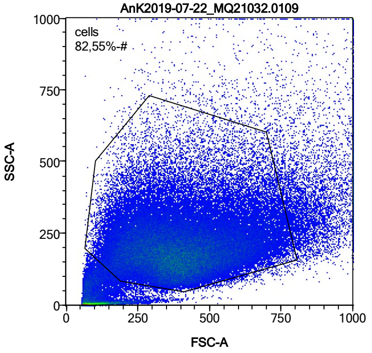


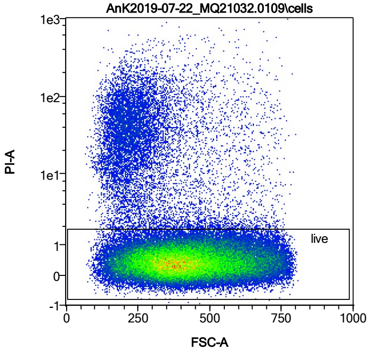


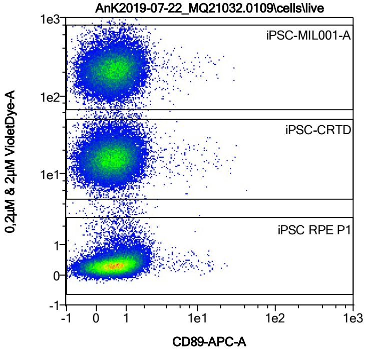


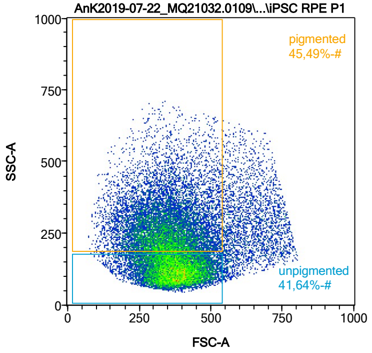


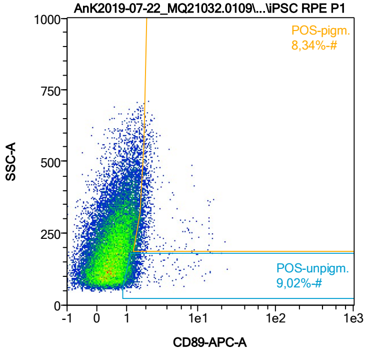


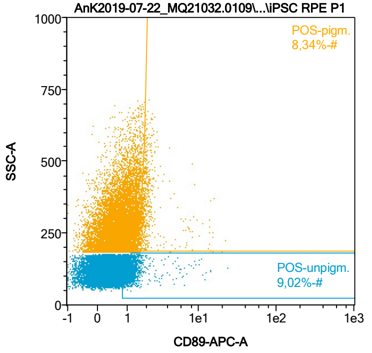


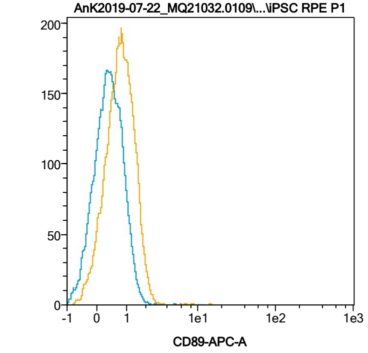


| Name | %-# | CD89-APC-A Mean | CD89-APC-A Median |
|---|---|---|---|
| POS-unpigm. | 9,02 | 1,52 | 1,21 |
| POS-pigm. | 8,34 | 2,21 | 1,80 |

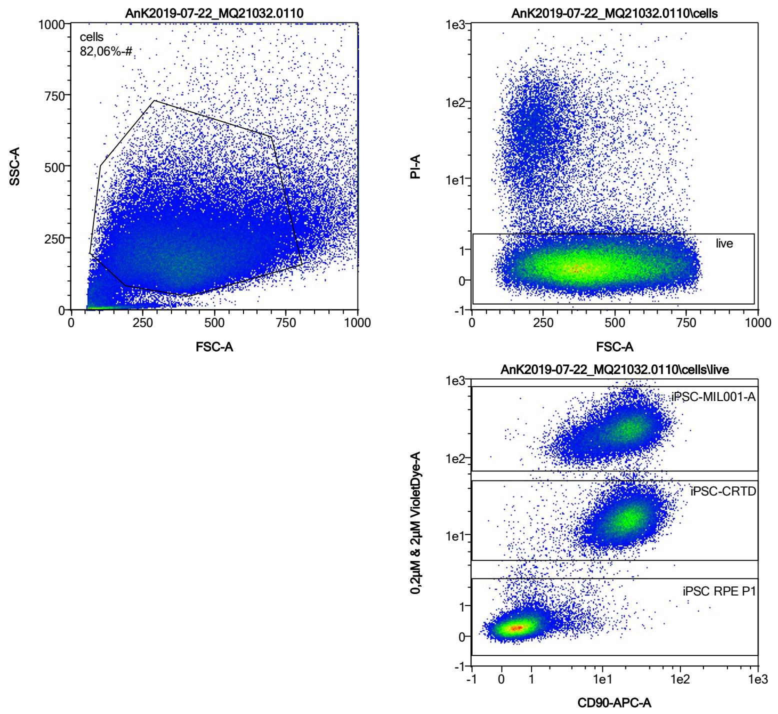


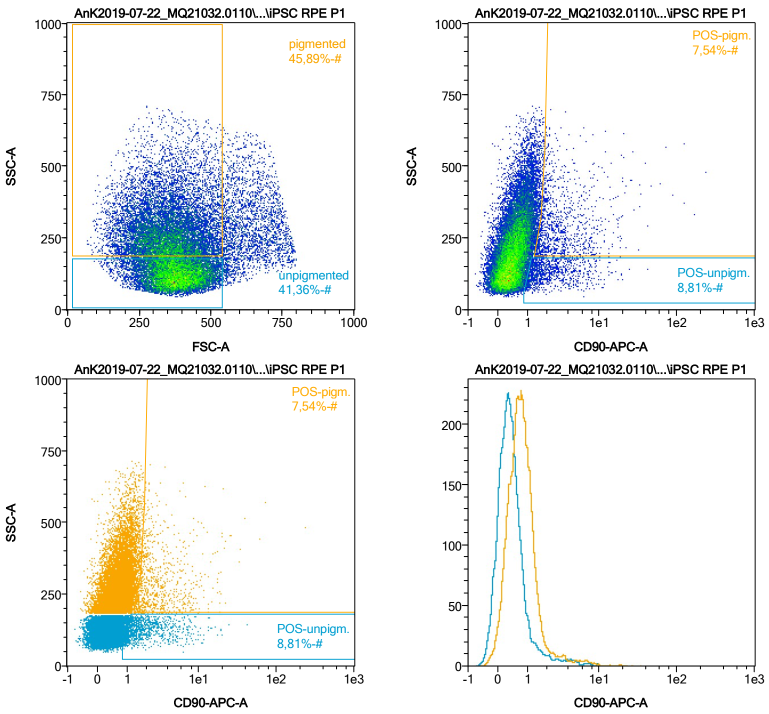


| Name | %-# | CD90-APC-A Mean | CD90-APC-A Median |
|---|---|---|---|
| POS-unpigm. | 8,81 | 2,65 | 1,68 |
| POS-pigm. | 7,54 | 4,20 | 2,40 |

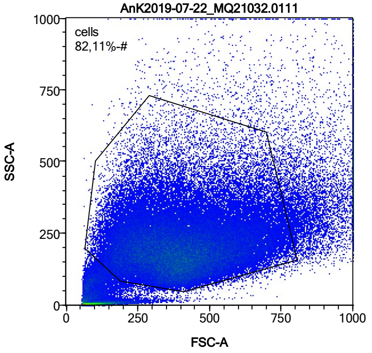


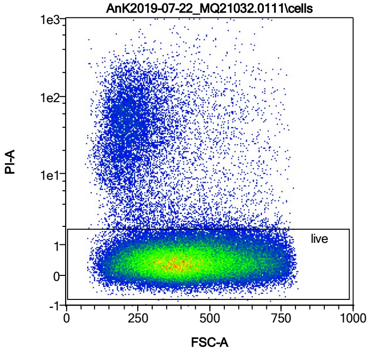


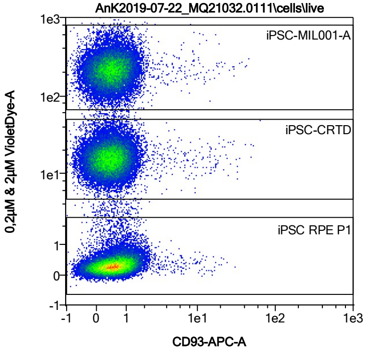


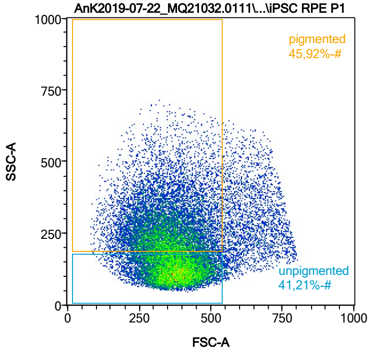


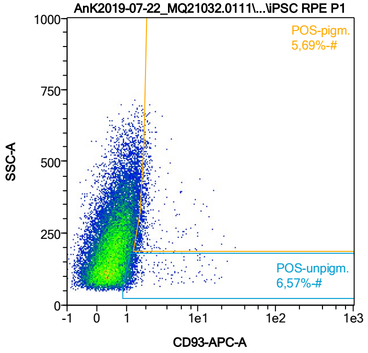


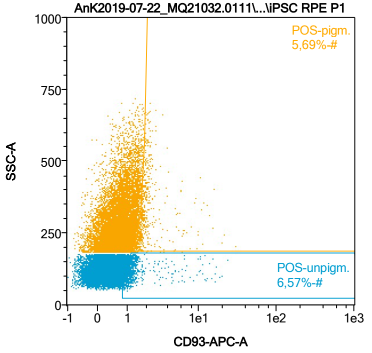


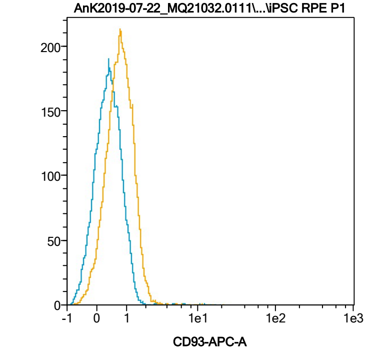


| Name | %-# | CD93-APC-A Mean | CD93-APC-A Median |
|---|---|---|---|
| POS-unpigm. | 6,57 | 1,81 | 1,18 |
| POS-pigm. | 5,69 | 2,28 | 1,80 |

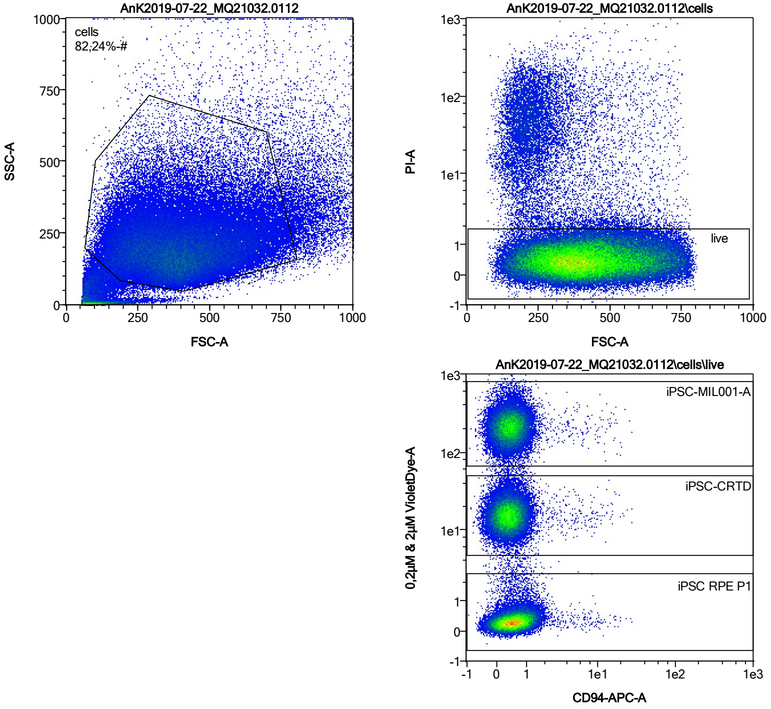


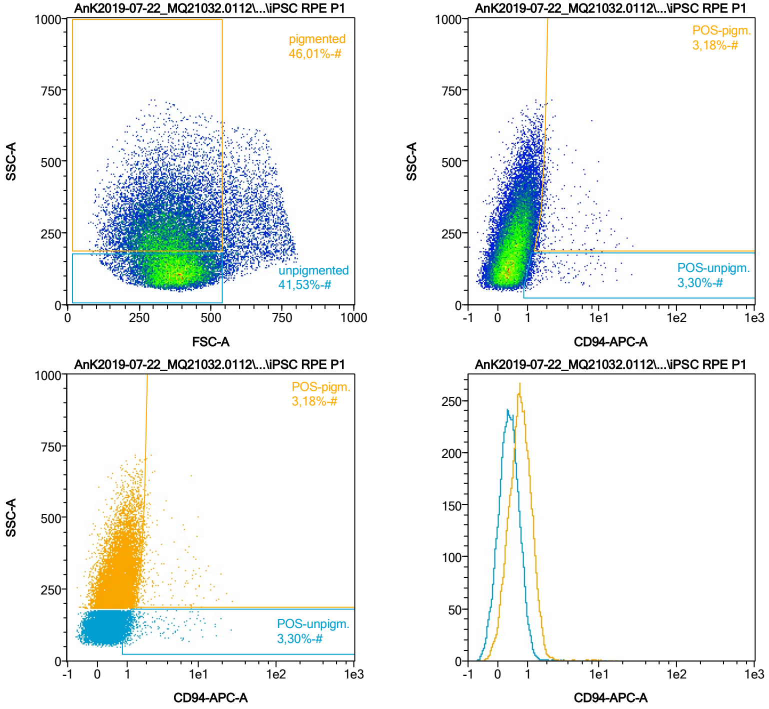


| Name | %-# | CD94-APC-A Mean | CD94-APC-A Median |
|---|---|---|---|
| POS-unpigm. | 3,30 | 2,15 | 1,18 |
| POS-pigm. | 3,18 | 2,94 | 1,84 |

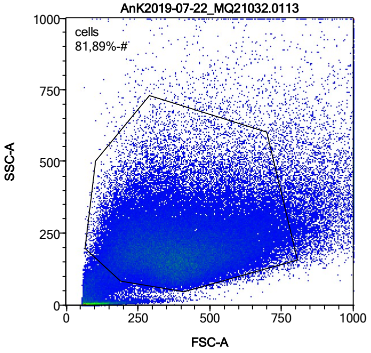


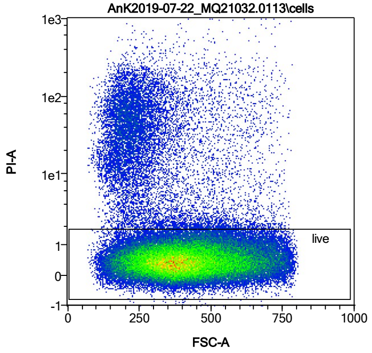


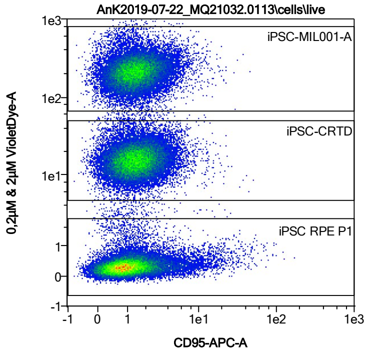


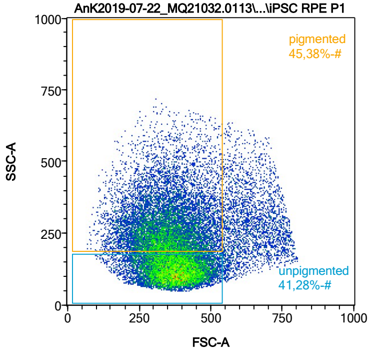


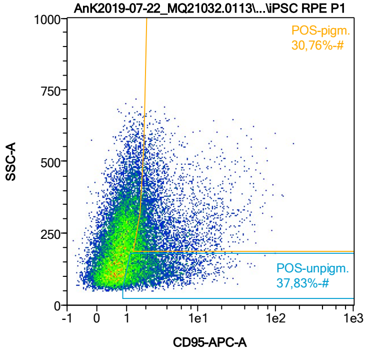


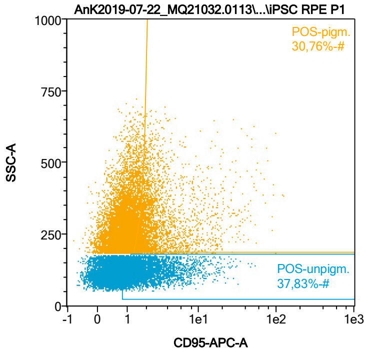


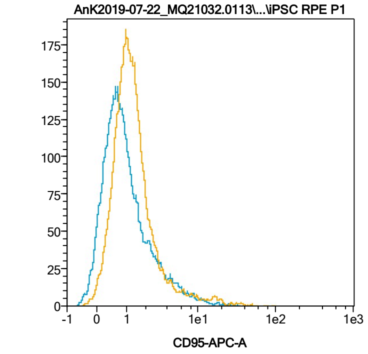


| Name | %-# | CD95-APC-A Mean | CD95-APC-A Median |
|---|---|---|---|
| POS-unpigm. | 37,83 | 2,87 | 1,82 |
| POS-pigm. | 30,76 | 4,44 | 2,26 |

Date:2024-06-14

Operator: Sebastian Knoebel

Version:2.13.0+54.g95abf0f4

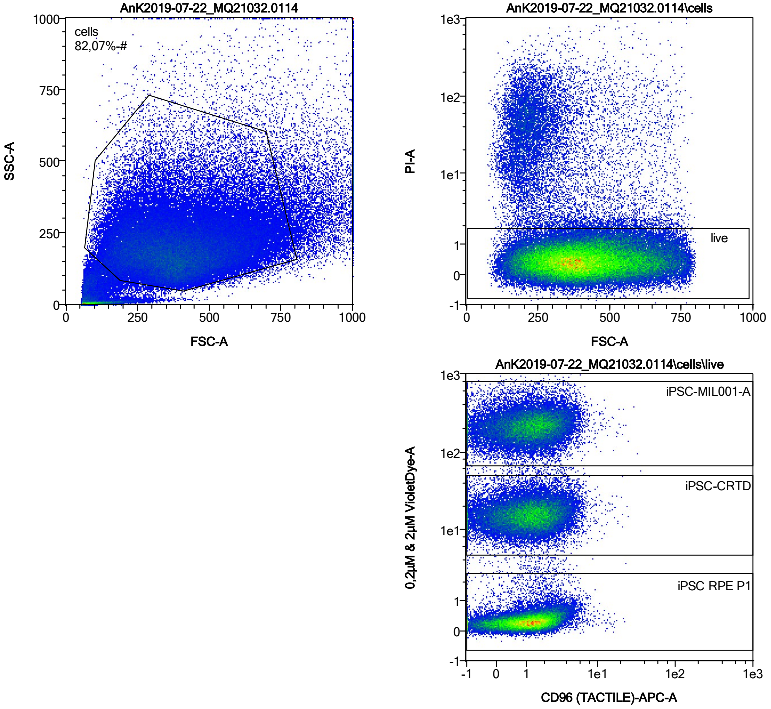


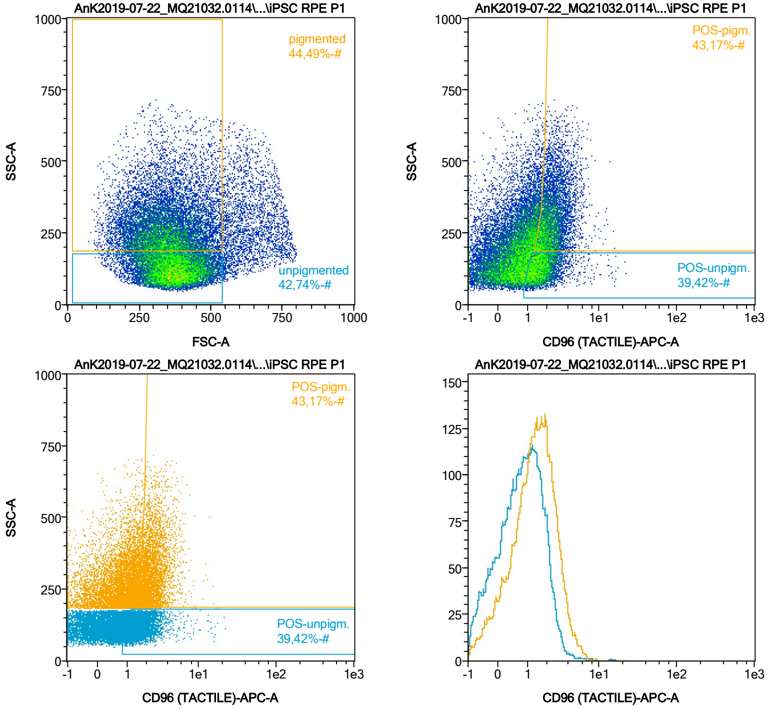


| Name | %-# | CD96 (TACTILE)-APC-A Mean | CD96 (TACTILE)-APC-A Median |
|---|---|---|---|
| POS-unpigm. | 39,42 | 1,81 | 1,57 |
| POS-pigm. | 43,17 | 2,48 | 2,22 |

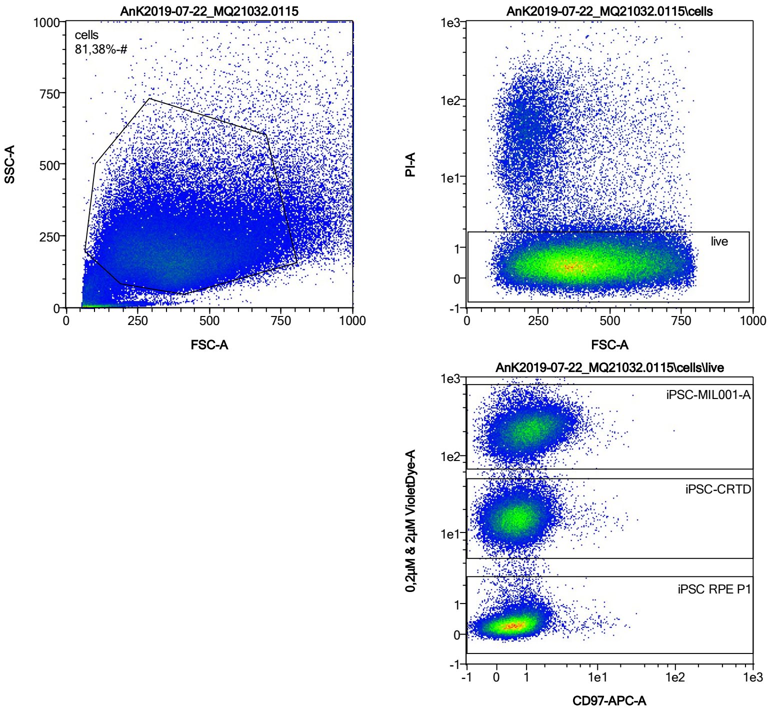


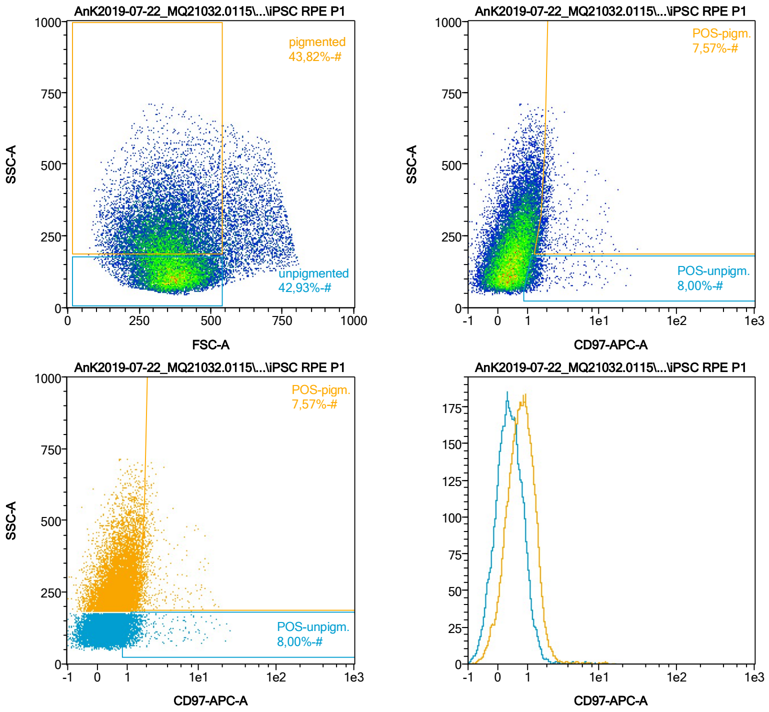


| Name | %-# | CD97-APC-A Mean | CD97-APC-A Median |
|---|---|---|---|
| POS-unpigm. | 8,00 | 1,81 | 1,23 |
| POS-pigm. | 7,57 | 2,40 | 1,83 |

Date:2024-06-14

Operator: Sebastian Knoebel

Version:2.13.0+54.g95abf0f4

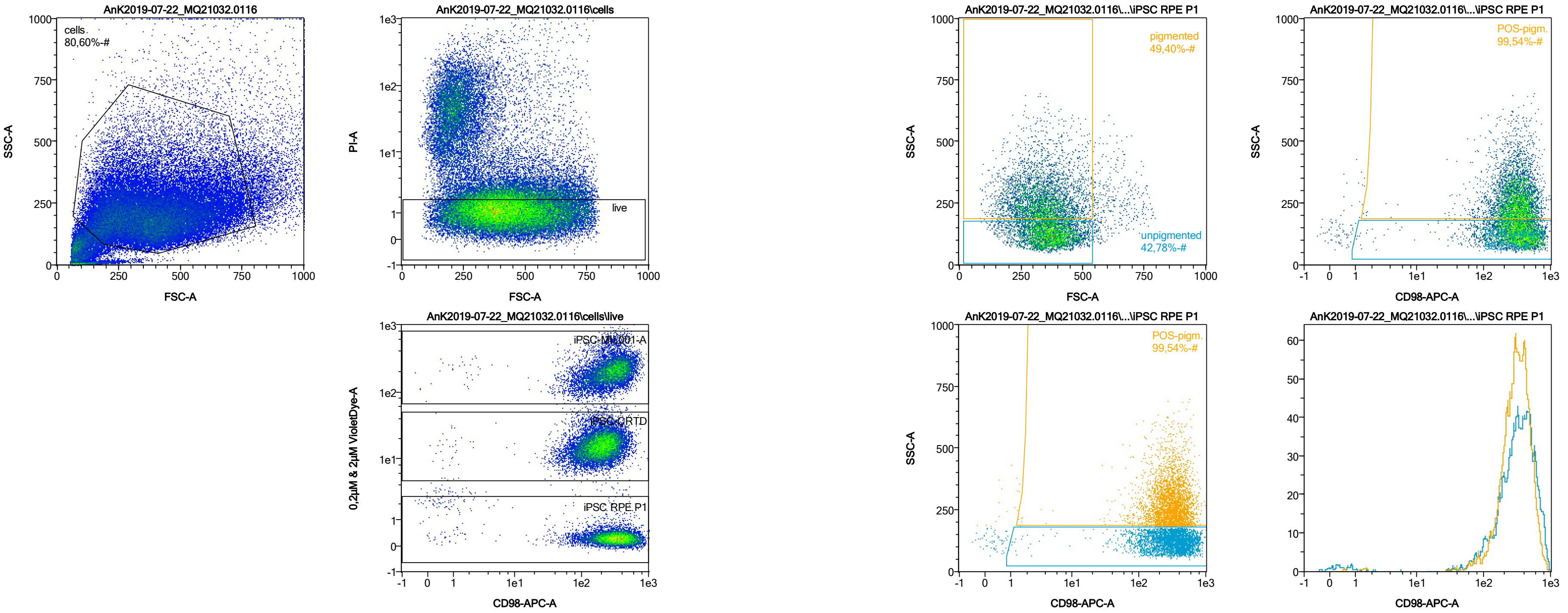


| Name | %-# | CD98-APC-A Mean | CD98-APC-A Median |
|---|---|---|---|
| POS-unpigm. | 98,27 | 354,90 | 330,95 |
| POS-pigm. | 99,54 | 327,07 | 309,44 |

Date:2024-06-14

Operator: Sebastian Knoebel

Version:2.13.0+54.g95abf0f4

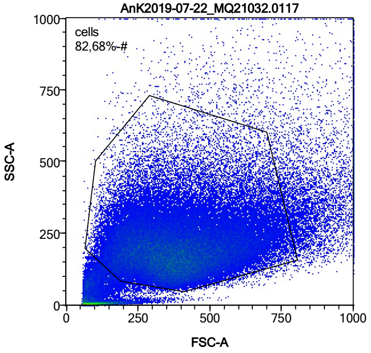


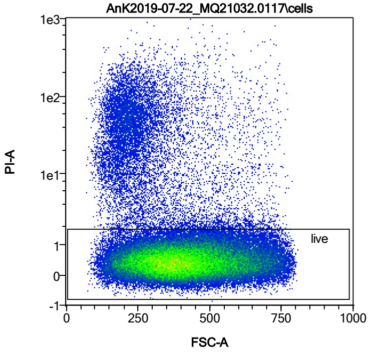


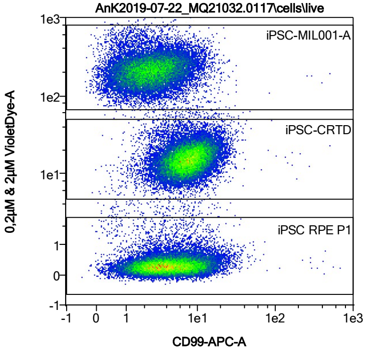


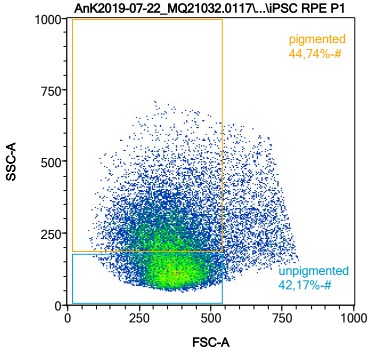


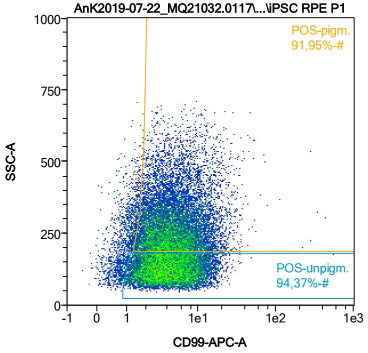


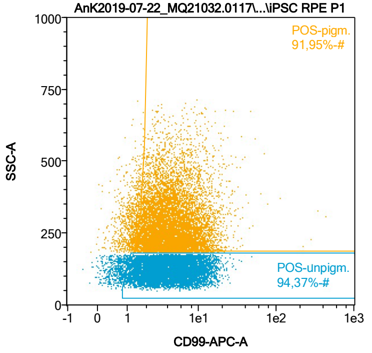


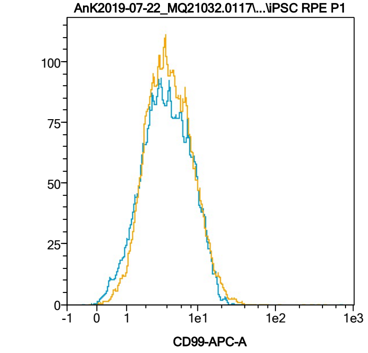


| Name | %-# | CD99-APC-A Mean | CD99-APC-A Median |
|---|---|---|---|
| POS-unpigm. | 94,37 | 5,31 | 4,05 |
| POS-pigm. | 91,95 | 5,74 | 4,29 |

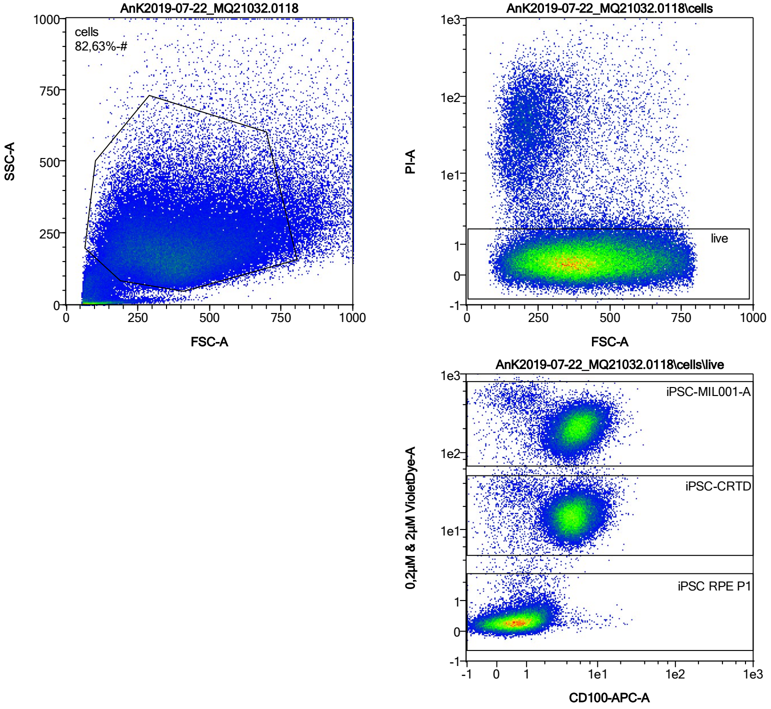


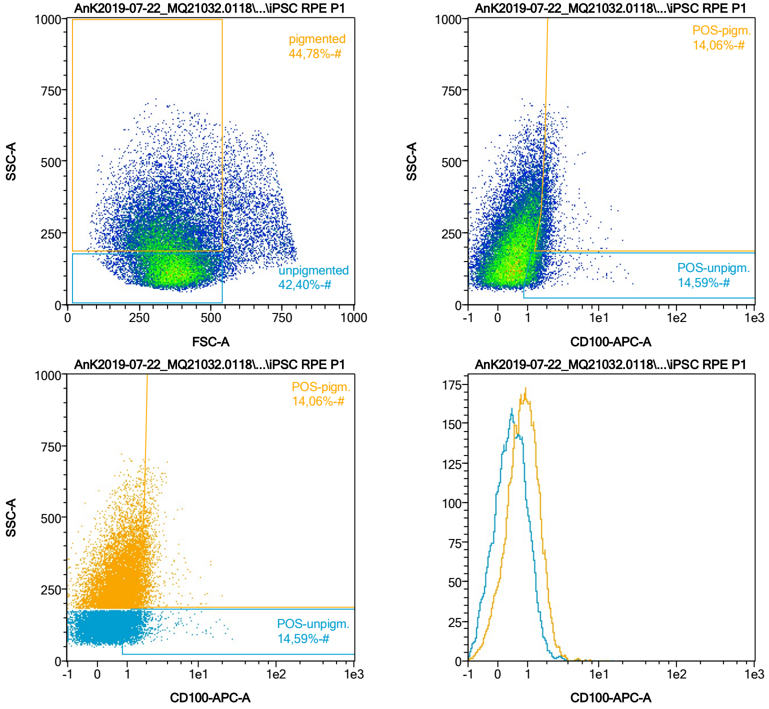


| Name | %-# | CD100-APC-A Mean | CD100-APC-A Median |
|---|---|---|---|
| POS-unpigm. | 14,59 | 1,61 | 1,29 |
| POS-pigm. | 14,06 | 2,09 | 1,88 |

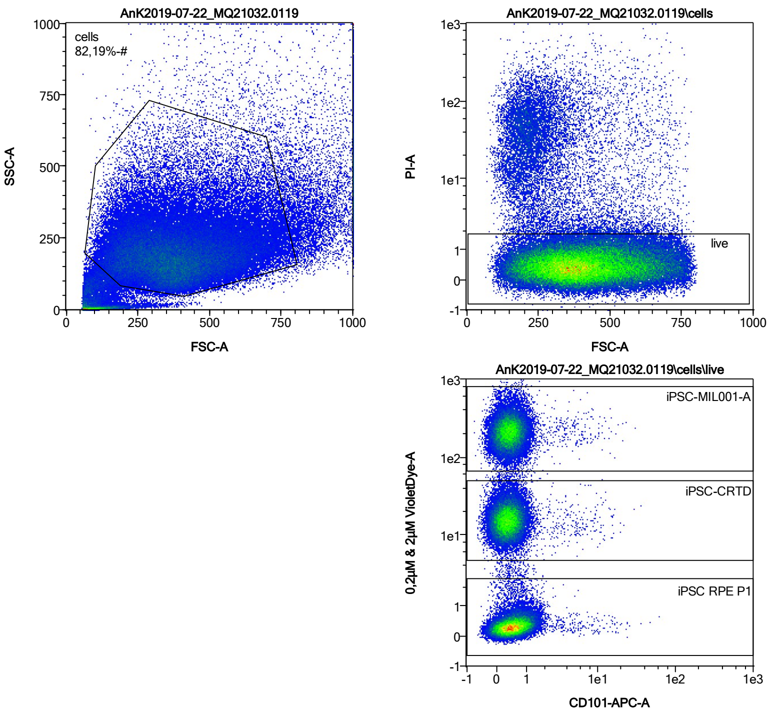


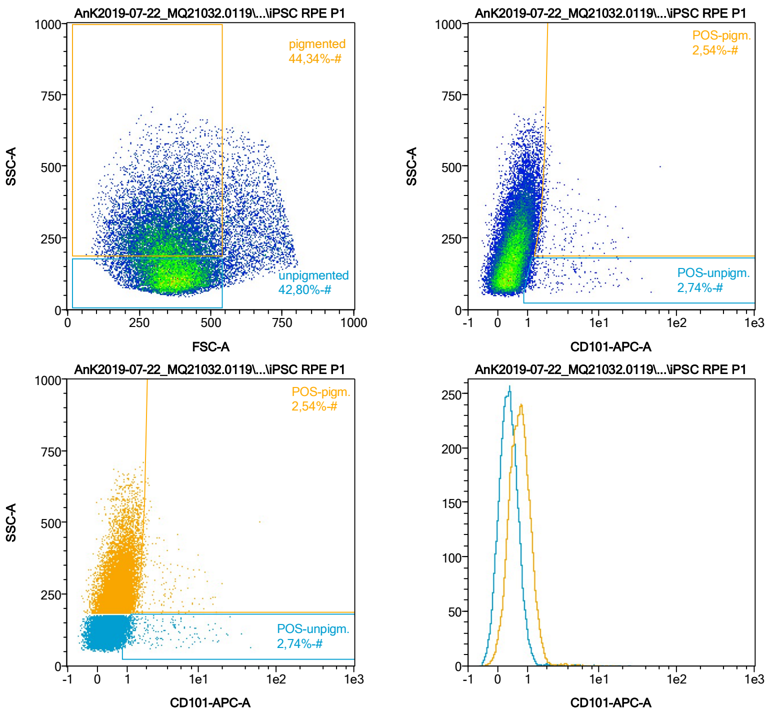


| Name | %-# | CD101-APC-A Mean | CD101-APC-A Median |
|---|---|---|---|
| POS-unpigm. | 2,74 | 3,31 | 1,31 |
| POS-pigm. | 2,54 | 3,25 | 1,89 |

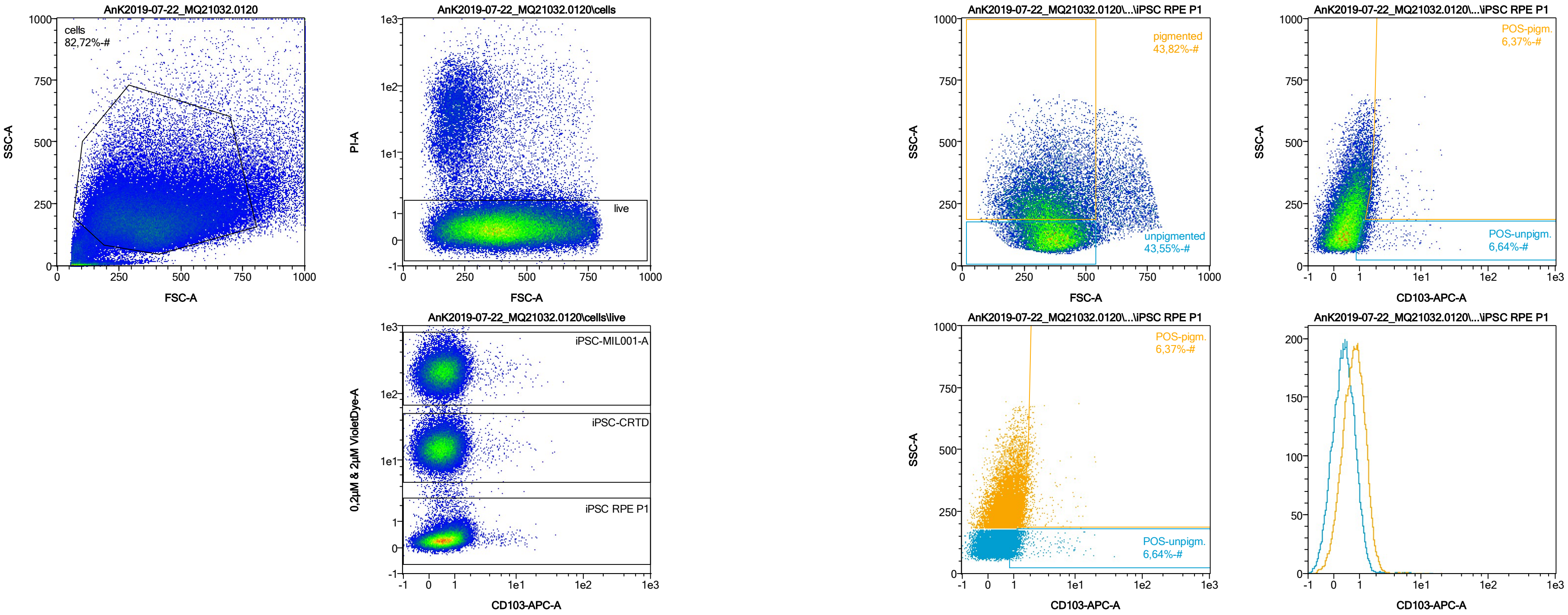


| Name | %-# | CD103-APC-A Mean | CD103-APC-A Median |
|---|---|---|---|
| POS-unpigm. | 6,64 | 1,78 | 1,20 |
| POS-pigm. | 6,37 | 2,21 | 1,80 |

Date:2024-06-14

Operator: Sebastian Knoebel

Version:2.13.0+54.g95abf0f4

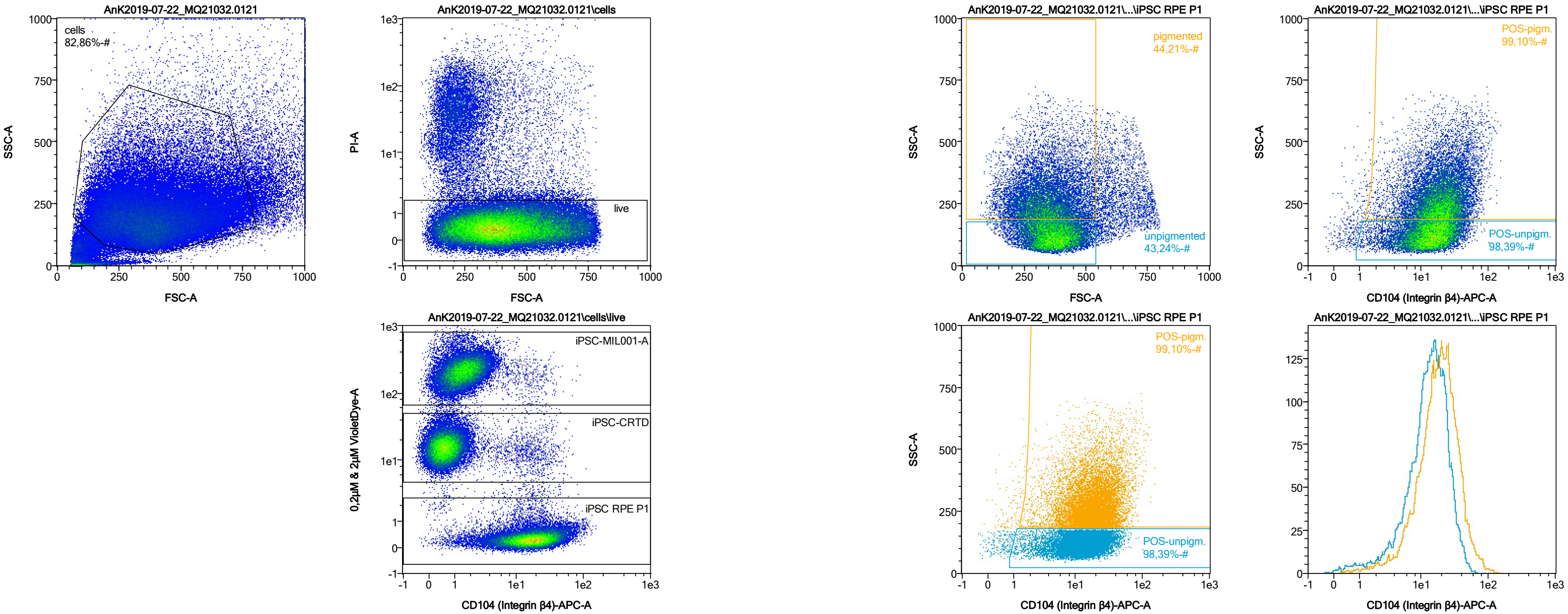


| Name | %-# | CD104 (Integrin β4)-APC-A Mean | CD104 (Integrin β4)-APC-A Median |
|---|---|---|---|
| POS-unpigm. | 98,39 | 15,17 | 13,69 |
| POS-pigm. | 99,10 | 21,60 | 18,69 |

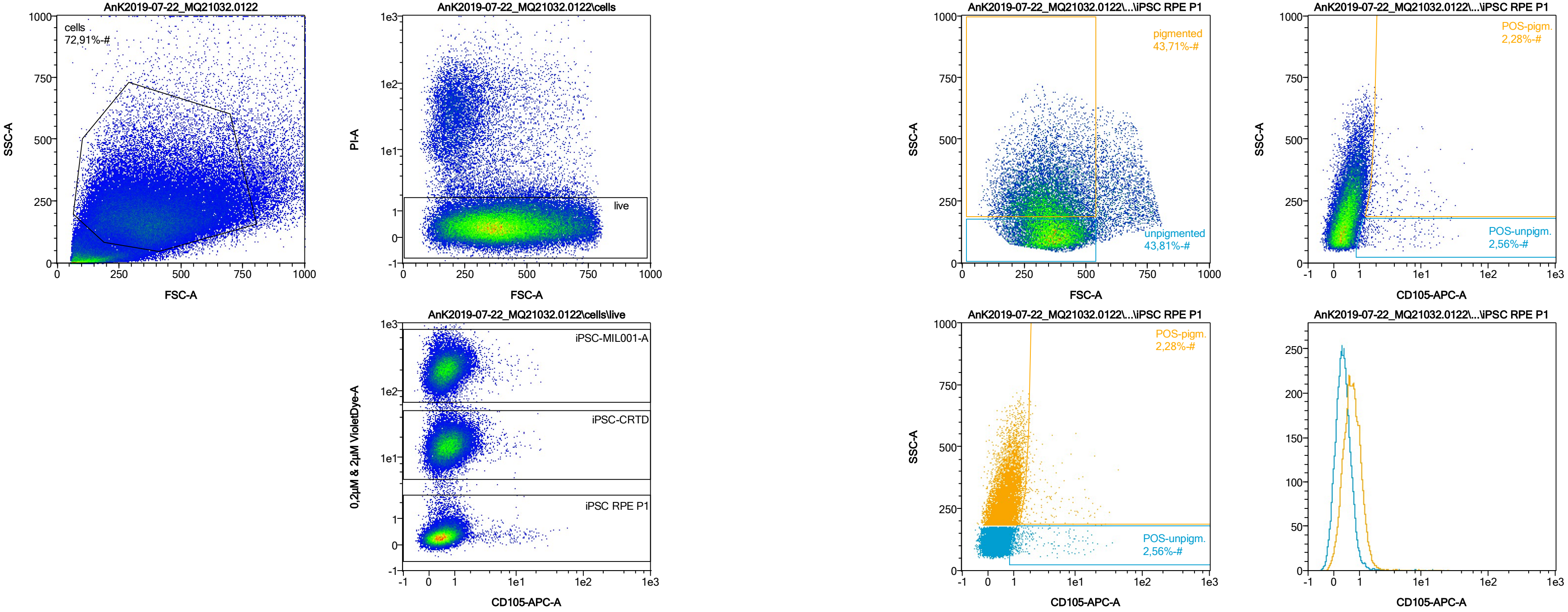


| Name | %-# | CD105-APC-A Mean | CD105-APC-A Median |
|---|---|---|---|
| POS-unpigm. | 2,56 | 3,51 | 1,44 |
| POS-pigm. | 2,28 | 4,73 | 2,15 |

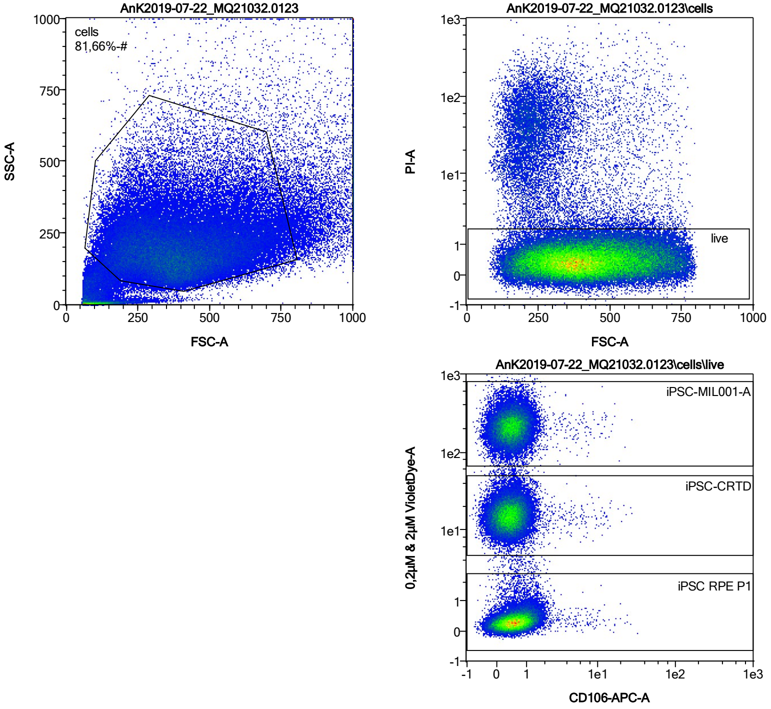


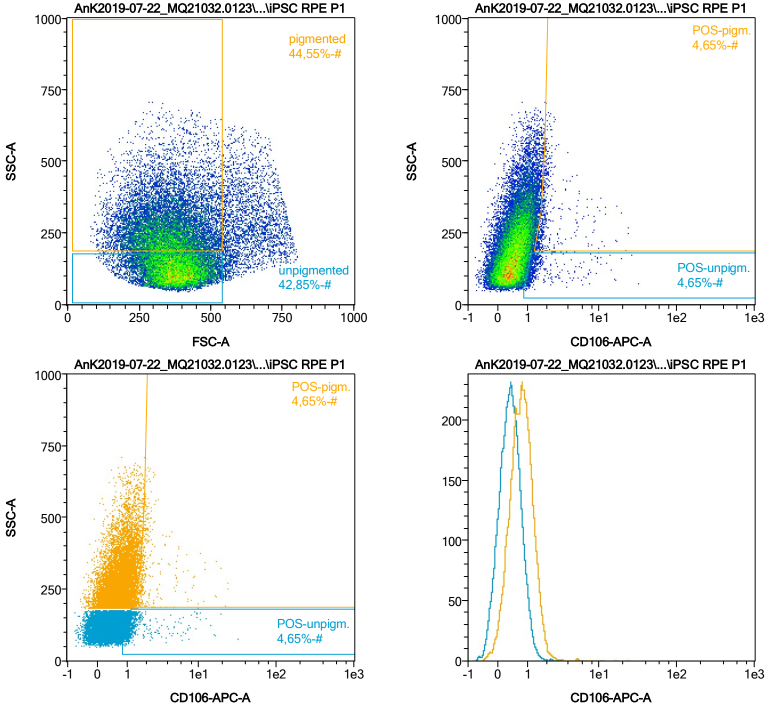


| Name | %-# | CD106-APC-A Mean | CD106-APC-A Median |
|---|---|---|---|
| POS-unpigm. | 4,65 | 1,83 | 1,19 |
| POS-pigm. | 4,65 | 2,43 | 1,80 |

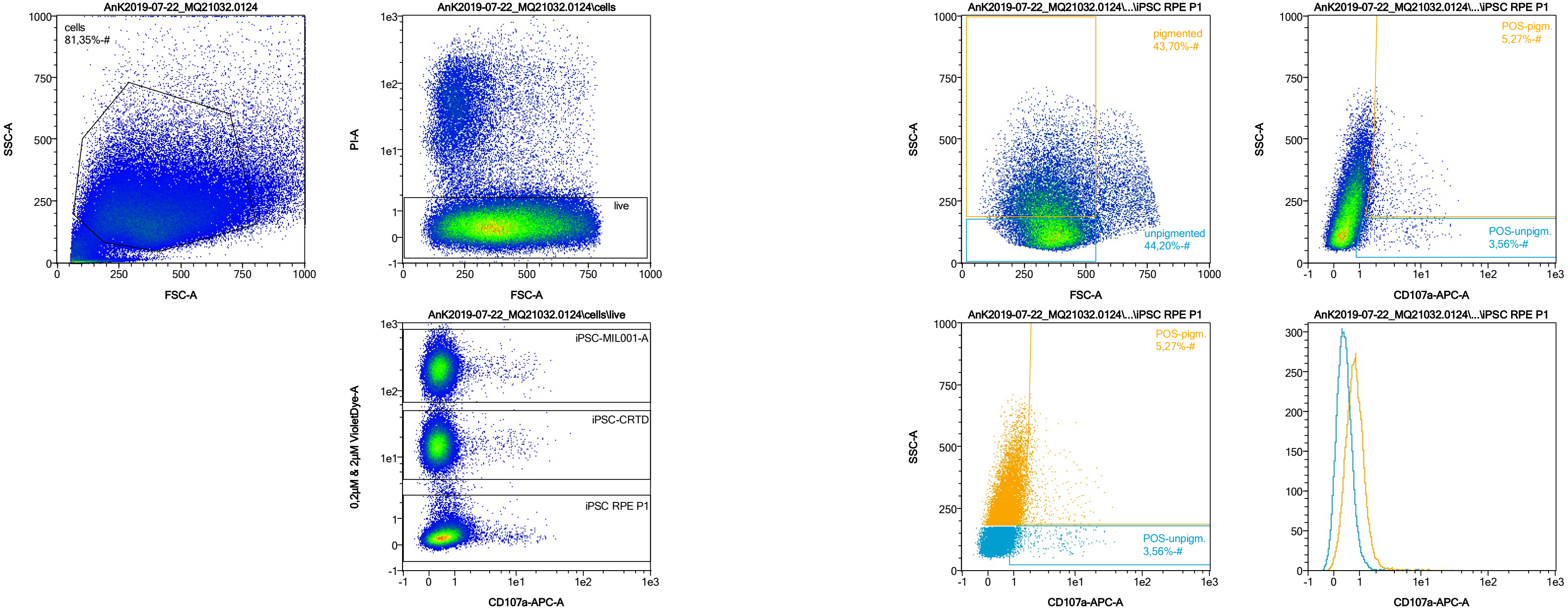


| Name | %-# | CD107a-APC-A Mean | CD107a-APC-A Median |
|---|---|---|---|
| POS-unpigm. | 3,56 | 3,39 | 1,35 |
| POS-pigm. | 5,27 | 3,31 | 2,00 |

Date:2024-06-14

Operator: Sebastian Knoebel

Version:2.13.0+54.g95abf0f4

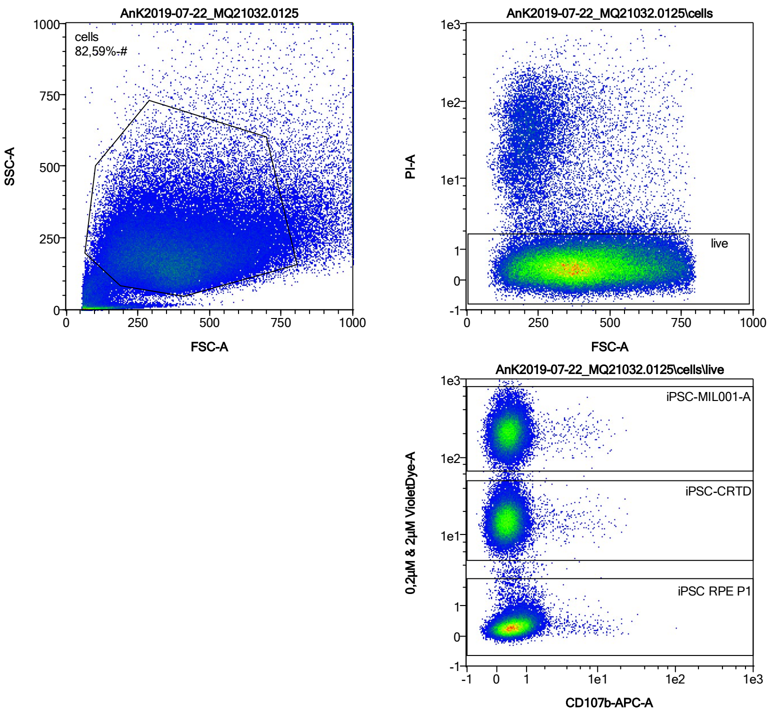


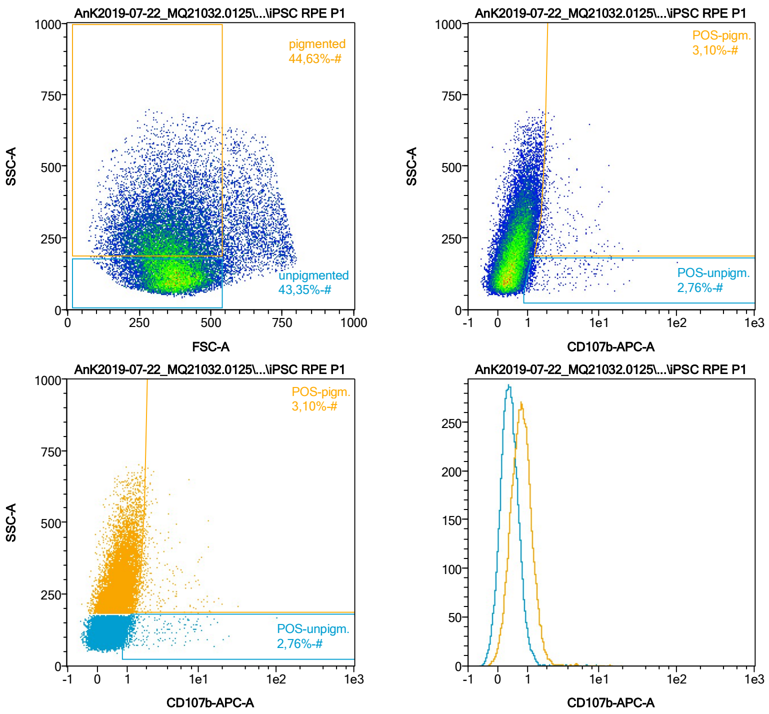


| Name | %-# | CD107b-APC-A Mean | CD107b-APC-A Median |
|---|---|---|---|
| POS-unpigm. | 2,76 | 2,95 | 1,28 |
| POS-pigm. | 3,10 | 3,05 | 1,95 |

Date:2024-06-14
Operator: Sebastian Knoebel
Version:2.13.0+54.g95abf0f4

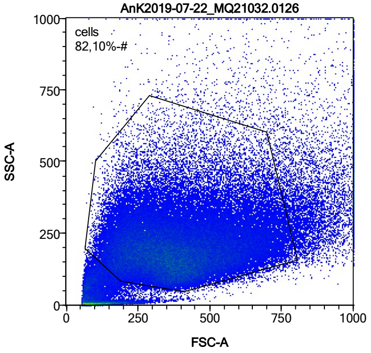


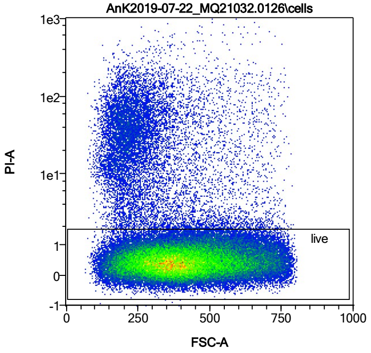


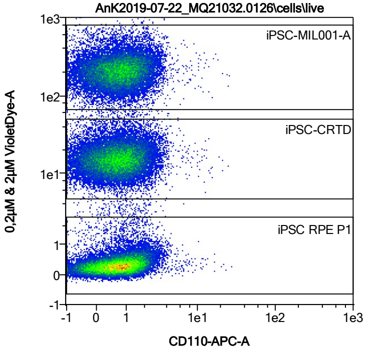


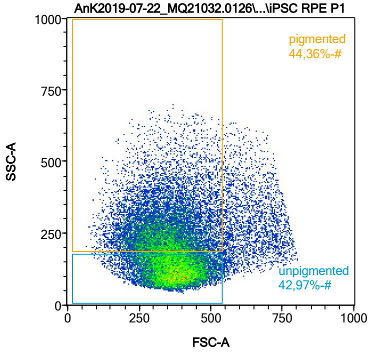


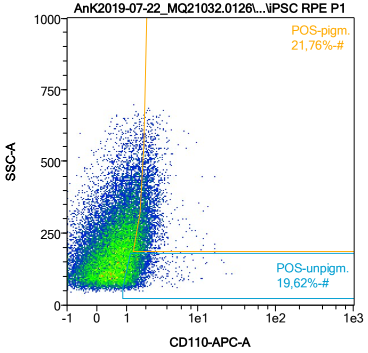


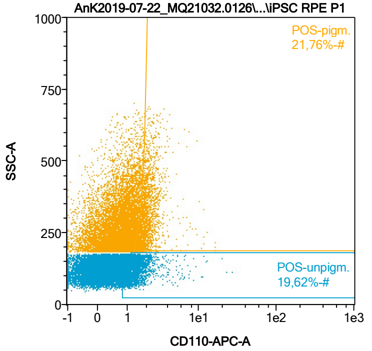


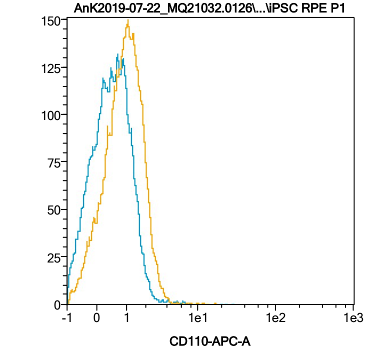


| Name | %-# | CD110-APC-A Mean | CD110-APC-A Median |
|---|---|---|---|
| POS-unpigm. | 19,62 | 1,58 | 1,33 |
| POS-pigm. | 21,76 | 2,11 | 1,91 |

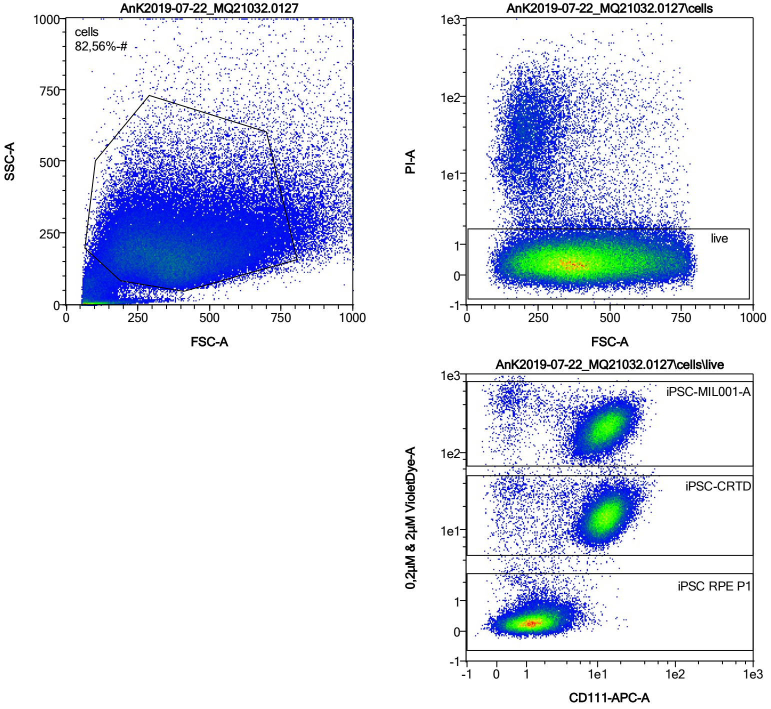


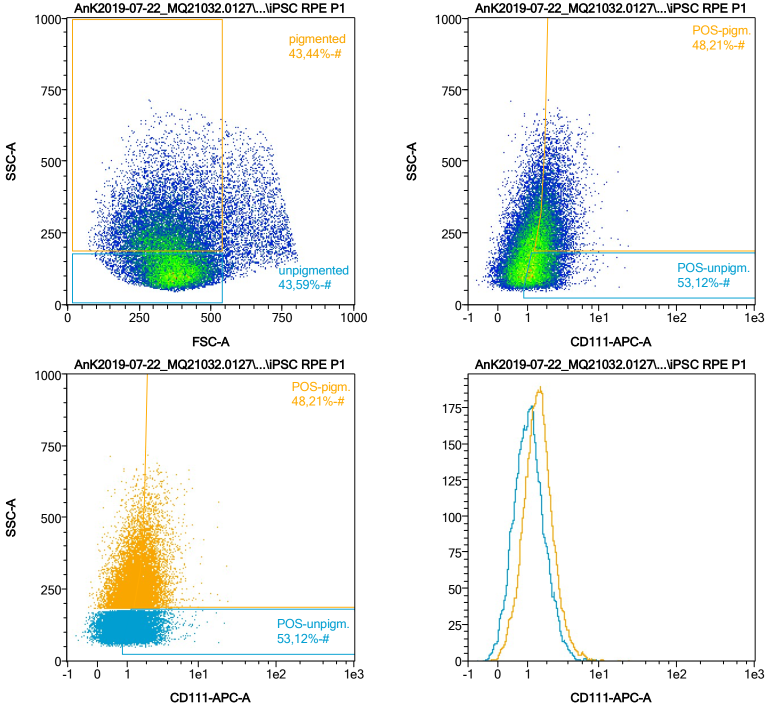


| Name | %-# | CD111-APC-A Mean | CD111-APC-A Median |
|---|---|---|---|
| POS-unpigm. | 53,12 | 1,73 | 1,50 |
| POS-pigm. | 48,21 | 2,33 | 2,04 |

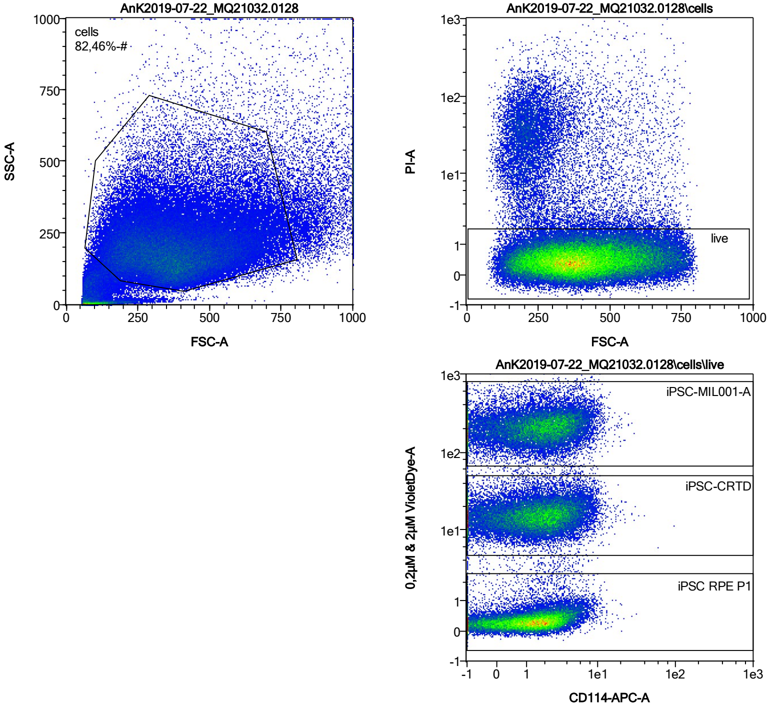


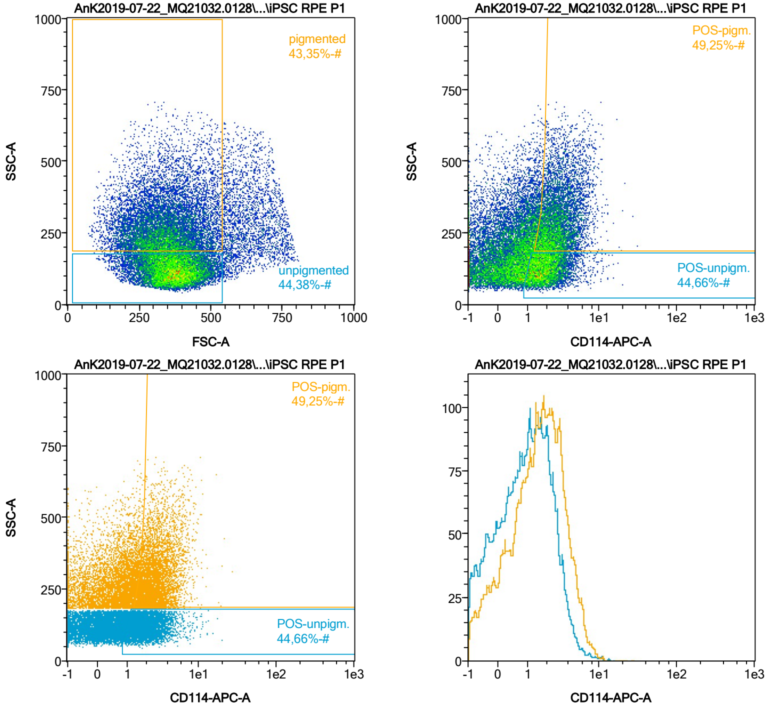


| Name | %-# | CD114-APC-A Mean | CD114-APC-A Median |
|---|---|---|---|
| POS-unpigm. | 44,66 | 2,15 | 1,84 |
| POS-pigm. | 49,25 | 3,02 | 2,62 |

Date:2024-06-14

Operator: Sebastian Knoebel

Version:2.13.0+54.g95abf0f4

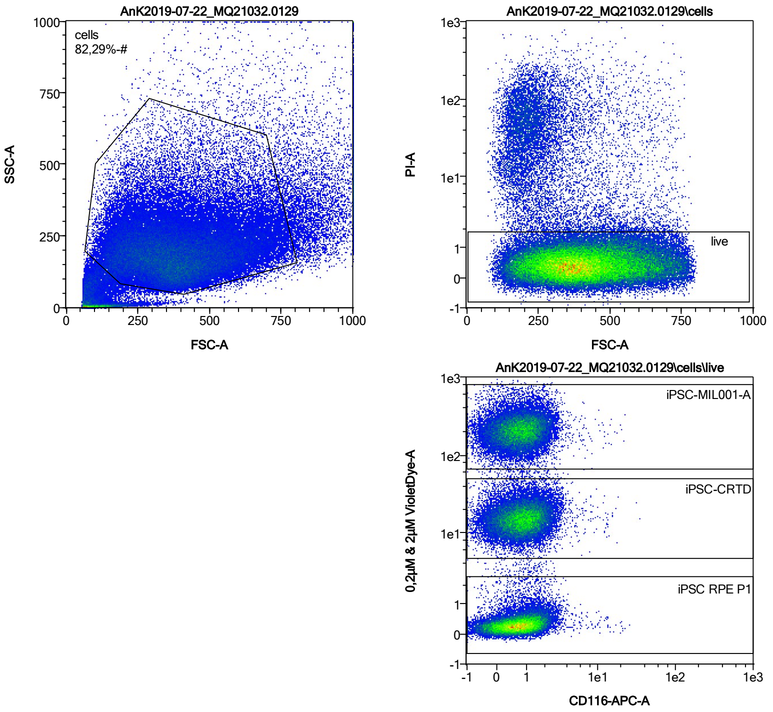


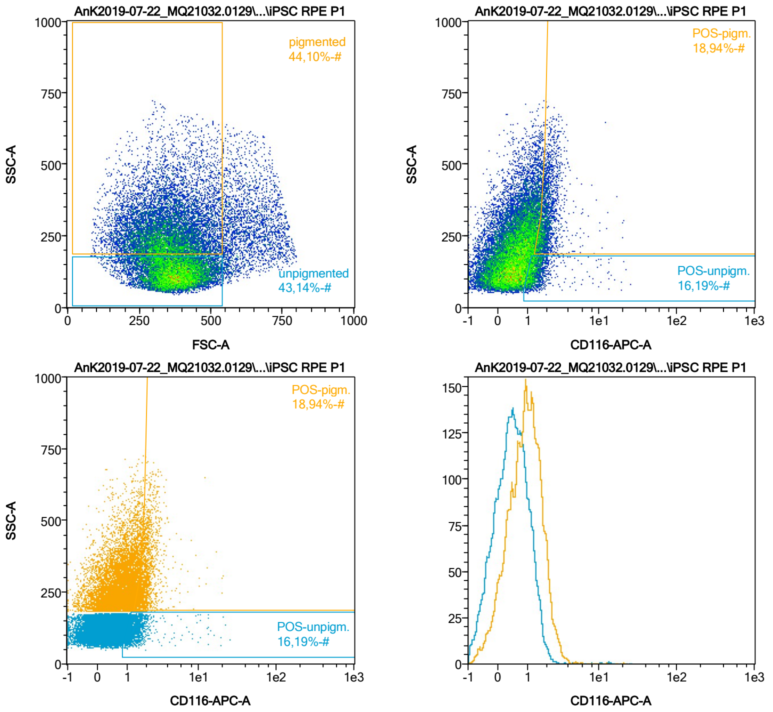


| Name | %-# | CD116-APC-A Mean | CD116-APC-A Median |
|---|---|---|---|
| POS-unpigm. | 16,19 | 1,56 | 1,27 |
| POS-pigm. | 18,94 | 2,08 | 1,88 |

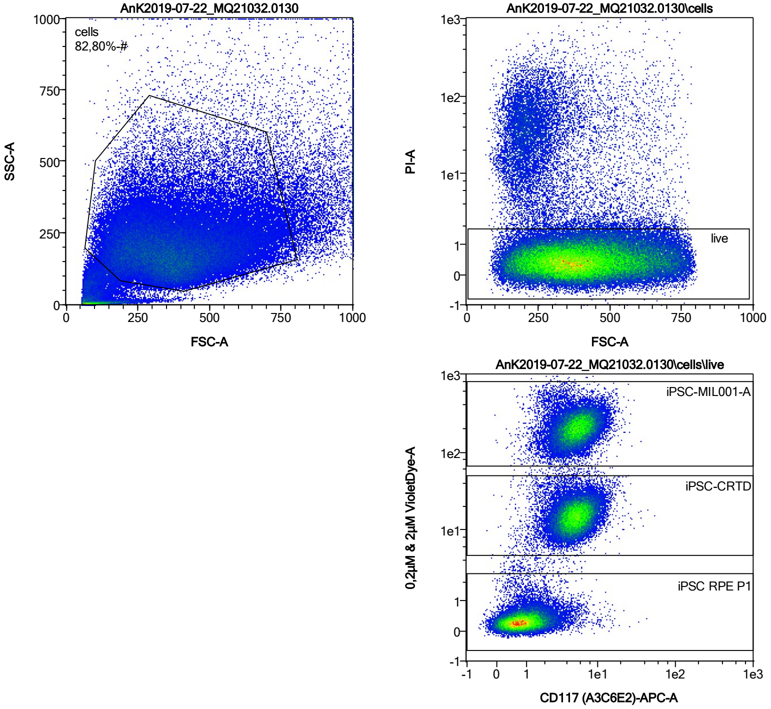


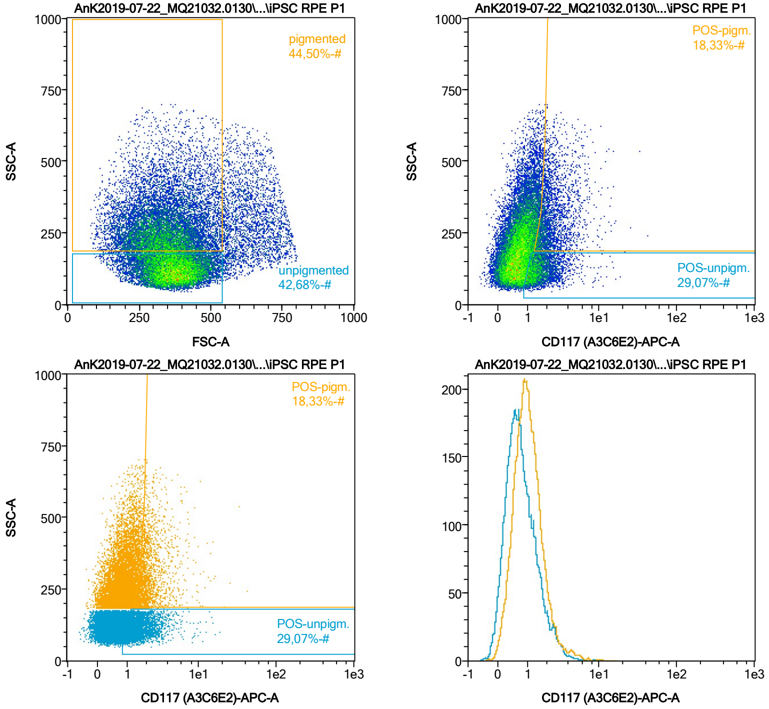


| Name | %-# | CD117 (A3C6E2)-APC-A Mean | CD117 (A3C6E2)-APC-A Median |
|---|---|---|---|
| POS-unpigm. | 29,07 | 1,74 | 1,45 |
| POS-pigm. | 18,33 | 2,46 | 1,98 |

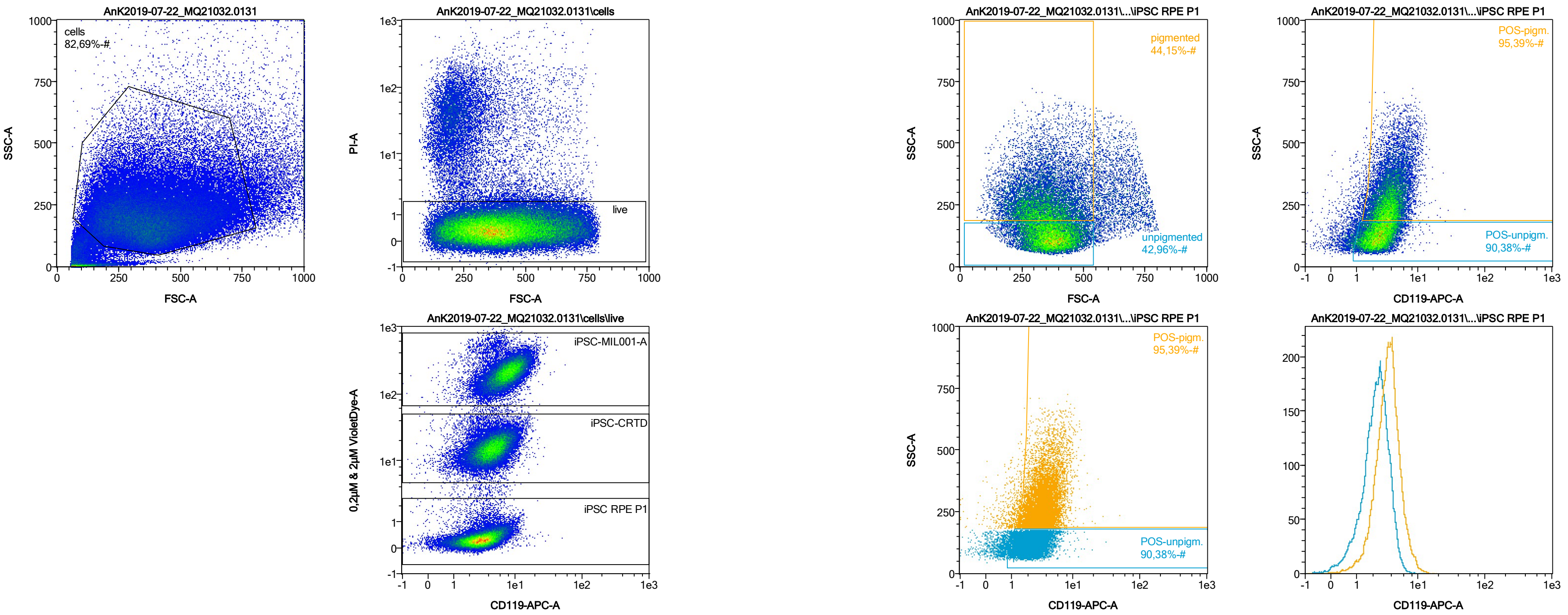


| Name | %-# | CD119-APC-A Mean | CD119-APC-A Median |
|---|---|---|---|
| POS-unpigm. | 90,38 | 2,59 | 2,43 |
| POS-pigm. | 95,39 | 3,85 | 3,56 |

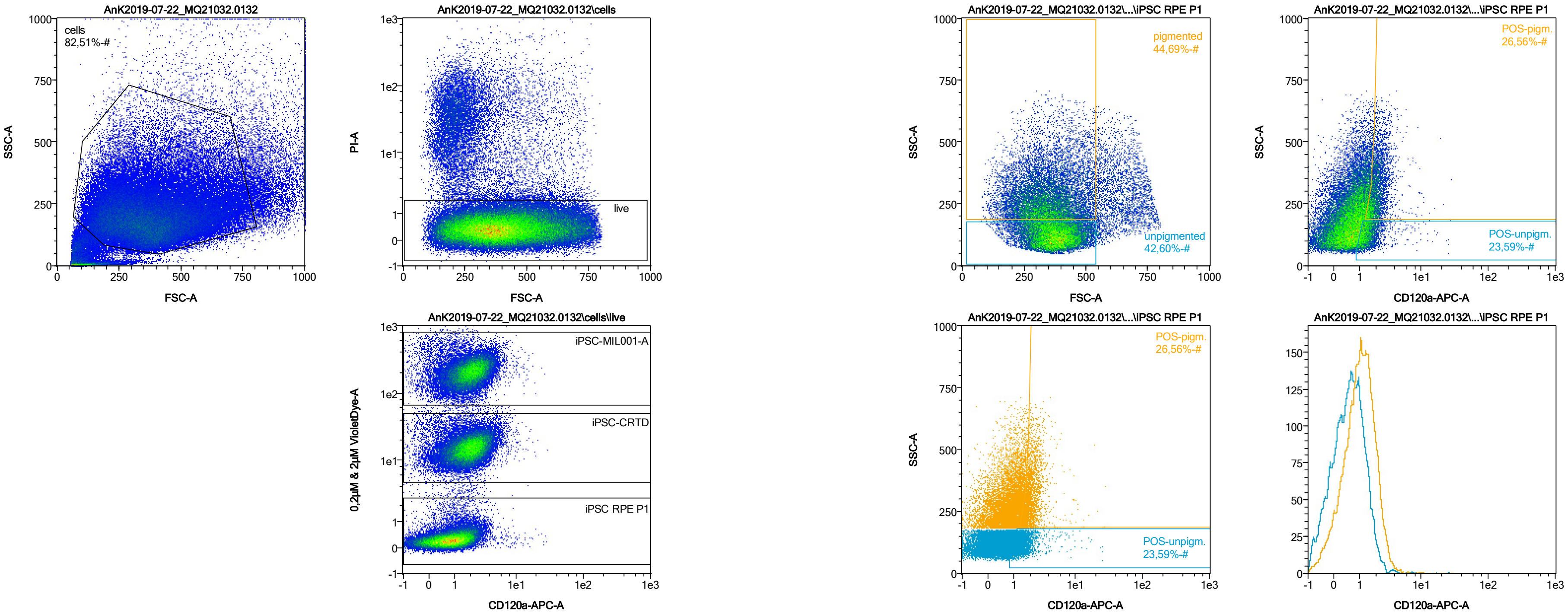


| Name | %-# | CD120a-APC-A Mean | CD120a-APC-A Median |
|---|---|---|---|
| POS-unpigm. | 23,59 | 1,60 | 1,36 |
| POS-pigm. | 26,56 | 2,15 | 1,94 |

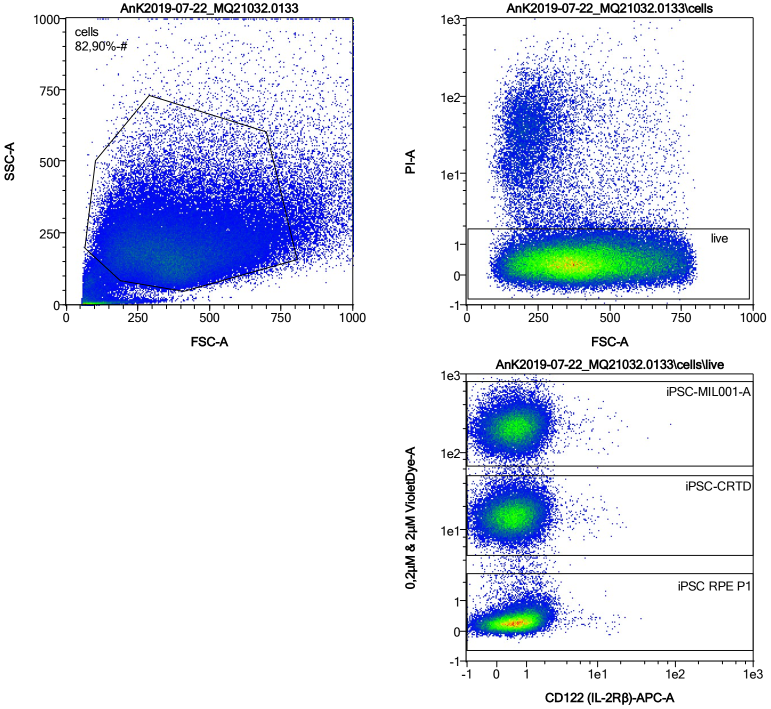


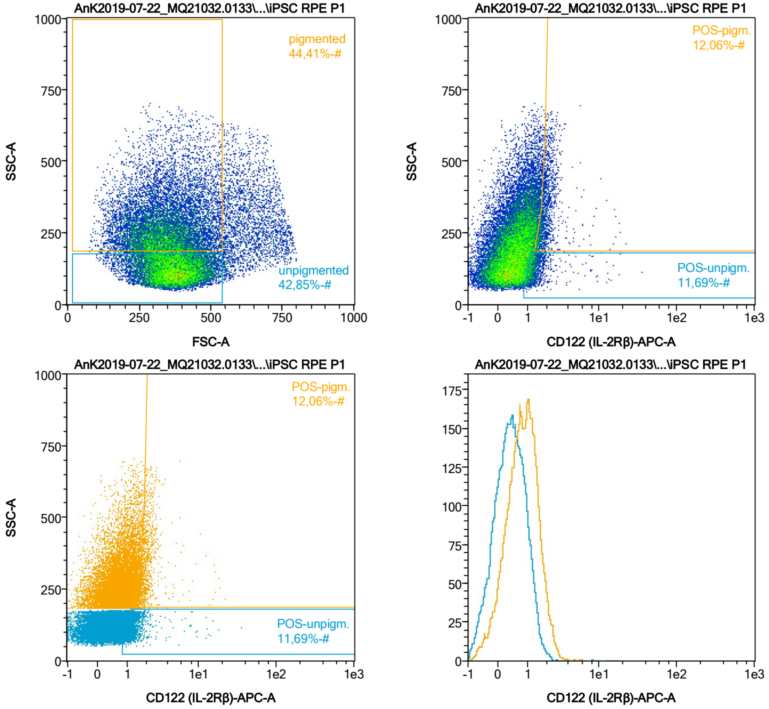


| Name | %-# | CD122 (IL-2Rβ)-APC-A Mean | CD122 (IL-2Rβ)-APC-A Median |
|---|---|---|---|
| POS-unpigm. | 11,69 | 1,48 | 1,24 |
| POS-pigm. | 12,06 | 2,07 | 1,84 |

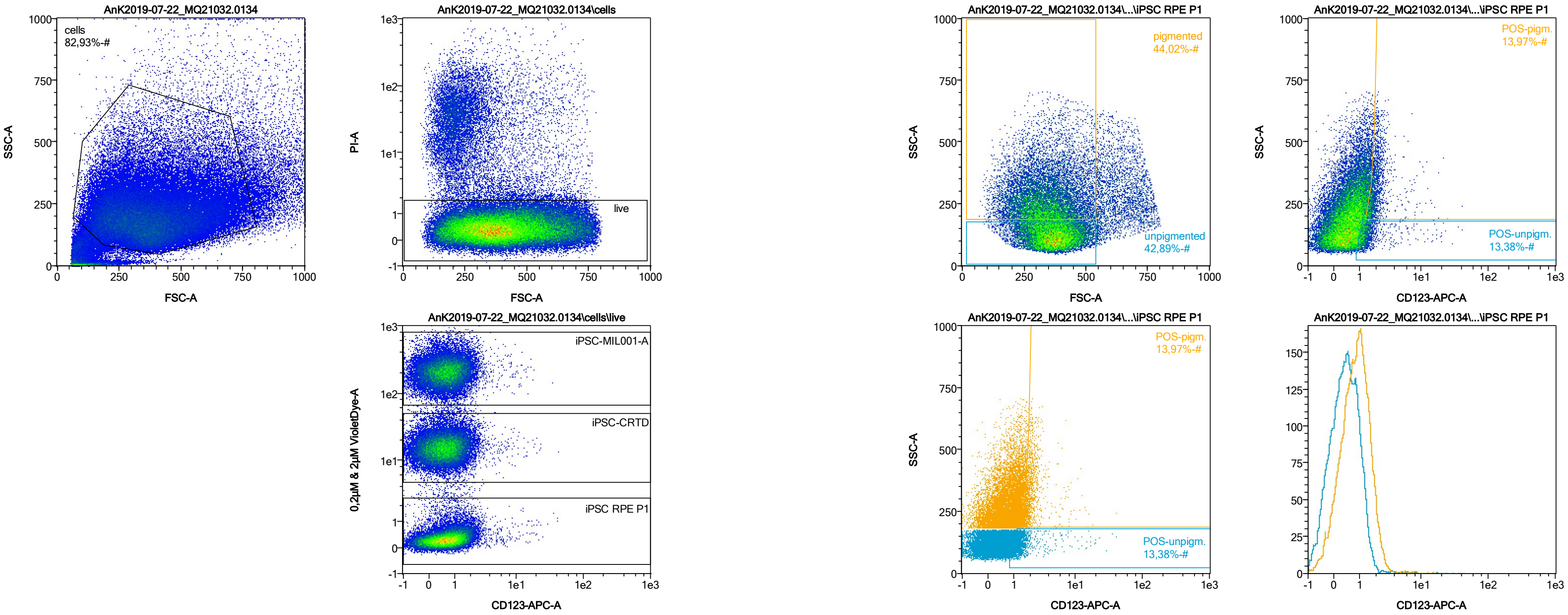


| Name | %-# | CD123-APC-A Mean | CD123-APC-A Median |
|---|---|---|---|
| POS-unpigm. | 13,38 | 1,67 | 1,25 |
| POS-pigm. | 13,97 | 2,15 | 1,86 |

Date:2024-06-14

Operator: Sebastian Knoebel

Version:2.13.0+54.g95abf0f4

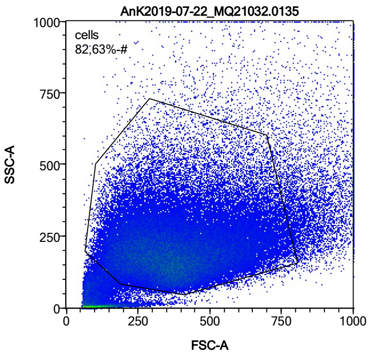


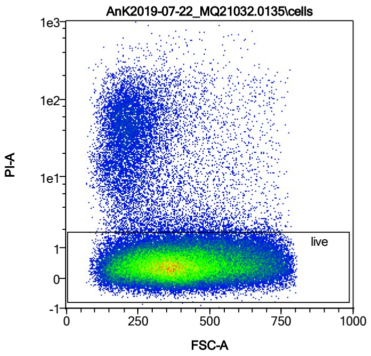


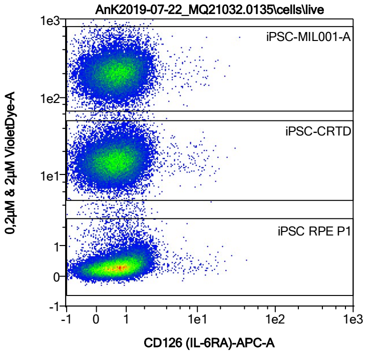


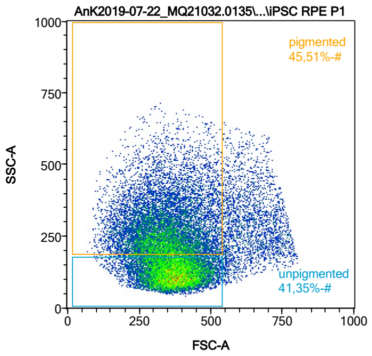


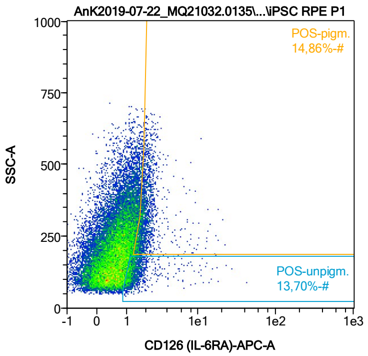


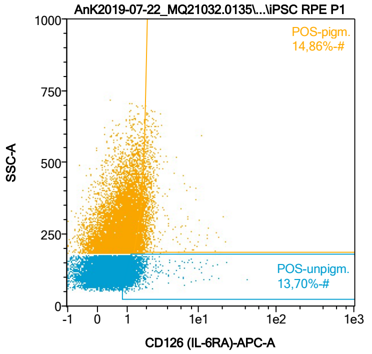


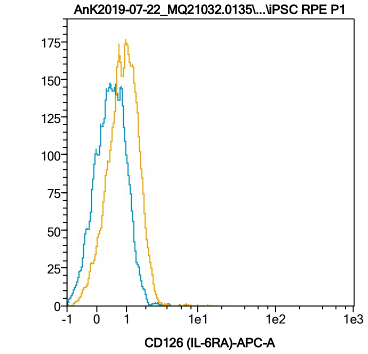


| Name | %-# | CD126 (IL-6RA)-APC-A Mean | CD126 (IL-6RA)-APC-A Median |
|---|---|---|---|
| POS-unpigm. | 13,70 | 1,56 | 1,24 |
| POS-pigm. | 14,86 | 2,11 | 1,86 |

Date:2024-06-14

Operator: Sebastian Knoebel

Version:2.13.0+54.g95abf0f4

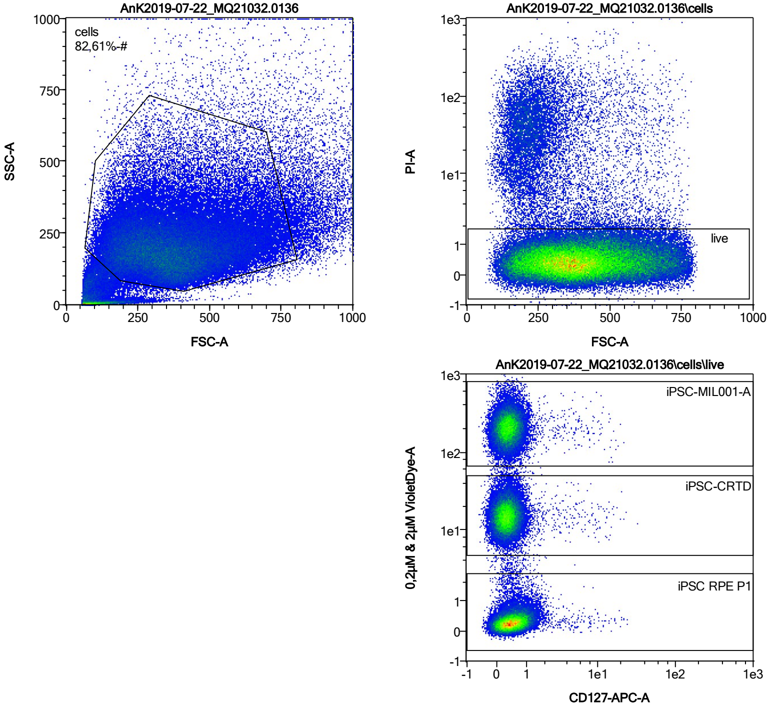


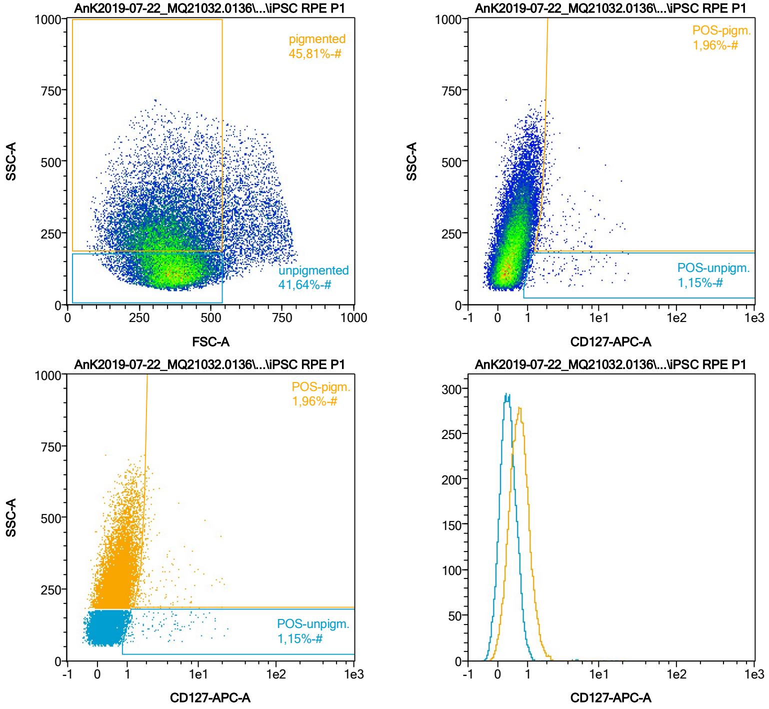


| Name | %-# | CD127-APC-A Mean | CD127-APC-A Median |
|---|---|---|---|
| POS-unpigm. | 1,15 | 4,00 | 2,22 |
| POS-pigm. | 1,96 | 3,20 | 1,99 |

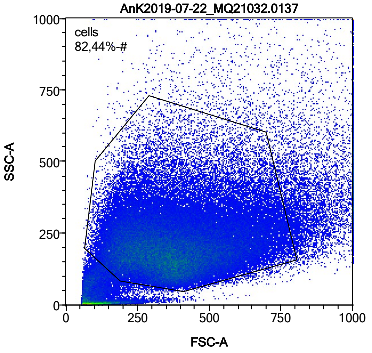


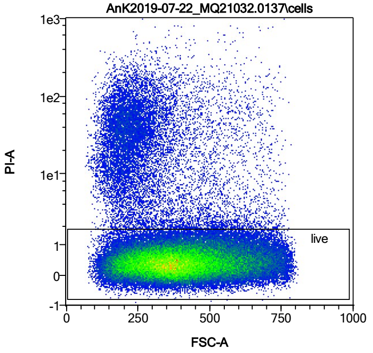


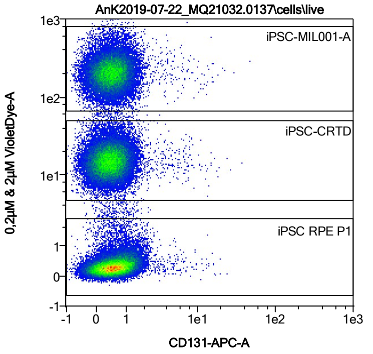


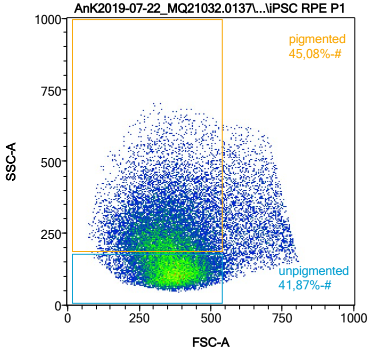


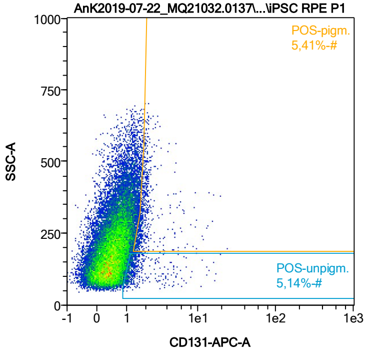


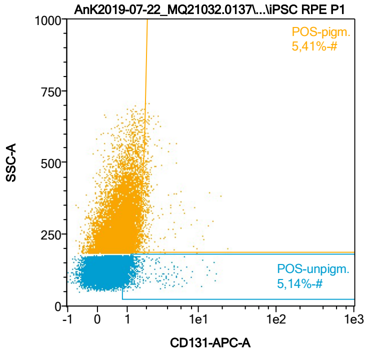


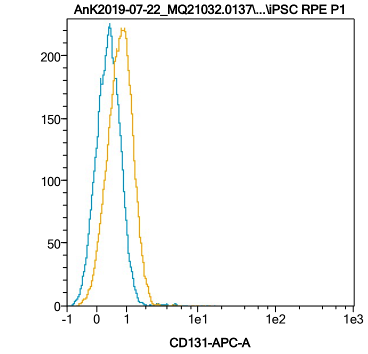


| Name | %-# | CD131-APC-A Mean | CD131-APC-A Median |
|---|---|---|---|
| POS-unpigm. | 5,14 | 1,72 | 1,22 |
| POS-pigm. | 5,41 | 2,17 | 1,80 |

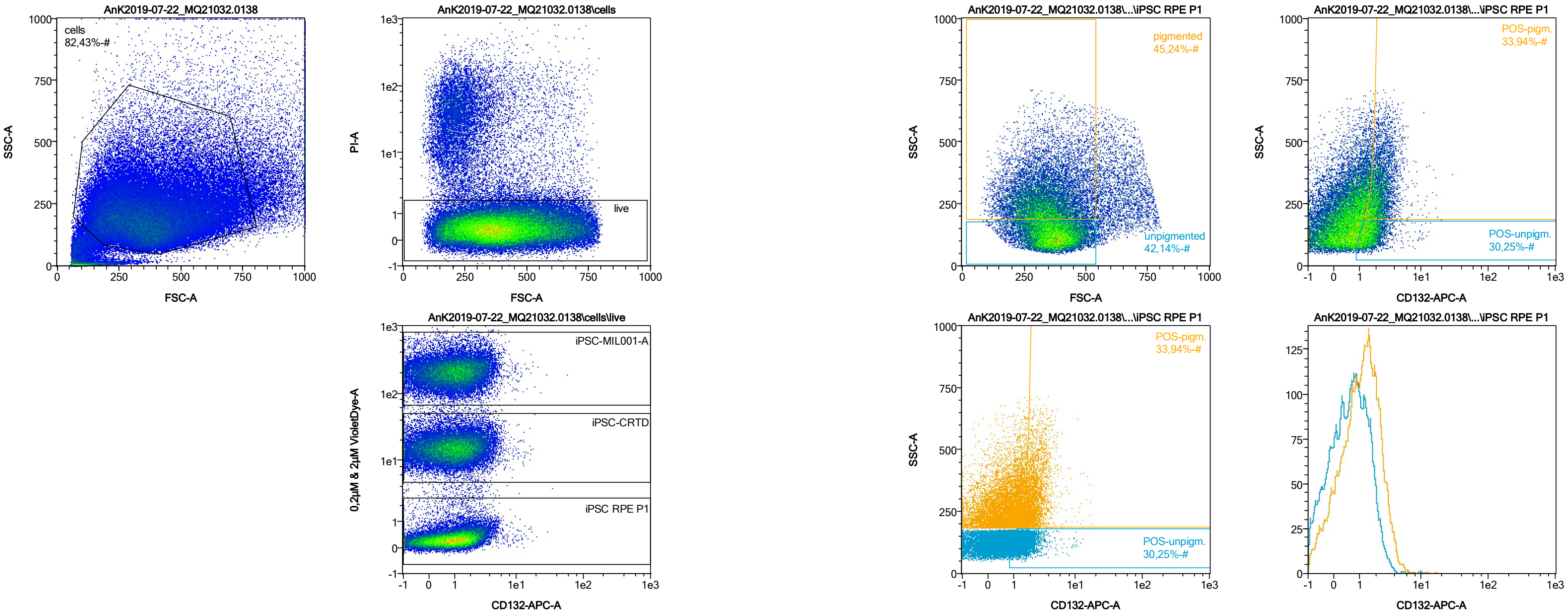


| Name | %-# | CD132-APC-A Mean | CD132-APC-A Median |
|---|---|---|---|
| POS-unpigm. | 30,25 | 1,69 | 1,50 |
| POS-pigm. | 33,94 | 2,37 | 2,11 |

Date:2024-06-14
Operator: Sebastian Knoebel
Version:2.13.0+54.g95abf0f4

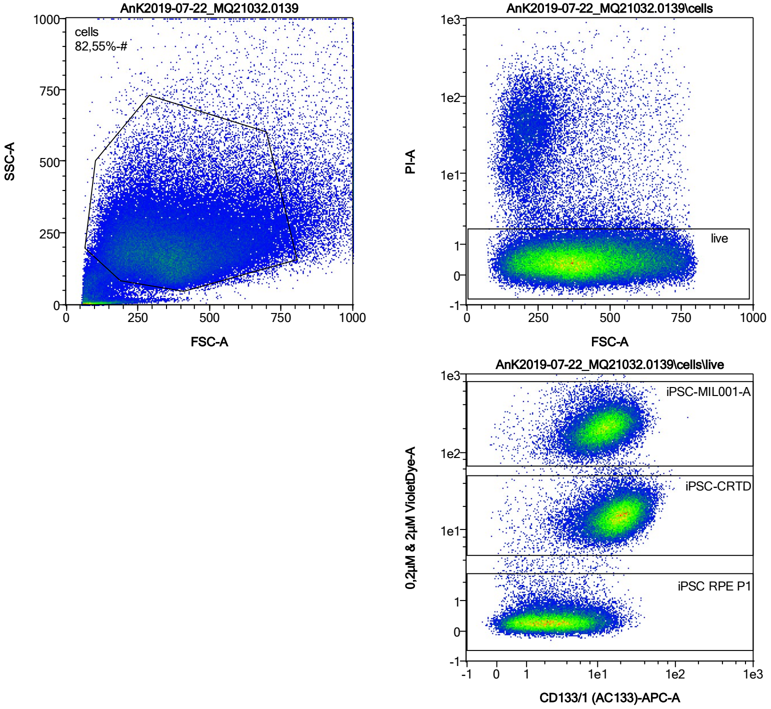


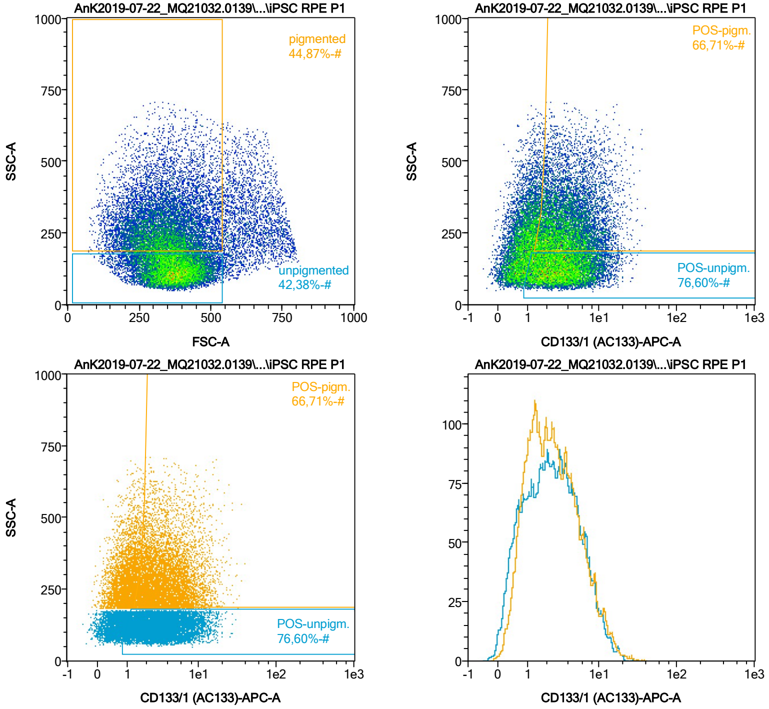


| Name | %-# | CD133/1 (AC133)-APC-A Mean | CD133/1 (AC133)-APC-A Median |
|---|---|---|---|
| POS-unpigm. | 76,60 | 3,90 | 3,03 |
| POS-pigm. | 66,71 | 4,40 | 3,36 |

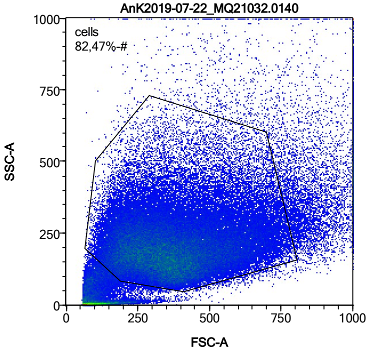


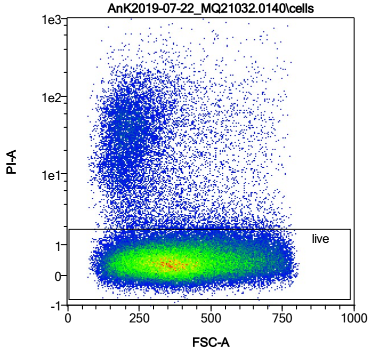


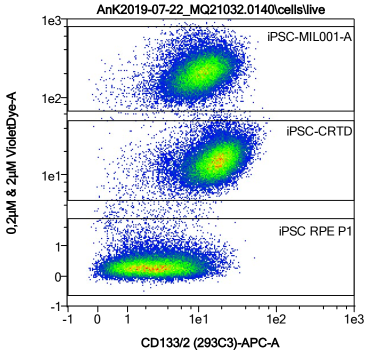


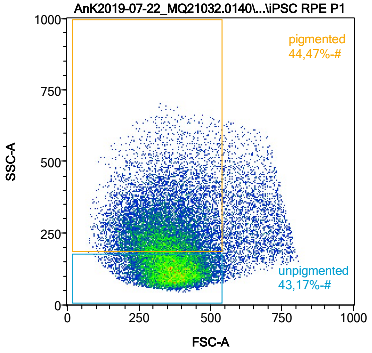


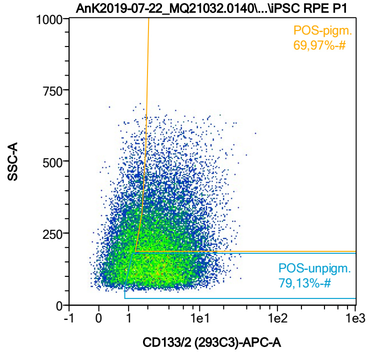


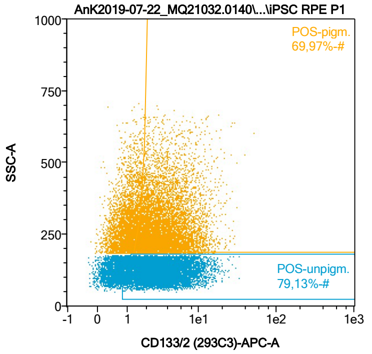


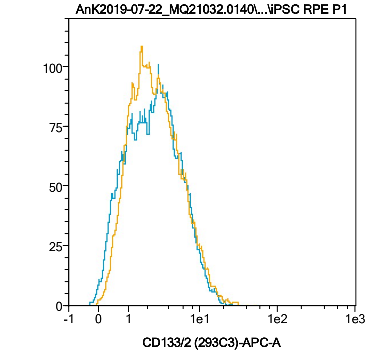


| Name | %-# | CD133/2 (293C3)-APC-A Mean | CD133/2 (293C3)-APC-A Median |
|---|---|---|---|
| POS-unpigm. | 79,13 | 3,88 | 3,05 |
| POS-pigm. | 69,97 | 4,36 | 3,32 |

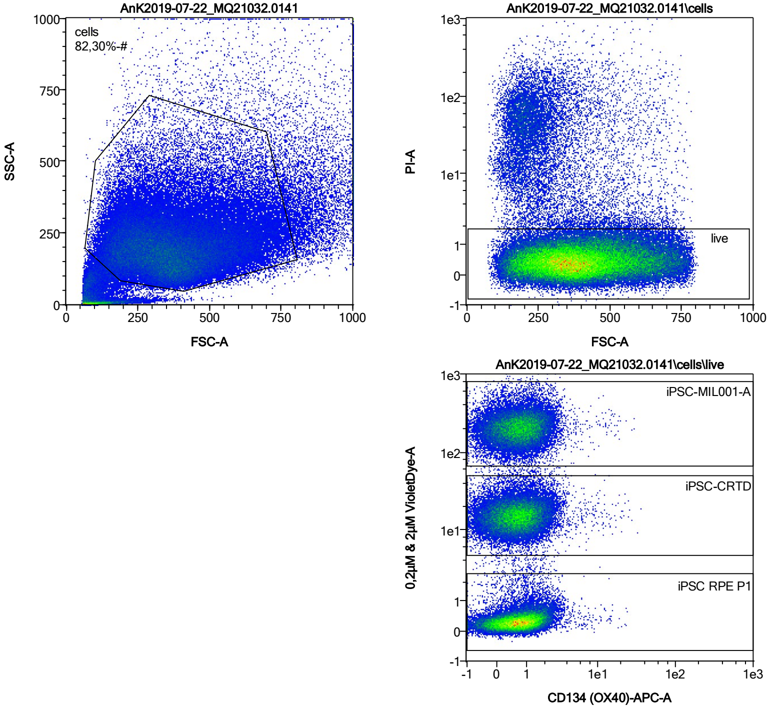


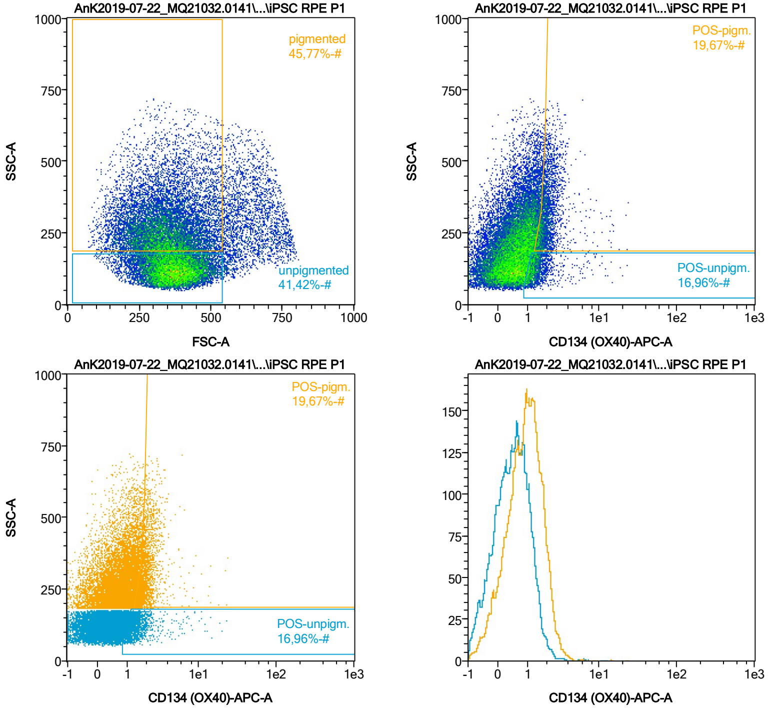


| Name | %-# | CD134 (OX40)-APC-A Mean | CD134 (OX40)-APC-A Median |
|---|---|---|---|
| POS-unpigm. | 16,96 | 1,55 | 1,28 |
| POS-pigm. | 19,67 | 2,12 | 1,90 |

Date:2024-06-14

Operator: Sebastian Knoebel

Version:2.13.0+54.g95abf0f4

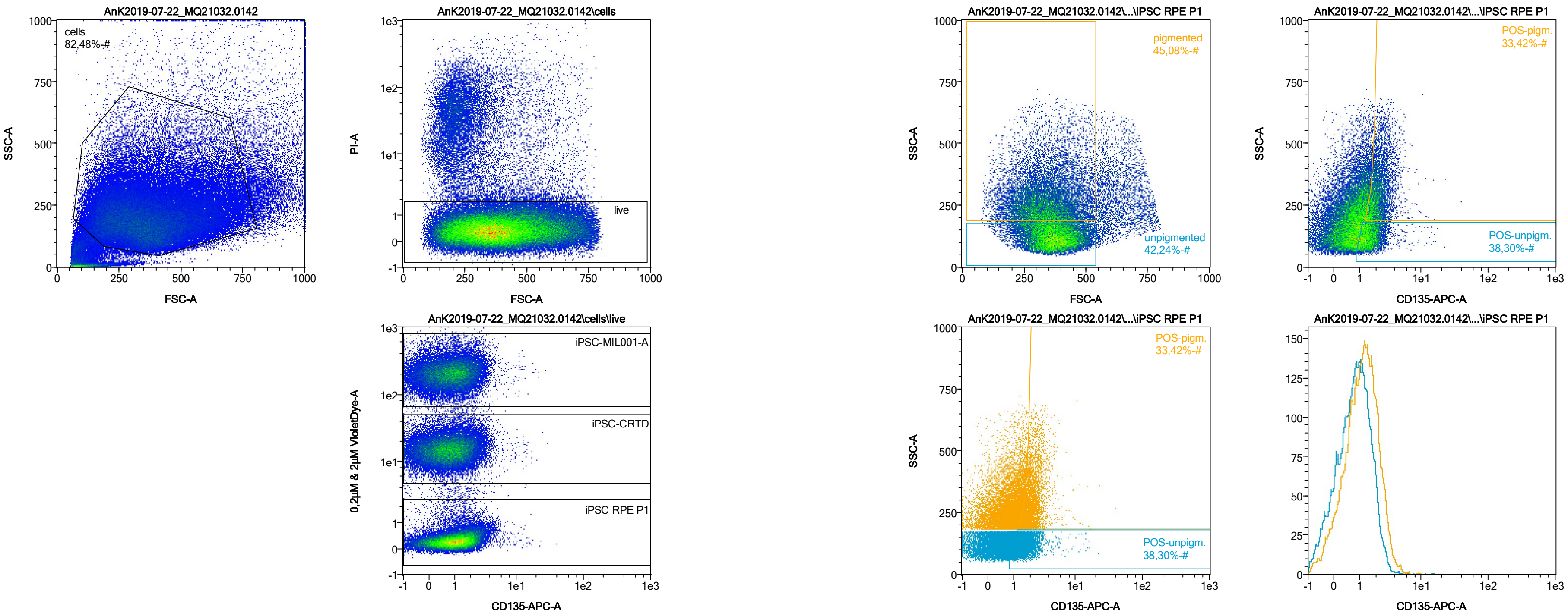


| Name | %-# | CD135-APC-A Mean | CD135-APC-A Median |
|---|---|---|---|
| POS-unpigm. | 38,30 | 1,67 | 1,48 |
| POS-pigm. | 33,42 | 2,25 | 2,04 |

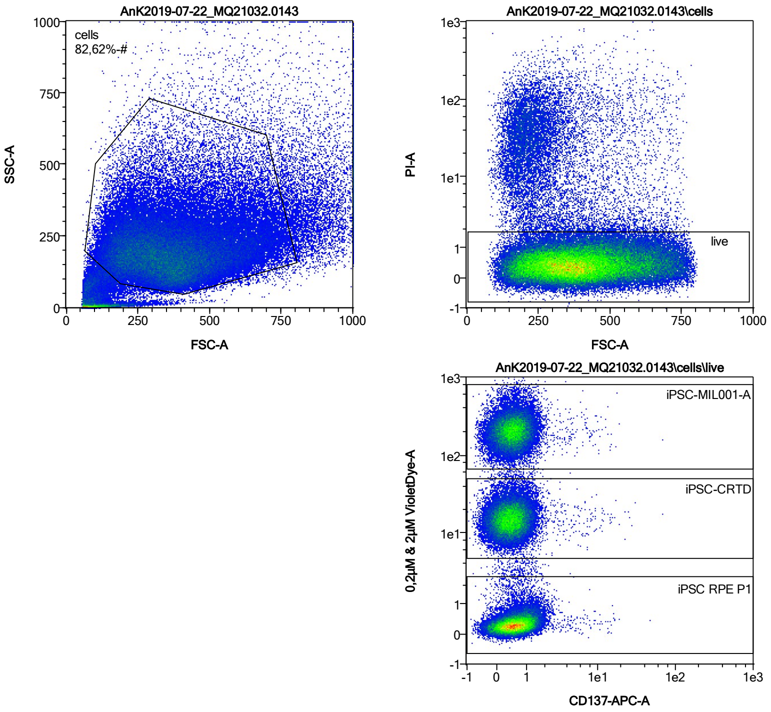


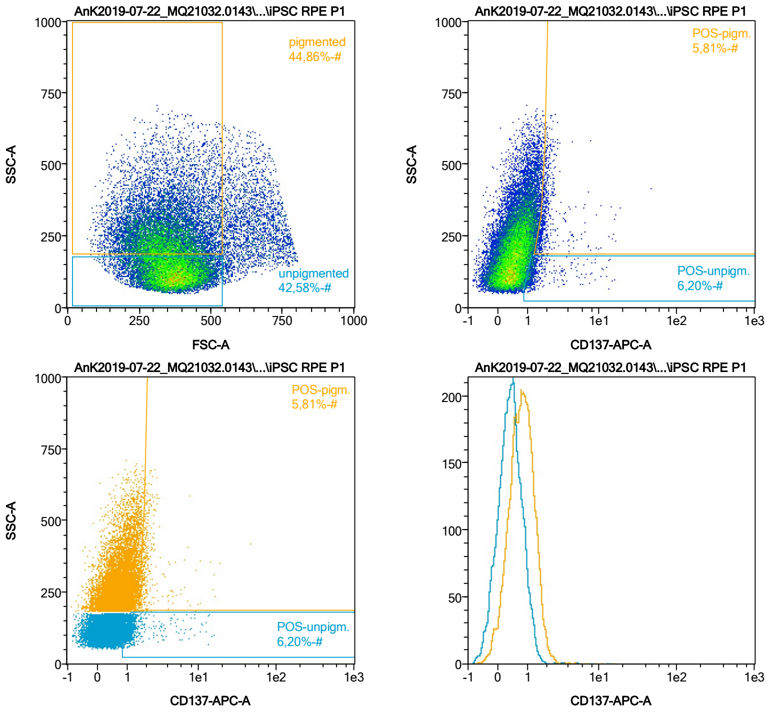


| Name | %-# | CD137-APC-A Mean | CD137-APC-A Median |
|---|---|---|---|
| POS-unpigm. | 6,20 | 1,65 | 1,19 |
| POS-pigm. | 5,81 | 2,19 | 1,78 |

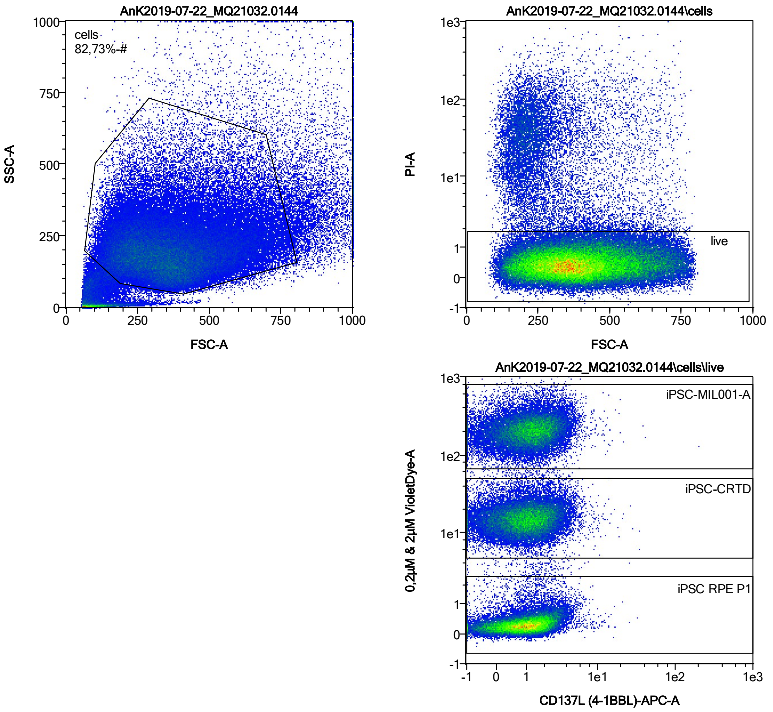


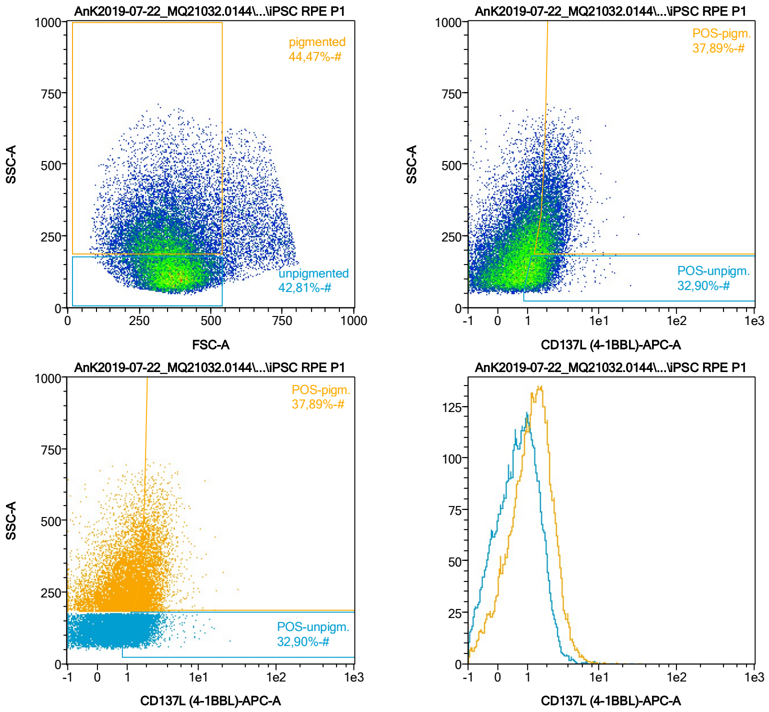


| Name | %-# | CD137L (4-1BBL)-APC-A Mean | CD137L (4-1BBL)-APC-A Median |
|---|---|---|---|
| POS-unpigm. | 32,90 | 1,69 | 1,48 |
| POS-pigm. | 37,89 | 2,41 | 2,12 |

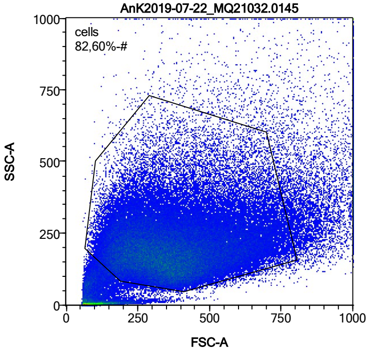


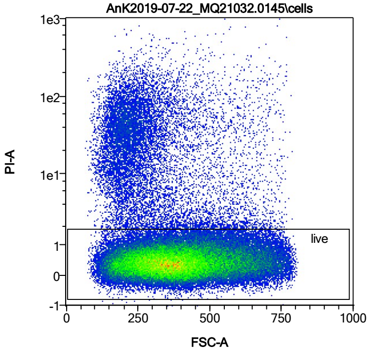


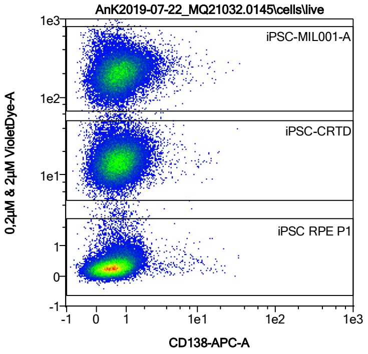


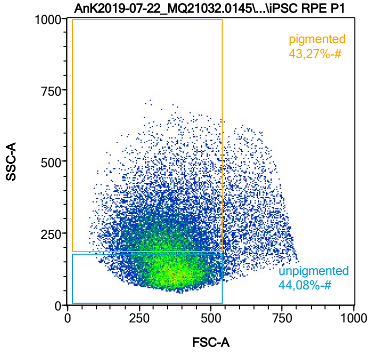


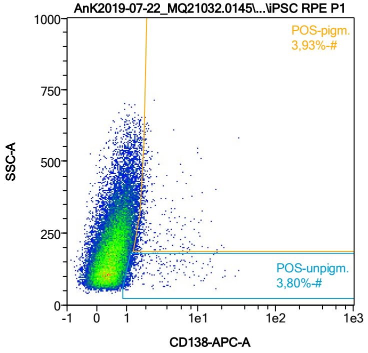


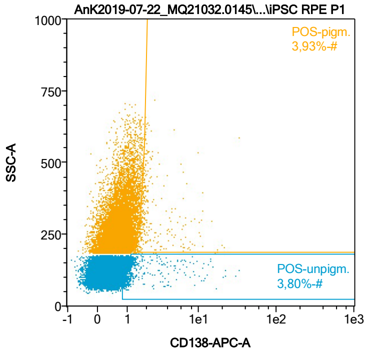


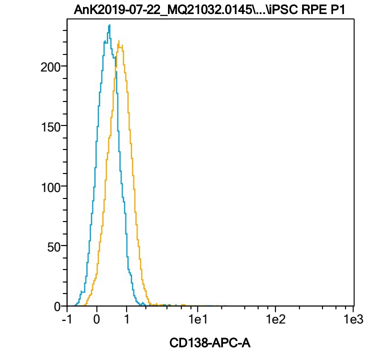


| Name | %-# | CD138-APC-A Mean | CD138-APC-A Median |
|---|---|---|---|
| POS-unpigm. | 3,80 | 2,37 | 1,21 |
| POS-pigm. | 3,93 | 2,74 | 1,85 |

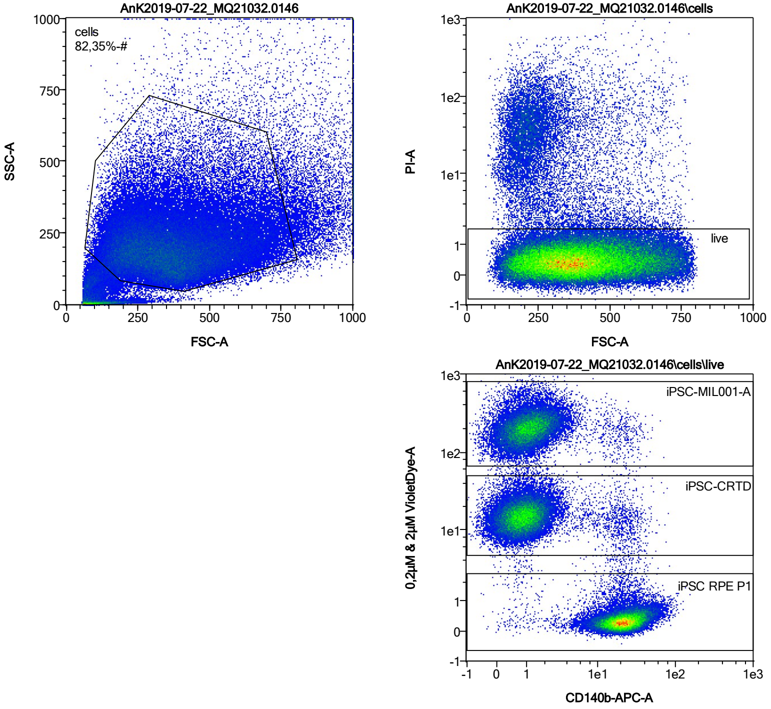


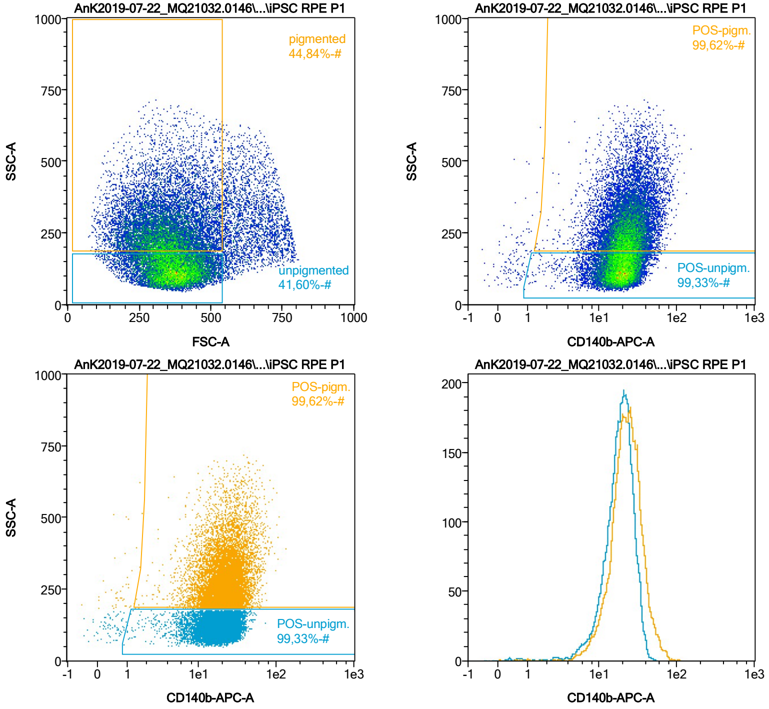


| Name | %-# | CD140b-APC-A Mean | CD140b-APC-A Median |
|---|---|---|---|
| POS-unpigm. | 99,33 | 19,26 | 18,83 |
| POS-pigm. | 99,62 | 24,40 | 22,55 |

Date:2024-06-14
Operator: Sebastian Knoebel
Version:2.13.0+54.g95abf0f4

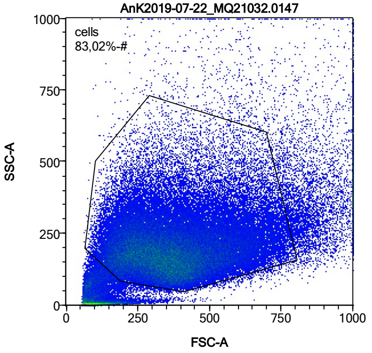


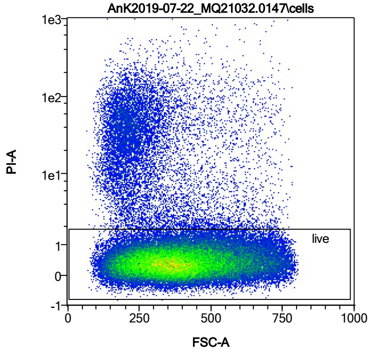


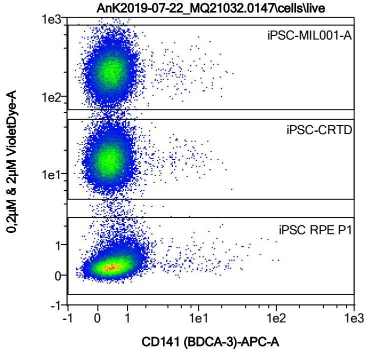


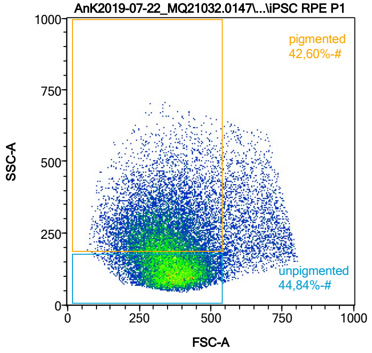


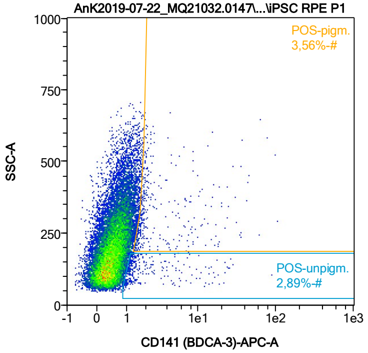


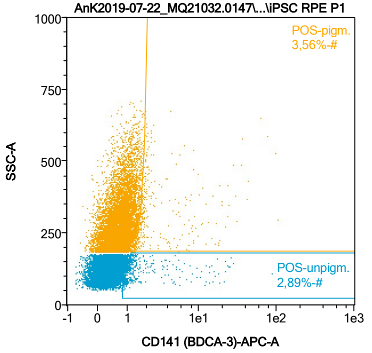


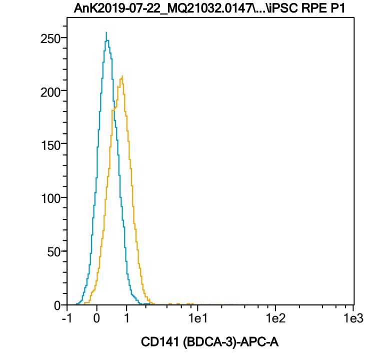


| Name | %-# | CD141 (BDCA-3)-APC-A Mean | CD141 (BDCA-3)-APC-A Median |
|---|---|---|---|
| POS-unpigm. | 2,89 | 3,26 | 1,25 |
| POS-pigm. | 3,56 | 4,92 | 1,88 |

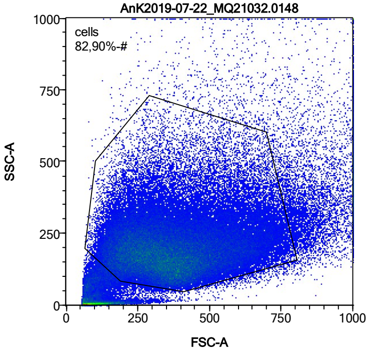


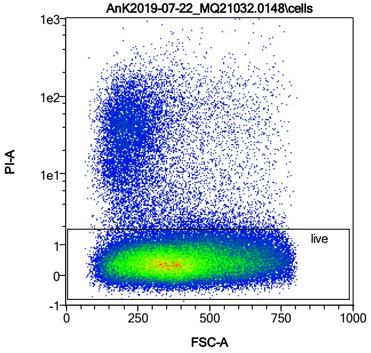


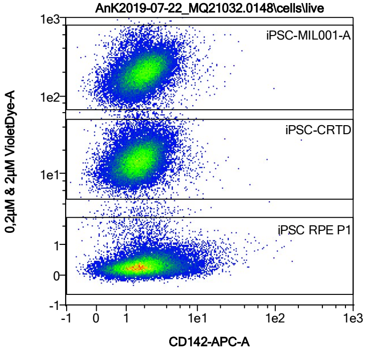


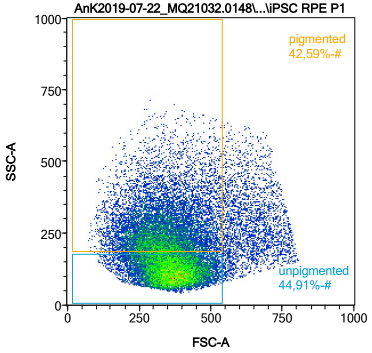


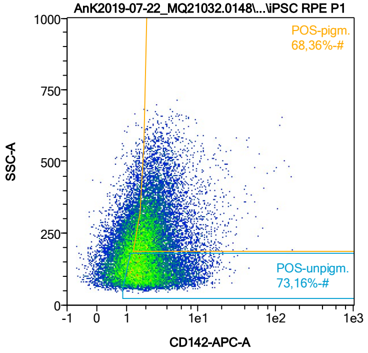


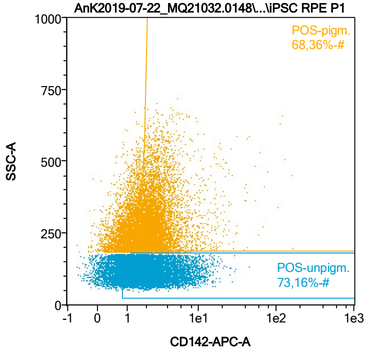


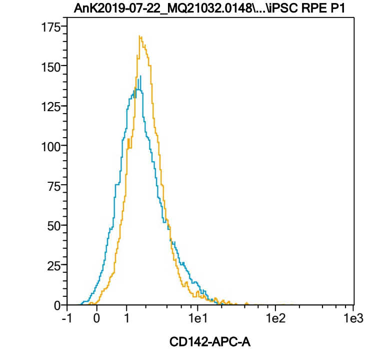


| Name | %-# | CD142-APC-A Mean | CD142-APC-A Median |
|---|---|---|---|
| POS-unpigm. | 73,16 | 2,86 | 2,05 |
| POS-pigm. | 68,36 | 3,23 | 2,37 |

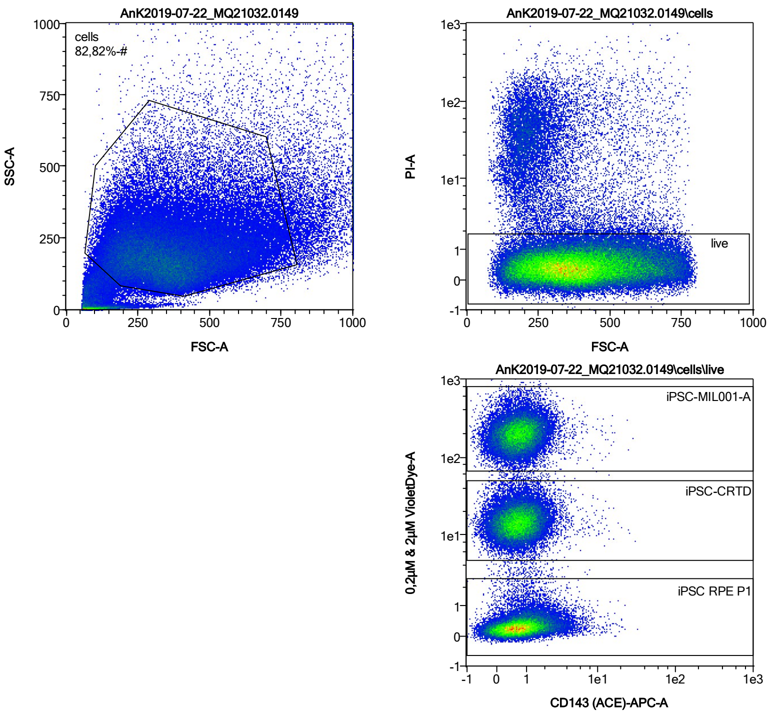


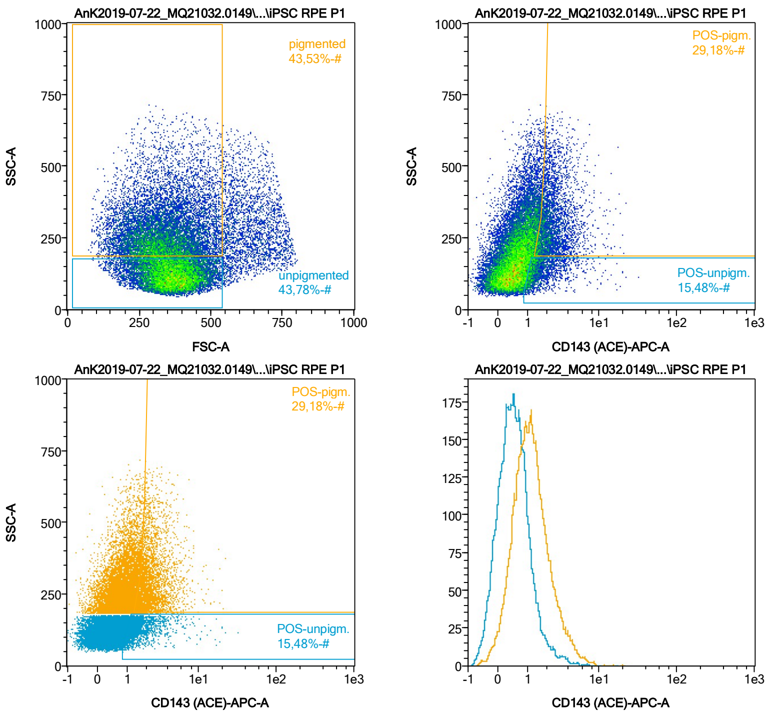


| Name | %-# | CD143 (ACE)-APC-A Mean | CD143 (ACE)-APC-A Median |
|---|---|---|---|
| POS-unpigm. | 15,48 | 1,83 | 1,36 |
| POS-pigm. | 29,18 | 2,63 | 2,19 |

Date:2024-06-14

Operator: Sebastian Knoebel

Version:2.13.0+54.g95abf0f4

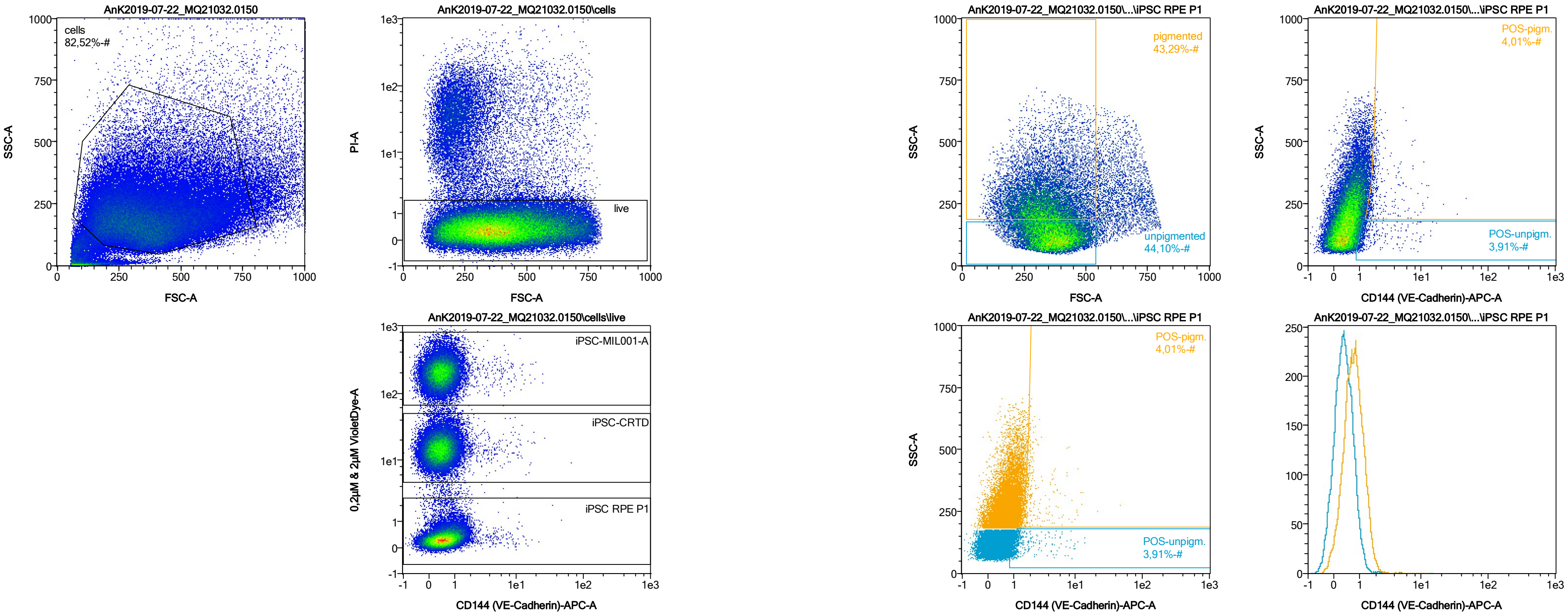


| Name | %-# | CD144 (VE-Cadherin)-APC-A Mean | CD144 (VE-Cadherin)-APC-A Median |
|---|---|---|---|
| POS-unpigm. | 3,91 | 1,78 | 1,18 |
| POS-pigm. | 4,01 | 2,47 | 1,82 |

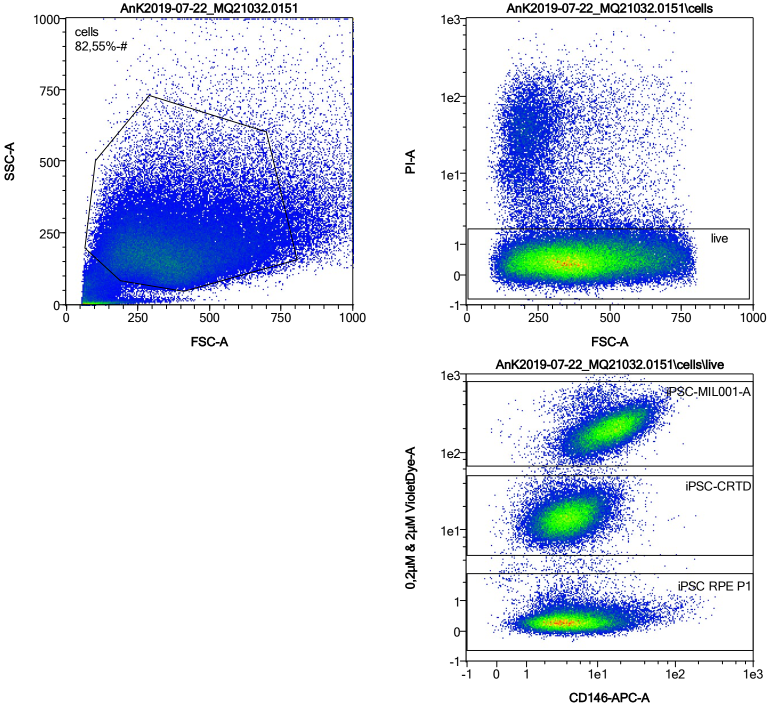


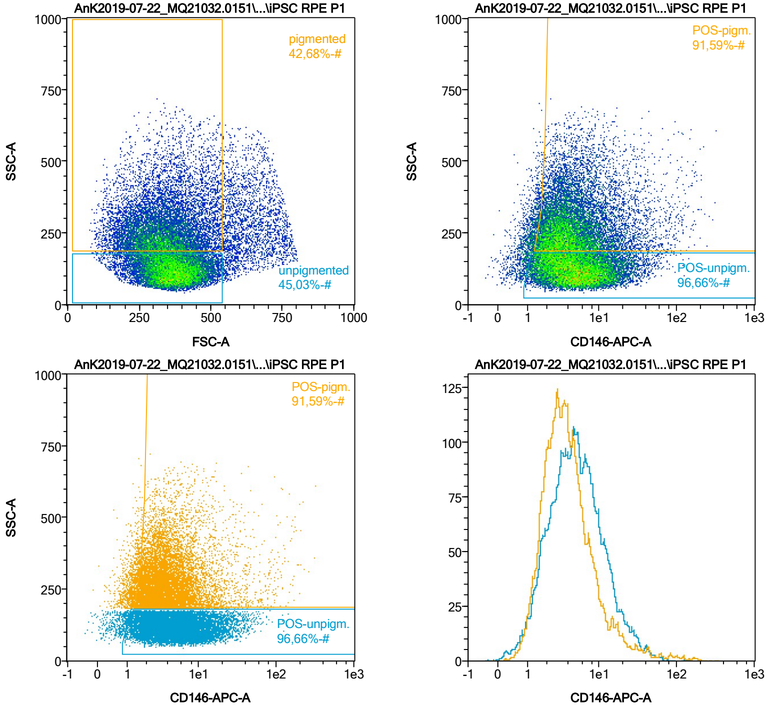


| Name | %-# | CD146-APC-A Mean | CD146-APC-A Median |
|---|---|---|---|
| POS-unpigm. | 96,66 | 7,14 | 4,98 |
| POS-pigm. | 91,59 | 6,88 | 3,75 |

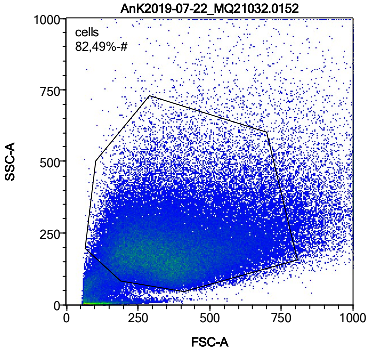


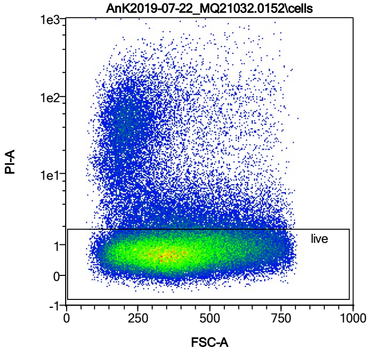


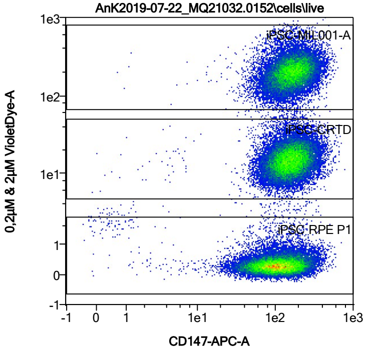


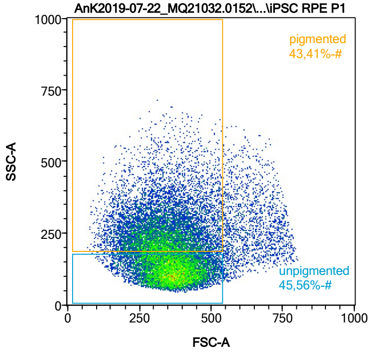


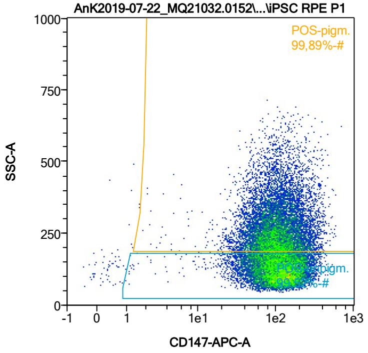


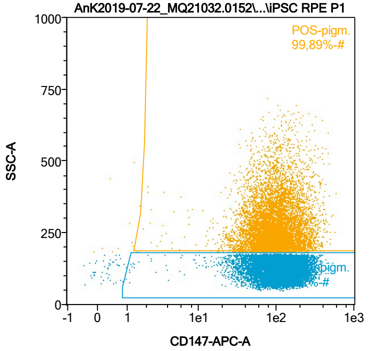


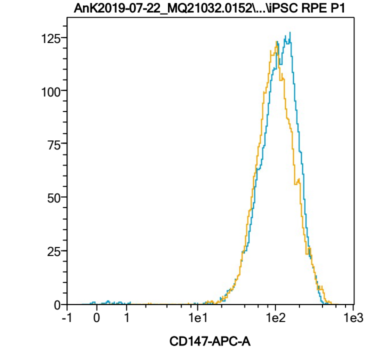


| Name | %-# | CD147-APC-A Mean | CD147-APC-A Median |
|---|---|---|---|
| POS-unpigm. | 99,52 | 122,10 | 111,69 |
| POS-pigm. | 99,89 | 112,97 | 96,92 |

Date:2024-06-14

Operator: Sebastian Knoebel

Version:2.13.0+54.g95abf0f4

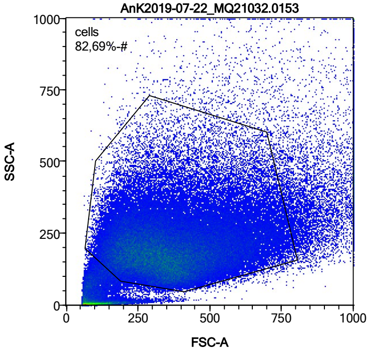


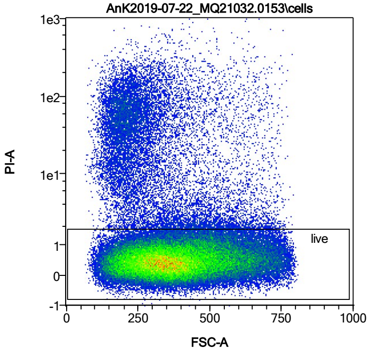


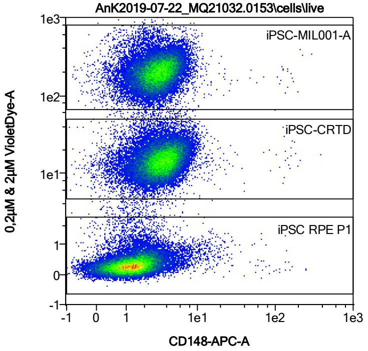


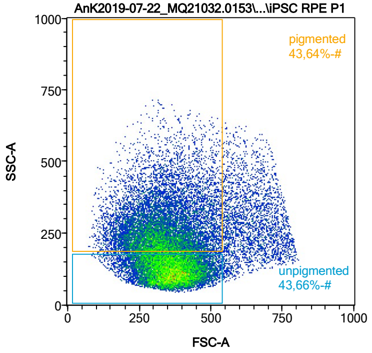


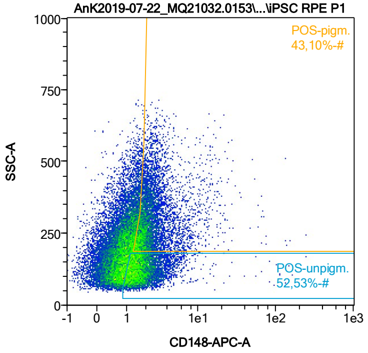


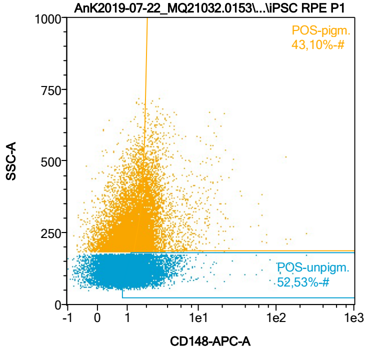


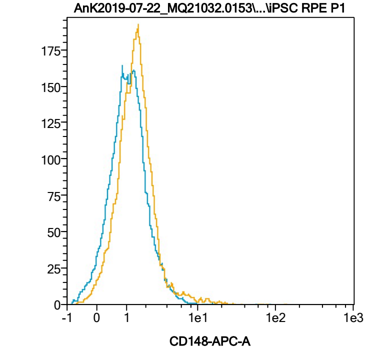


| Name | %-# | CD148-APC-A Mean | CD148-APC-A Median |
|---|---|---|---|
| POS-unpigm. | 52,53 | 2,11 | 1,56 |
| POS-pigm. | 43,10 | 2,95 | 2,04 |

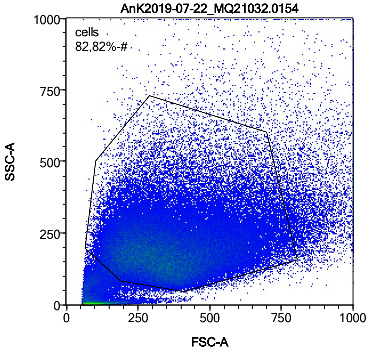


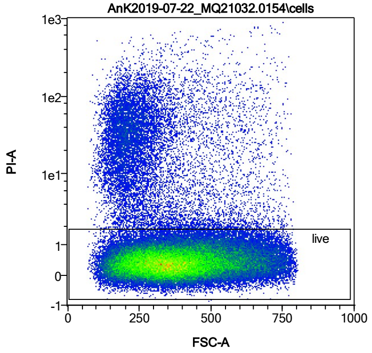


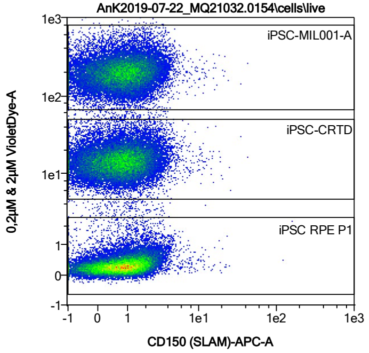


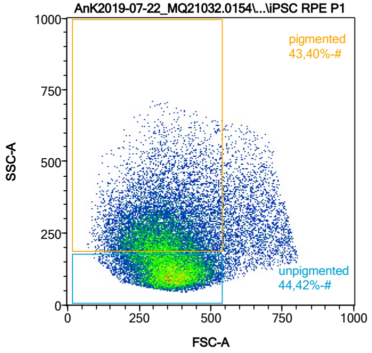


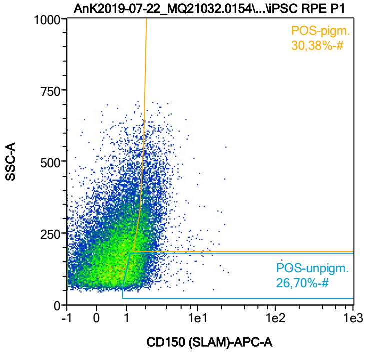


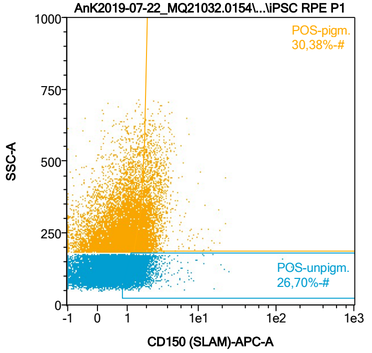


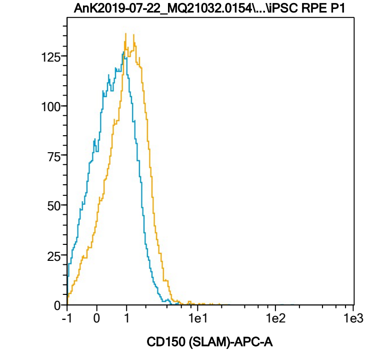


| Name | %-# | CD150 (SLAM)-APC-A Mean | CD150 (SLAM)-APC-A Median |
|---|---|---|---|
| POS-unpigm. | 26,70 | 1,70 | 1,41 |
| POS-pigm. | 30,38 | 2,27 | 2,02 |

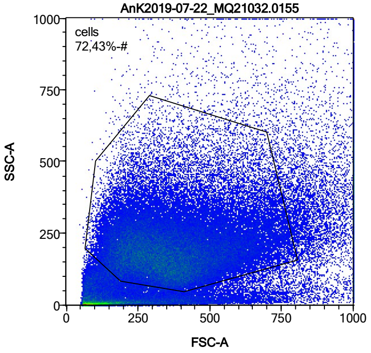


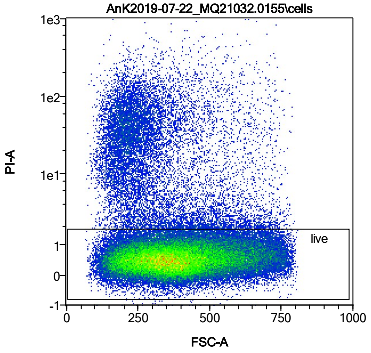


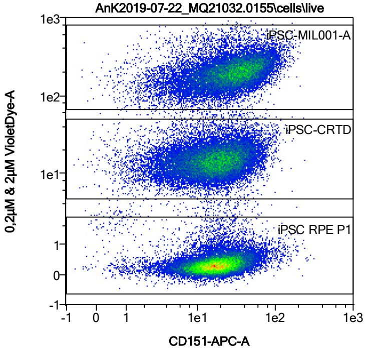


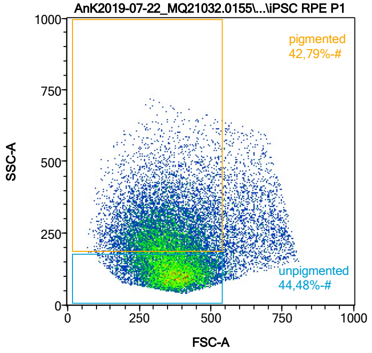


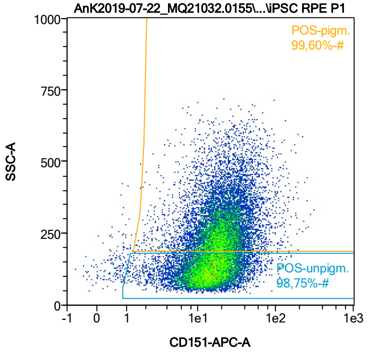


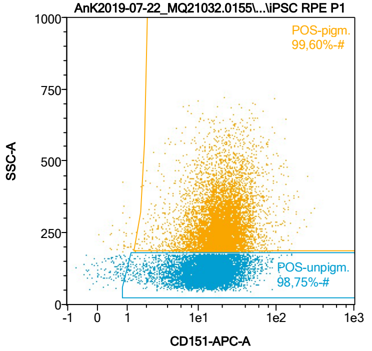


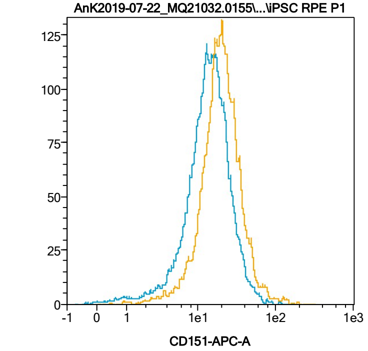


| Name | %-# | CD151-APC-A Mean | CD151-APC-A Median |
|---|---|---|---|
| POS-unpigm. | 98,75 | 16,08 | 13,97 |
| POS-pigm. | 99,60 | 22,93 | 19,47 |

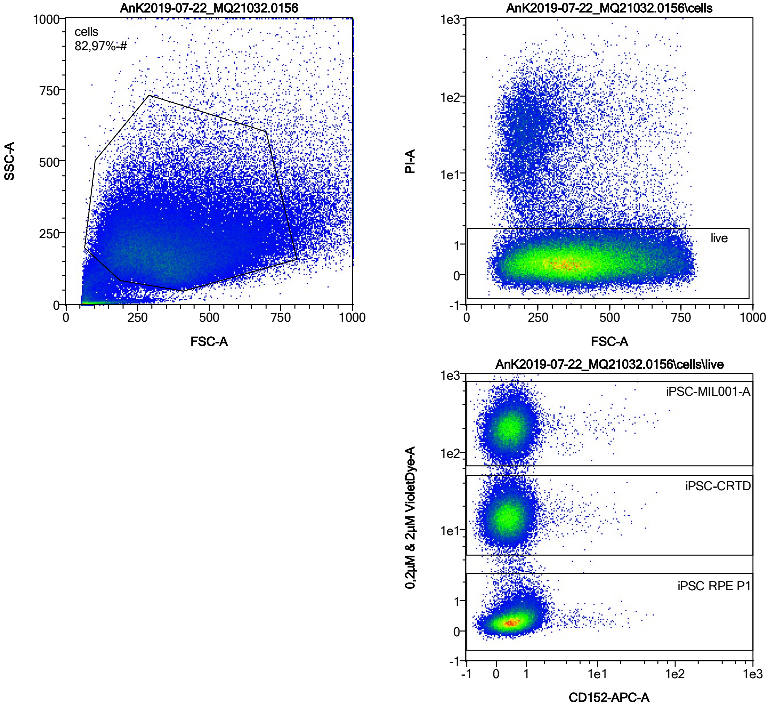


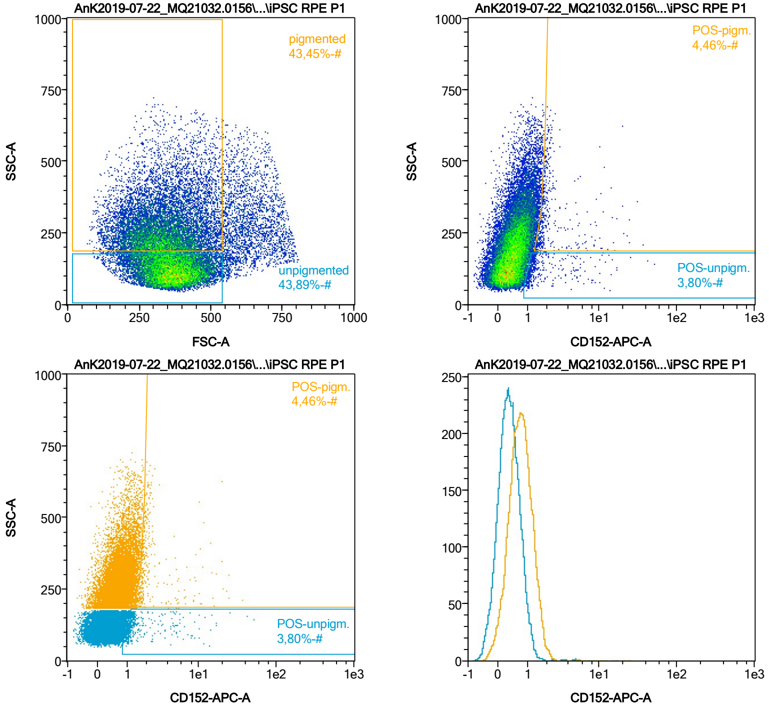


| Name | %-# | CD152-APC-A Mean | CD152-APC-A Median |
|---|---|---|---|
| POS-unpigm. | 3,80 | 2,23 | 1,18 |
| POS-pigm. | 4,46 | 2,68 | 1,80 |

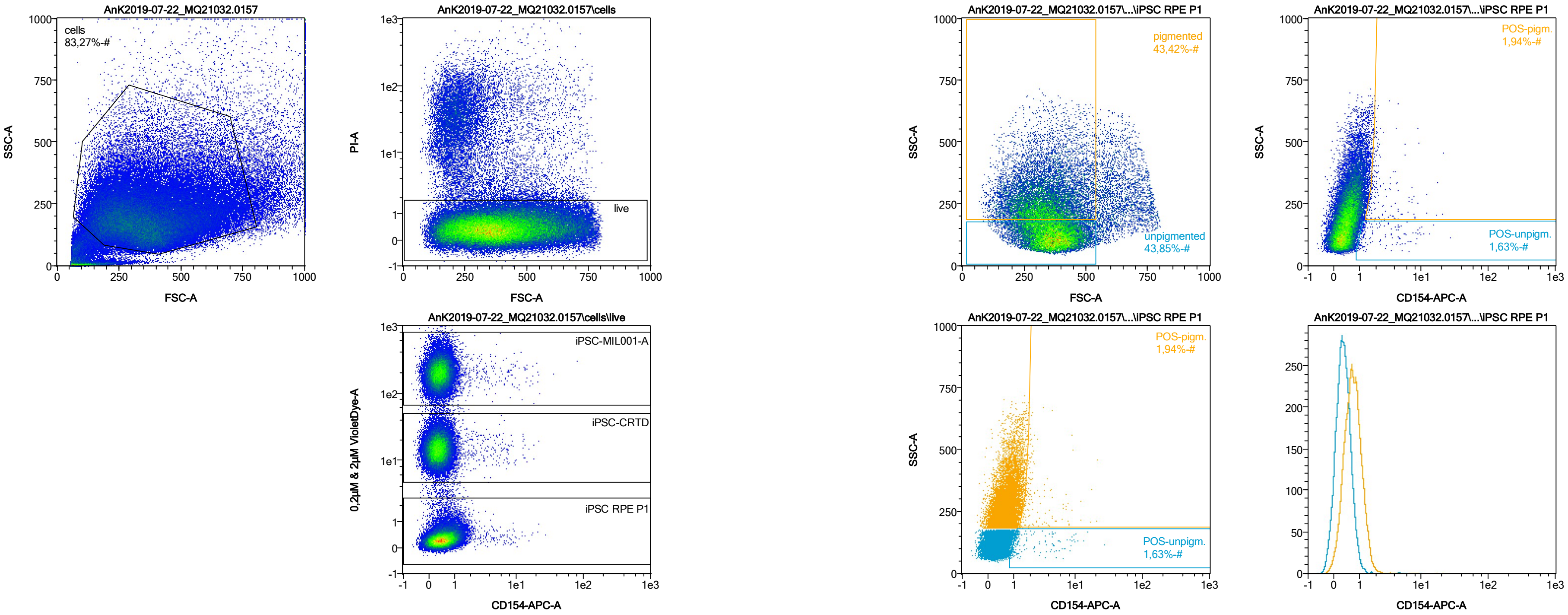


| Name | %-# | CD154-APC-A Mean | CD154-APC-A Median |
|---|---|---|---|
| POS-unpigm. | 1,63 | 2,72 | 1,26 |
| POS-pigm. | 1,94 | 2,86 | 1,91 |

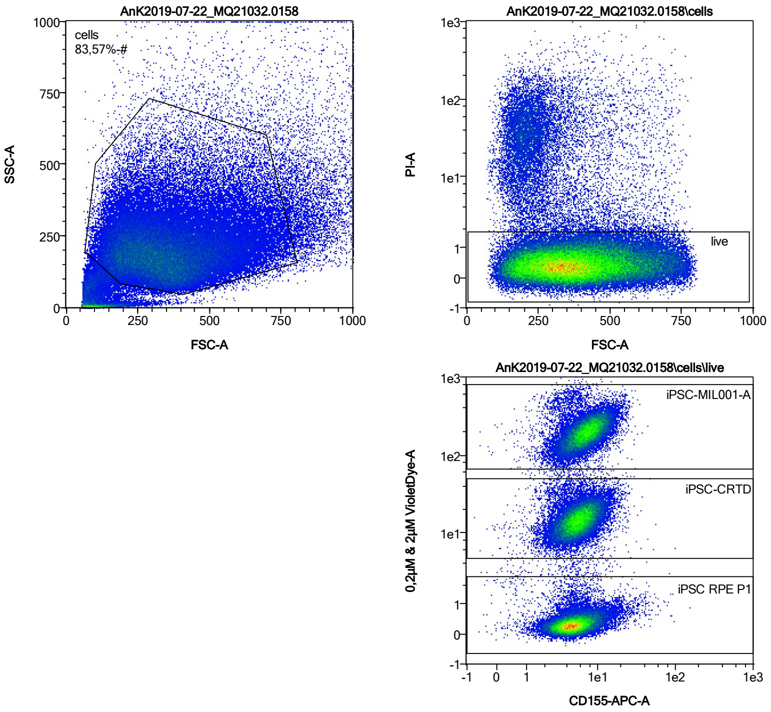


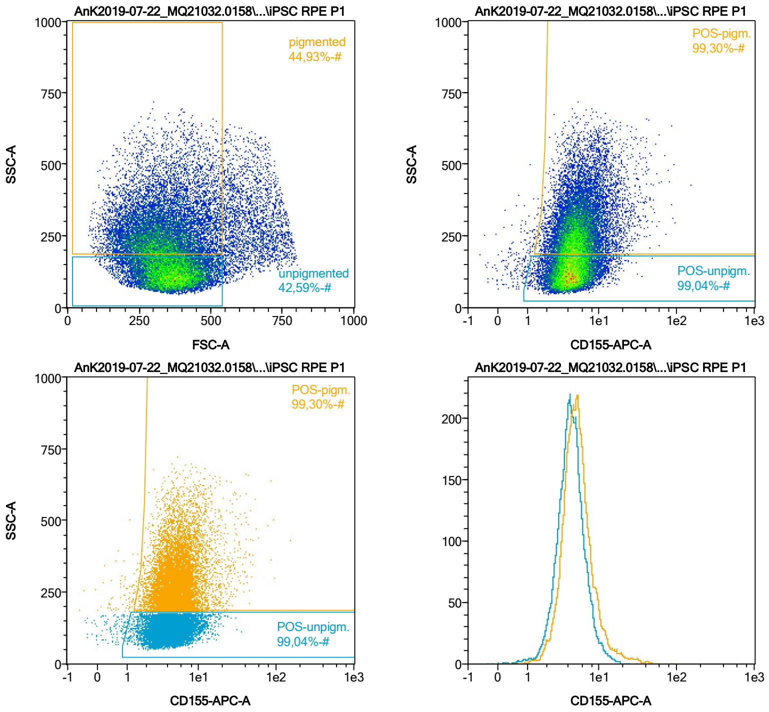


| Name | %-# | CD155-APC-A Mean | CD155-APC-A Median |
|---|---|---|---|
| POS-unpigm. | 99,04 | 4,63 | 4,23 |
| POS-pigm. | 99,30 | 5,95 | 5,02 |

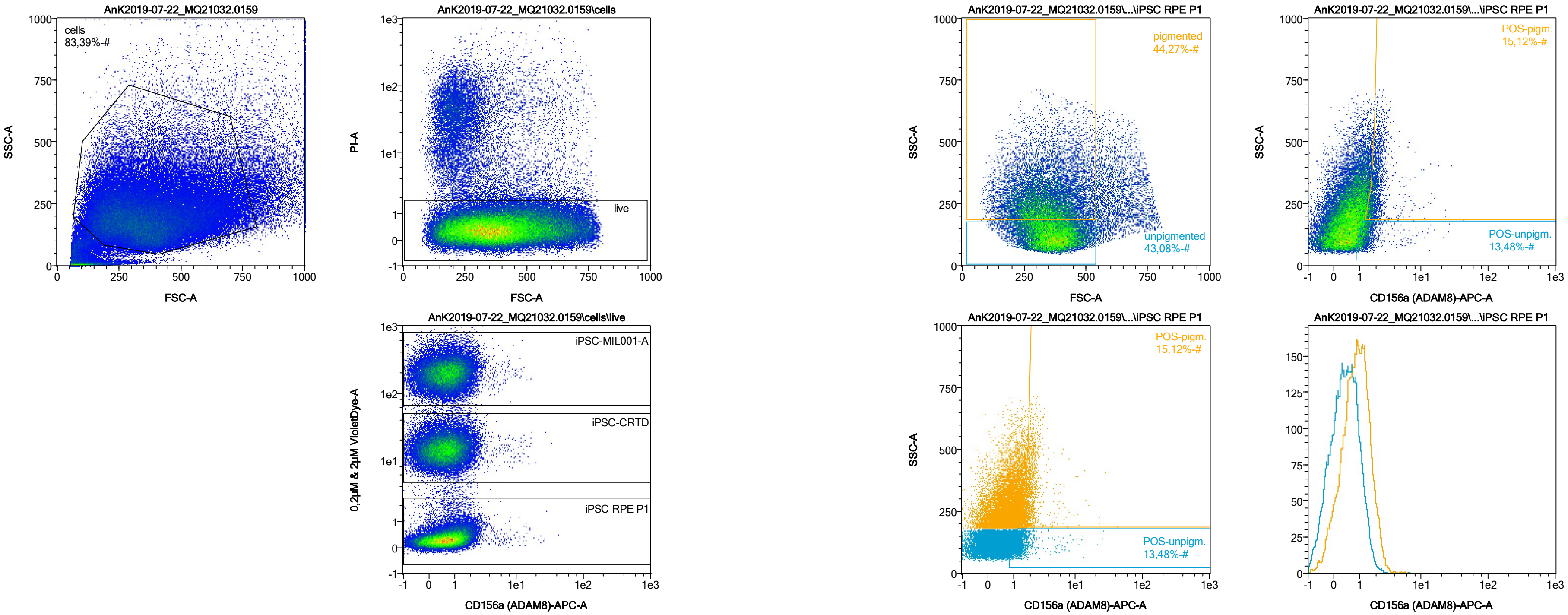


| Name | %-# | CD156a (ADAM8)-APC-A Mean | CD156a (ADAM8)-APC-A Median |
|---|---|---|---|
| POS-unpigm. | 13,48 | 1,51 | 1,26 |
| POS-pigm. | 15,12 | 2,14 | 1,88 |

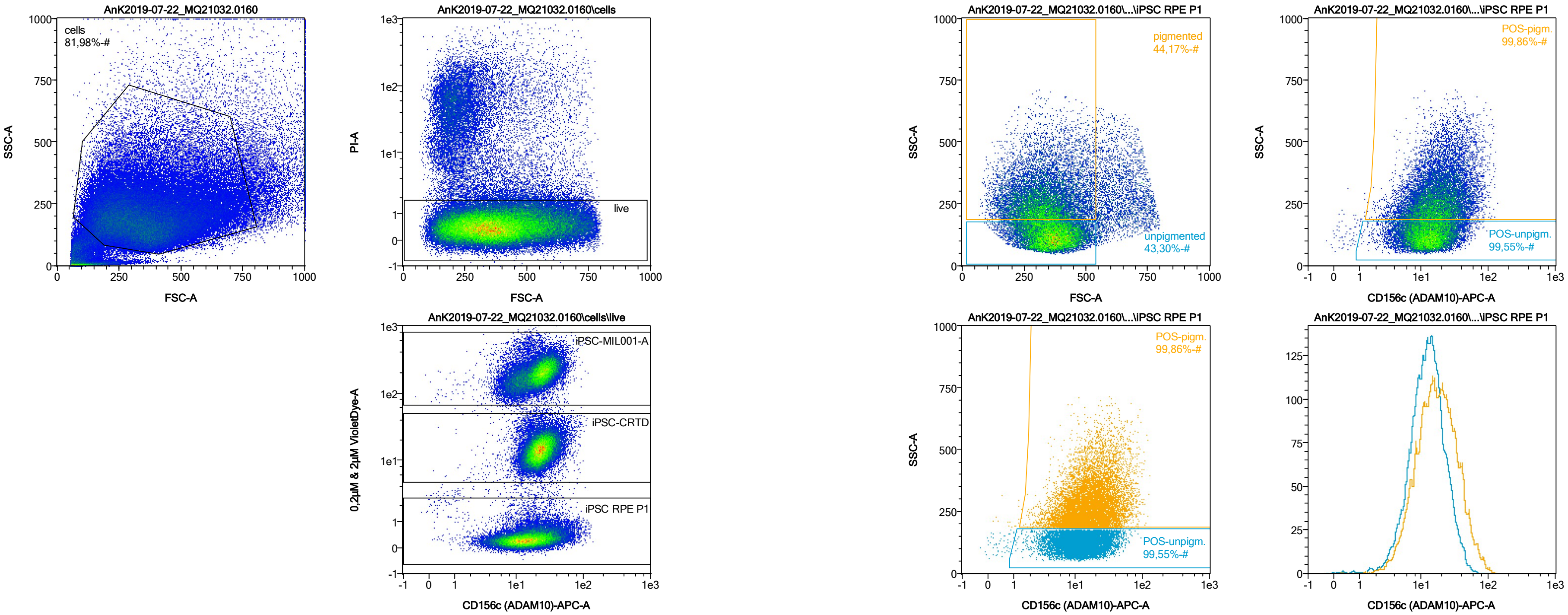


| Name | %-# | CD156c (ADAM10)-APC-A Mean | CD156c (ADAM10)-APC-A Median |
|---|---|---|---|
| POS-unpigm. | 99,55 | 14,34 | 12,48 |
| POS-pigm. | 99,86 | 20,92 | 16,67 |

Date:2024-06-14
Operator: Sebastian Knoebel
Version:2.13.0+54.g95abf0f4

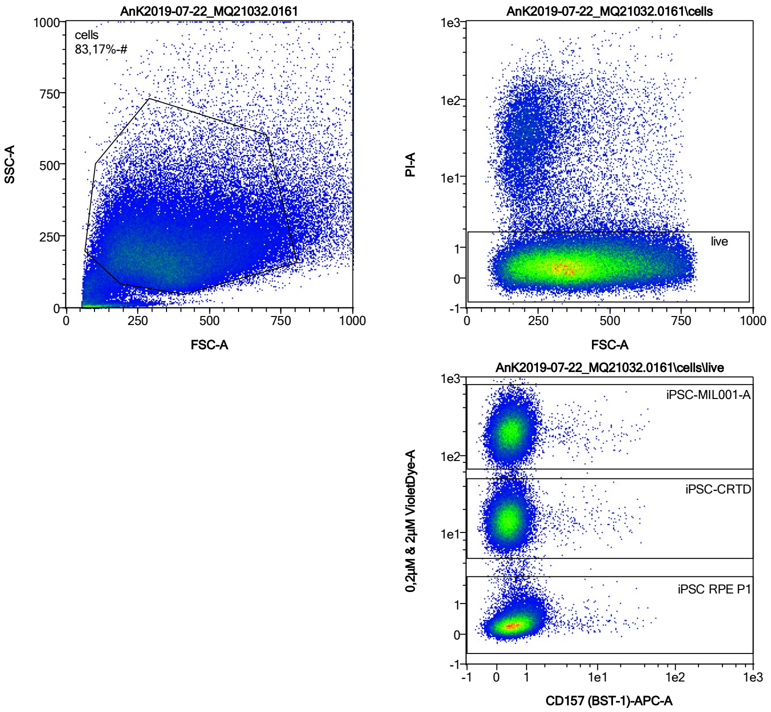


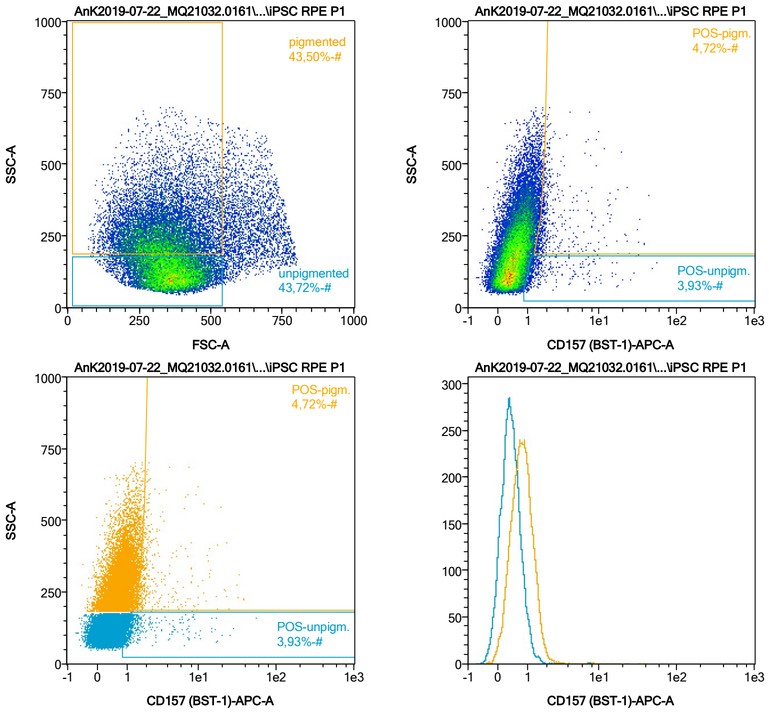


| Name | %-# | CD157 (BST-1)-APC-A Mean | CD157 (BST-1)-APC-A Median |
|---|---|---|---|
| POS-unpigm. | 3,93 | 2,31 | 1,20 |
| POS-pigm. | 4,72 | 3,16 | 1,87 |

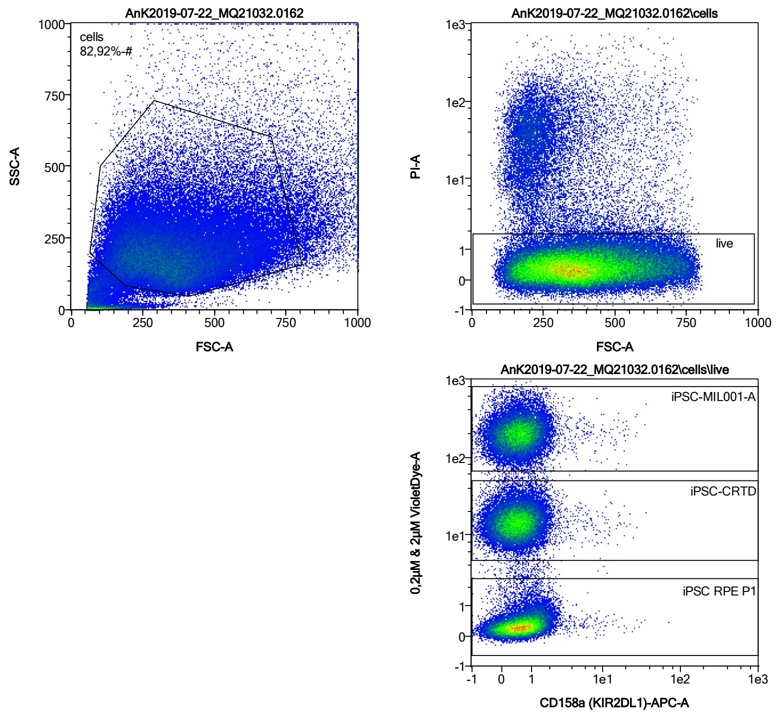


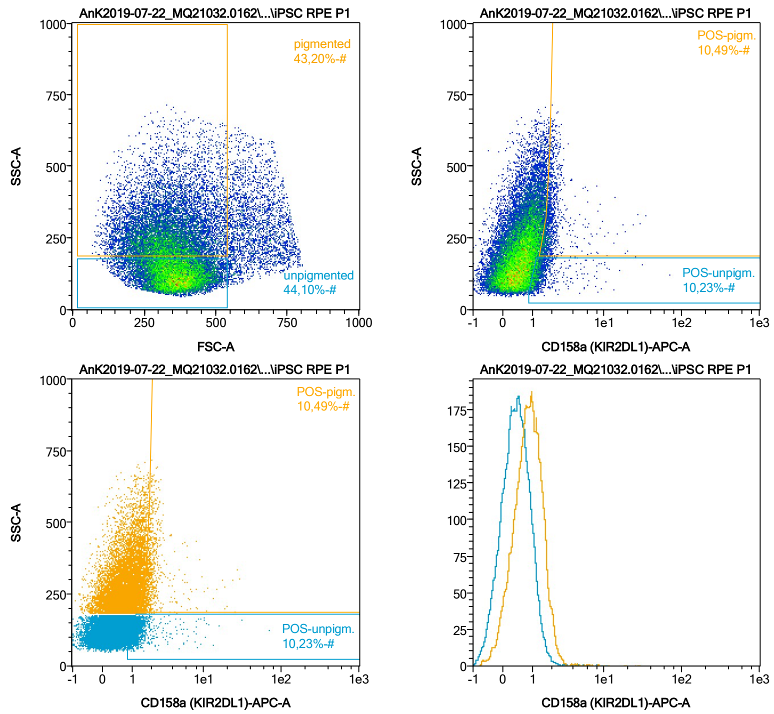


| Name | %-# | CD158a (KIR2DL1)-APC-A Mean | CD158a (KIR2DL1)-APC-A Median |
|---|---|---|---|
| POS-unpigm. | 10,23 | 1,56 | 1,21 |
| POS-pigm. | 10,49 | 2,18 | 1,81 |

Date:2024-06-14
Operator: Sebastian Knoebel
Version:2.13.0+54.g95abf0f4

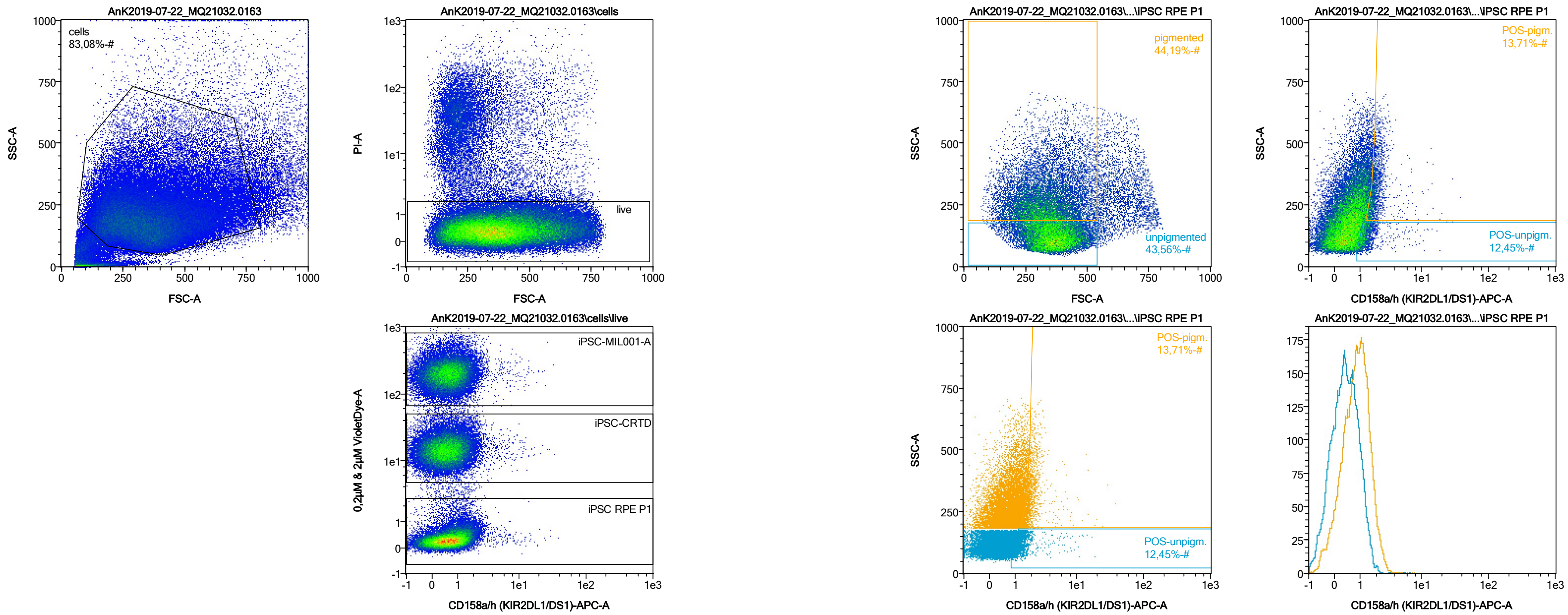


| Name | %-# | CD158a/h (KIR2DL1/DS1)-APC-A Mean | CD158a/h (KIR2DL1/DS1)-APC-A Median |
|---|---|---|---|
| POS-unpigm. | 12,45 | 1,49 | 1,24 |
| POS-pigm. | 13,71 | 2,10 | 1,83 |

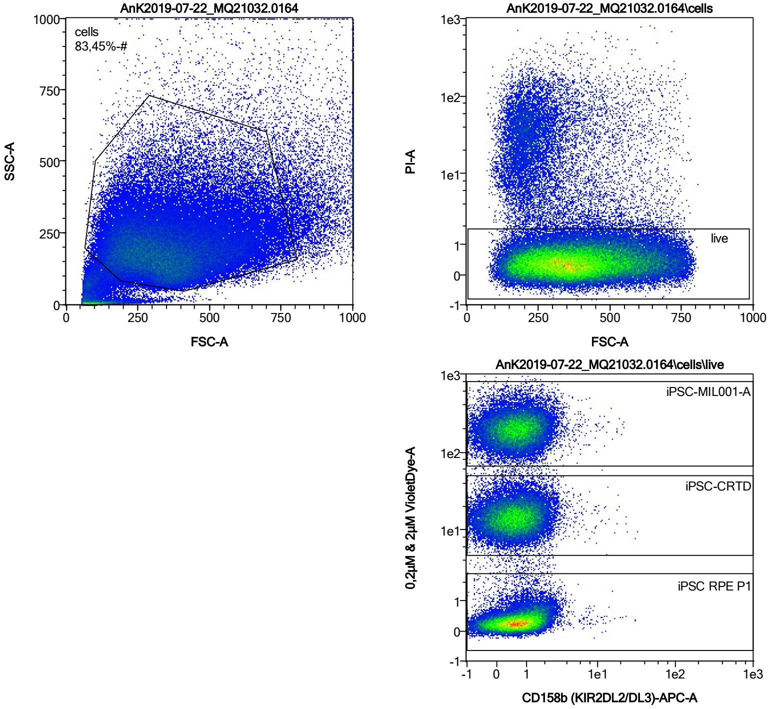


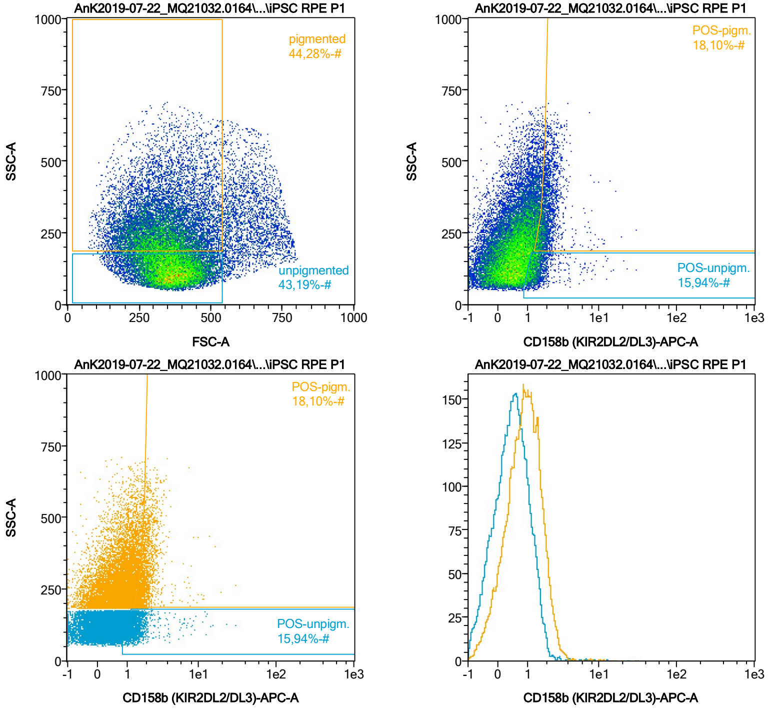


| Name | %-# | CD158b (KIR2DL2/DL3)-APC-A Mean | CD158b (KIR2DL2/DL3)-APC-A Median |
|---|---|---|---|
| POS-unpigm. | 15,94 | 1,54 | 1,30 |
| POS-pigm. | 18,10 | 2,10 | 1,89 |

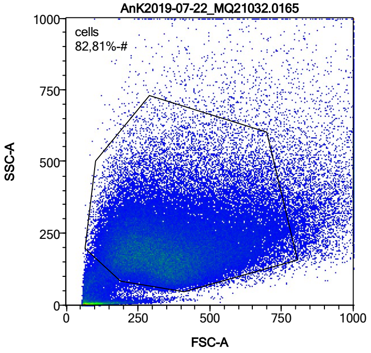


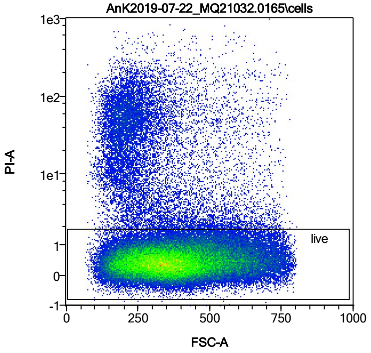


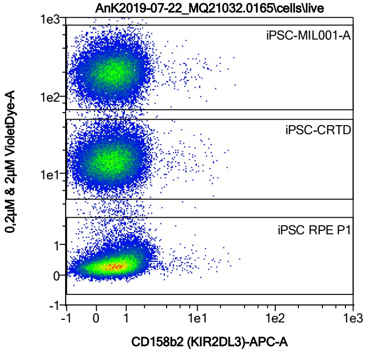


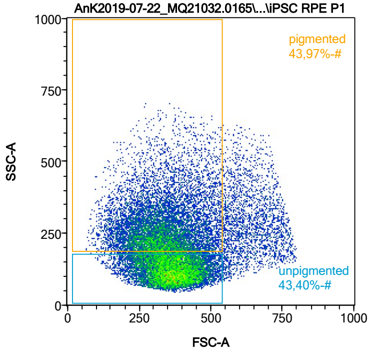


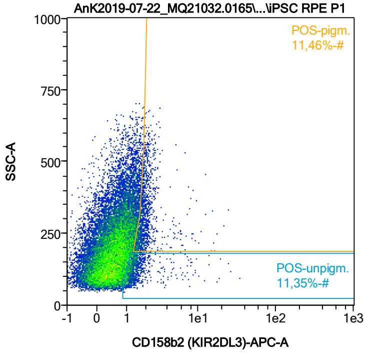


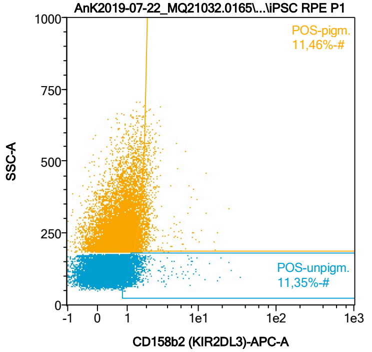


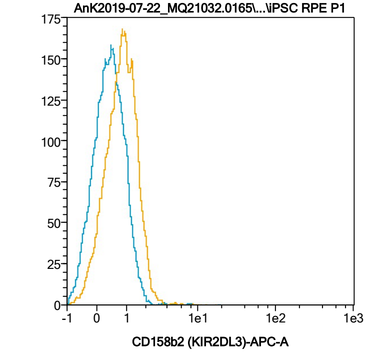


| Name | %-# | CD158b2 (KIR2DL3)-APC-A Mean | CD158b2 (KIR2DL3)-APC-A Median |
|---|---|---|---|
| POS-unpigm. | 11,35 | 1,58 | 1,23 |
| POS-pigm. | 11,46 | 2,10 | 1,83 |

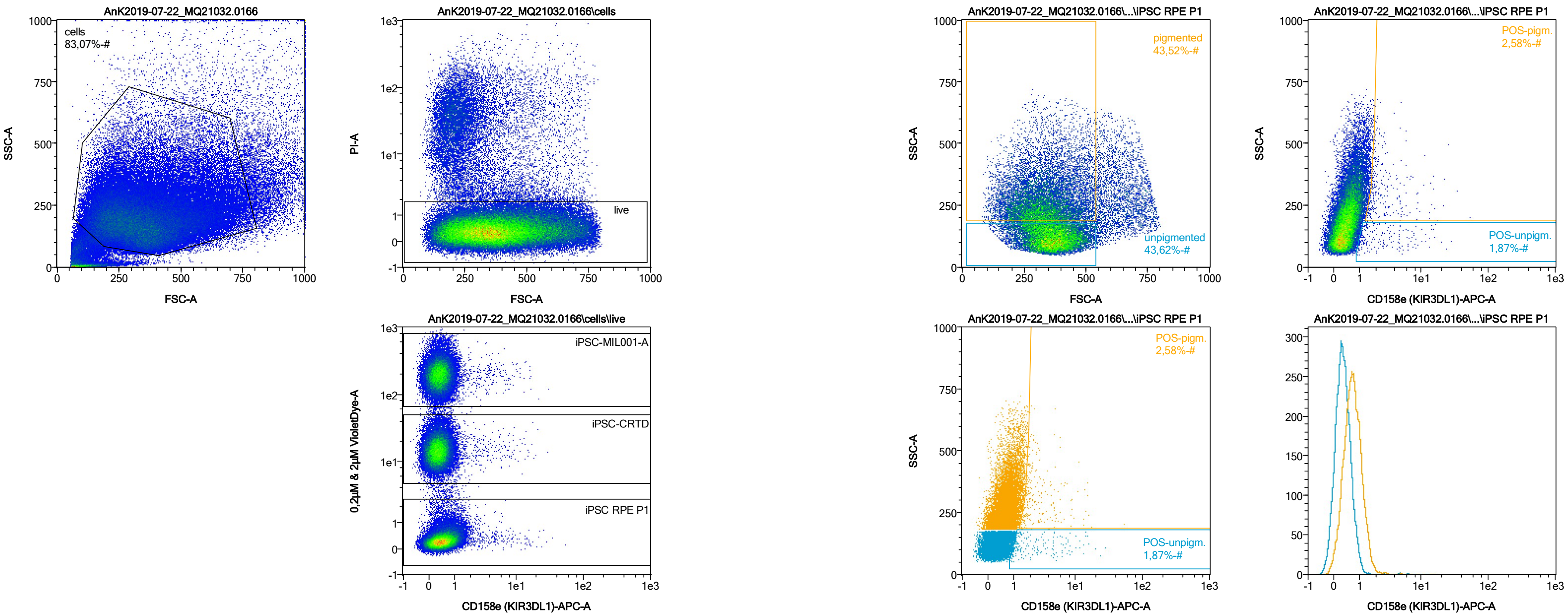


| Name | %-# | CD158e (KIR3DL1)-APC-A Mean | CD158e (KIR3DL1)-APC-A Median |
|---|---|---|---|
| POS-unpigm. | 1,87 | 3,18 | 1,40 |
| POS-pigm. | 2,58 | 3,16 | 1,96 |

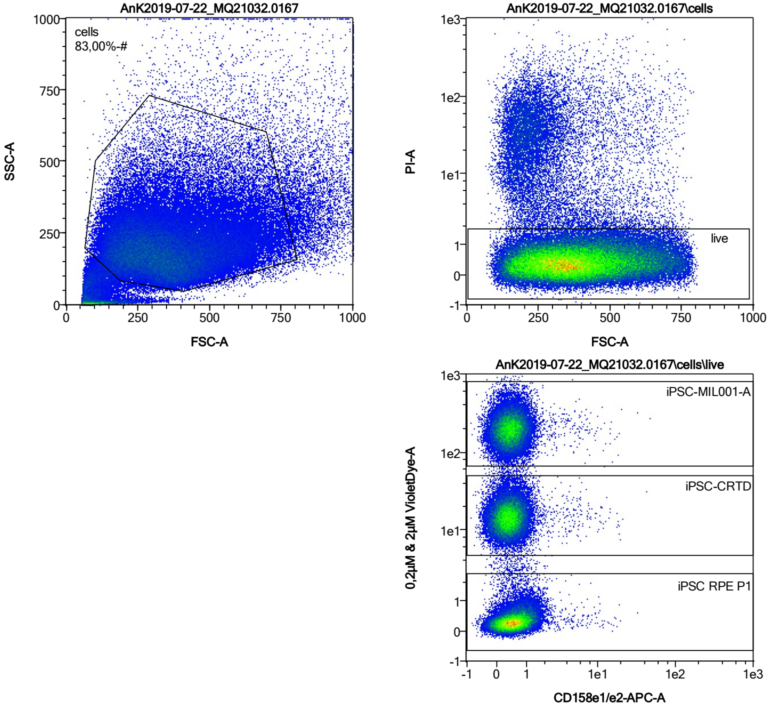


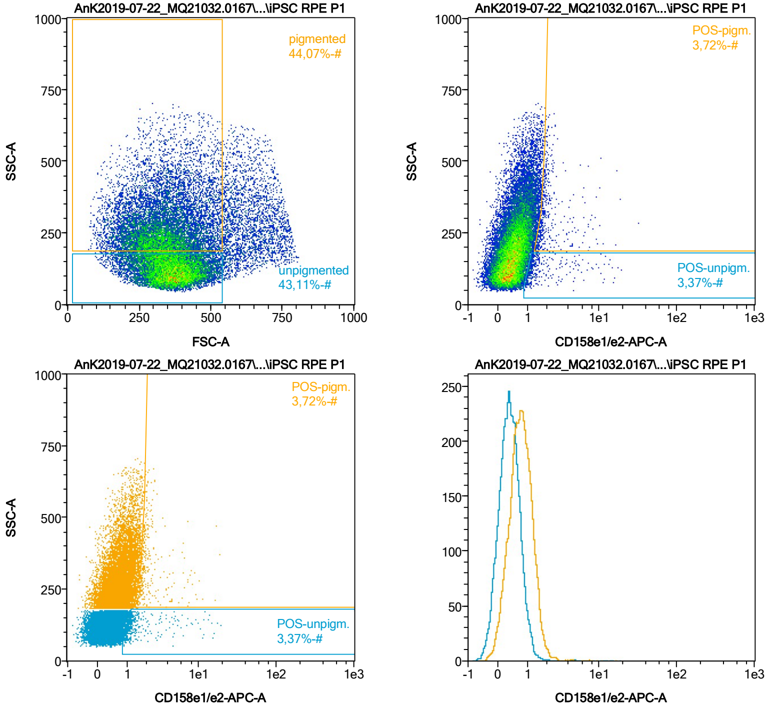


| Name | %-# | CD158e1/e2-APC-A Mean | CD158e1/e2-APC-A Median |
|---|---|---|---|
| POS-unpigm. | 3,37 | 2,02 | 1,18 |
| POS-pigm. | 3,72 | 2,36 | 1,84 |

Date:2024-06-14

Operator: Sebastian Knoebel

Version:2.13.0+54.g95abf0f4

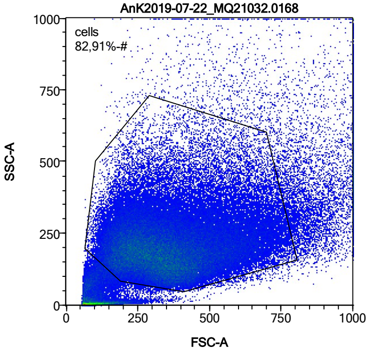


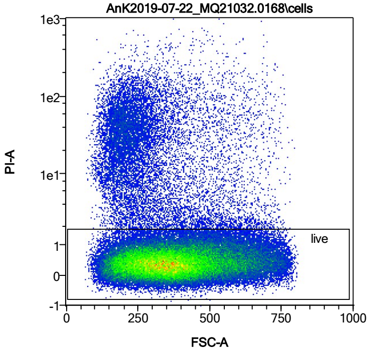


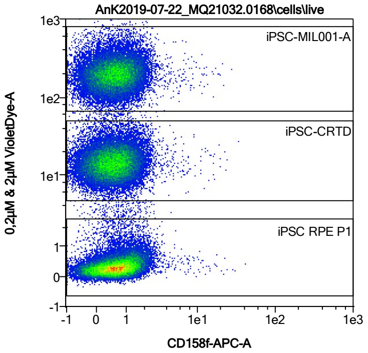


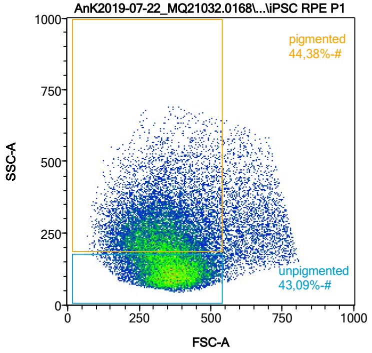


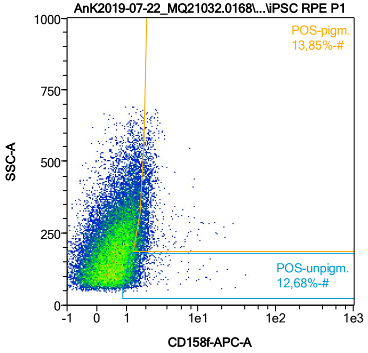


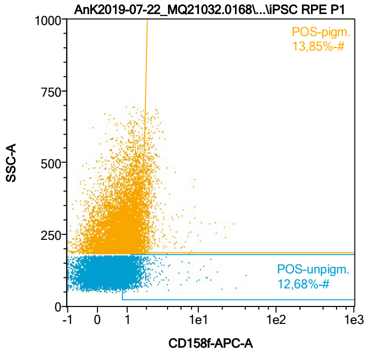


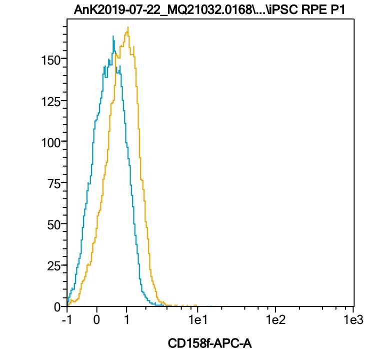


| Name | %-# | CD158f-APC-A Mean | CD158f-APC-A Median |
|---|---|---|---|
| POS-unpigm. | 12,68 | 1,50 | 1,24 |
| POS-pigm. | 13,85 | 2,09 | 1,87 |

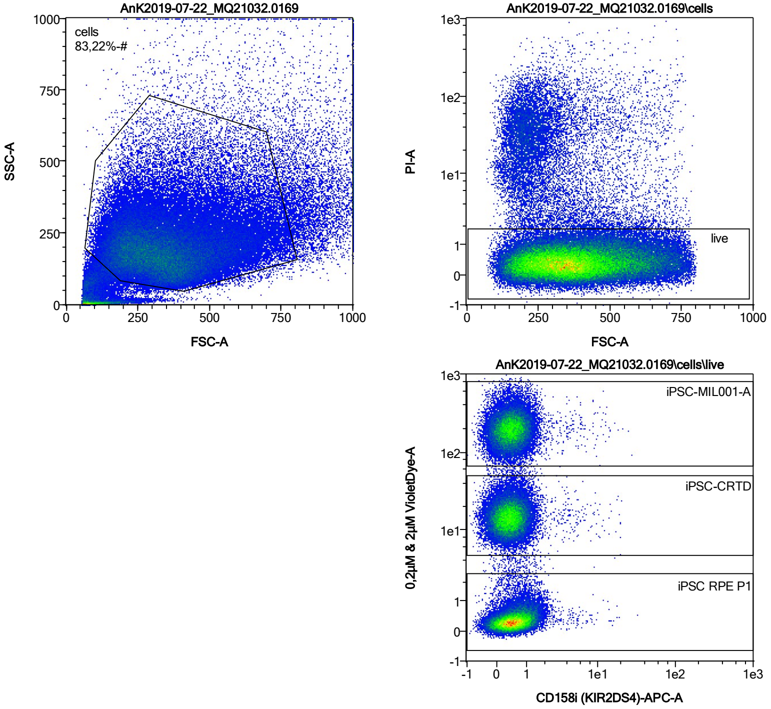


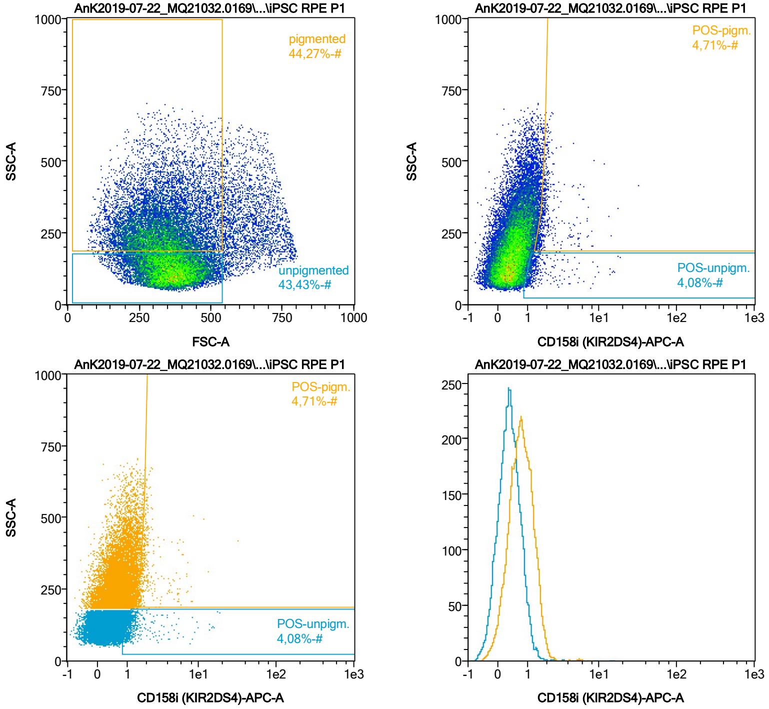


| Name | %-# | CD158i (KIR2DS4)-APC-A Mean | CD158i (KIR2DS4)-APC-A Median |
|---|---|---|---|
| POS-unpigm. | 4,08 | 1,72 | 1,17 |
| POS-pigm. | 4,71 | 2,19 | 1,79 |

Date:2024-06-14

Operator: Sebastian Knoebel

Version:2.13.0+54.g95abf0f4

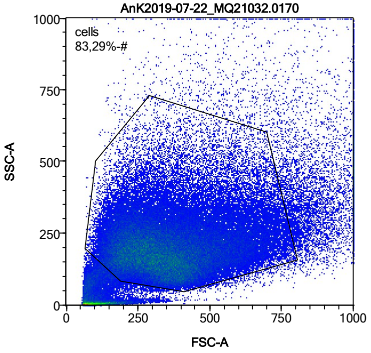


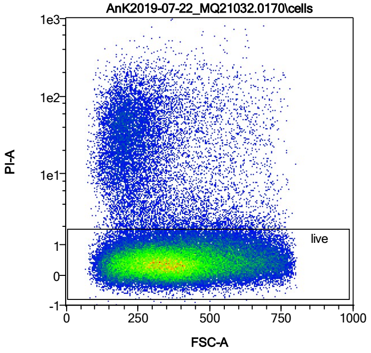


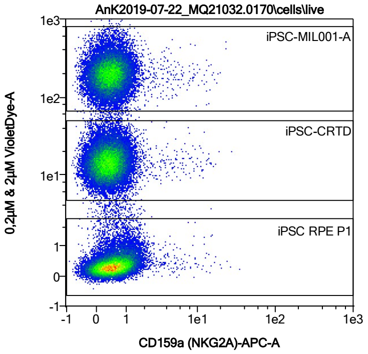


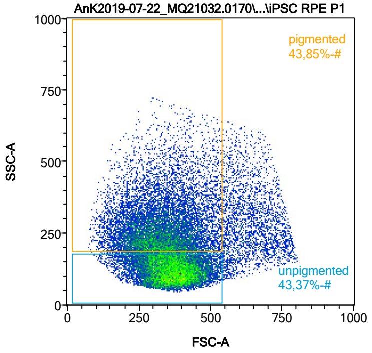


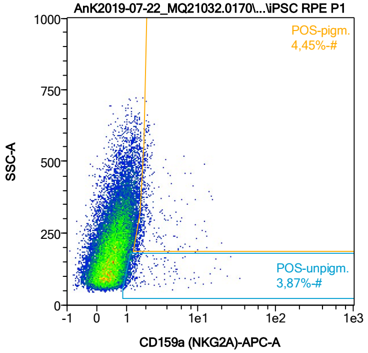


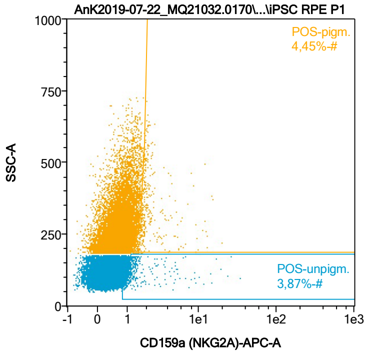


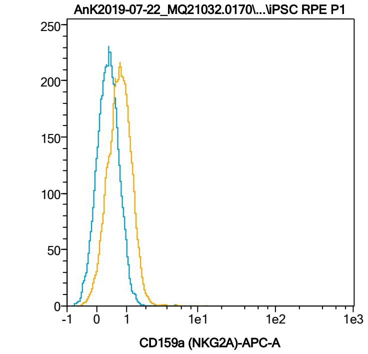


| Name | %-# | CD159a (NKG2A)-APC-A Mean | CD159a (NKG2A)-APC-A Median |
|---|---|---|---|
| POS-unpigm. | 3,87 | 1,93 | 1,18 |
| POS-pigm. | 4,45 | 2,36 | 1,83 |

Date:2024-06-14
Operator: Sebastian Knoebel
Version:2.13.0+54.g95abf0f4

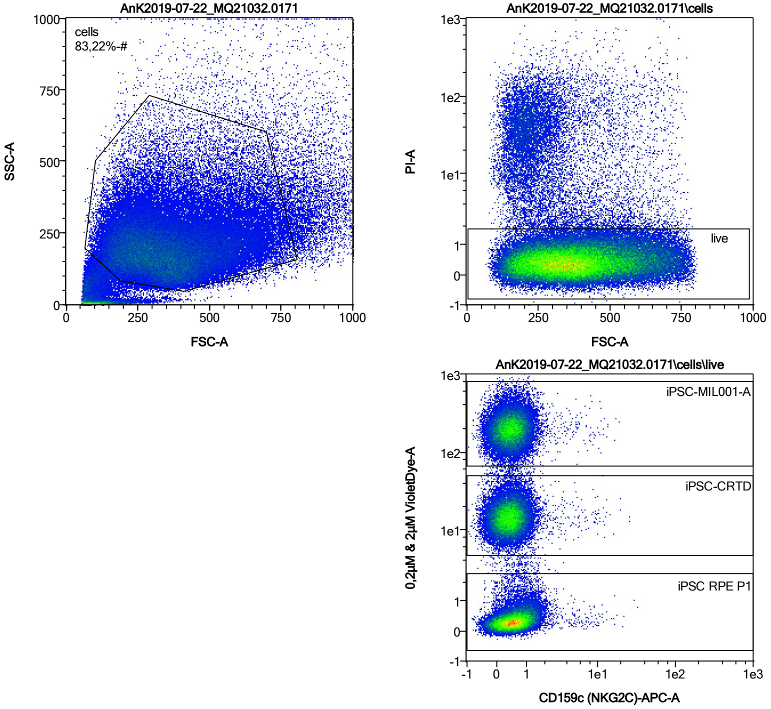


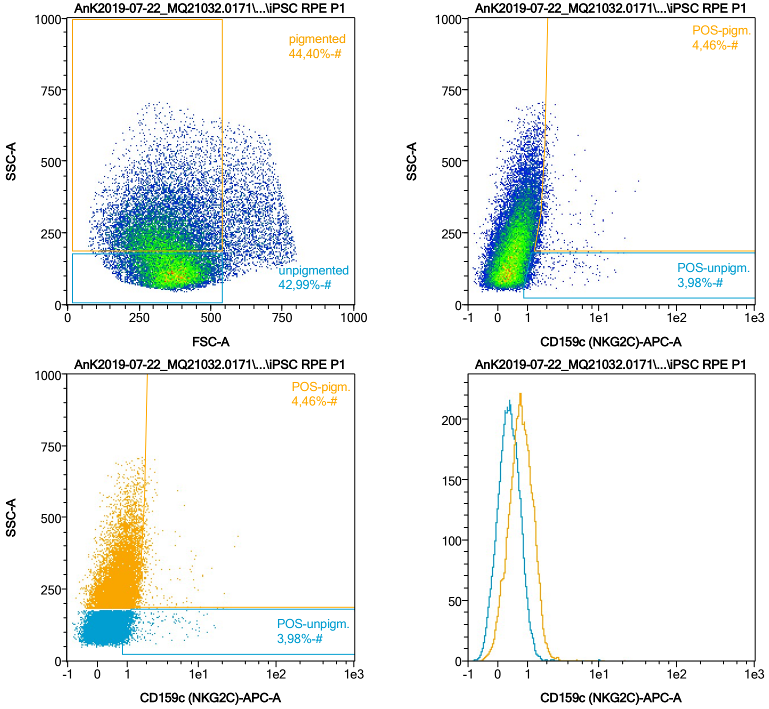


| Name | %-# | CD159c (NKG2C)-APC-A Mean | CD159c (NKG2C)-APC-A Median |
|---|---|---|---|
| POS-unpigm. | 3,98 | 1,88 | 1,19 |
| POS-pigm. | 4,46 | 2,45 | 1,79 |

Date:2024-06-14
Operator: Sebastian Knoebel
Version:2.13.0+54.g95abf0f4

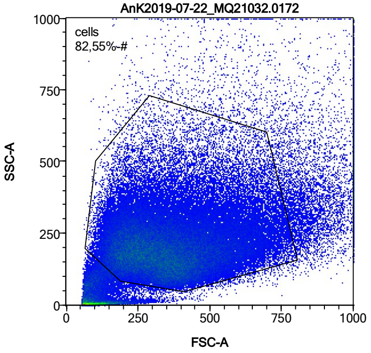


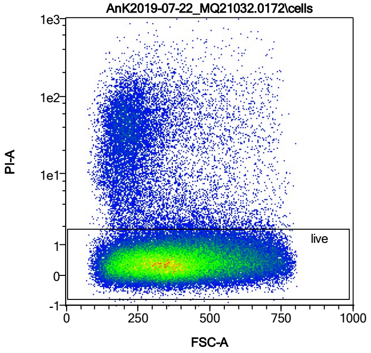


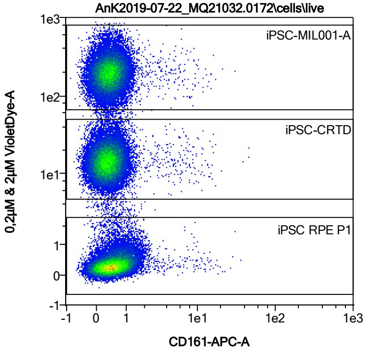


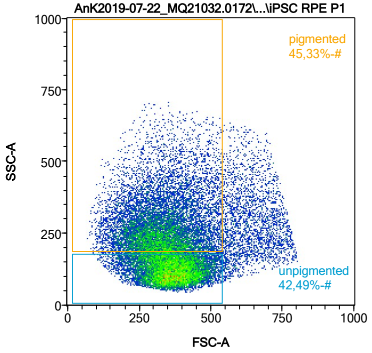


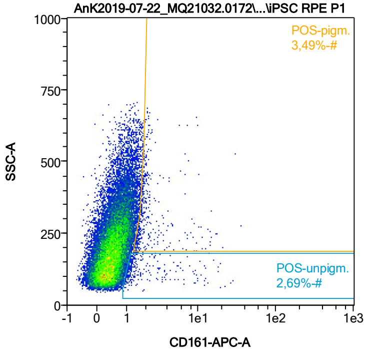


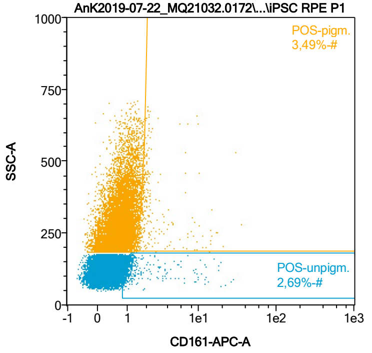


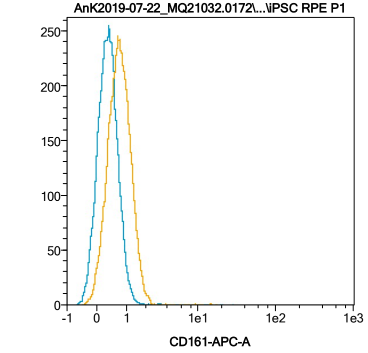


| Name | %-# | CD161-APC-A Mean | CD161-APC-A Median |
|---|---|---|---|
| POS-unpigm. | 2,69 | 2,62 | 1,22 |
| POS-pigm. | 3,49 | 2,79 | 1,83 |

Date:2024-06-14

Operator: Sebastian Knoebel

Version:2.13.0+54.g95abf0f4

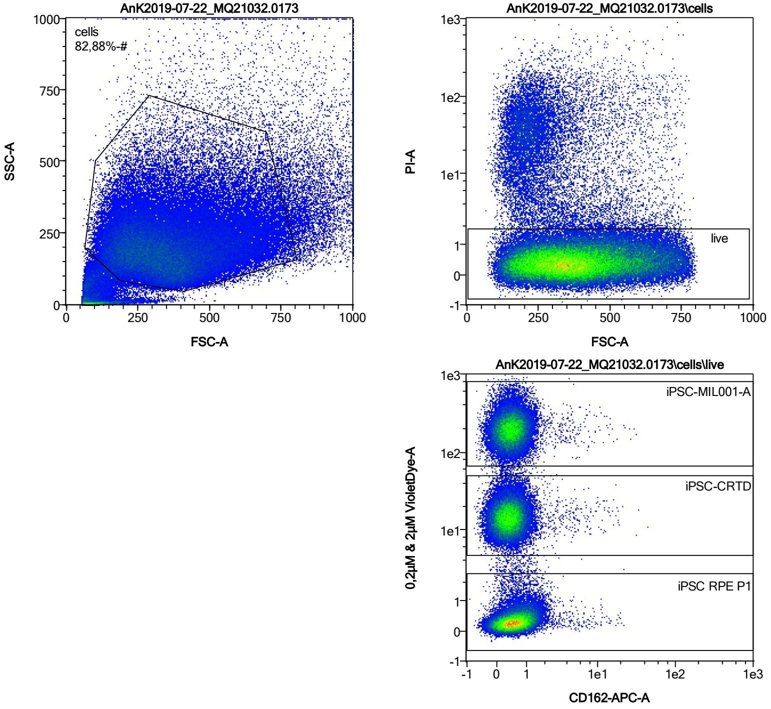


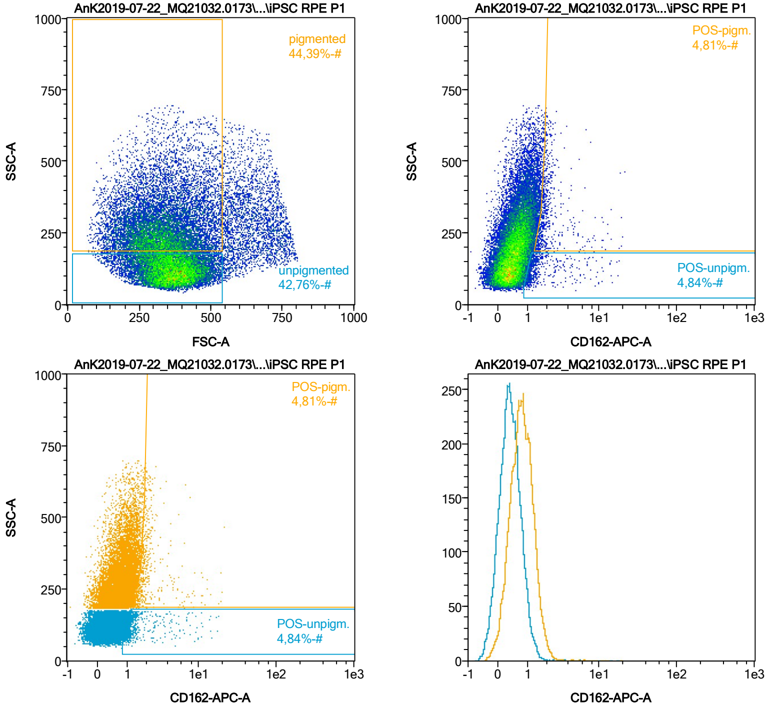


| Name | %-# | CD162-APC-A Mean | CD162-APC-A Median |
|---|---|---|---|
| POS-unpigm. | 4,84 | 1,95 | 1,22 |
| POS-pigm. | 4,81 | 2,48 | 1,85 |

Date:2024-06-14

Operator: Sebastian Knoebel

Version:2.13.0+54.g95abf0f4

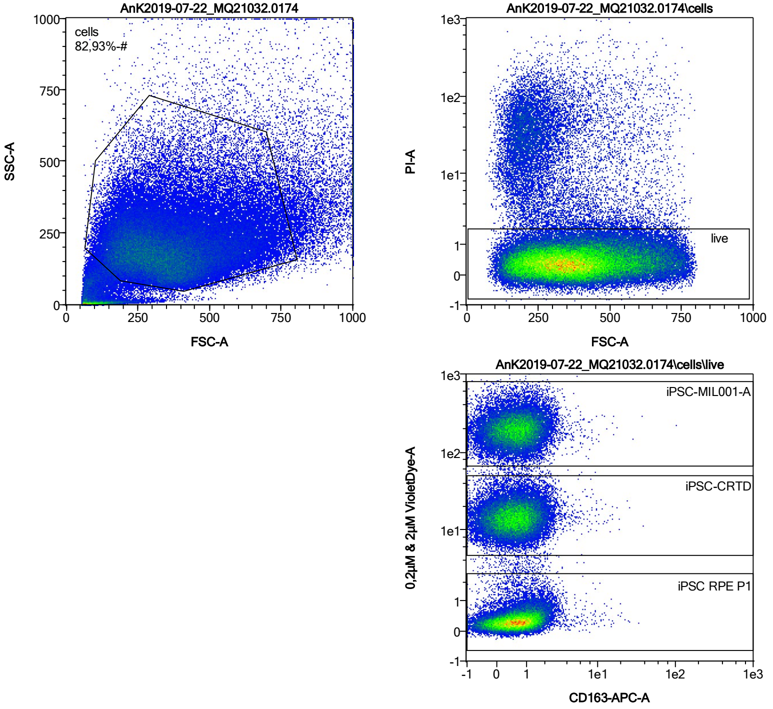


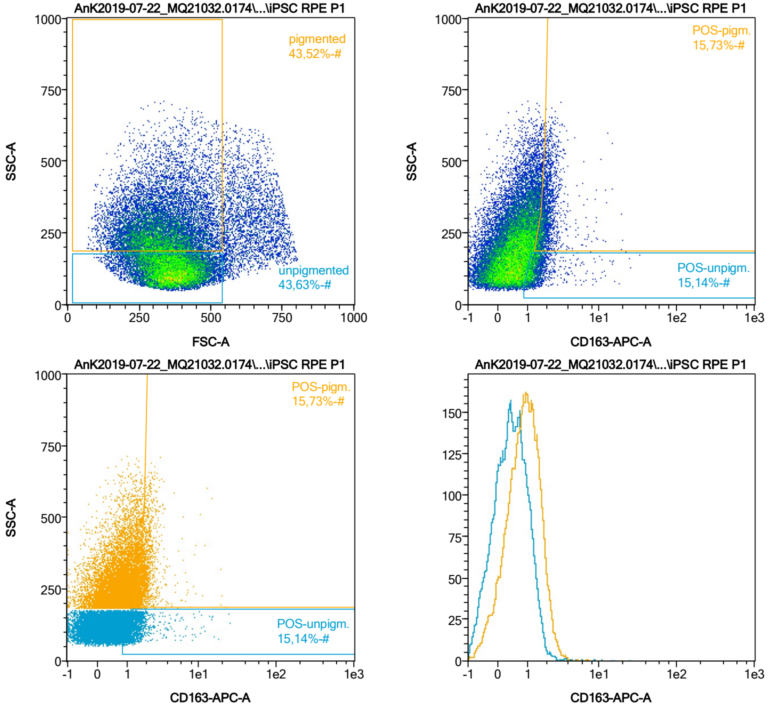


| Name | %-# | CD163-APC-A Mean | CD163-APC-A Median |
|---|---|---|---|
| POS-unpigm. | 15,14 | 1,55 | 1,26 |
| POS-pigm. | 15,73 | 2,08 | 1,85 |

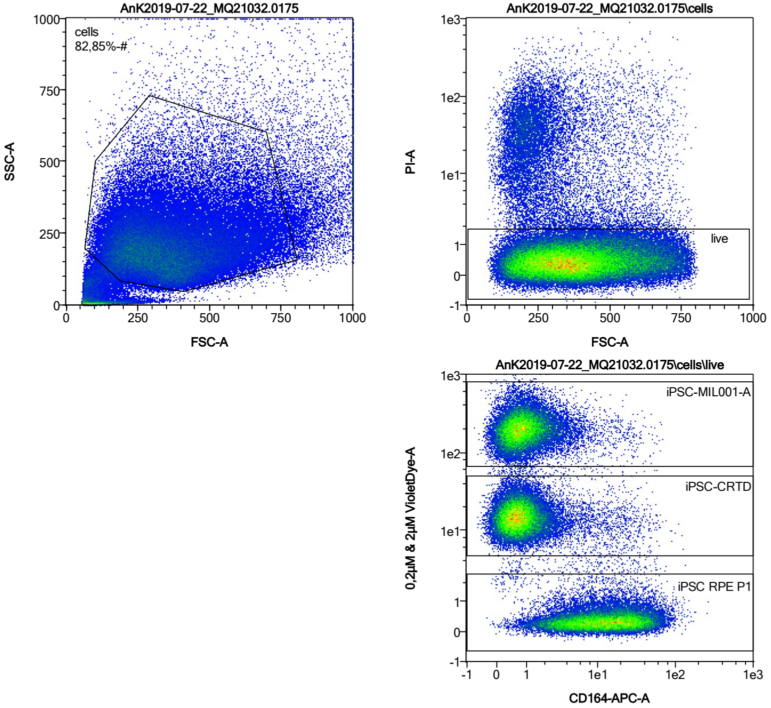


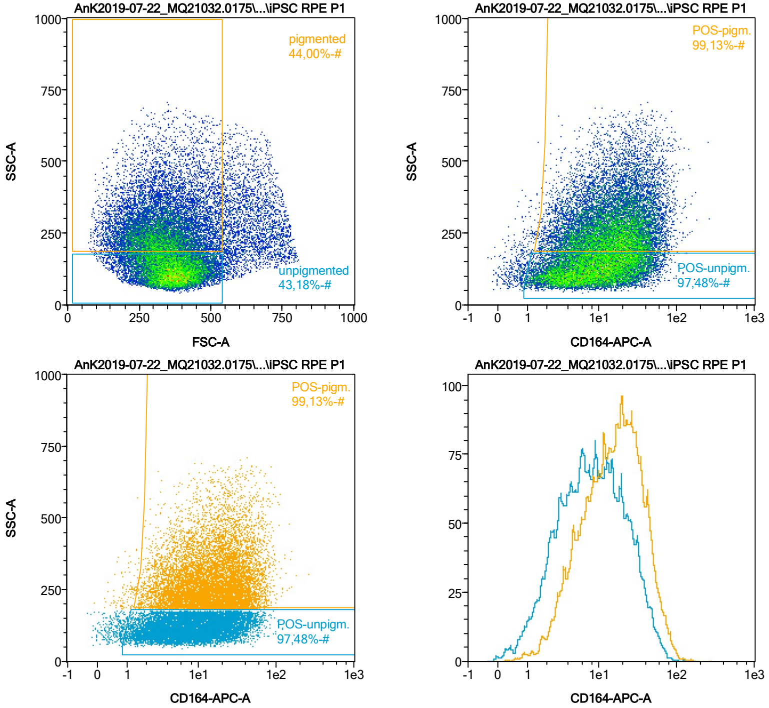


| Name | %-# | CD164-APC-A Mean | CD164-APC-A Median |
|---|---|---|---|
| POS-unpigm. | 97,48 | 12,49 | 8,39 |
| POS-pigm. | 99,13 | 20,30 | 15,76 |

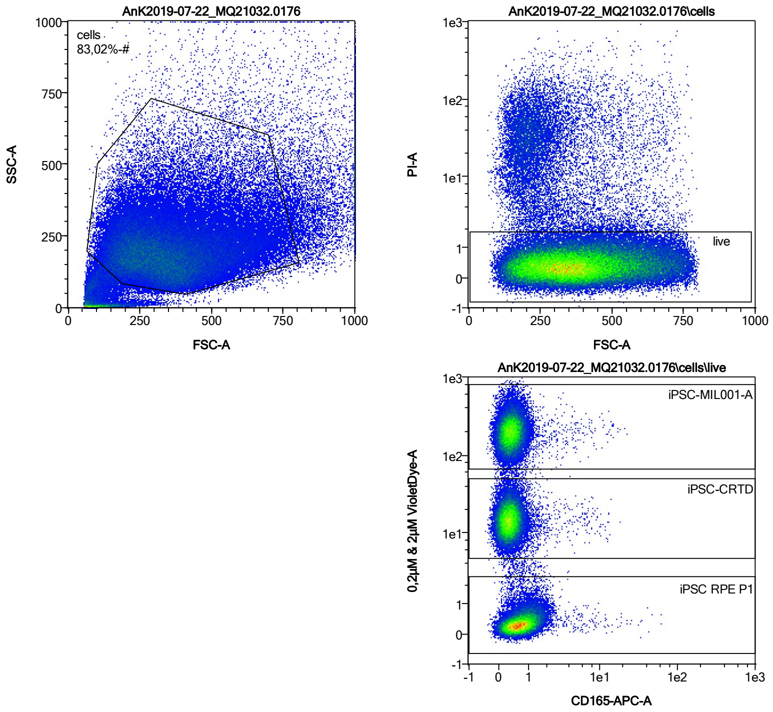


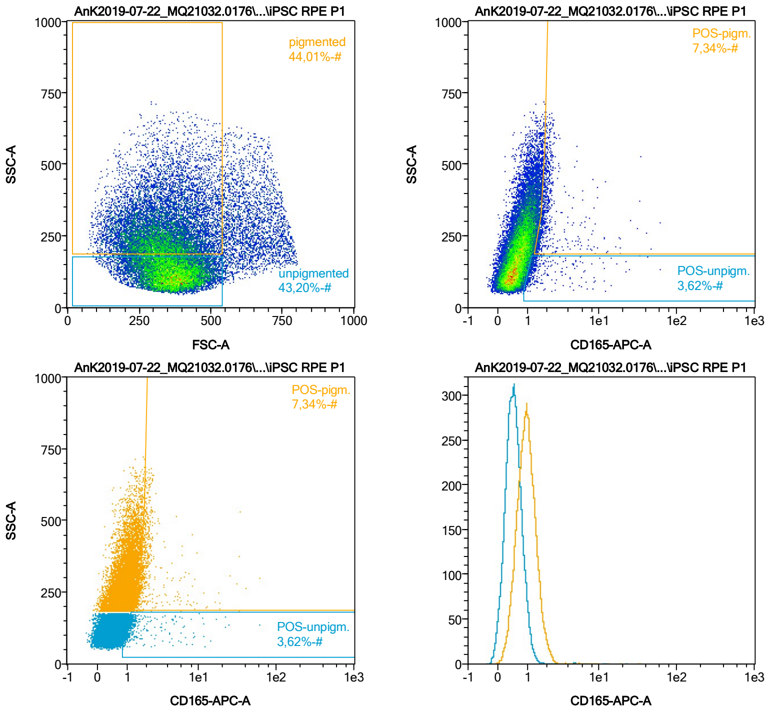


| Name | %-# | CD165-APC-A Mean | CD165-APC-A Median |
|---|---|---|---|
| POS-unpigm. | 3,62 | 2,12 | 1,20 |
| POS-pigm. | 7,34 | 2,48 | 1,83 |

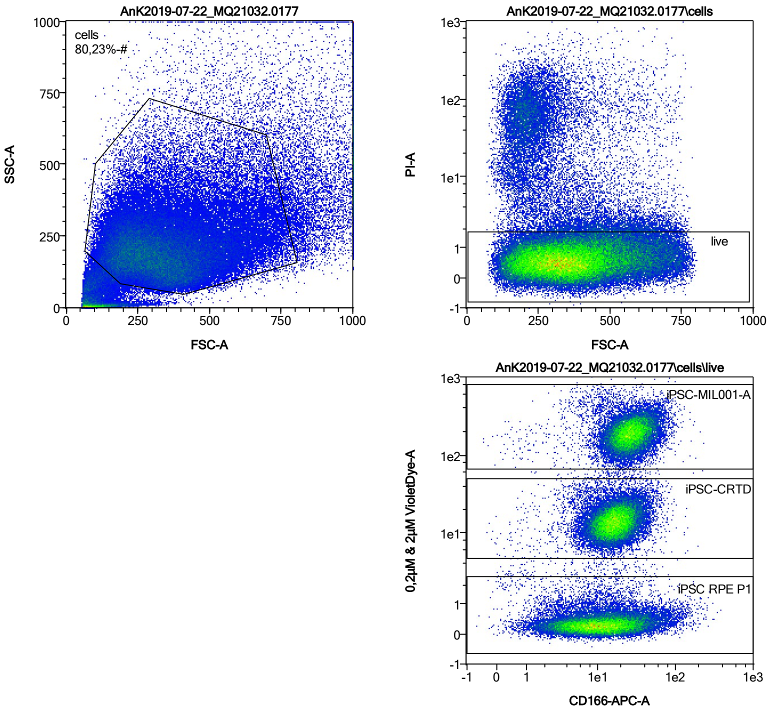


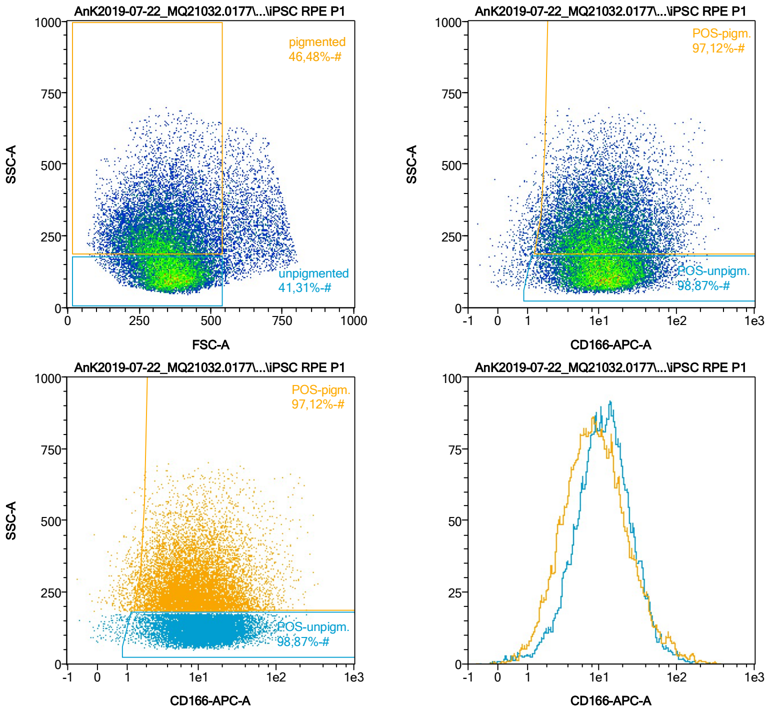


| Name | %-# | CD166-APC-A Mean | CD166-APC-A Median |
|---|---|---|---|
| POS-unpigm. | 98,87 | 15,68 | 11,57 |
| POS-pigm. | 97,12 | 14,91 | 8,97 |

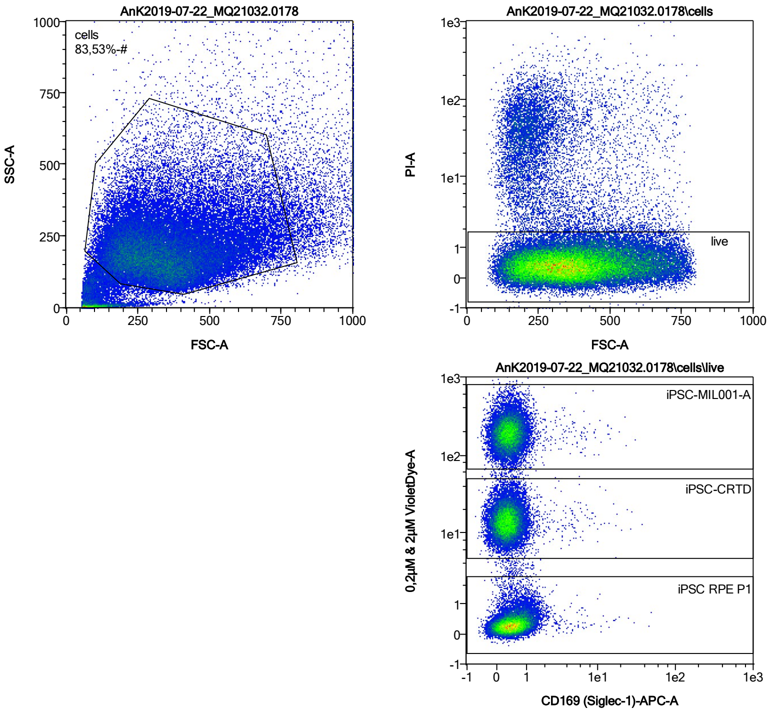


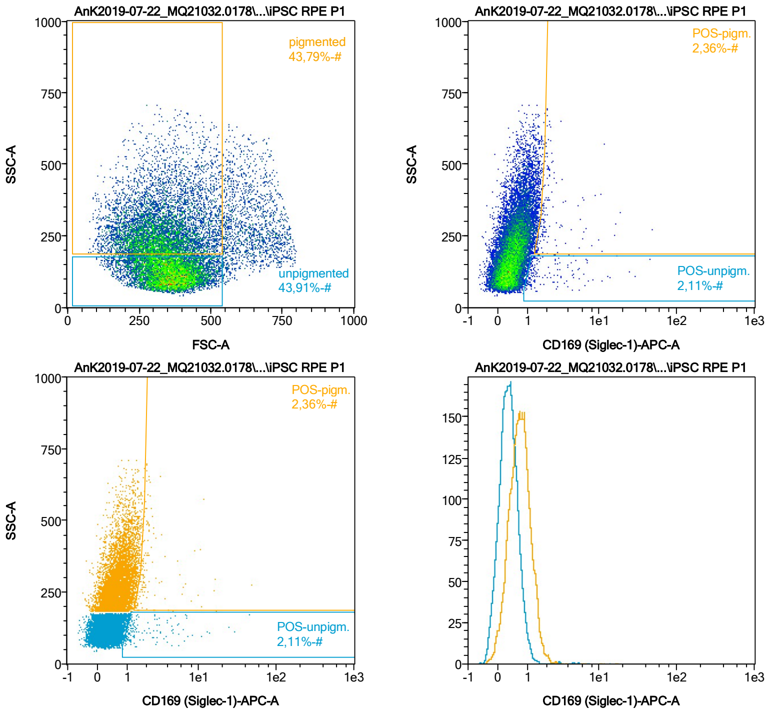


| Name | %-# | CD169 (Siglec-1)-APC-A Mean | CD169 (Siglec-1)-APC-A Median |
|---|---|---|---|
| POS-unpigm. | 2,11 | 2,94 | 1,30 |
| POS-pigm. | 2,36 | 2,93 | 1,87 |

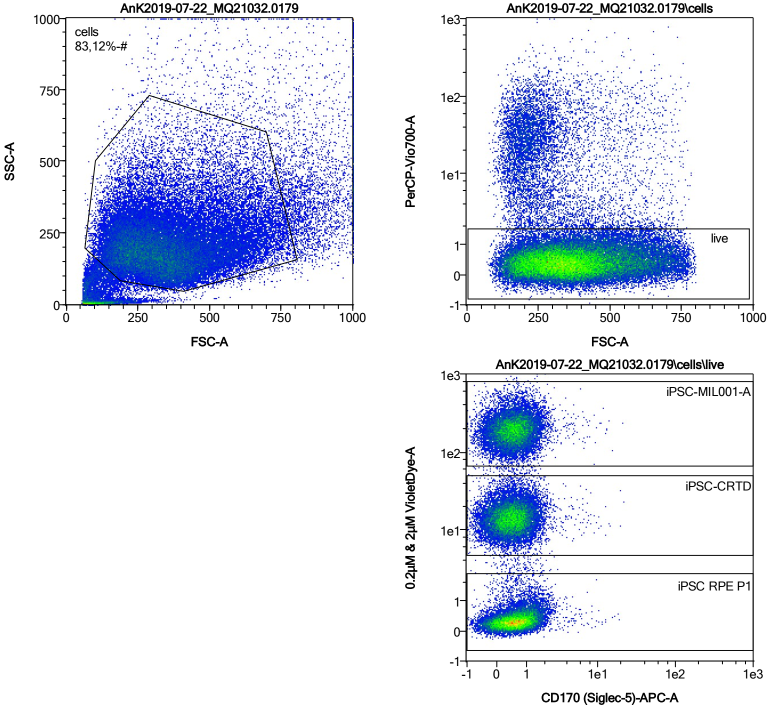


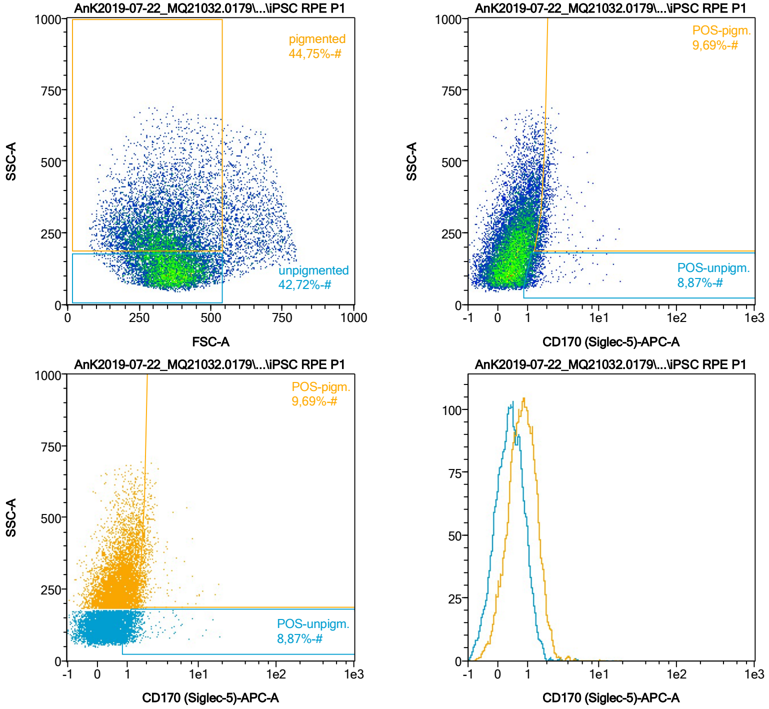


| Name | %-# | CD170 (Siglec-5)-APC-A Mean | CD170 (Siglec-5)-APC-A Median |
|---|---|---|---|
| POS-unpigm. | 8,87 | 1,51 | 1,22 |
| POS-pigm. | 9,69 | 2,10 | 1,84 |

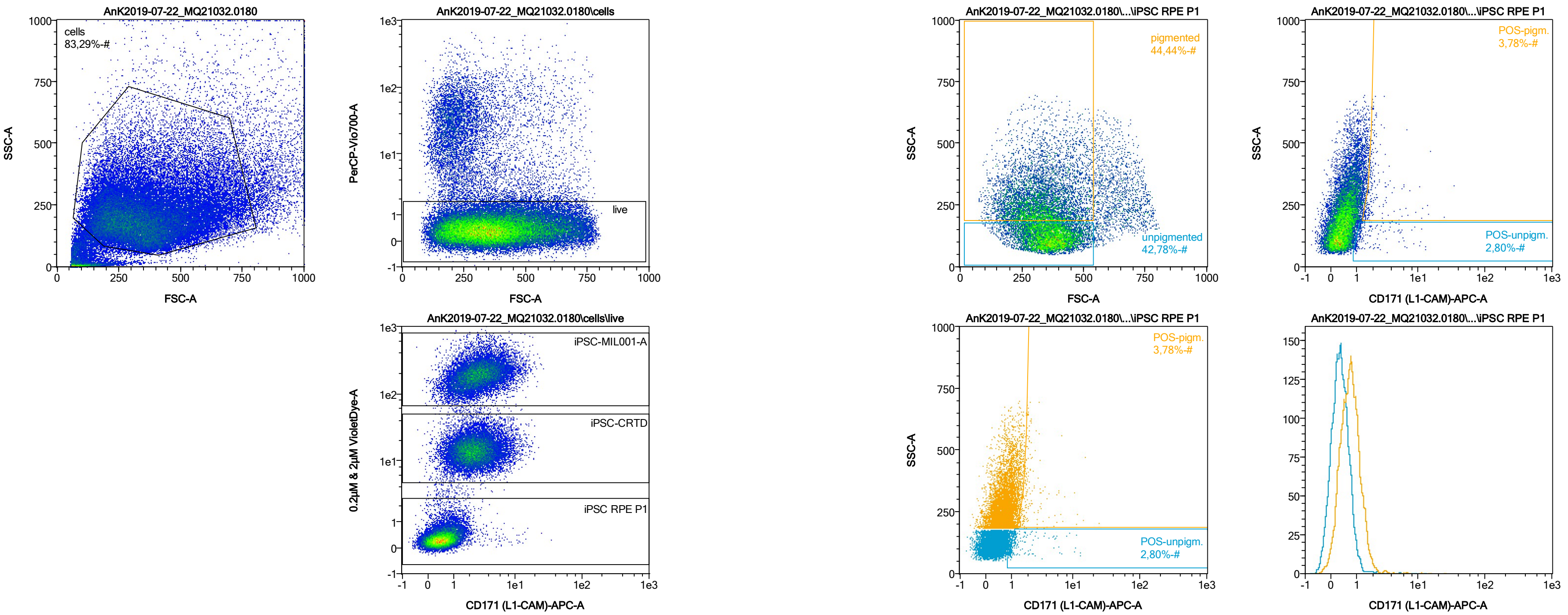


| Name | %-# | CD171 (L1-CAM)-APC-A Mean | CD171 (L1-CAM)-APC-A Median |
|---|---|---|---|
| POS-unpigm. | 2,80 | 2,00 | 1,20 |
| POS-pigm. | 3,78 | 2,46 | 1,86 |

Date:2024-06-14

Operator: Sebastian Knoebel

Version:2.13.0+54.g95abf0f4

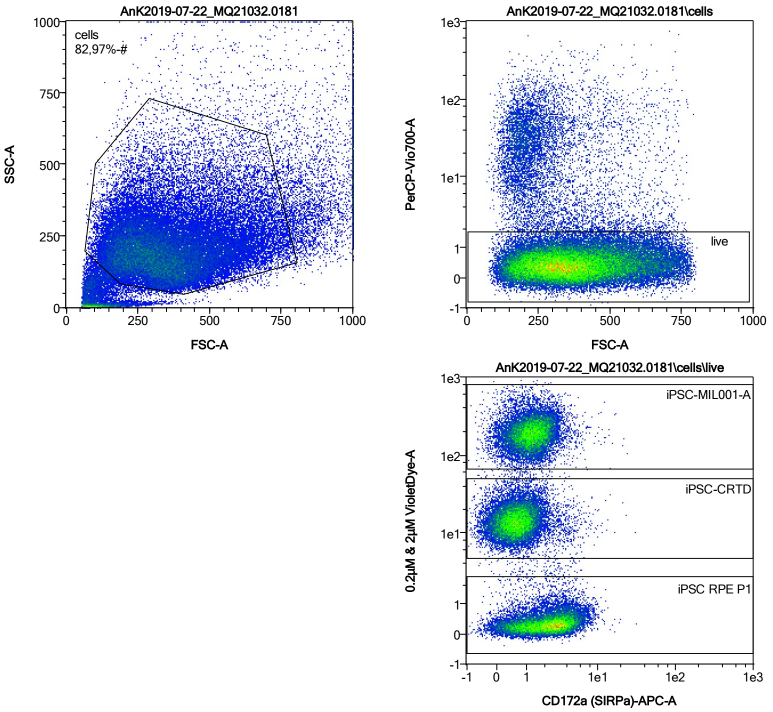


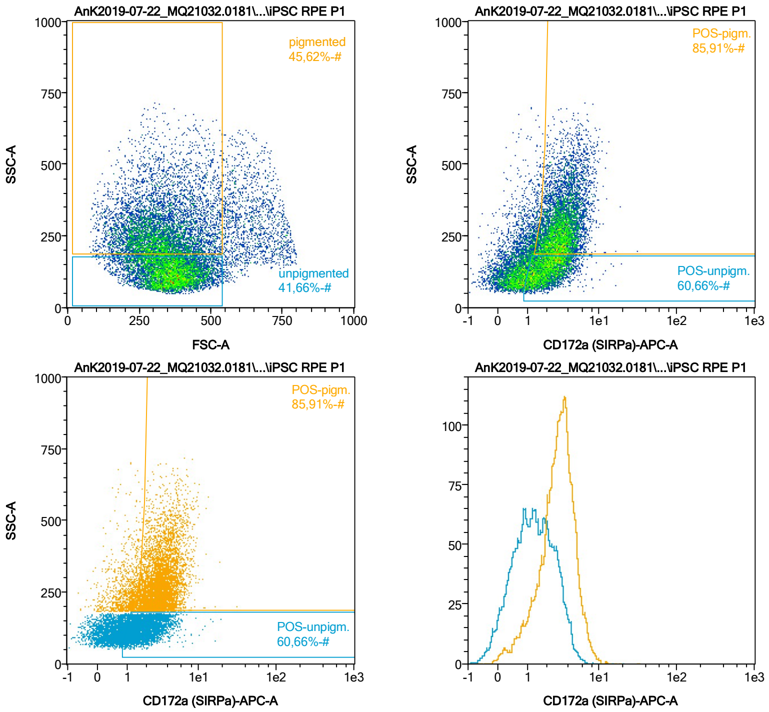


| Name | %-# | CD172a (SIRPa)-APC-A Mean | CD172a (SIRPa)-APC-A Median |
|---|---|---|---|
| POS-unpigm. | 60,66 | 2,15 | 1,91 |
| POS-pigm. | 85,91 | 3,44 | 3,19 |

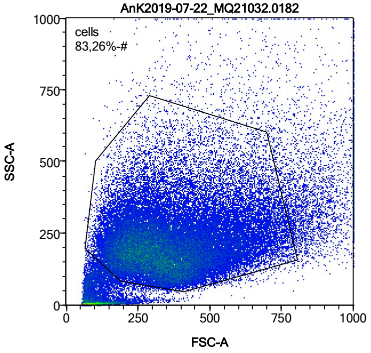


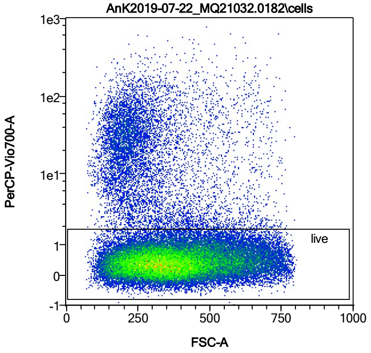


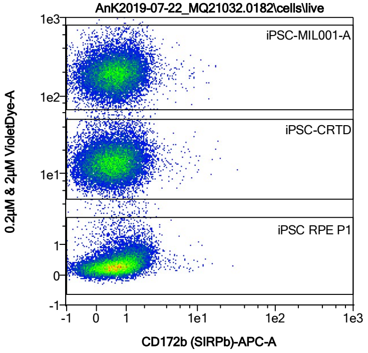


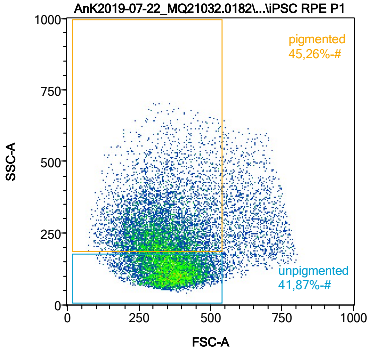


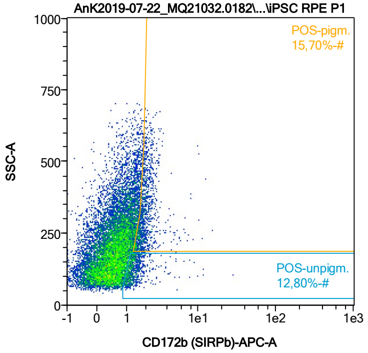


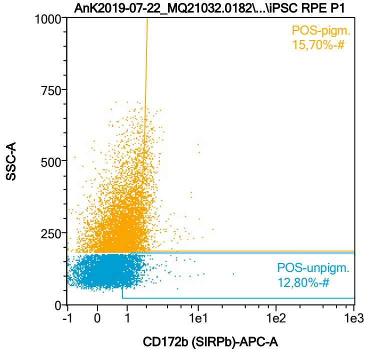


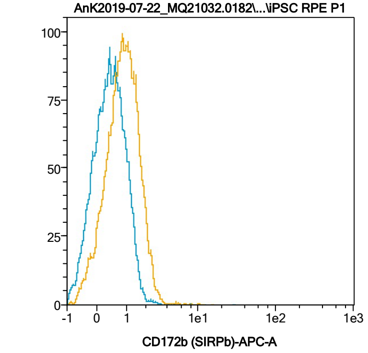


| Name | %-# | CD172b (SIRPb)-APC-A Mean | CD172b (SIRPb)-APC-A Median |
|---|---|---|---|
| POS-unpigm. | 12,80 | 1,46 | 1,24 |
| POS-pigm. | 15,70 | 2,08 | 1,88 |

Date:2024-06-14

Operator: Sebastian Knoebel

Version:2.13.0+54.g95abf0f4

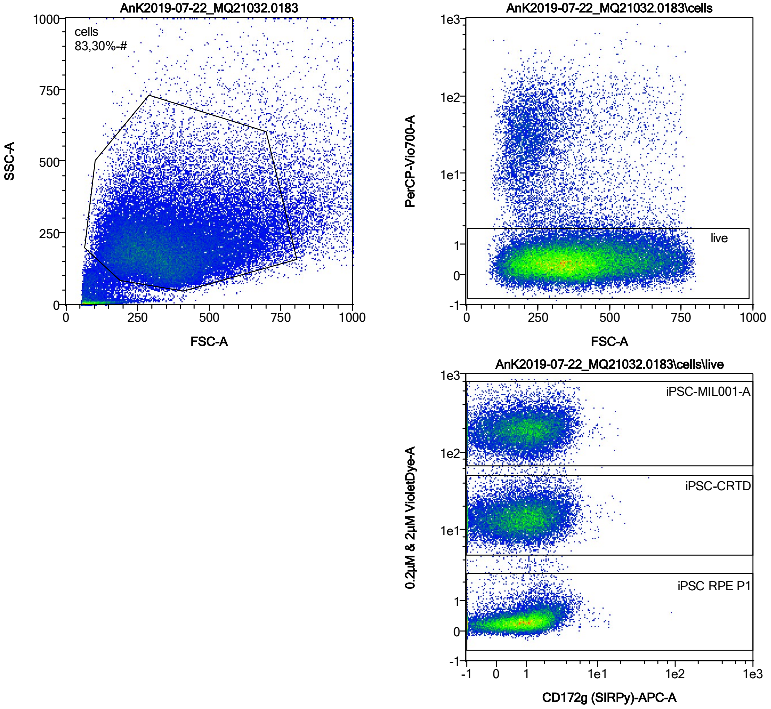


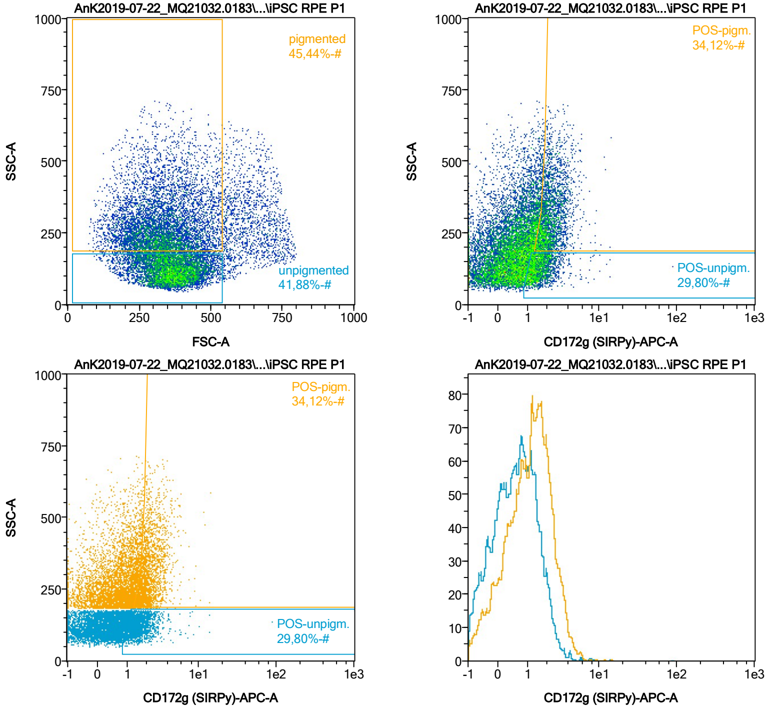


| Name | %-# | CD172g (SIRPγ)-APC-A Mean | CD172g (SIRPγ)-APC-A Median |
|---|---|---|---|
| POS-unpigm. | 29,80 | 1,70 | 1,45 |
| POS-pigm. | 34,12 | 2,36 | 2,11 |

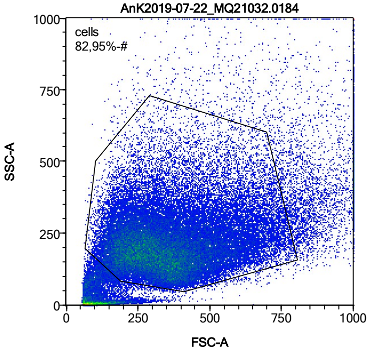


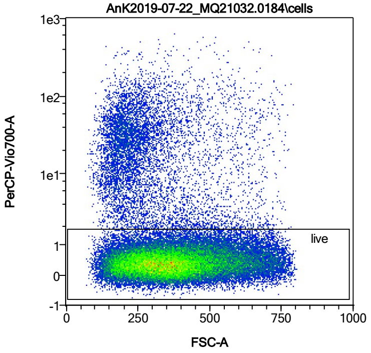


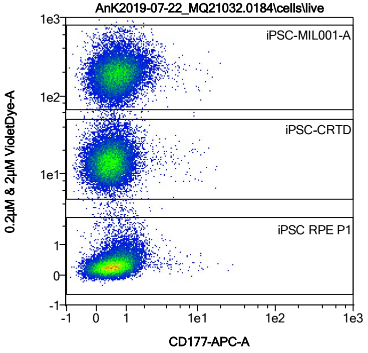


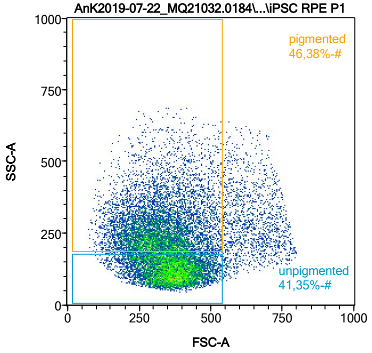


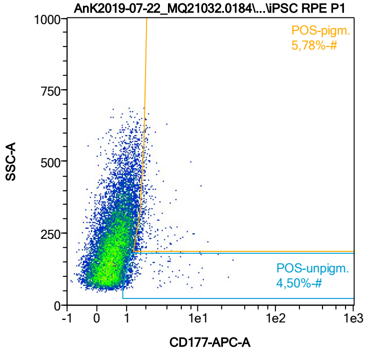


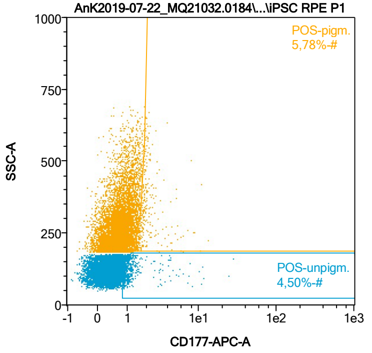


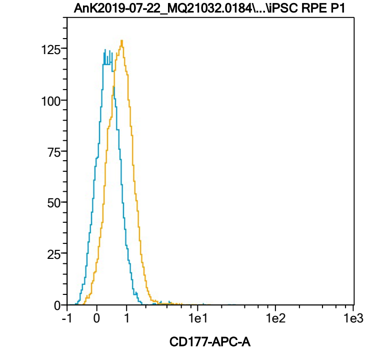


| Name | %-# | CD177-APC-A Mean | CD177-APC-A Median |
|---|---|---|---|
| POS-unpigm. | 4,50 | 1,91 | 1,19 |
| POS-pigm. | 5,78 | 2,16 | 1,81 |

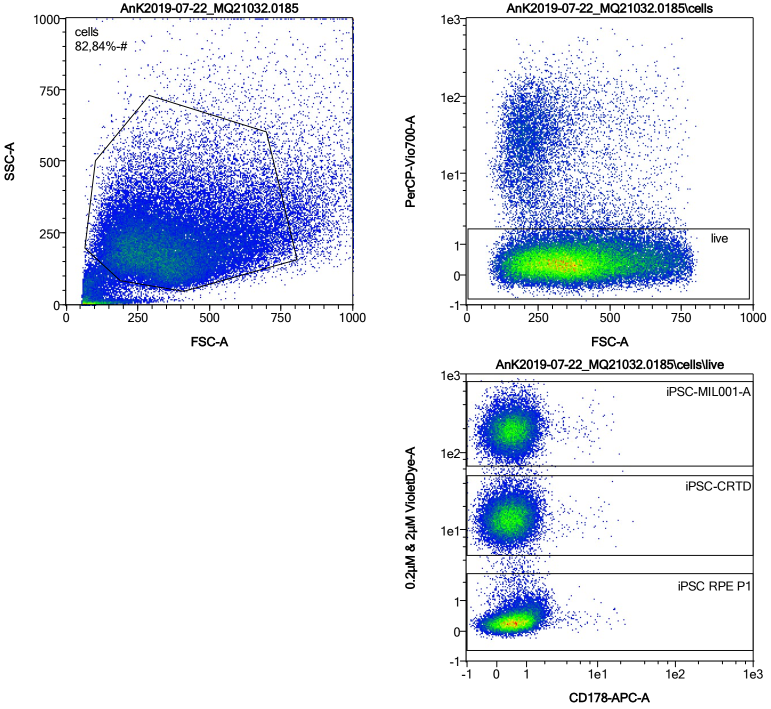


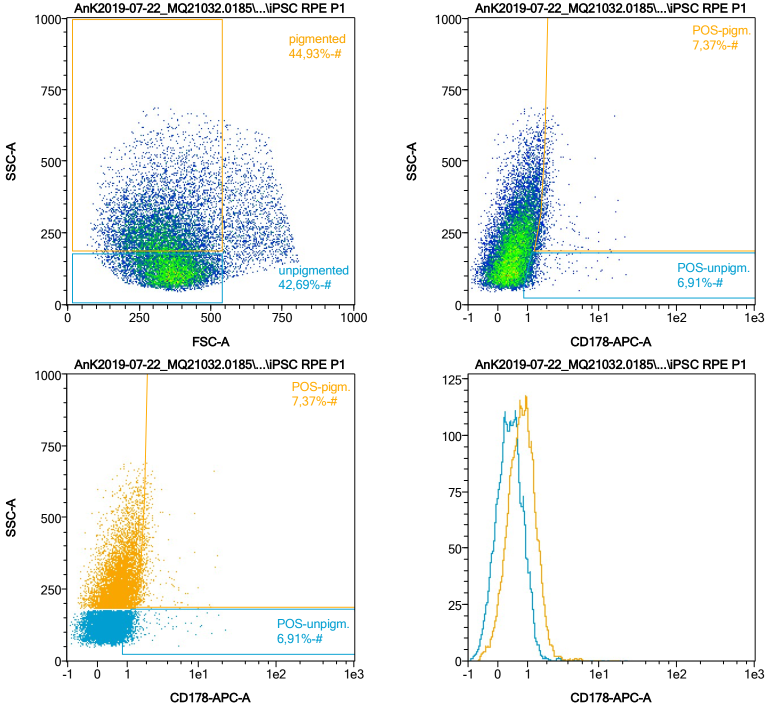


| Name | %-# | CD178-APC-A Mean | CD178-APC-A Median |
|---|---|---|---|
| POS-unpigm. | 6,91 | 1,61 | 1,18 |
| POS-pigm. | 7,37 | 2,26 | 1,84 |

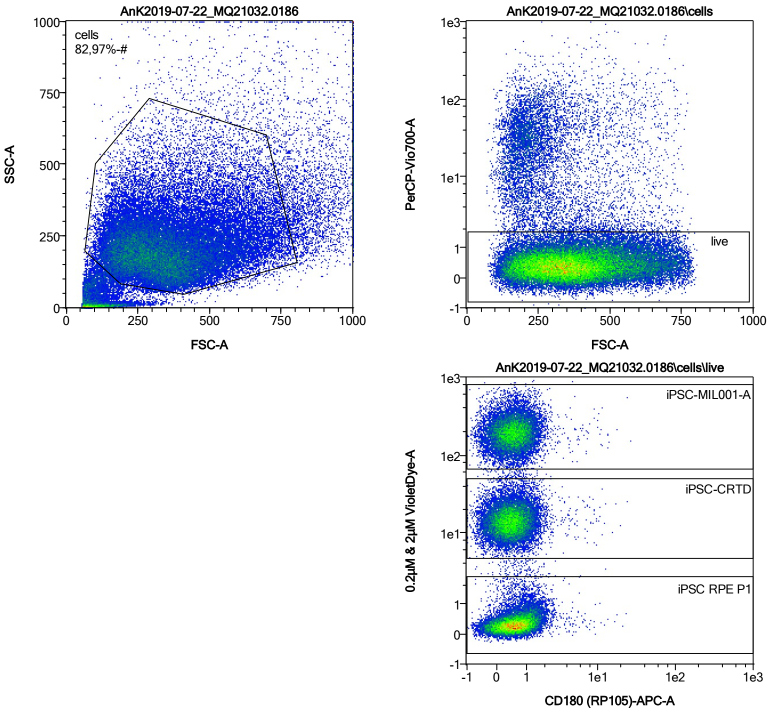


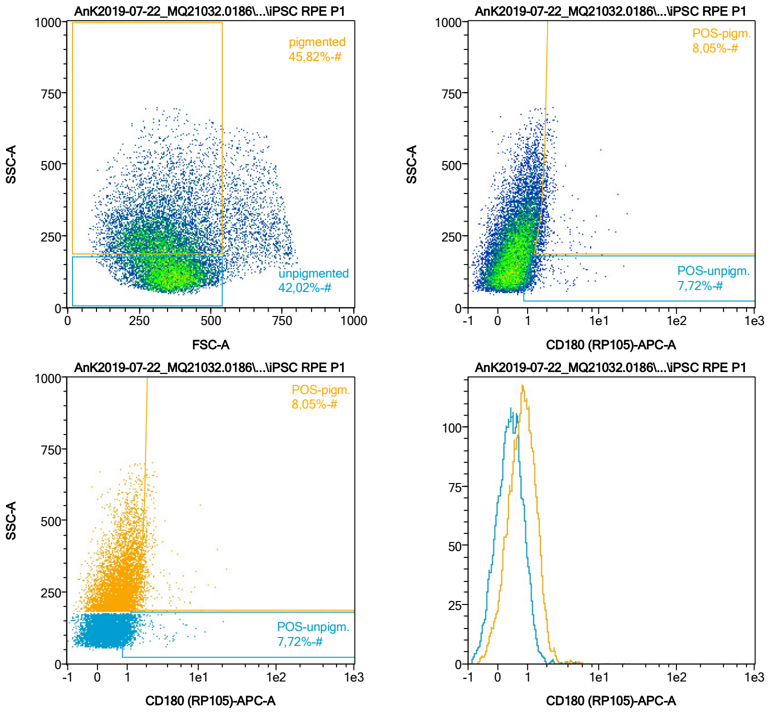


| Name | %-# | CD180 (RP105)-APC-A Mean | CD180 (RP105)-APC-A Median |
|---|---|---|---|
| POS-unpigm. | 7,72 | 1,54 | 1,20 |
| POS-pigm. | 8,05 | 2,21 | 1,84 |

Date:2024-06-14

Operator: Sebastian Knoebel

Version:2.13.0+54.g95abf0f4

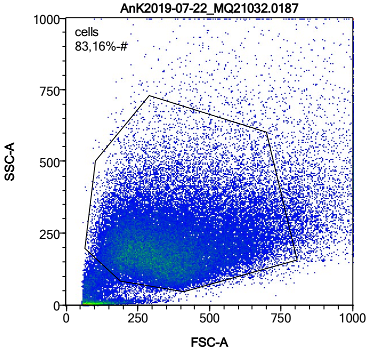


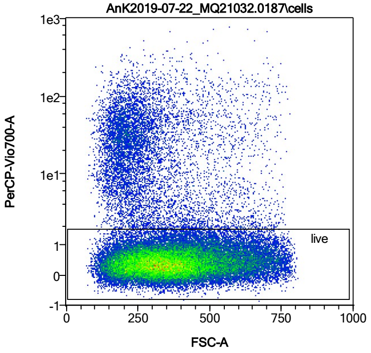


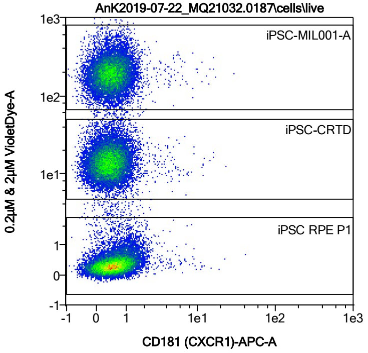


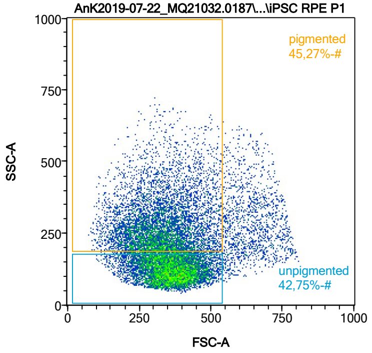


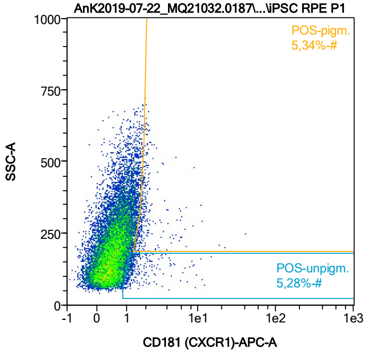


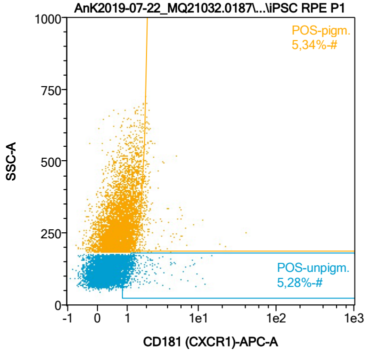


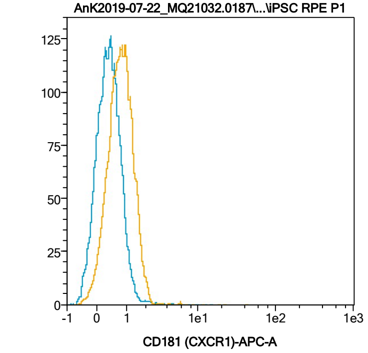


| Name | %-# | CD181 (CXCR1)-APC-A Mean | CD181 (CXCR1)-APC-A Median |
|---|---|---|---|
| POS-unpigm. | 5,28 | 1,78 | 1,22 |
| POS-pigm. | 5,34 | 2,34 | 1,85 |

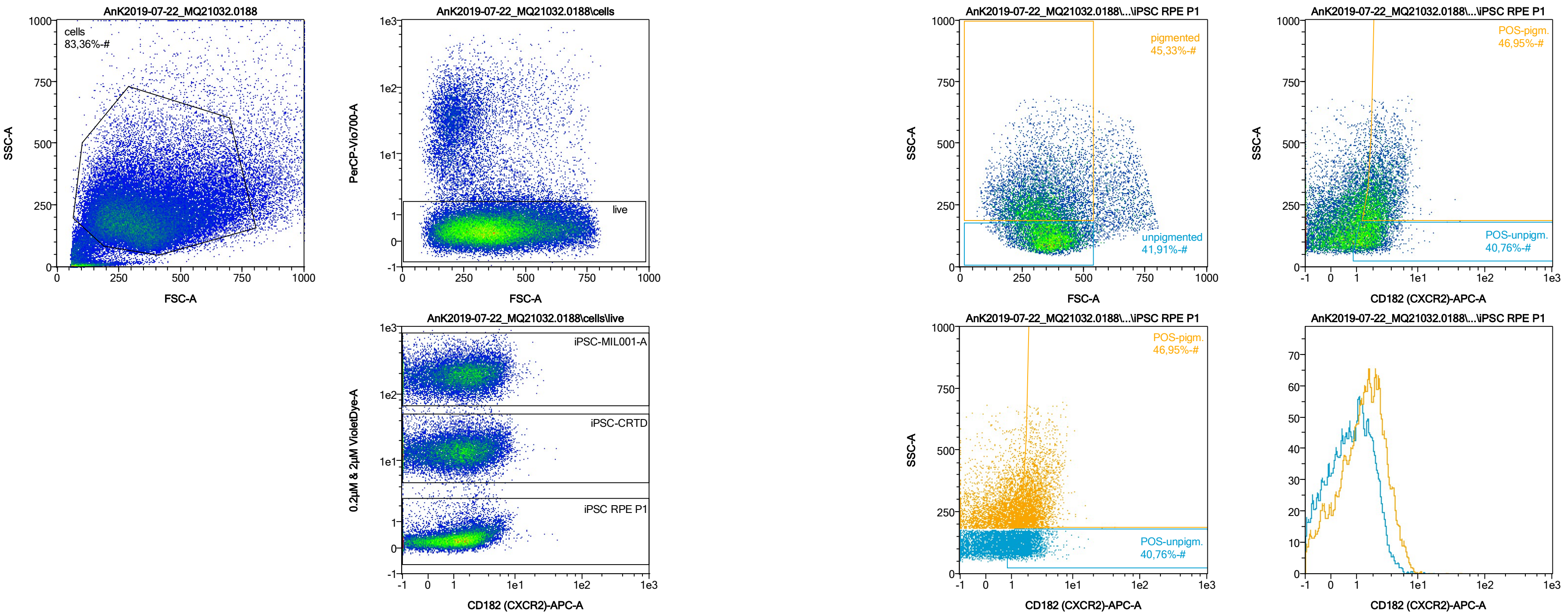


| Name | %-# | CD182 (CXCR2)-APC-A Mean | CD182 (CXCR2)-APC-A Median |
|---|---|---|---|
| POS-unpigm. | 40,76 | 1,97 | 1,73 |
| POS-pigm. | 46,95 | 2,88 | 2,48 |

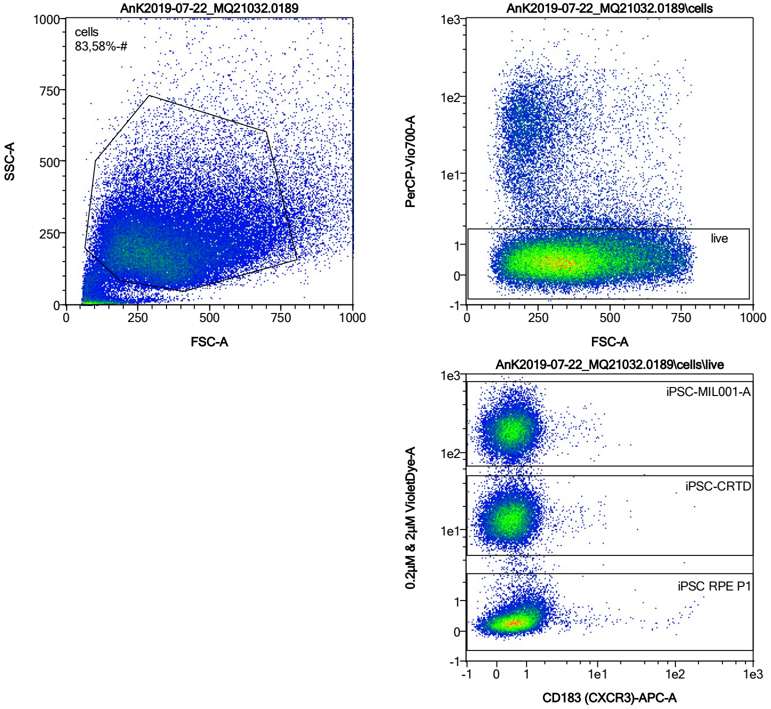


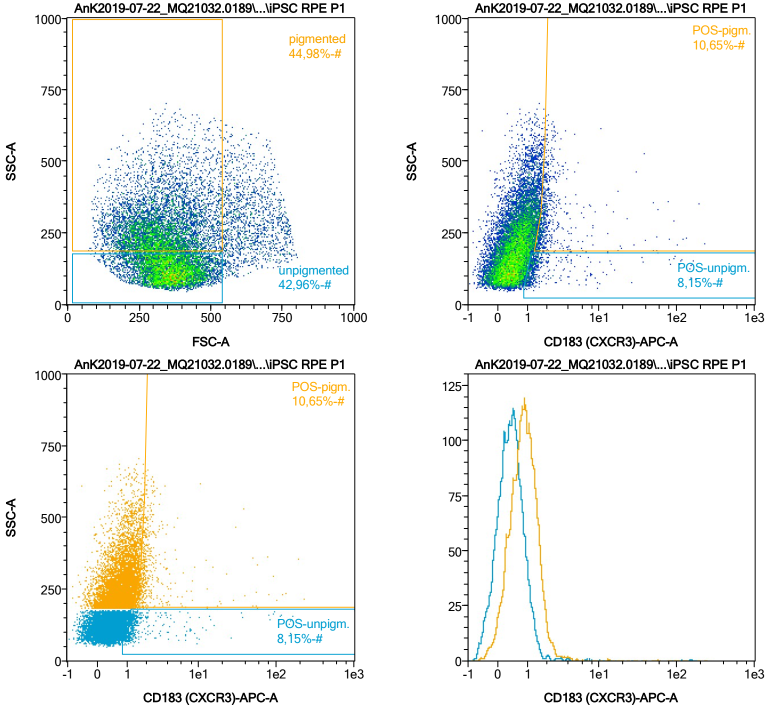


| Name | %-# | CD183 (CXCR3)-APC-A Mean | CD183 (CXCR3)-APC-A Median |
|---|---|---|---|
| POS-unpigm. | 8,15 | 5,21 | 1,25 |
| POS-pigm. | 10,65 | 4,39 | 1,85 |

Date:2024-06-14

Operator: Sebastian Knoebel

Version:2.13.0+54.g95abf0f4

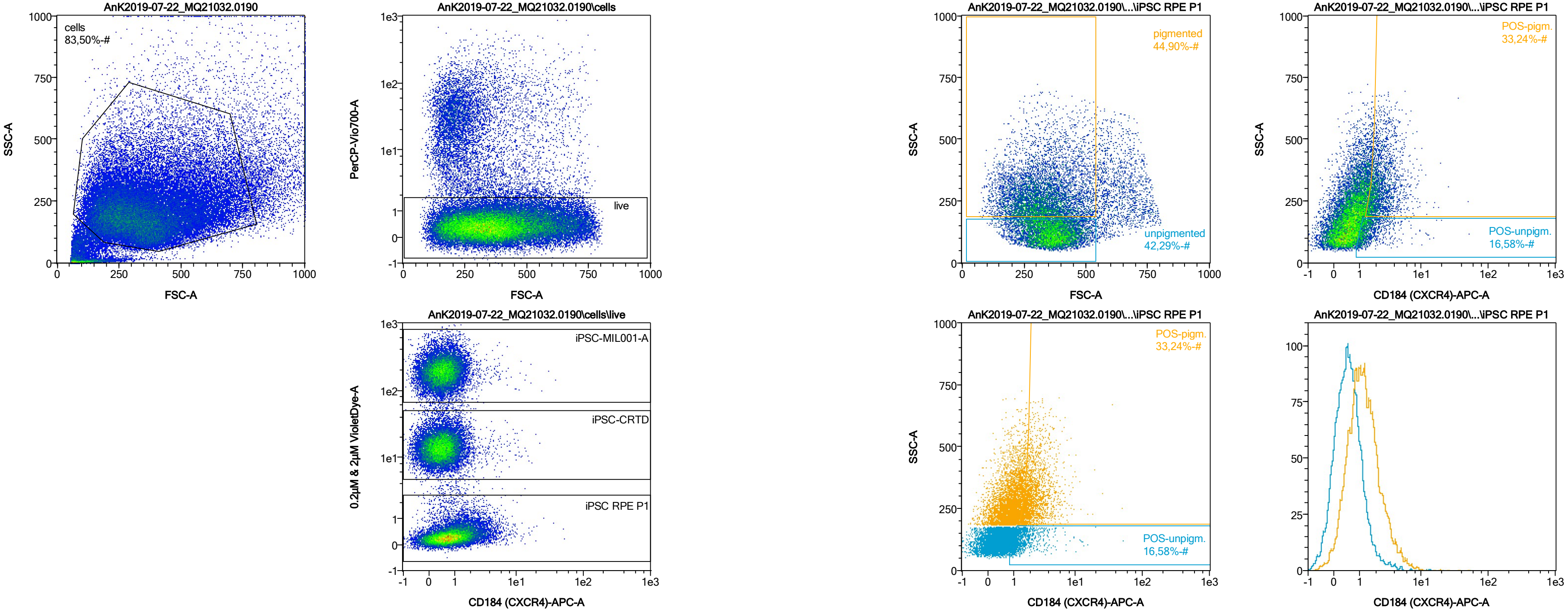


| Name | %-# | CD184 (CXCR4)-APC-A Mean | CD184 (CXCR4)-APC-A Median |
|---|---|---|---|
| POS-unpigm. | 16,58 | 1,77 | 1,35 |
| POS-pigm. | 33,24 | 2,61 | 2,13 |

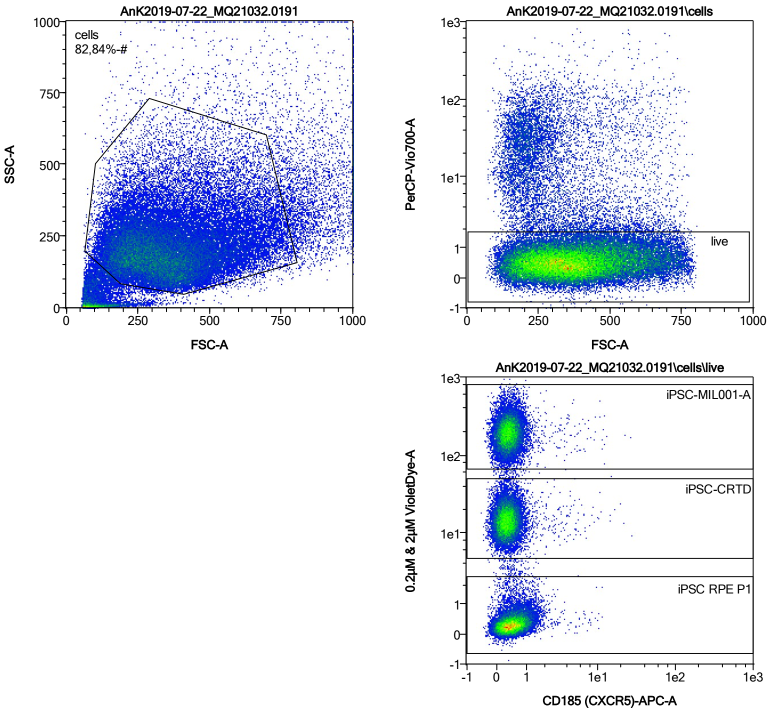


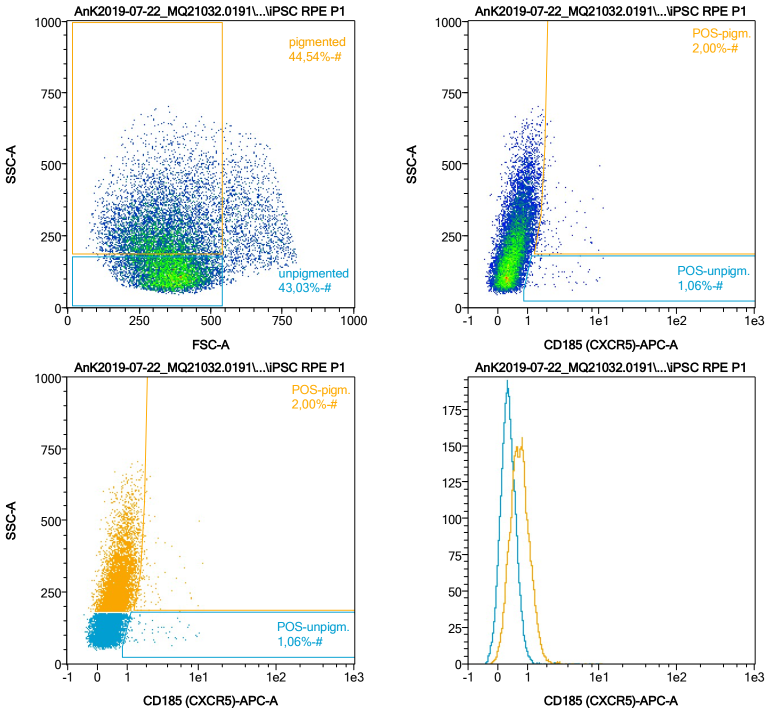


| Name | %-# | CD185 (CXCR5)-APC-A Mean | CD185 (CXCR5)-APC-A Median |
|---|---|---|---|
| POS-unpigm. | 1,06 | 2,73 | 1,62 |
| POS-pigm. | 2,00 | 2,50 | 1,88 |

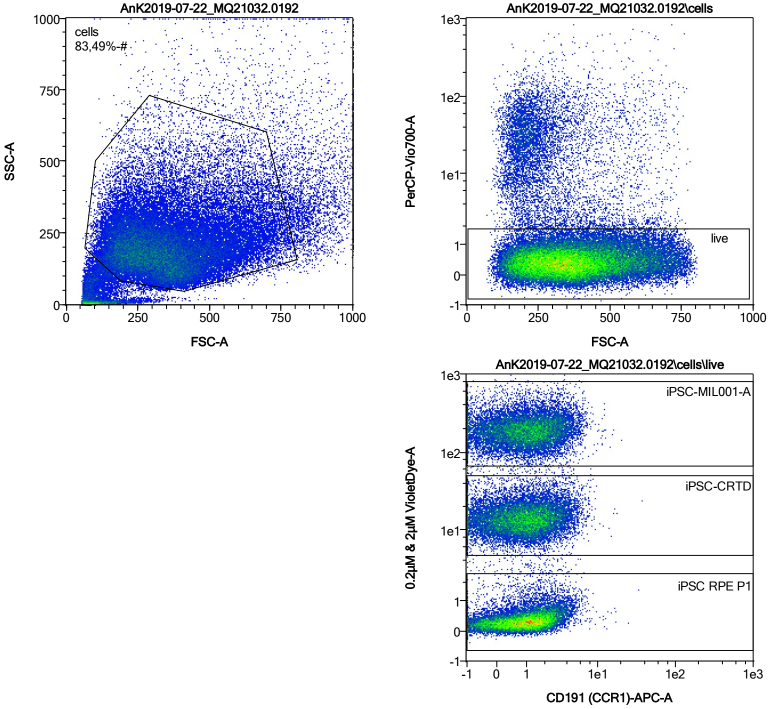


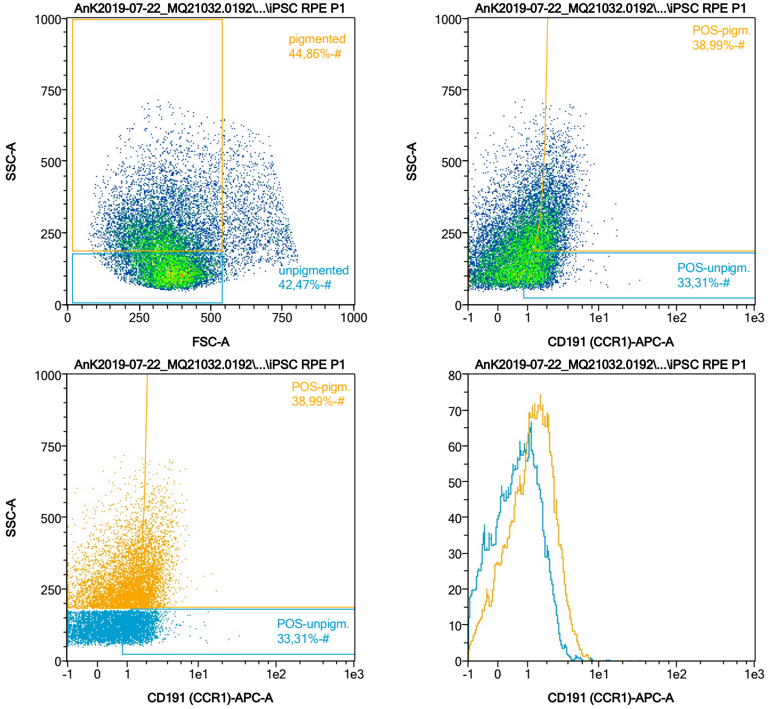


| Name | %-# | CD191 (CCR1)-APC-A Mean | CD191 (CCR1)-APC-A Median |
|---|---|---|---|
| POS-unpigm. | 33,31 | 1,76 | 1,53 |
| POS-pigm. | 38,99 | 2,48 | 2,21 |

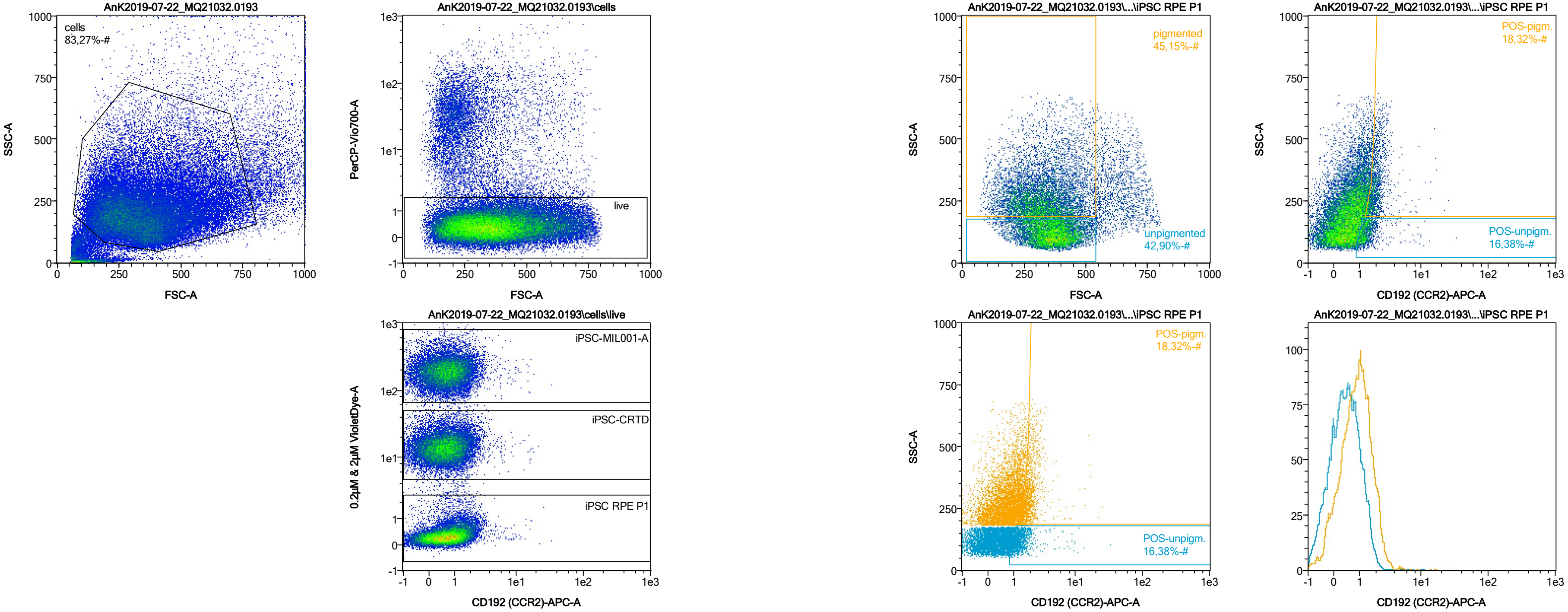


| Name | %-# | CD192 (CCR2)-APC-A Mean | CD192 (CCR2)-APC-A Median |
|---|---|---|---|
| POS-unpigm. | 16,38 | 1,54 | 1,30 |
| POS-pigm. | 18,32 | 2,14 | 1,91 |

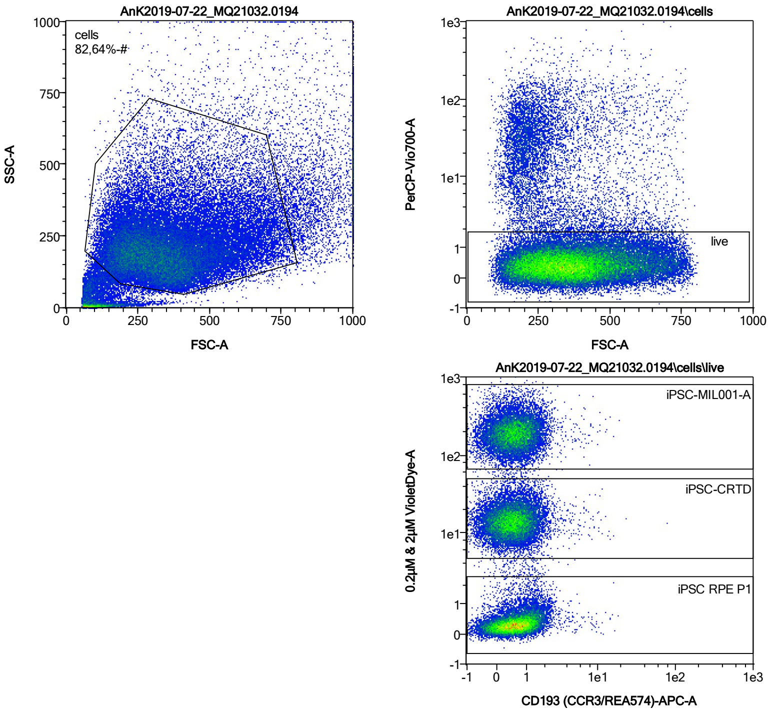


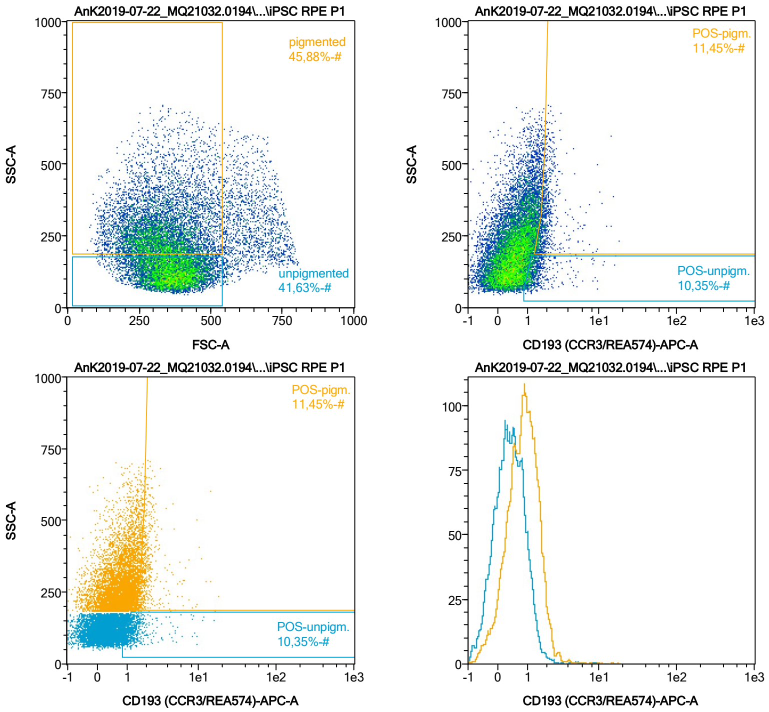


| Name | %-# | CD193 (CCR3/REA574)-APC-A Mean | CD193 (CCR3/REA574)-APC-A Median |
|---|---|---|---|
| POS-unpigm. | 10,35 | 1,53 | 1,23 |
| POS-pigm. | 11,45 | 2,15 | 1,85 |

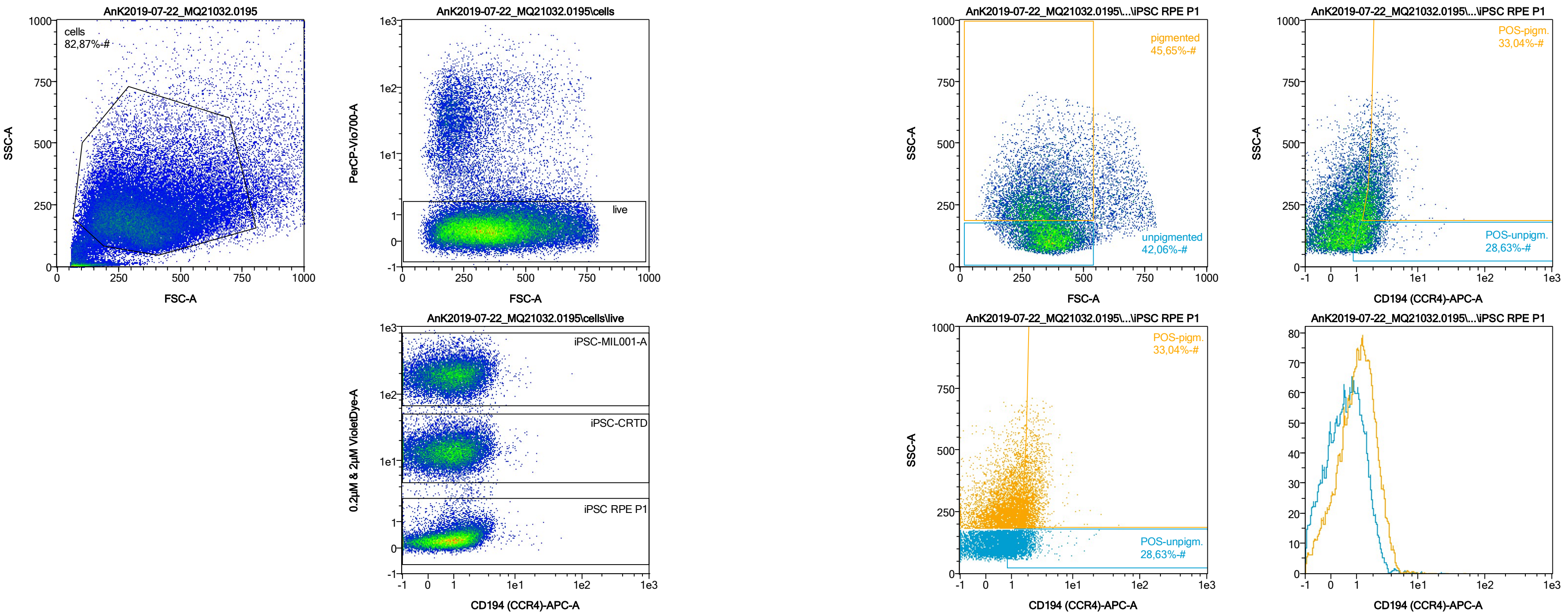


| Name | %-# | CD194 (CCR4)-APC-A Mean | CD194 (CCR4)-APC-A Median |
|---|---|---|---|
| POS-unpigm. | 28,63 | 1,69 | 1,46 |
| POS-pigm. | 33,04 | 2,39 | 2,12 |

Date:2024-06-14
Operator: Sebastian Knoebel
Version:2.13.0+54.g95abf0f4

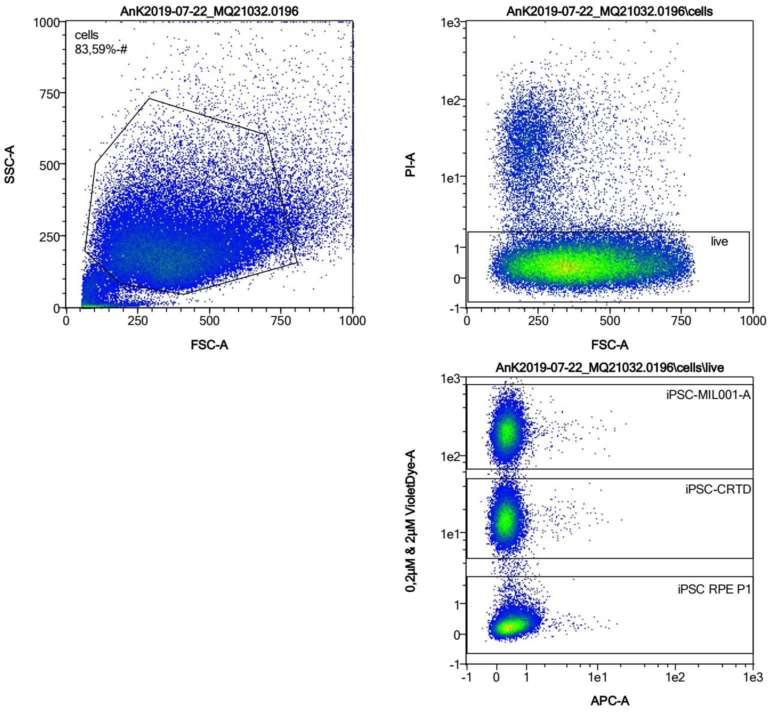


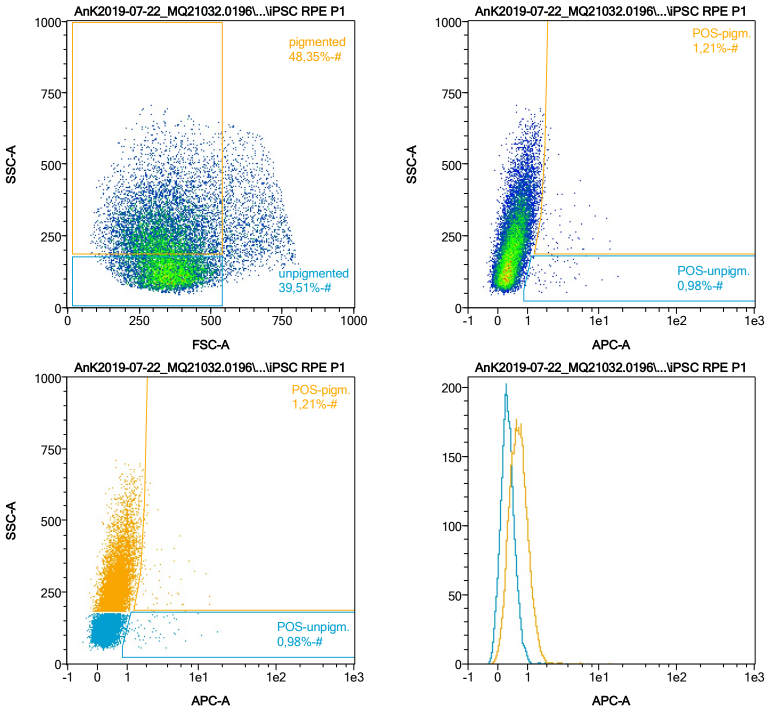


| Name | %-# | APC-A Mean | APC-A Median |
|---|---|---|---|
| POS-unpigm. | 0,98 | 2,95 | 1,98 |
| POS-pigm. | 1,21 | 3,12 | 2,06 |

Date:2024-06-14

Operator: Sebastian Knoebel

Version:2.13.0+54.g95abf0f4

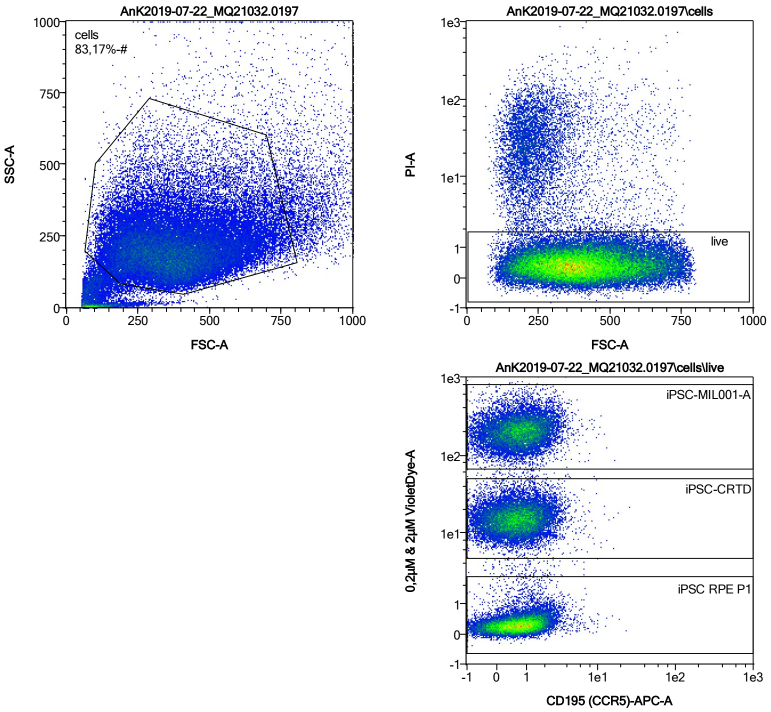


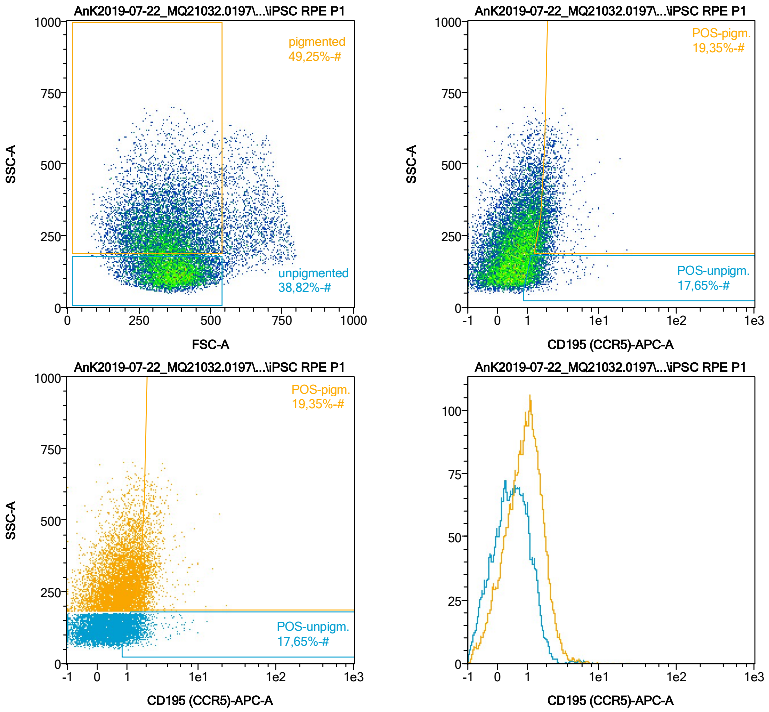


| Name | %-# | CD195 (CCR5)-APC-A Mean | CD195 (CCR5)-APC-A Median |
|---|---|---|---|
| POS-unpigm. | 17,65 | 1,56 | 1,35 |
| POS-pigm. | 19,35 | 2,25 | 1,97 |

Date:2024-06-14

Operator: Sebastian Knoebel

Version:2.13.0+54.g95abf0f4

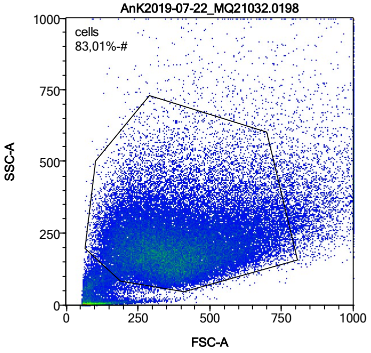


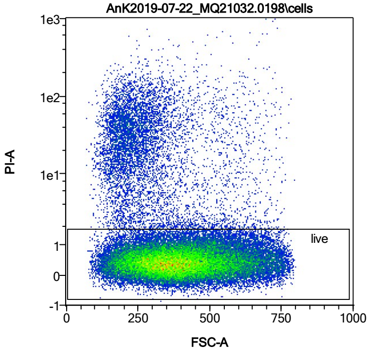


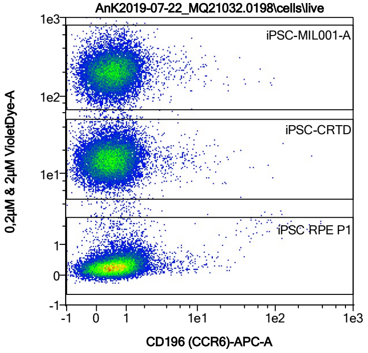


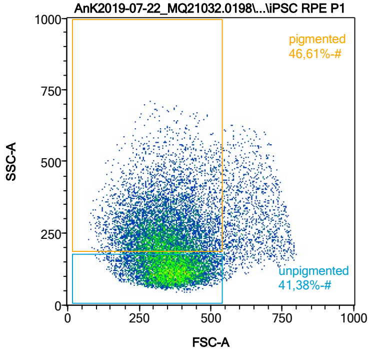


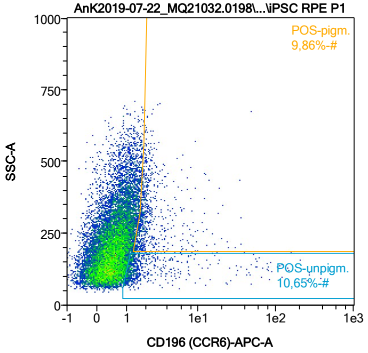


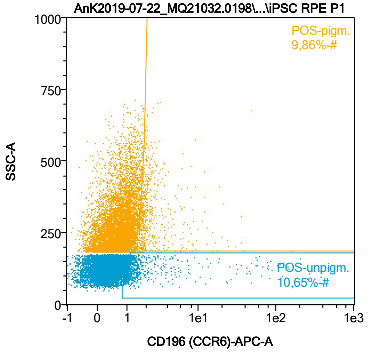


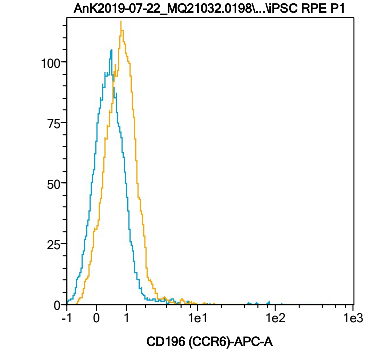


| Name | %-# | CD196 (CCR6)-APC-A Mean | CD196 (CCR6)-APC-A Median |
|---|---|---|---|
| POS-unpigm. | 10,65 | 7,95 | 1,30 |
| POS-pigm. | 9,86 | 3,17 | 1,94 |

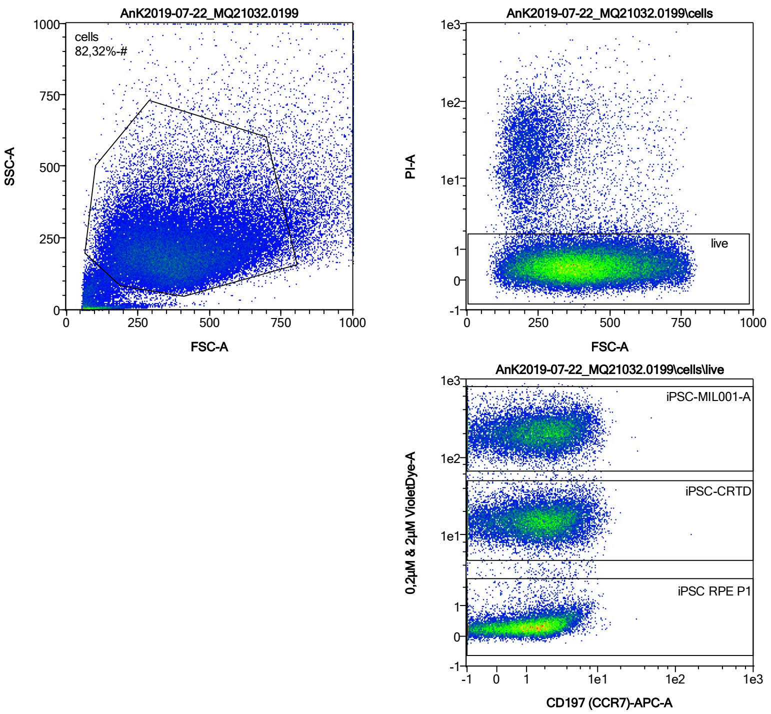


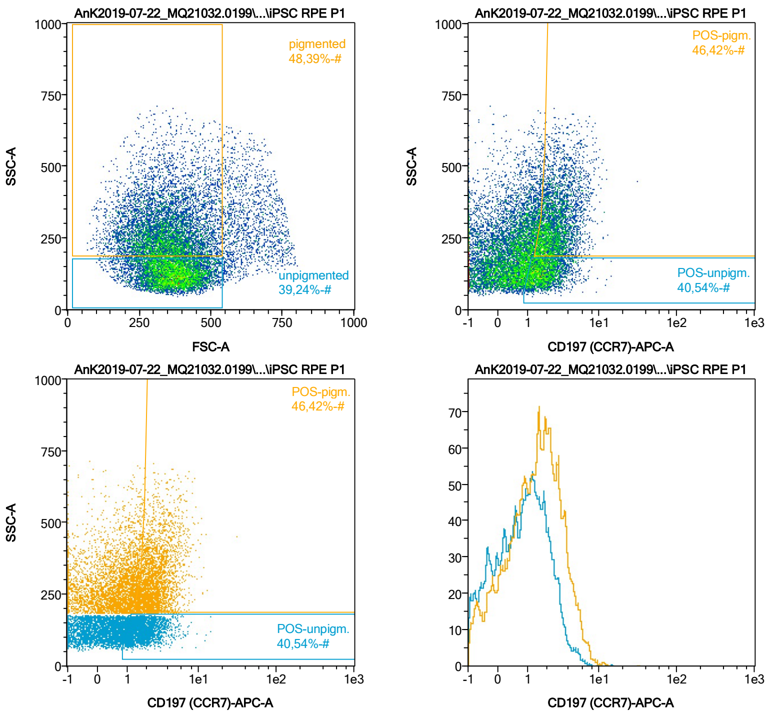


| Name | %-# | CD197 (CCR7)-APC-A Mean | CD197 (CCR7)-APC-A Median |
|---|---|---|---|
| POS-unpigm. | 40,54 | 2,03 | 1,78 |
| POS-pigm. | 46,42 | 2,93 | 2,55 |

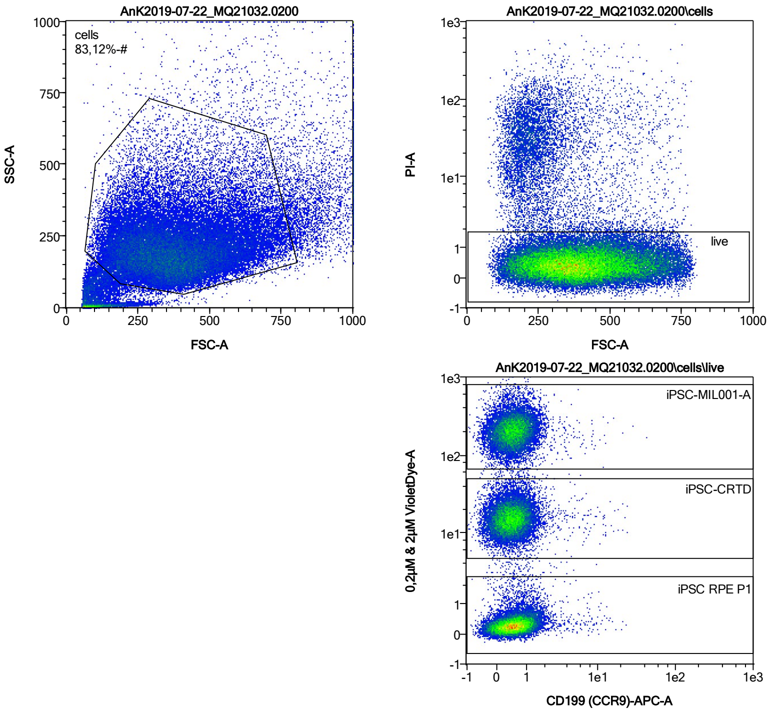


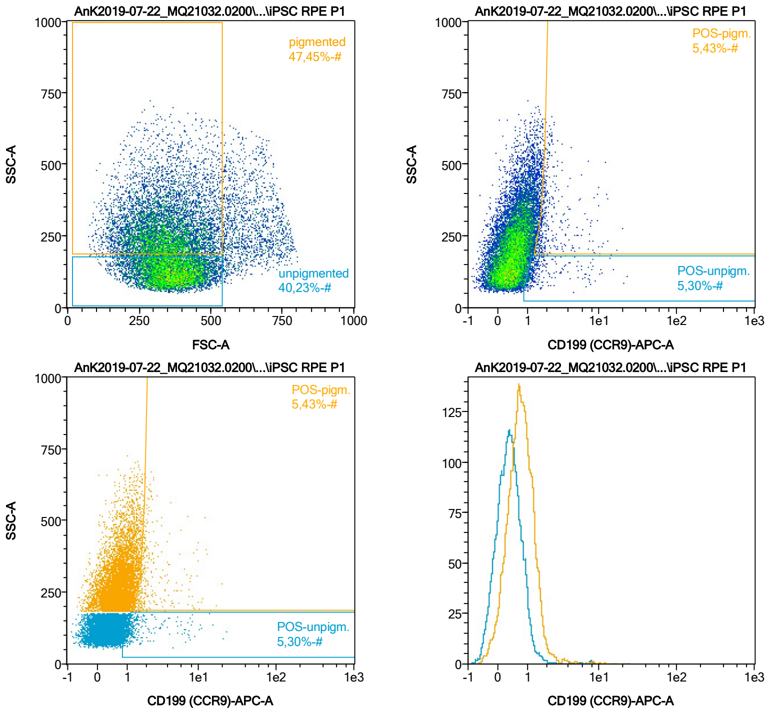


| Name | %-# | CD199 (CCR9)-APC-A Mean | CD199 (CCR9)-APC-A Median |
|---|---|---|---|
| POS-unpigm. | 5,30 | 2,09 | 1,22 |
| POS-pigm. | 5,43 | 2,72 | 1,91 |

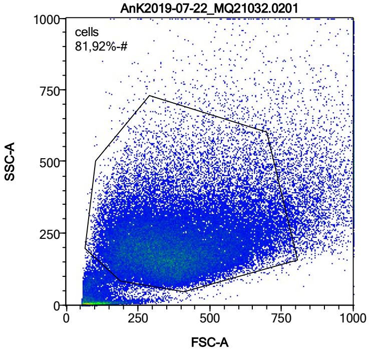


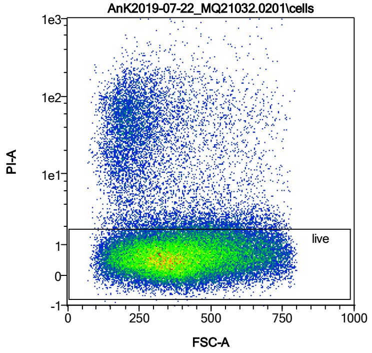


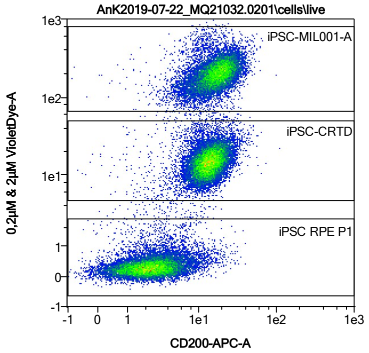


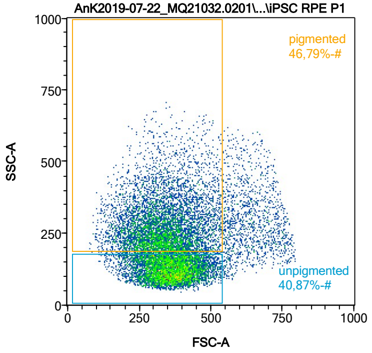


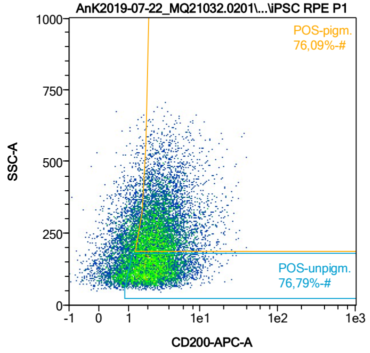


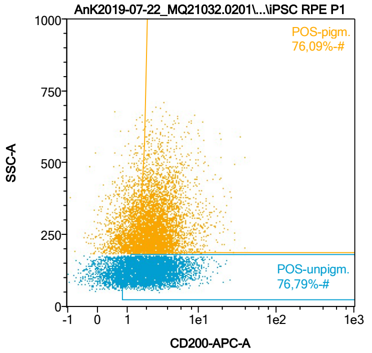


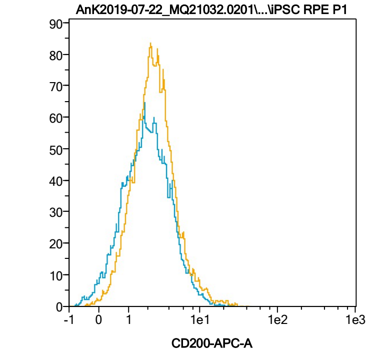


| Name | %-# | CD200-APC-A Mean | CD200-APC-A Median |
|---|---|---|---|
| POS-unpigm. | 76,79 | 2,97 | 2,41 |
| POS-pigm. | 76,09 | 3,57 | 2,84 |

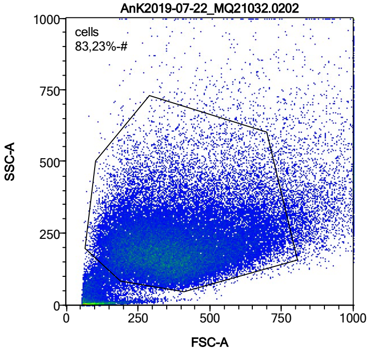


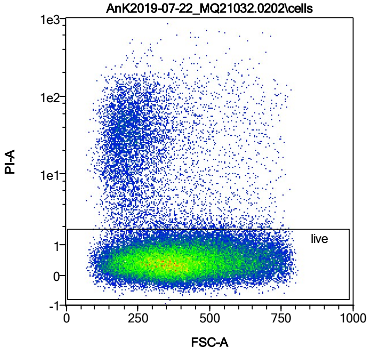


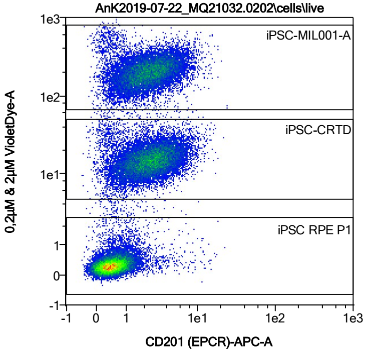


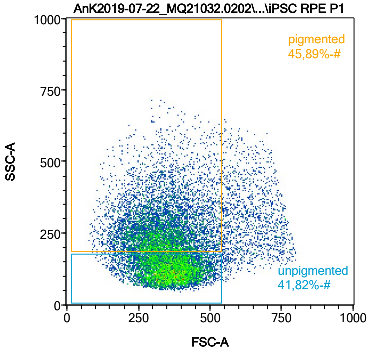


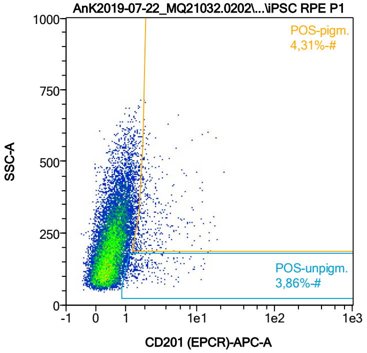


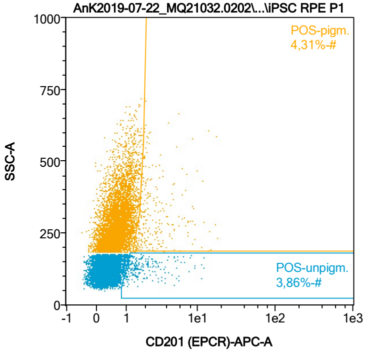


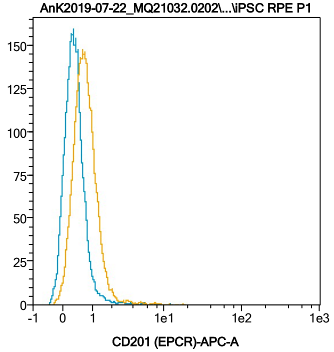


| Name | %-# | CD201 (EPCR)-APC-A Mean | CD201 (EPCR)-APC-A Median |
|---|---|---|---|
| POS-unpigm. | 3,86 | 2,24 | 1,52 |
| POS-pigm. | 4,31 | 3,18 | 2,12 |

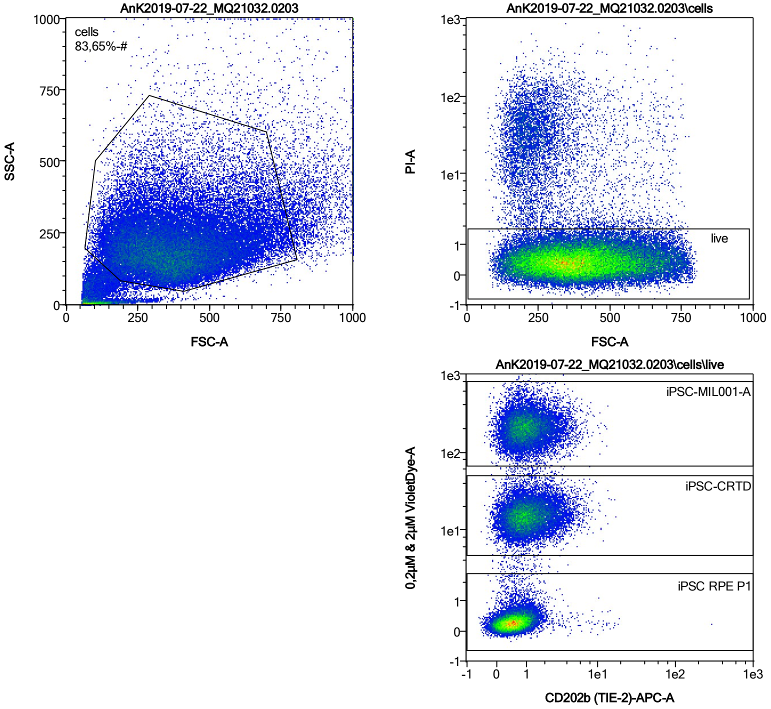


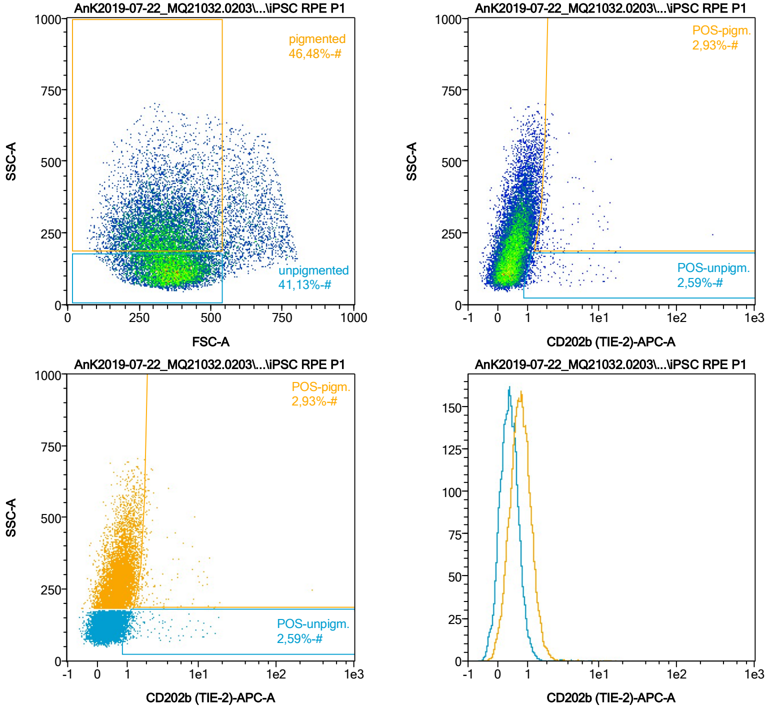


| Name | %-# | CD202b (TIE-2)-APC-A Mean | CD202b (TIE-2)-APC-A Median |
|---|---|---|---|
| POS-unpigm. | 2,59 | 2,49 | 1,26 |
| POS-pigm. | 2,93 | 4,32 | 1,94 |

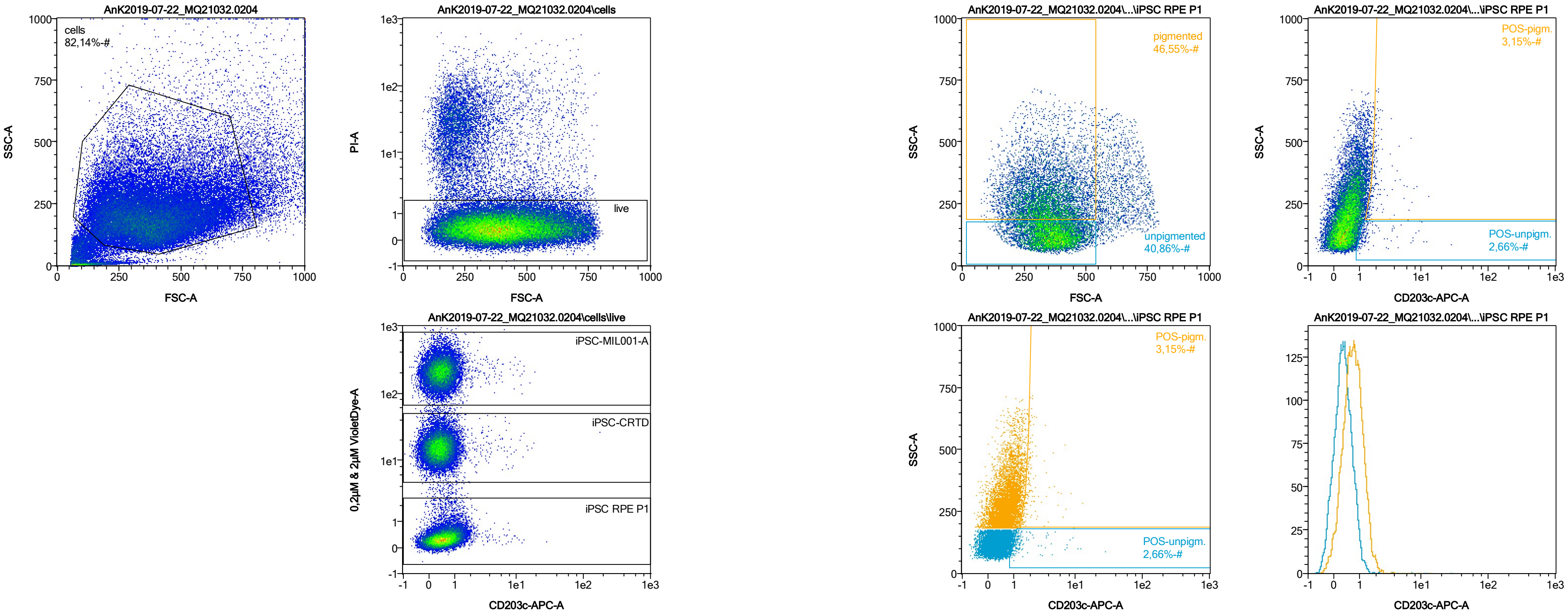


| Name | %-# | CD203c-APC-A Mean | CD203c-APC-A Median |
|---|---|---|---|
| POS-unpigm. | 2,66 | 1,95 | 1,21 |
| POS-pigm. | 3,15 | 2,17 | 1,86 |

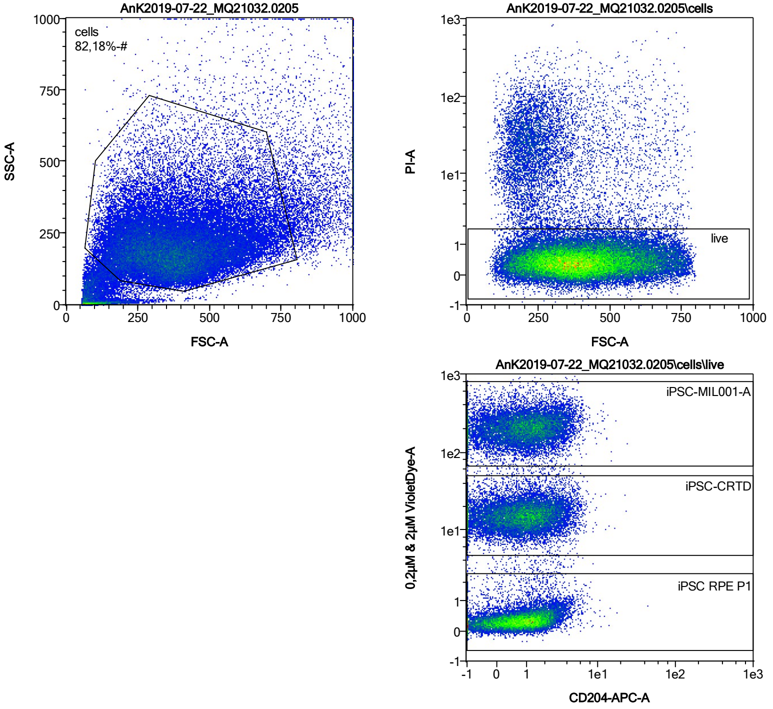


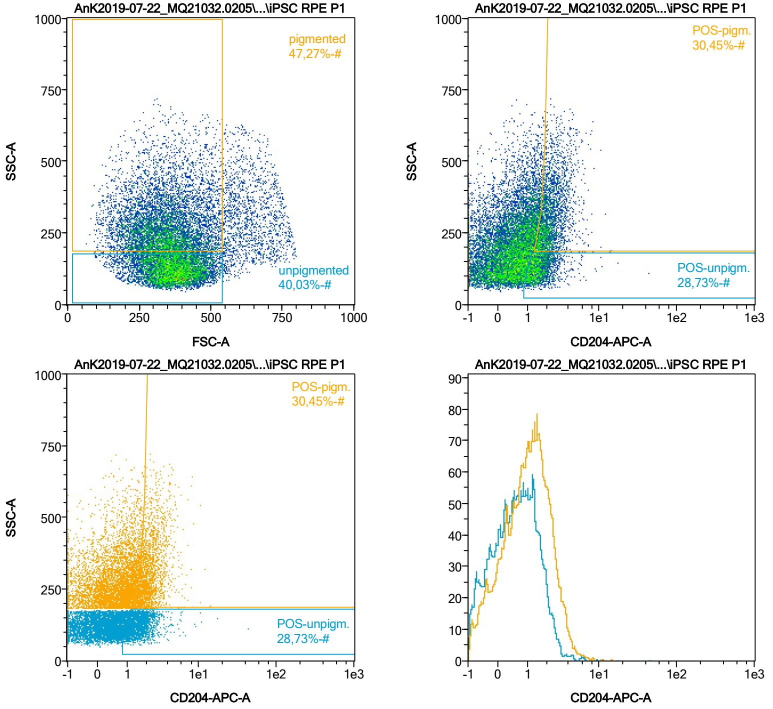


| Name | %-# | CD204-APC-A Mean | CD204-APC-A Median |
|---|---|---|---|
| POS-unpigm. | 28,73 | 1,71 | 1,48 |
| POS-pigm. | 30,45 | 2,39 | 2,14 |

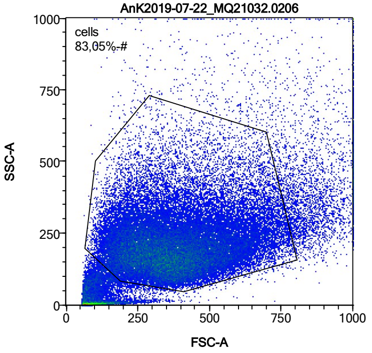


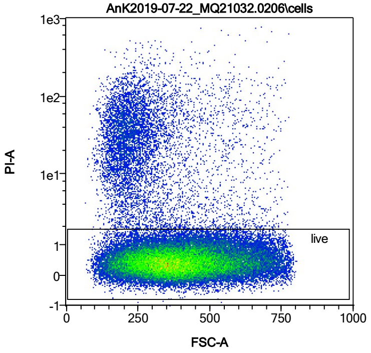


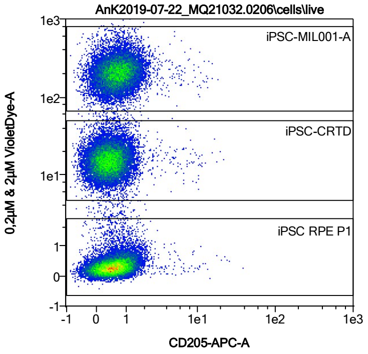


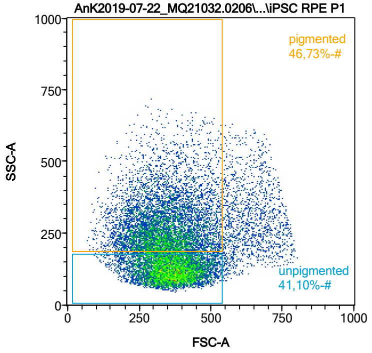


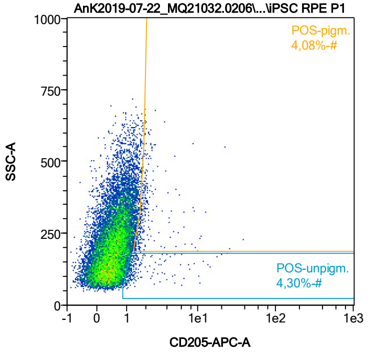


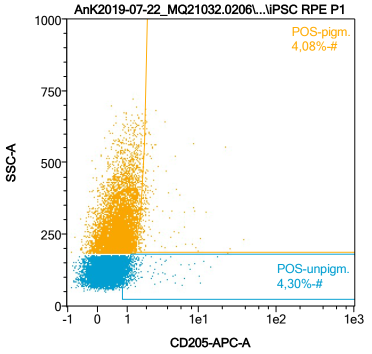


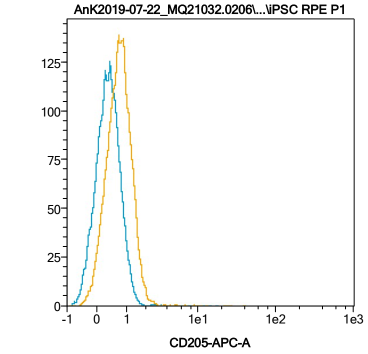


| Name | %-# | CD205-APC-A Mean | CD205-APC-A Median |
|---|---|---|---|
| POS-unpigm. | 4,30 | 1,73 | 1,19 |
| POS-pigm. | 4,08 | 2,59 | 1,82 |

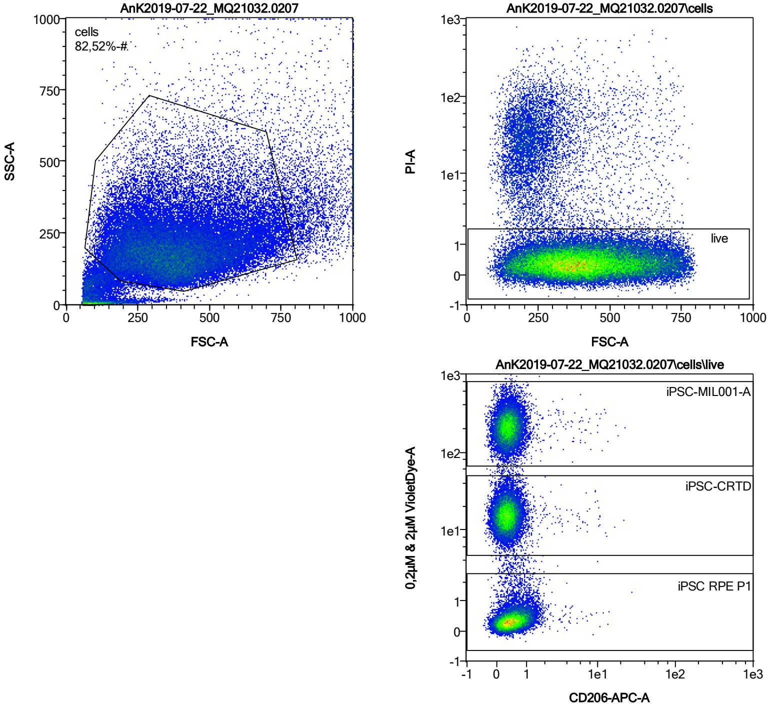


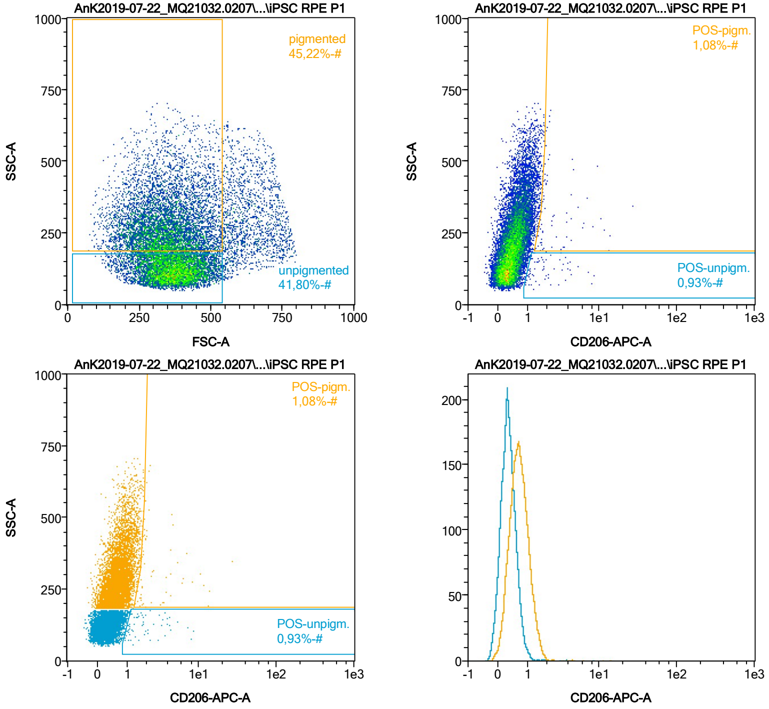


| Name | %-# | CD206-APC-A Mean | CD206-APC-A Median |
|---|---|---|---|
| POS-unpigm. | 0,93 | 2,28 | 1,71 |
| POS-pigm. | 1,08 | 3,40 | 1,91 |

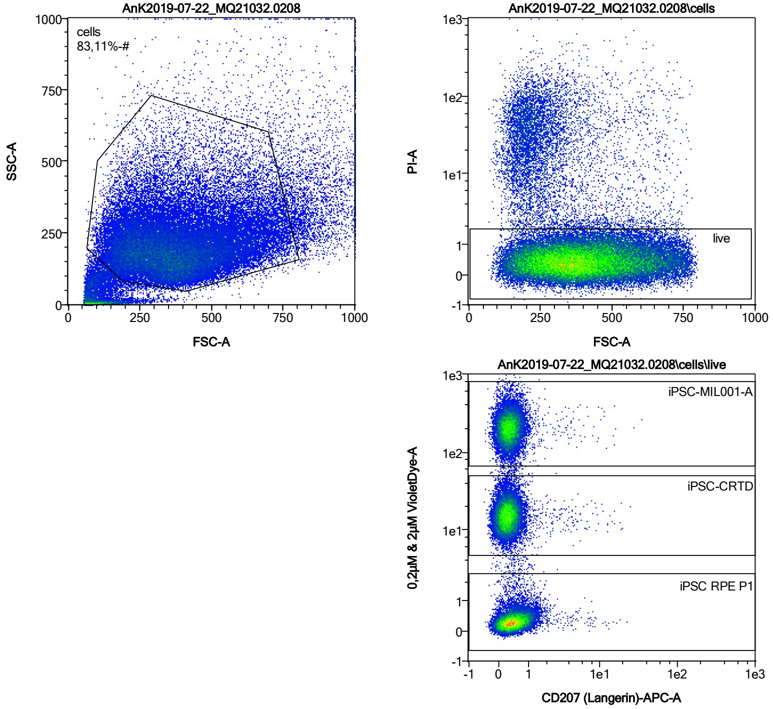


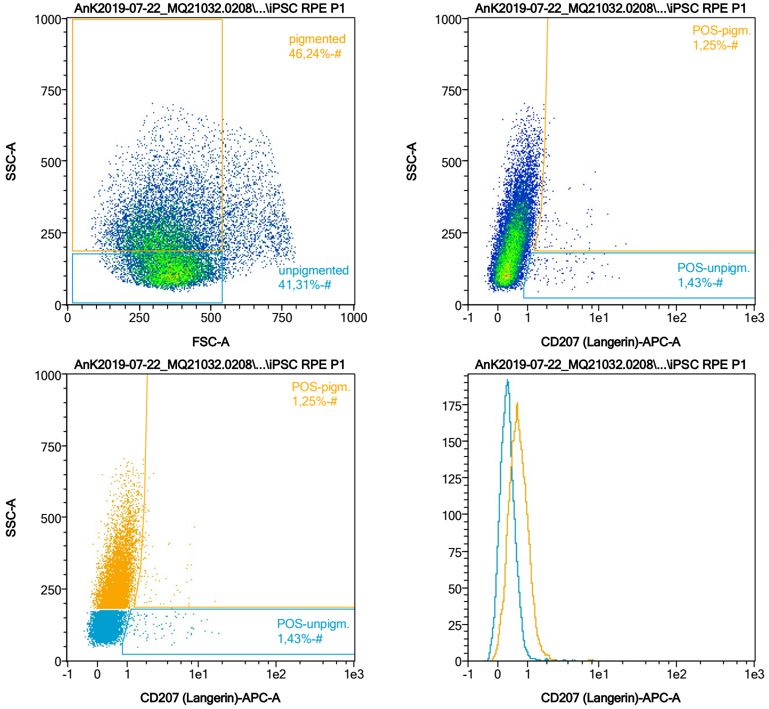


| Name | %-# | CD207 (Langerin)-APC-A Mean | CD207 (Langerin)-APC-A Median |
|---|---|---|---|
| POS-unpigm. | 1,43 | 3,62 | 2,07 |
| POS-pigm. | 1,25 | 3,16 | 2,10 |

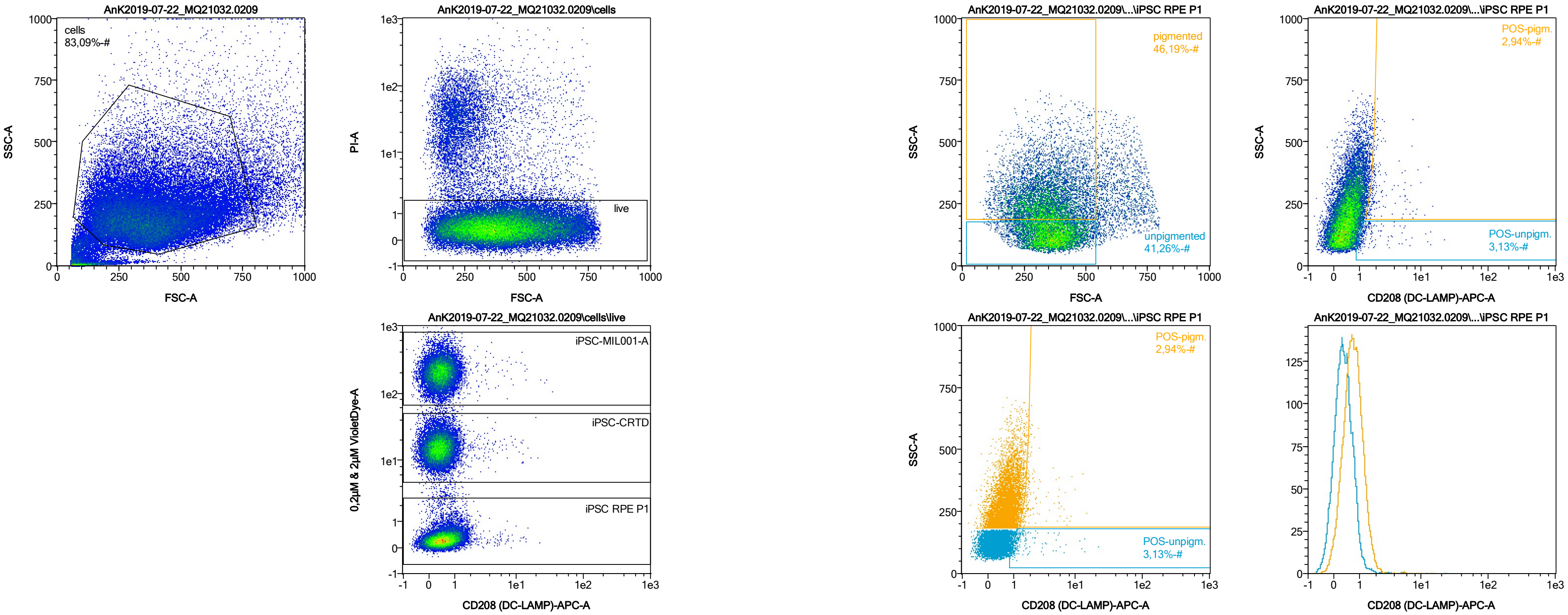


| Name | %-# | CD208 (DC-LAMP)-APC-A Mean | CD208 (DC-LAMP)-APC-A Median |
|---|---|---|---|
| POS-unpigm. | 3,13 | 1,93 | 1,22 |
| POS-pigm. | 2,94 | 2,50 | 1,84 |

Date:2024-06-14
Operator: Sebastian Knoebel
Version:2.13.0+54.g95abf0f4

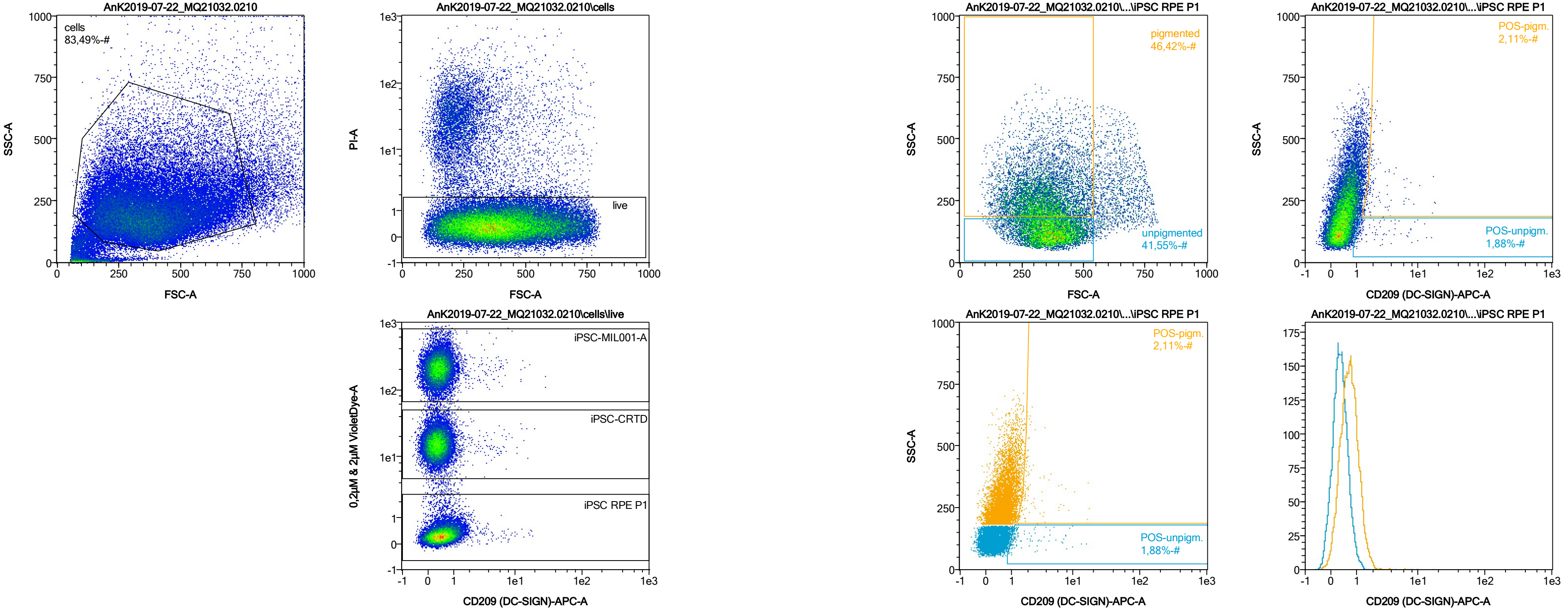


| Name | %-# | CD209 (DC-SIGN)-APC-A Mean | CD209 (DC-SIGN)-APC-A Median |
|---|---|---|---|
| POS-unpigm. | 1,88 | 2,38 | 1,25 |
| POS-pigm. | 2,11 | 2,67 | 1,90 |

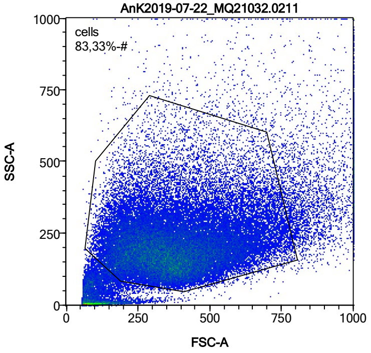


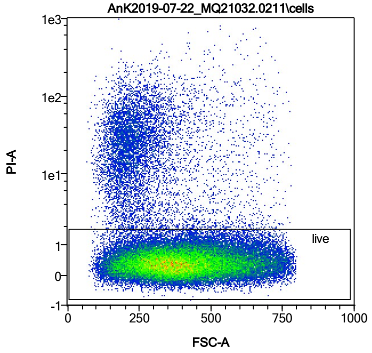


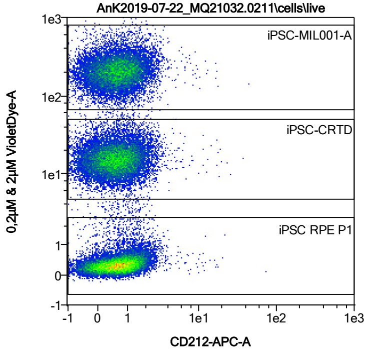


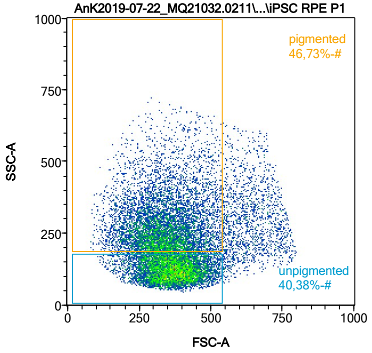


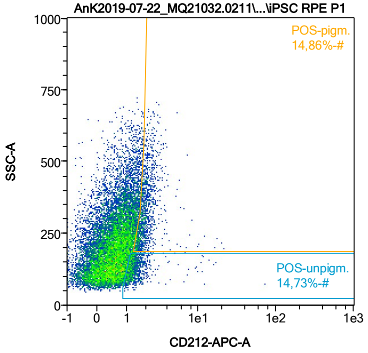


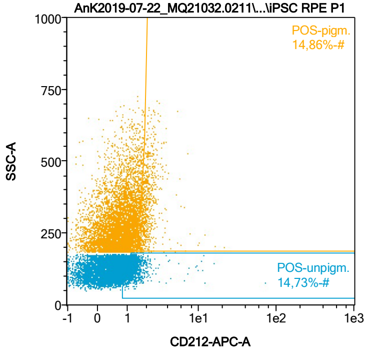


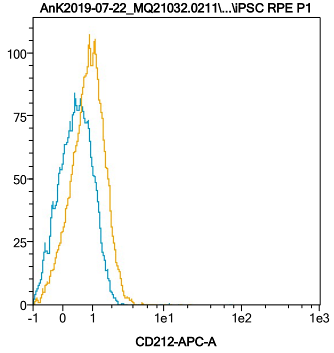


| Name | %-# | CD212-APC-A Mean | CD212-APC-A Median |
|---|---|---|---|
| POS-unpigm. | 14,73 | 1,56 | 1,27 |
| POS-pigm. | 14,86 | 2,09 | 1,89 |

Date:2024-06-14

Operator: Sebastian Knoebel

Version:2.13.0+54.g95abf0f4

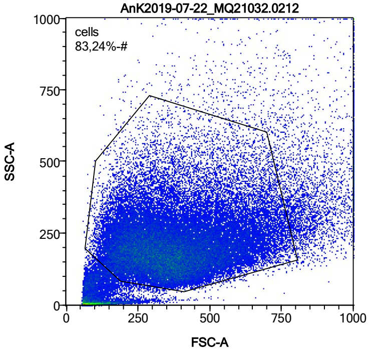


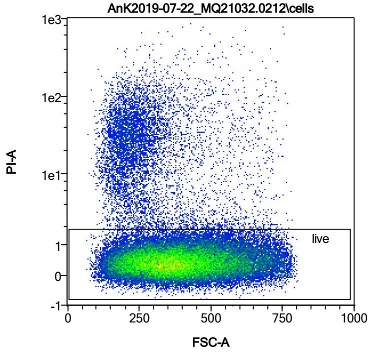


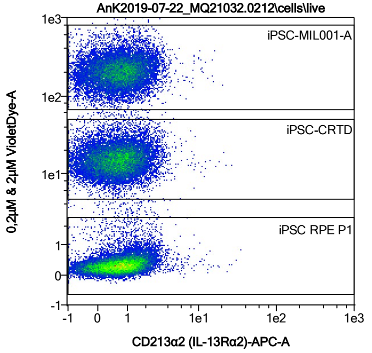


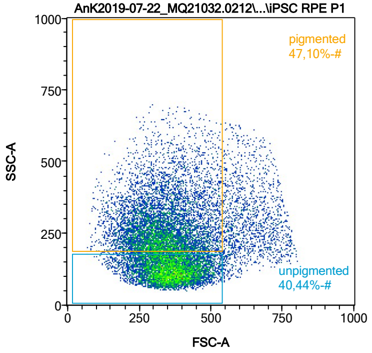


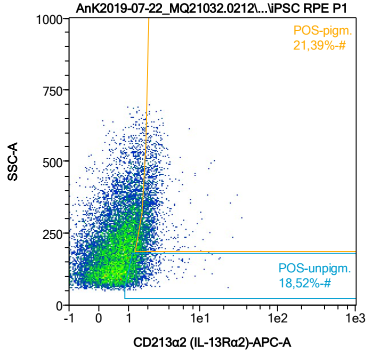


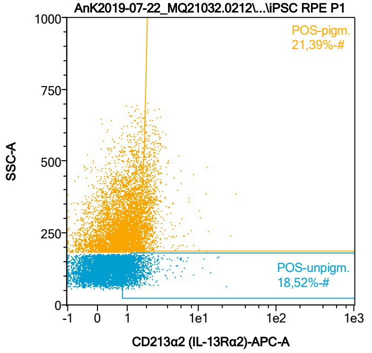


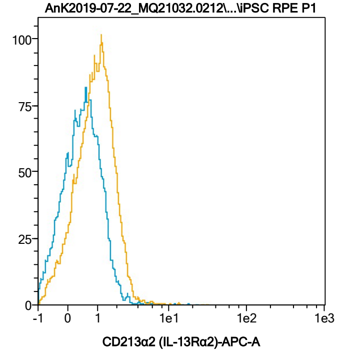


| Name | %-# | CD213α2 (IL-13Rα2)-APC-A Mean | CD213α2 (IL-13Rα2)-APC-A Median |
|---|---|---|---|
| POS-unpigm. | 18,52 | 1,60 | 1,33 |
| POS-pigm. | 21,39 | 2,21 | 1,96 |

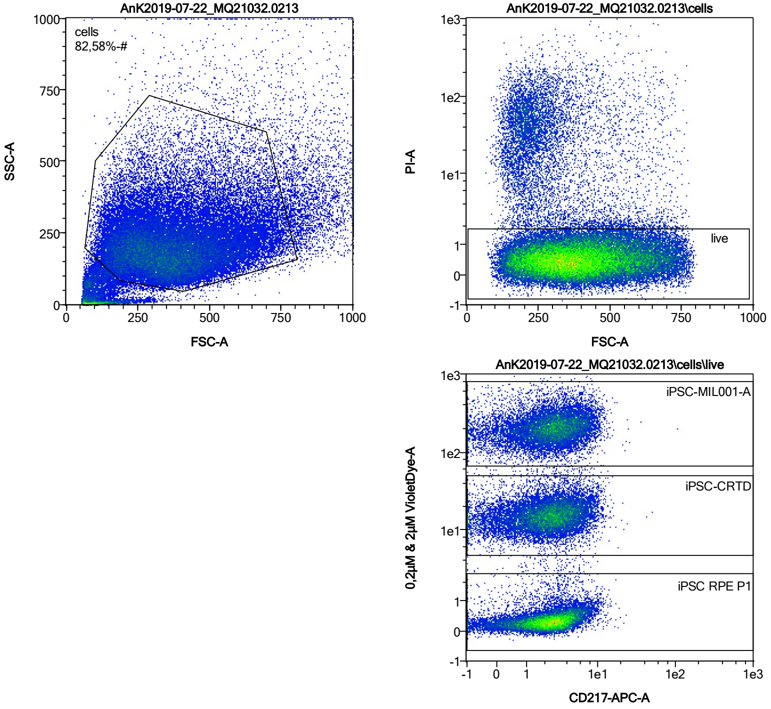


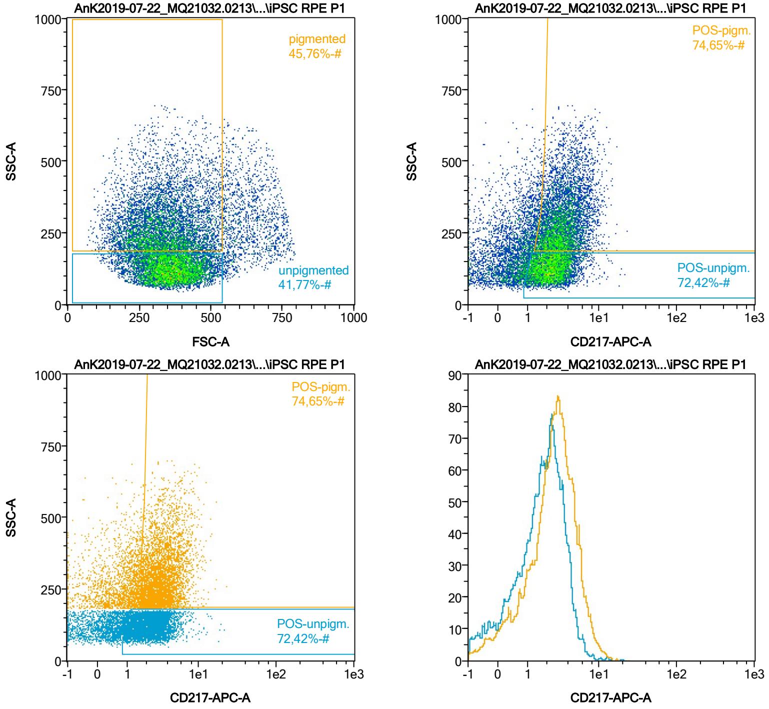


| Name | %-# | CD217-APC-A Mean | CD217-APC-A Median |
|---|---|---|---|
| POS-unpigm. | 72,42 | 2,54 | 2,28 |
| POS-pigm. | 74,65 | 3,54 | 3,10 |

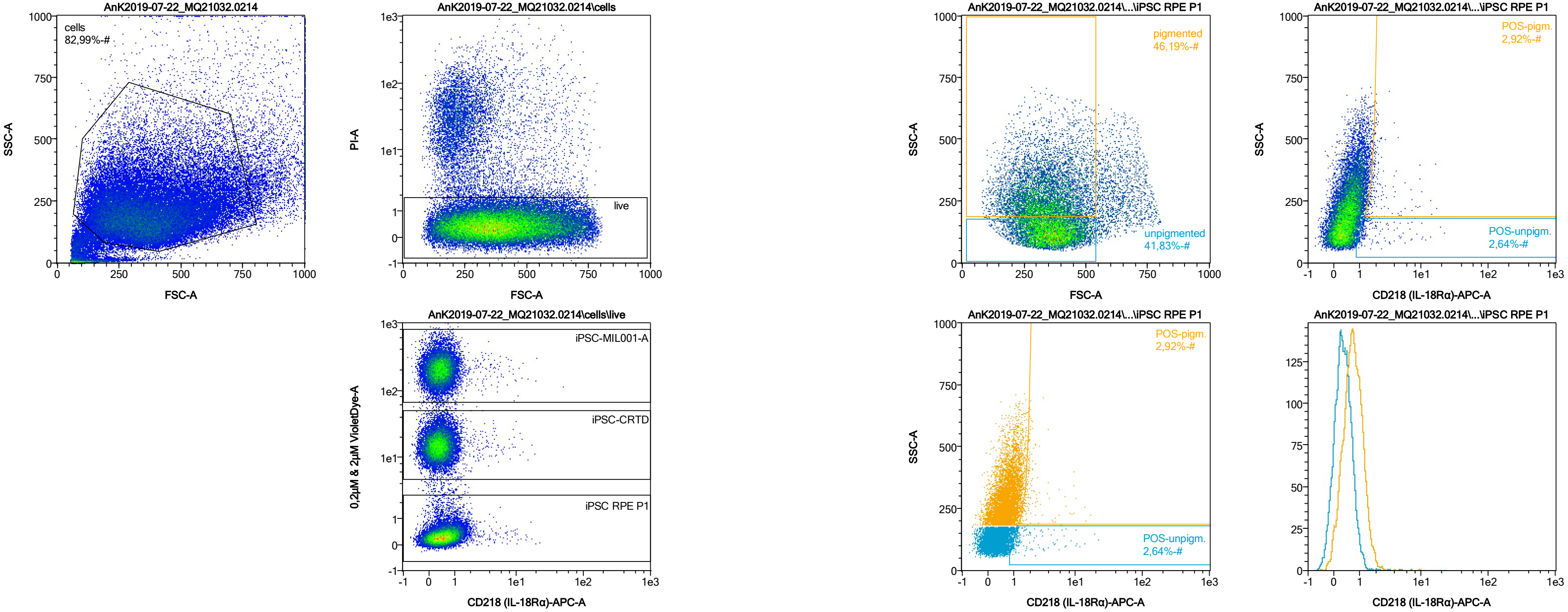


| Name | %-# | CD218 (IL-18Rα)-APC-A Mean | CD218 (IL-18Rα)-APC-A Median |
|---|---|---|---|
| POS-unpigm. | 2,64 | 2,38 | 1,24 |
| POS-pigm. | 2,92 | 2,51 | 1,87 |

Date:2024-06-14
Operator: Sebastian Knoebel
Version:2.13.0+54.g95abf0f4

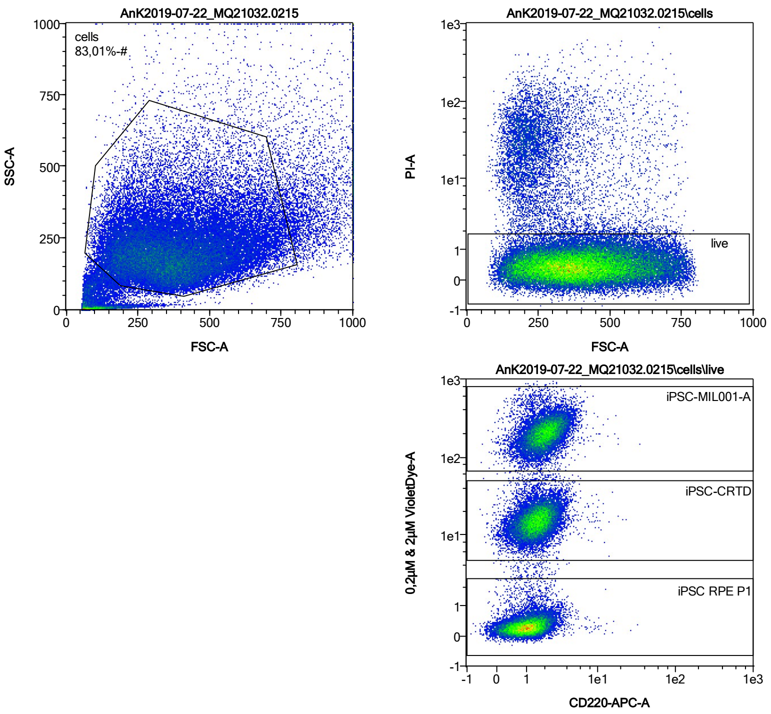


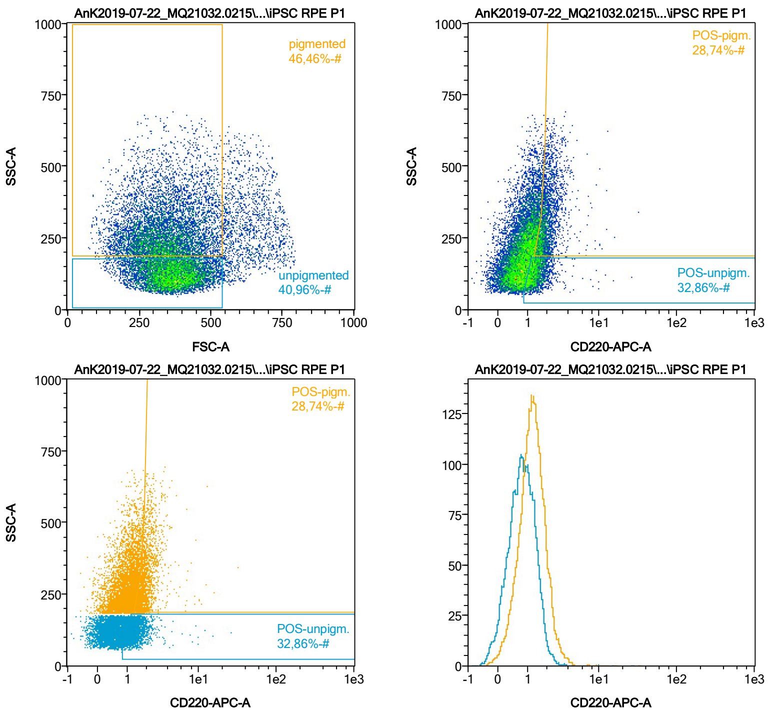


| Name | %-# | CD220-APC-A Mean | CD220-APC-A Median |
|---|---|---|---|
| POS-unpigm. | 32,86 | 1,45 | 1,31 |
| POS-pigm. | 28,74 | 2,08 | 1,89 |

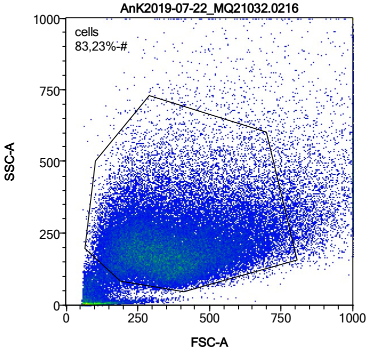


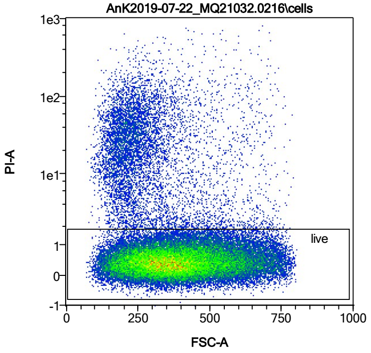


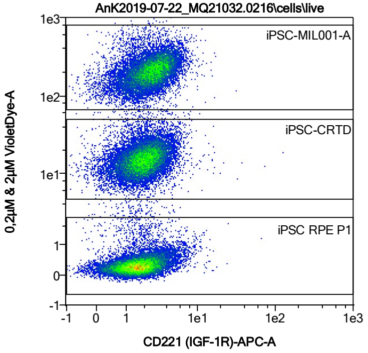


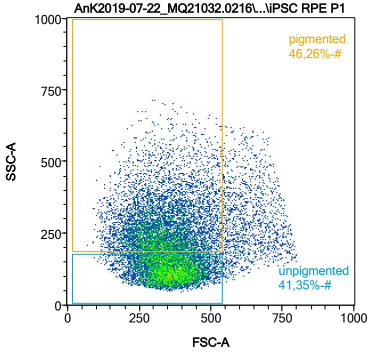


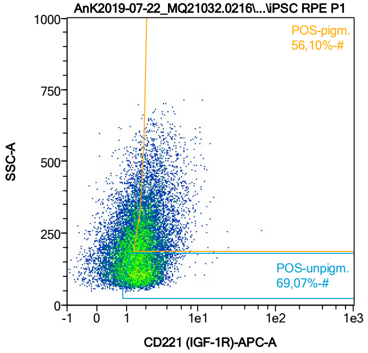


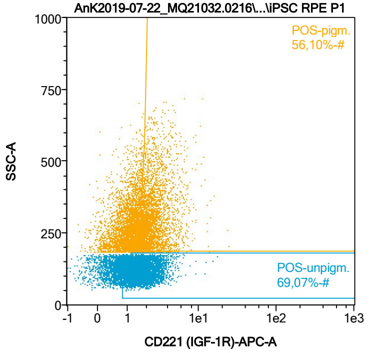


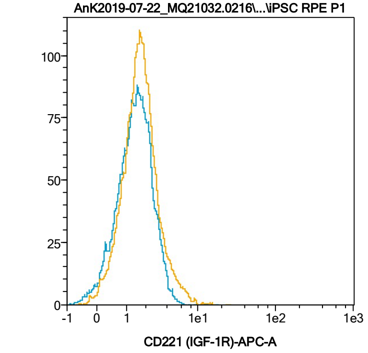


| Name | %-# | CD221 (IGF-1R)-APC-A Mean | CD221 (IGF-1R)-APC-A Median |
|---|---|---|---|
| POS-unpigm. | 69,07 | 1,96 | 1,76 |
| POS-pigm. | 56,10 | 2,59 | 2,19 |

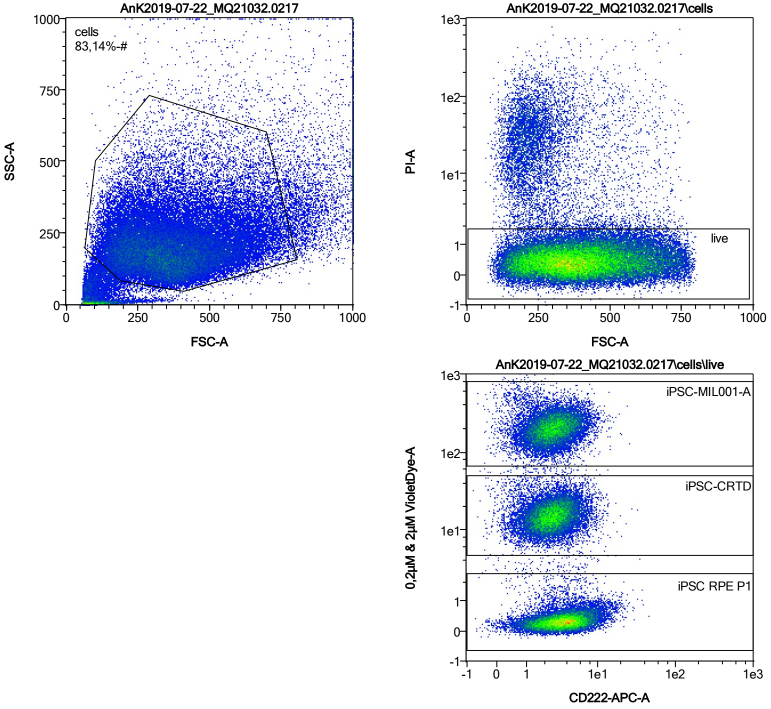


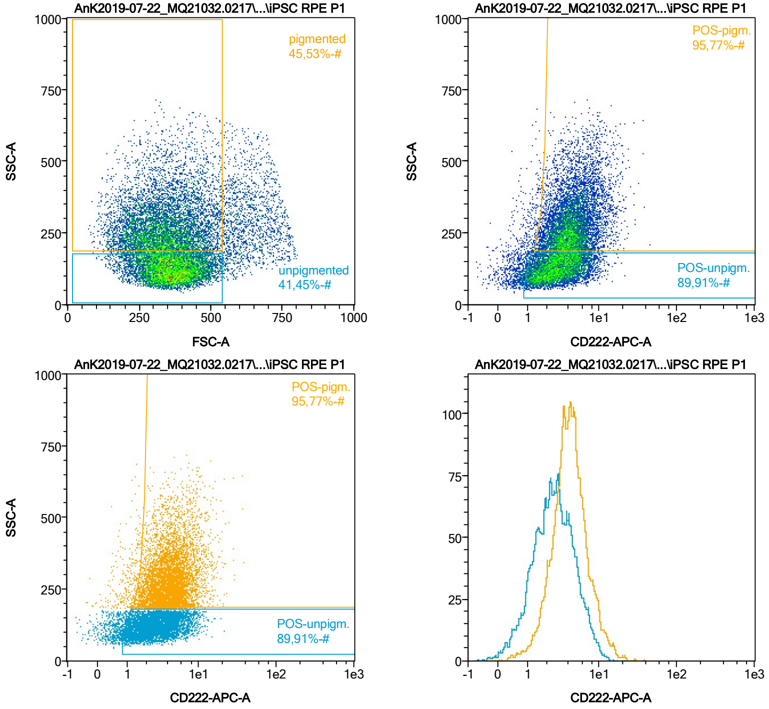


| Name | %-# | CD222-APC-A Mean | CD222-APC-A Median |
|---|---|---|---|
| POS-unpigm. | 89,91 | 3,24 | 2,74 |
| POS-pigm. | 95,77 | 4,81 | 4,16 |

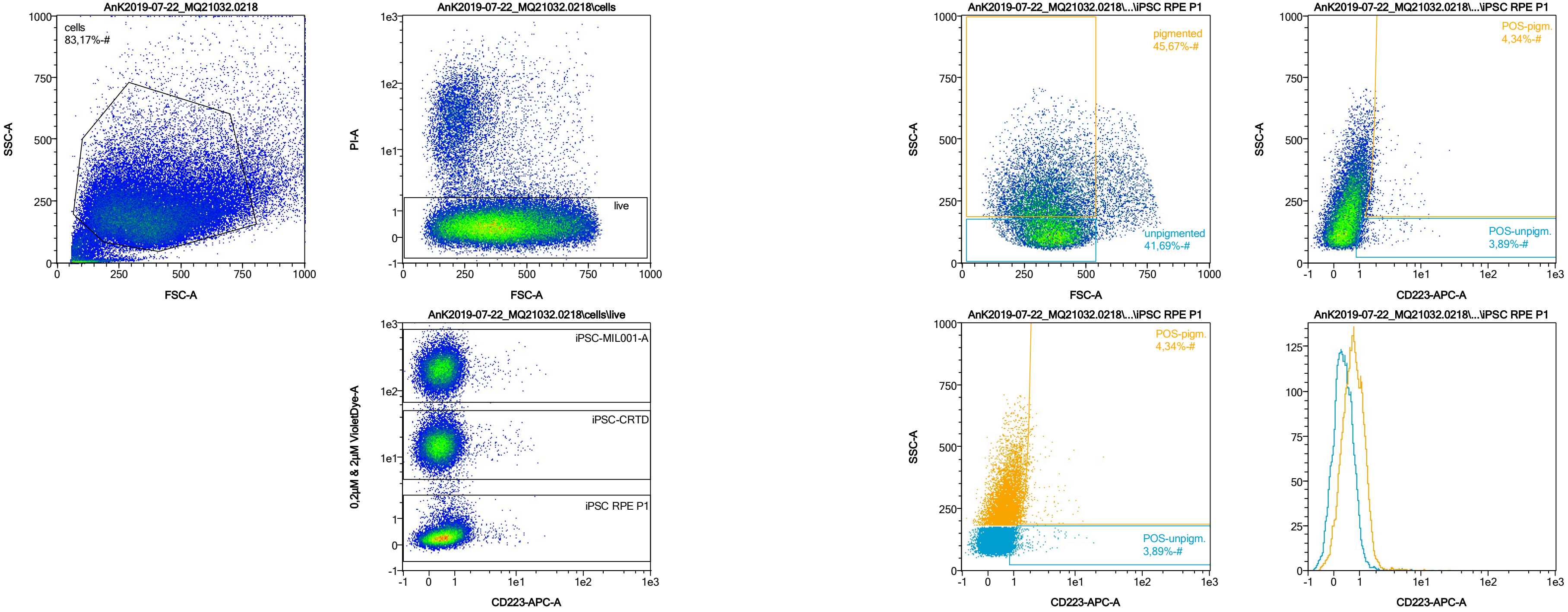


| Name | %-# | CD223-APC-A Mean | CD223-APC-A Median |
|---|---|---|---|
| POS-unpigm. | 3,89 | 1,86 | 1,23 |
| POS-pigm. | 4,34 | 2,49 | 1,82 |

Date:2024-06-14
Operator: Sebastian Knoebel
Version:2.13.0+54.g95abf0f4

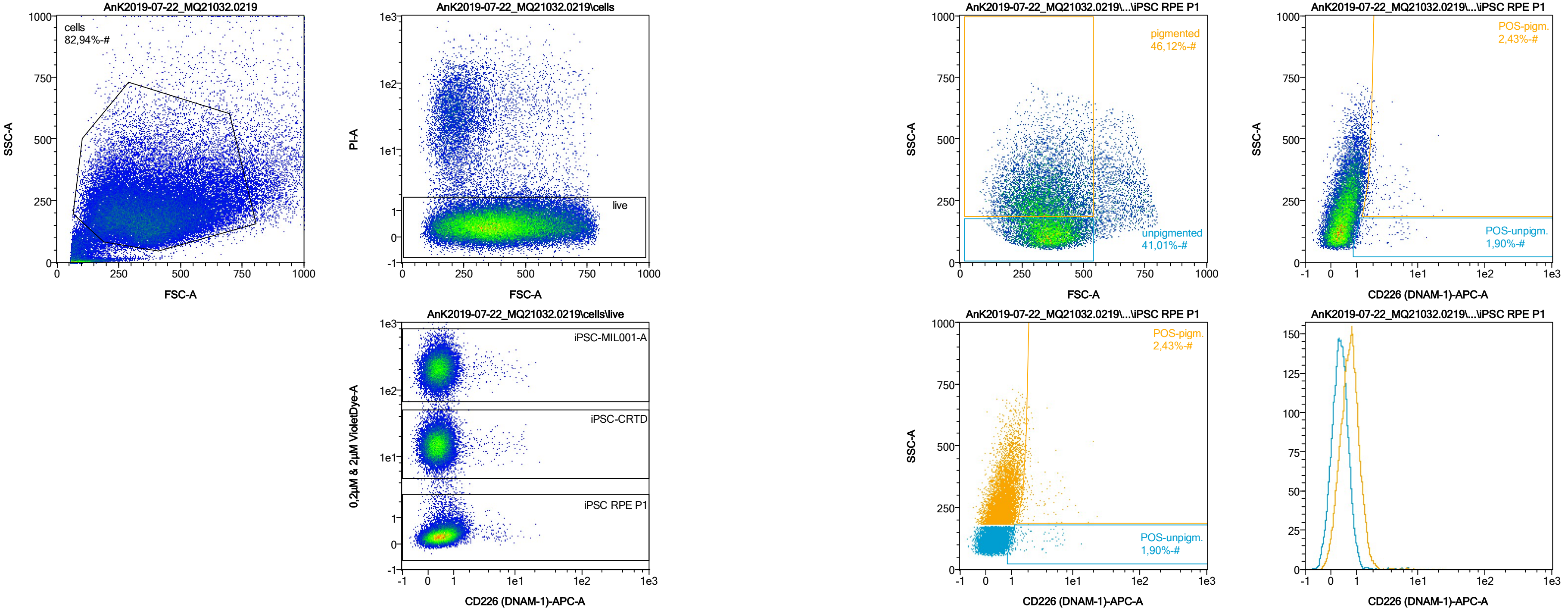


| Name | %-# | CD226 (DNAM-1)-APC-A Mean | CD226 (DNAM-1)-APC-A Median |
|---|---|---|---|
| POS-unpigm. | 1,90 | 2,34 | 1,24 |
| POS-pigm. | 2,43 | 3,04 | 1,97 |

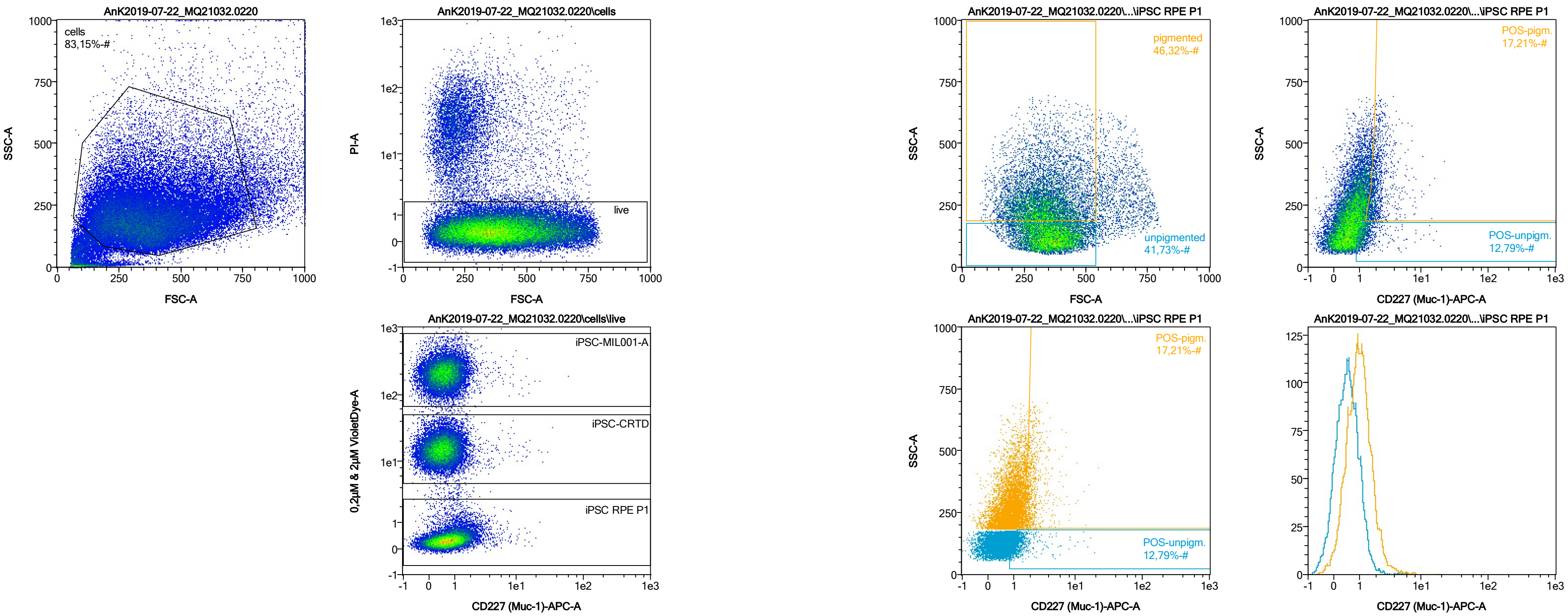


| Name | %-# | CD227 (Muc-1)-APC-A Mean | CD227 (Muc-1)-APC-A Median |
|---|---|---|---|
| POS-unpigm. | 12,79 | 1,55 | 1,24 |
| POS-pigm. | 17,21 | 2,23 | 1,92 |

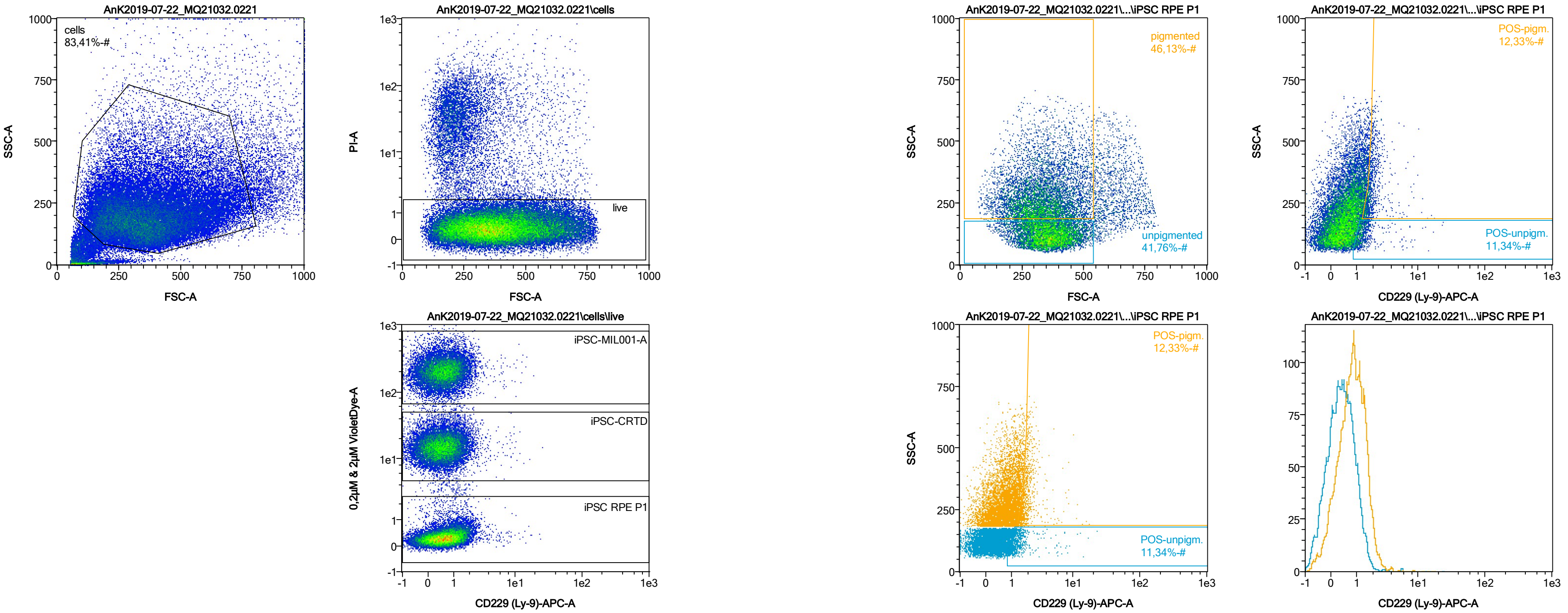


| Name | %-# | CD229 (Ly-9)-APC-A Mean | CD229 (Ly-9)-APC-A Median |
|---|---|---|---|
| POS-unpigm. | 11,34 | 1,57 | 1,24 |
| POS-pigm. | 12,33 | 2,08 | 1,81 |

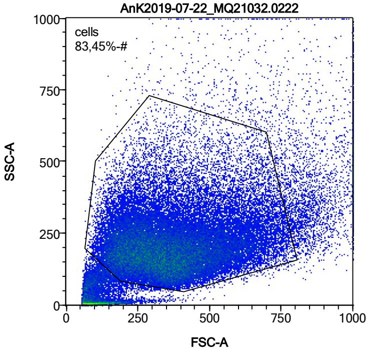


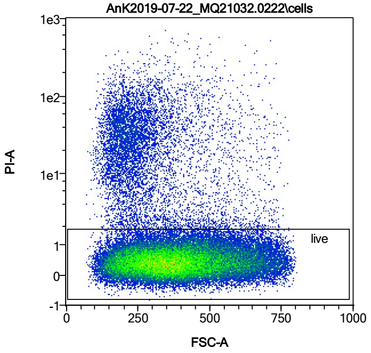


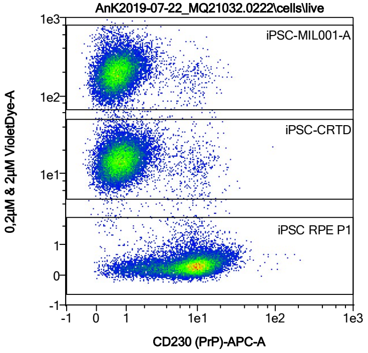


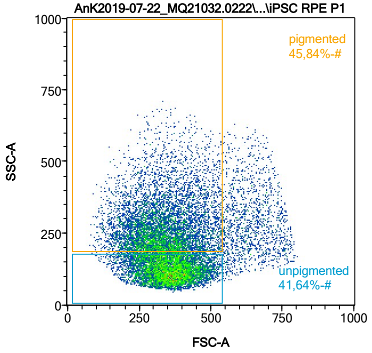


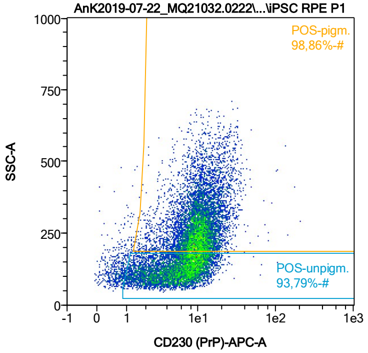


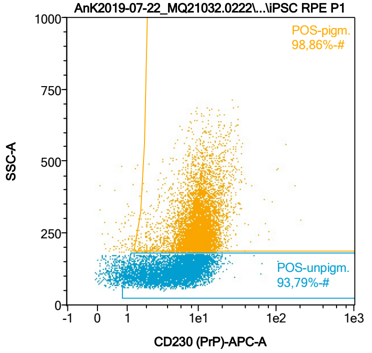


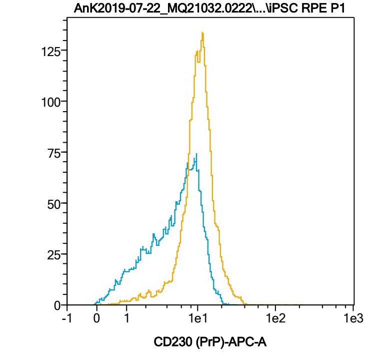


| Name | %-# | CD230 (PrP)-APC-A Mean | CD230 (PrP)-APC-A Median |
|---|---|---|---|
| POS-unpigm. | 93,79 | 6,43 | 5,95 |
| POS-pigm. | 98,86 | 11,10 | 10,24 |

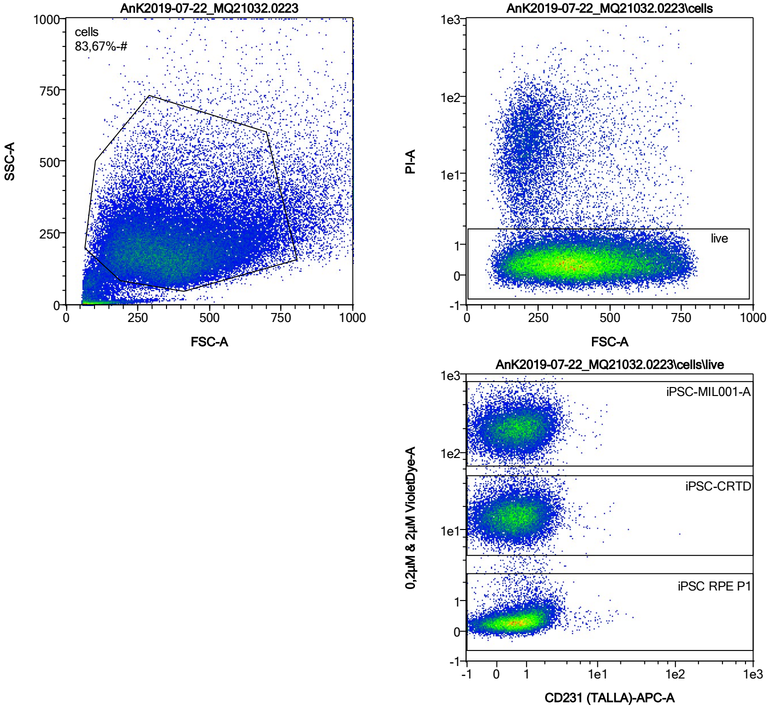


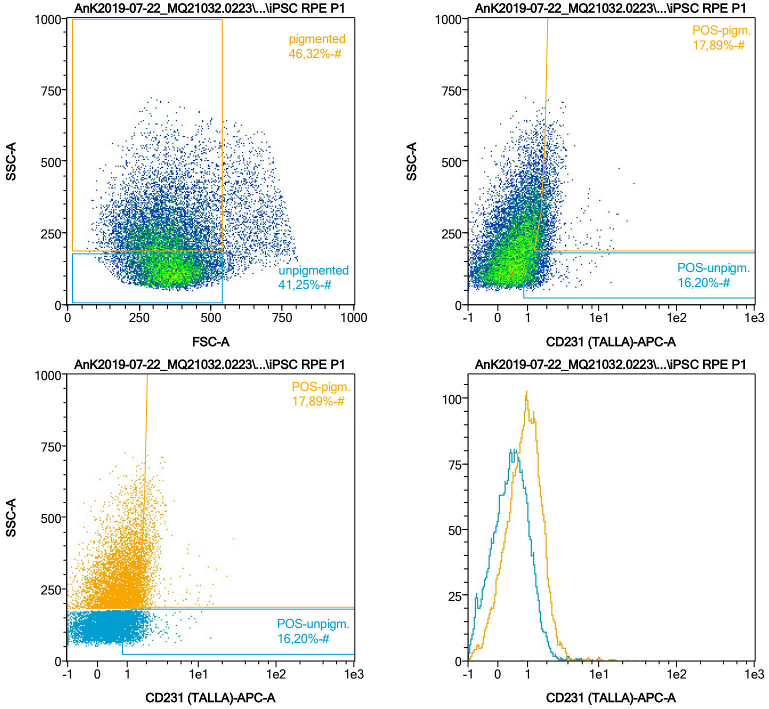


| Name | %-# | CD231 (TALLA)-APC-A Mean | CD231 (TALLA)-APC-A Median |
|---|---|---|---|
| POS-unpigm. | 16,20 | 1,53 | 1,30 |
| POS-pigm. | 17,89 | 2,22 | 1,90 |

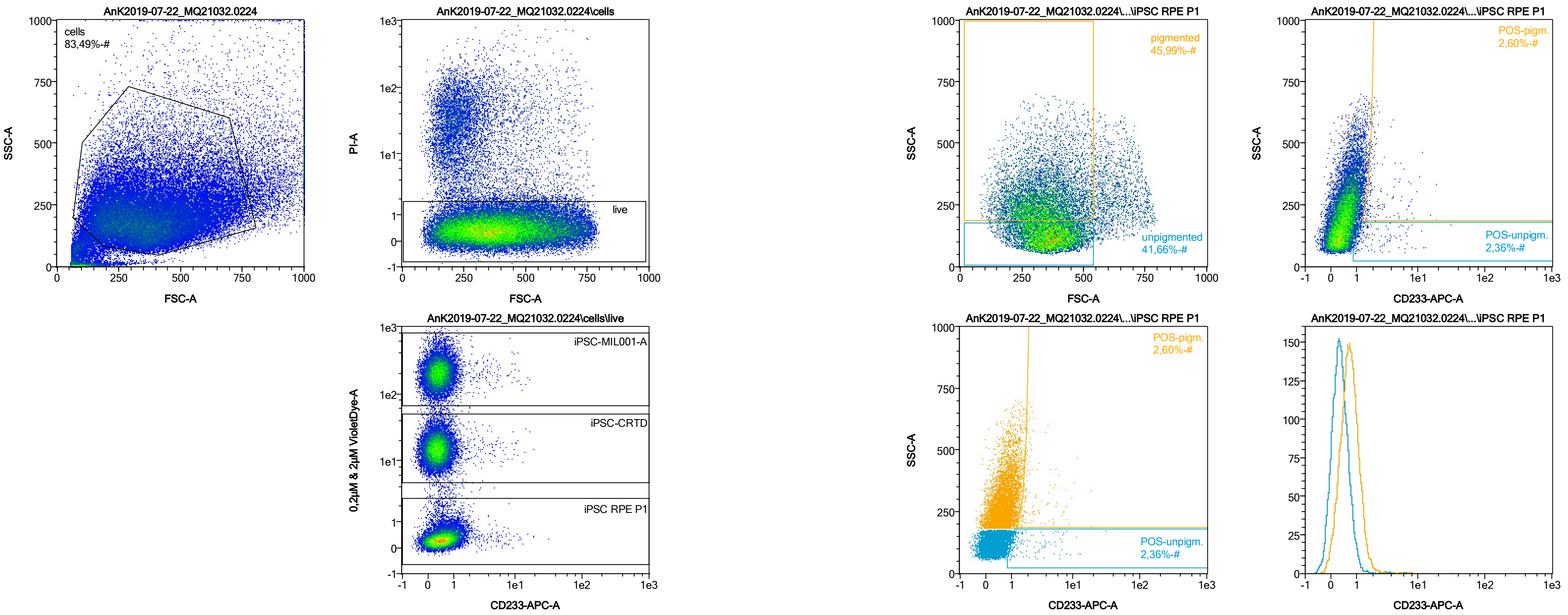


| Name | %-# | CD233-APC-A Mean | CD233-APC-A Median |
|---|---|---|---|
| POS-unpigm. | 2,36 | 2,69 | 1,38 |
| POS-pigm. | 2,60 | 2,90 | 1,94 |

Date:2024-06-14

Operator: Sebastian Knoebel

Version:2.13.0+54.g95abf0f4

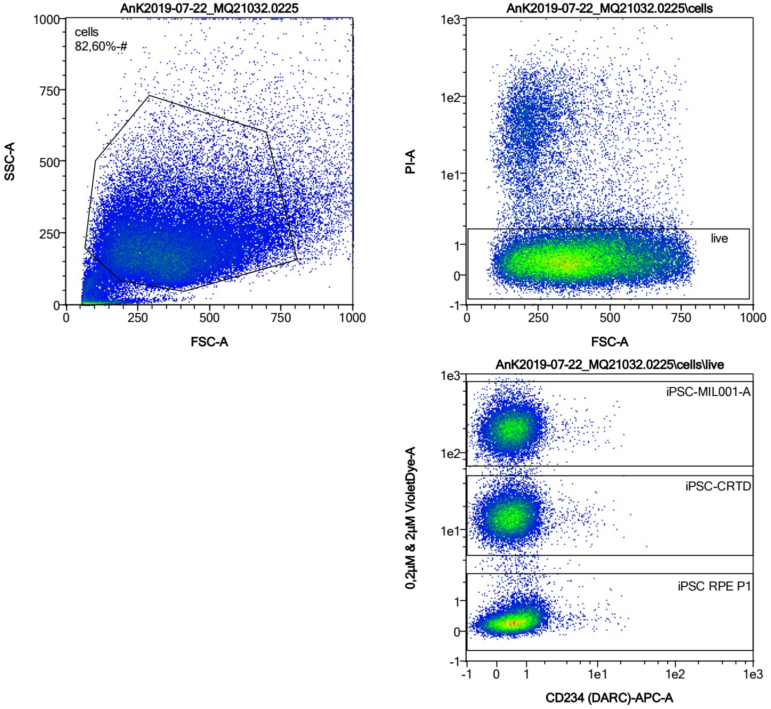


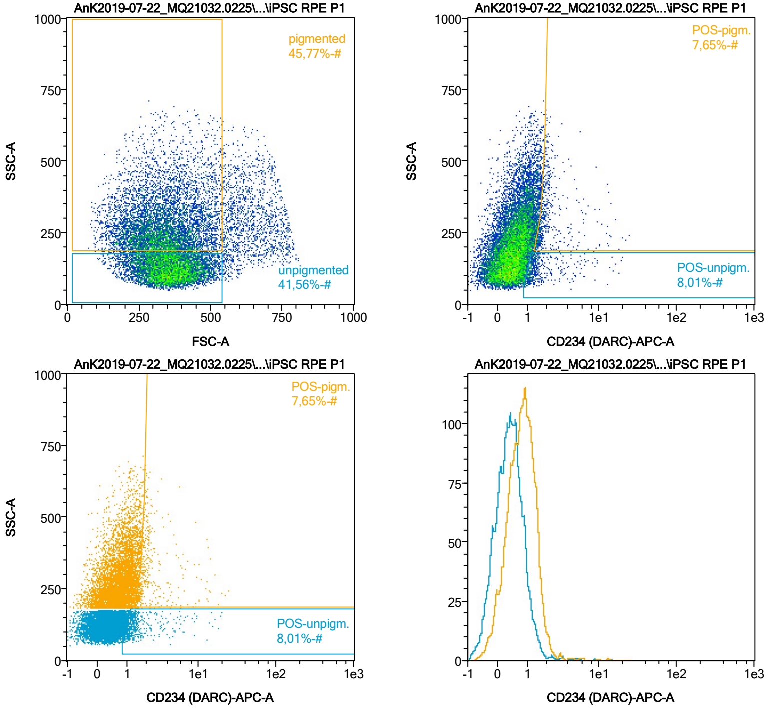


| Name | %-# | CD234 (DARC)-APC-A Mean | CD234 (DARC)-APC-A Median |
|---|---|---|---|
| POS-unpigm. | 8,01 | 1,95 | 1,28 |
| POS-pigm. | 7,65 | 2,64 | 1,85 |

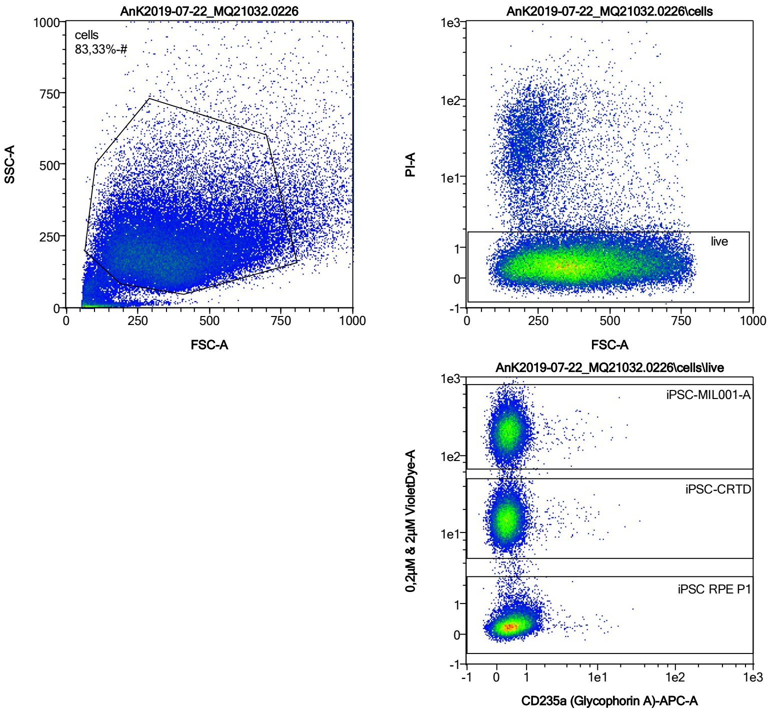


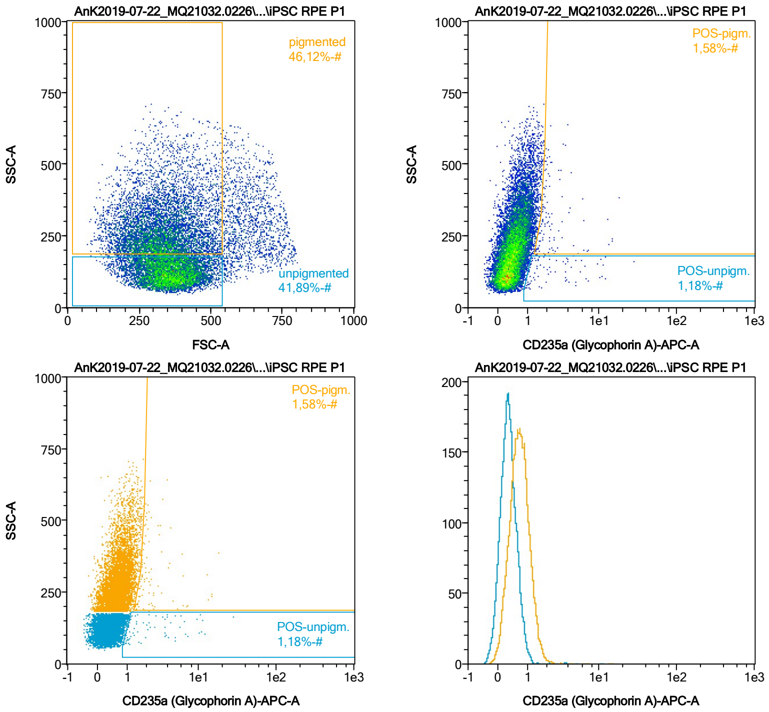


| Name | %-# | CD235a (Glycophorin A)-APC-A Mean | CD235a (Glycophorin A)-APC-A Median |
|---|---|---|---|
| POS-unpigm. | 1,18 | 3,32 | 1,49 |
| POS-pigm. | 1,58 | 2,89 | 1,94 |

Date:2024-06-14

Operator: Sebastian Knoebel

Version:2.13.0+54.g95abf0f4

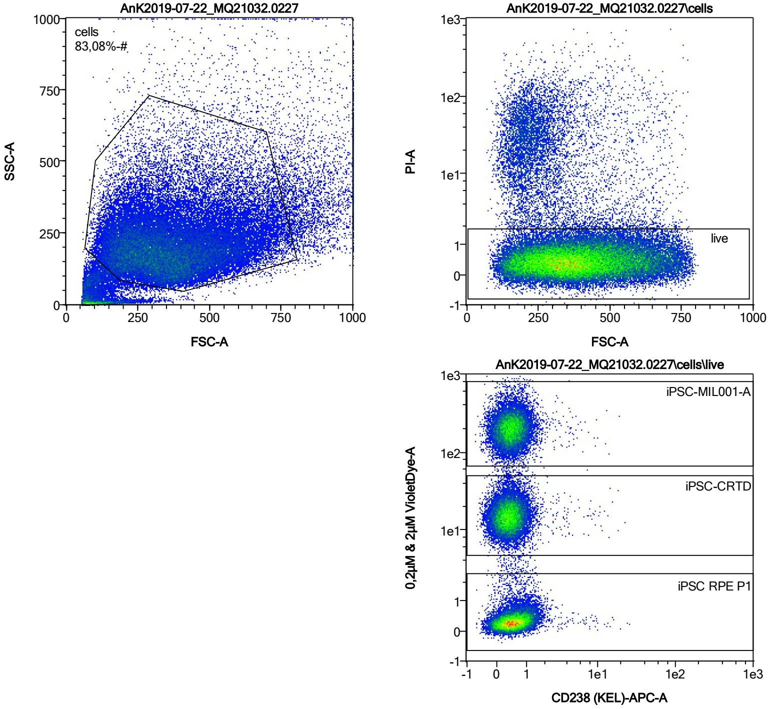


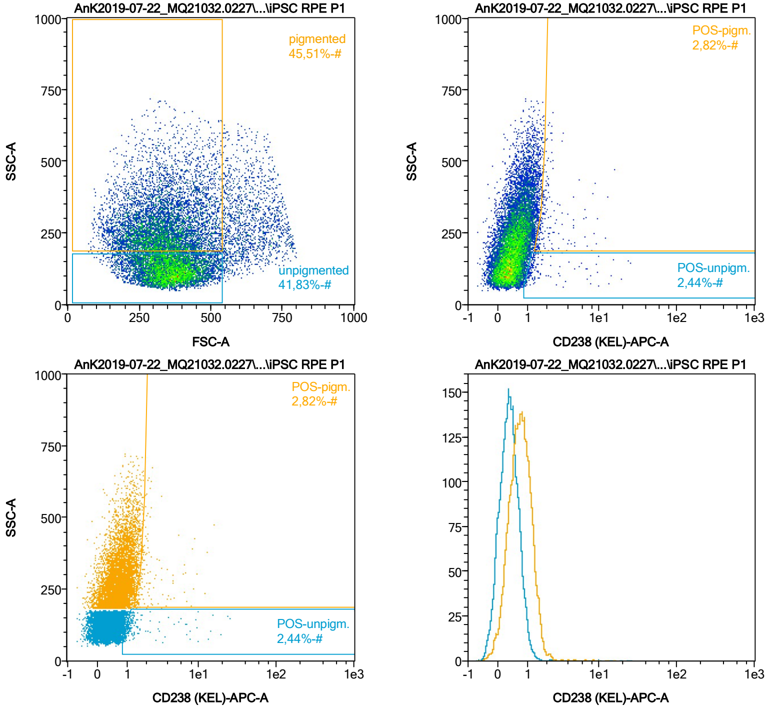


| Name | %-# | CD238 (KEL)-APC-A Mean | CD238 (KEL)-APC-A Median |
|---|---|---|---|
| POS-unpigm. | 2,44 | 2,53 | 1,21 |
| POS-pigm. | 2,82 | 2,49 | 1,81 |

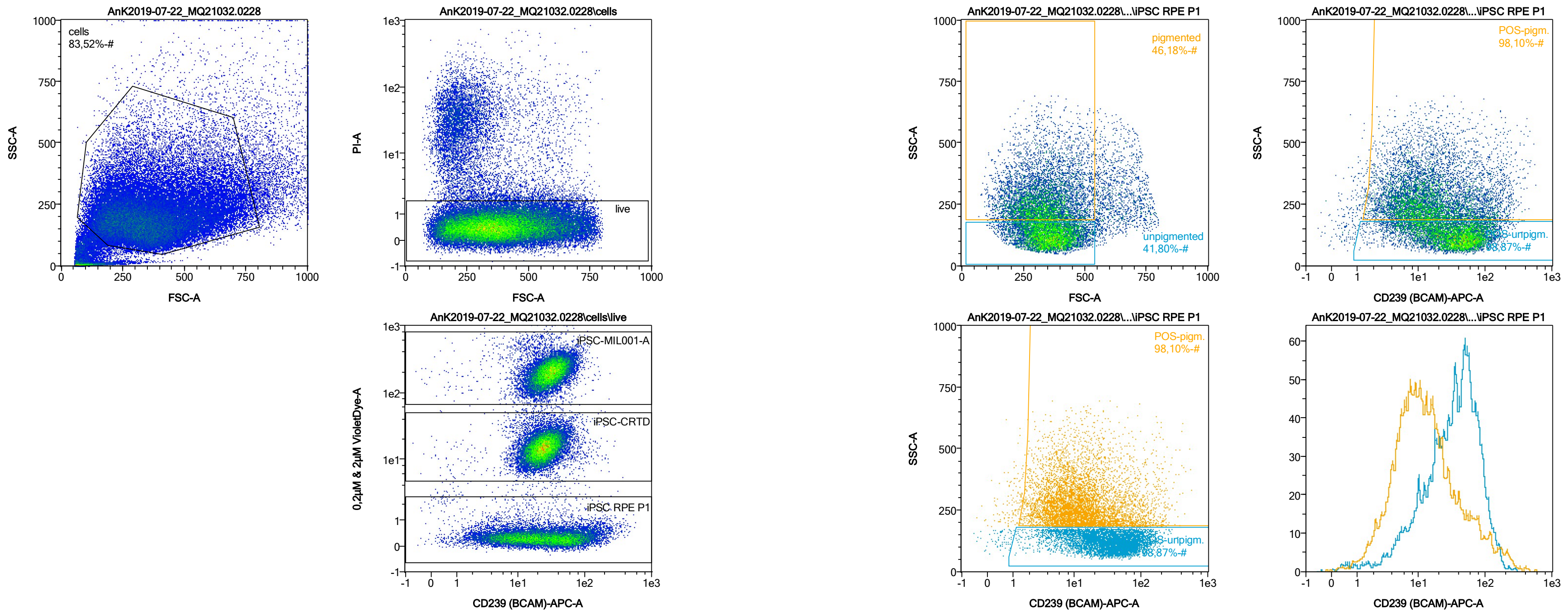


| Name | %-# | CD239 (BCAM)-APC-A Mean | CD239 (BCAM)-APC-A Median |
|---|---|---|---|
| POS-unpigm. | 98,87 | 43,32 | 35,65 |
| POS-pigm. | 98,10 | 26,11 | 11,83 |

Date:2024-06-14

Operator: Sebastian Knoebel

Version:2.13.0+54.g95abf0f4

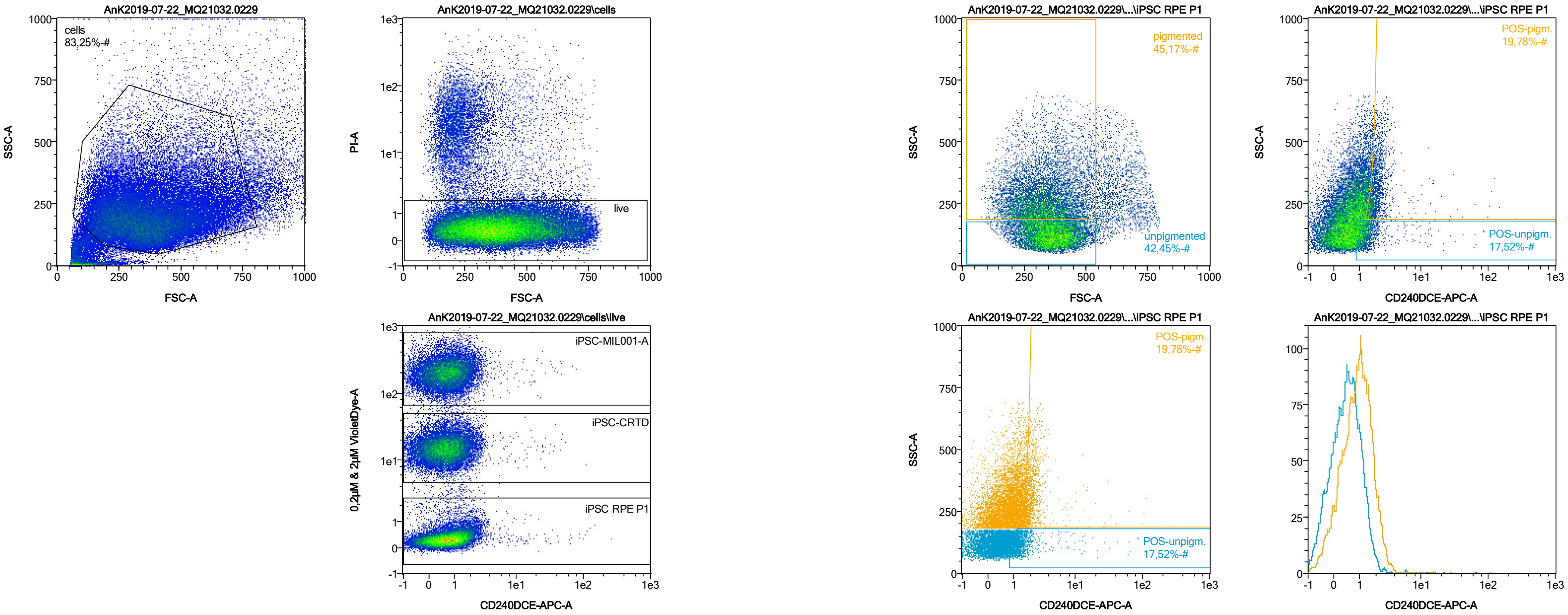


| Name | %-# | CD240DCE-APC-A Mean | CD240DCE-APC-A Median |
|---|---|---|---|
| POS-unpigm. | 17,52 | 2,04 | 1,30 |
| POS-pigm. | 19,78 | 2,78 | 1,90 |

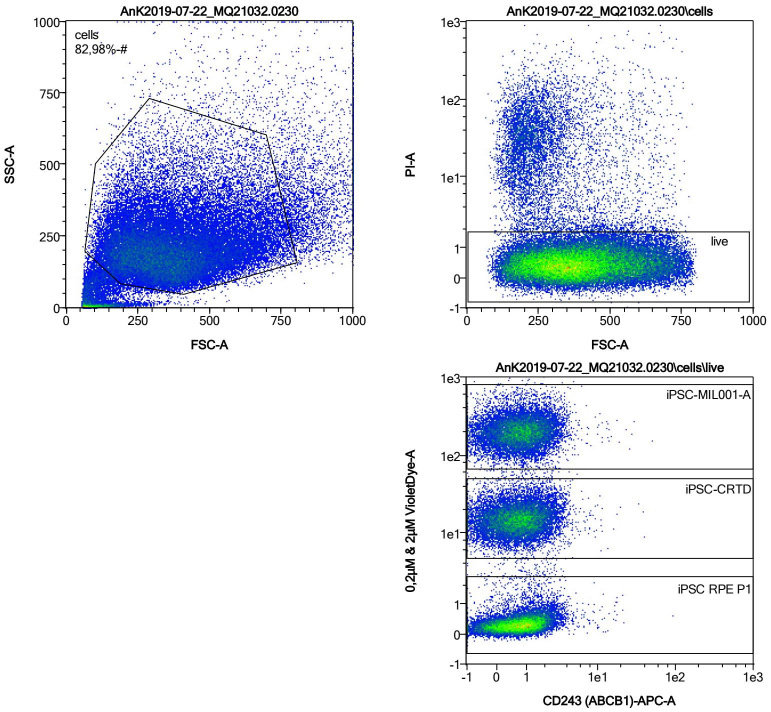


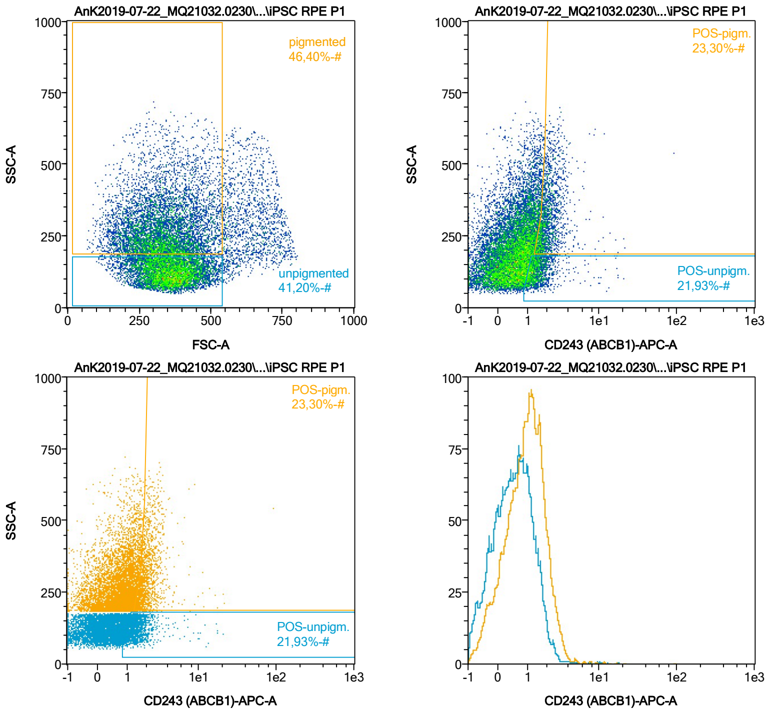


| Name | %-# | CD243 (ABCB1)-APC-A Mean | CD243 (ABCB1)-APC-A Median |
|---|---|---|---|
| POS-unpigm. | 21,93 | 1,63 | 1,37 |
| POS-pigm. | 23,30 | 2,26 | 1,95 |

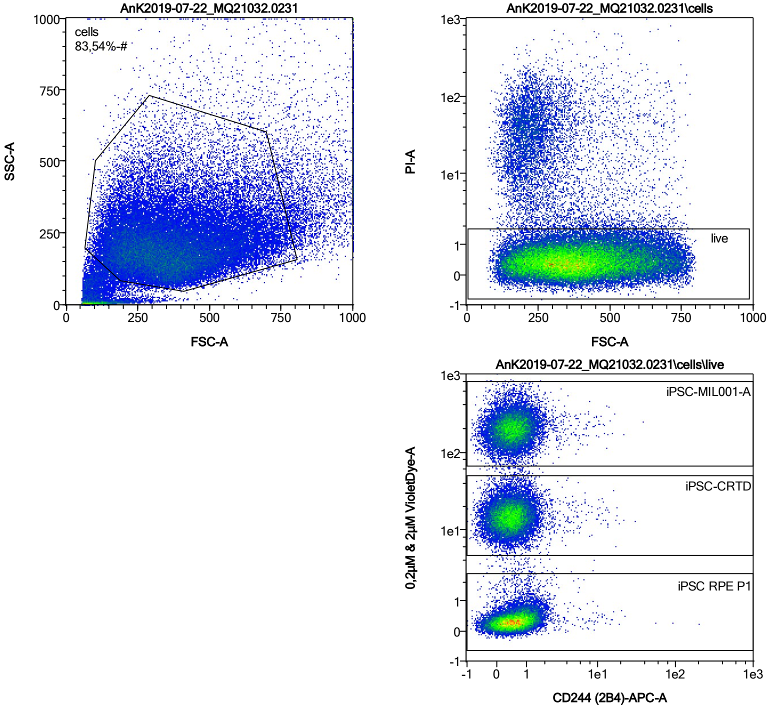


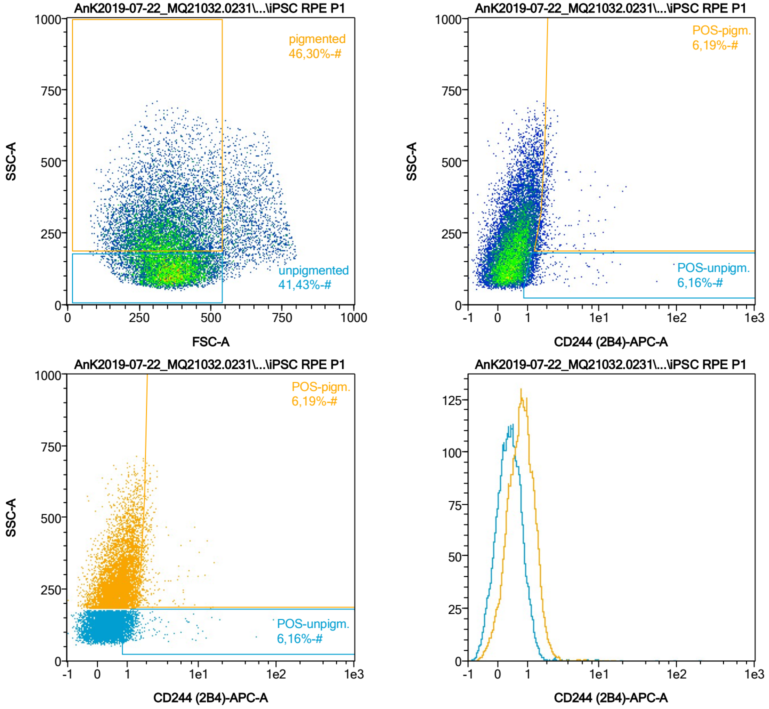


| Name | %-# | CD244 (2B4)-APC-A Mean | CD244 (2B4)-APC-A Median |
|---|---|---|---|
| POS-unpigm. | 6,16 | 2,59 | 1,20 |
| POS-pigm. | 6,19 | 2,26 | 1,85 |

Date:2024-06-14

Operator: Sebastian Knoebel

Version:2.13.0+54.g95abf0f4

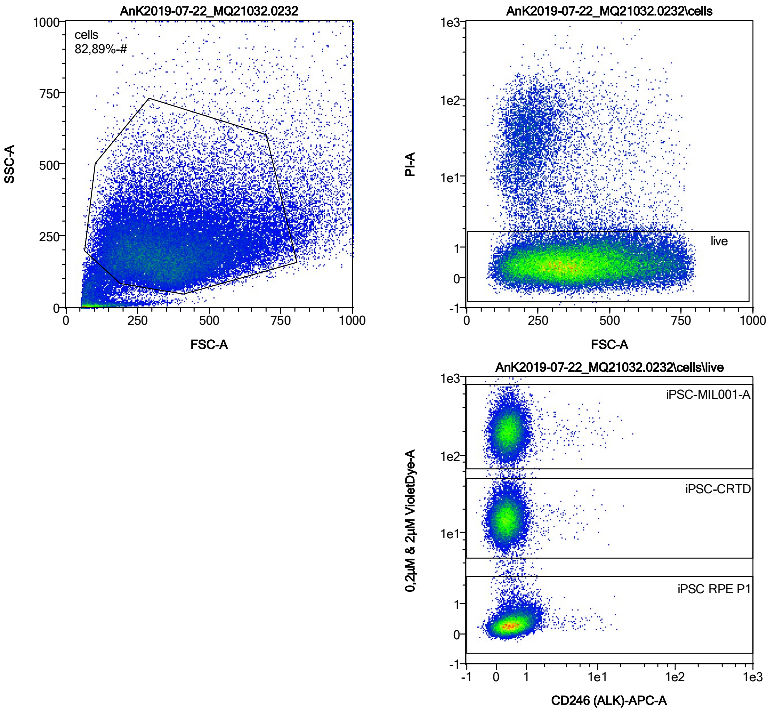


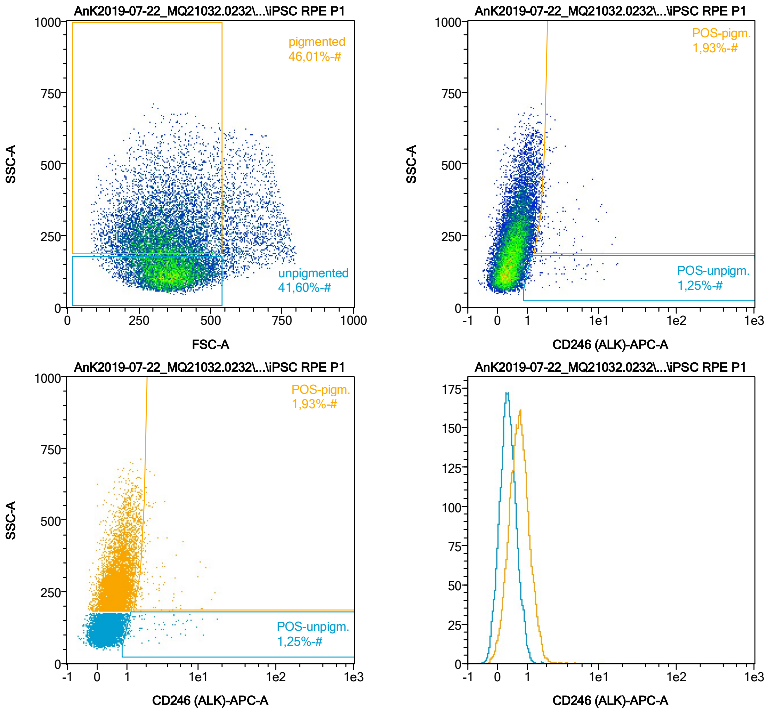


| Name | %-# | CD246 (ALK)-APC-A Mean | CD246 (ALK)-APC-A Median |
|---|---|---|---|
| POS-unpigm. | 1,25 | 3,13 | 1,73 |
| POS-pigm. | 1,93 | 2,93 | 1,98 |

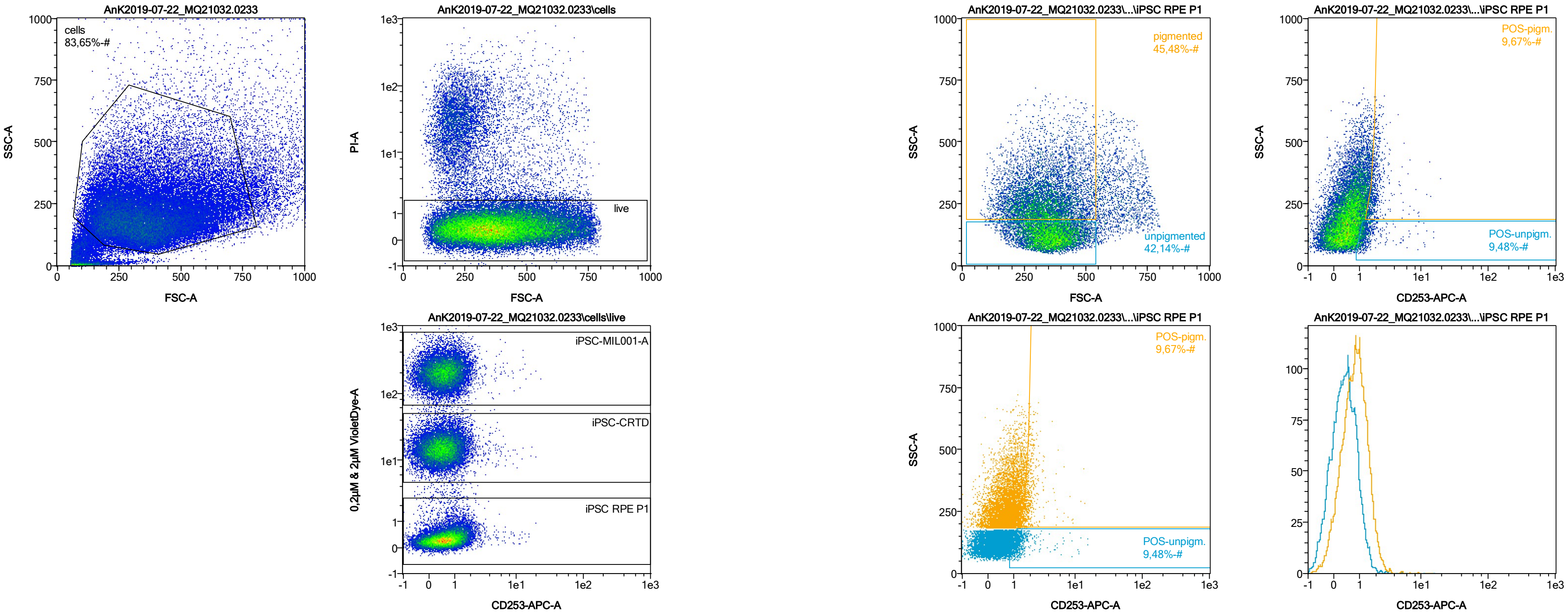


| Name | %-# | CD253-APC-A Mean | CD253-APC-A Median |
|---|---|---|---|
| POS-unpigm. | 9,48 | 1,55 | 1,23 |
| POS-pigm. | 9,67 | 2,06 | 1,83 |

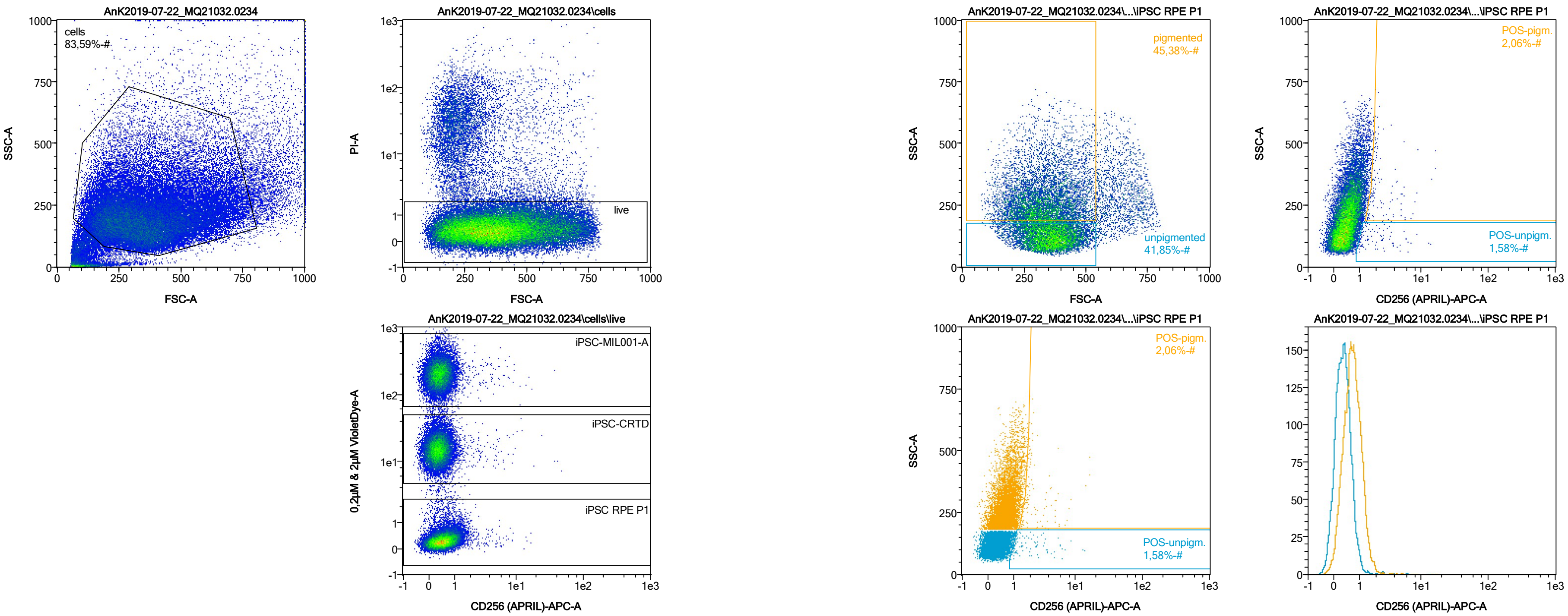


| Name | %-# | CD256 (APRIL)-APC-A Mean | CD256 (APRIL)-APC-A Median |
|---|---|---|---|
| POS-unpigm. | 1,58 | 2,40 | 1,48 |
| POS-pigm. | 2,06 | 2,74 | 1,91 |

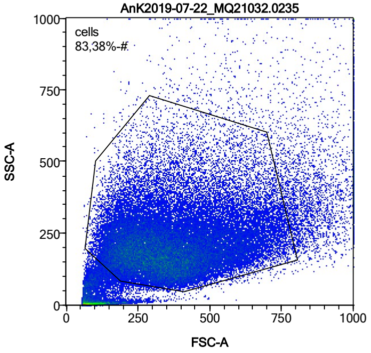


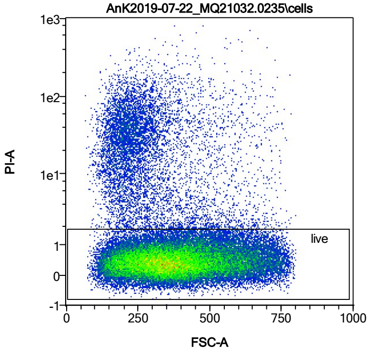


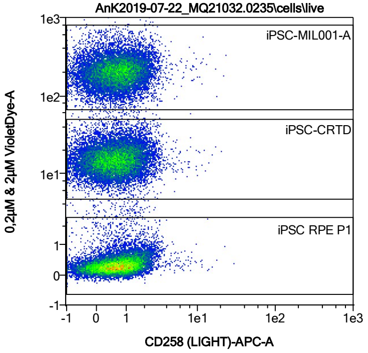


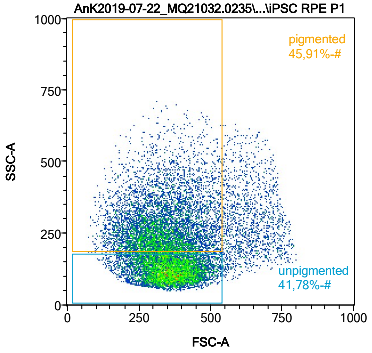


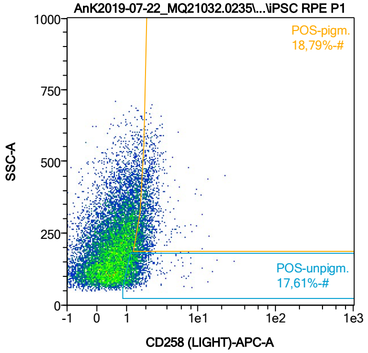


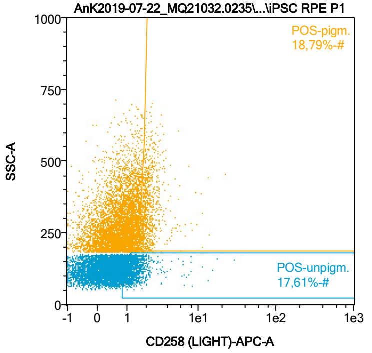


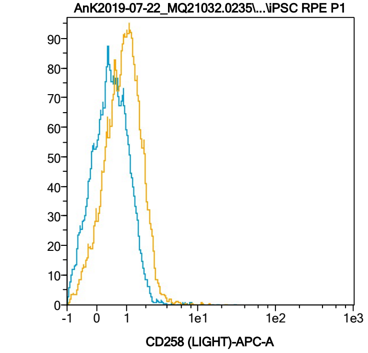


| Name | %-# | CD258 (LIGHT)-APC-A Mean | CD258 (LIGHT)-APC-A Median |
|---|---|---|---|
| POS-unpigm. | 17,61 | 1,55 | 1,31 |
| POS-pigm. | 18,79 | 2,17 | 1,94 |

Date:2024-06-14

Operator: Sebastian Knoebel

Version:2.13.0+54.g95abf0f4

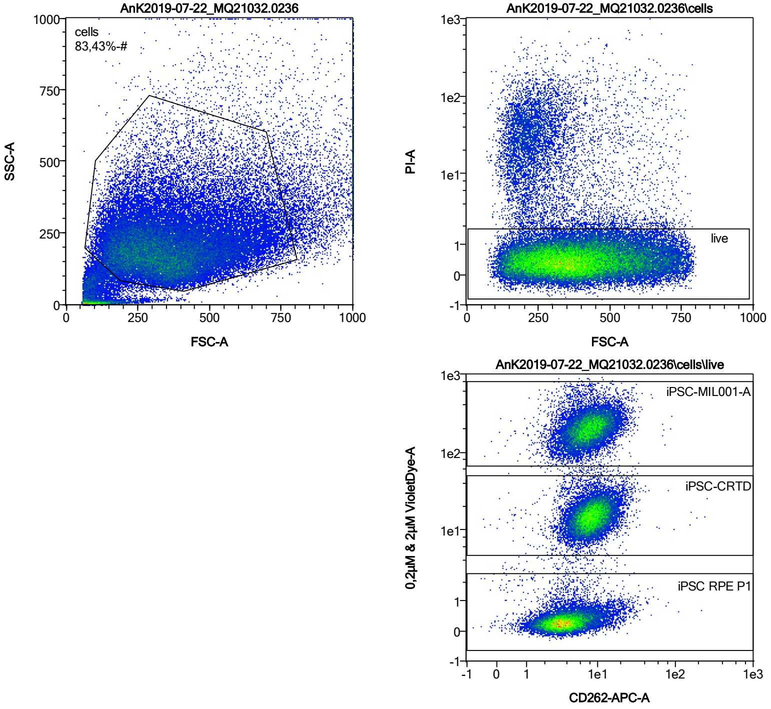


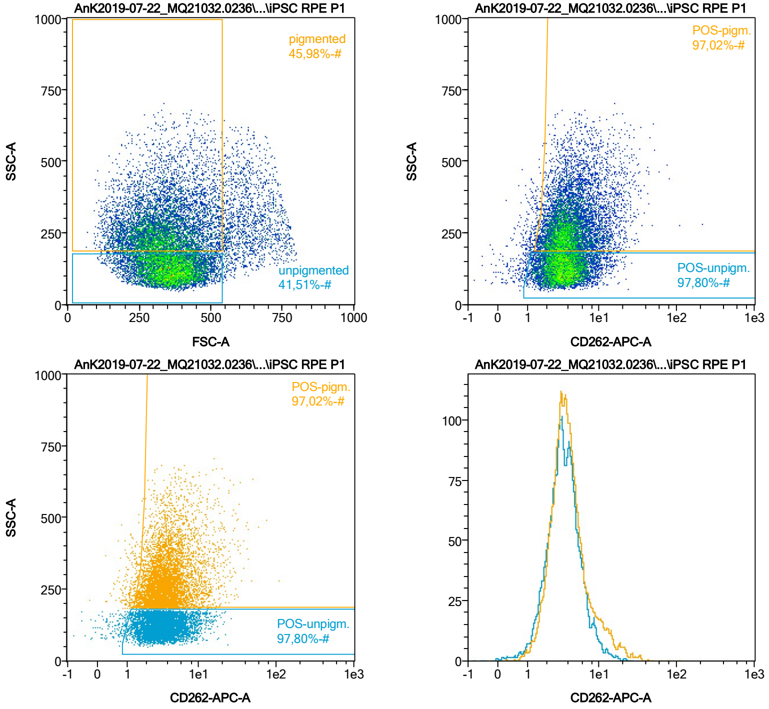


| Name | %-# | CD262-APC-A Mean | CD262-APC-A Median |
|---|---|---|---|
| POS-unpigm. | 97,80 | 3,91 | 3,41 |
| POS-pigm. | 97,02 | 4,86 | 3,72 |

Date:2024-06-14

Operator: Sebastian Knoebel

Version:2.13.0+54.g95abf0f4

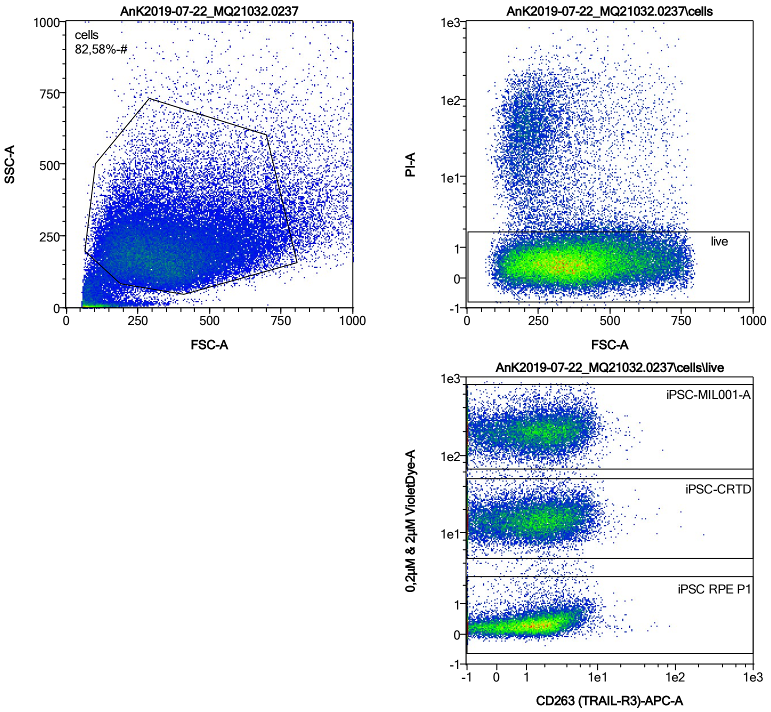


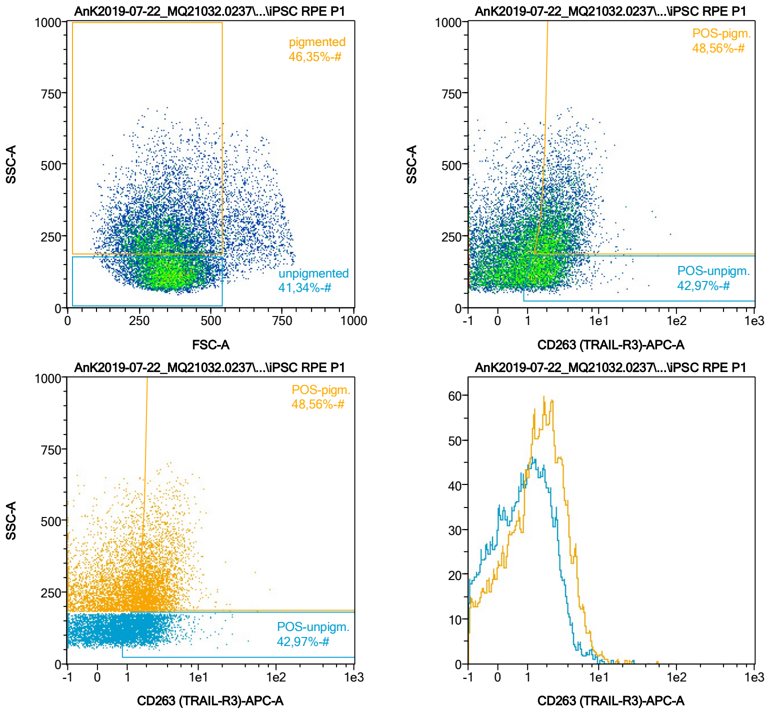


| Name | %-# | CD263 (TRAIL-R3)-APC-A Mean | CD263 (TRAIL-R3)-APC-A Median |
|---|---|---|---|
| POS-unpigm. | 42,97 | 2,34 | 1,92 |
| POS-pigm. | 48,56 | 3,25 | 2,68 |

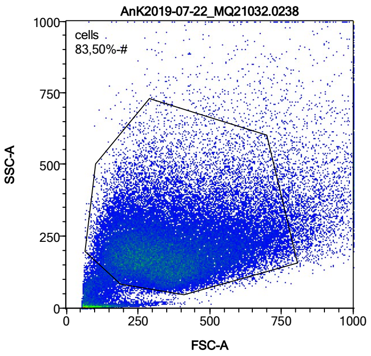


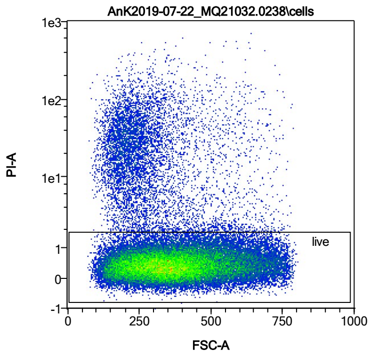


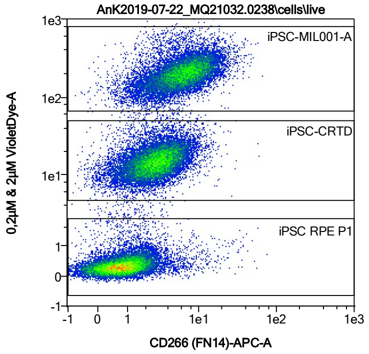


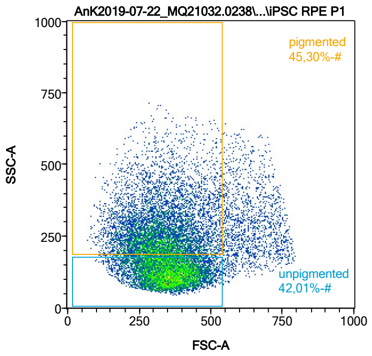


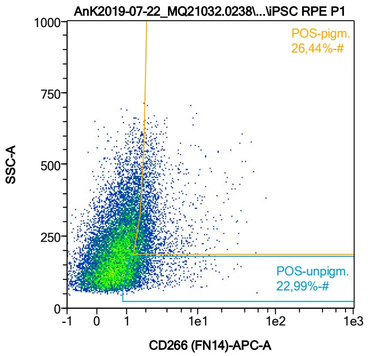


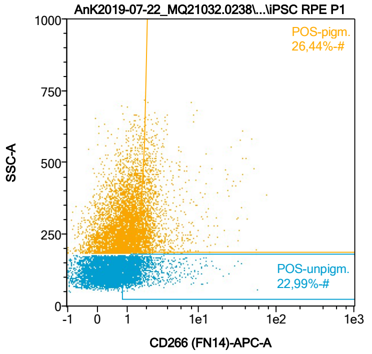


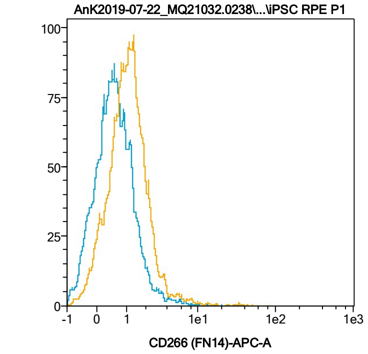


| Name | %-# | CD266 (FN14)-APC-A Mean | CD266 (FN14)-APC-A Median |
|---|---|---|---|
| POS-unpigm. | 22,99 | 2,08 | 1,42 |
| POS-pigm. | 26,44 | 3,13 | 2,09 |

Date:2024-06-14
Operator: Sebastian Knoebel
Version:2.13.0+54.g95abf0f4

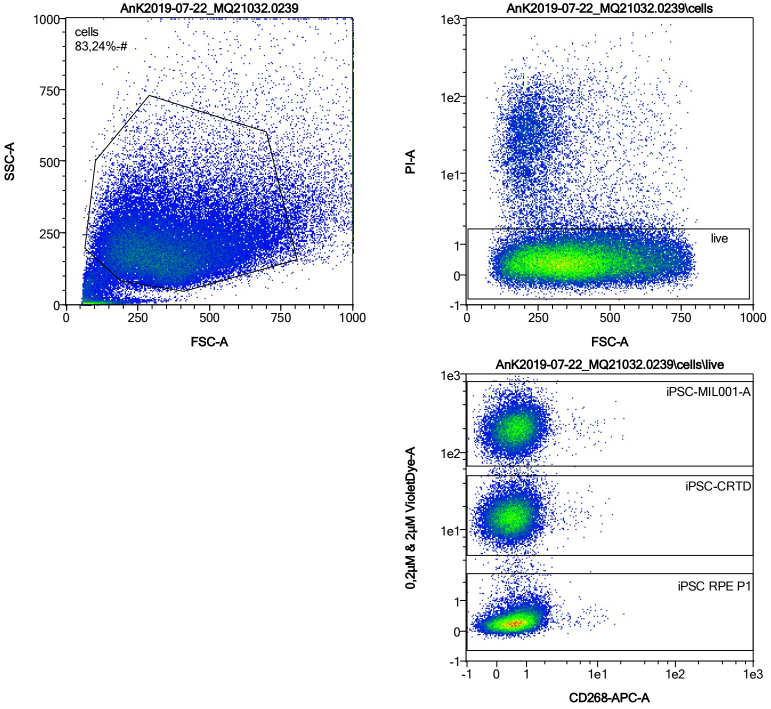


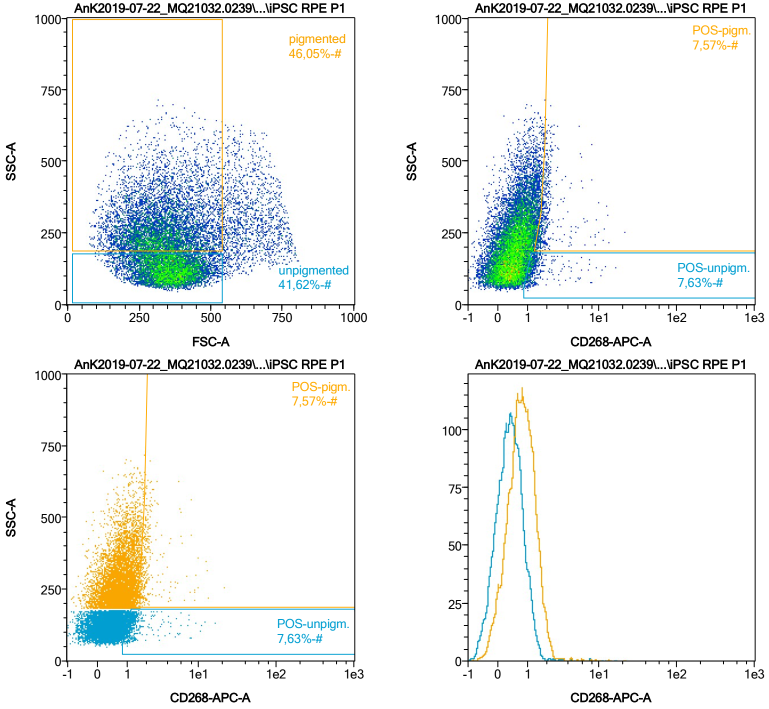


| Name | %-# | CD268-APC-A Mean | CD268-APC-A Median |
|---|---|---|---|
| POS-unpigm. | 7,63 | 1,58 | 1,21 |
| POS-pigm. | 7,57 | 2,19 | 1,80 |

Date:2024-06-14

Operator: Sebastian Knoebel

Version:2.13.0+54.g95abf0f4

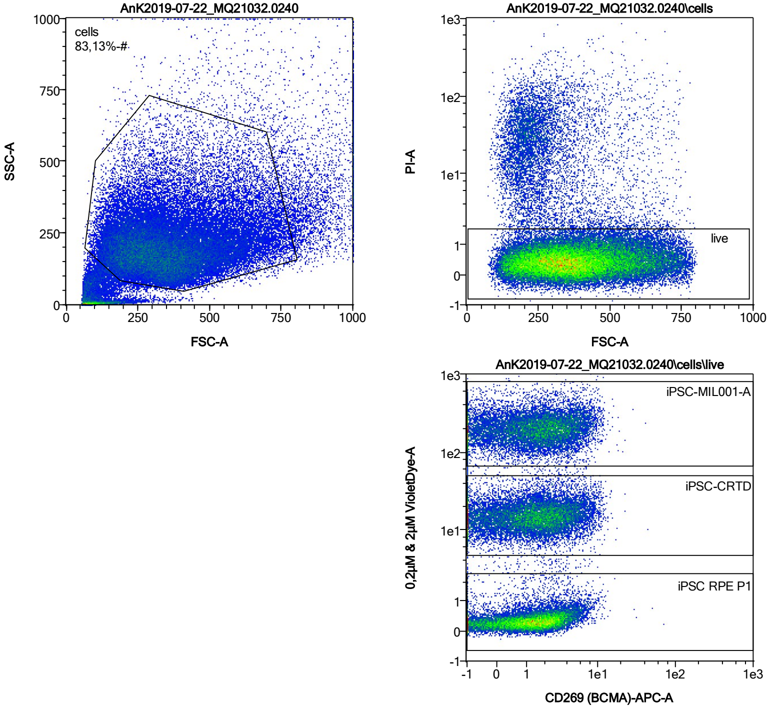


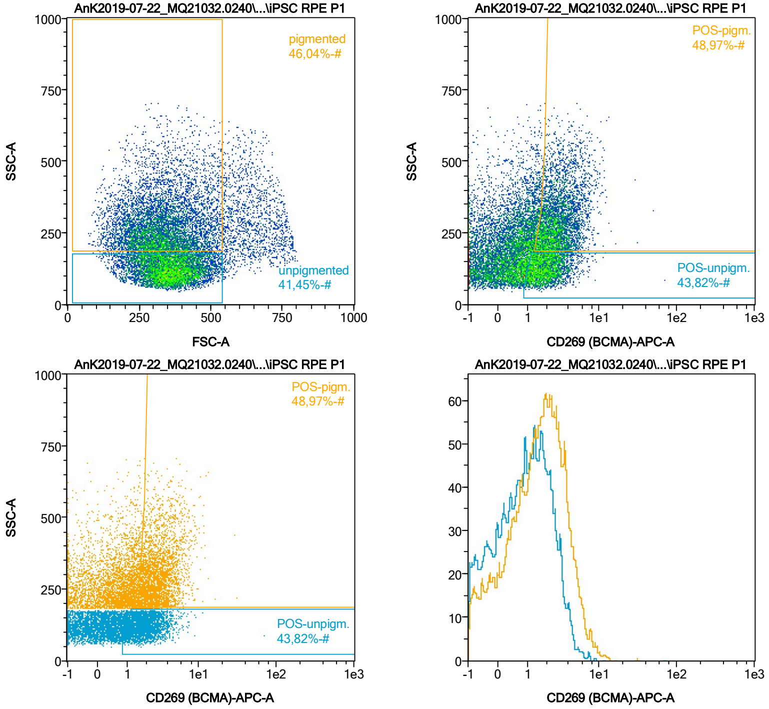


| Name | %-# | CD269 (BCMA)-APC-A Mean | CD269 (BCMA)-APC-A Median |
|---|---|---|---|
| POS-unpigm. | 43,82 | 2,11 | 1,84 |
| POS-pigm. | 48,97 | 3,14 | 2,69 |

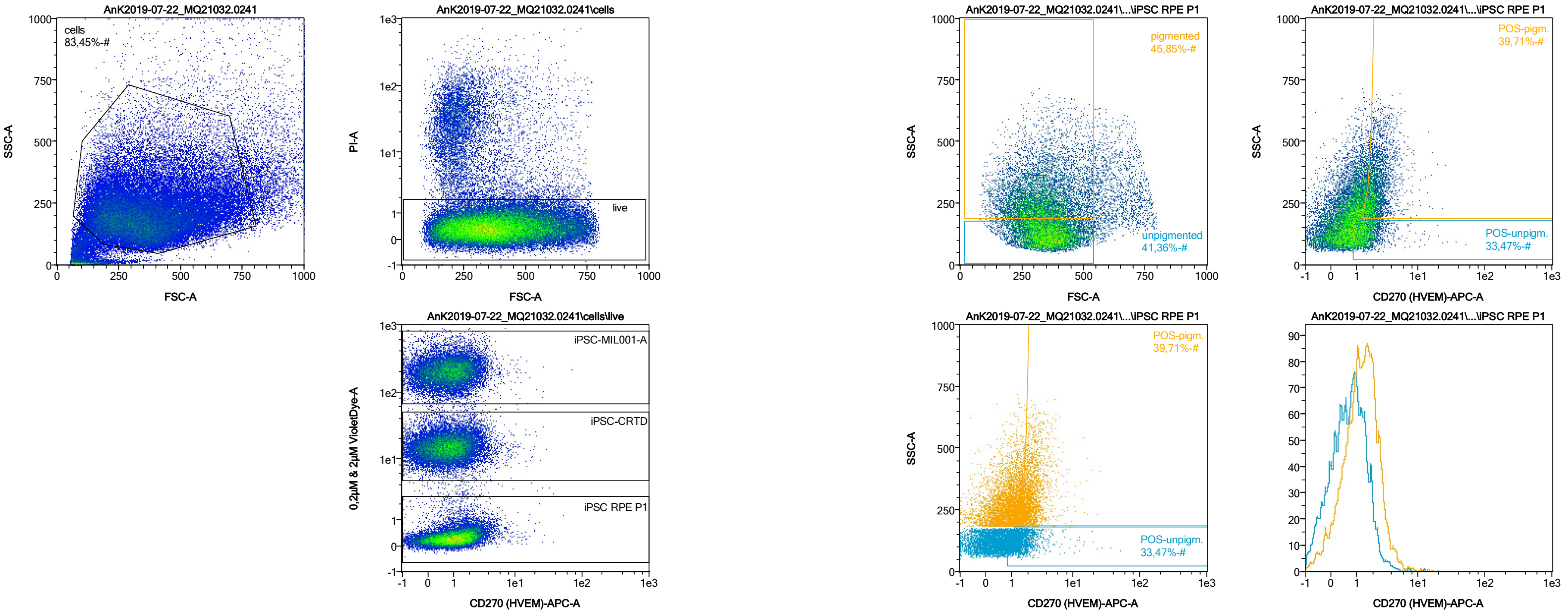


| Name | %-# | CD270 (HVEM)-APC-A Mean | CD270 (HVEM)-APC-A Median |
|---|---|---|---|
| POS-unpigm. | 33,47 | 1,76 | 1,46 |
| POS-pigm. | 39,71 | 2,43 | 2,12 |

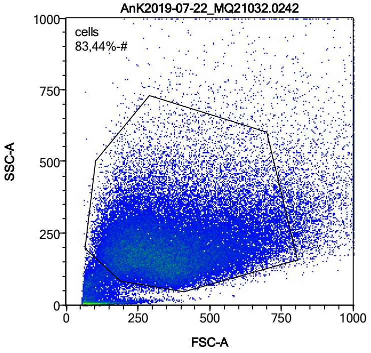


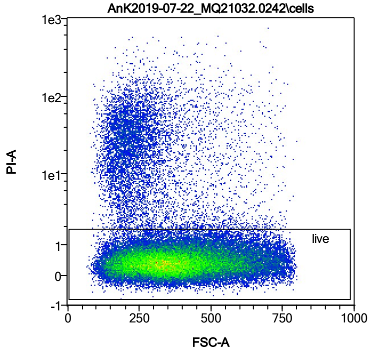


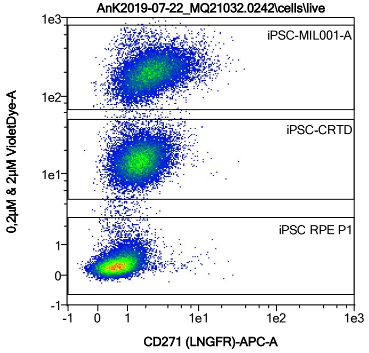


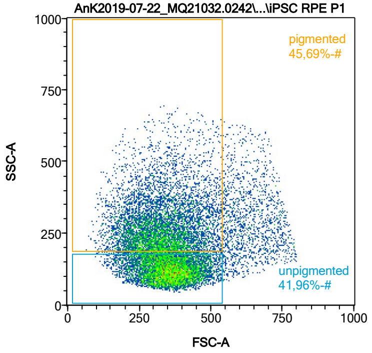


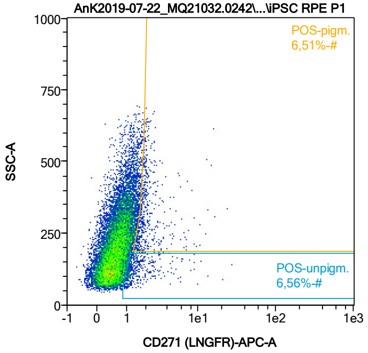


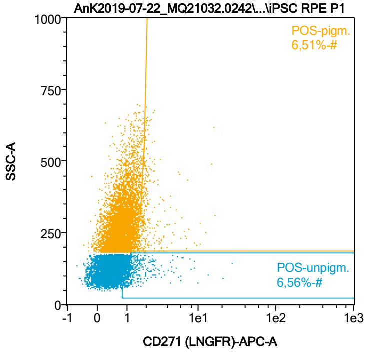


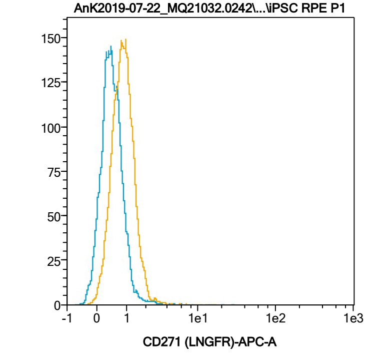


| Name | %-# | CD271 (LNGFR)-APC-A Mean | CD271 (LNGFR)-APC-A Median |
|---|---|---|---|
| POS-unpigm. | 6,56 | 1,76 | 1,23 |
| POS-pigm. | 6,51 | 2,27 | 1,85 |

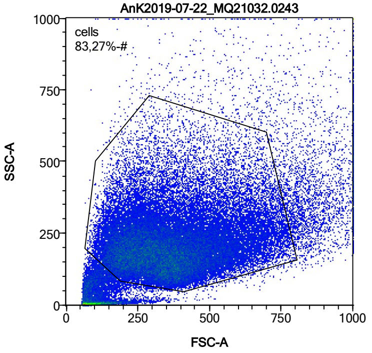


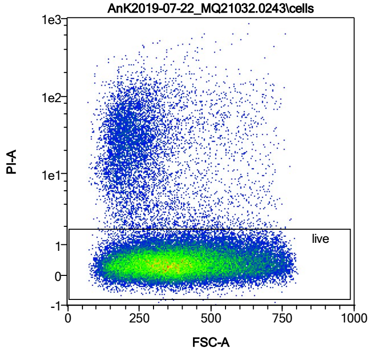


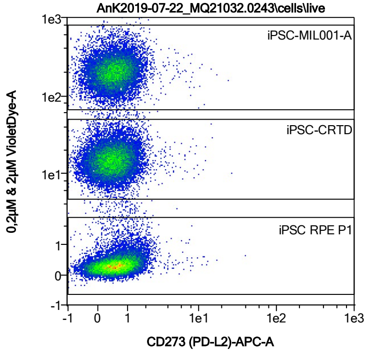


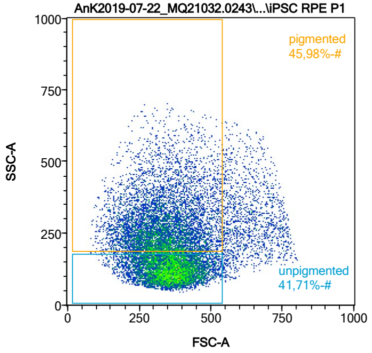


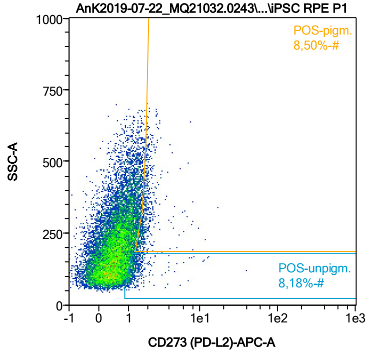


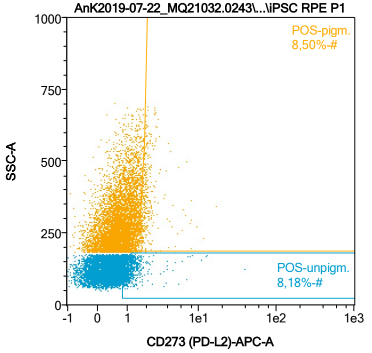


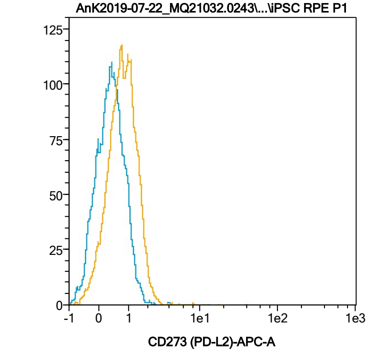


| Name | %-# | CD273 (PD-L2)-APC-A Mean | CD273 (PD-L2)-APC-A Median |
|---|---|---|---|
| POS-unpigm. | 8,18 | 1,52 | 1,20 |
| POS-pigm. | 8,50 | 2,04 | 1,82 |

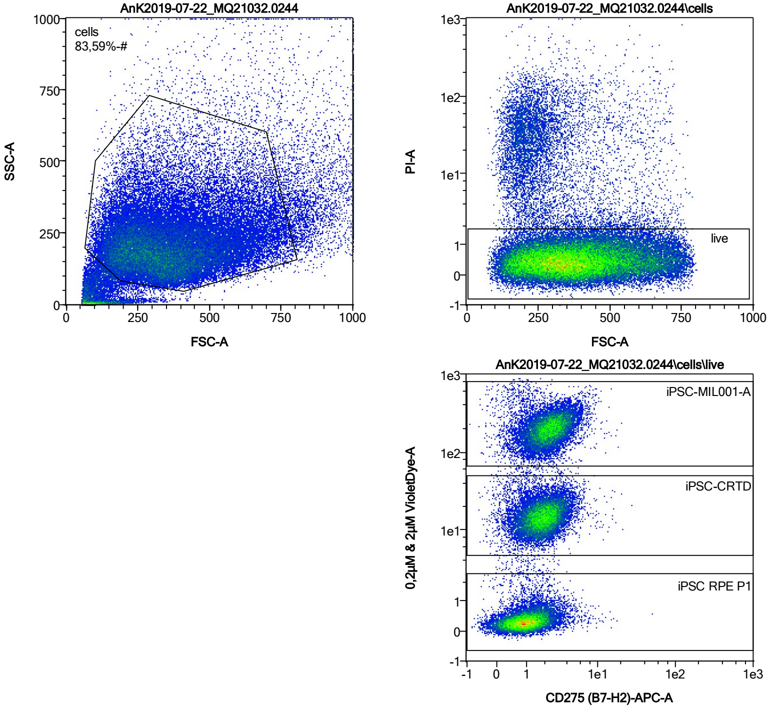


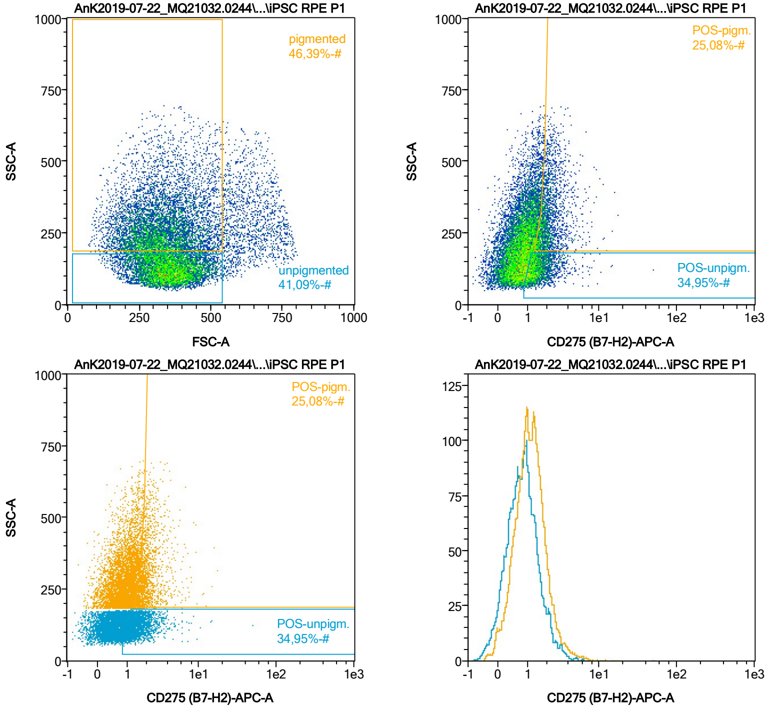


| Name | %-# | CD275 (B7-H2)-APC-A Mean | CD275 (B7-H2)-APC-A Median |
|---|---|---|---|
| POS-unpigm. | 34,95 | 1,65 | 1,39 |
| POS-pigm. | 25,08 | 2,26 | 1,95 |

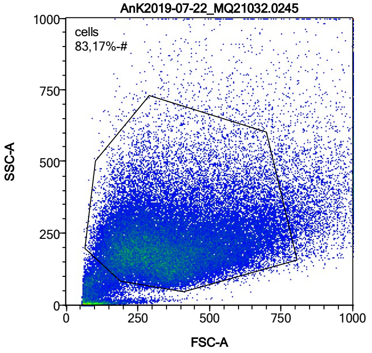


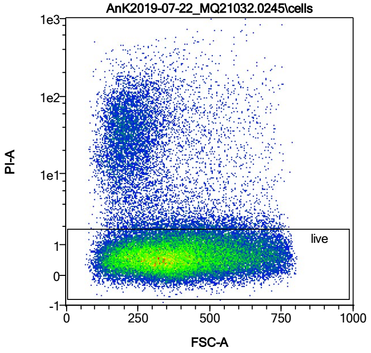


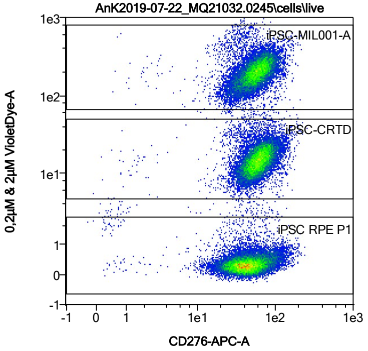


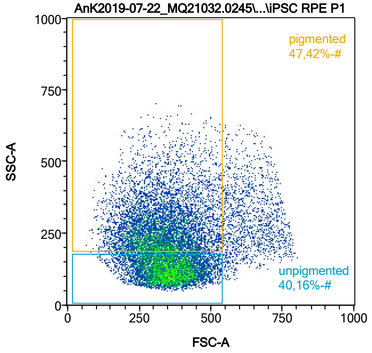


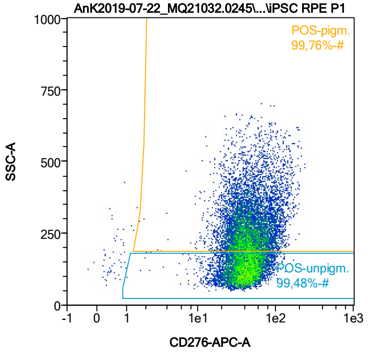


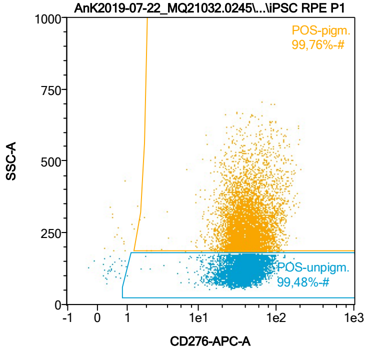


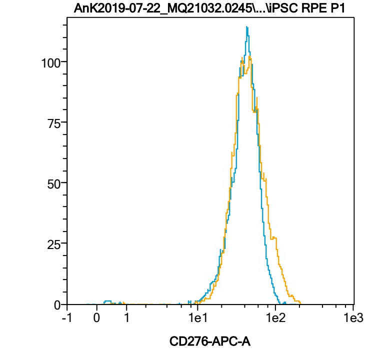


| Name | %-# | CD276-APC-A Mean | CD276-APC-A Median |
|---|---|---|---|
| POS-unpigm. | 99,48 | 42,20 | 40,49 |
| POS-pigm. | 99,76 | 49,12 | 43,15 |

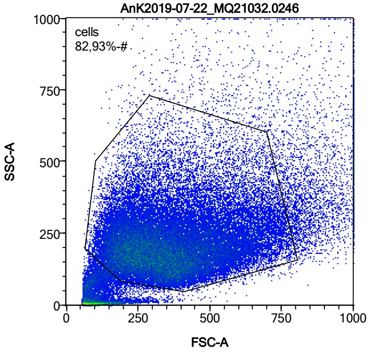


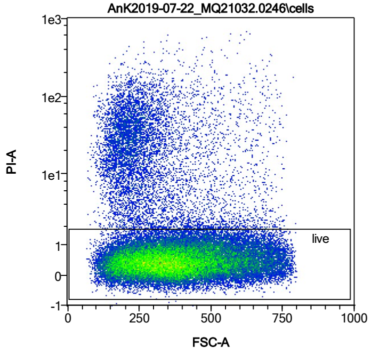


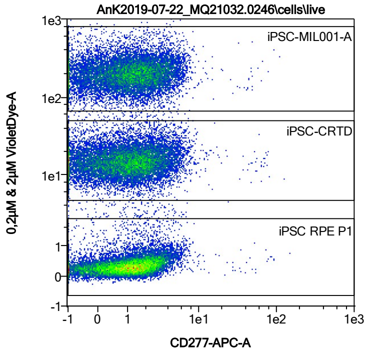


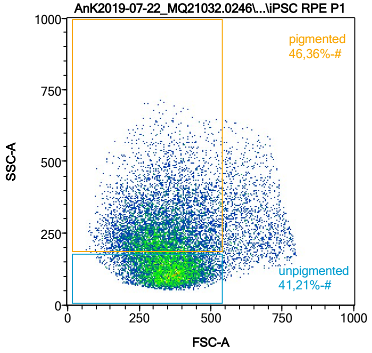


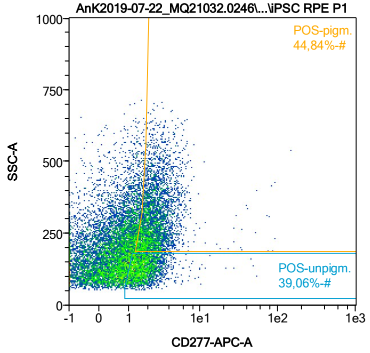


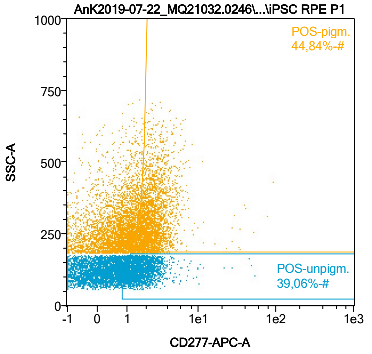


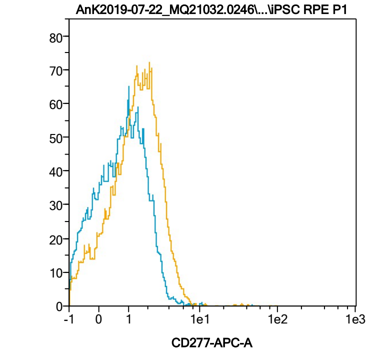


| Name | %-# | CD277-APC-A Mean | CD277-APC-A Median |
|---|---|---|---|
| POS-unpigm. | 39,06 | 1,93 | 1,65 |
| POS-pigm. | 44,84 | 2,80 | 2,36 |

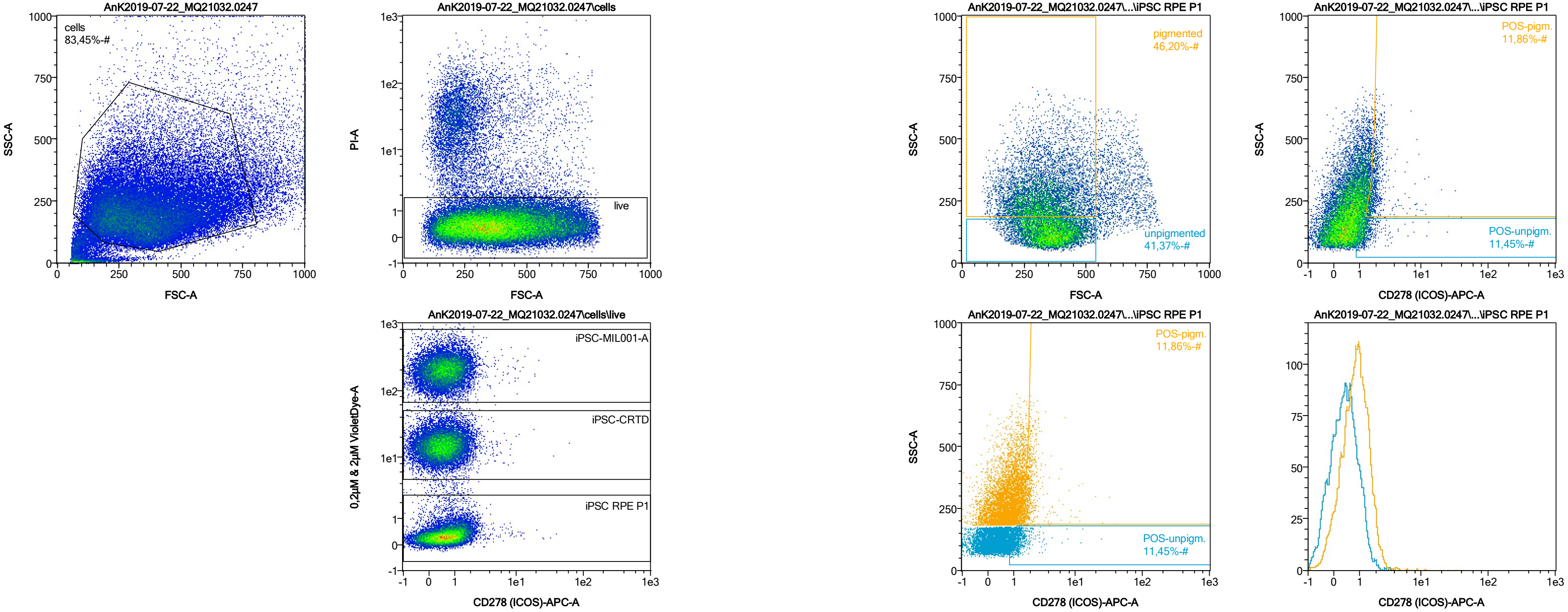


| Name | %-# | CD278 (ICOS)-APC-A Mean | CD278 (ICOS)-APC-A Median |
|---|---|---|---|
| POS-unpigm. | 11,45 | 1,43 | 1,24 |
| POS-pigm. | 11,86 | 2,23 | 1,86 |

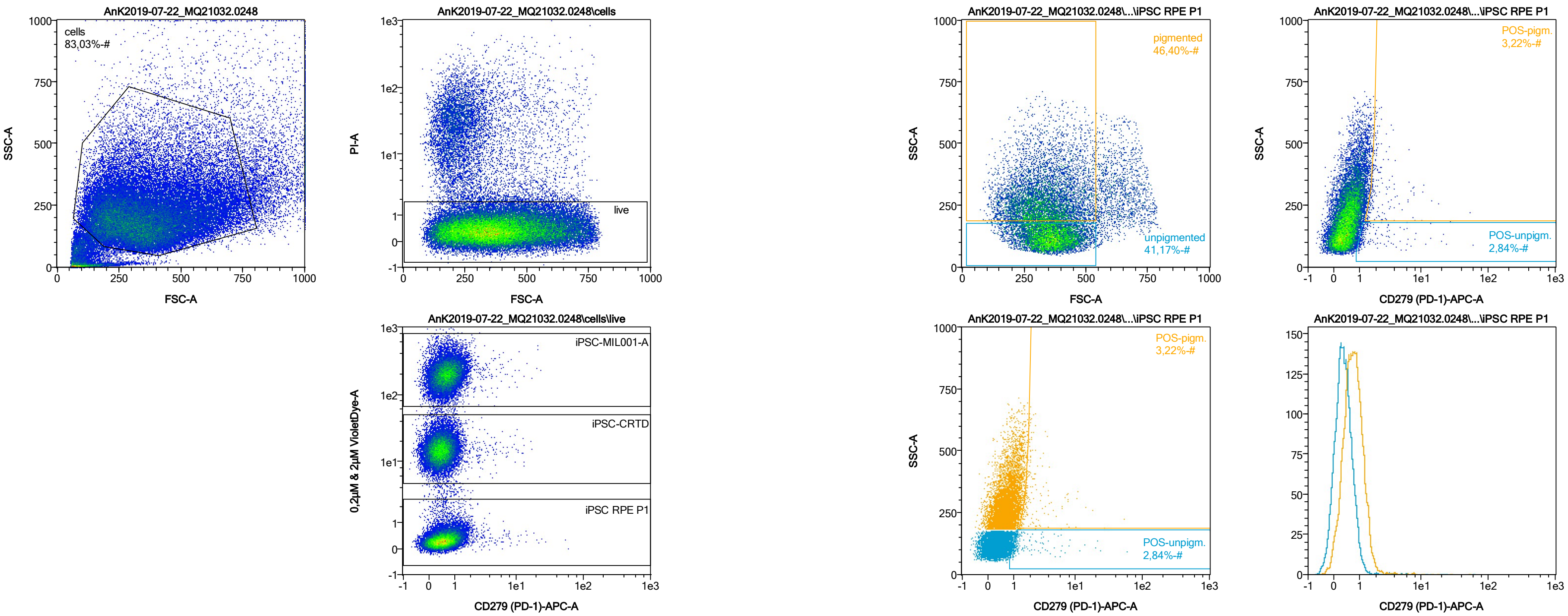


| Name | %-# | CD279 (PD-1)-APC-A Mean | CD279 (PD-1)-APC-A Median |
|---|---|---|---|
| POS-unpigm. | 2,84 | 2,61 | 1,20 |
| POS-pigm. | 3,22 | 2,88 | 1,87 |

Date:2024-06-14

Operator: Sebastian Knoebel

Version:2.13.0+54.g95abf0f4

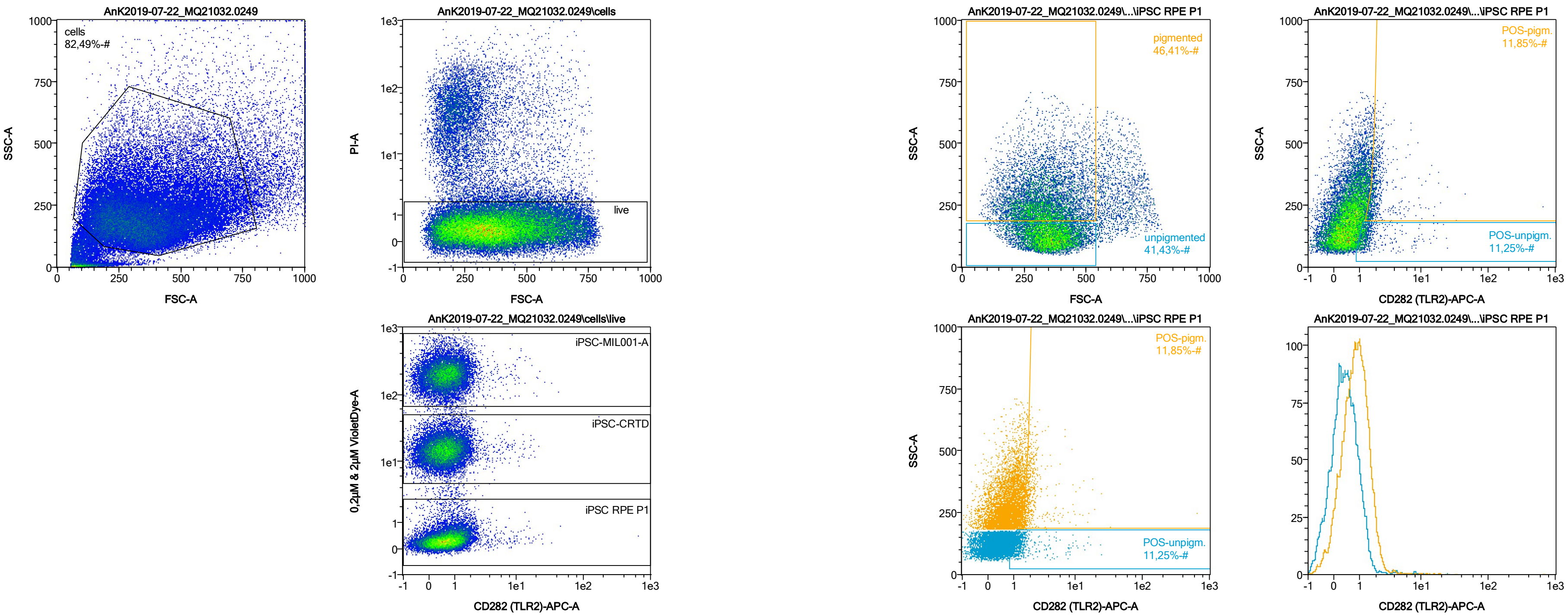


| Name | %-# | CD282 (TLR2)-APC-A Mean | CD282 (TLR2)-APC-A Median |
|---|---|---|---|
| POS-unpigm. | 11,25 | 1,80 | 1,27 |
| POS-pigm. | 11,85 | 3,22 | 1,86 |

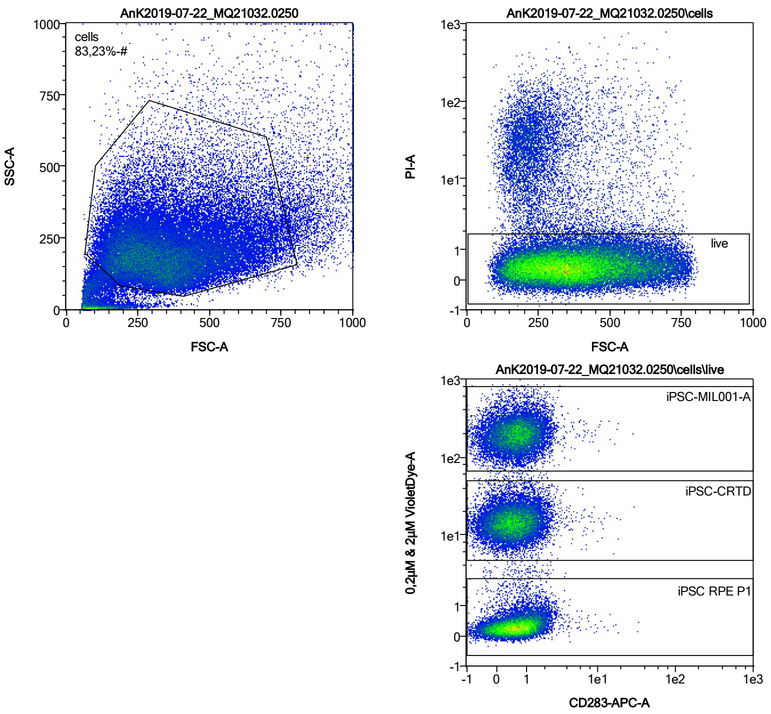


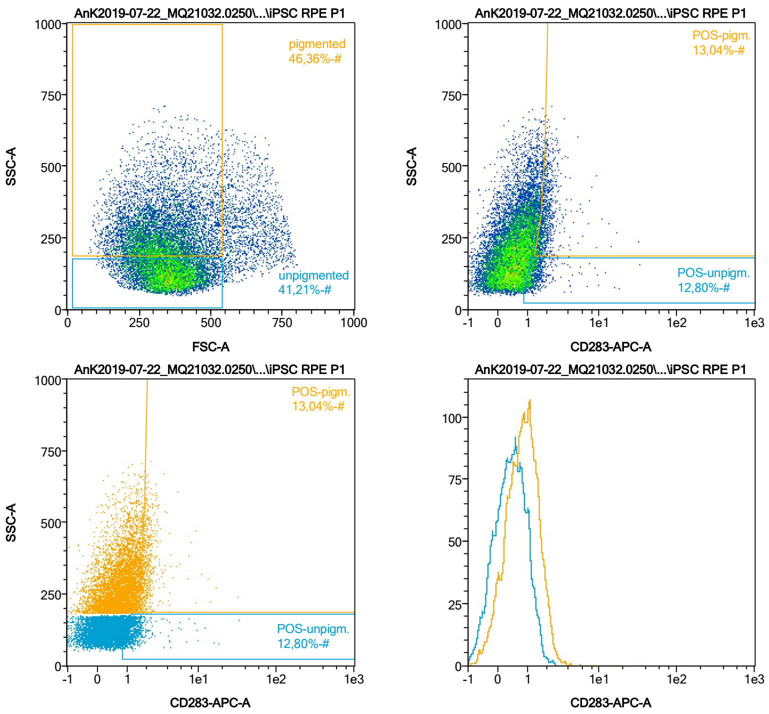


| Name | %-# | CD283-APC-A Mean | CD283-APC-A Median |
|---|---|---|---|
| POS-unpigm. | 12,80 | 1,49 | 1,24 |
| POS-pigm. | 13,04 | 2,09 | 1,87 |

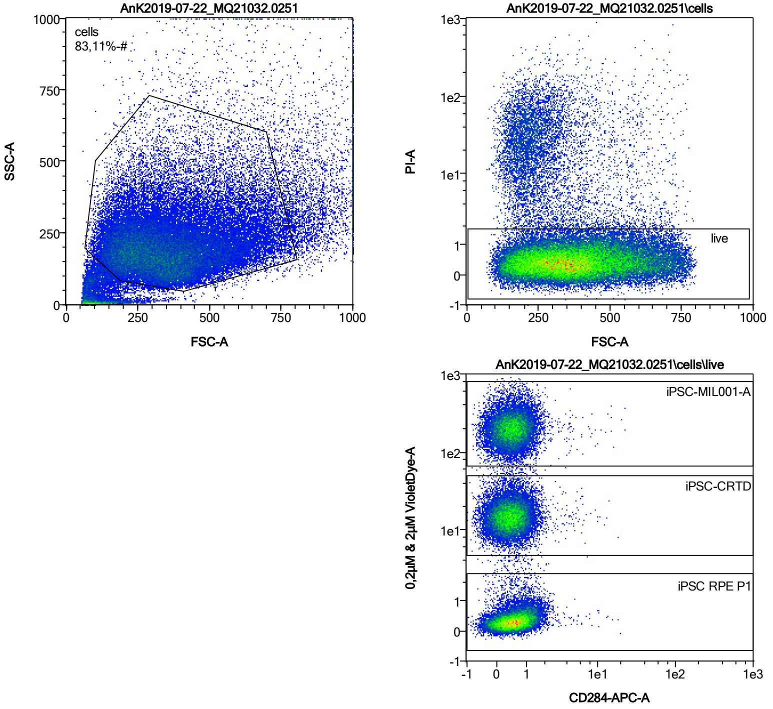


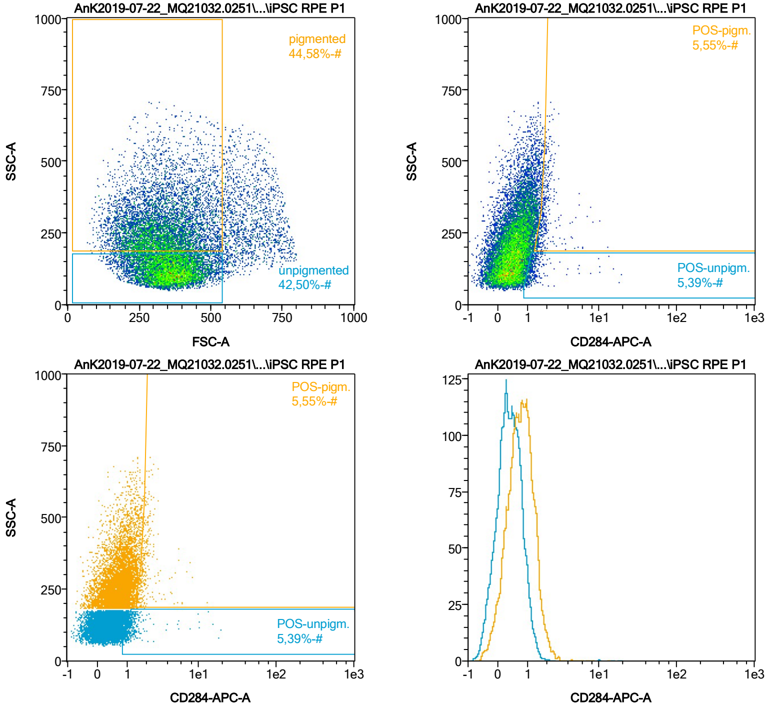


| Name | %-# | CD284-APC-A Mean | CD284-APC-A Median |
|---|---|---|---|
| POS-unpigm. | 5,39 | 1,56 | 1,18 |
| POS-pigm. | 5,55 | 2,04 | 1,82 |

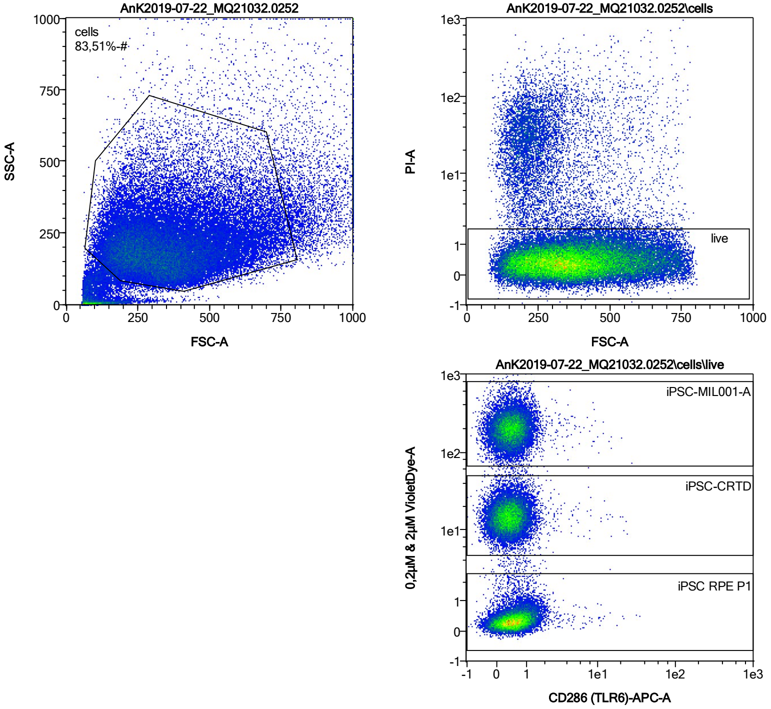


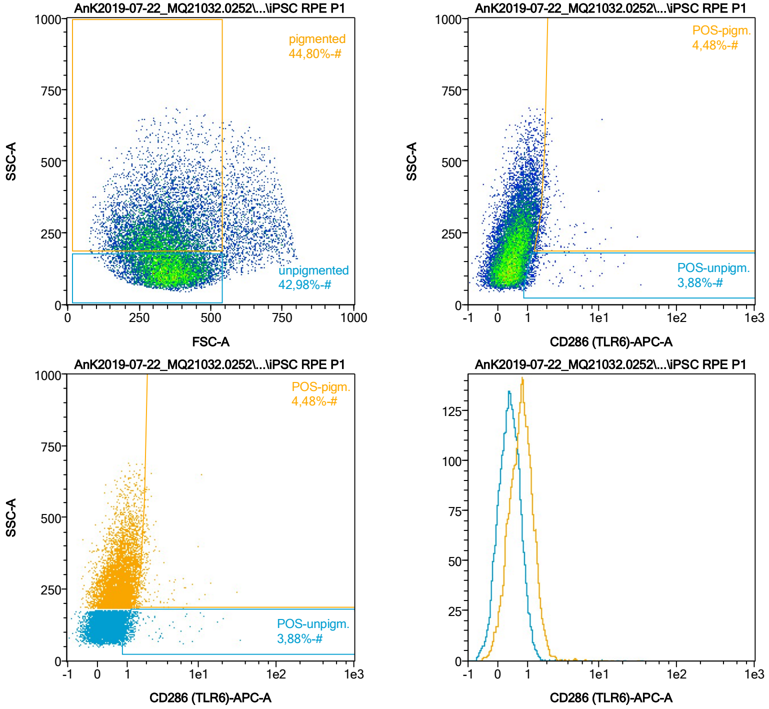


| Name | %-# | CD286 (TLR6)-APC-A Mean | CD286 (TLR6)-APC-A Median |
|---|---|---|---|
| POS-unpigm. | 3,88 | 1,92 | 1,17 |
| POS-pigm. | 4,48 | 2,32 | 1,81 |

Date:2024-06-14
Operator: Sebastian Knoebel
Version:2.13.0+54.g95abf0f4

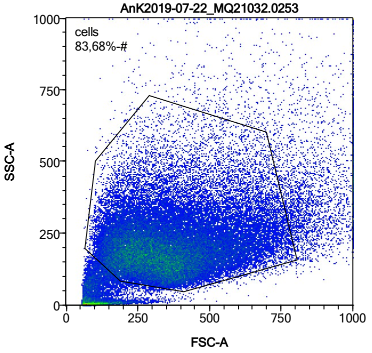


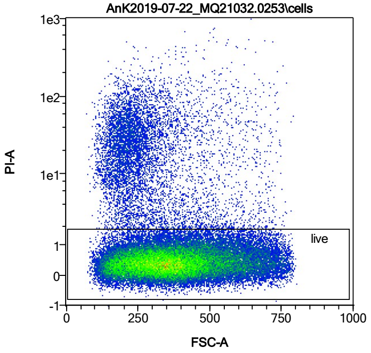


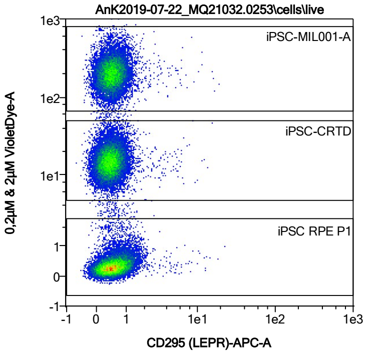


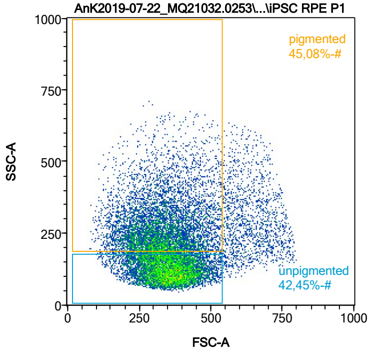


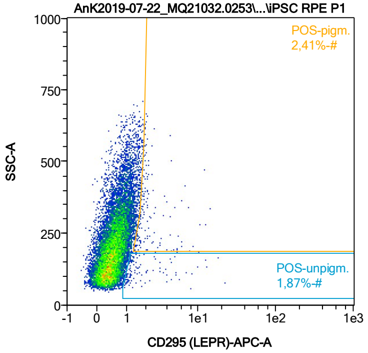


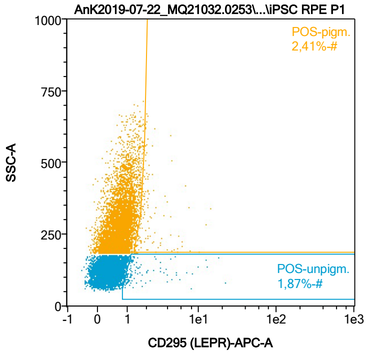


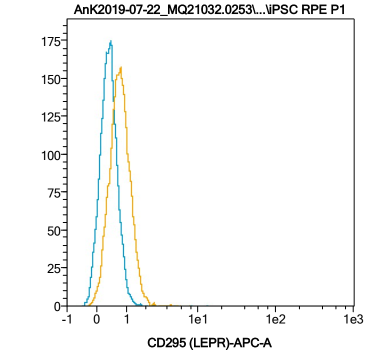


| Name | %-# | CD295 (LEPR)-APC-A Mean | CD295 (LEPR)-APC-A Median |
|---|---|---|---|
| POS-unpigm. | 1,87 | 2,37 | 1,23 |
| POS-pigm. | 2,41 | 2,45 | 1,90 |

Date:2024-06-14

Operator: Sebastian Knoebel

Version:2.13.0+54.g95abf0f4

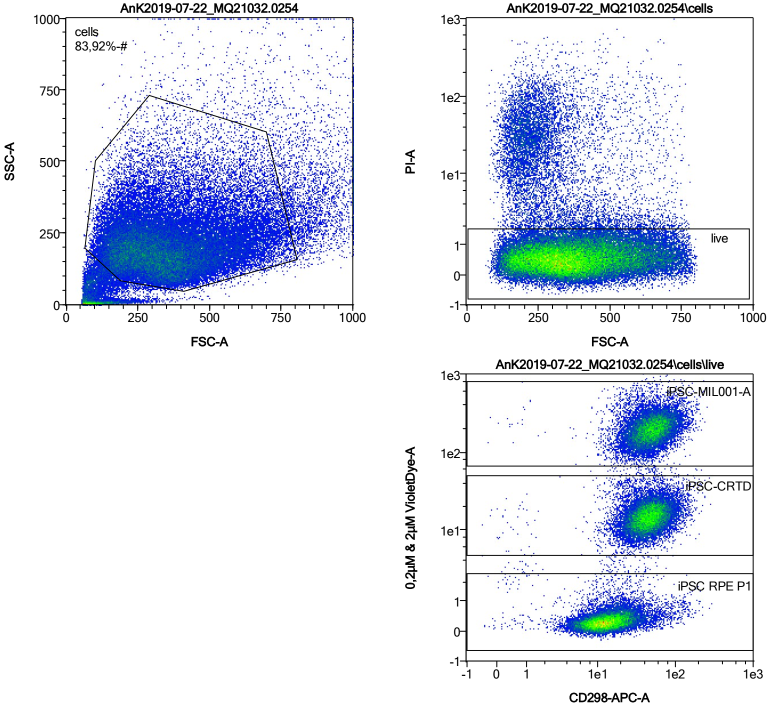


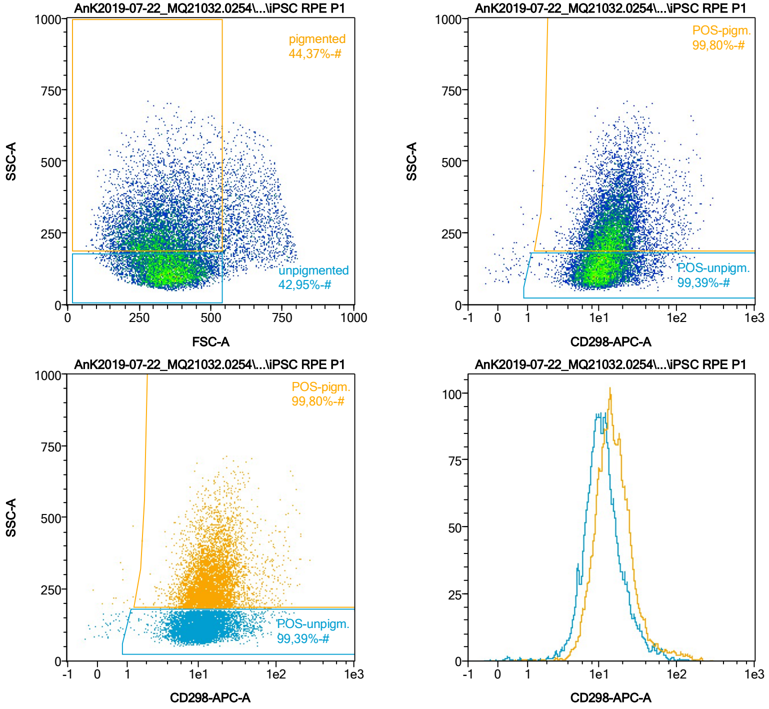


| Name | %-# | CD298-APC-A Mean | CD298-APC-A Median |
|---|---|---|---|
| POS-unpigm. | 99,39 | 13,36 | 10,72 |
| POS-pigm. | 99,80 | 18,43 | 14,62 |

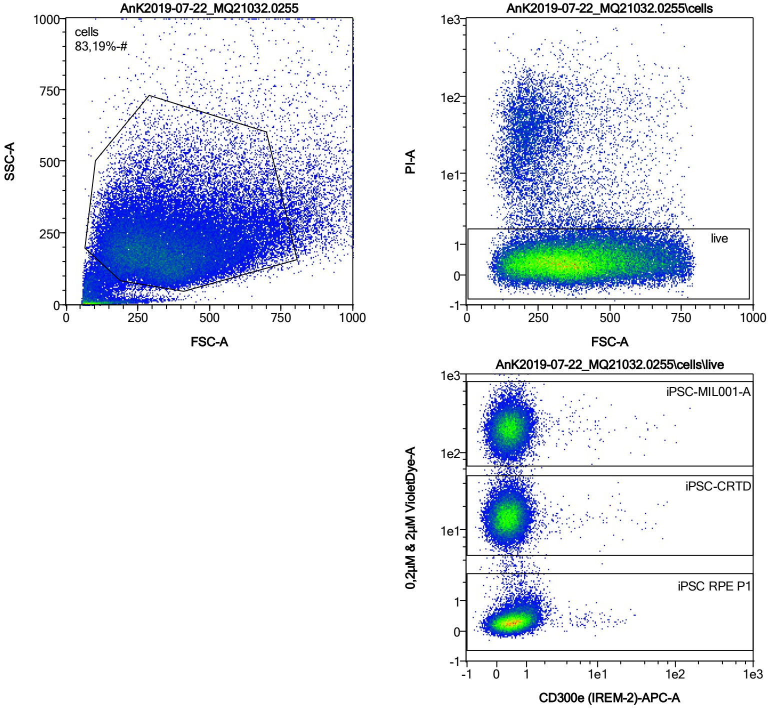


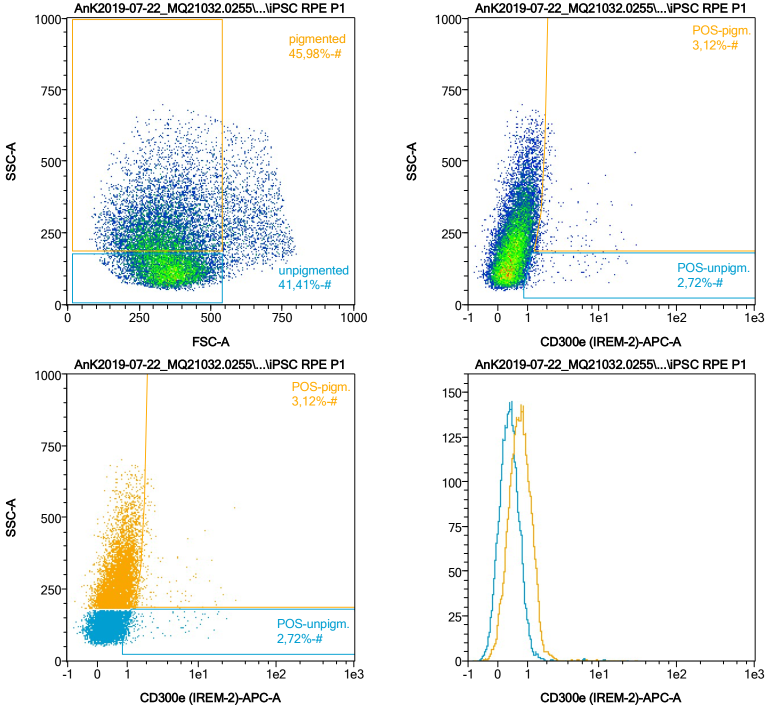


| Name | %-# | CD300e (IREM-2)-APC-A Mean | CD300e (IREM-2)-APC-A Median |
|---|---|---|---|
| POS-unpigm. | 2,72 | 2,77 | 1,29 |
| POS-pigm. | 3,12 | 3,21 | 1,87 |

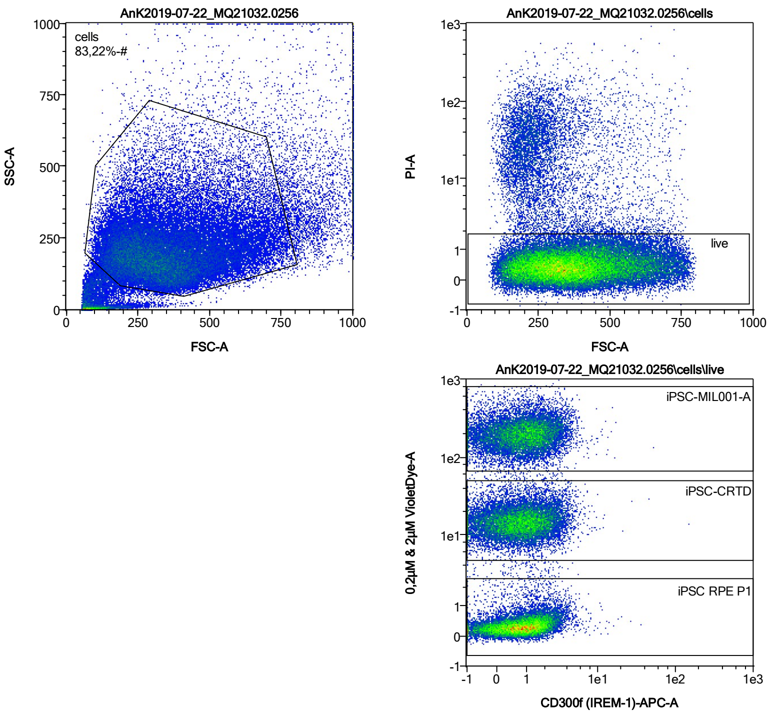


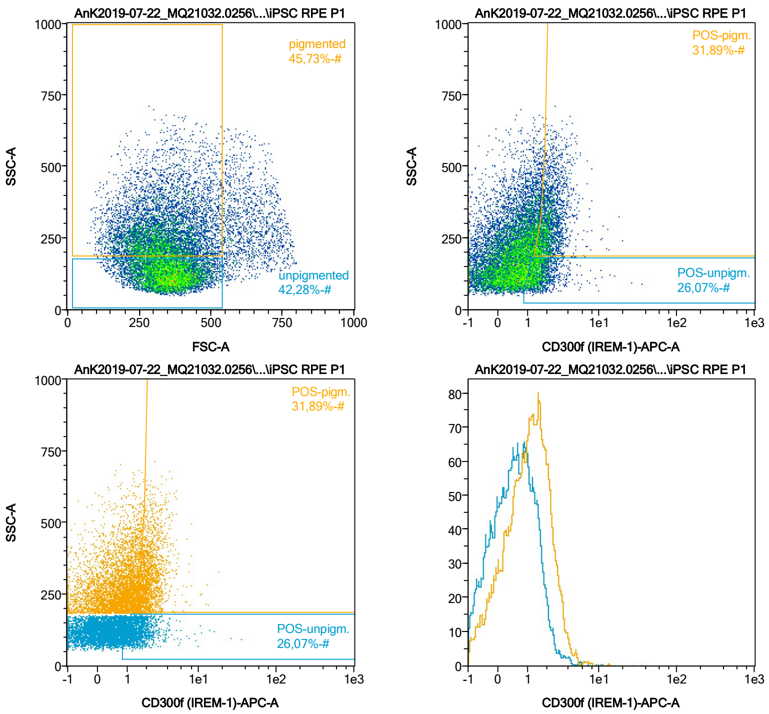


| Name | %-# | CD300f (IREM-1)-APC-A Mean | CD300f (IREM-1)-APC-A Median |
|---|---|---|---|
| POS-unpigm. | 26,07 | 1,68 | 1,44 |
| POS-pigm. | 31,89 | 2,32 | 2,08 |

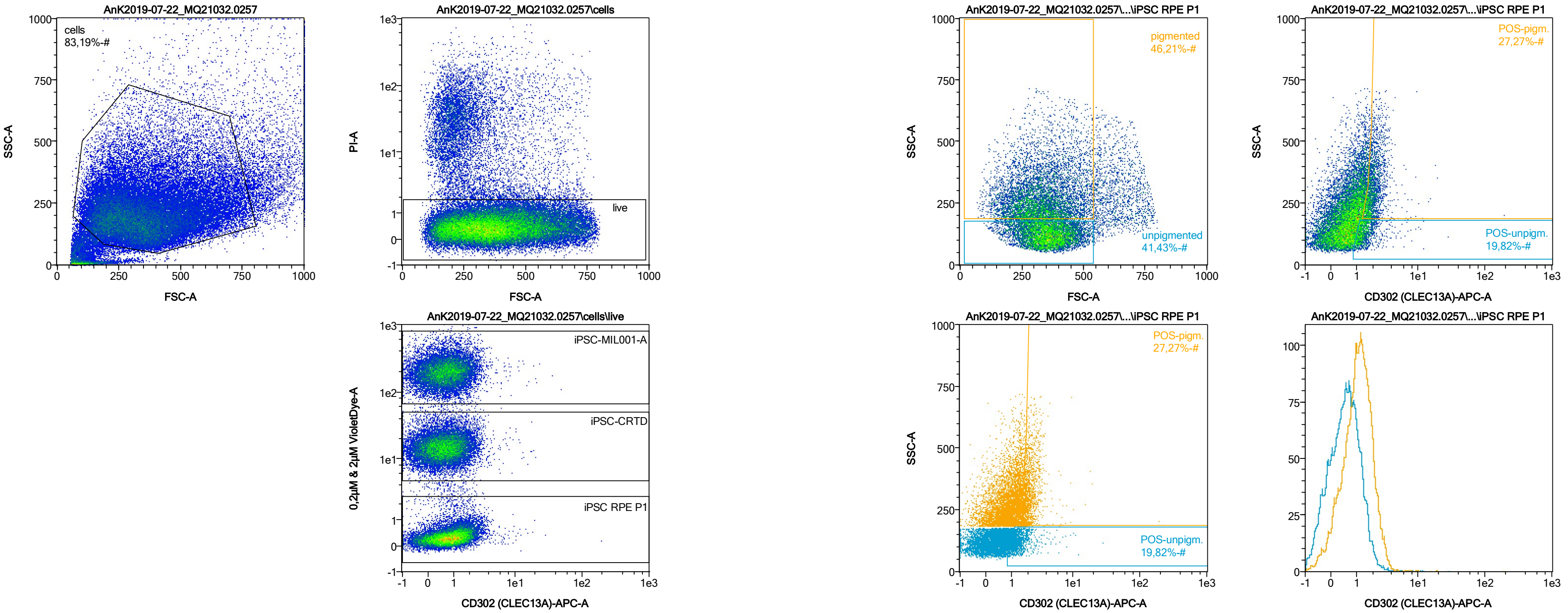


| Name | %-# | CD302 (CLEC13A)-APC-A Mean | CD302 (CLEC13A)-APC-A Median |
|---|---|---|---|
| POS-unpigm. | 19,82 | 1,53 | 1,30 |
| POS-pigm. | 27,27 | 2,22 | 1,92 |

Date:2024-06-14

Operator: Sebastian Knoebel

Version:2.13.0+54.g95abf0f4

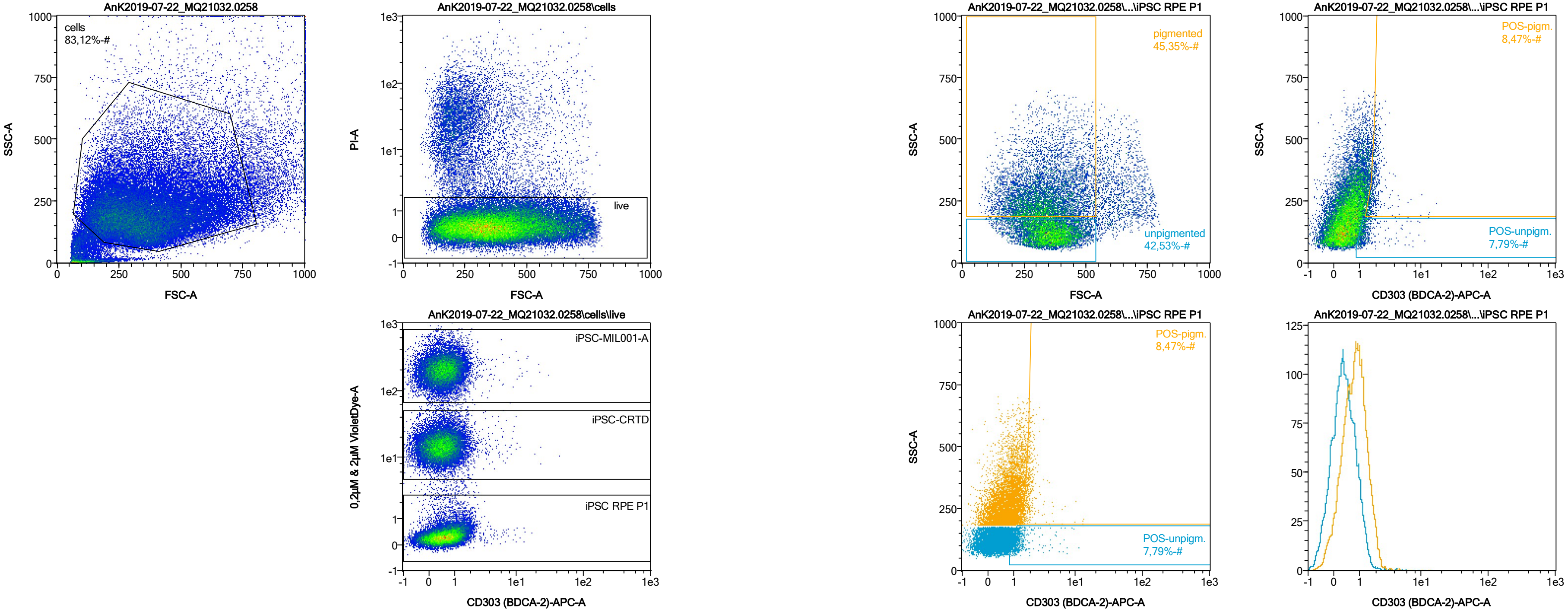


| Name | %-# | CD303 (BDCA-2)-APC-A Mean | CD303 (BDCA-2)-APC-A Median |
|---|---|---|---|
| POS-unpigm. | 7,79 | 1,50 | 1,22 |
| POS-pigm. | 8,47 | 2,00 | 1,82 |

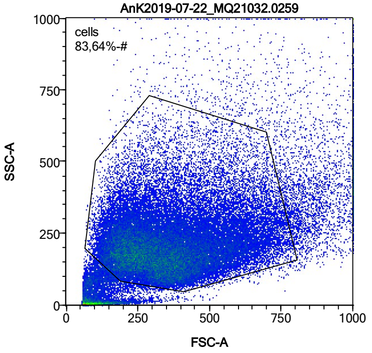


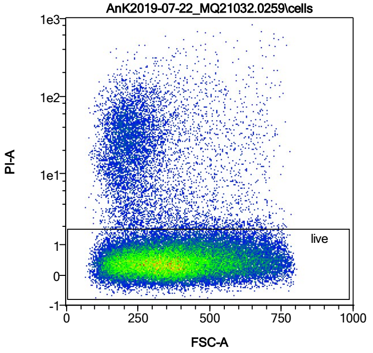


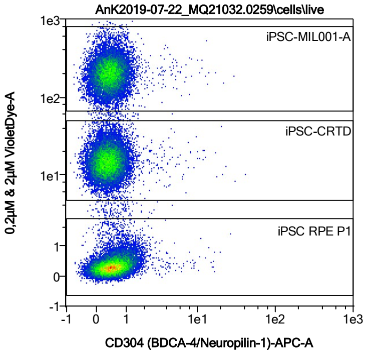


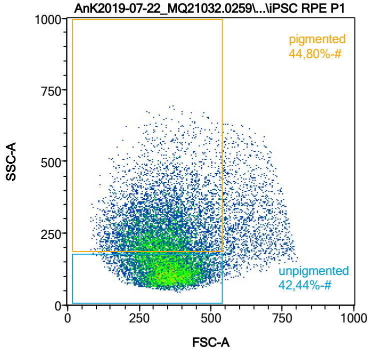


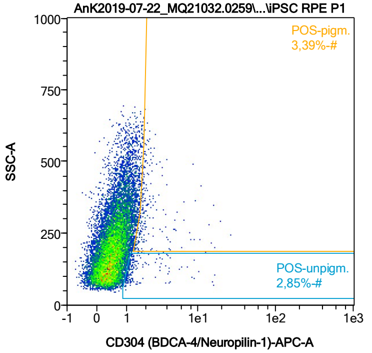


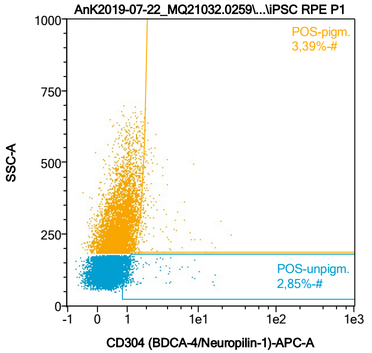


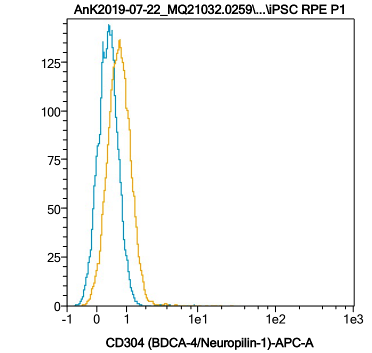


| Name | %-# | CD304 (BDCA-4/Neuropilin-1)-APC-A Mean | CD304 (BDCA-4/Neuropilin-1)-APC-A Median |
|---|---|---|---|
| POS-unpigm. | 2,85 | 1,92 | 1,17 |
| POS-pigm. | 3,39 | 2,64 | 1,86 |

Date:2024-06-14
Operator: Sebastian Knoebel
Version:2.13.0+54.g95abf0f4

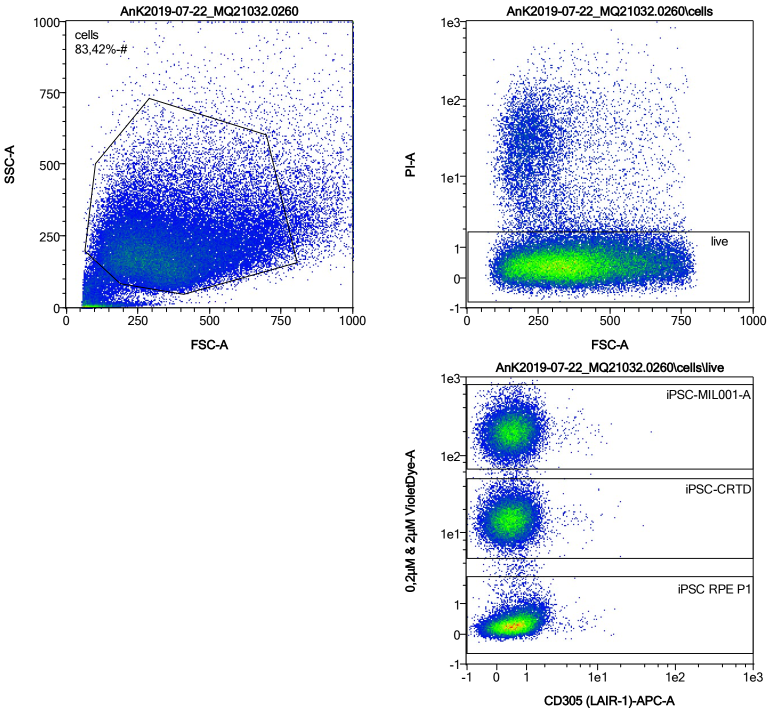


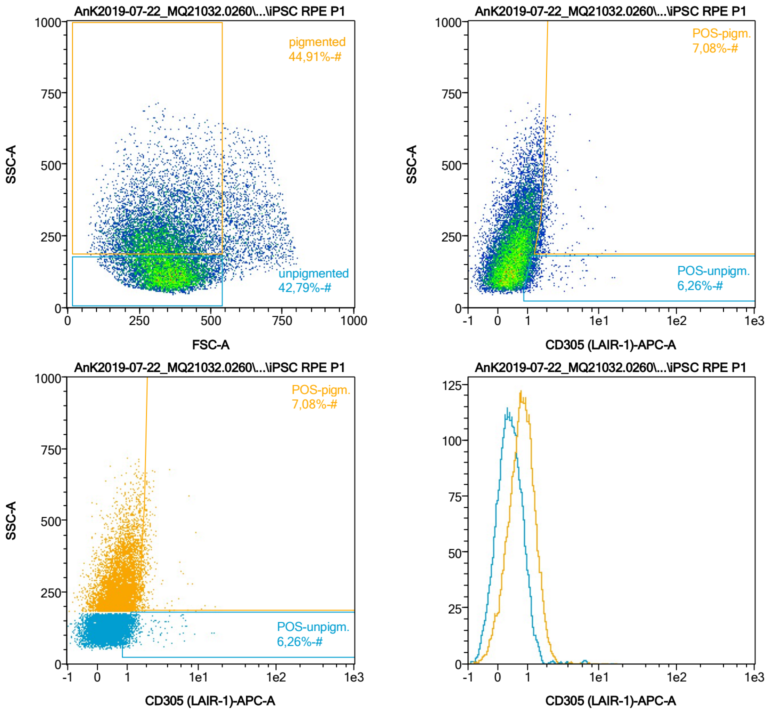


| Name | %-# | CD305 (LAIR-1)-APC-A Mean | CD305 (LAIR-1)-APC-A Median |
|---|---|---|---|
| POS-unpigm. | 6,26 | 1,63 | 1,21 |
| POS-pigm. | 7,08 | 2,07 | 1,81 |

Date:2024-06-14
Operator: Sebastian Knoebel
Version:2.13.0+54.g95abf0f4

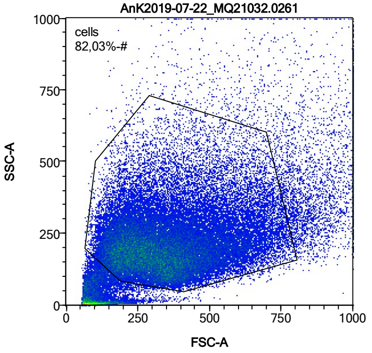


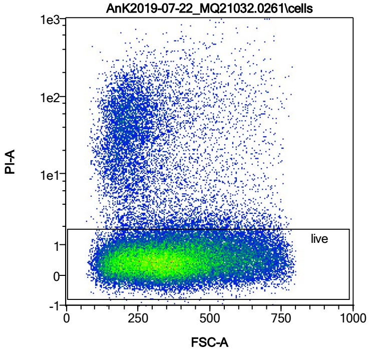


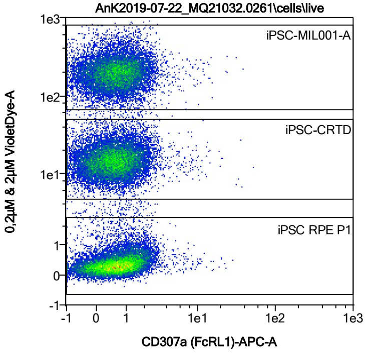


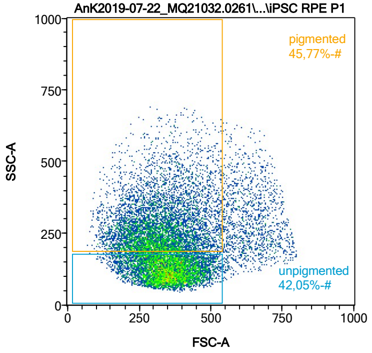


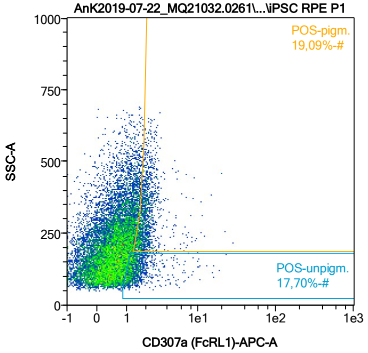


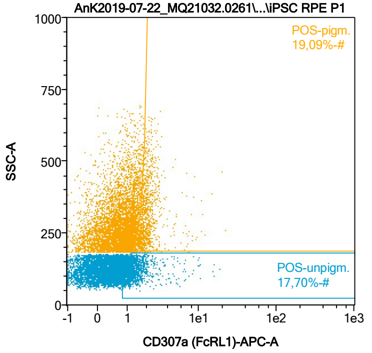


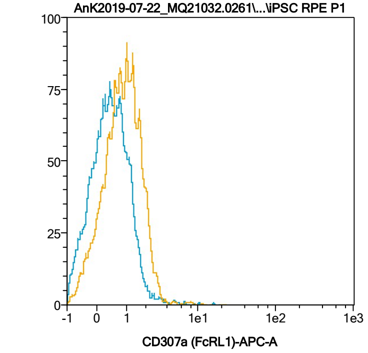


| Name | %-# | CD307a (FcRL1)-APC-A Mean | CD307a (FcRL1)-APC-A Median |
|---|---|---|---|
| POS-unpigm. | 17,70 | 1,67 | 1,31 |
| POS-pigm. | 19,09 | 2,20 | 1,93 |

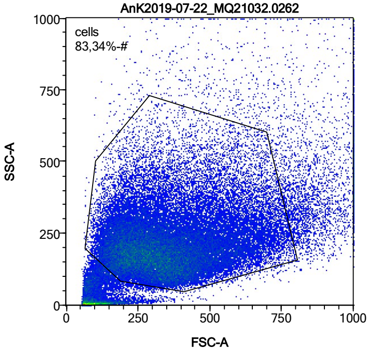


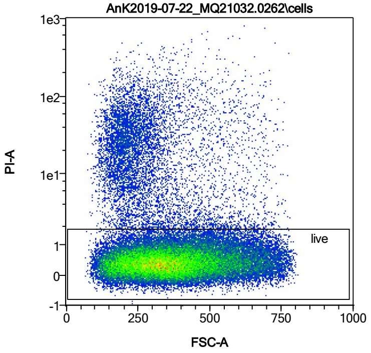


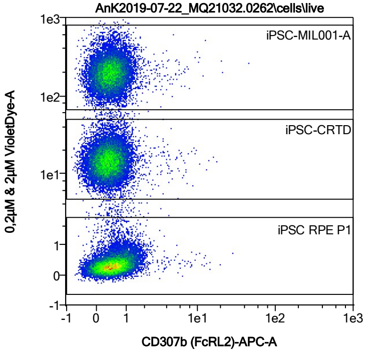


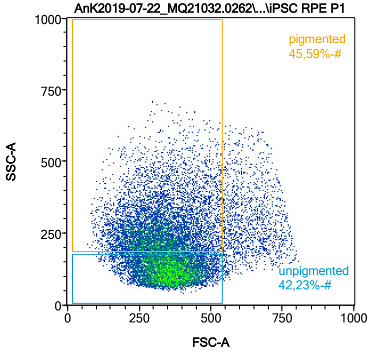


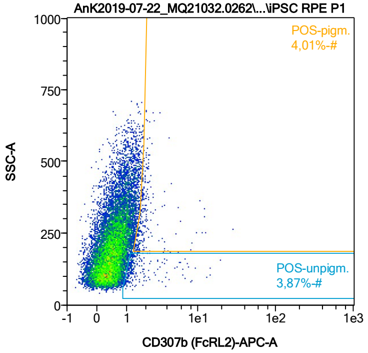


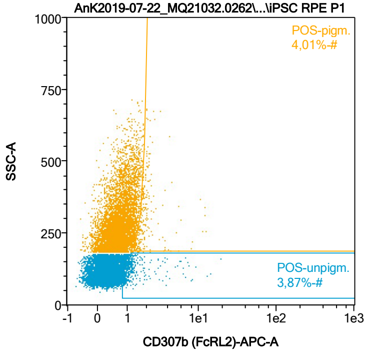


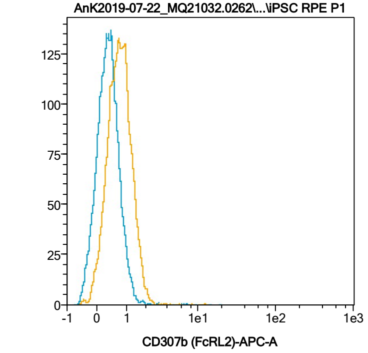


| Name | %-# | CD307b (FcRL2)-APC-A Mean | CD307b (FcRL2)-APC-A Median |
|---|---|---|---|
| POS-unpigm. | 3,87 | 1,88 | 1,23 |
| POS-pigm. | 4,01 | 2,22 | 1,81 |

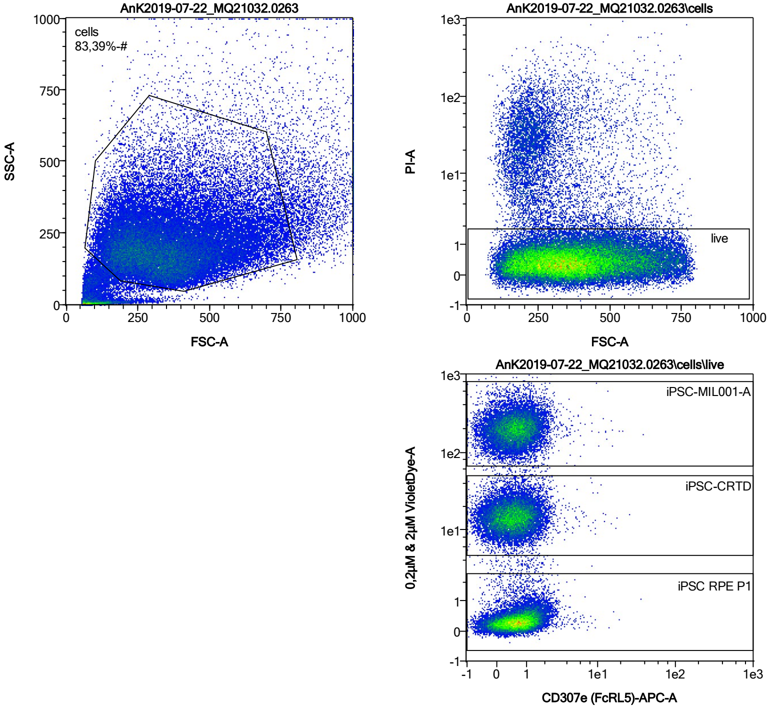


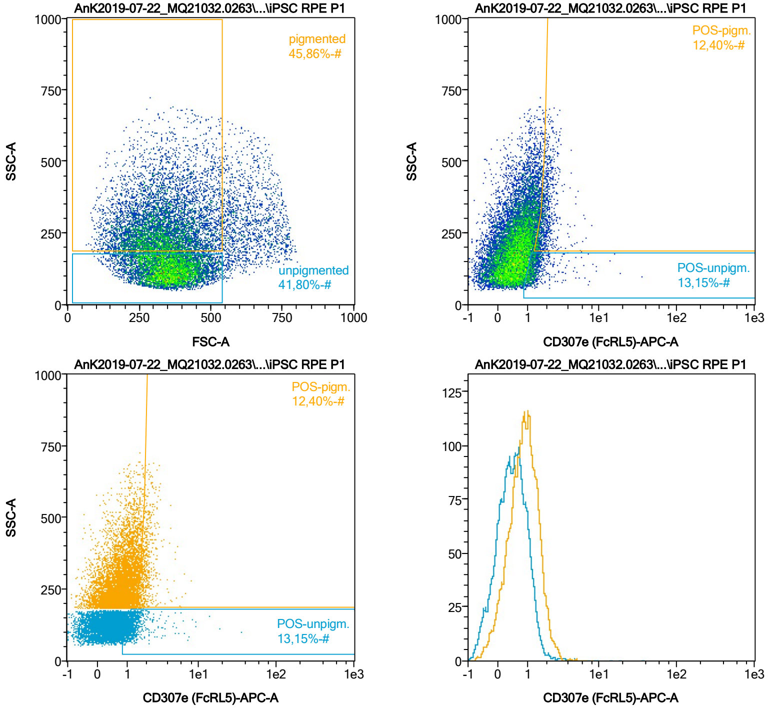


| Name | %-# | CD307e (FcRL5)-APC-A Mean | CD307e (FcRL5)-APC-A Median |
|---|---|---|---|
| POS-unpigm. | 13,15 | 1,50 | 1,24 |
| POS-pigm. | 12,40 | 1,97 | 1,83 |

Date:2024-06-14
Operator: Sebastian Knoebel
Version:2.13.0+54.g95abf0f4

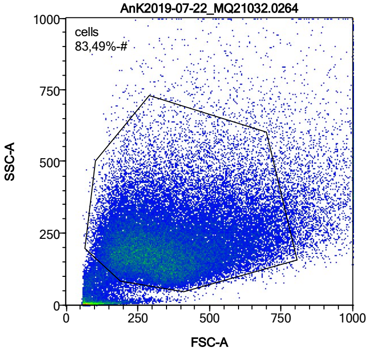


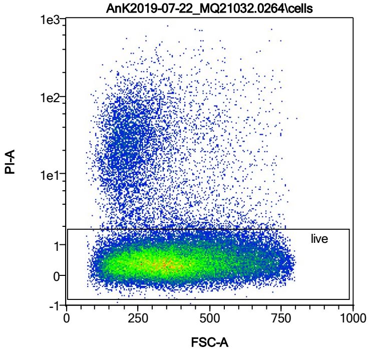


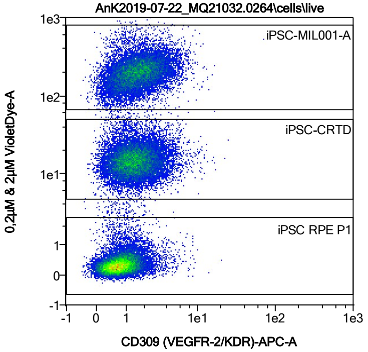


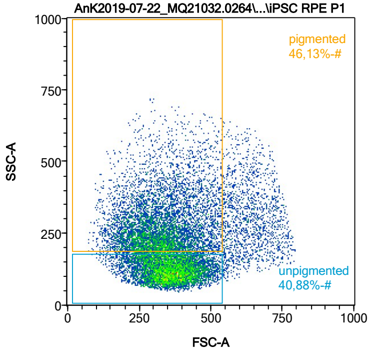


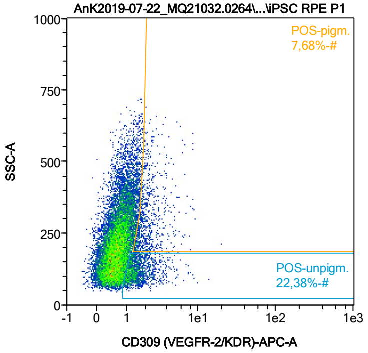


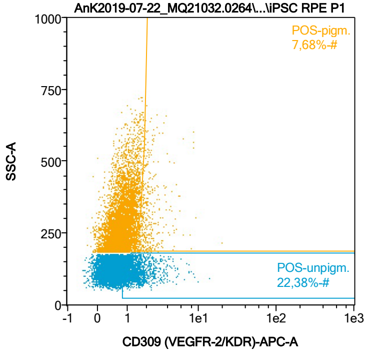


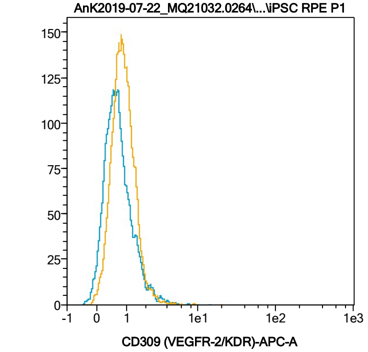


| Name | %-# | CD309 (VEGFR-2/KDR)-APC-A Mean | CD309 (VEGFR-2/KDR)-APC-A Median |
|---|---|---|---|
| POS-unpigm. | 22,38 | 1,65 | 1,39 |
| POS-pigm. | 7,68 | 2,24 | 1,90 |

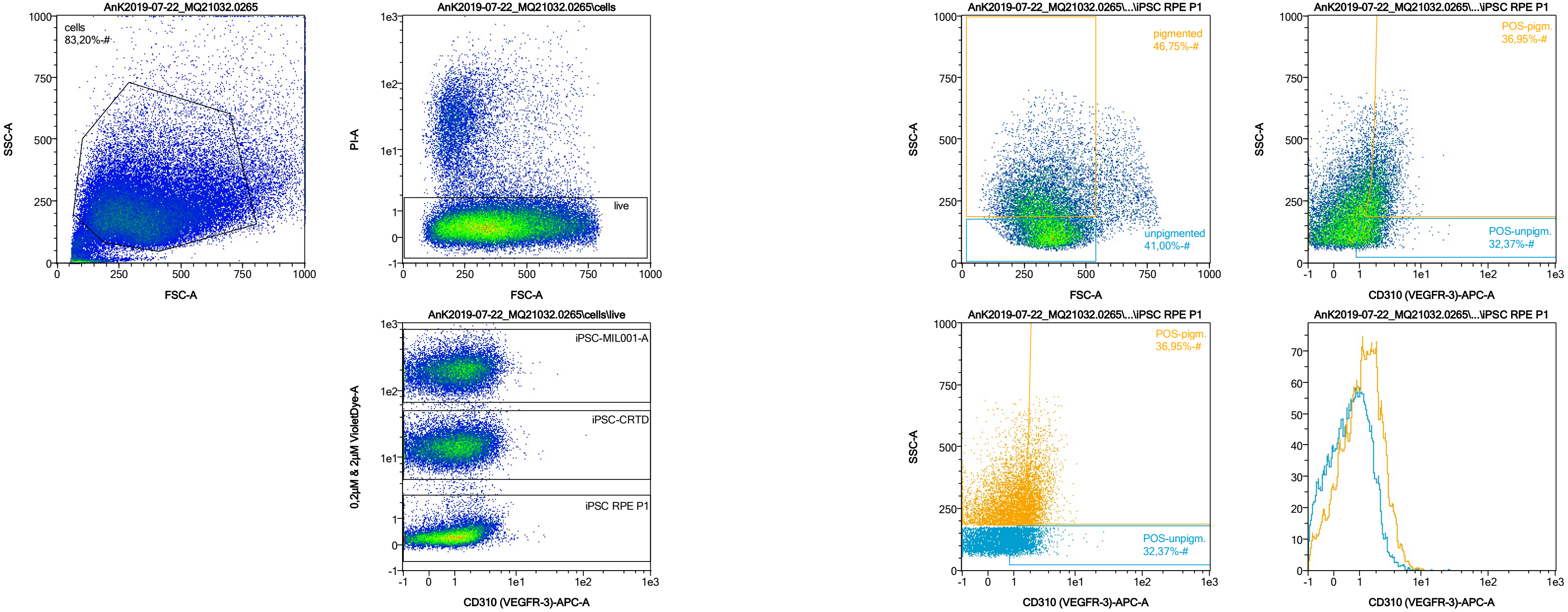


| Name | %-# | CD310 (VEGFR-3)-APC-A Mean | CD310 (VEGFR-3)-APC-A Median |
|---|---|---|---|
| POS-unpigm. | 32,37 | 1,81 | 1,55 |
| POS-pigm. | 36,95 | 2,51 | 2,21 |

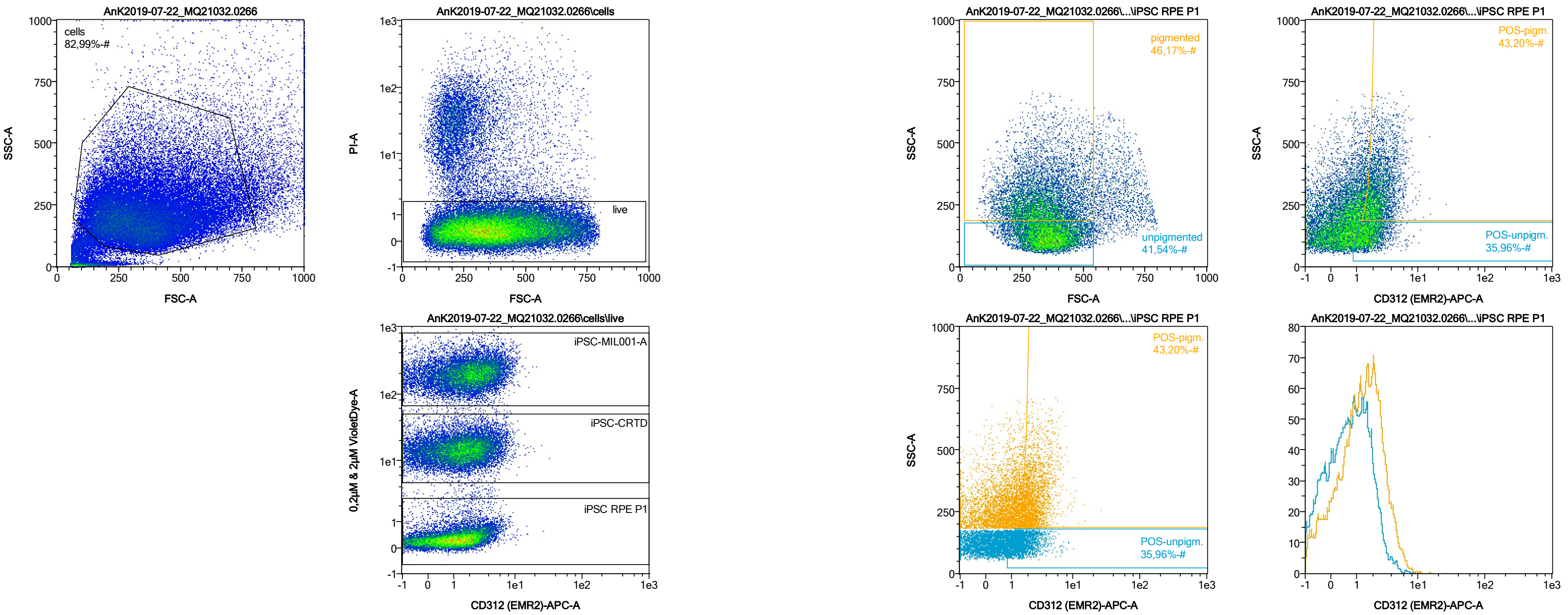


| Name | %-# | CD312 (EMR2)-APC-A Mean | CD312 (EMR2)-APC-A Median |
|---|---|---|---|
| POS-unpigm. | 35,96 | 1,85 | 1,62 |
| POS-pigm. | 43,20 | 2,68 | 2,35 |

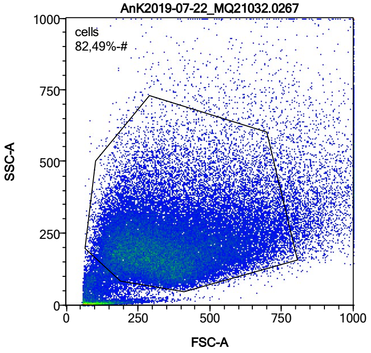


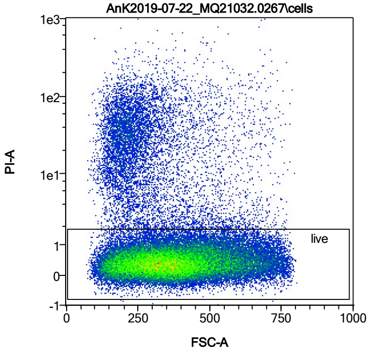


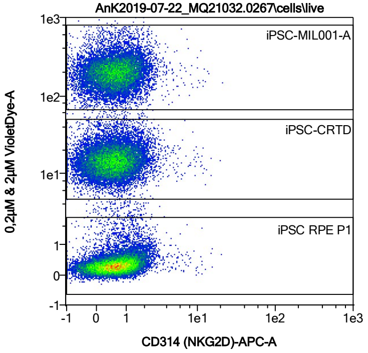


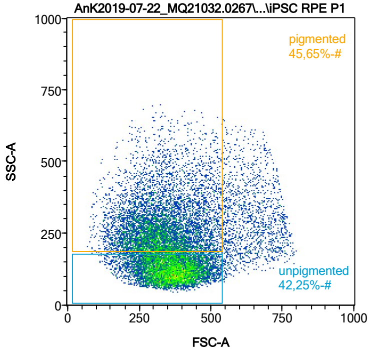


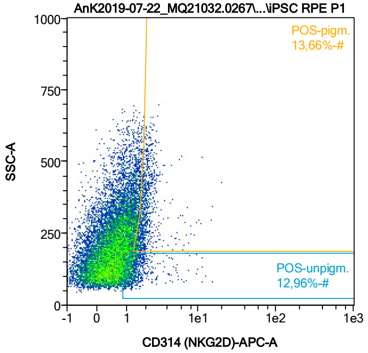


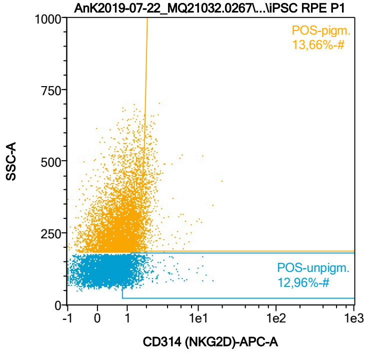


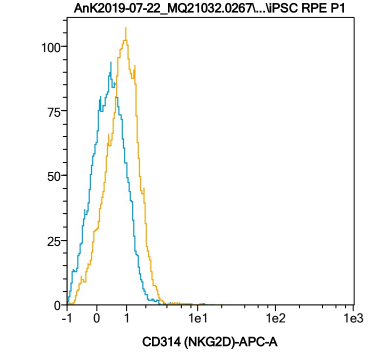


| Name | %-# | CD314 (NKG2D)-APC-A Mean | CD314 (NKG2D)-APC-A Median |
|---|---|---|---|
| POS-unpigm. | 12,96 | 1,54 | 1,27 |
| POS-pigm. | 13,66 | 2,10 | 1,87 |

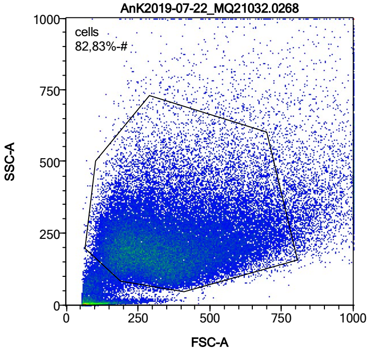


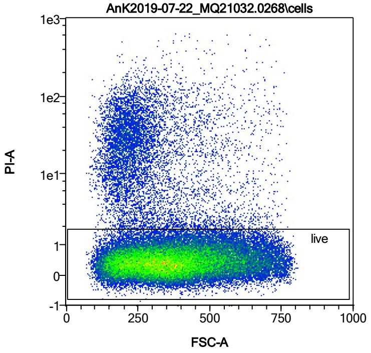


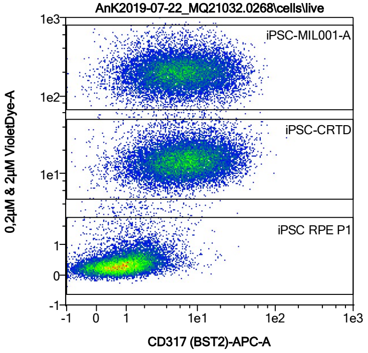


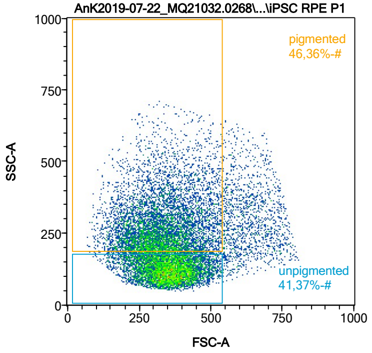


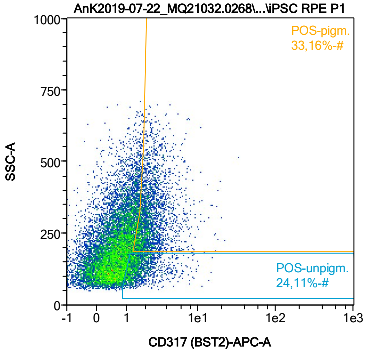


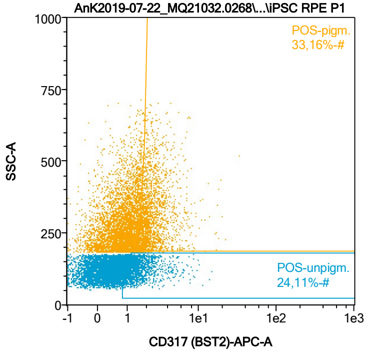


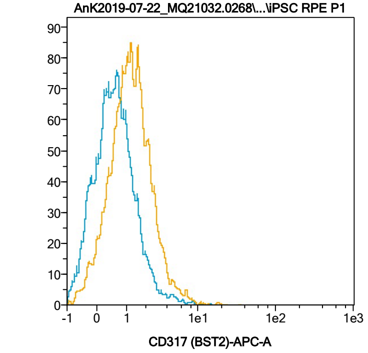


| Name | %-# | CD317 (BST2)-APC-A Mean | CD317 (BST2)-APC-A Median |
|---|---|---|---|
| POS-unpigm. | 24,11 | 1,84 | 1,47 |
| POS-pigm. | 33,16 | 2,67 | 2,19 |

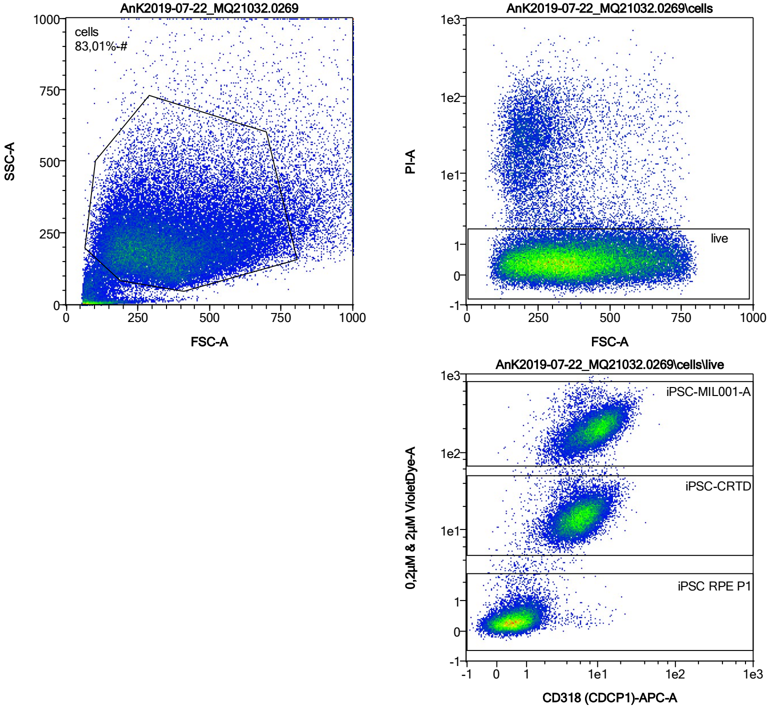


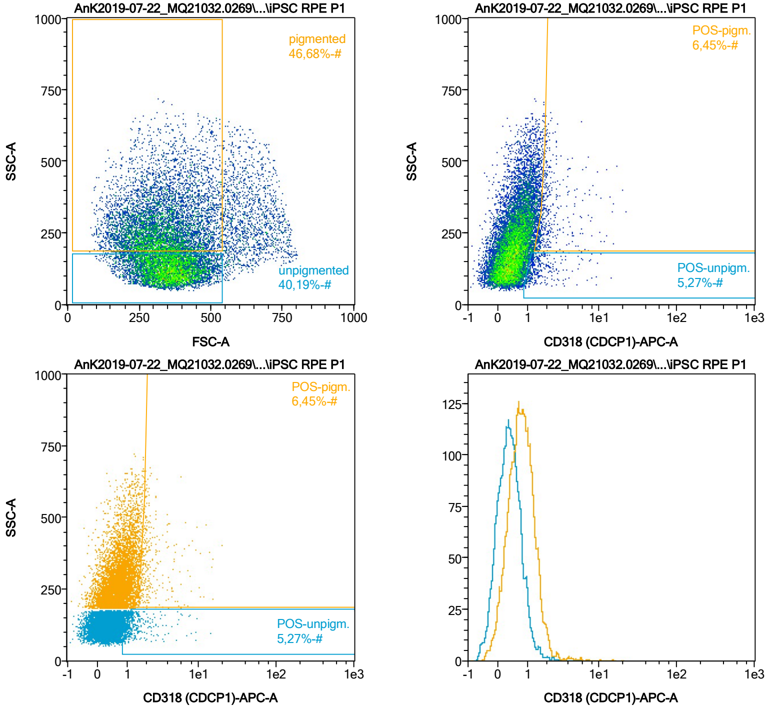


| Name | %-# | CD318 (CDCP1)-APC-A Mean | CD318 (CDCP1)-APC-A Median |
|---|---|---|---|
| POS-unpigm. | 5,27 | 1,74 | 1,26 |
| POS-pigm. | 6,45 | 2,55 | 1,93 |

Date:2024-06-14

Operator: Sebastian Knoebel

Version:2.13.0+54.g95abf0f4

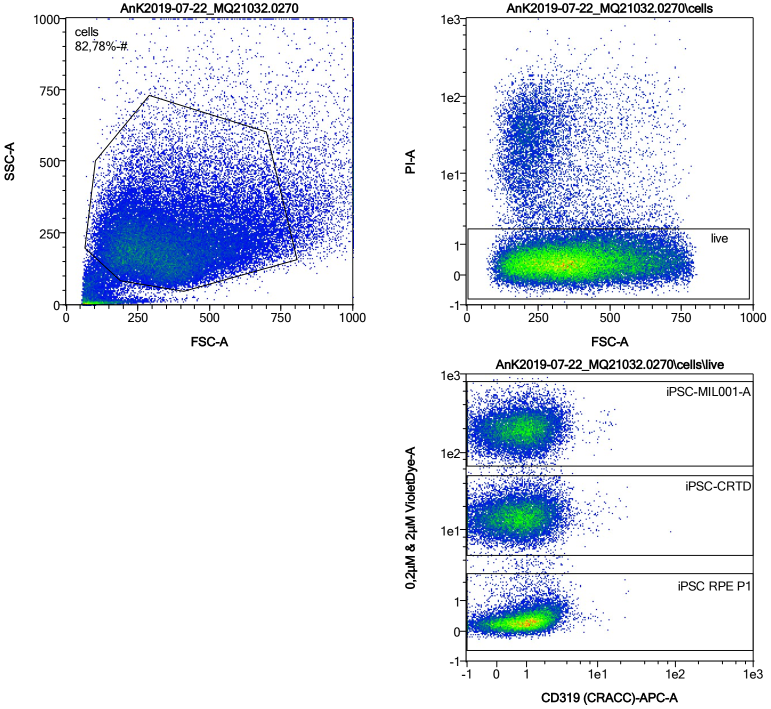


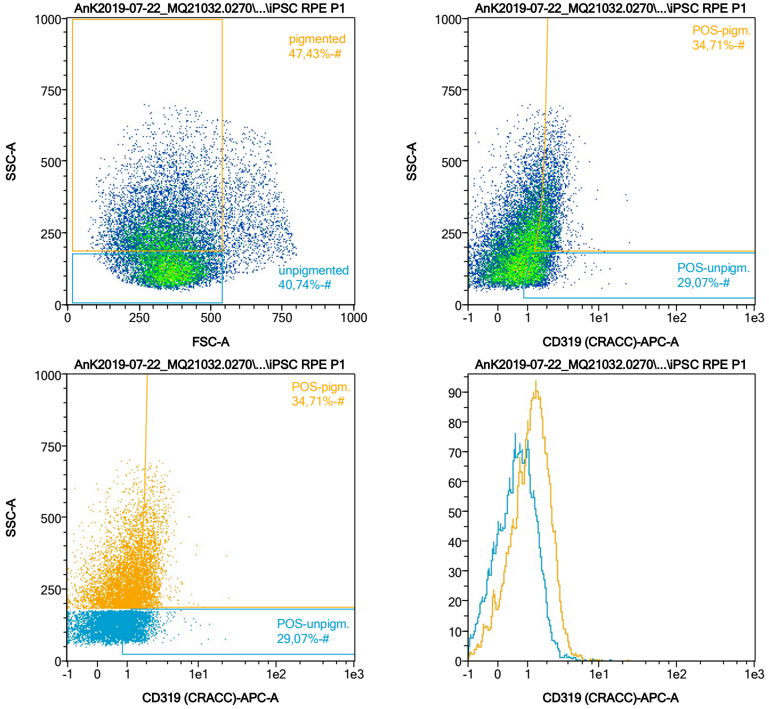


| Name | %-# | CD319 (CRACC)-APC-A Mean | CD319 (CRACC)-APC-A Median |
|---|---|---|---|
| POS-unpigm. | 29,07 | 1,60 | 1,41 |
| POS-pigm. | 34,71 | 2,25 | 2,06 |

Date:2024-06-14

Operator: Sebastian Knoebel

Version:2.13.0+54.g95abf0f4

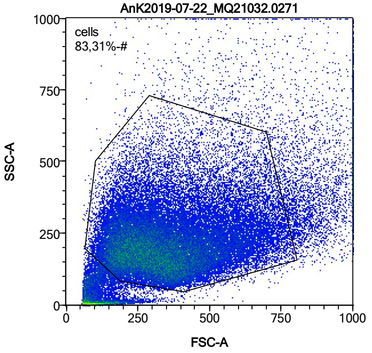


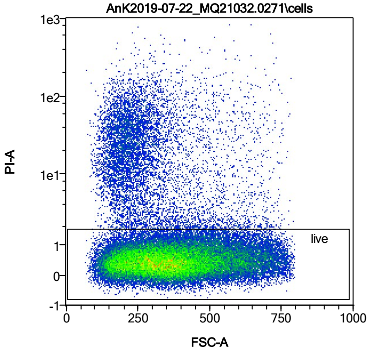


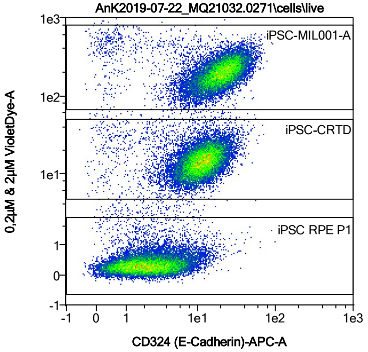


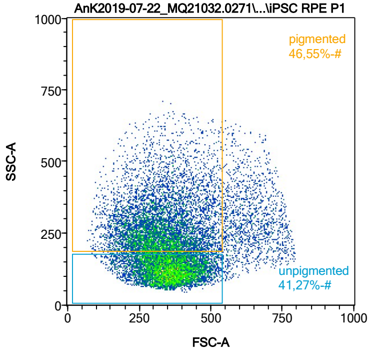


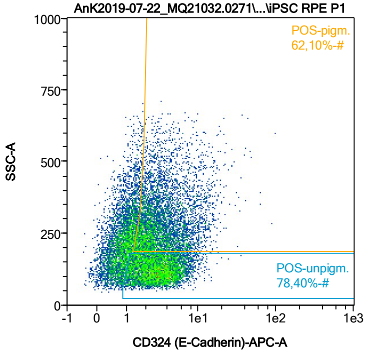


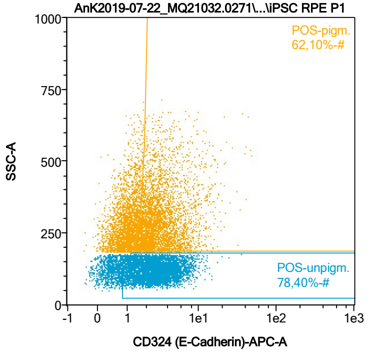


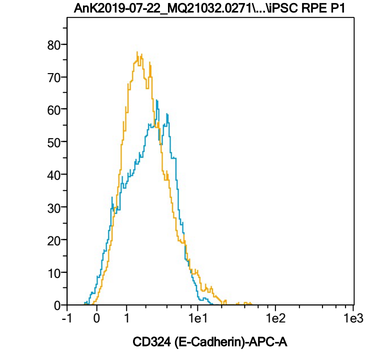


| Name | %-# | CD324 (E-Cadherin)-APC-A Mean | CD324 (E-Cadherin)-APC-A Median |
|---|---|---|---|
| POS-unpigm. | 78,40 | 3,27 | 2,83 |
| POS-pigm. | 62,10 | 3,76 | 2,74 |

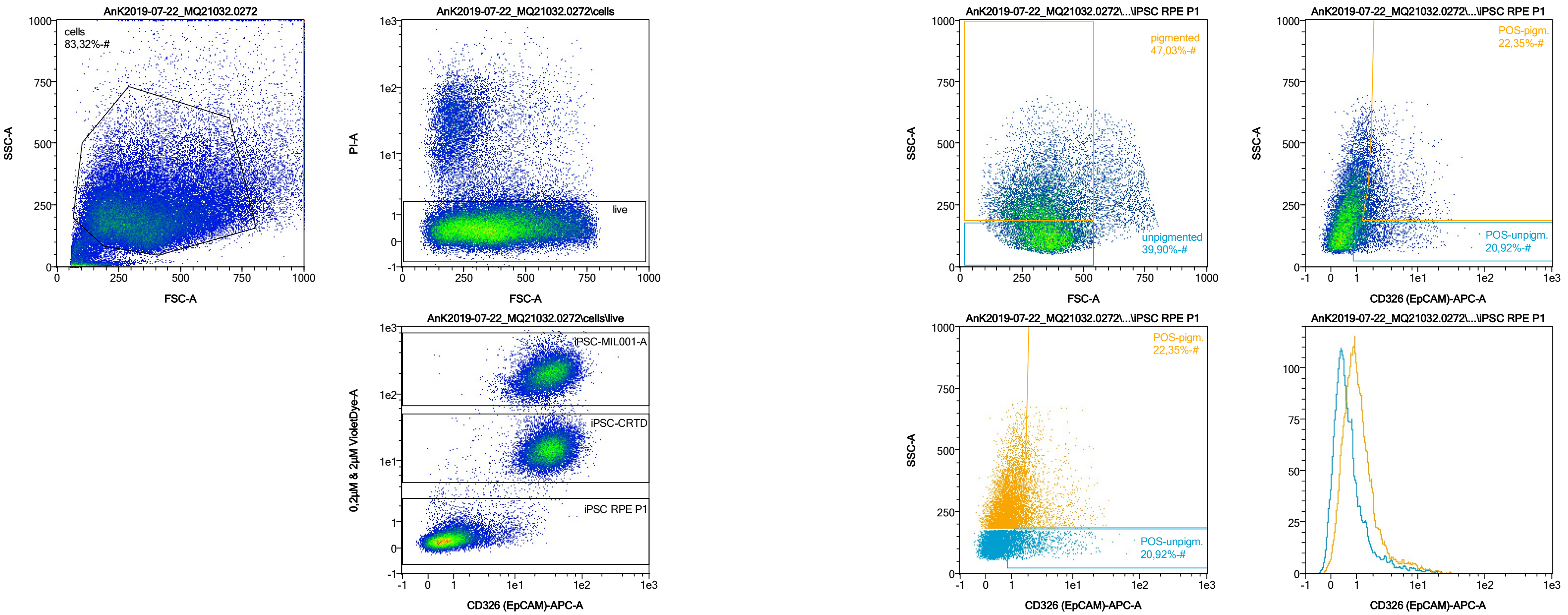


| Name | %-# | CD326 (EpCAM)-APC-A Mean | CD326 (EpCAM)-APC-A Median |
|---|---|---|---|
| POS-unpigm. | 20,92 | 3,10 | 1,66 |
| POS-pigm. | 22,35 | 3,63 | 2,28 |

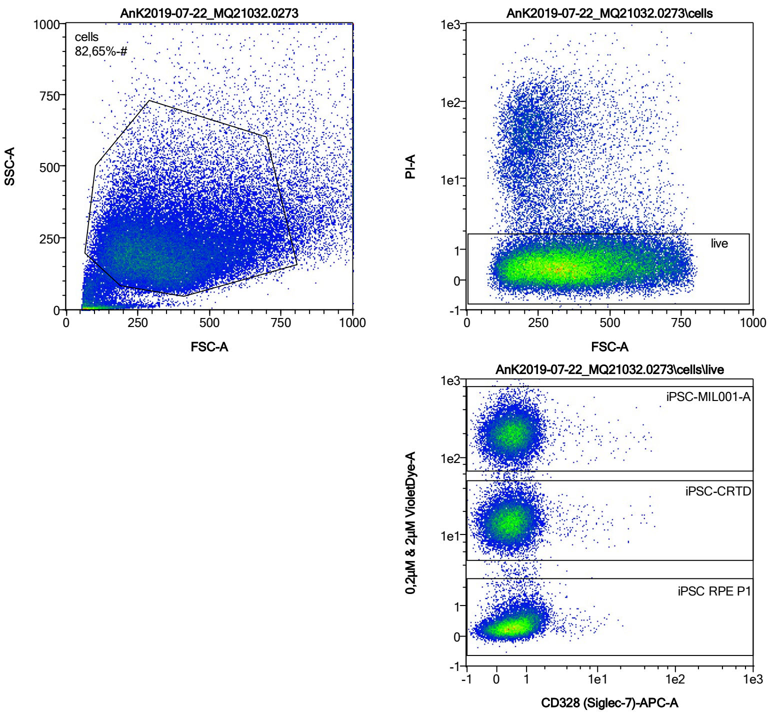


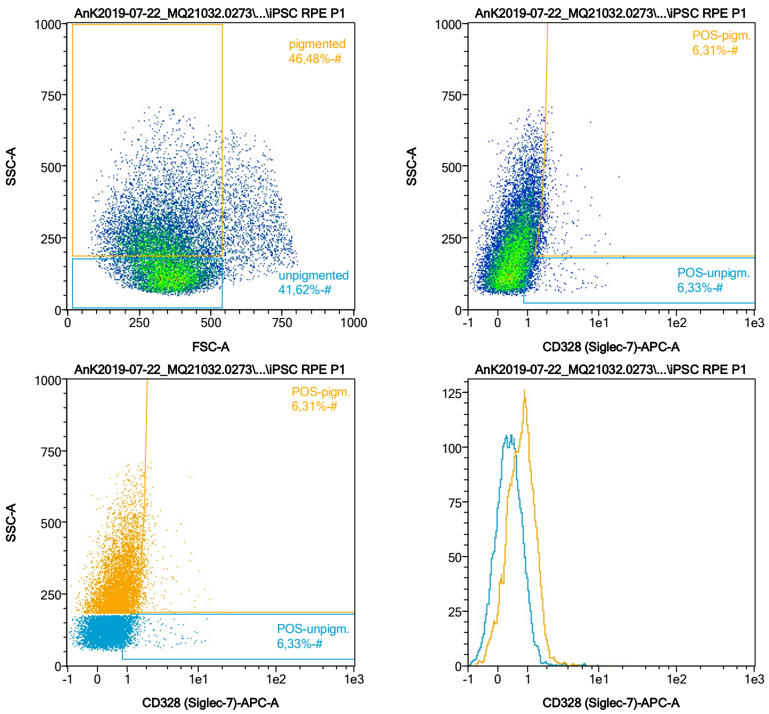


| Name | %-# | CD328 (Siglec-7)-APC-A Mean | CD328 (Siglec-7)-APC-A Median |
|---|---|---|---|
| POS-unpigm. | 6,33 | 1,72 | 1,21 |
| POS-pigm. | 6,31 | 2,24 | 1,84 |

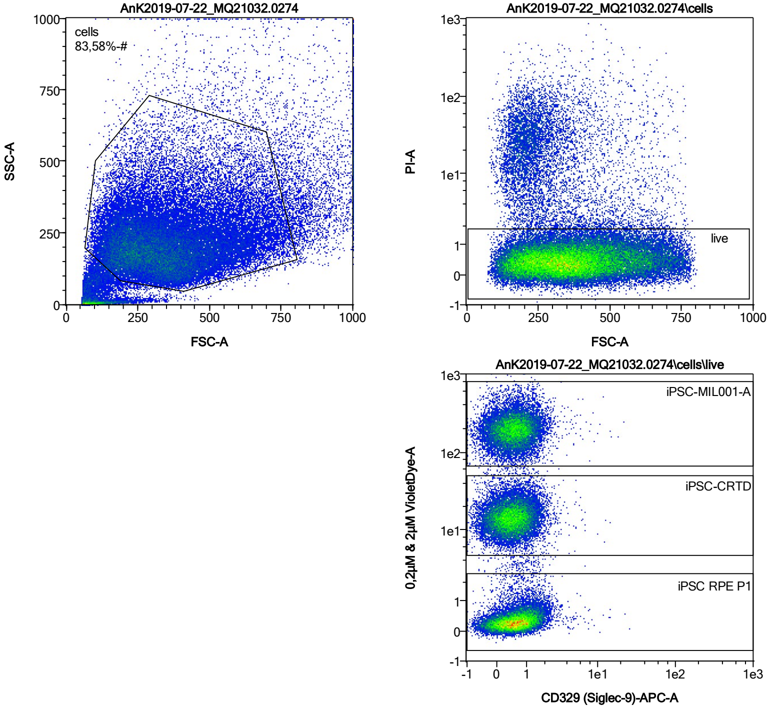


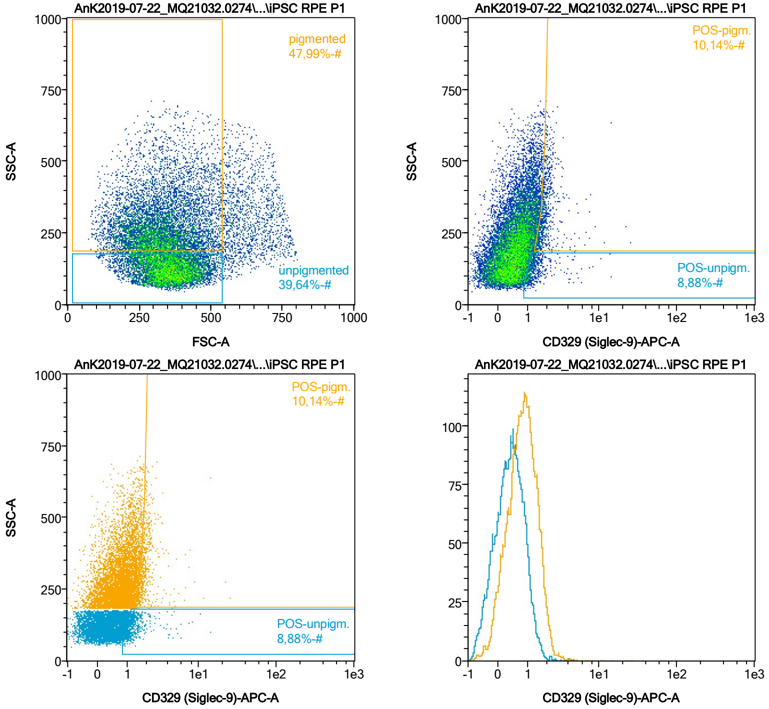


| Name | %-# | CD329 (Siglec-9)-APC-A Mean | CD329 (Siglec-9)-APC-A Median |
|---|---|---|---|
| POS-unpigm. | 8,88 | 1,46 | 1,20 |
| POS-pigm. | 10,14 | 2,06 | 1,82 |

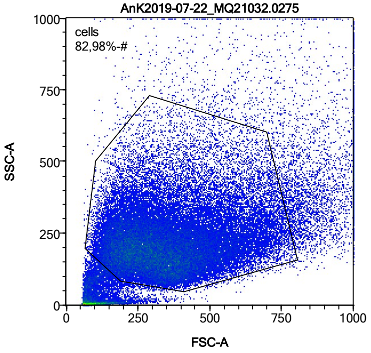


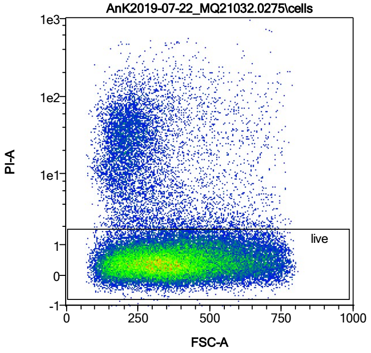


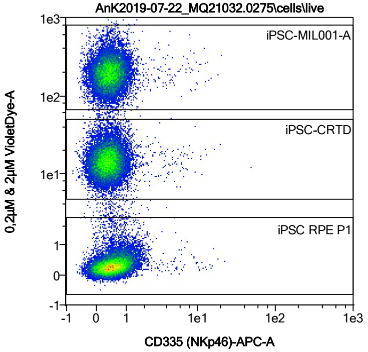


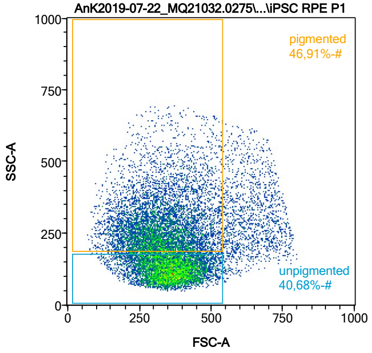


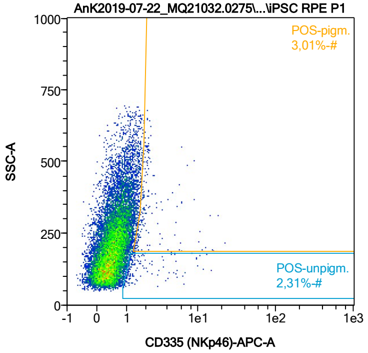


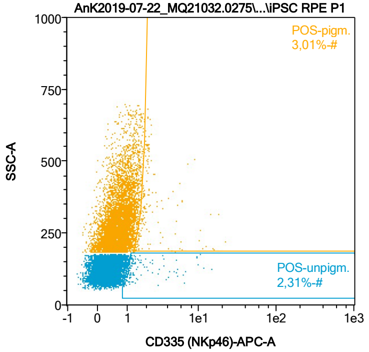


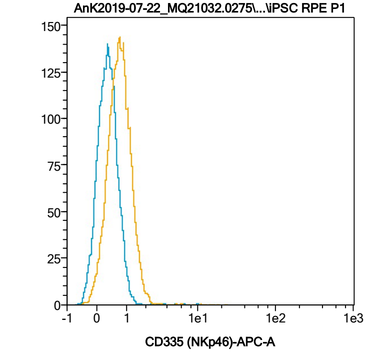


| Name | %-# | CD335 (NKp46)-APC-A Mean | CD335 (NKp46)-APC-A Median |
|---|---|---|---|
| POS-unpigm. | 2,31 | 2,15 | 1,22 |
| POS-pigm. | 3,01 | 2,75 | 1,87 |

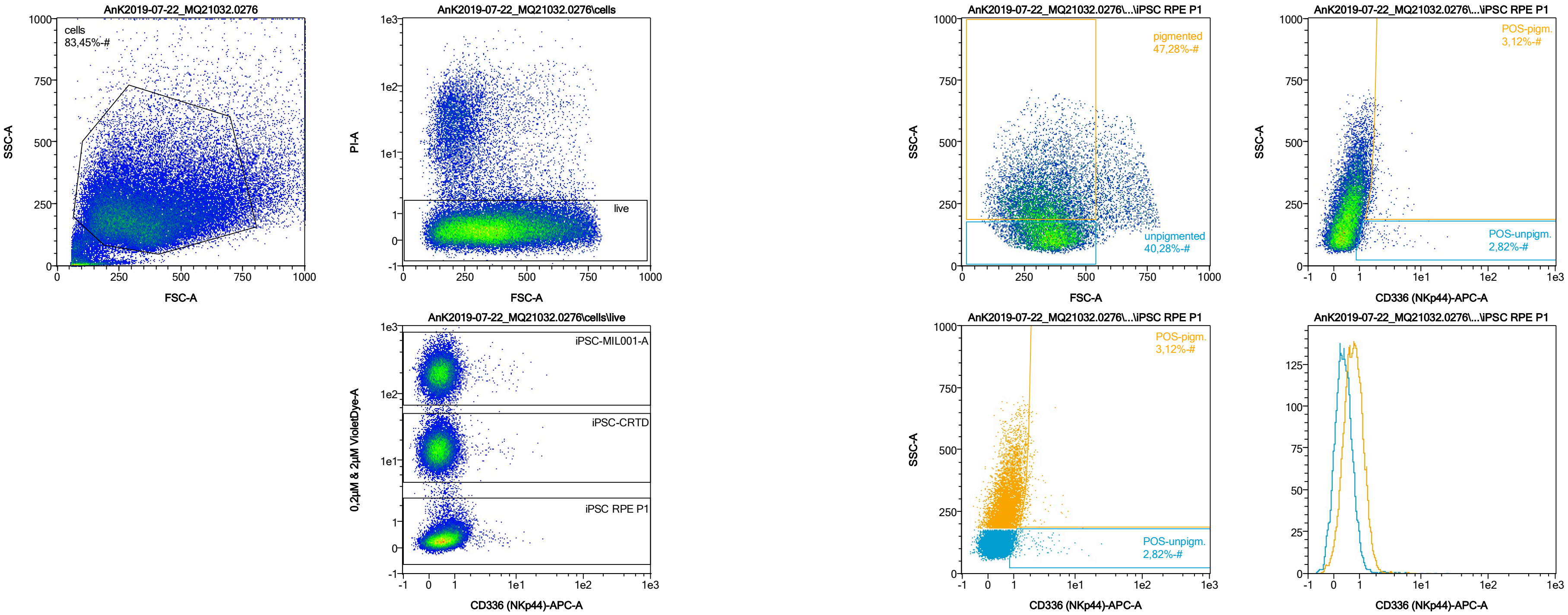


| Name | %-# | CD336 (NKp44)-APC-A Mean | CD336 (NKp44)-APC-A Median |
|---|---|---|---|
| POS-unpigm. | 2,82 | 2,21 | 1,26 |
| POS-pigm. | 3,12 | 2,13 | 1,85 |

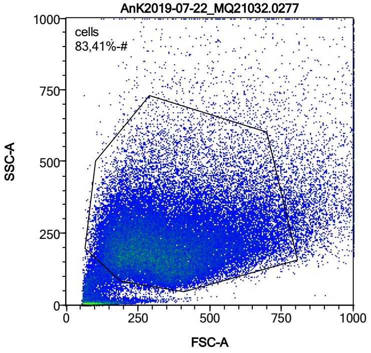


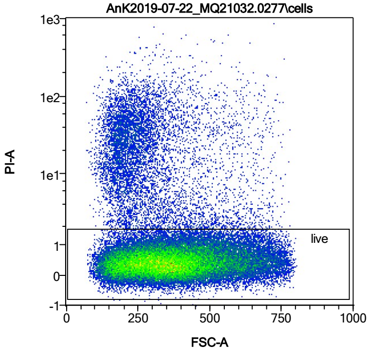


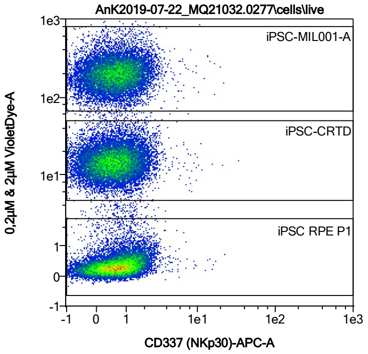


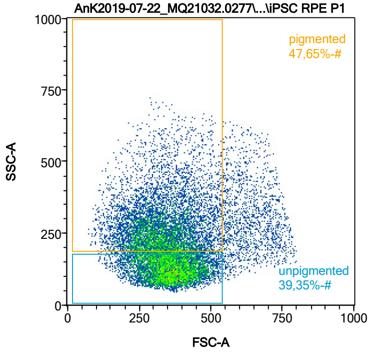


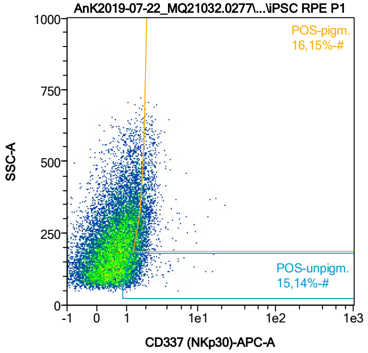


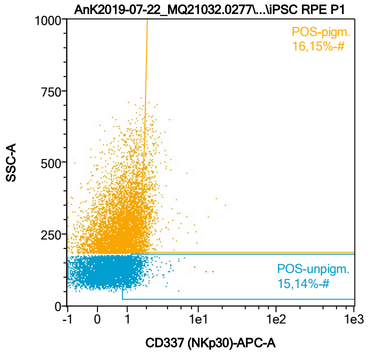


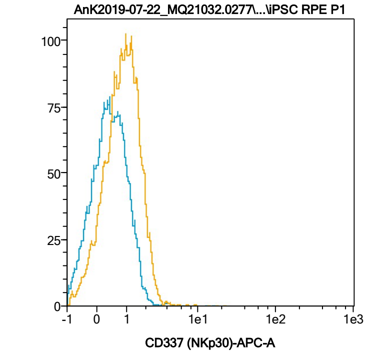


| Name | %-# | CD337 (NKp30)-APC-A Mean | CD337 (NKp30)-APC-A Median |
|---|---|---|---|
| POS-unpigm. | 15,14 | 1,45 | 1,29 |
| POS-pigm. | 16,15 | 2,07 | 1,88 |

Date:2024-06-14

Operator: Sebastian Knoebel

Version:2.13.0+54.g95abf0f4

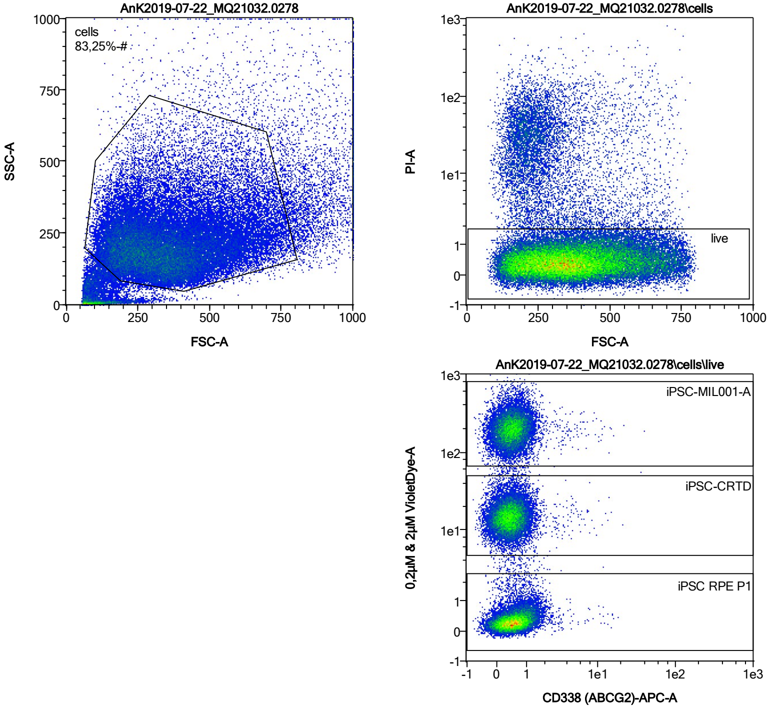


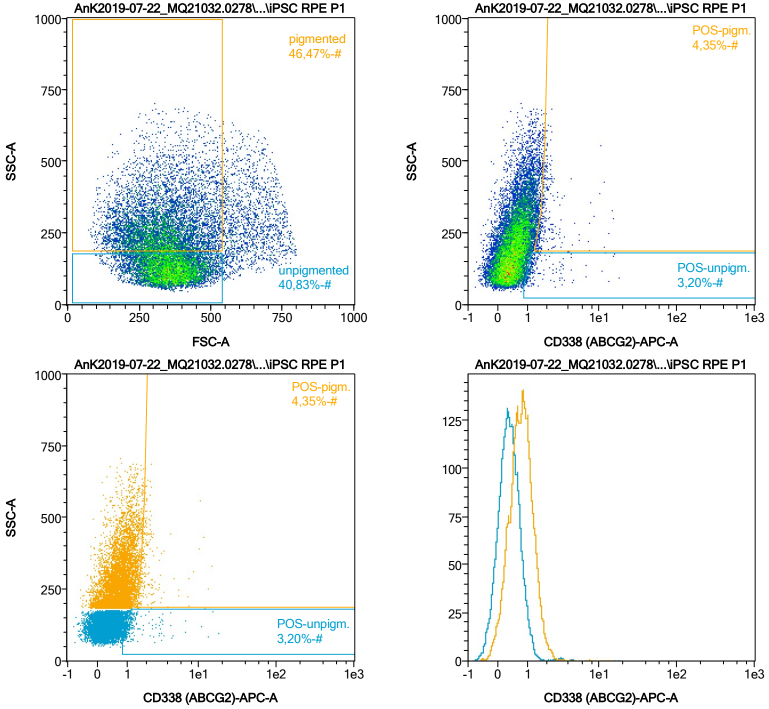


| Name | %-# | CD338 (ABCG2)-APC-A Mean | CD338 (ABCG2)-APC-A Median |
|---|---|---|---|
| POS-unpigm. | 3,20 | 1,80 | 1,19 |
| POS-pigm. | 4,35 | 2,25 | 1,84 |

Date:2024-06-14

Operator: Sebastian Knoebel

Version:2.13.0+54.g95abf0f4

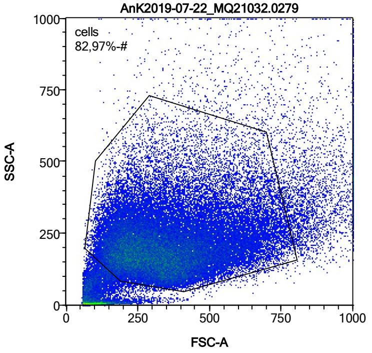


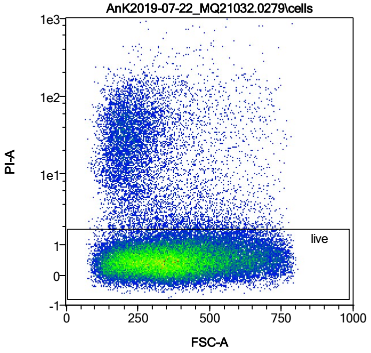


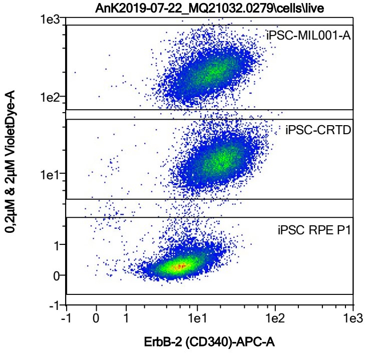


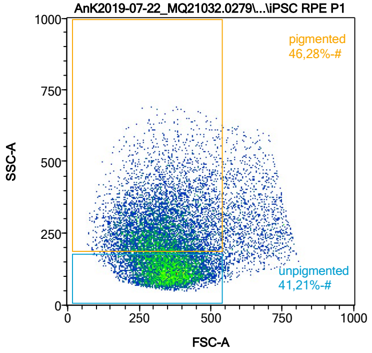


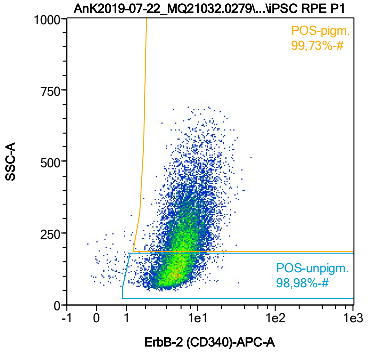


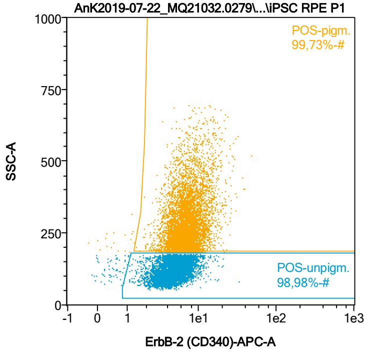


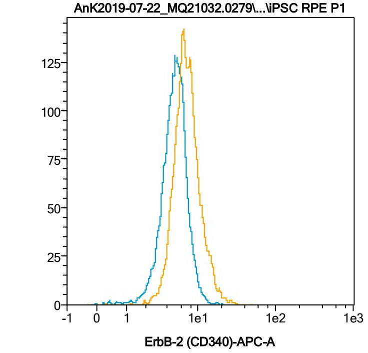


| Name | %-# | ErbB-2 (CD340)-APC-A Mean | ErbB-2 (CD340)-APC-A Median |
|---|---|---|---|
| POS-unpigm. | 98,98 | 5,18 | 4,97 |
| POS-pigm. | 99,73 | 7,62 | 6,91 |

Date:2024-06-14

Operator: Sebastian Knoebel

Version:2.13.0+54.g95abf0f4

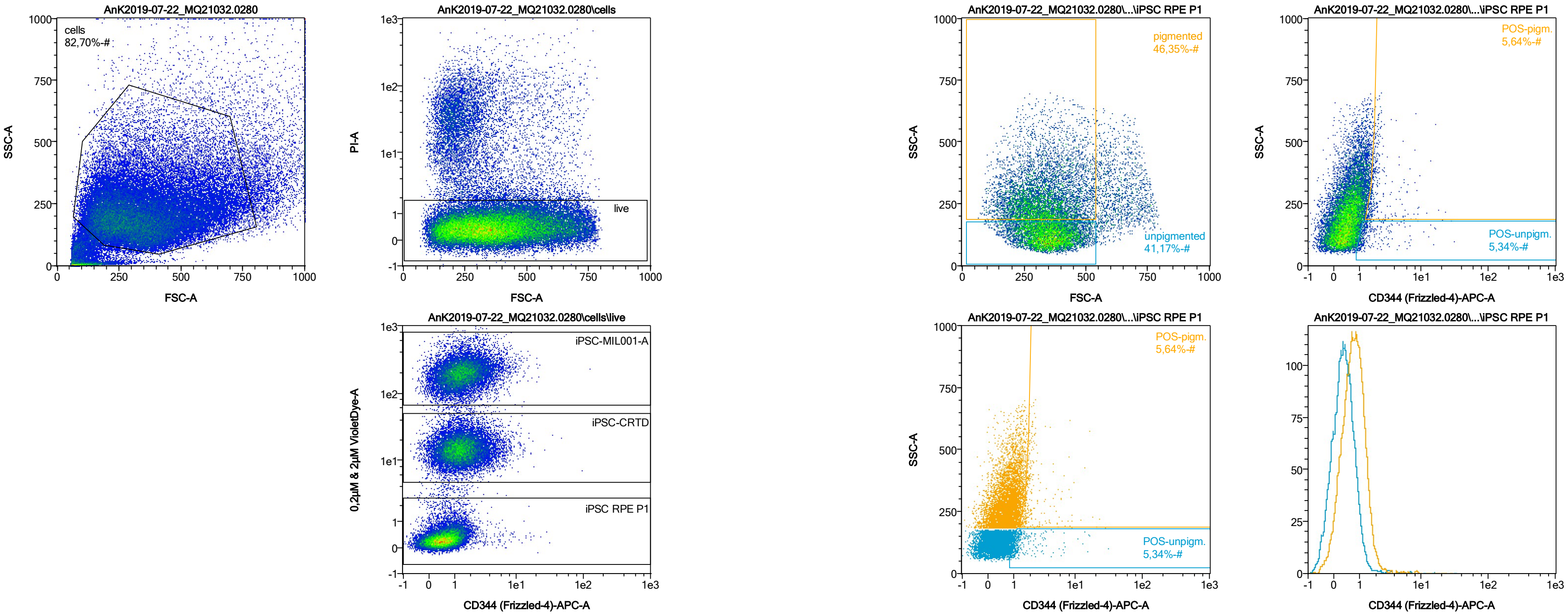


| Name | %-# | CD344 (Frizzled-4)-APC-A Mean | CD344 (Frizzled-4)-APC-A Median |
|---|---|---|---|
| POS-unpigm. | 5,34 | 1,91 | 1,22 |
| POS-pigm. | 5,64 | 2,61 | 1,85 |

Date:2024-06-14

Operator: Sebastian Knoebel

Version:2.13.0+54.g95abf0f4

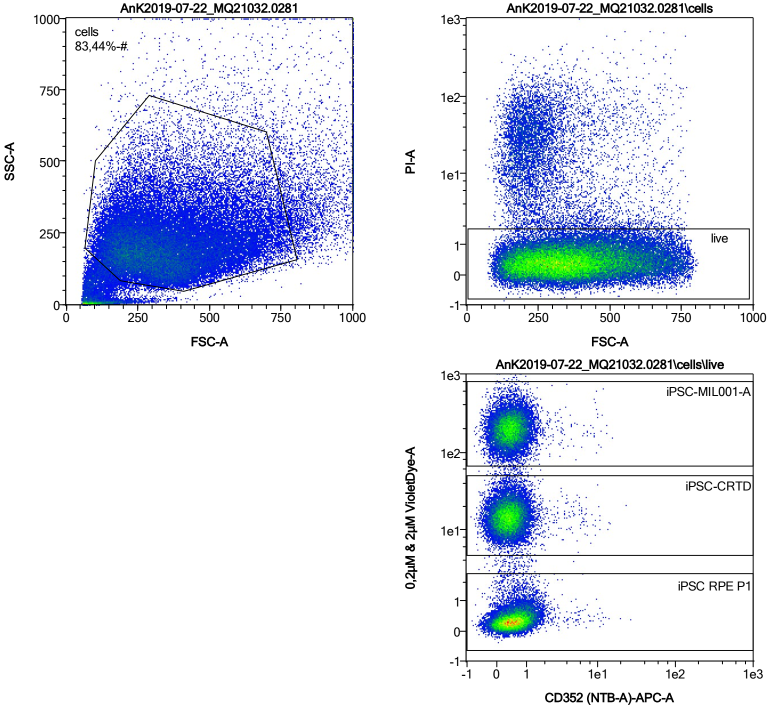


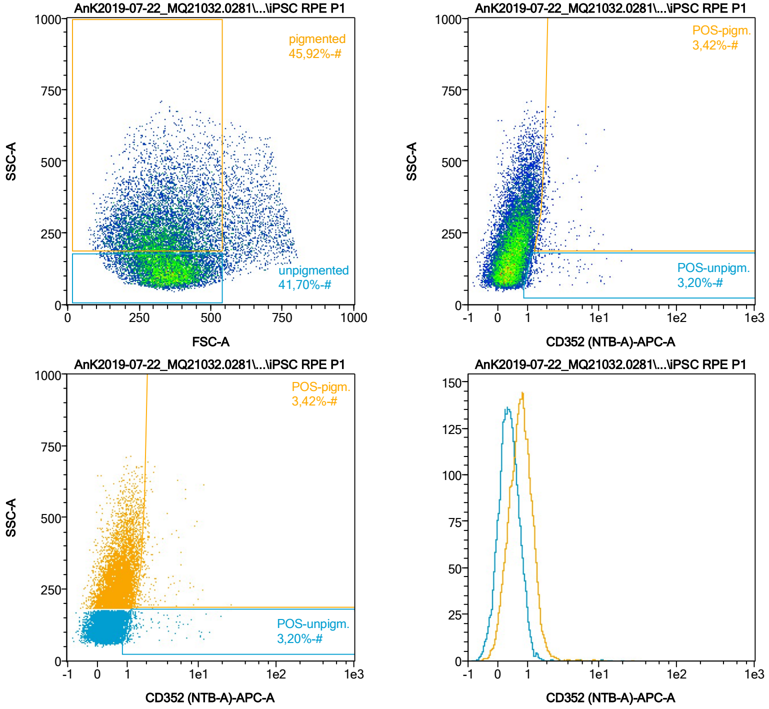


| Name | %-# | CD352 (NTB-A)-APC-A Mean | CD352 (NTB-A)-APC-A Median |
|---|---|---|---|
| POS-unpigm. | 3,20 | 2,08 | 1,18 |
| POS-pigm. | 3,42 | 2,58 | 1,91 |

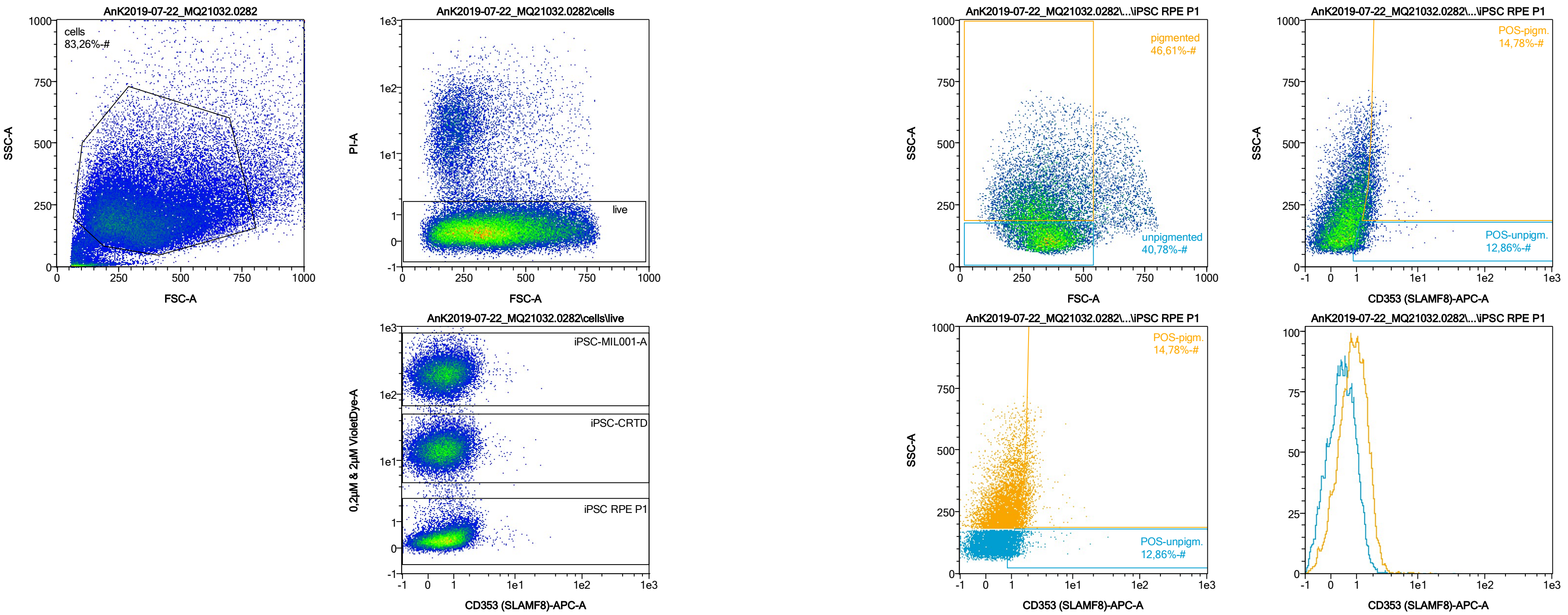


| Name | %-# | CD353 (SLAMF8)-APC-A Mean | CD353 (SLAMF8)-APC-A Median |
|---|---|---|---|
| POS-unpigm. | 12,86 | 1,47 | 1,25 |
| POS-pigm. | 14,78 | 2,05 | 1,87 |

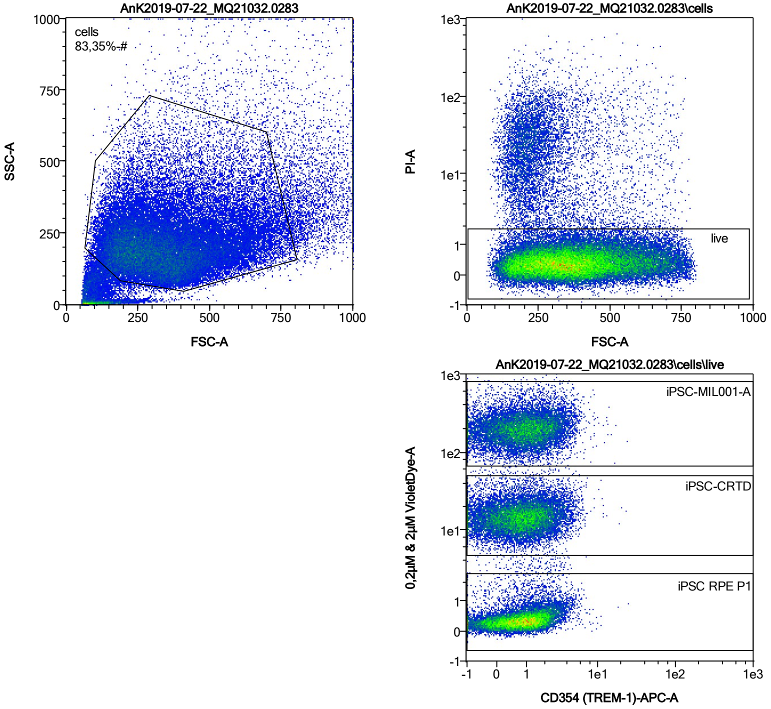


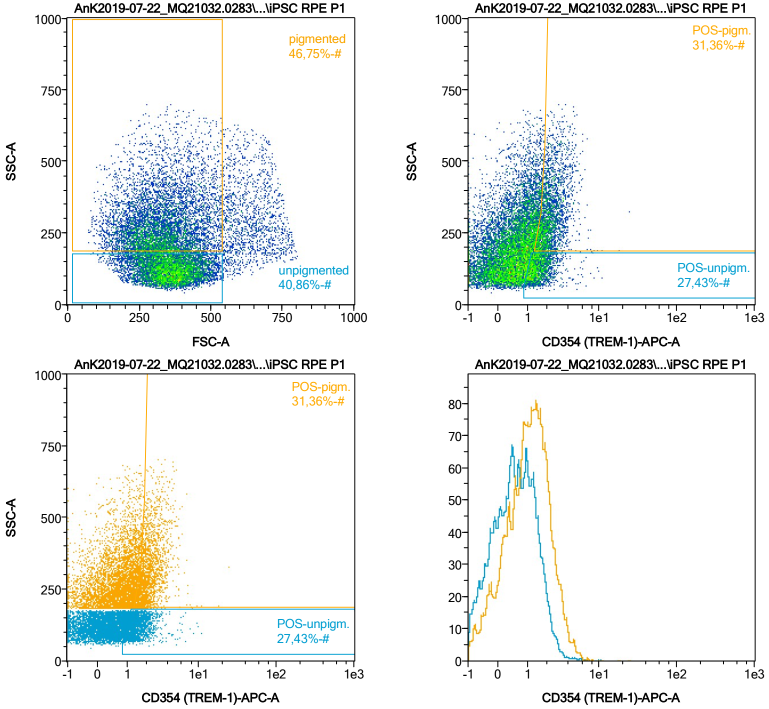


| Name | %-# | CD354 (TREM-1)-APC-A Mean | CD354 (TREM-1)-APC-A Median |
|---|---|---|---|
| POS-unpigm. | 27,43 | 1,62 | 1,43 |
| POS-pigm. | 31,36 | 2,35 | 2,09 |

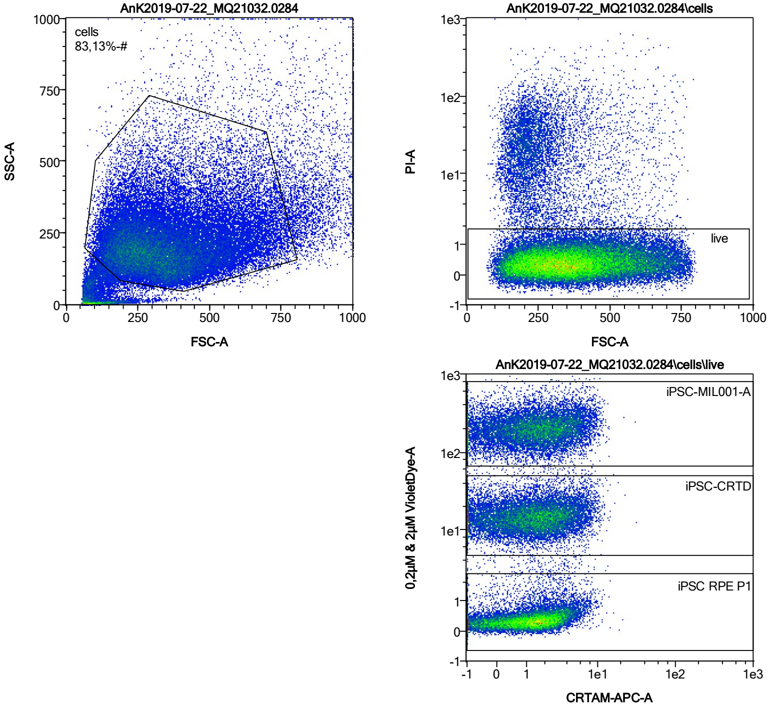


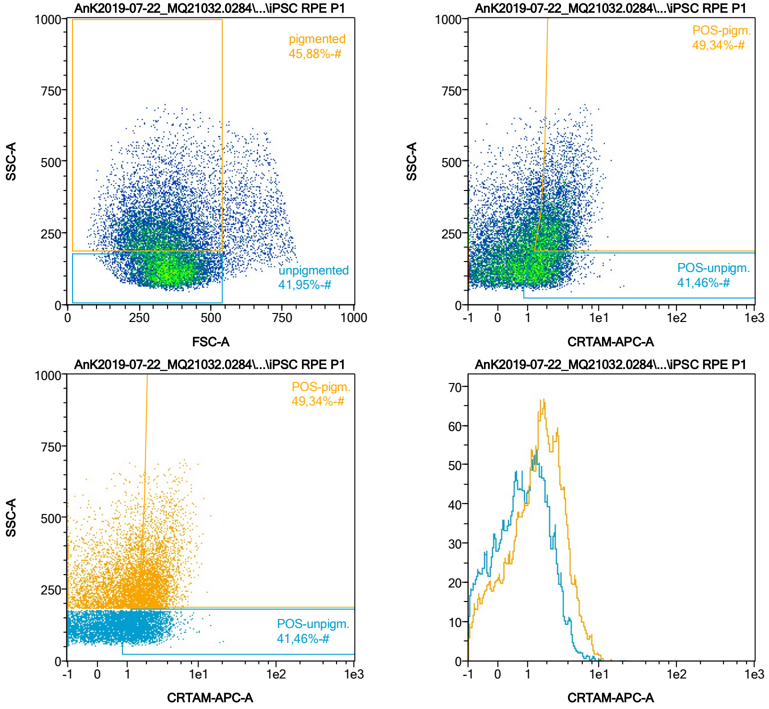


| Name | %-# | CRTAM-APC-A Mean | CRTAM-APC-A Median |
|---|---|---|---|
| POS-unpigm. | 41,46 | 2,08 | 1,78 |
| POS-pigm. | 49,34 | 2,97 | 2,58 |

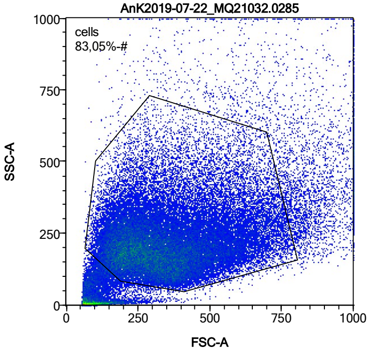


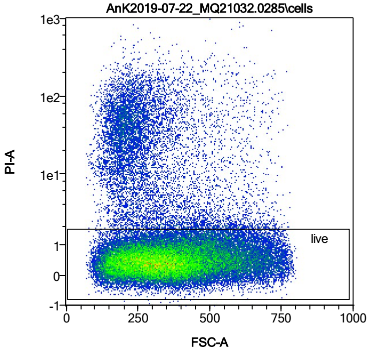


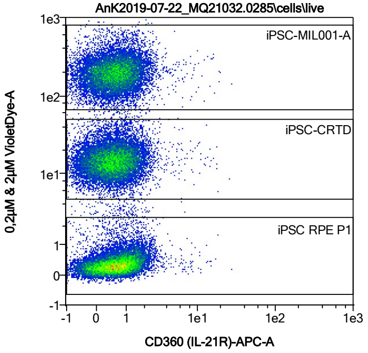


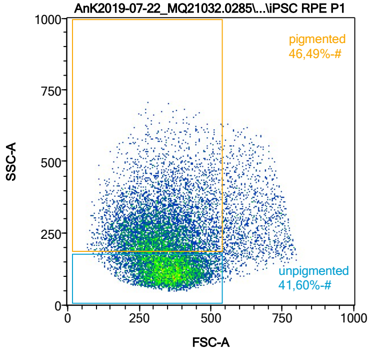


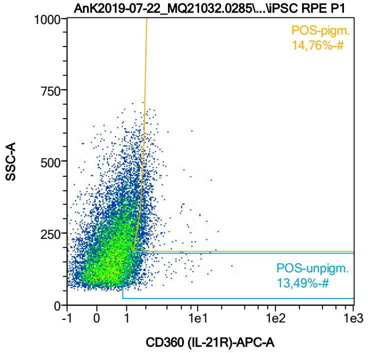


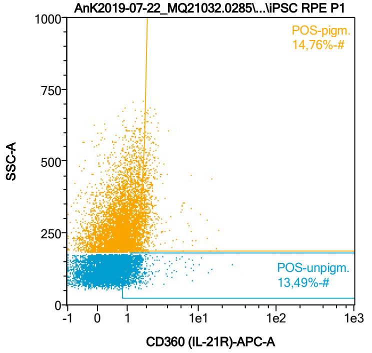


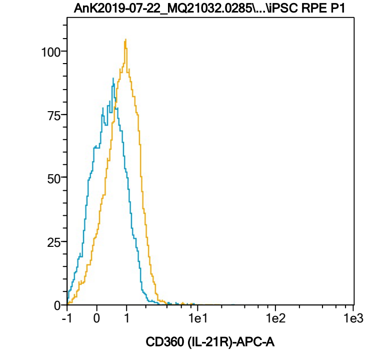


| Name | %-# | CD360 (IL-21R)-APC-A Mean | CD360 (IL-21R)-APC-A Median |
|---|---|---|---|
| POS-unpigm. | 13,49 | 1,59 | 1,29 |
| POS-pigm. | 14,76 | 2,11 | 1,86 |

Date:2024-06-14

Operator: Sebastian Knoebel

Version:2.13.0+54.g95abf0f4

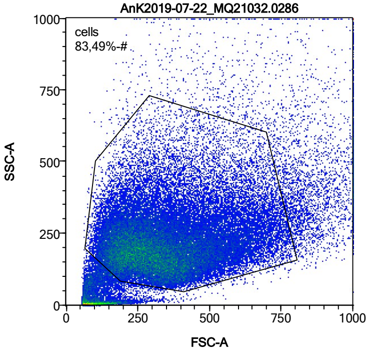


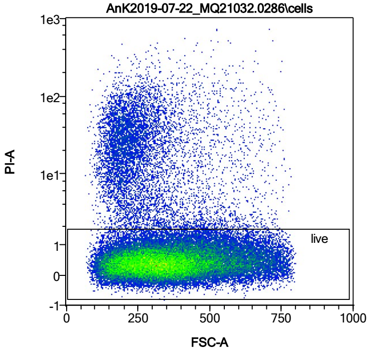


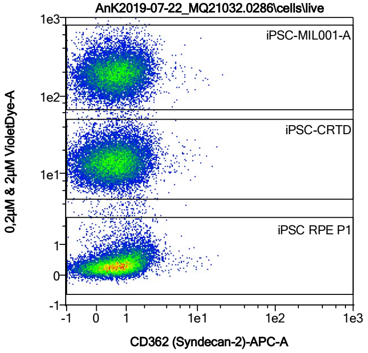


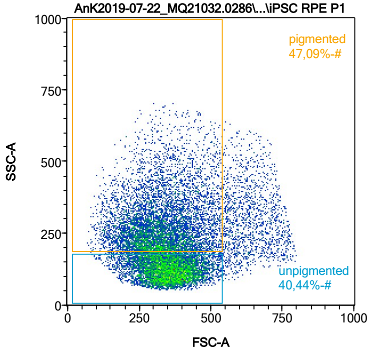


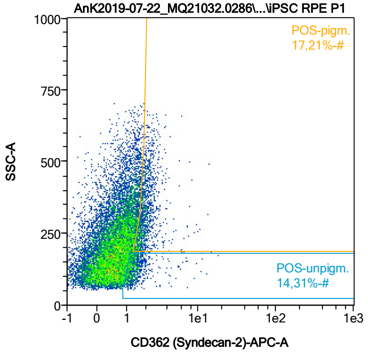


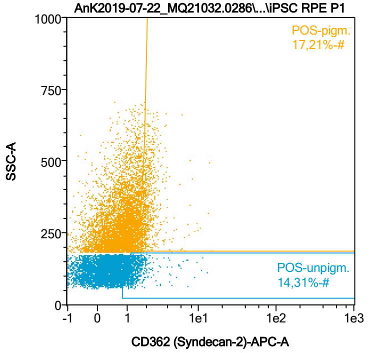


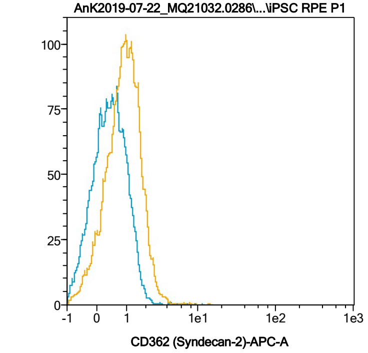


| Name | %-# | CD362 (Syndecan-2)-APC-A Mean | CD362 (Syndecan-2)-APC-A Median |
|---|---|---|---|
| POS-unpigm. | 14,31 | 1,50 | 1,27 |
| POS-pigm. | 17,21 | 2,12 | 1,91 |

Date:2024-06-14

Operator: Sebastian Knoebel

Version:2.13.0+54.g95abf0f4

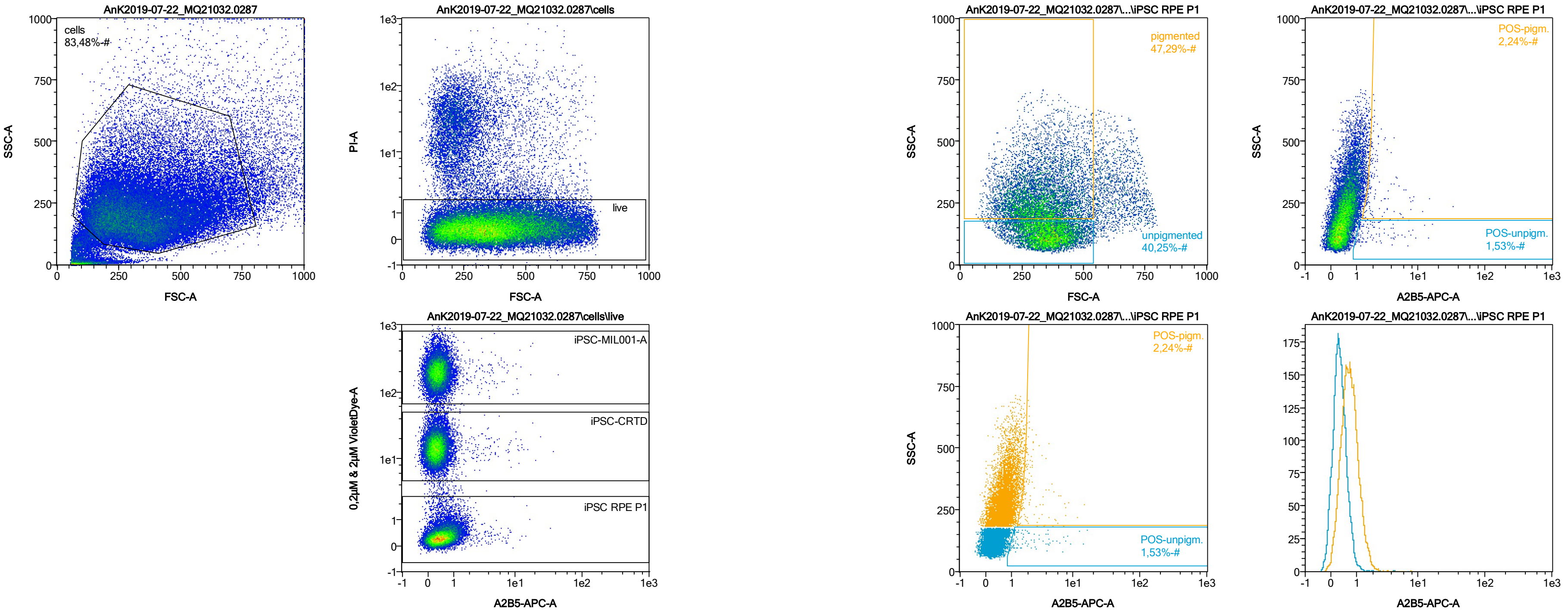


| Name | %-# | A2B5-APC-A Mean | A2B5-APC-A Median |
|---|---|---|---|
| POS-unpigm. | 1,53 | 2,77 | 1,60 |
| POS-pigm. | 2,24 | 2,59 | 1,92 |

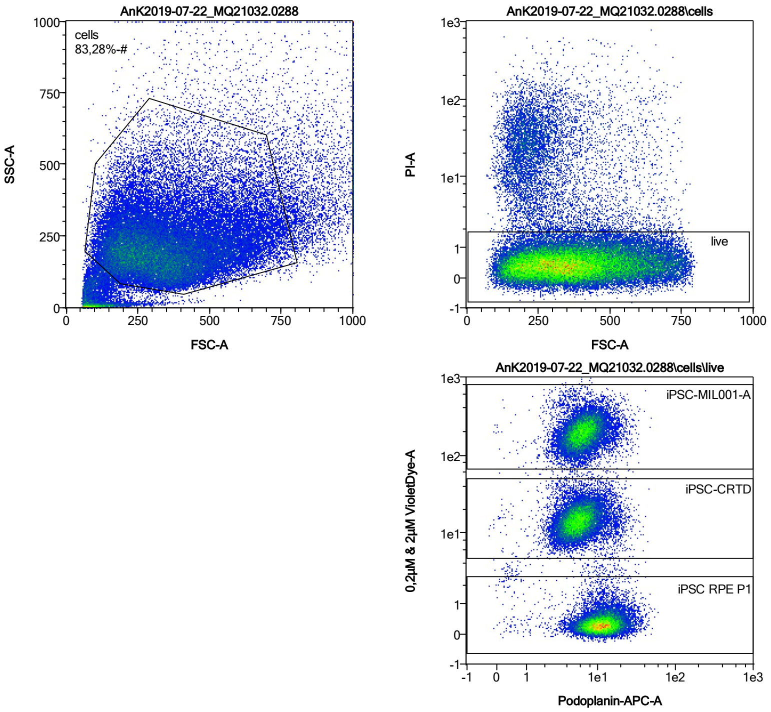


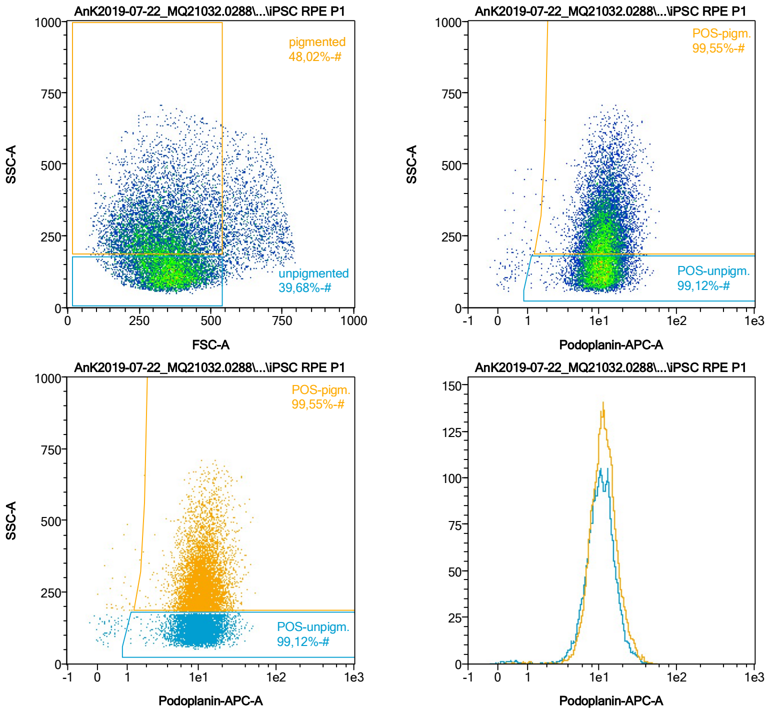


| Name | %-# | Podoplanin-APC-A Mean | Podoplanin-APC-A Median |
|---|---|---|---|
| POS-unpigm. | 99,12 | 11,21 | 10,47 |
| POS-pigm. | 99,55 | 12,20 | 11,24 |

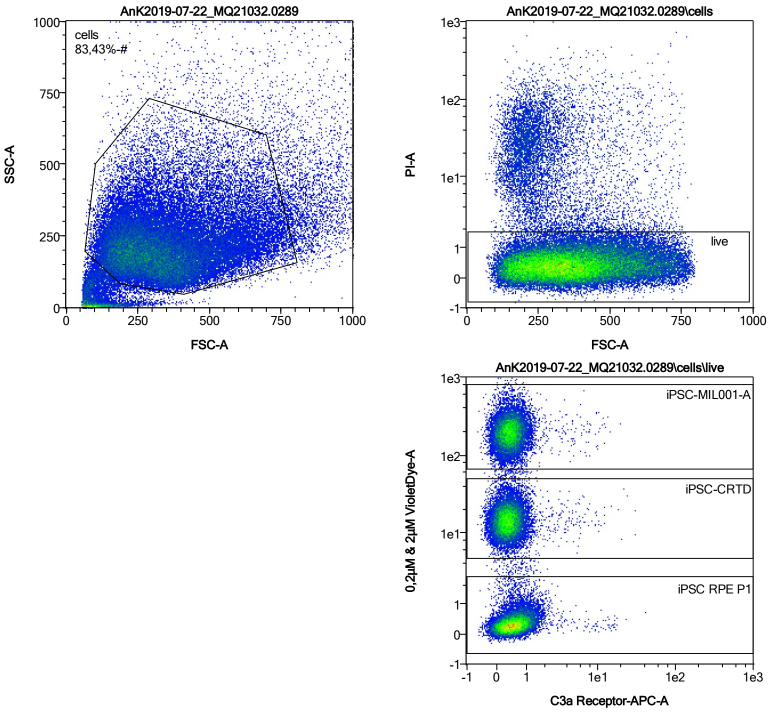


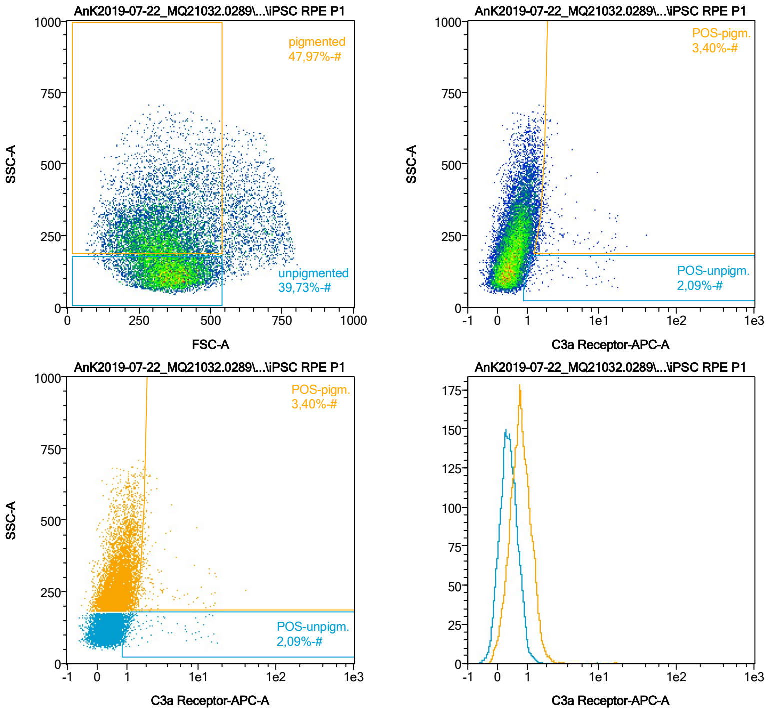


| Name | %-# | C3a Receptor-APC-A Mean | C3a Receptor-APC-A Median |
|---|---|---|---|
| POS-unpigm. | 2,09 | 3,01 | 1,29 |
| POS-pigm. | 3,40 | 3,07 | 1,89 |

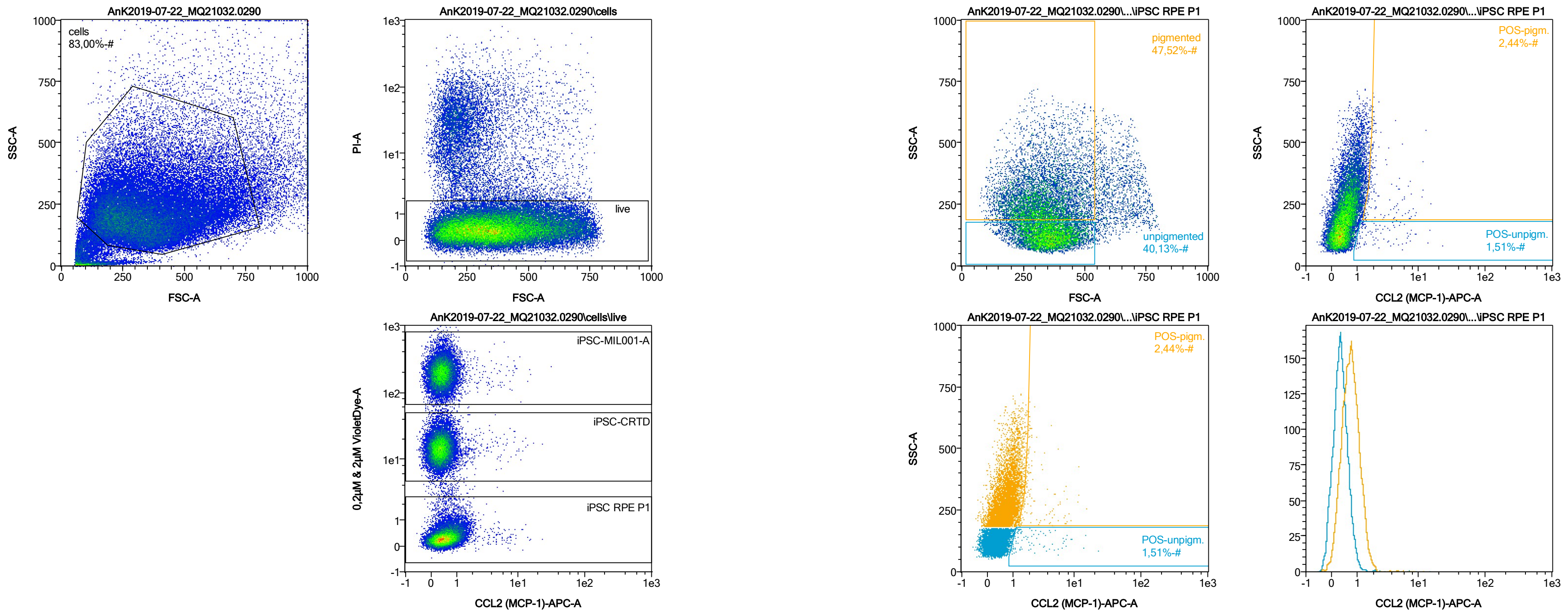


| Name | %-# | CCL2 (MCP-1)-APC-A Mean | CCL2 (MCP-1)-APC-A Median |
|---|---|---|---|
| POS-unpigm. | 1,51 | 3,17 | 1,81 |
| POS-pigm. | 2,44 | 2,87 | 1,95 |

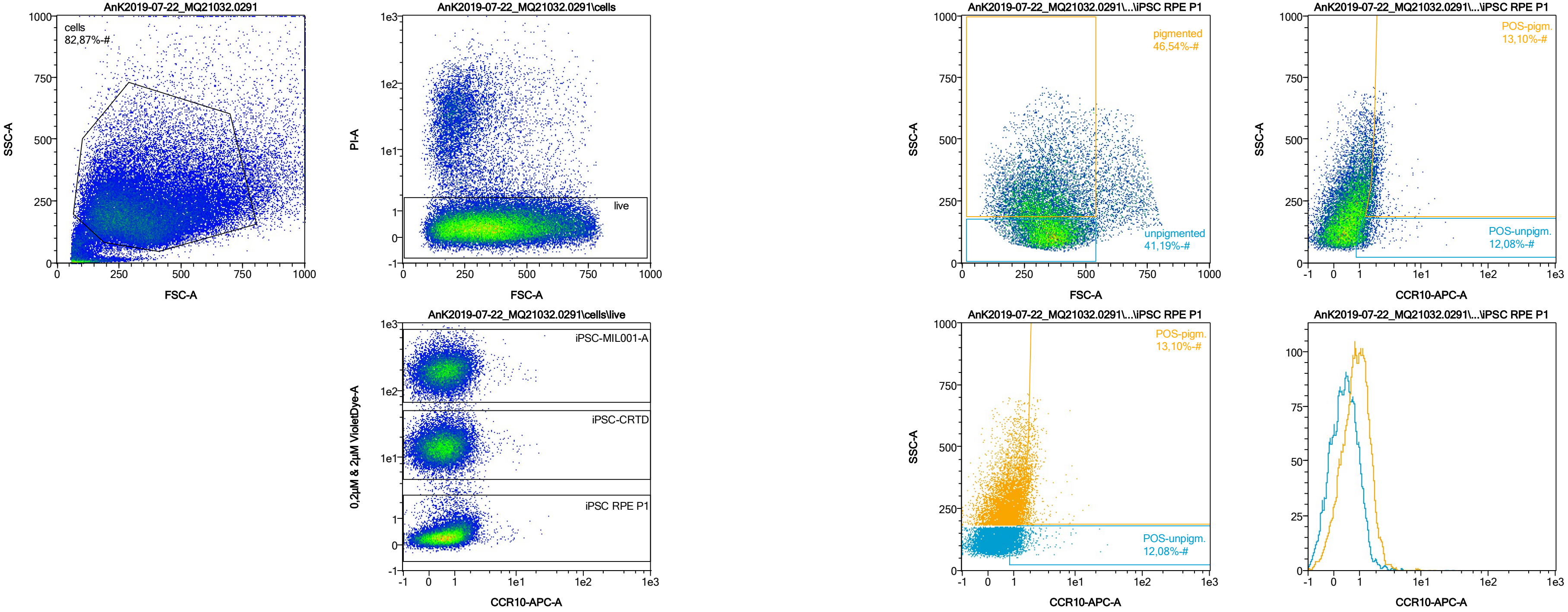


| Name | %-# | CCR10-APC-A Mean | CCR10-APC-A Median |
|---|---|---|---|
| POS-unpigm. | 12,08 | 1,59 | 1,25 |
| POS-pigm. | 13,10 | 2,12 | 1,89 |

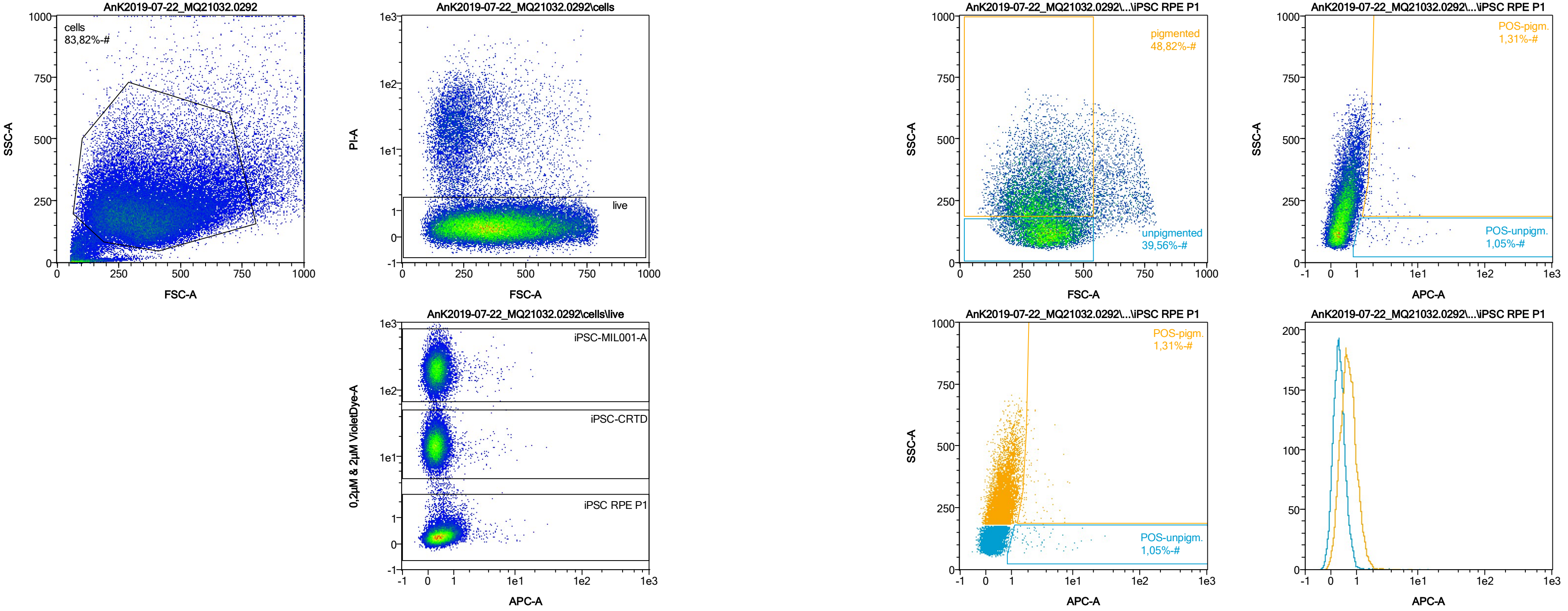


| Name | %-# | APC-A Mean | APC-A Median |
|---|---|---|---|
| POS-unpigm. | 1,05 | 4,11 | 2,28 |
| POS-pigm. | 1,31 | 2,72 | 1,99 |

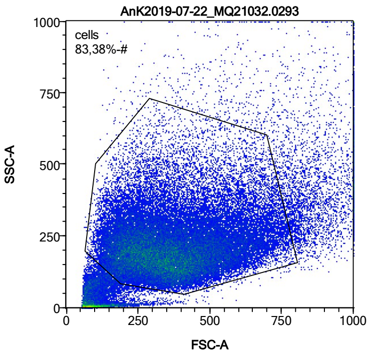


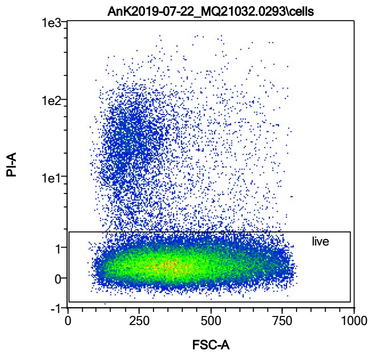


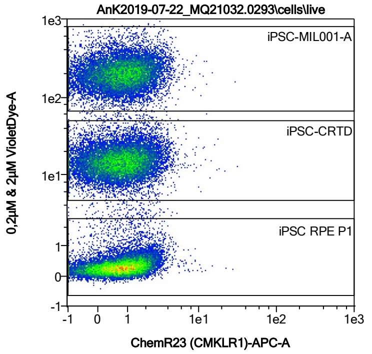


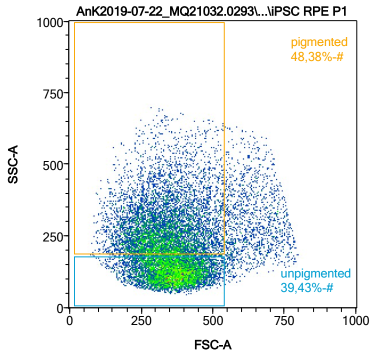


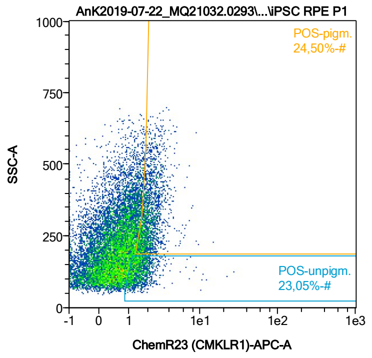


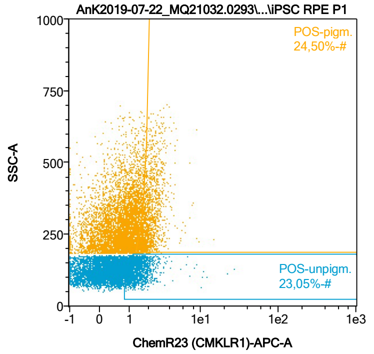


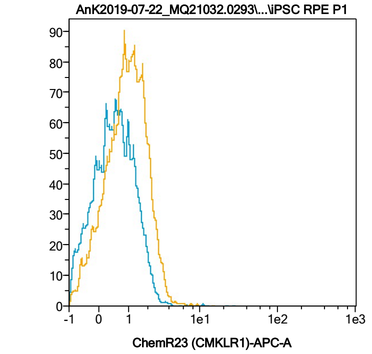


| Name | %-# | ChemR23 (CMKLR1)-APC-A Mean | ChemR23 (CMKLR1)-APC-A Median |
|---|---|---|---|
| POS-unpigm. | 23,05 | 1,61 | 1,41 |
| POS-pigm. | 24,50 | 2,19 | 2,00 |

Date:2024-06-14
Operator: Sebastian Knoebel
Version:2.13.0+54.g95abf0f4

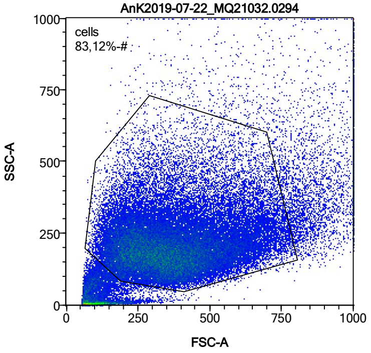


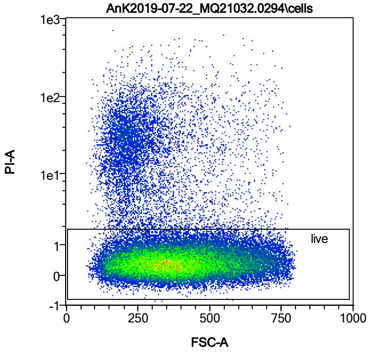


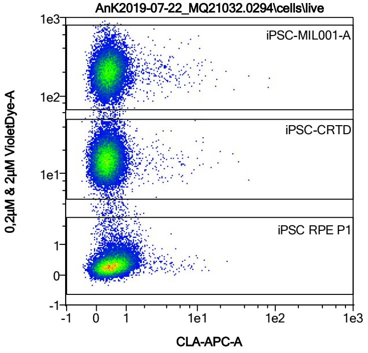


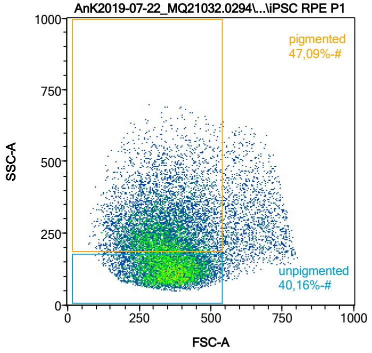


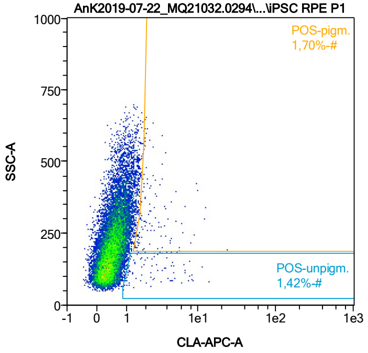


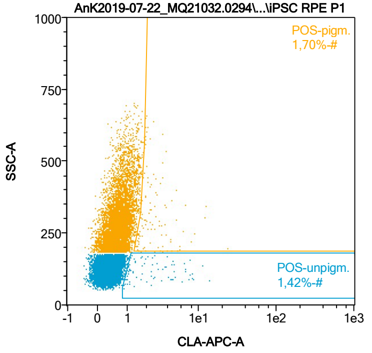


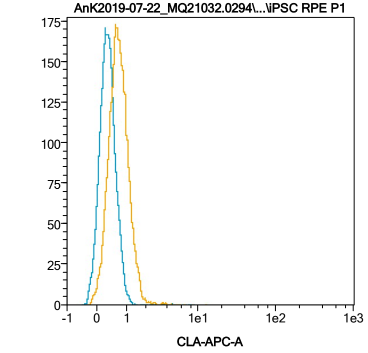


| Name | %-# | CLA-APC-A Mean | CLA-APC-A Median |
|---|---|---|---|
| POS-unpigm. | 1,42 | 2,98 | 1,91 |
| POS-pigm. | 1,70 | 3,01 | 2,13 |

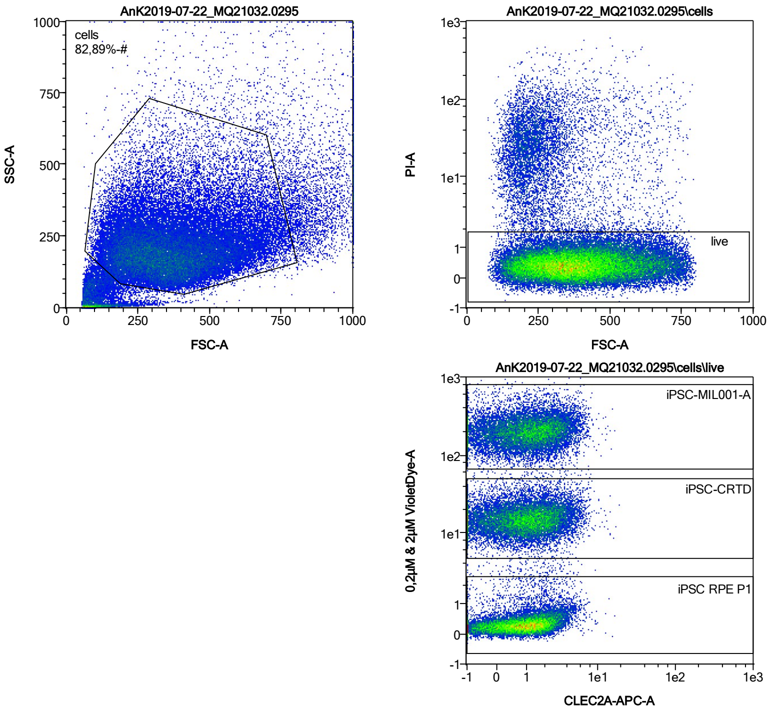


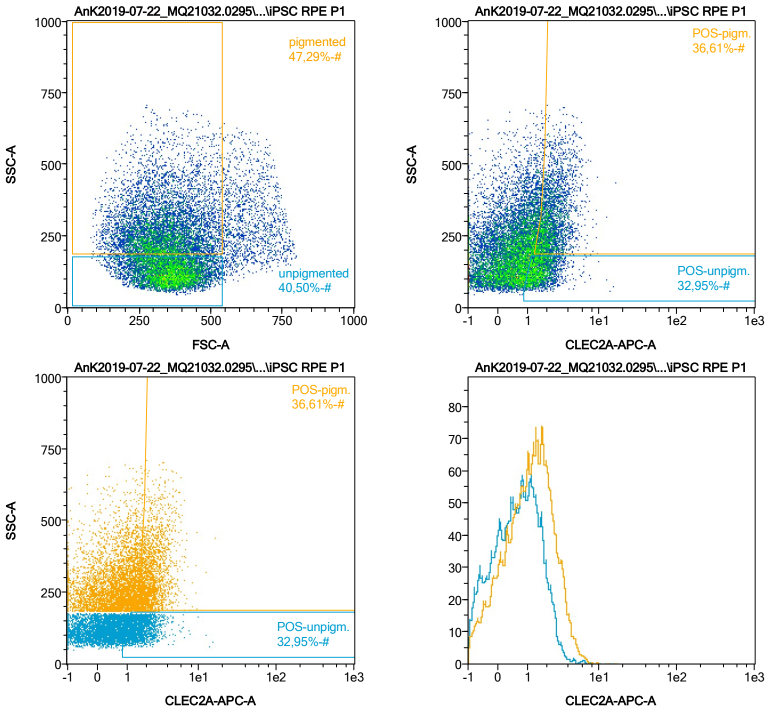


| Name | %-# | CLEC2A-APC-A Mean | CLEC2A-APC-A Median |
|---|---|---|---|
| POS-unpigm. | 32,95 | 1,74 | 1,55 |
| POS-pigm. | 36,61 | 2,50 | 2,22 |

Date:2024-06-14

Operator: Sebastian Knoebel

Version:2.13.0+54.g95abf0f4

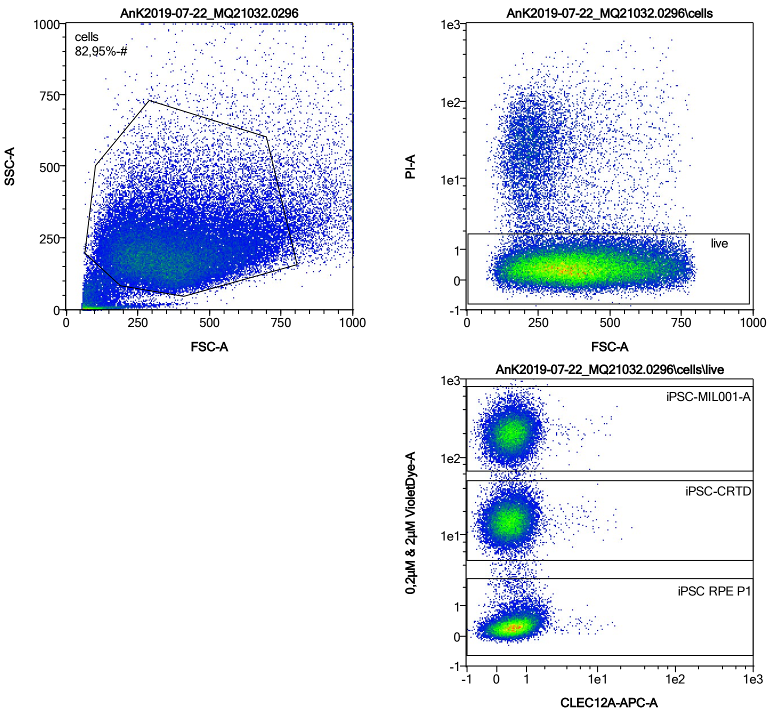


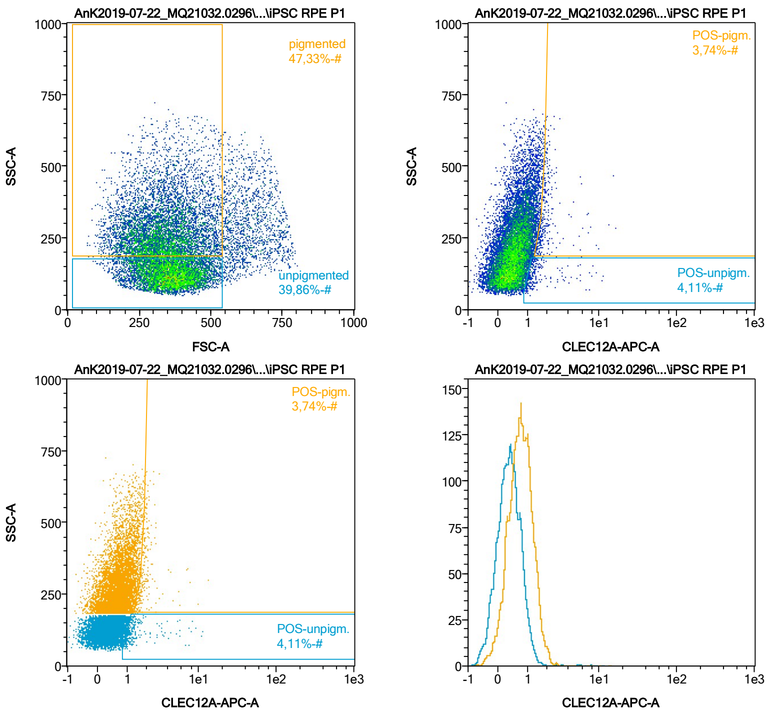


| Name | %-# | CLEC12A-APC-A Mean | CLEC12A-APC-A Median |
|---|---|---|---|
| POS-unpigm. | 4,11 | 1,71 | 1,19 |
| POS-pigm. | 3,74 | 2,06 | 1,83 |

Date:2024-06-14

Operator: Sebastian Knoebel

Version:2.13.0+54.g95abf0f4

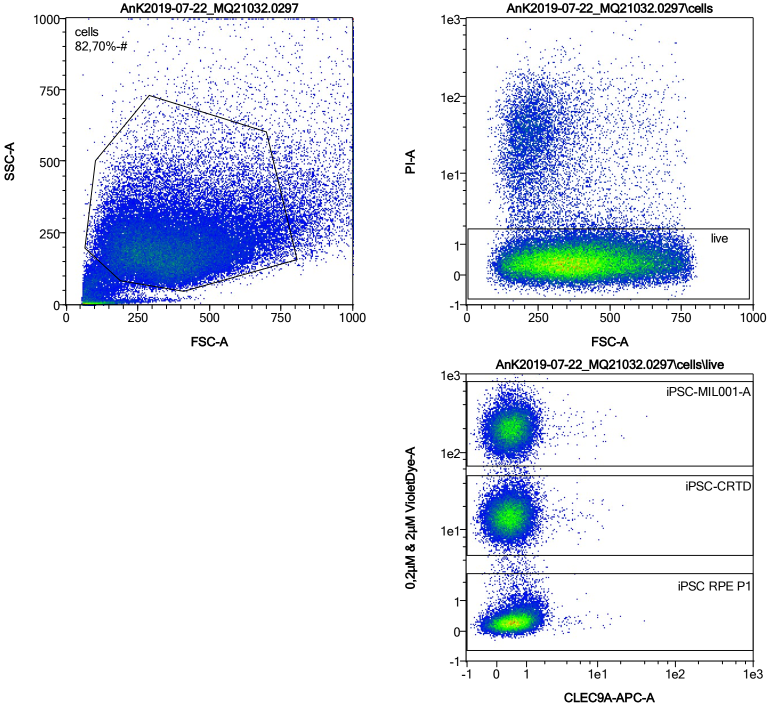


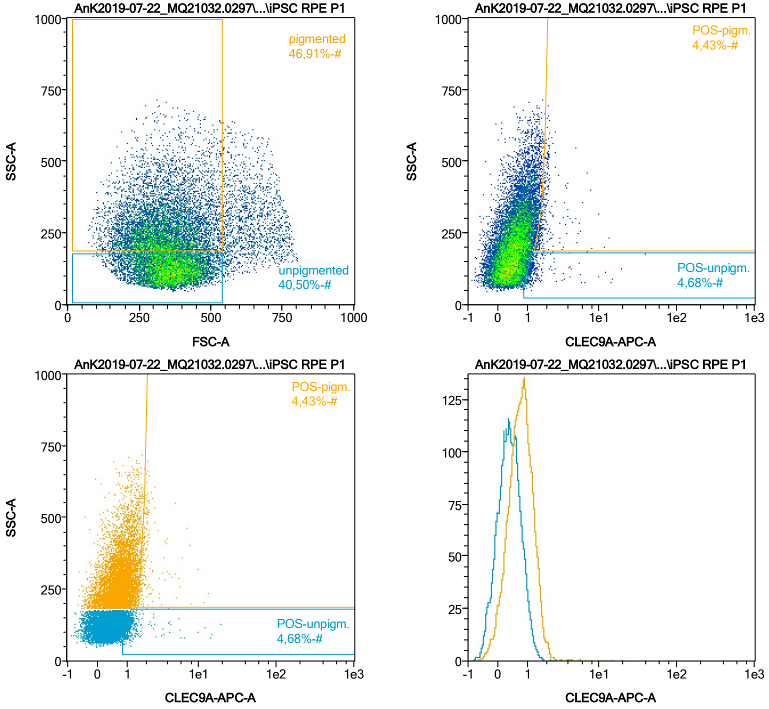


| Name | %-# | CLEC9A-APC-A Mean | CLEC9A-APC-A Median |
|---|---|---|---|
| POS-unpigm. | 4,68 | 1,60 | 1,16 |
| POS-pigm. | 4,43 | 2,14 | 1,79 |

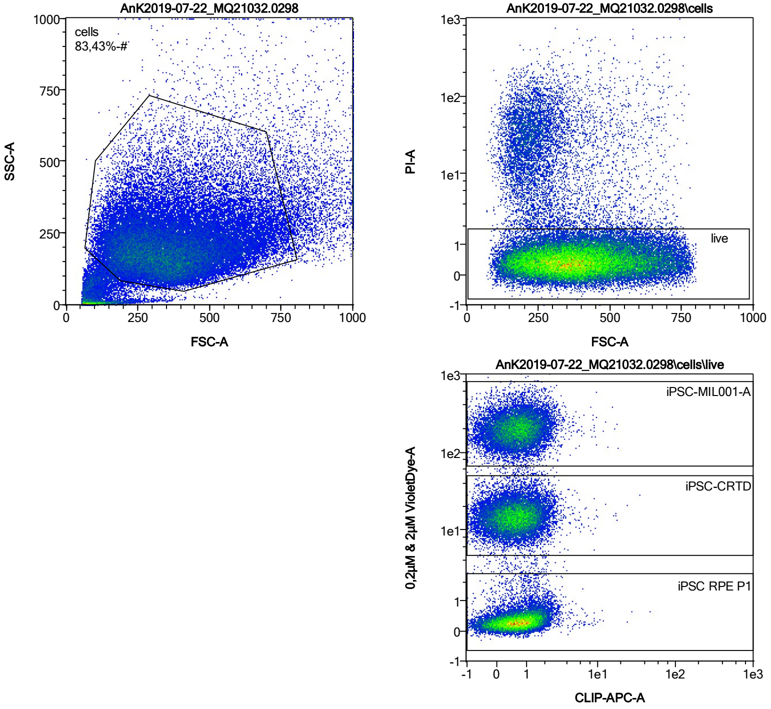


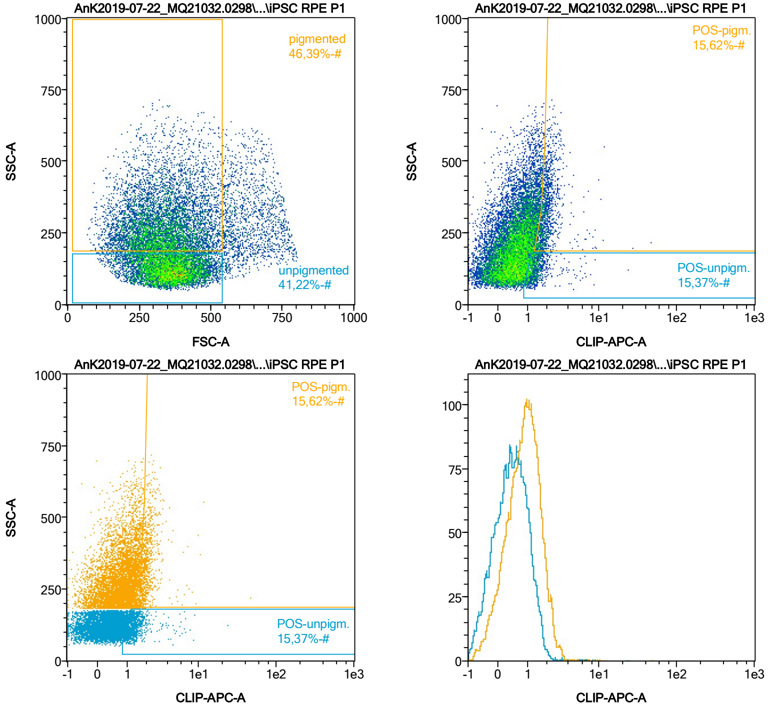


| Name | %-# | CLIP-APC-A Mean | CLIP-APC-A Median |
|---|---|---|---|
| POS-unpigm. | 15,37 | 1,47 | 1,26 |
| POS-pigm. | 15,62 | 2,10 | 1,91 |

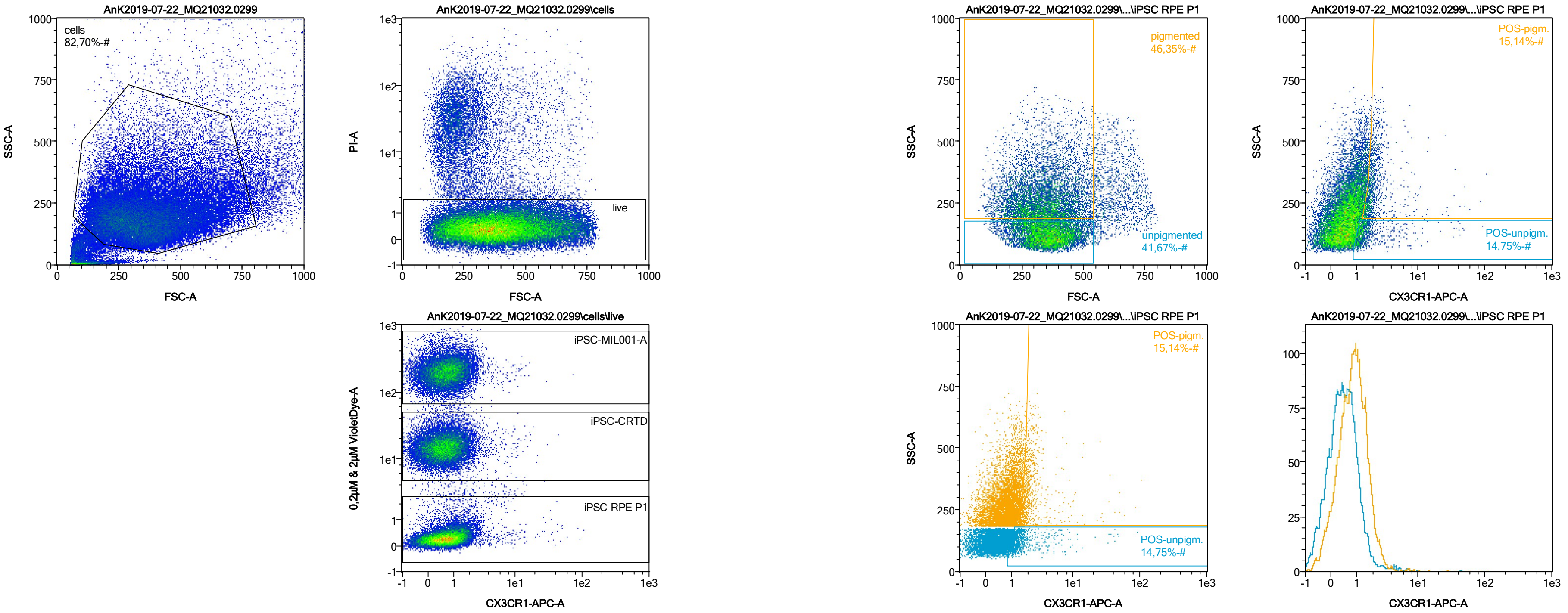


| Name | %-# | CX3CR1-APC-A Mean | CX3CR1-APC-A Median |
|---|---|---|---|
| POS-unpigm. | 14,75 | 2,22 | 1,32 |
| POS-pigm. | 15,14 | 2,81 | 1,90 |

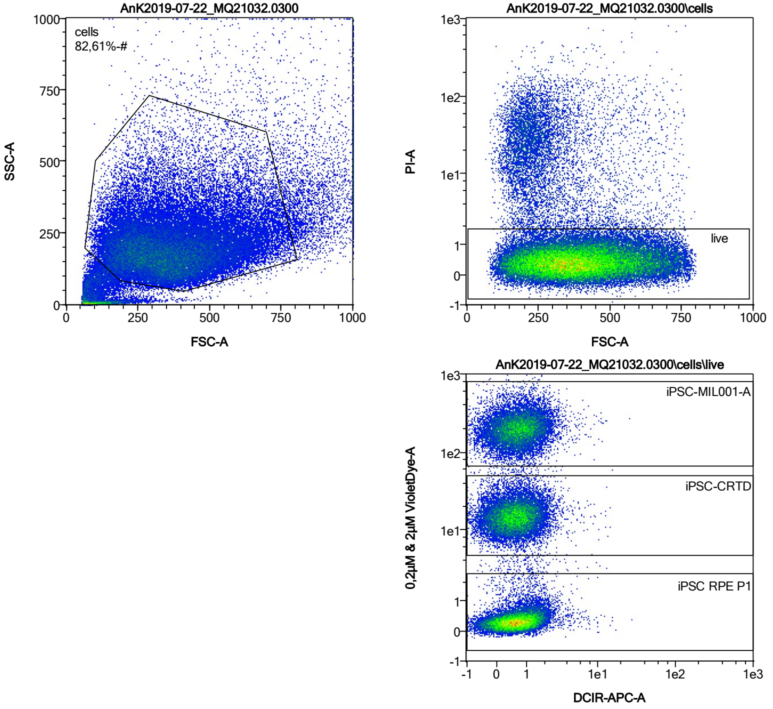


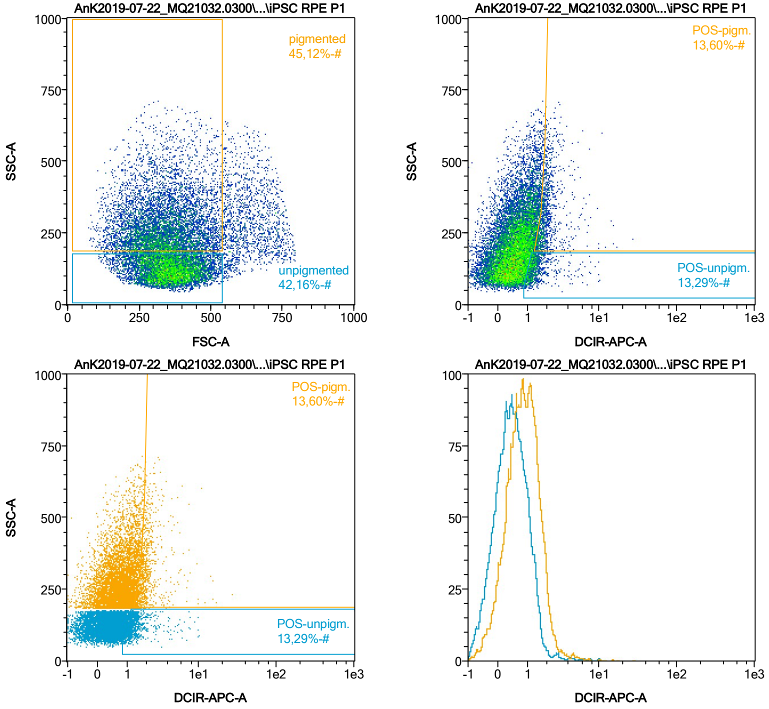


| Name | %-# | DCIR-APC-A Mean | DCIR-APC-A Median |
|---|---|---|---|
| POS-unpigm. | 13,29 | 1,54 | 1,28 |
| POS-pigm. | 13,60 | 2,23 | 1,87 |

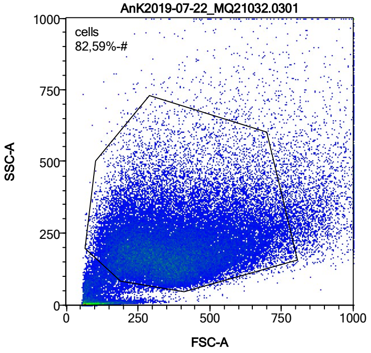


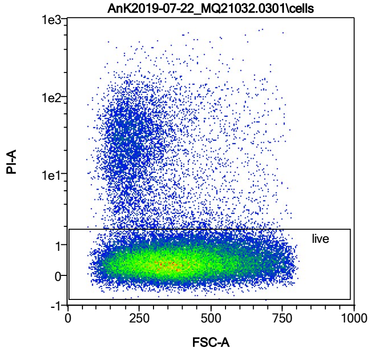


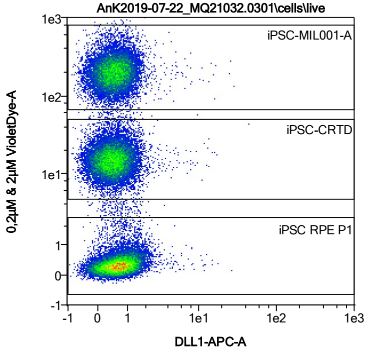


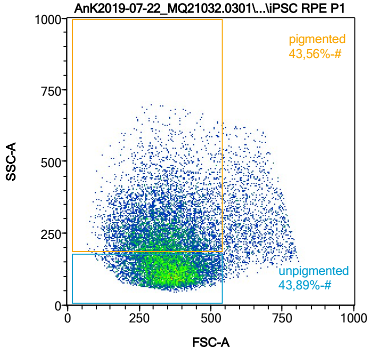


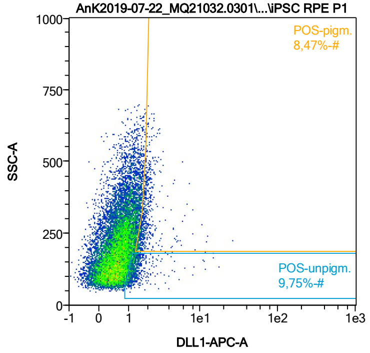


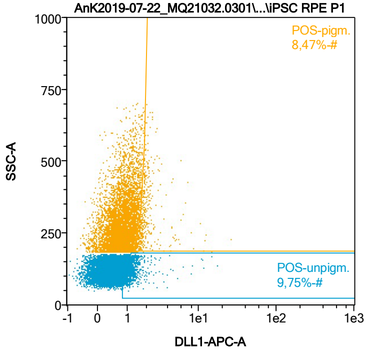


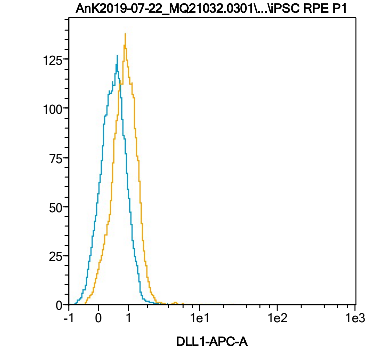


| Name | %-# | DLL1-APC-A Mean | DLL1-APC-A Median |
|---|---|---|---|
| POS-unpigm. | 9,75 | 1,45 | 1,22 |
| POS-pigm. | 8,47 | 2,11 | 1,77 |

Date:2024-06-14

Operator: Sebastian Knoebel

Version:2.13.0+54.g95abf0f4

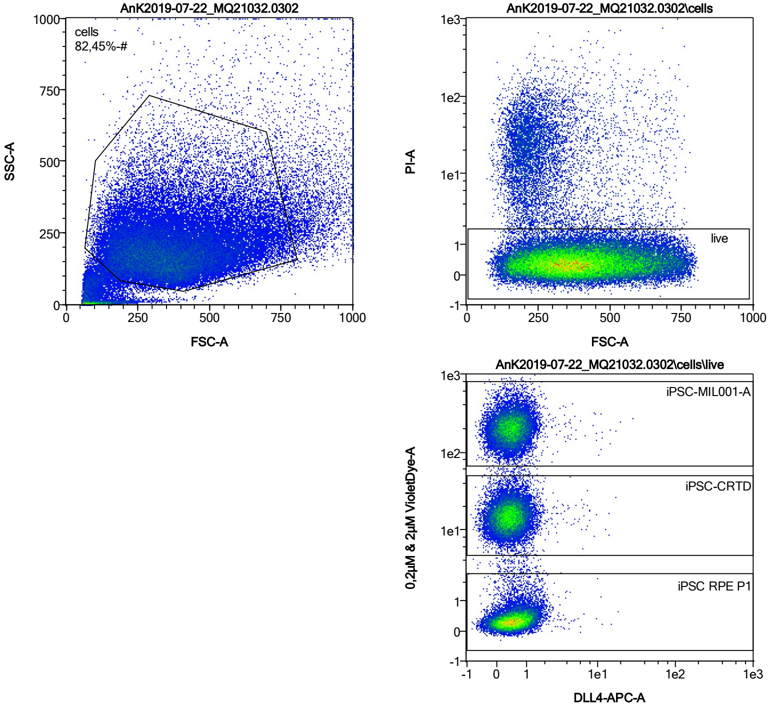


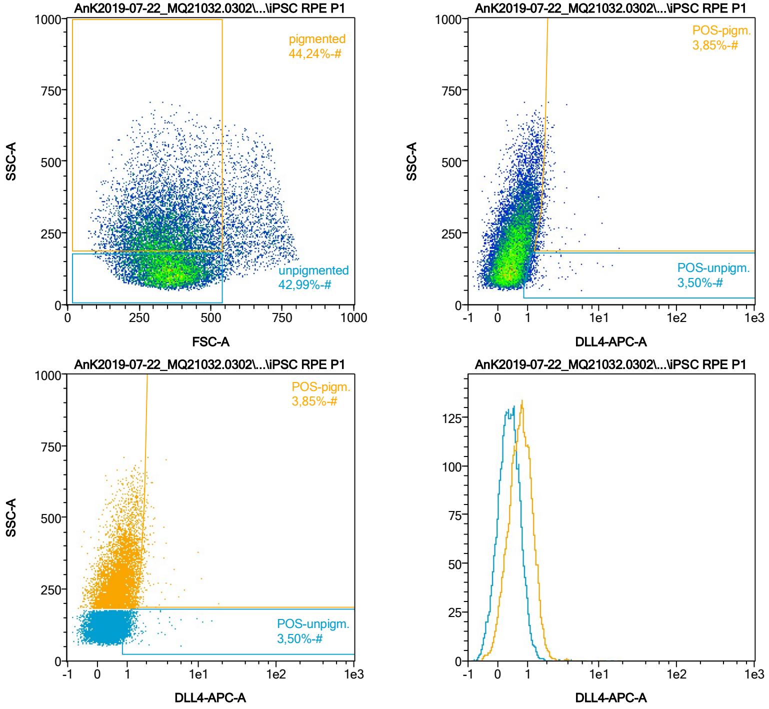


| Name | %-# | DLL4-APC-A Mean | DLL4-APC-A Median |
|---|---|---|---|
| POS-unpigm. | 3,50 | 1,51 | 1,16 |
| POS-pigm. | 3,85 | 2,03 | 1,78 |

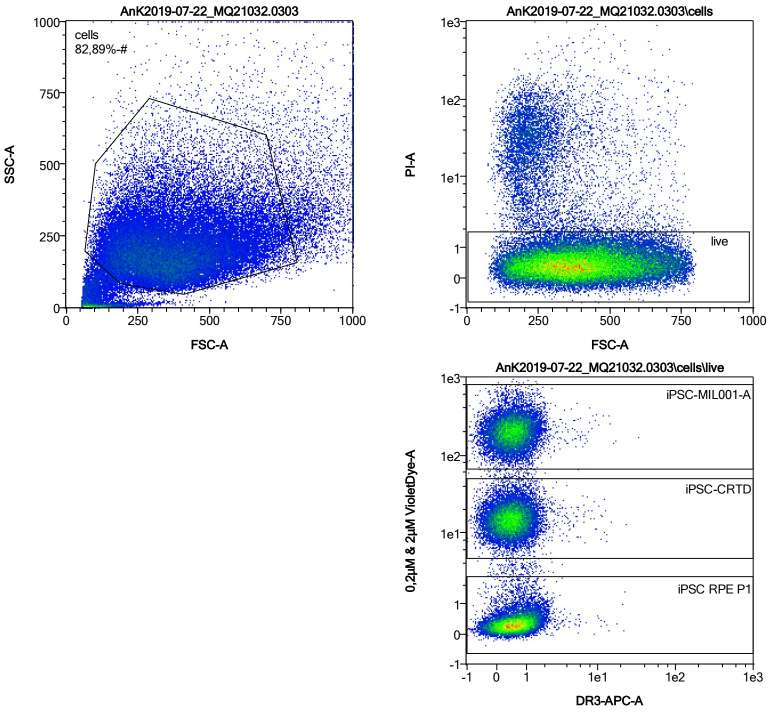


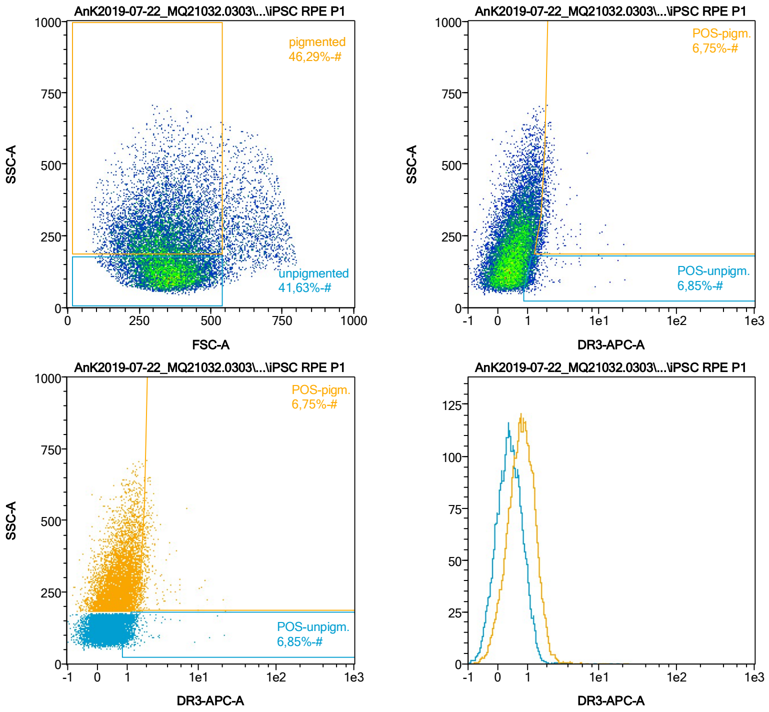


| Name | %-# | DR3-APC-A Mean | DR3-APC-A Median |
|---|---|---|---|
| POS-unpigm. | 6,85 | 1,42 | 1,20 |
| POS-pigm. | 6,75 | 2,18 | 1,85 |

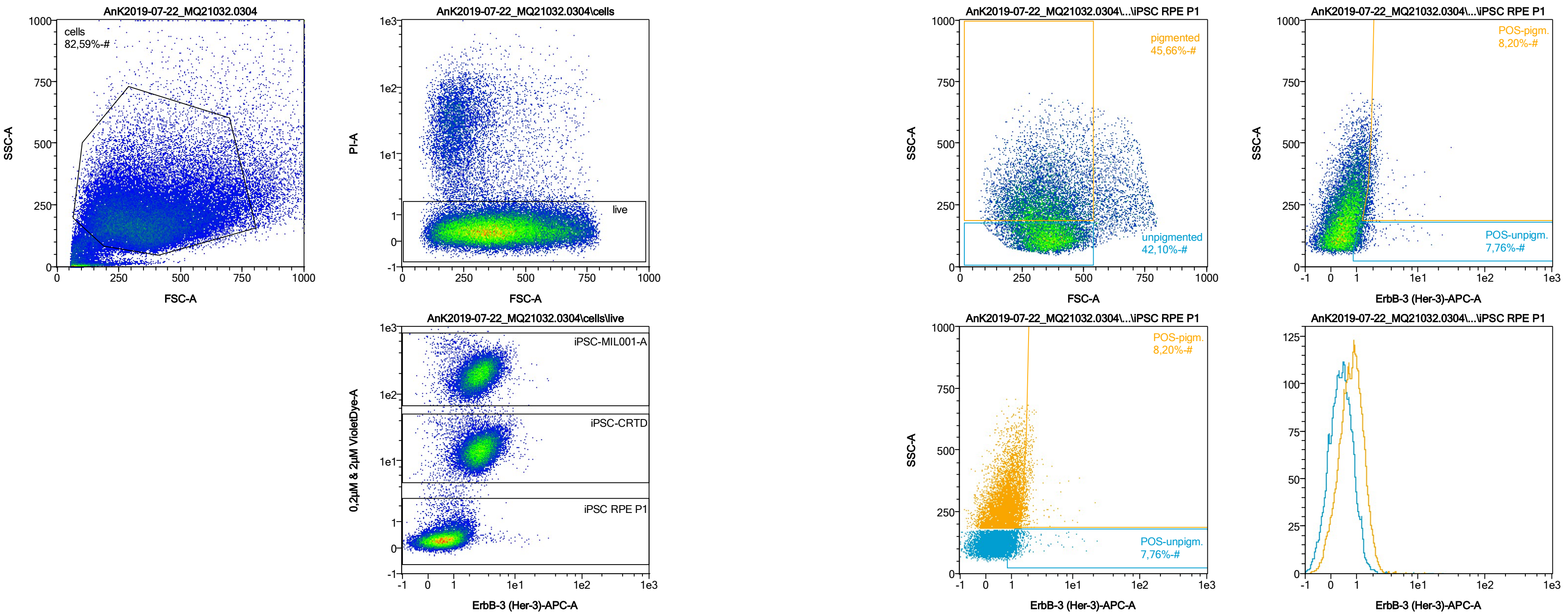


| Name | %-# | ErbB-3 (Her-3)-APC-A Mean | ErbB-3 (Her-3)-APC-A Median |
|---|---|---|---|
| POS-unpigm. | 7,76 | 1,61 | 1,22 |
| POS-pigm. | 8,20 | 2,05 | 1,81 |

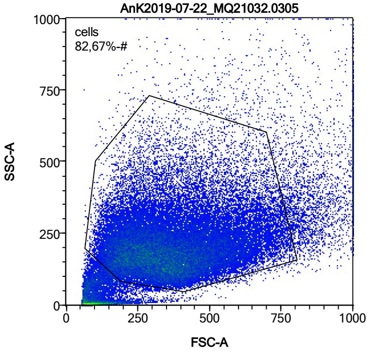


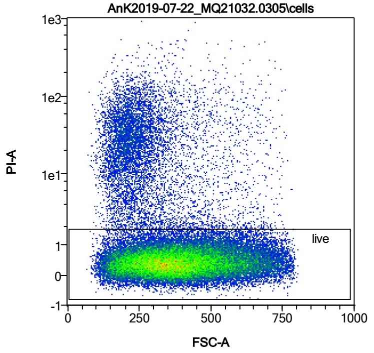


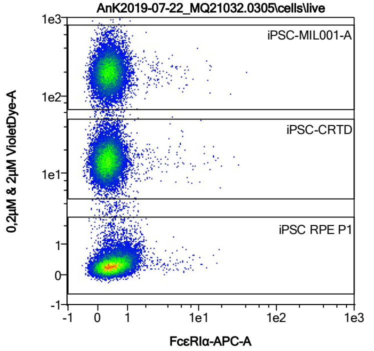


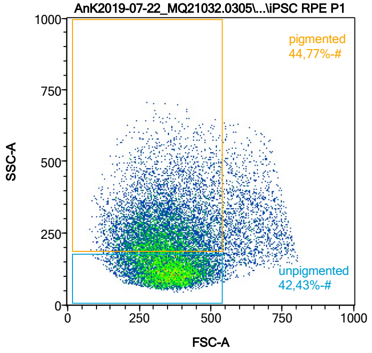


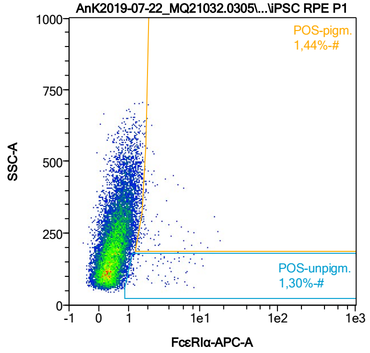


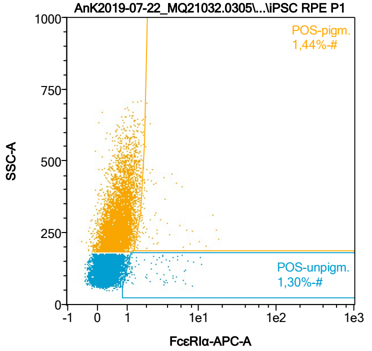


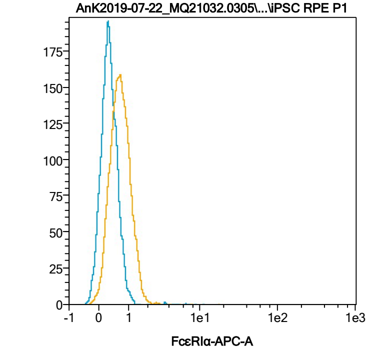


| Name | %-# | FcεRIα-APC-A Mean | FcεRIα-APC-A Median |
|---|---|---|---|
| POS-unpigm. | 1,30 | 2,88 | 2,00 |
| POS-pigm. | 1,44 | 3,08 | 1,92 |

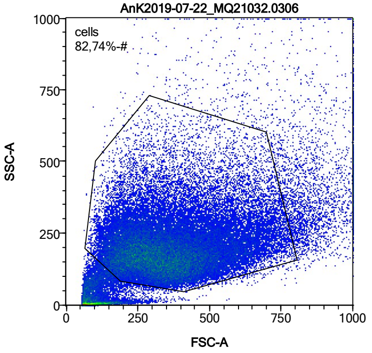


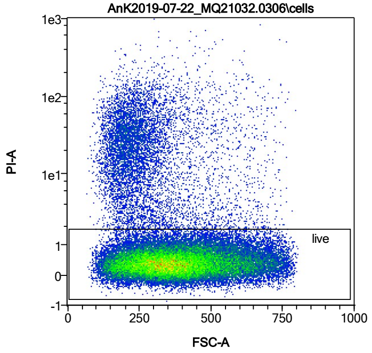


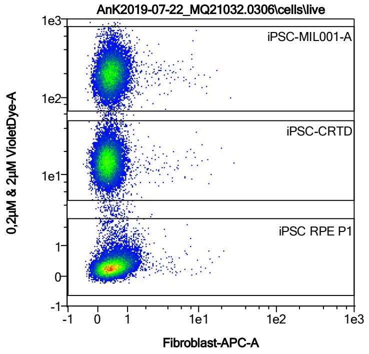


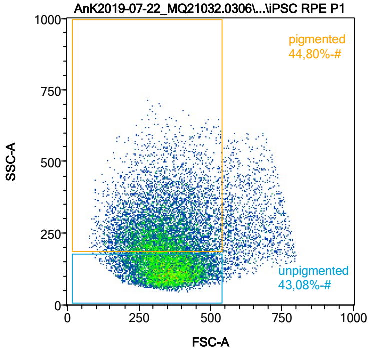


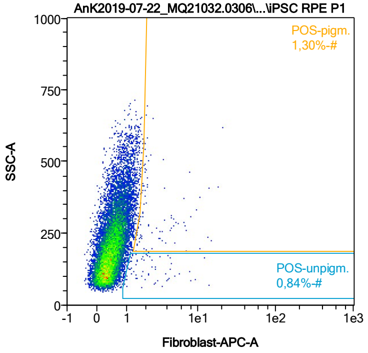


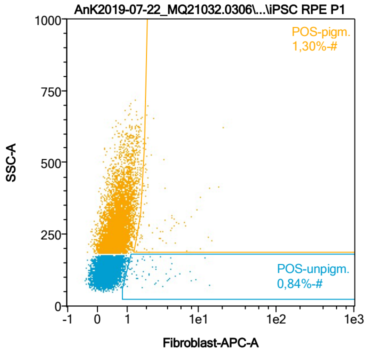


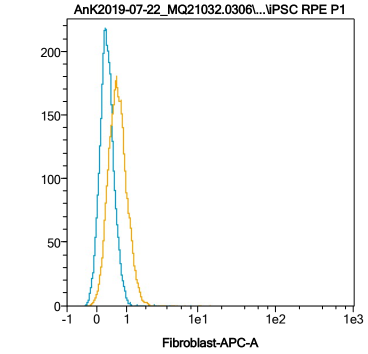


| Name | %-# | Fibroblast-APC-A Mean | Fibroblast-APC-A Median |
|---|---|---|---|
| POS-unpigm. | 0,84 | 2,93 | 1,89 |
| POS-pigm. | 1,30 | 3,64 | 2,08 |

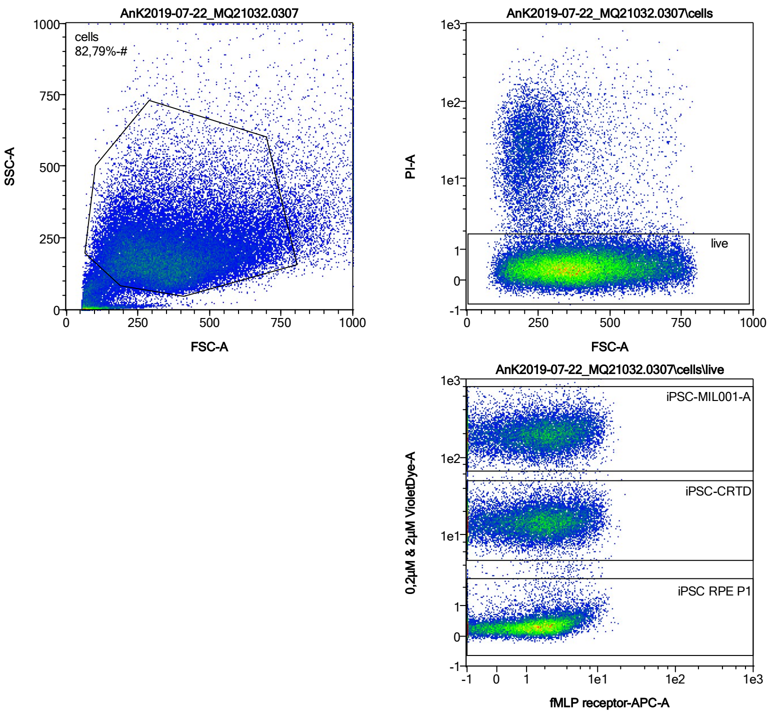


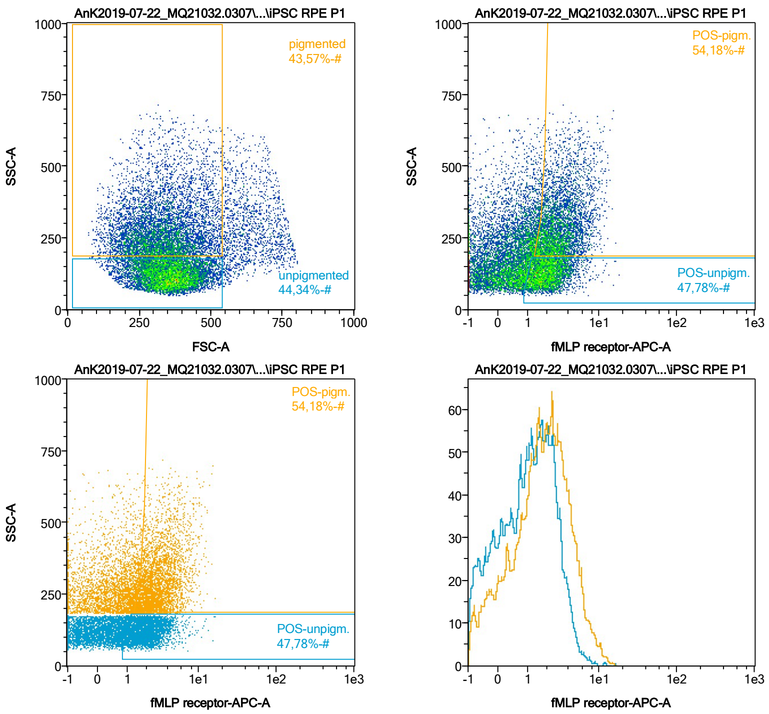


| Name | %-# | fMLP receptor-APC-A Mean | fMLP receptor-APC-A Median |
|---|---|---|---|
| POS-unpigm. | 47,78 | 2,25 | 1,97 |
| POS-pigm. | 54,18 | 3,37 | 2,84 |

Date:2024-06-14
Operator: Sebastian Knoebel
Version:2.13.0+54.g95abf0f4

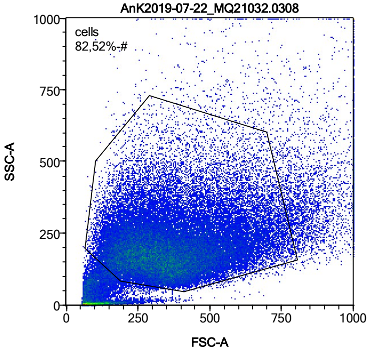


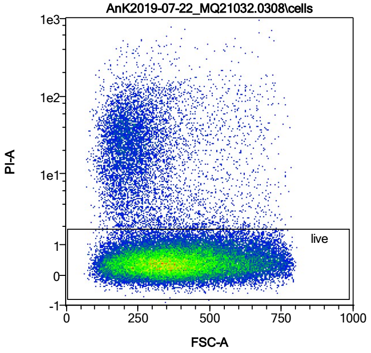


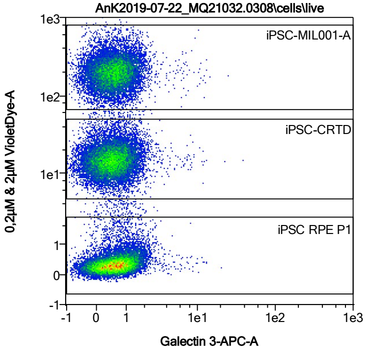


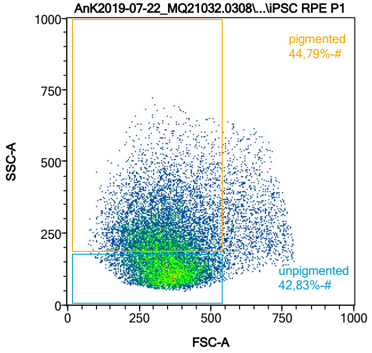


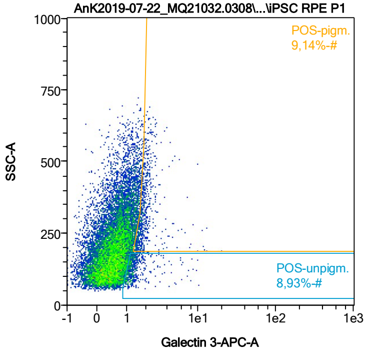


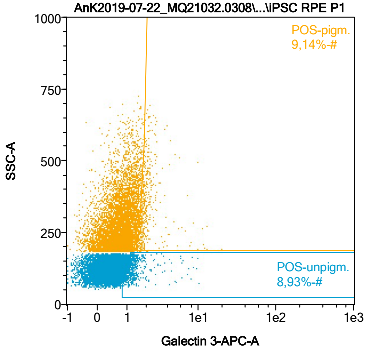


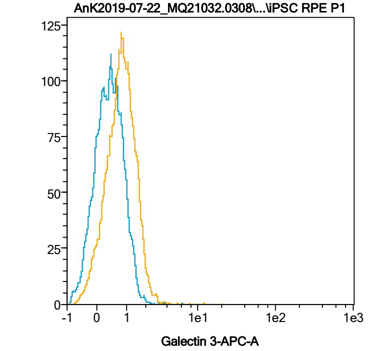


| Name | %-# | Galectin 3-APC-A Mean | Galectin 3-APC-A Median |
|---|---|---|---|
| POS-unpigm. | 8,93 | 1,49 | 1,22 |
| POS-pigm. | 9,14 | 2,14 | 1,83 |

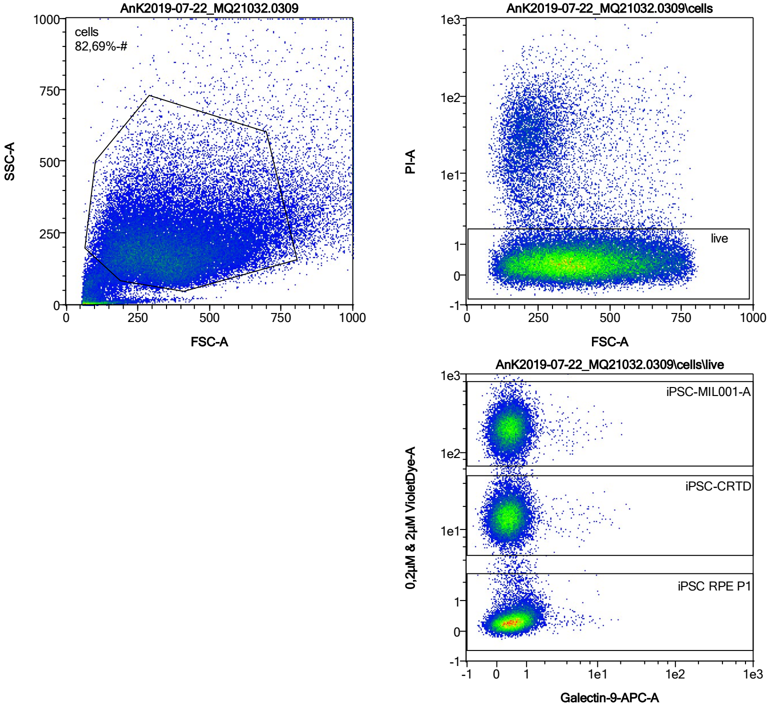


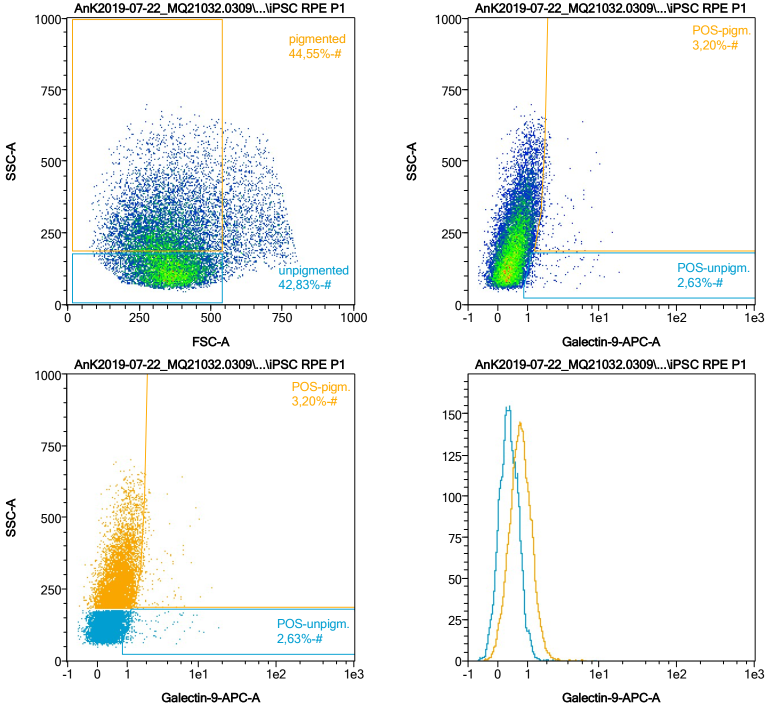


| Name | %-# | Galectin-9-APC-A Mean | Galectin-9-APC-A Median |
|---|---|---|---|
| POS-unpigm. | 2,63 | 2,06 | 1,21 |
| POS-pigm. | 3,20 | 2,50 | 1,88 |

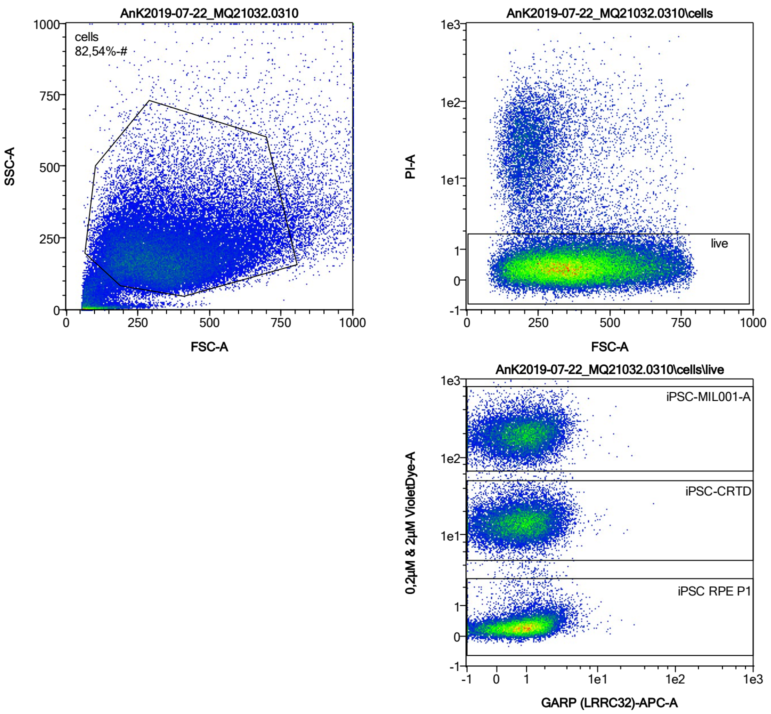


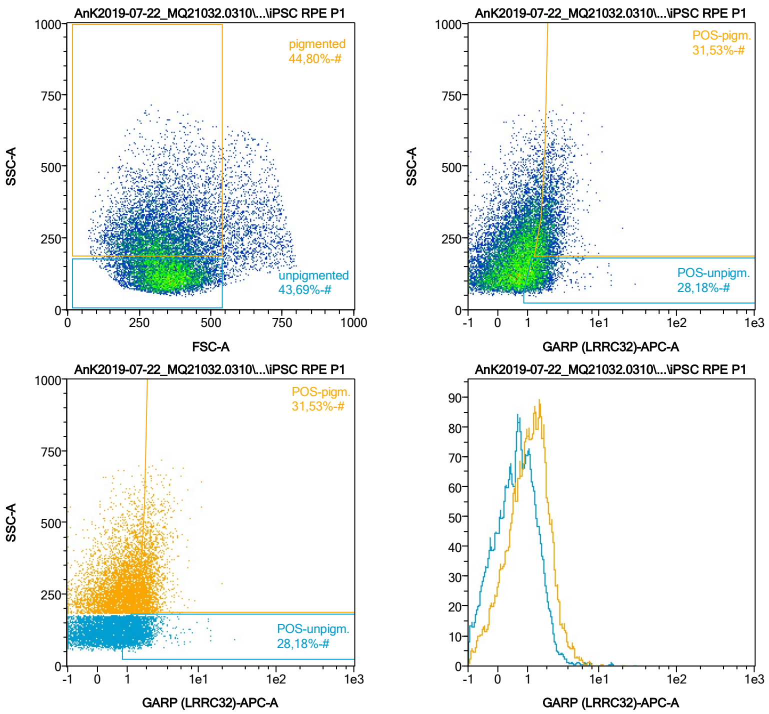


| Name | %-# | GARP (LRRC32)-APC-A Mean | GARP (LRRC32)-APC-A Median |
|---|---|---|---|
| POS-unpigm. | 28,18 | 1,68 | 1,43 |
| POS-pigm. | 31,53 | 2,28 | 2,03 |

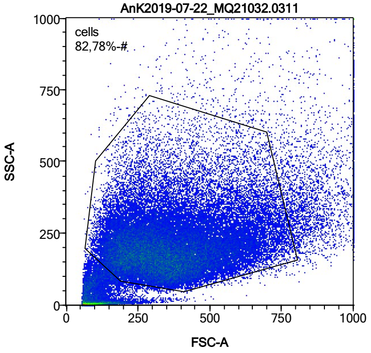


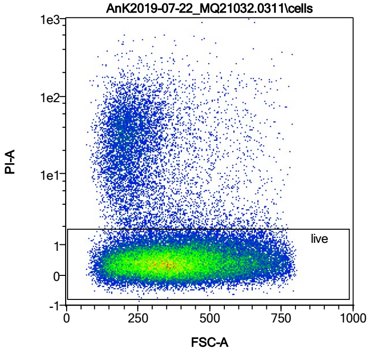


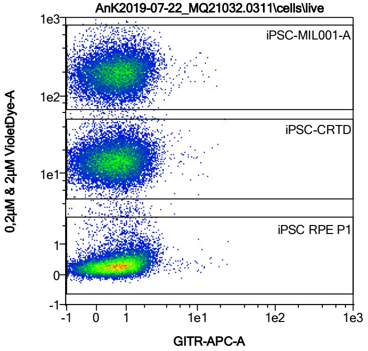


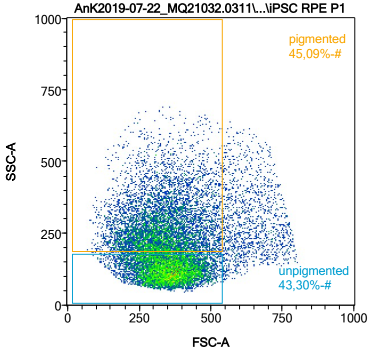


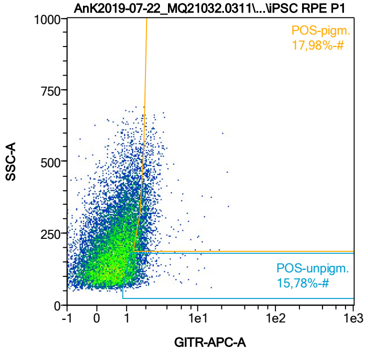


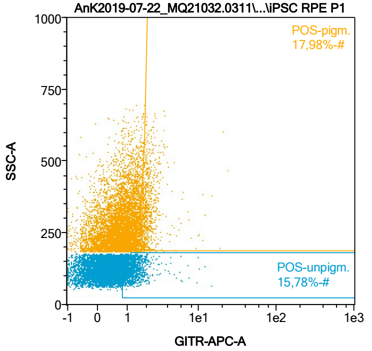


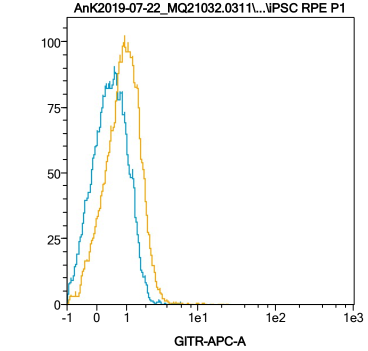


| Name | %-# | GITR-APC-A Mean | GITR-APC-A Median |
|---|---|---|---|
| POS-unpigm. | 15,78 | 1,47 | 1,29 |
| POS-pigm. | 17,98 | 2,11 | 1,87 |

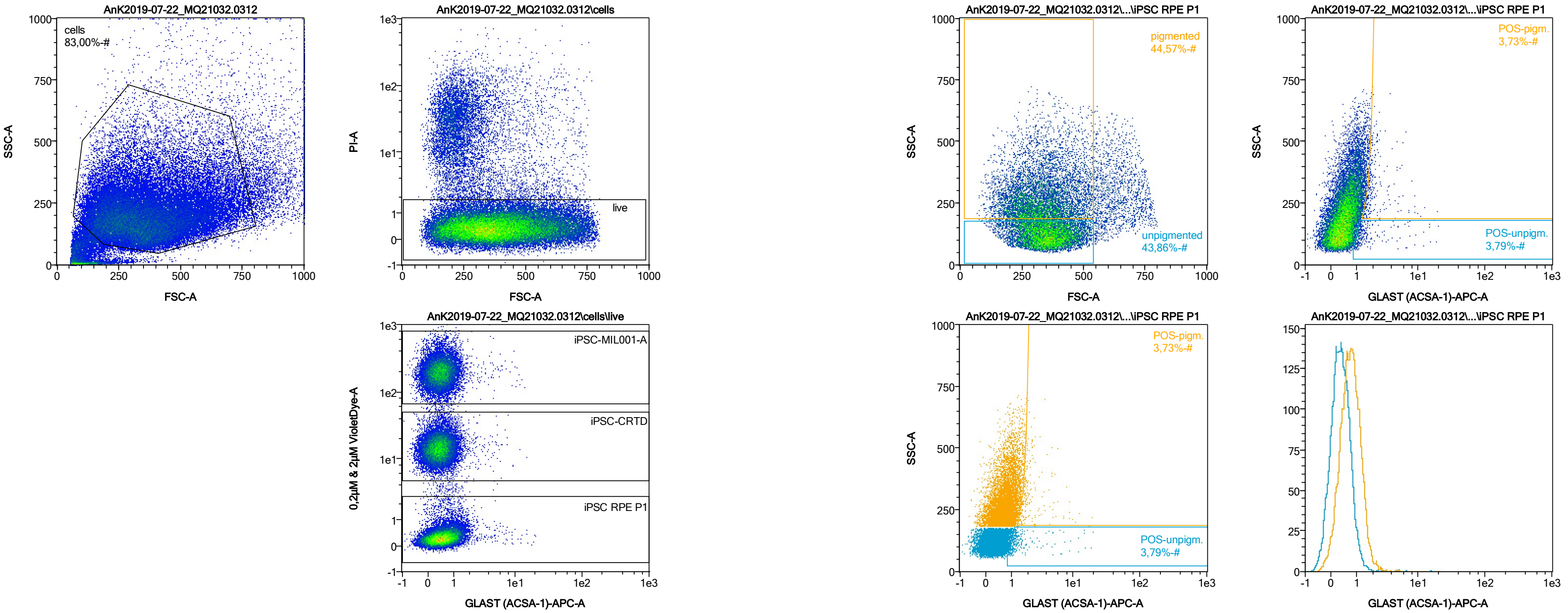


| Name | %-# | GLAST (ACSA-1)-APC-A Mean | GLAST (ACSA-1)-APC-A Median |
|---|---|---|---|
| POS-unpigm. | 3,79 | 1,96 | 1,21 |
| POS-pigm. | 3,73 | 2,47 | 1,86 |

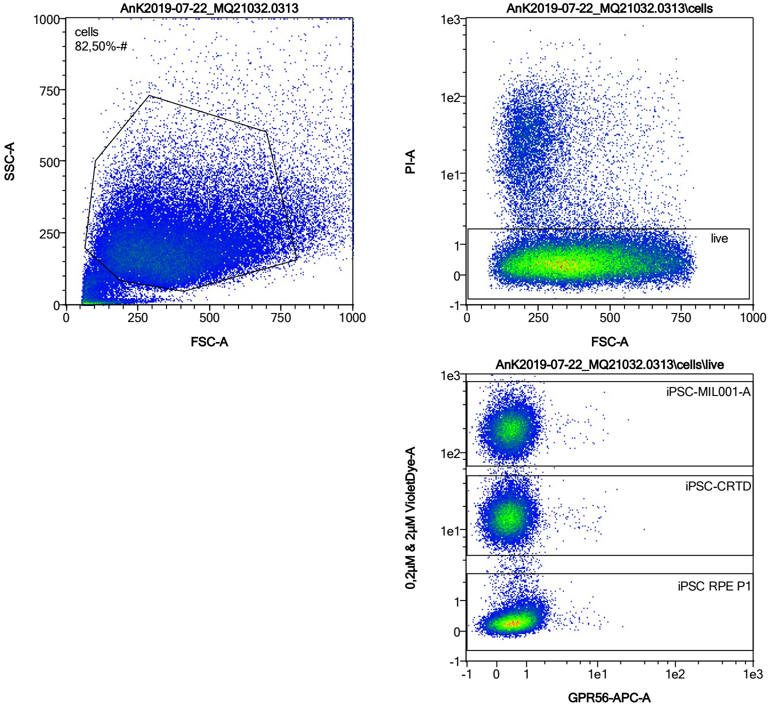


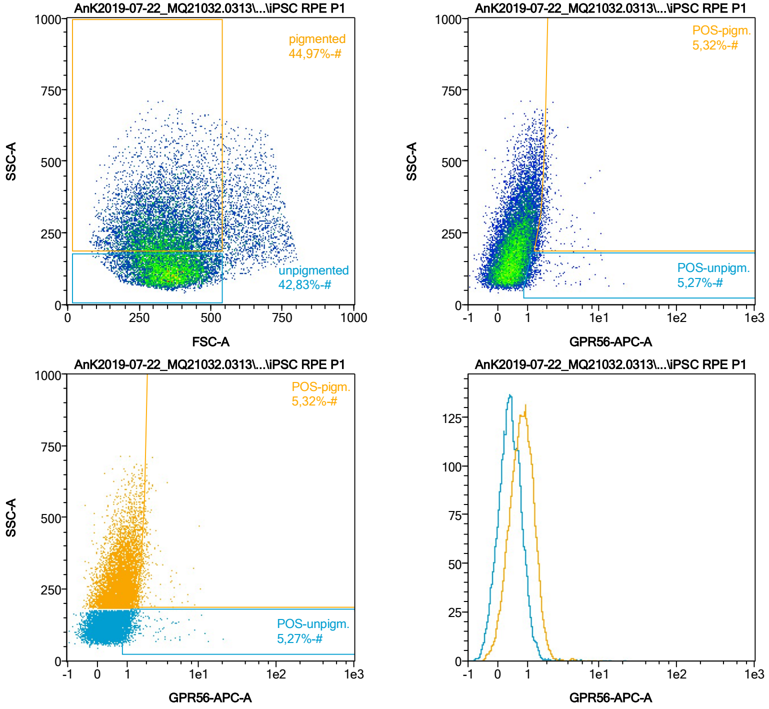


| Name | %-# | GPR56-APC-A Mean | GPR56-APC-A Median |
|---|---|---|---|
| POS-unpigm. | 5,27 | 1,65 | 1,19 |
| POS-pigm. | 5,32 | 2,05 | 1,80 |

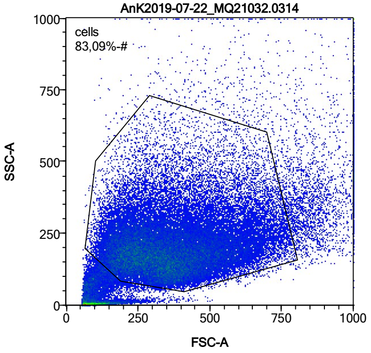


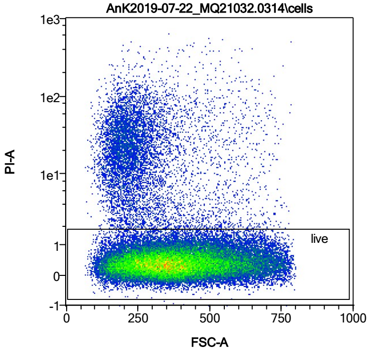


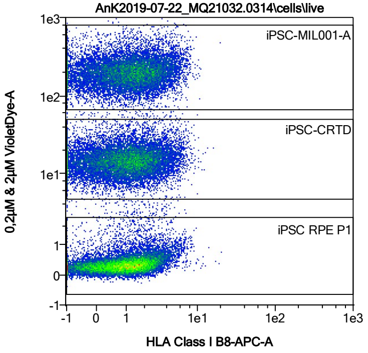


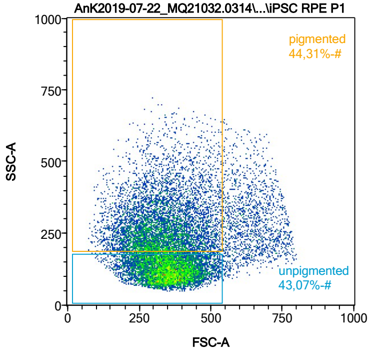


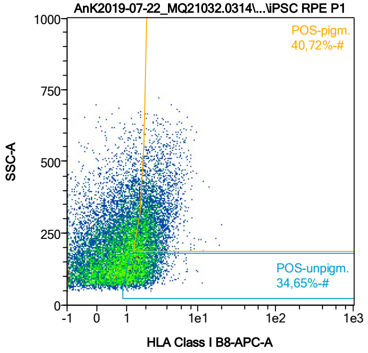


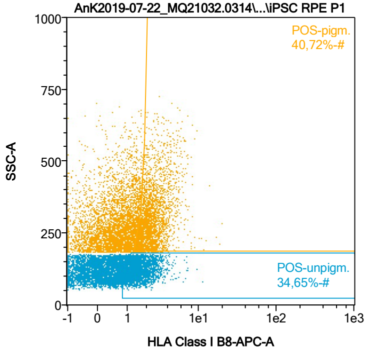


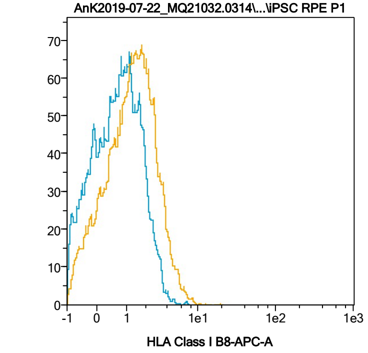


| Name | %-# | HLA Class I B8-APC-A Mean | HLA Class I B8-APC-A Median |
|---|---|---|---|
| POS-unpigm. | 34,65 | 1,78 | 1,60 |
| POS-pigm. | 40,72 | 2,65 | 2,29 |

Date:2024-06-14
Operator: Sebastian Knoebel
Version:2.13.0+54.g95abf0f4

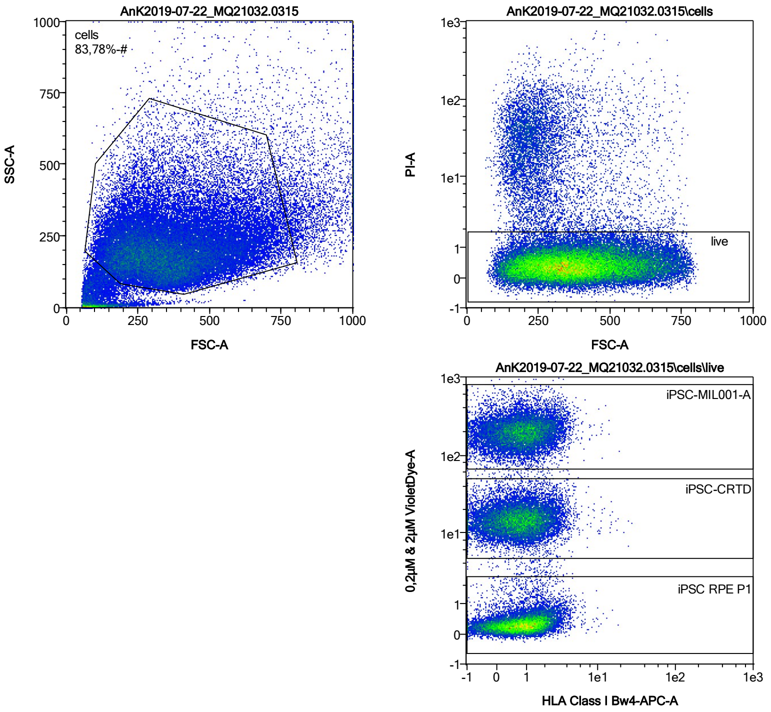


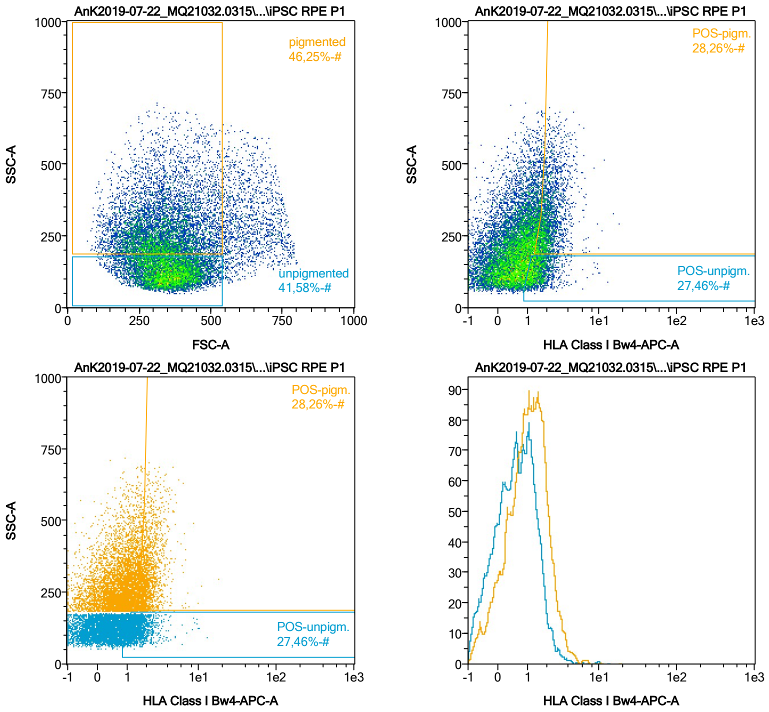


| Name | %-# | HLA Class I Bw4-APC-A Mean | HLA Class I Bw4-APC-A Median |
|---|---|---|---|
| POS-unpigm. | 27,46 | 1,57 | 1,39 |
| POS-pigm. | 28,26 | 2,20 | 1,99 |

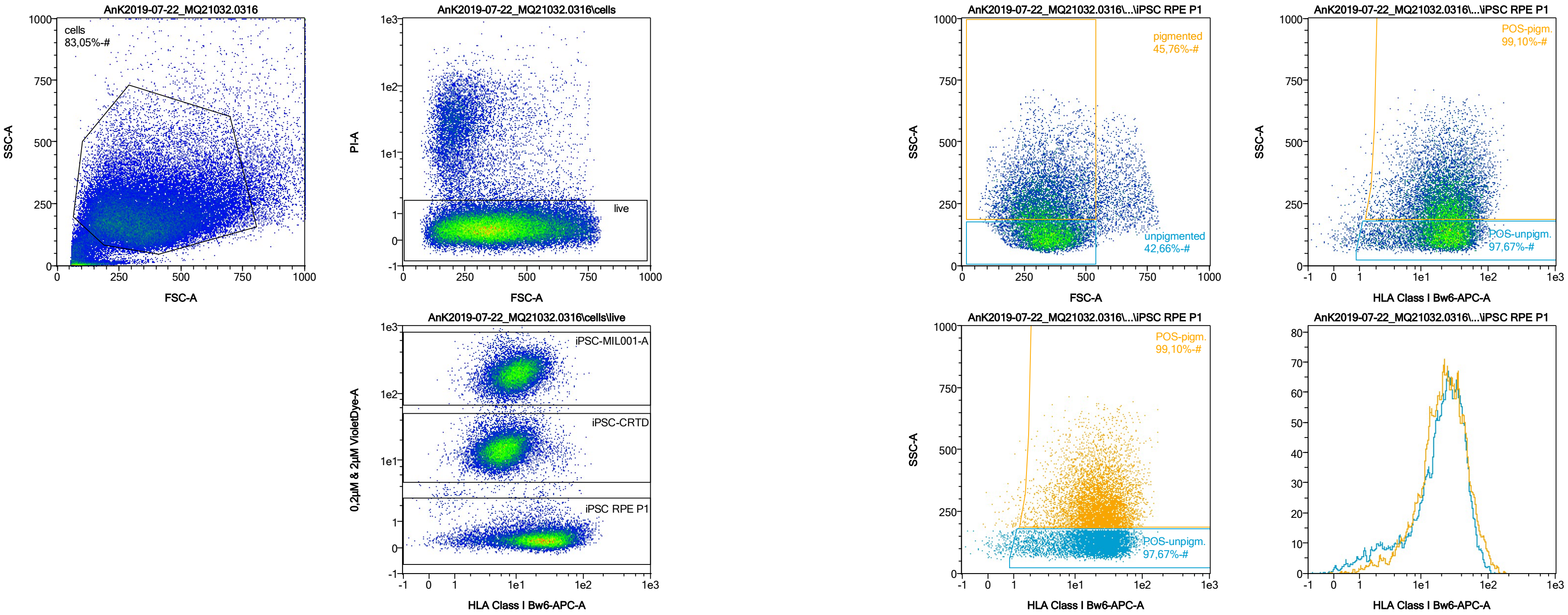


| Name | %-# | HLA Class I Bw6-APC-A Mean | HLA Class I Bw6-APC-A Median |
|---|---|---|---|
| POS-unpigm. | 97,67 | 24,84 | 22,25 |
| POS-pigm. | 99,10 | 27,52 | 22,66 |

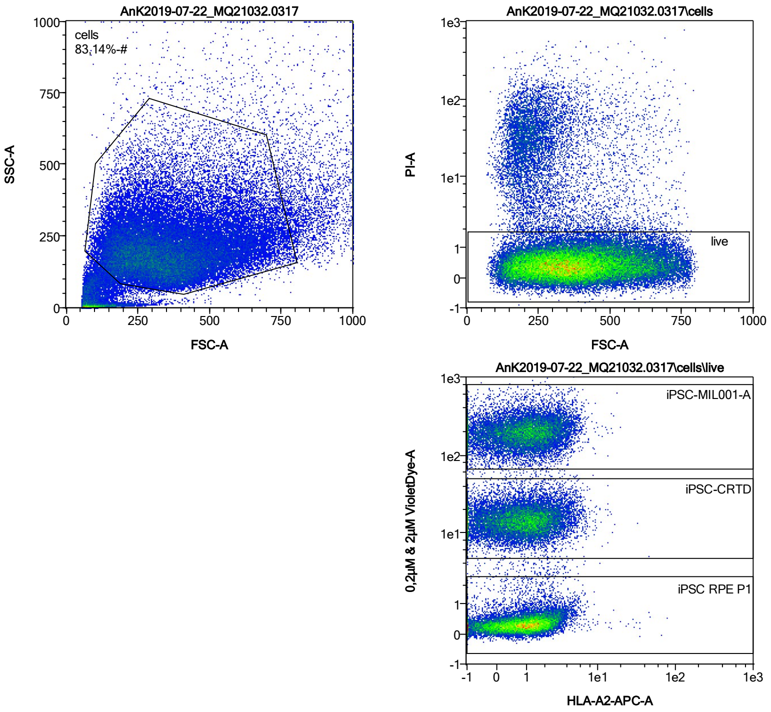


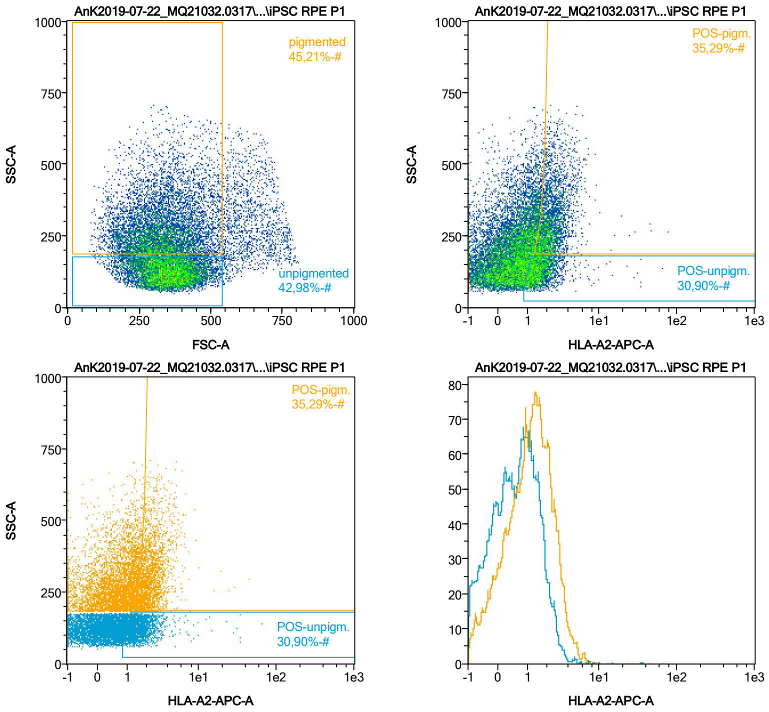


| Name | %-# | HLA-A2-APC-A Mean | HLA-A2-APC-A Median |
|---|---|---|---|
| POS-unpigm. | 30,90 | 1,82 | 1,49 |
| POS-pigm. | 35,29 | 2,48 | 2,19 |

Date:2024-06-14

Operator: Sebastian Knoebel

Version:2.13.0+54.g95abf0f4

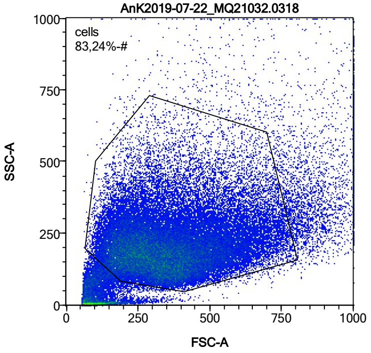


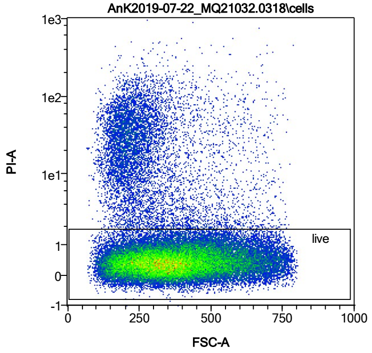


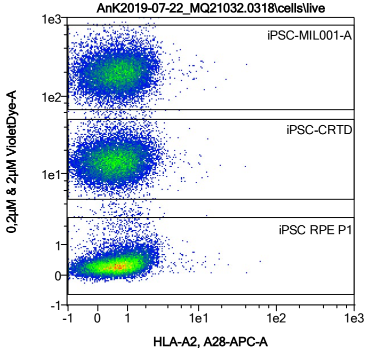


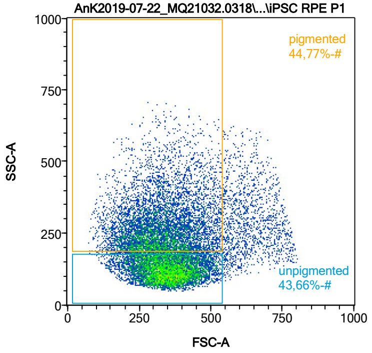


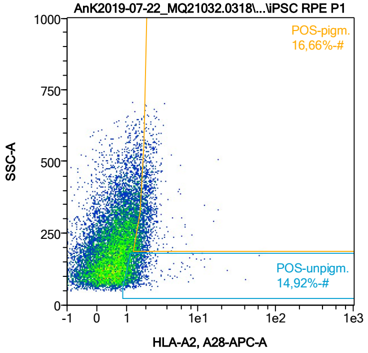


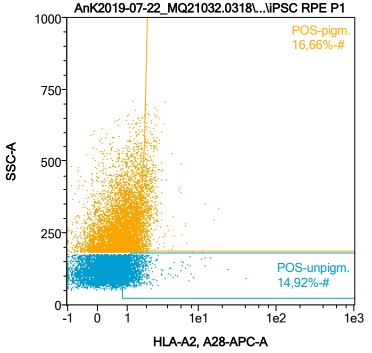


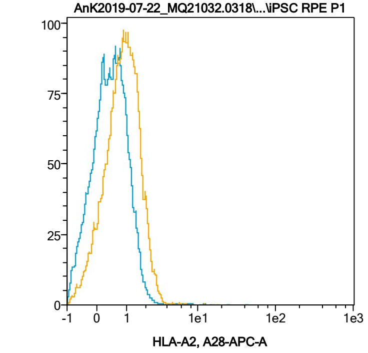


| Name | %-# | HLA-A2, A28-APC-A Mean | HLA-A2, A28-APC-A Median |
|---|---|---|---|
| POS-unpigm. | 14,92 | 1,57 | 1,29 |
| POS-pigm. | 16,66 | 2,05 | 1,88 |

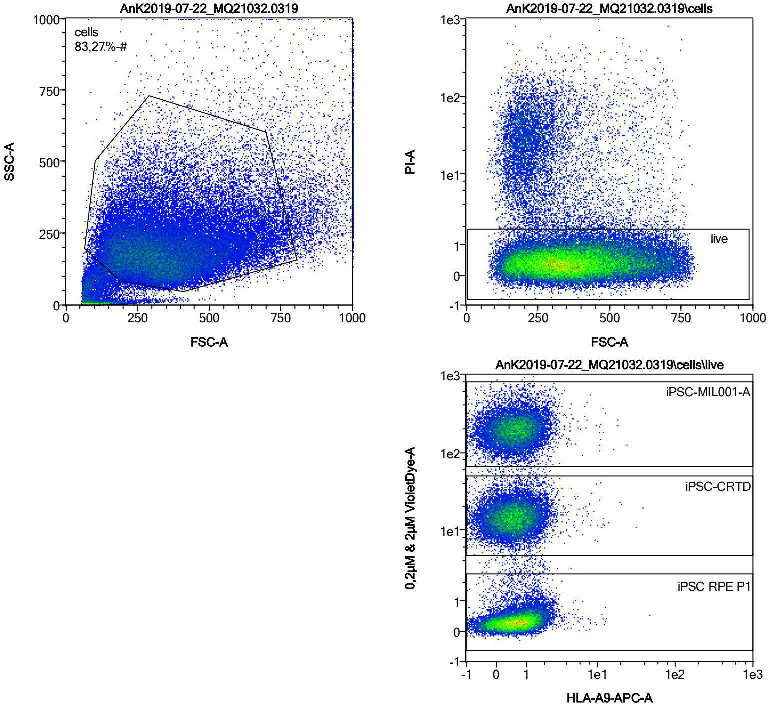


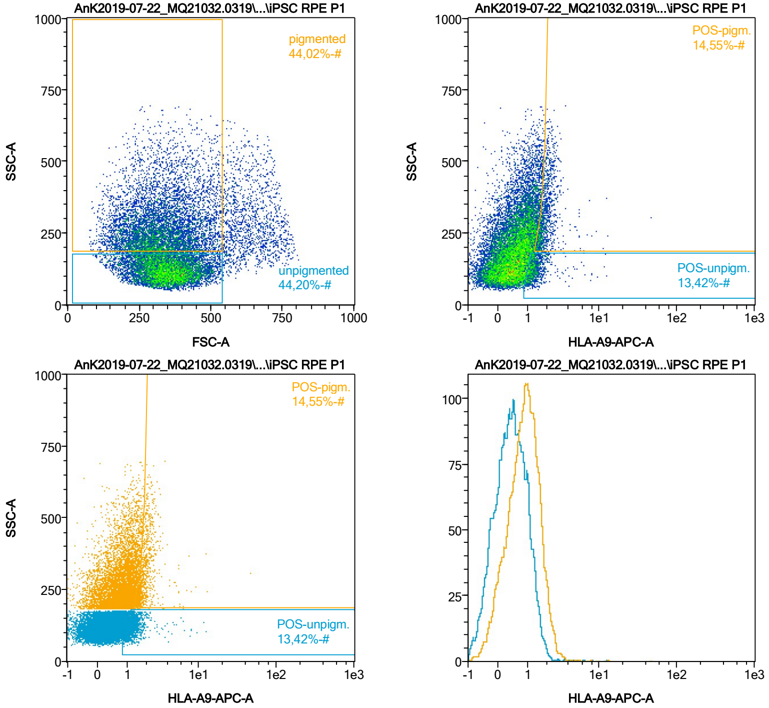


| Name | %-# | HLA-A9-APC-A Mean | HLA-A9-APC-A Median |
|---|---|---|---|
| POS-unpigm. | 13,42 | 1,42 | 1,25 |
| POS-pigm. | 14,55 | 2,06 | 1,84 |

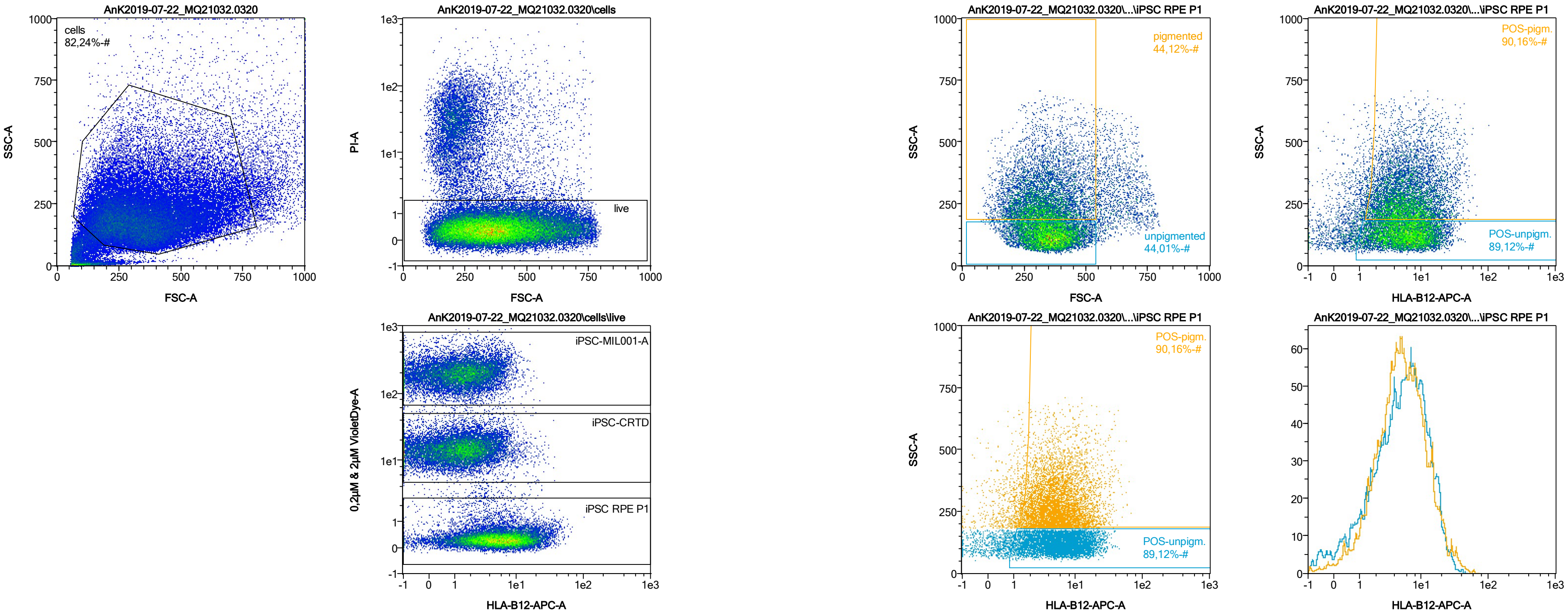


| Name | %-# | HLA-B12-APC-A Mean | HLA-B12-APC-A Median |
|---|---|---|---|
| POS-unpigm. | 89,12 | 7,36 | 5,96 |
| POS-pigm. | 90,16 | 7,41 | 5,46 |

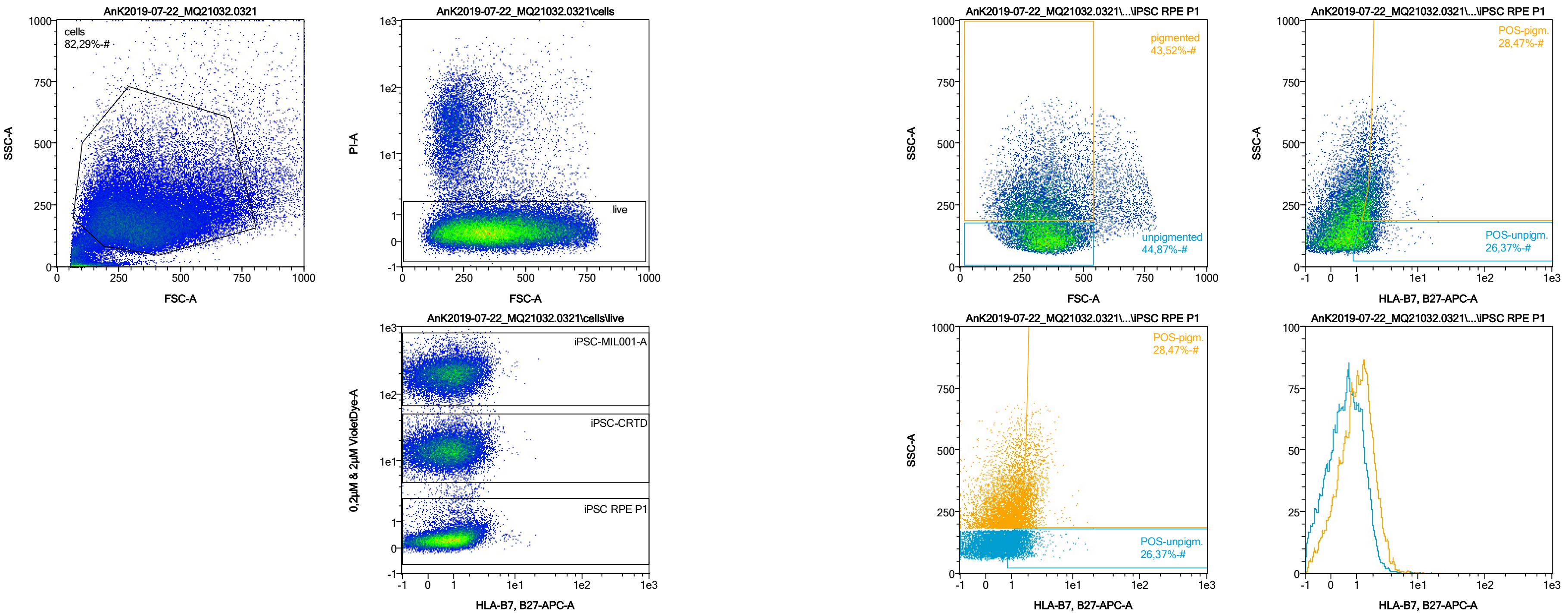


| Name | %-# | HLA-B7, B27-APC-A Mean | HLA-B7, B27-APC-A Median |
|---|---|---|---|
| POS-unpigm. | 26,37 | 1,60 | 1,40 |
| POS-pigm. | 28,47 | 2,25 | 2,01 |

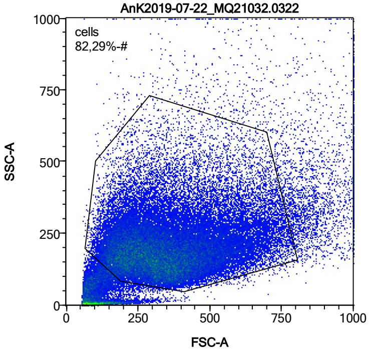


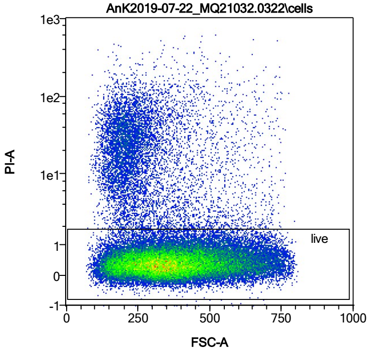


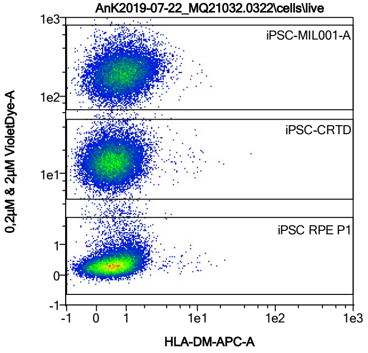


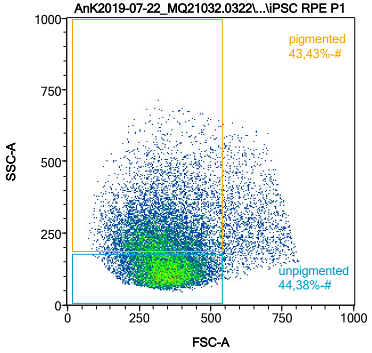


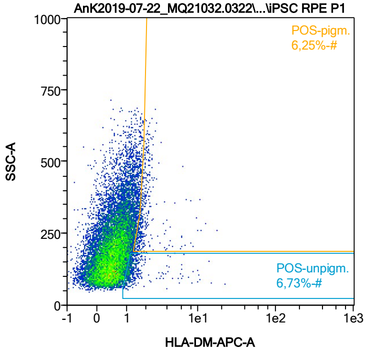


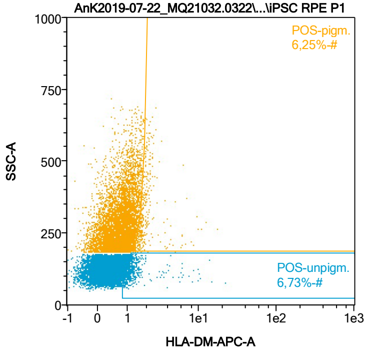


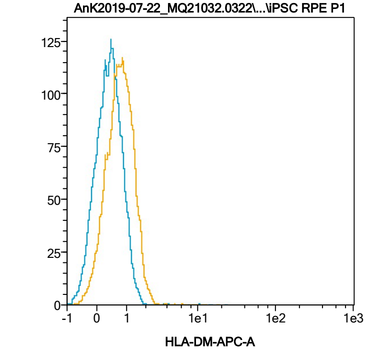


| Name | %-# | HLA-DM-APC-A Mean | HLA-DM-APC-A Median |
|---|---|---|---|
| POS-unpigm. | 6,73 | 1,60 | 1,19 |
| POS-pigm. | 6,25 | 2,13 | 1,79 |

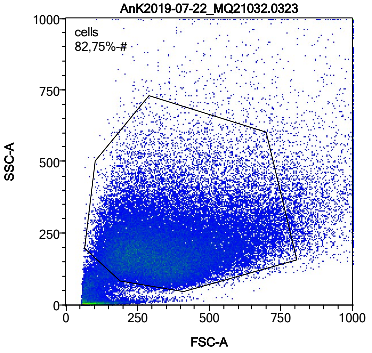


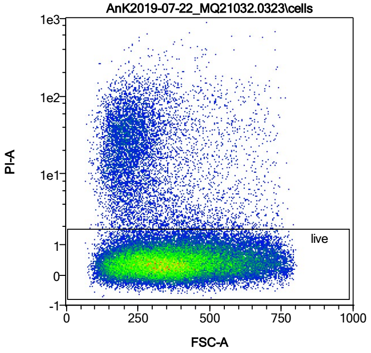


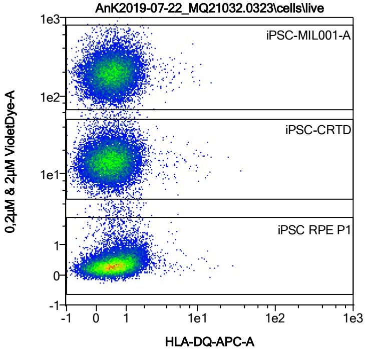


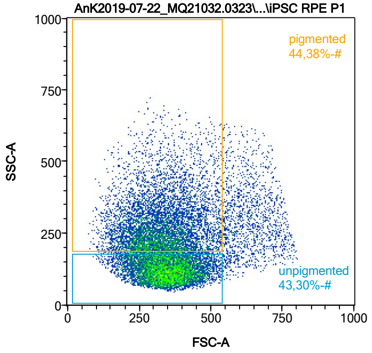


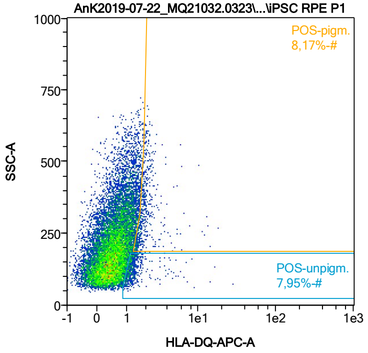


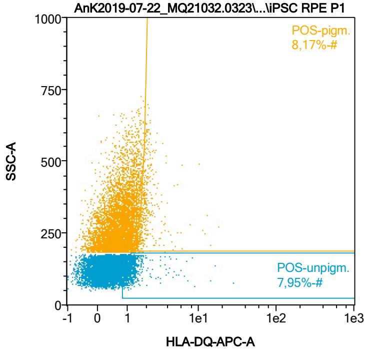


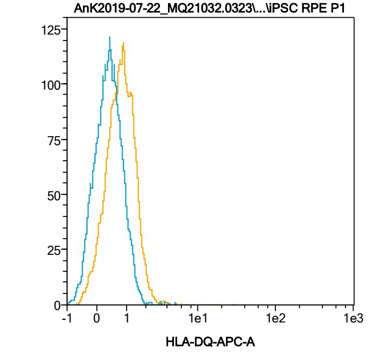


| Name | %-# | HLA-DQ-APC-A Mean | HLA-DQ-APC-A Median |
|---|---|---|---|
| POS-unpigm. | 7,95 | 1,54 | 1,22 |
| POS-pigm. | 8,17 | 2,09 | 1,82 |

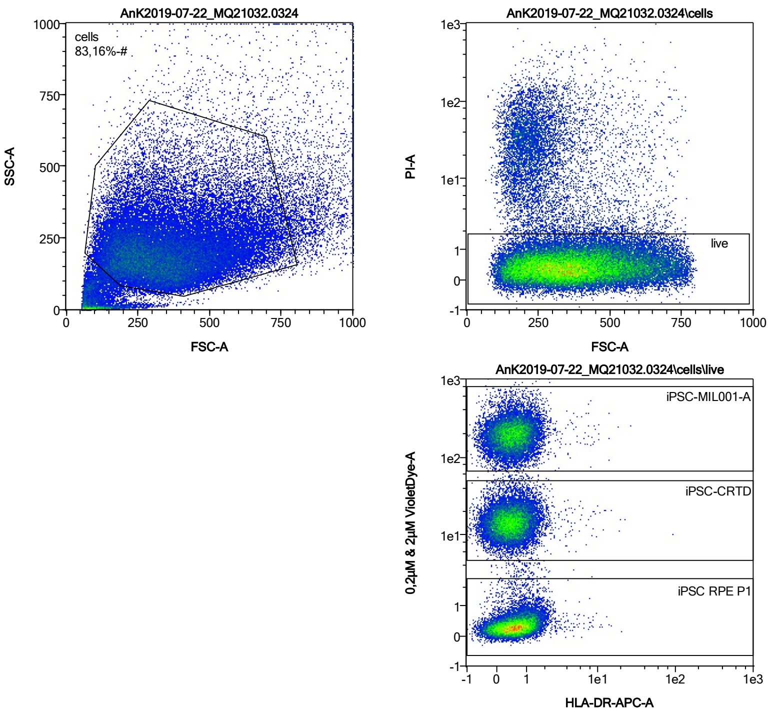


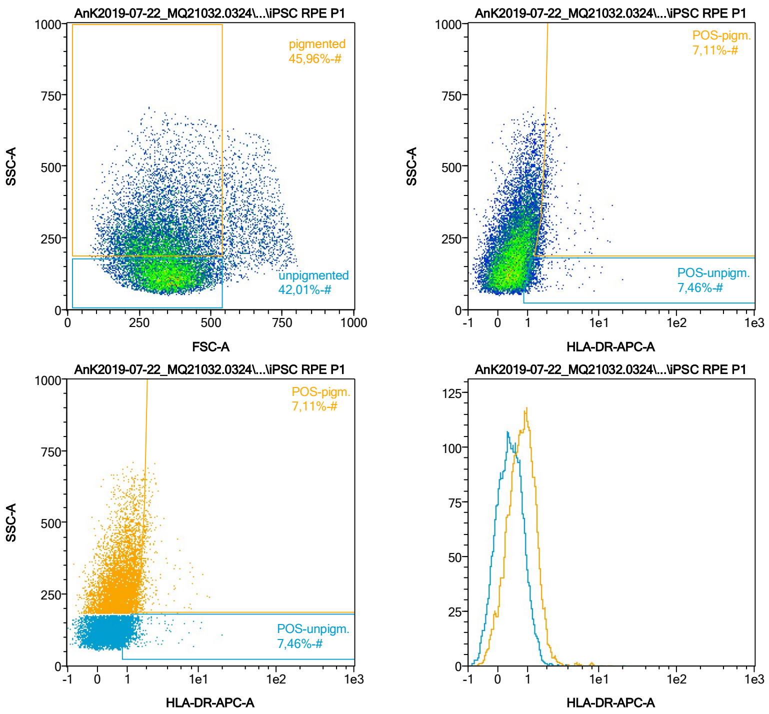


| Name | %-# | HLA-DR-APC-A Mean | HLA-DR-APC-A Median |
|---|---|---|---|
| POS-unpigm. | 7,46 | 1,51 | 1,22 |
| POS-pigm. | 7,11 | 2,16 | 1,84 |

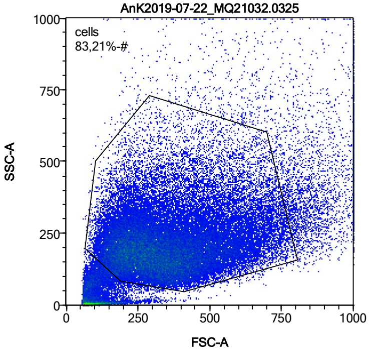


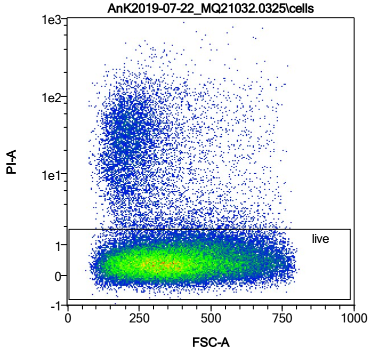


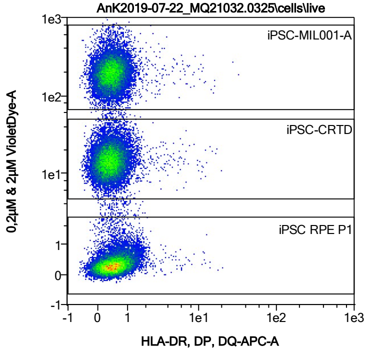


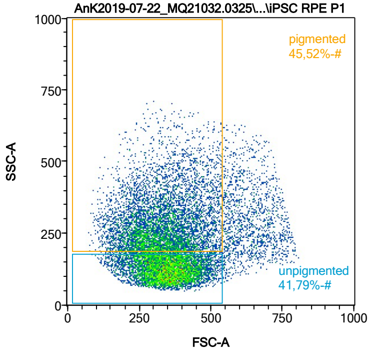


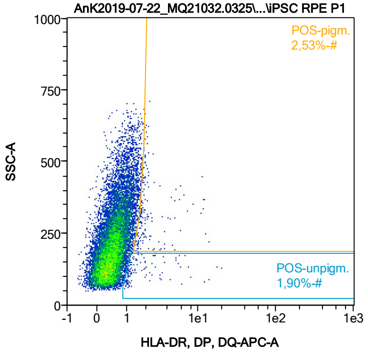


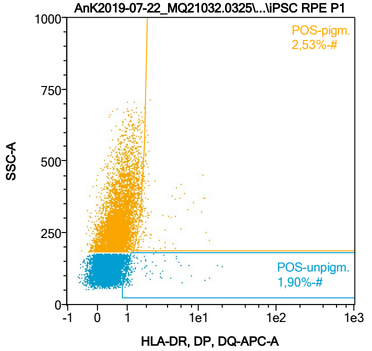


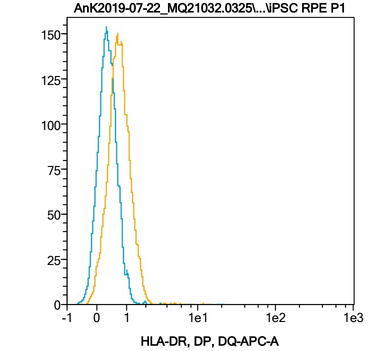


| Name | %-# | HLA-DR, DP, DQ-APC-A Mean | HLA-DR, DP, DQ-APC-A Median |
|---|---|---|---|
| POS-unpigm. | 1,90 | 2,26 | 1,18 |
| POS-pigm. | 2,53 | 2,57 | 1,84 |

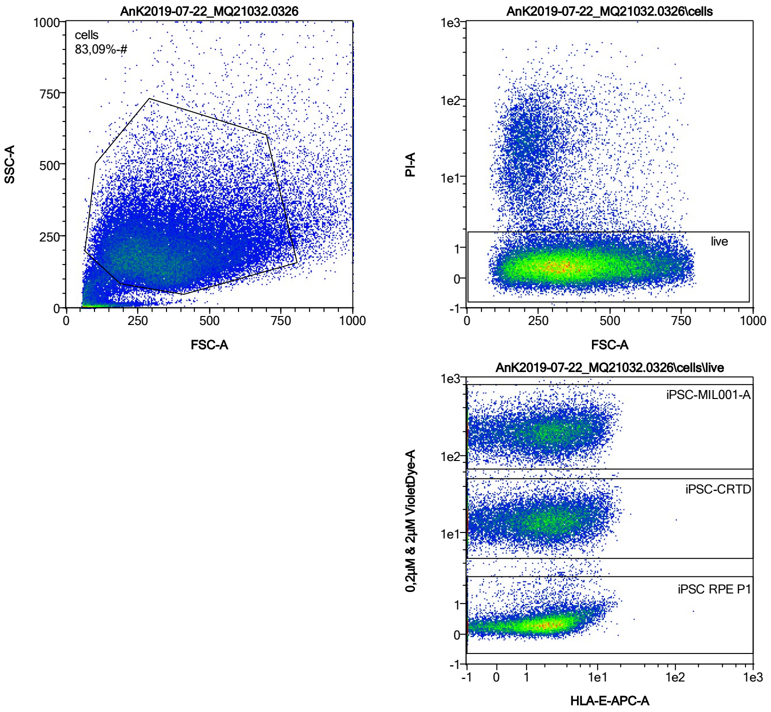


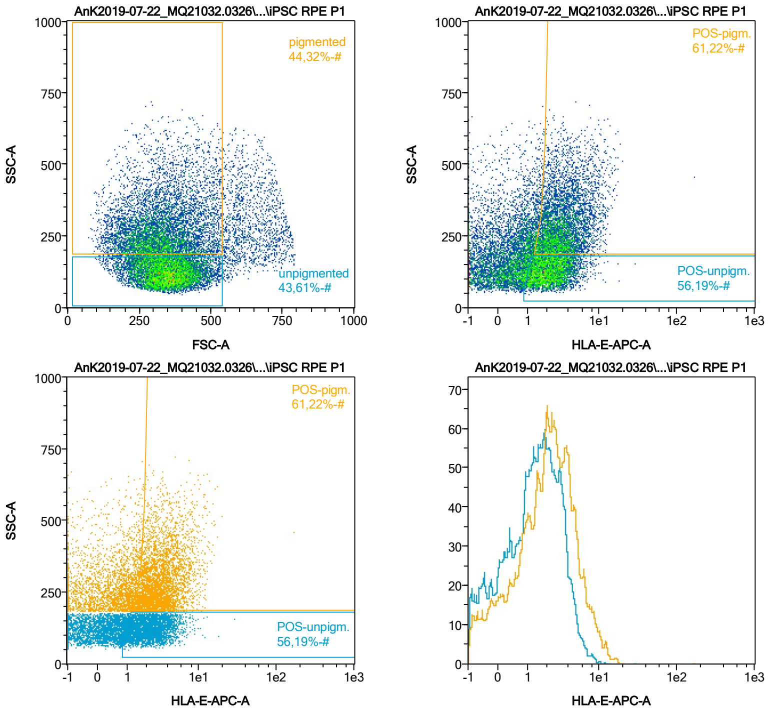


| Name | %-# | HLA-E-APC-A Mean | HLA-E-APC-A Median |
|---|---|---|---|
| POS-unpigm. | 56,19 | 2,48 | 2,15 |
| POS-pigm. | 61,22 | 3,73 | 3,07 |

Date:2024-06-14

Operator: Sebastian Knoebel

Version:2.13.0+54.g95abf0f4

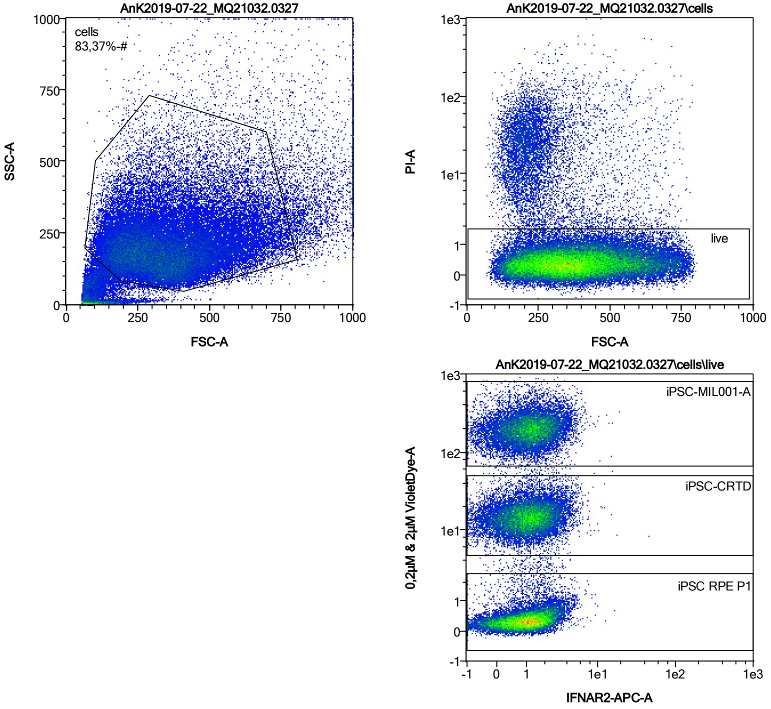


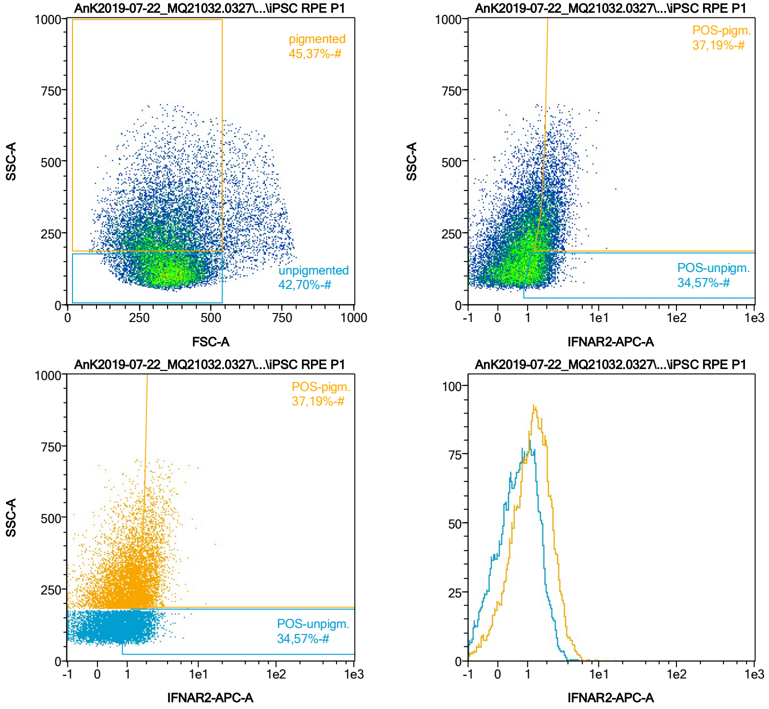


| Name | %-# | IFNAR2-APC-A Mean | IFNAR2-APC-A Median |
|---|---|---|---|
| POS-unpigm. | 34,57 | 1,57 | 1,42 |
| POS-pigm. | 37,19 | 2,30 | 2,08 |

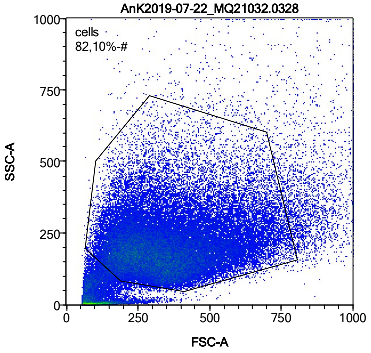


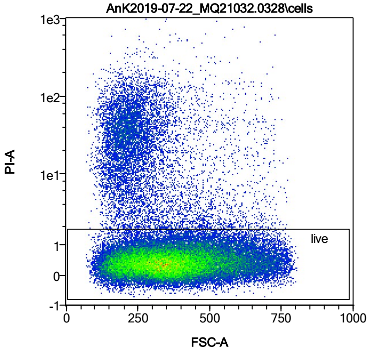


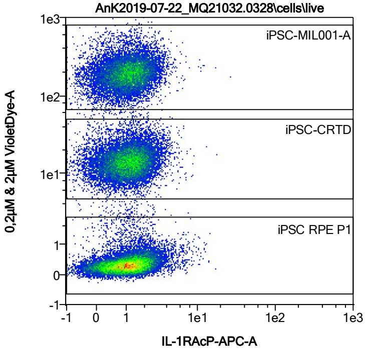


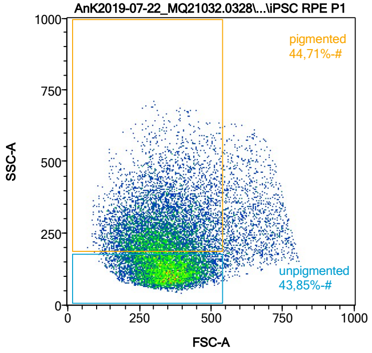


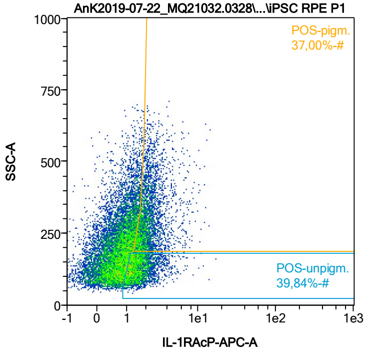


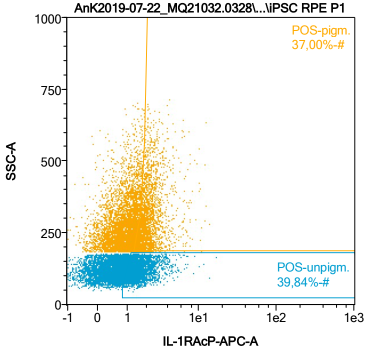


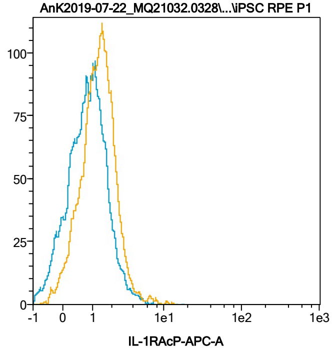


| Name | %-# | IL-1RAcP-APC-A Mean | IL-1RAcP-APC-A Median |
|---|---|---|---|
| POS-unpigm. | 39,84 | 1,70 | 1,47 |
| POS-pigm. | 37,00 | 2,32 | 2,02 |

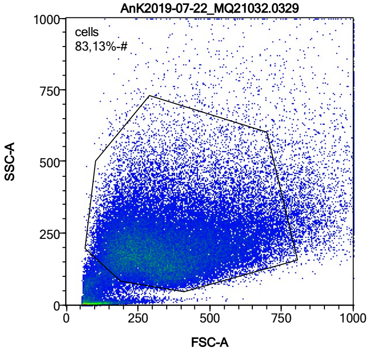


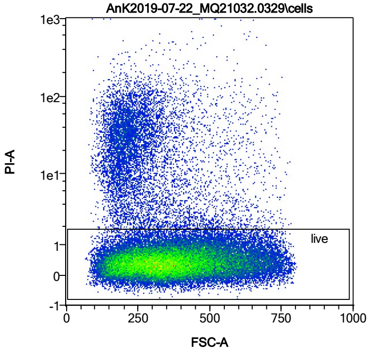


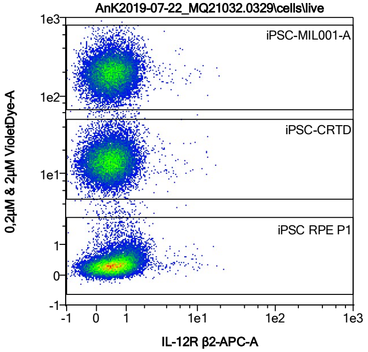


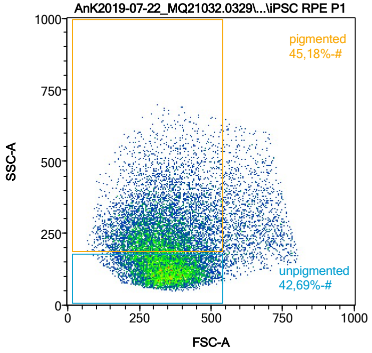


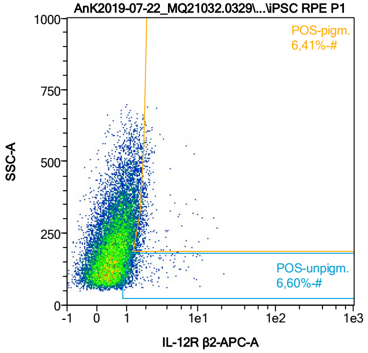


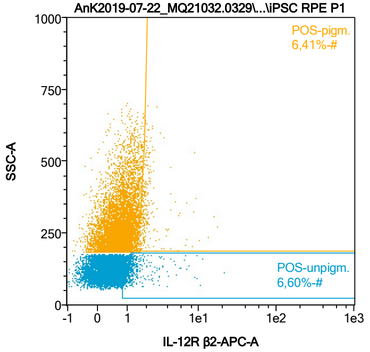


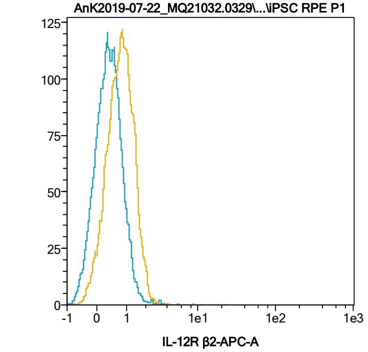


| Name | %-# | IL-12R β2-APC-A Mean | IL-12R β2-APC-A Median |
|---|---|---|---|
| POS-unpigm. | 6,60 | 1,63 | 1,21 |
| POS-pigm. | 6,41 | 2,13 | 1,81 |

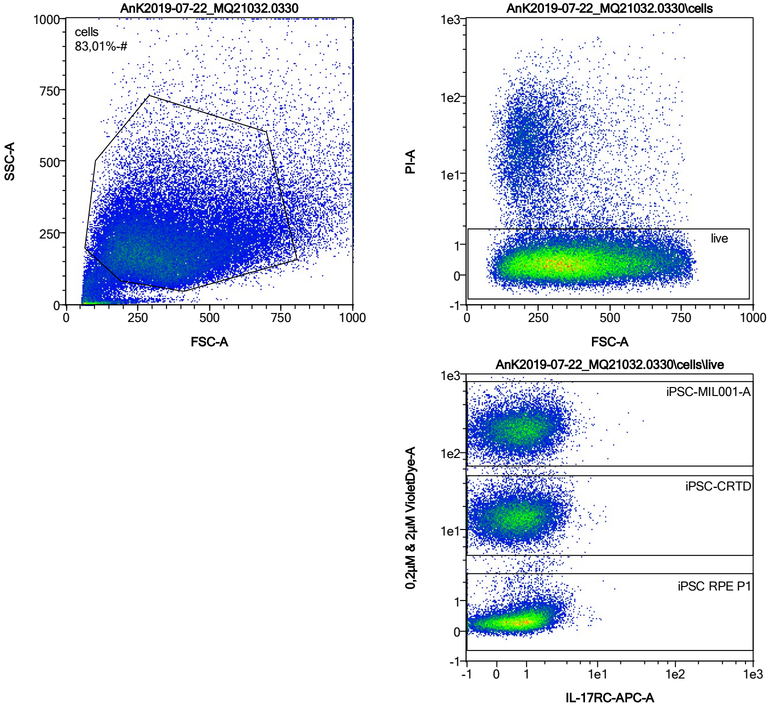


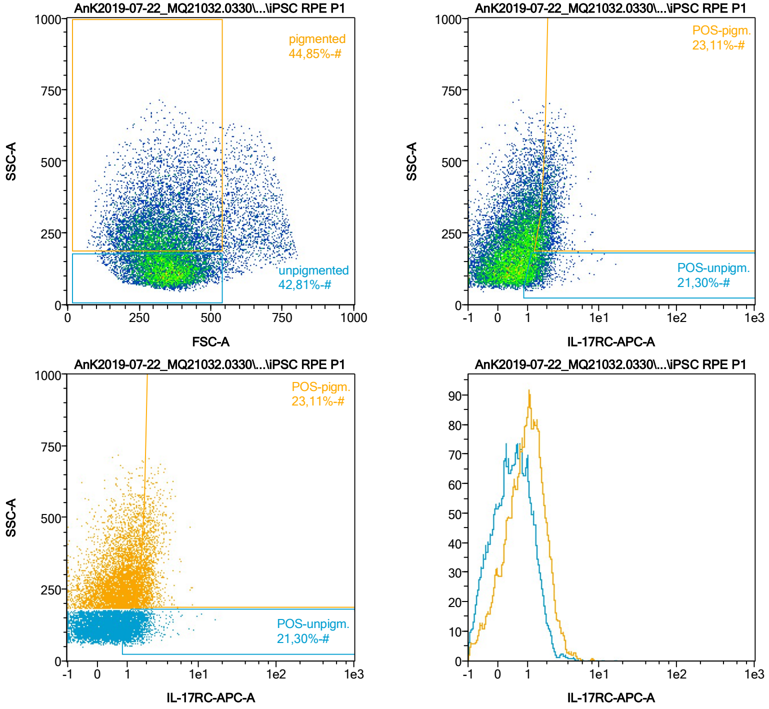


| Name | %-# | IL-17RC-APC-A Mean | IL-17RC-APC-A Median |
|---|---|---|---|
| POS-unpigm. | 21,30 | 1,53 | 1,35 |
| POS-pigm. | 23,11 | 2,16 | 1,99 |

Date:2024-06-14

Operator: Sebastian Knoebel

Version:2.13.0+54.g95abf0f4

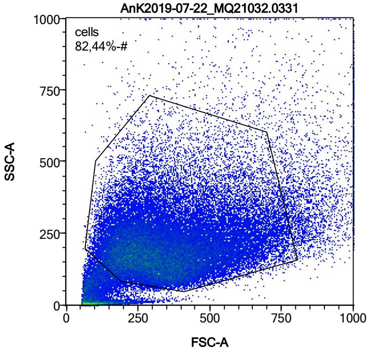


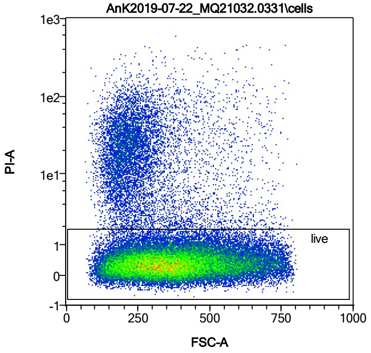


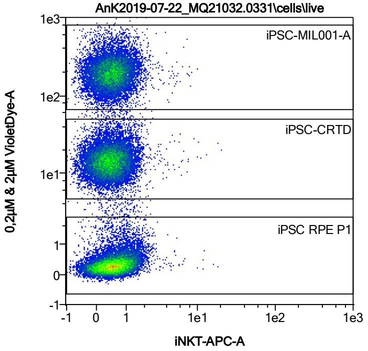


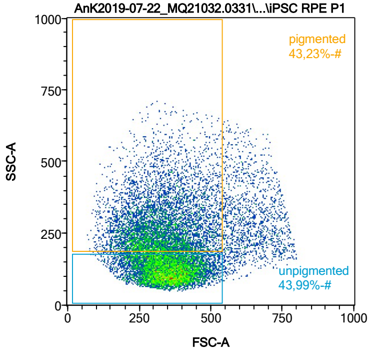


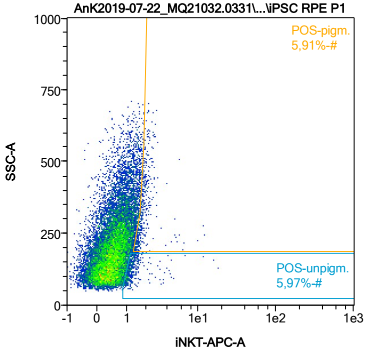


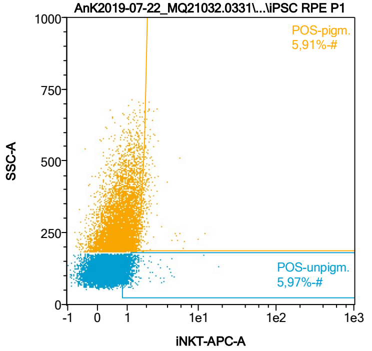


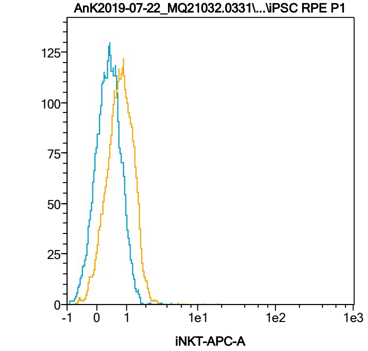


| Name | %-# | iNKT-APC-A Mean | iNKT-APC-A Median |
|---|---|---|---|
| POS-unpigm. | 5,97 | 1,36 | 1,17 |
| POS-pigm. | 5,91 | 1,94 | 1,77 |

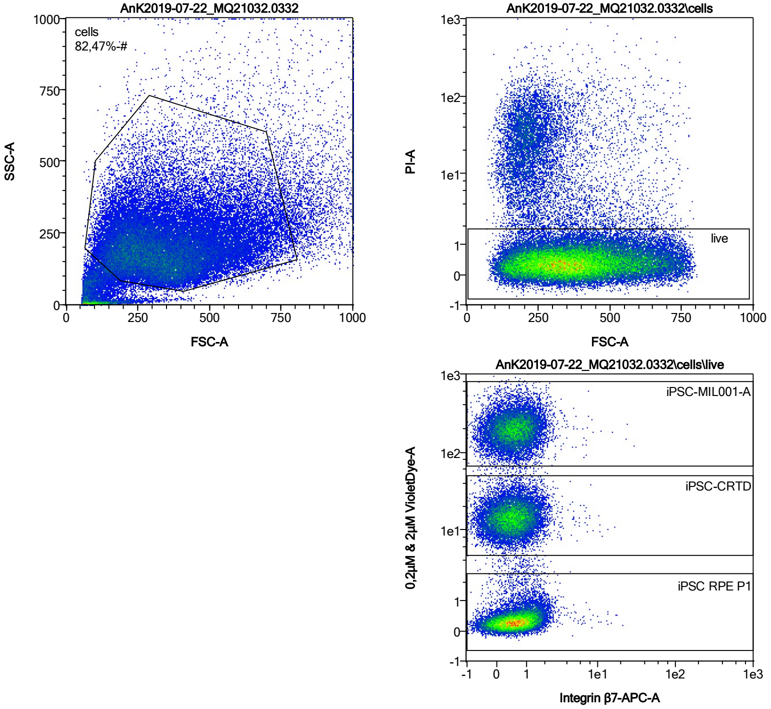


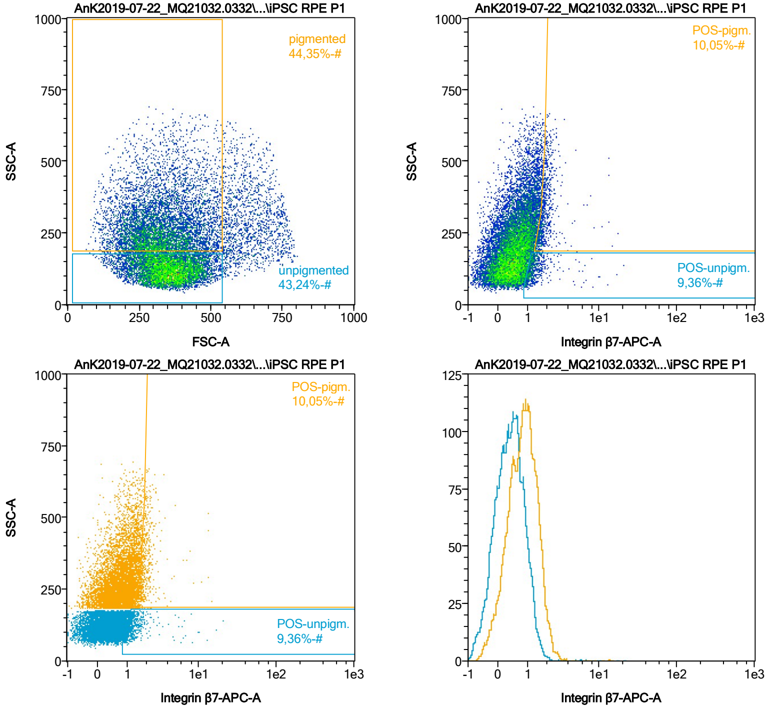


| Name | %-# | Integrin β7-APC-A Mean | Integrin β7-APC-A Median |
|---|---|---|---|
| POS-unpigm. | 9,36 | 1,49 | 1,21 |
| POS-pigm. | 10,05 | 1,99 | 1,79 |

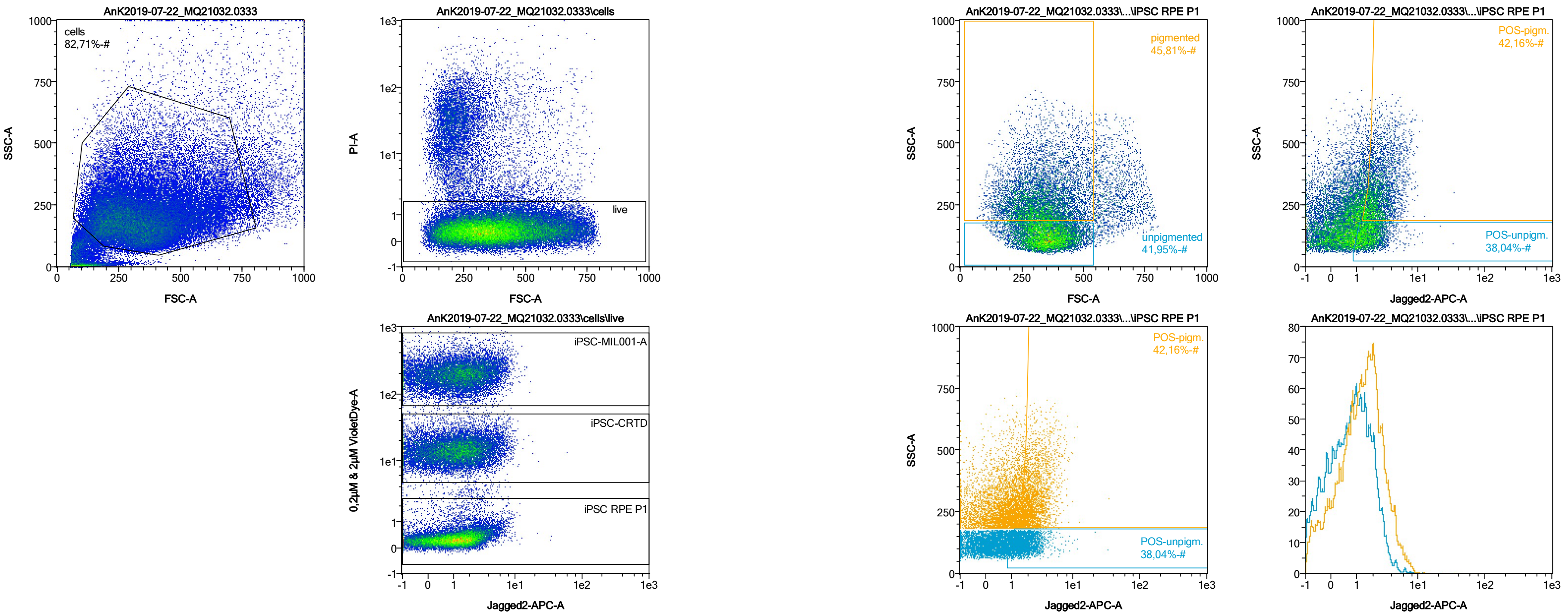


| Name | %-# | Jagged2-APC-A Mean | Jagged2-APC-A Median |
|---|---|---|---|
| POS-unpigm. | 38,04 | 1,91 | 1,67 |
| POS-pigm. | 42,16 | 2,71 | 2,34 |

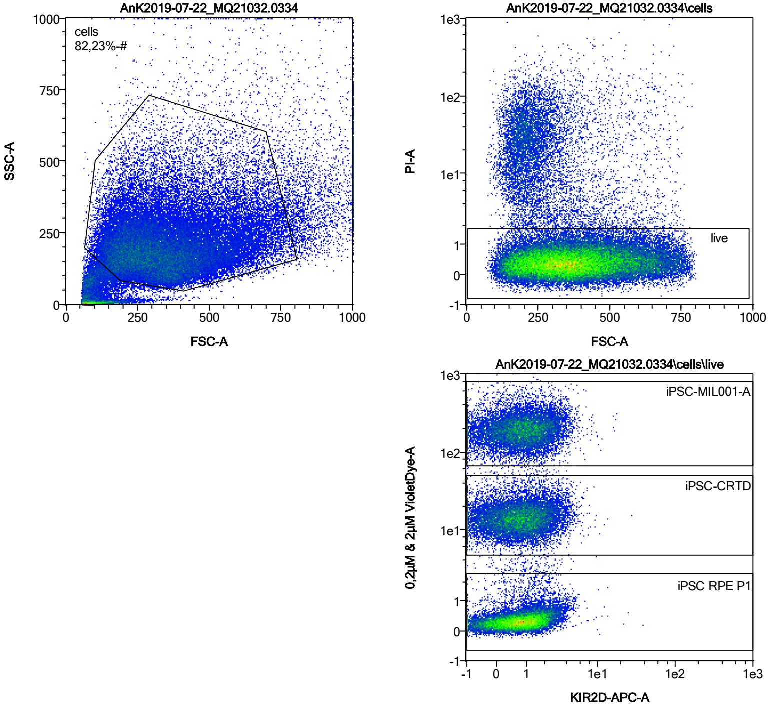


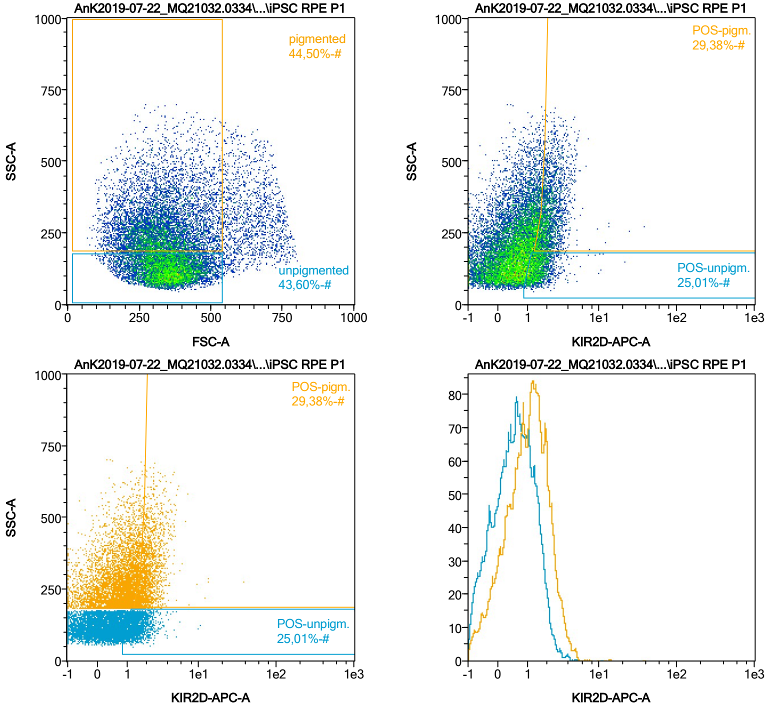


| Name | %-# | KIR2D-APC-A Mean | KIR2D-APC-A Median |
|---|---|---|---|
| POS-unpigm. | 25,01 | 1,55 | 1,42 |
| POS-pigm. | 29,38 | 2,27 | 2,05 |

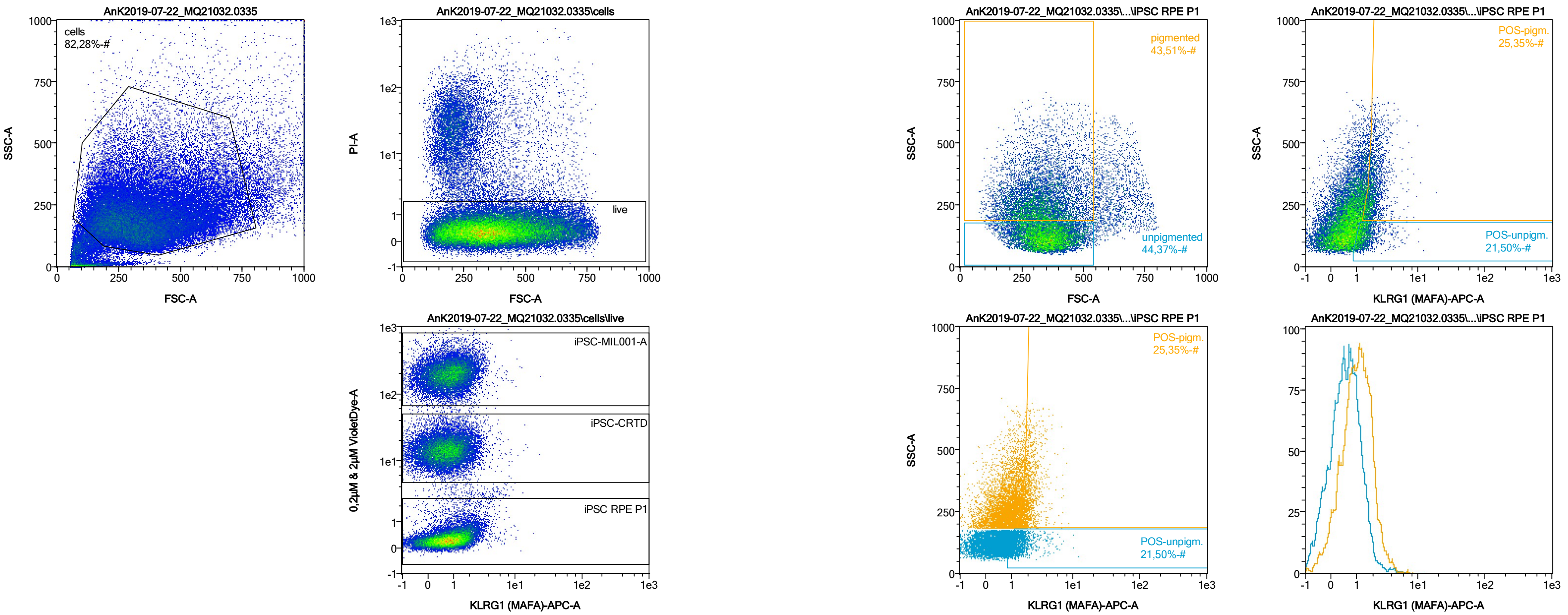


| Name | %-# | KLRG1 (MAFA)-APC-A Mean | KLRG1 (MAFA)-APC-A Median |
|---|---|---|---|
| POS-unpigm. | 21,50 | 1,54 | 1,33 |
| POS-pigm. | 25,35 | 2,12 | 1,94 |

Date:2024-06-14

Operator: Sebastian Knoebel

Version:2.13.0+54.g95abf0f4

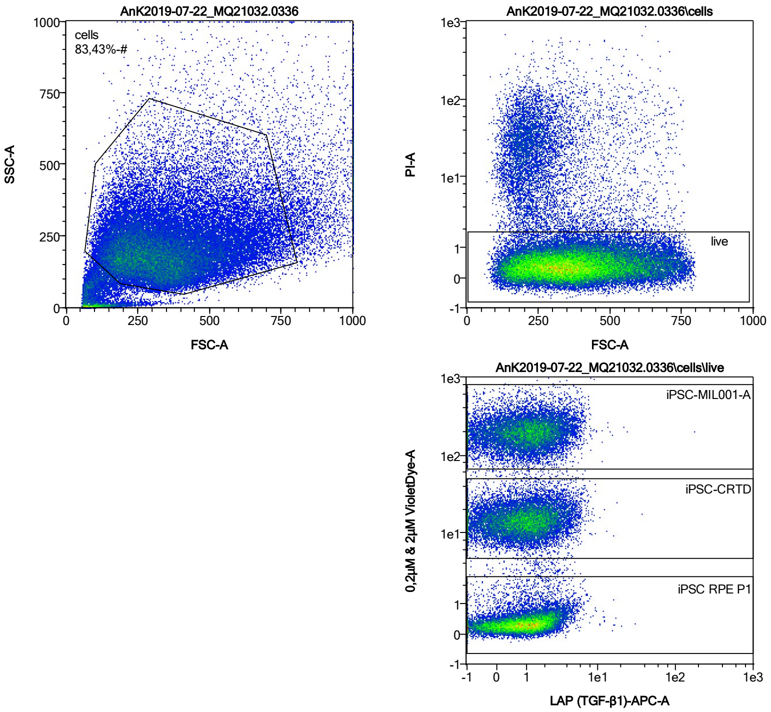


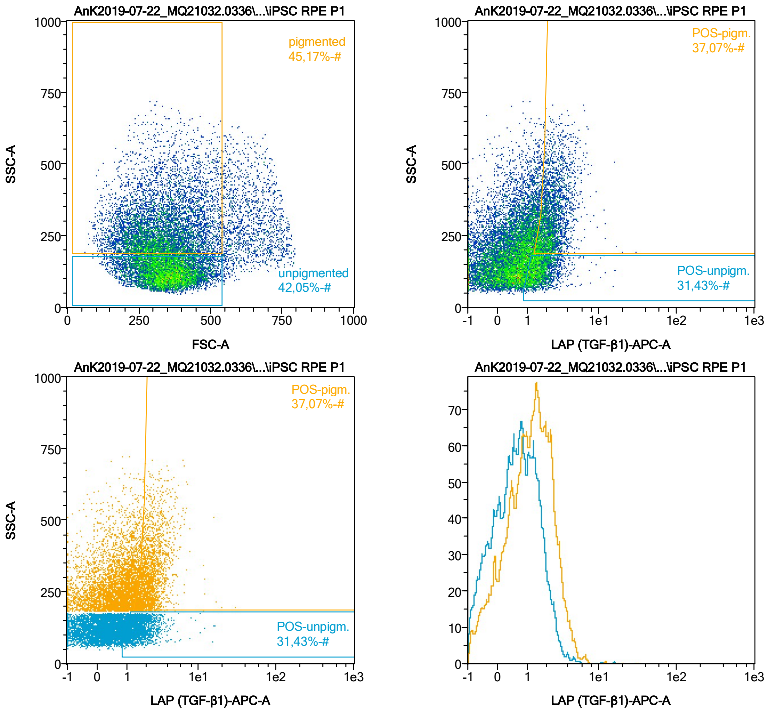


| Name | %-# | LAP (TGF-β1)-APC-A Mean | LAP (TGF-β1)-APC-A Median |
|---|---|---|---|
| POS-unpigm. | 31,43 | 1,70 | 1,50 |
| POS-pigm. | 37,07 | 2,45 | 2,21 |

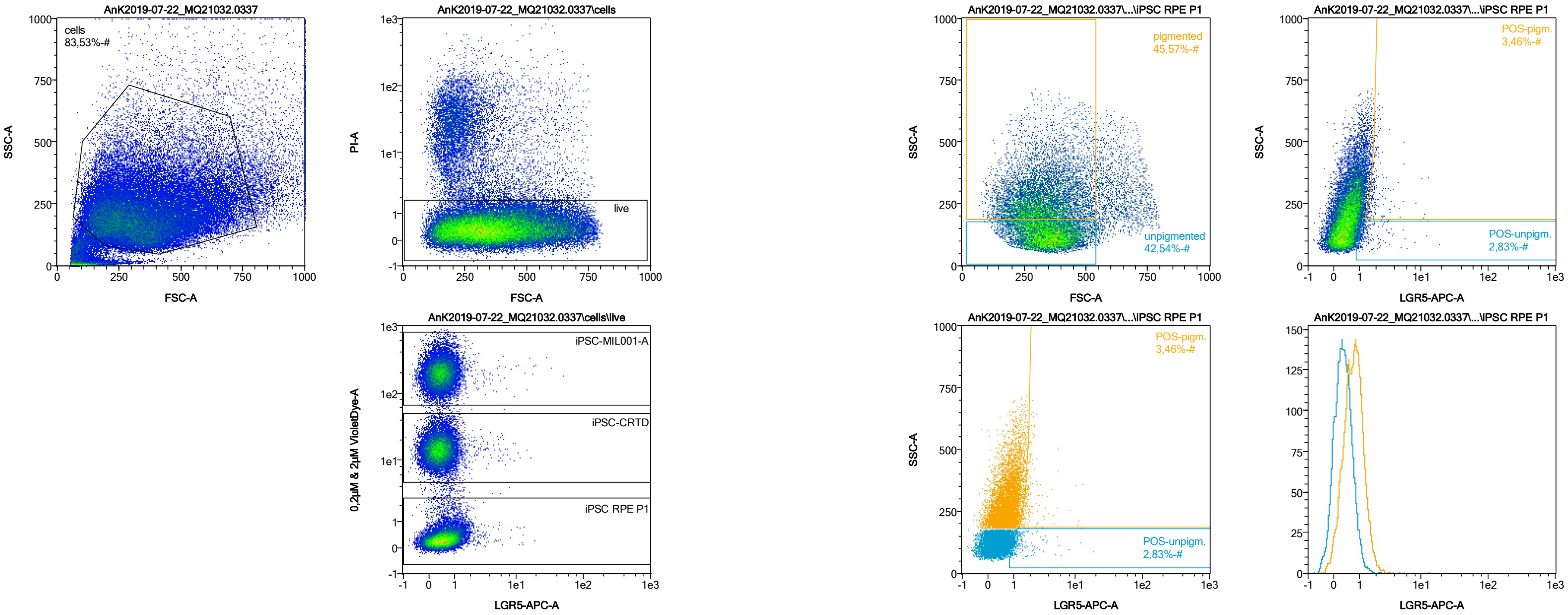


| Name | %-# | LGR5-APC-A Mean | LGR5-APC-A Median |
|---|---|---|---|
| POS-unpigm. | 2,83 | 1,68 | 1,21 |
| POS-pigm. | 3,46 | 2,25 | 1,82 |

Date:2024-06-14
Operator: Sebastian Knoebel
Version:2.13.0+54.g95abf0f4

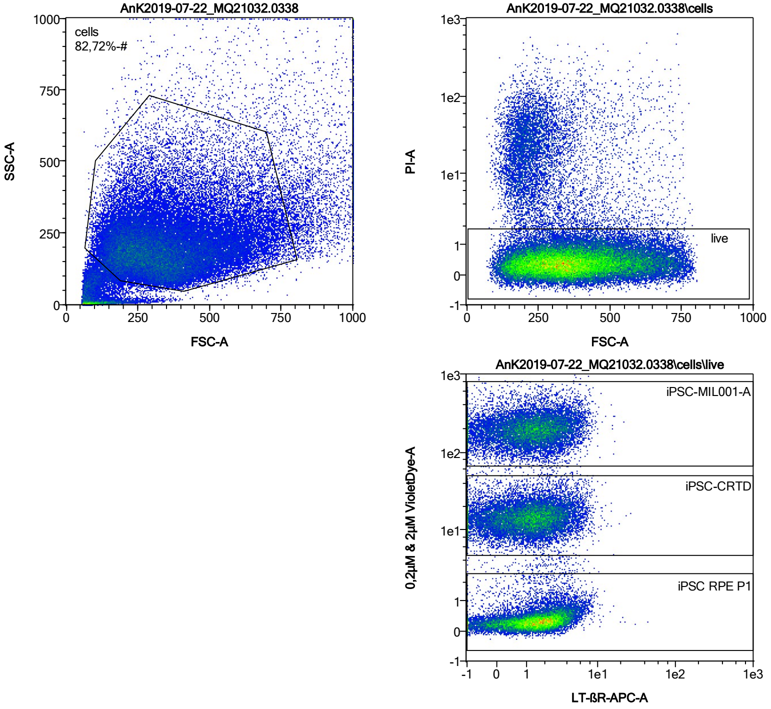


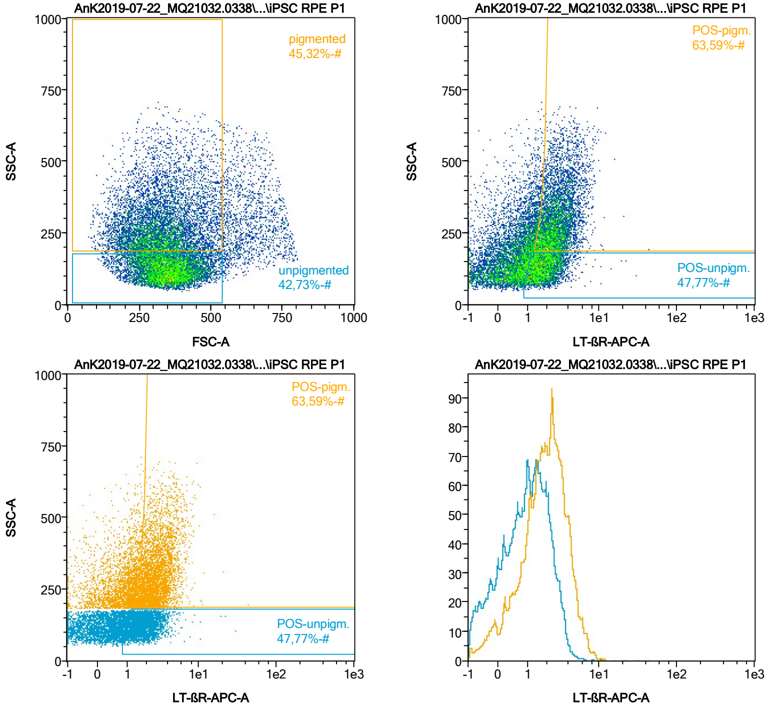


| Name | %-# | LT-ßR-APC-A Mean | LT-ßR-APC-A Median |
|---|---|---|---|
| POS-unpigm. | 47,77 | 1,95 | 1,73 |
| POS-pigm. | 63,59 | 2,97 | 2,63 |

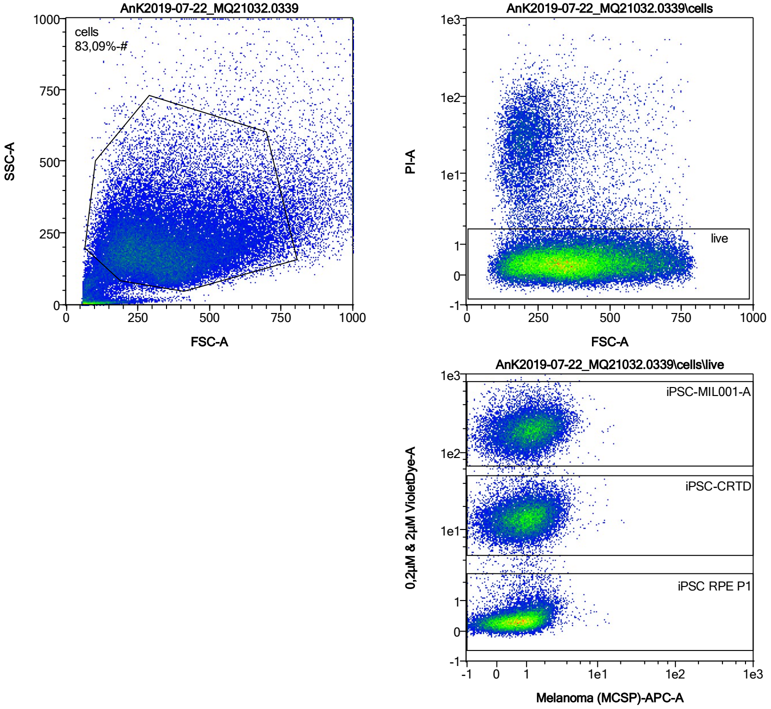


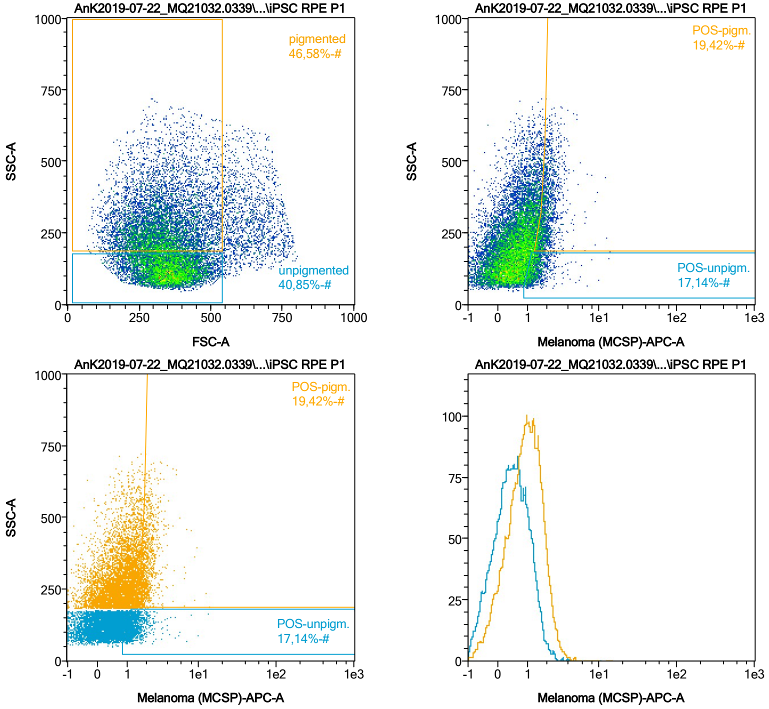


| Name | %-# | Melanoma (MCSP)-APC-A Mean | Melanoma (MCSP)-APC-A Median |
|---|---|---|---|
| POS-unpigm. | 17,14 | 1,44 | 1,29 |
| POS-pigm. | 19,42 | 2,07 | 1,91 |

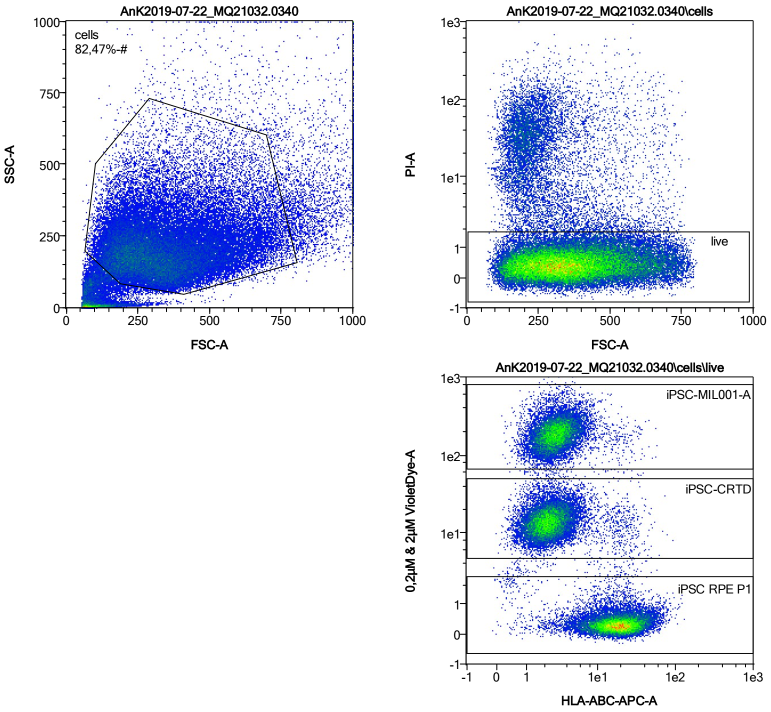


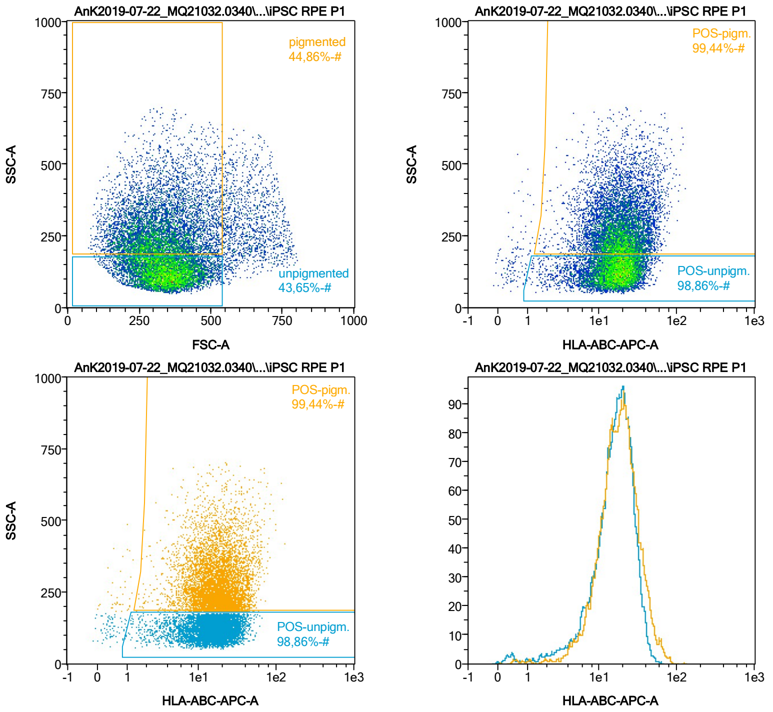


| Name | %-# | HLA-ABC-APC-A Mean | HLA-ABC-APC-A Median |
|---|---|---|---|
| POS-unpigm. | 98,86 | 17,64 | 16,85 |
| POS-pigm. | 99,44 | 20,34 | 18,02 |

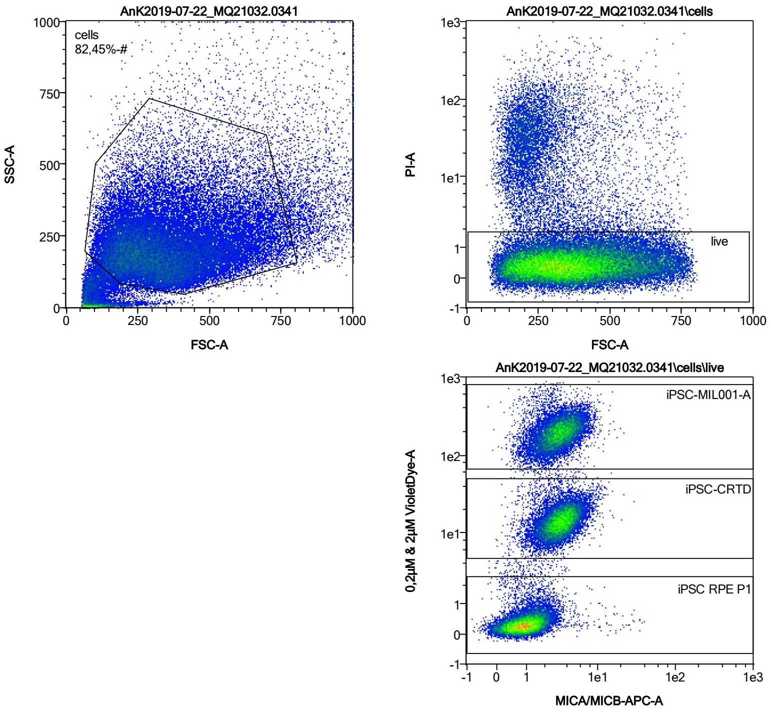


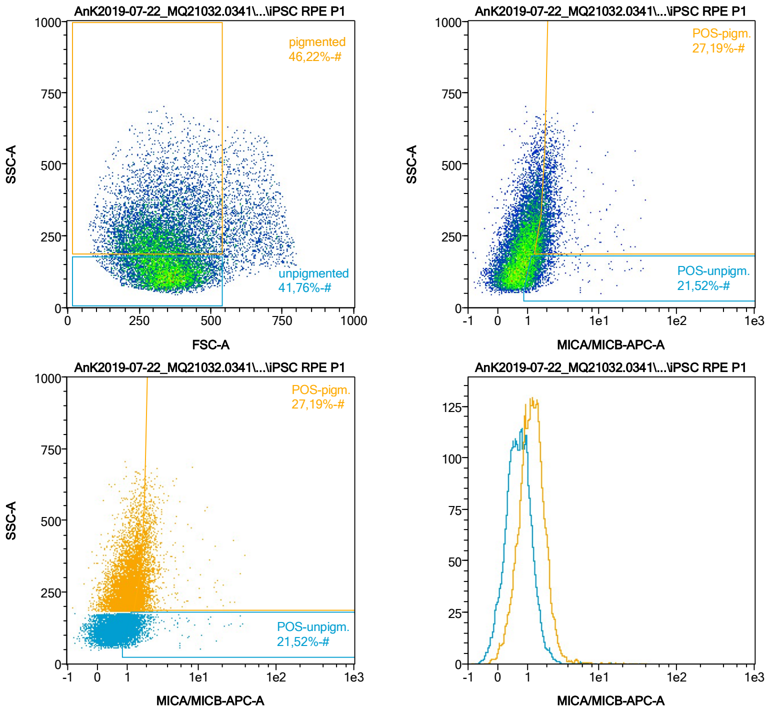


| Name | %-# | MICA/MICB-APC-A Mean | MICA/MICB-APC-A Median |
|---|---|---|---|
| POS-unpigm. | 21,52 | 1,60 | 1,26 |
| POS-pigm. | 27,19 | 2,22 | 1,88 |

Date:2024-06-14
Operator: Sebastian Knoebel
Version:2.13.0+54.g95abf0f4

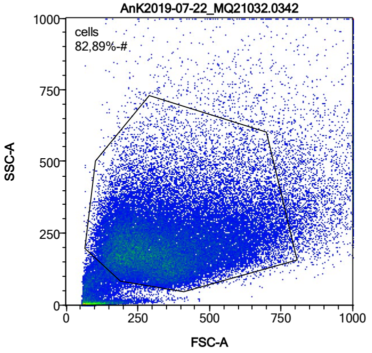


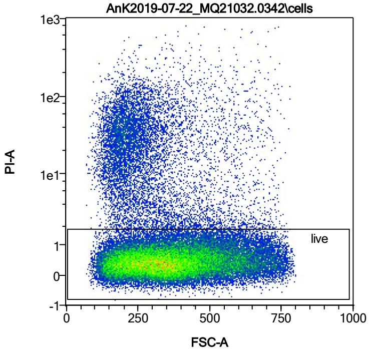


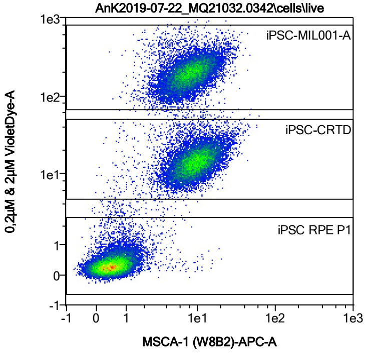


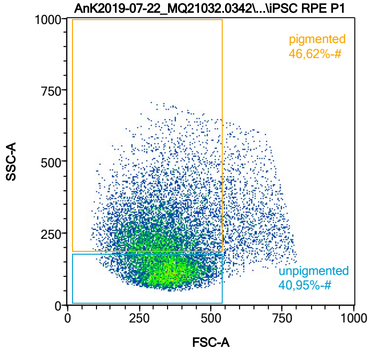


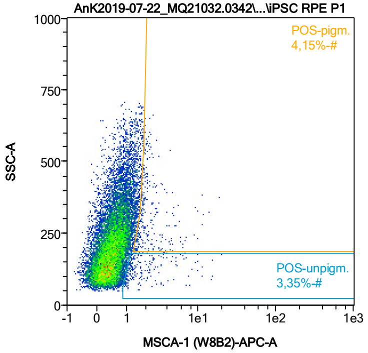


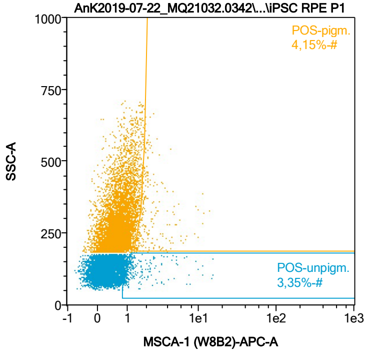


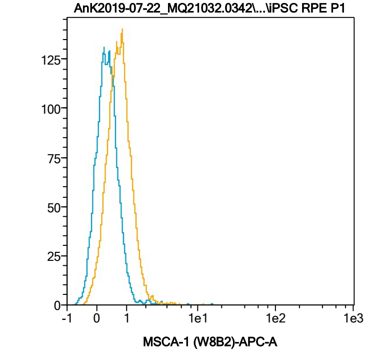


| Name | %-# | MSCA-1 (W8B2)-APC-A Mean | MSCA-1 (W8B2)-APC-A Median |
|---|---|---|---|
| POS-unpigm. | 3,35 | 2,20 | 1,30 |
| POS-pigm. | 4,15 | 2,51 | 1,93 |

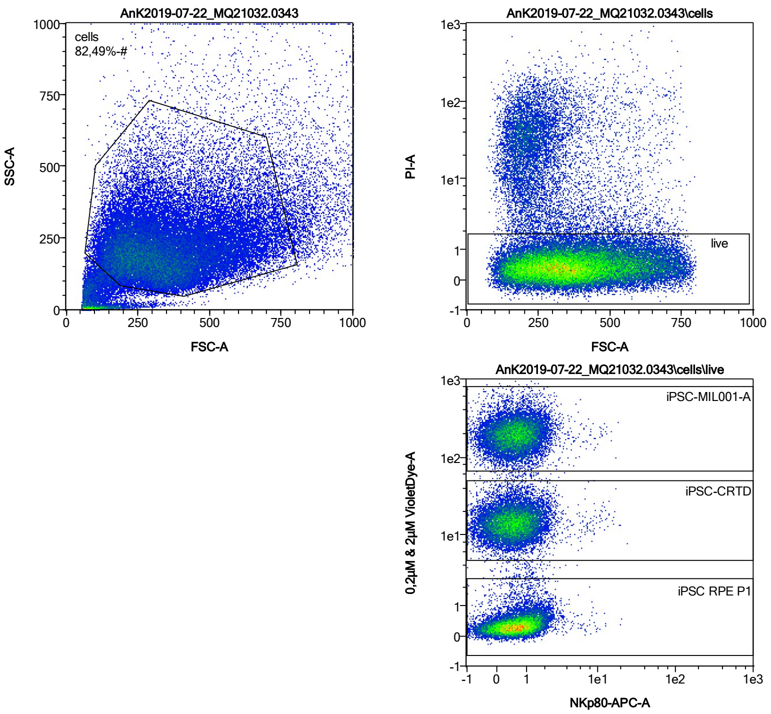


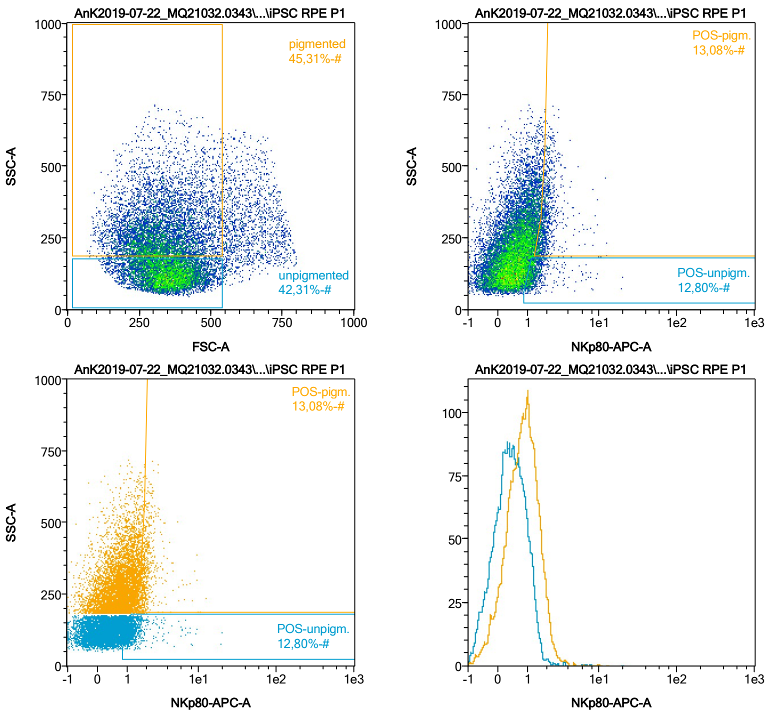


| Name | %-# | NKp80-APC-A Mean | NKp80-APC-A Median |
|---|---|---|---|
| POS-unpigm. | 12,80 | 1,50 | 1,23 |
| POS-pigm. | 13,08 | 2,05 | 1,86 |

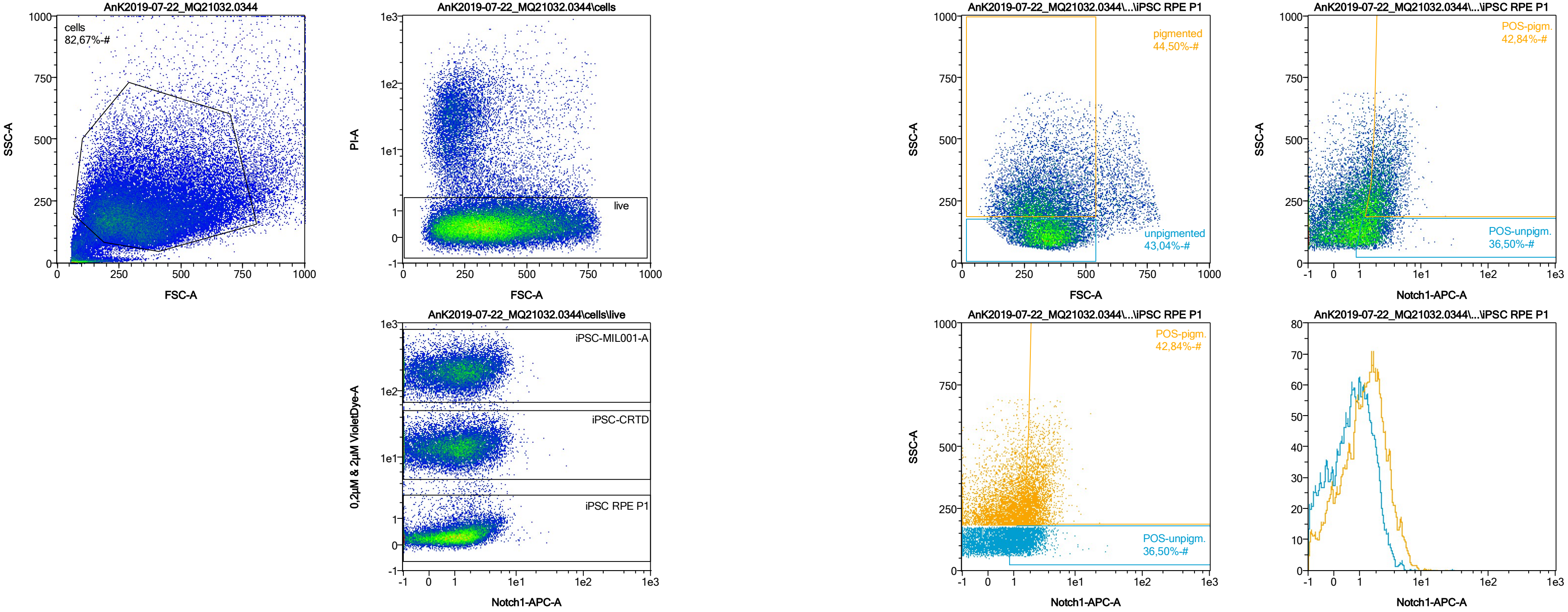


| Name | %-# | Notch1-APC-A Mean | Notch1-APC-A Median |
|---|---|---|---|
| POS-unpigm. | 36,50 | 1,85 | 1,62 |
| POS-pigm. | 42,84 | 2,72 | 2,35 |

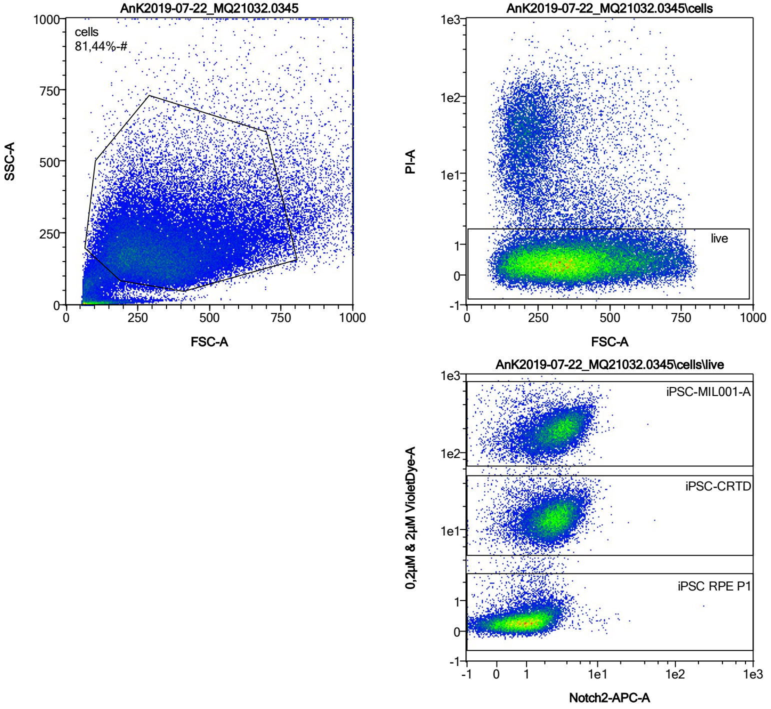


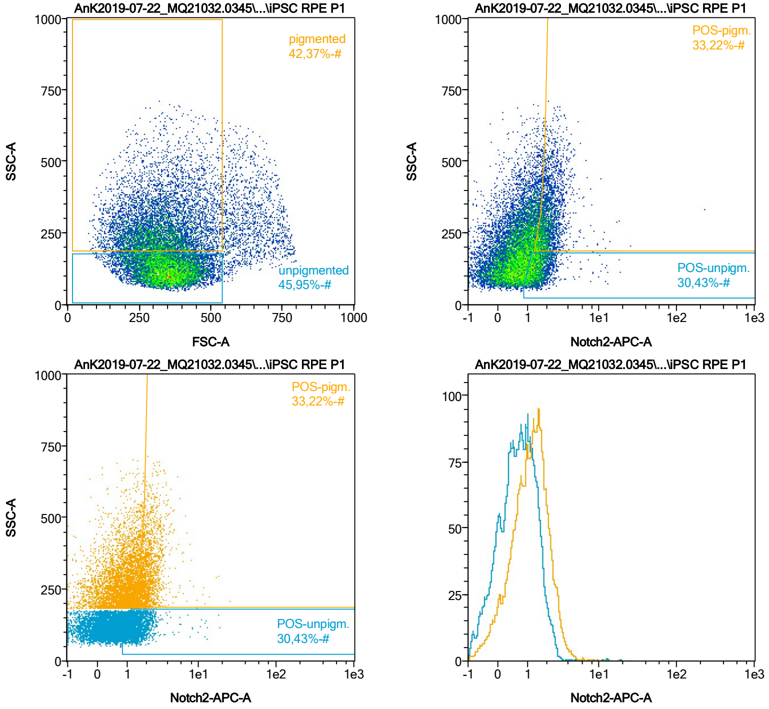


| Name | %-# | Notch2-APC-A Mean | Notch2-APC-A Median |
|---|---|---|---|
| POS-unpigm. | 30,43 | 1,56 | 1,39 |
| POS-pigm. | 33,22 | 2,20 | 1,97 |

Date:2024-06-14

Operator: Sebastian Knoebel

Version:2.13.0+54.g95abf0f4

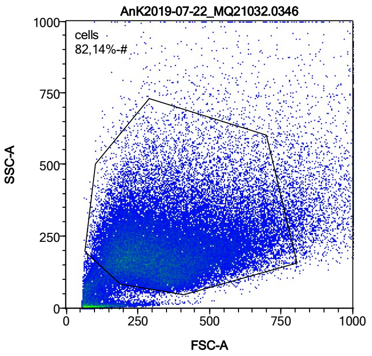


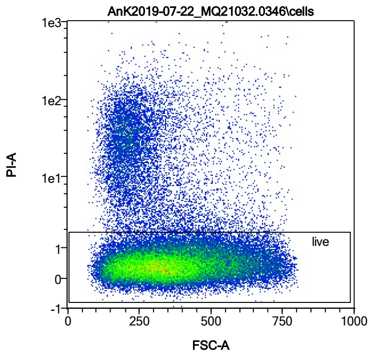


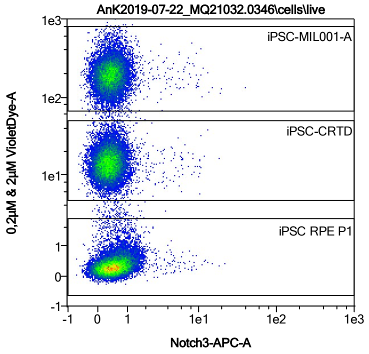


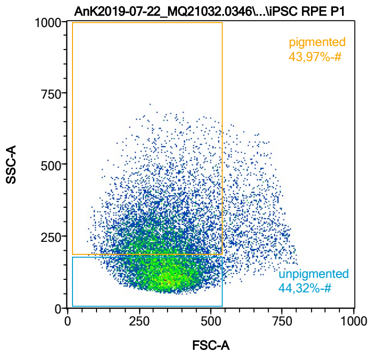


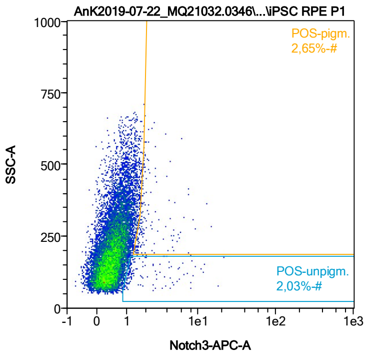


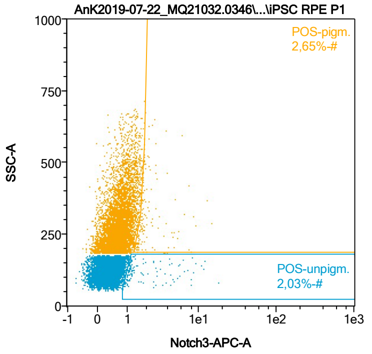


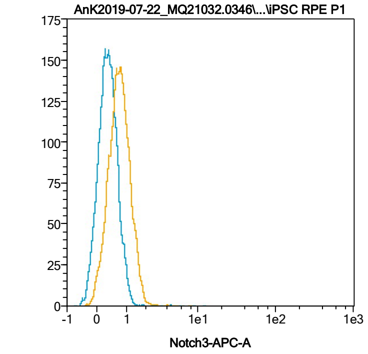


| Name | %-# | Notch3-APC-A Mean | Notch3-APC-A Median |
|---|---|---|---|
| POS-unpigm. | 2,03 | 2,44 | 1,20 |
| POS-pigm. | 2,65 | 2,43 | 1,81 |

Date:2024-06-14

Operator: Sebastian Knoebel

Version:2.13.0+54.g95abf0f4

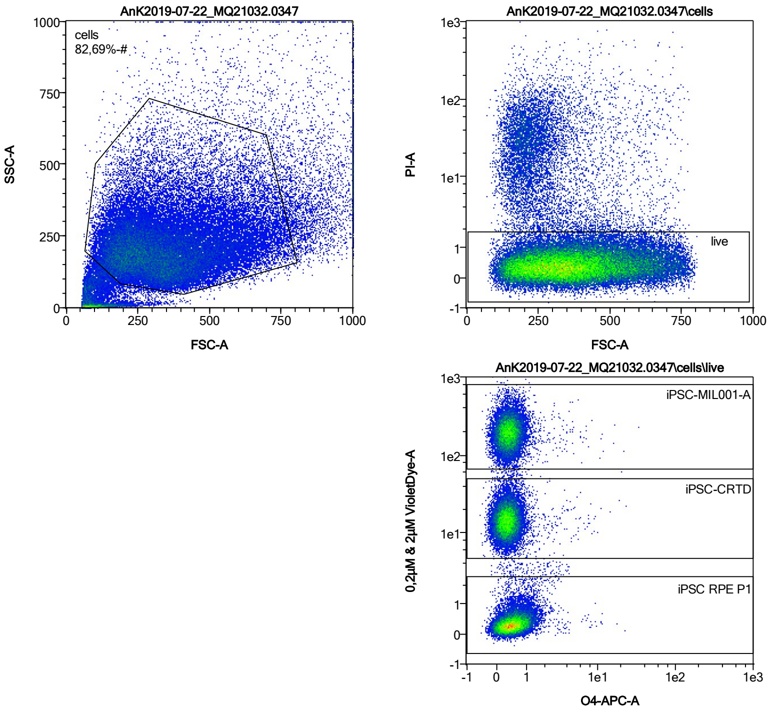


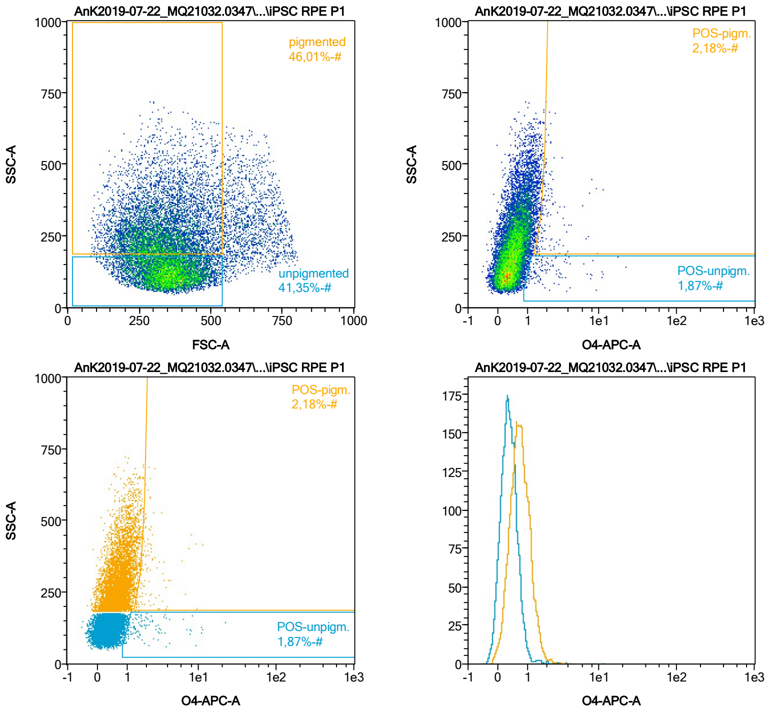


| Name | %-# | O4-APC-A Mean | O4-APC-A Median |
|---|---|---|---|
| POS-unpigm. | 1,87 | 2,64 | 1,61 |
| POS-pigm. | 2,18 | 2,38 | 1,92 |

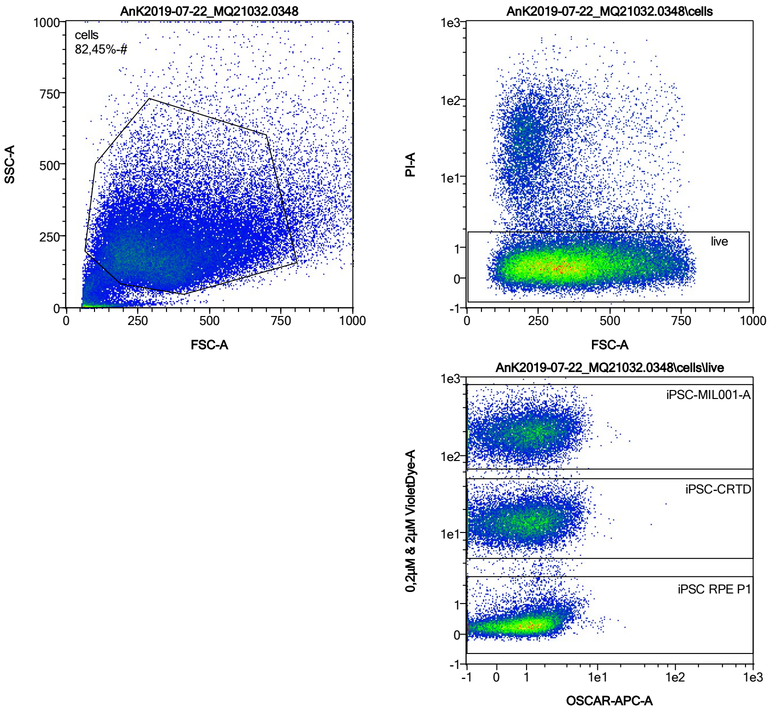


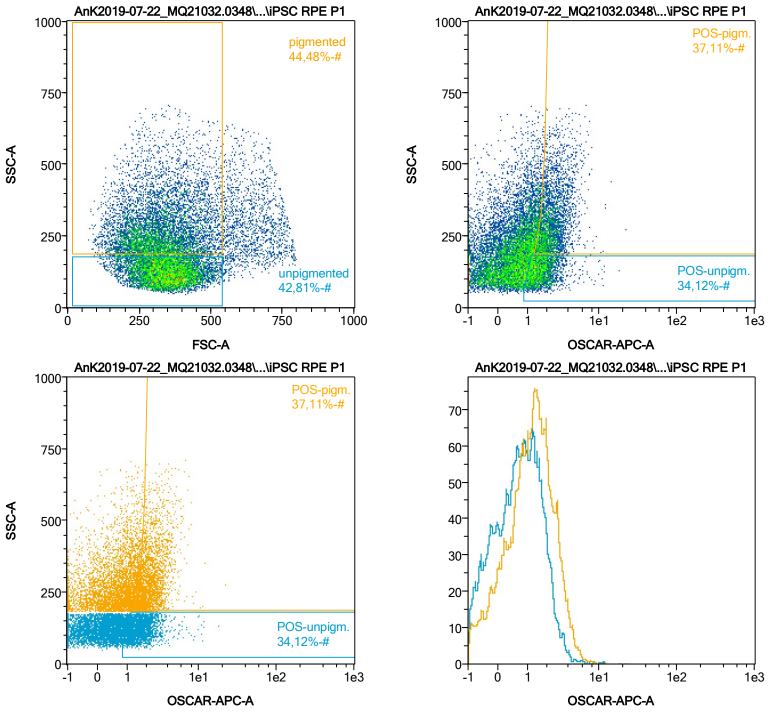


| Name | %-# | OSCAR-APC-A Mean | OSCAR-APC-A Median |
|---|---|---|---|
| POS-unpigm. | 34,12 | 1,80 | 1,57 |
| POS-pigm. | 37,11 | 2,48 | 2,20 |

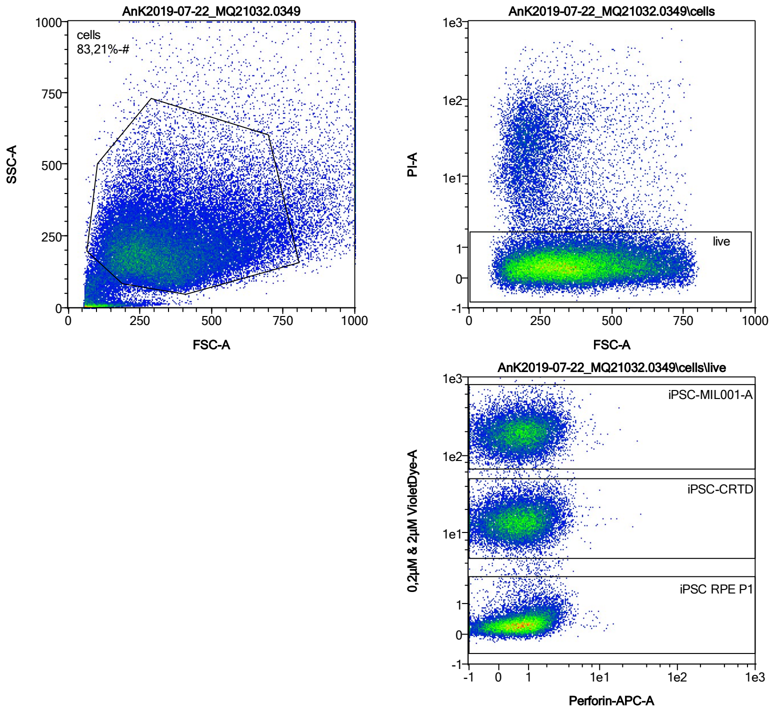


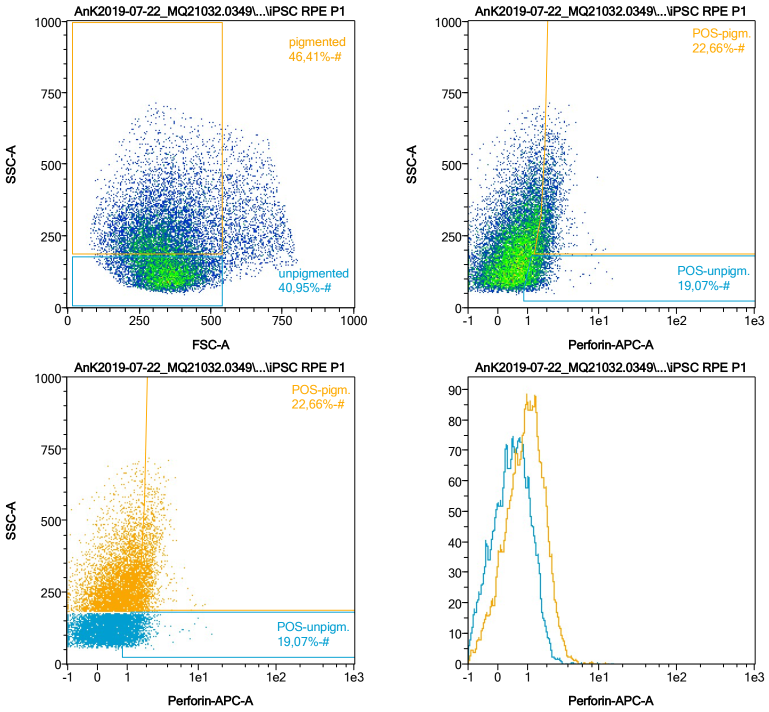


| Name | %-# | Perforin-APC-A Mean | Perforin-APC-A Median |
|---|---|---|---|
| POS-unpigm. | 19,07 | 1,49 | 1,34 |
| POS-pigm. | 22,66 | 2,19 | 2,00 |

Date:2024-06-14

Operator: Sebastian Knoebel

Version:2.13.0+54.g95abf0f4

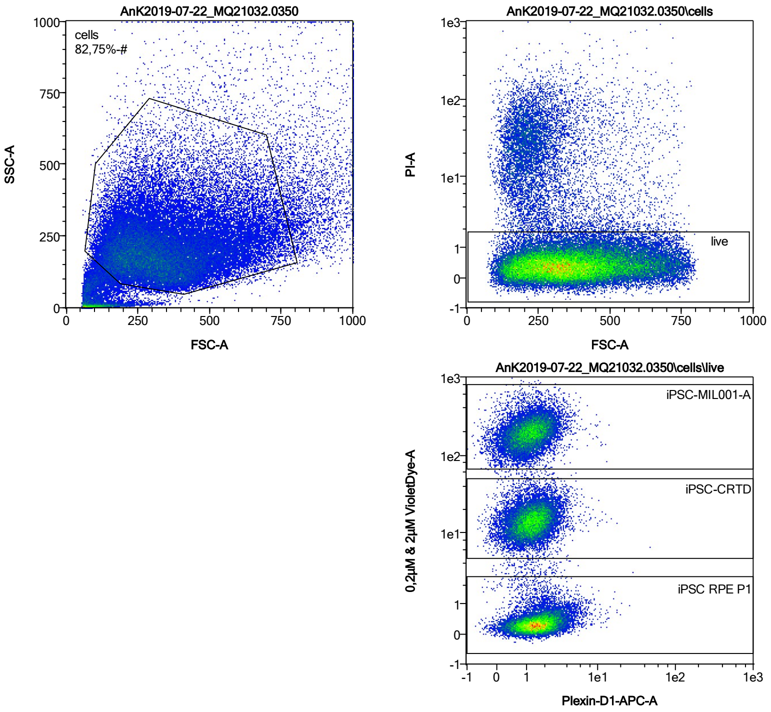


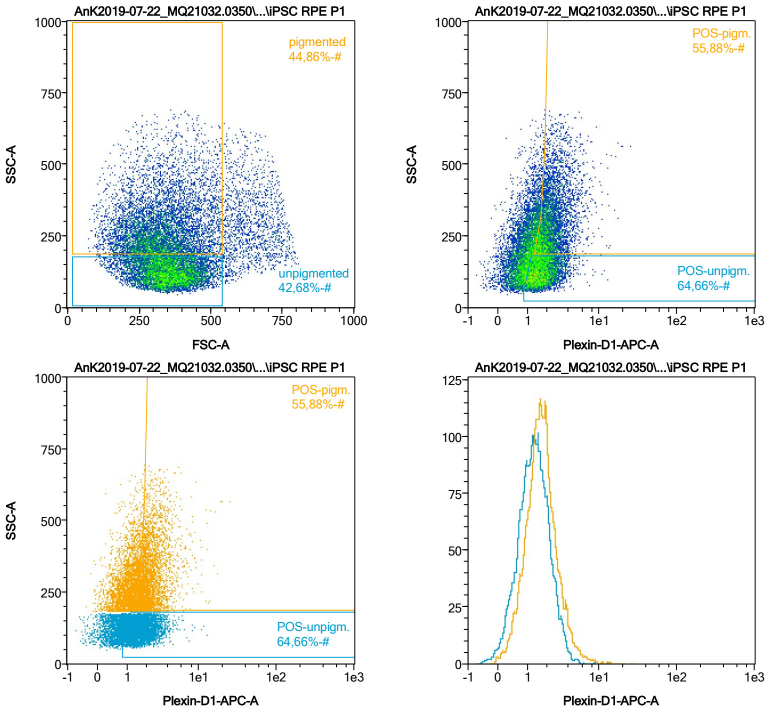


| Name | %-# | Plexin-D1-APC-A Mean | Plexin-D1-APC-A Median |
|---|---|---|---|
| POS-unpigm. | 64,66 | 1,81 | 1,62 |
| POS-pigm. | 55,88 | 2,46 | 2,12 |

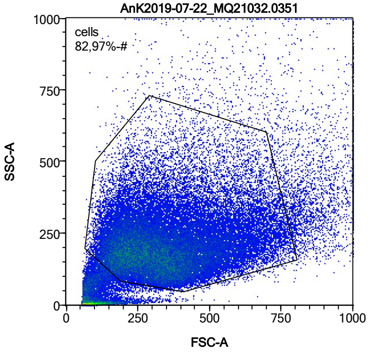


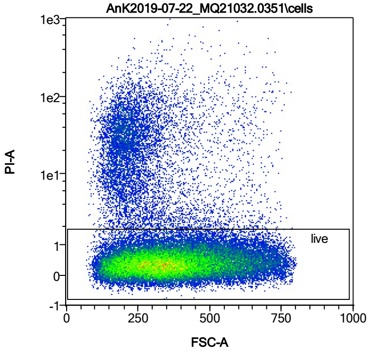


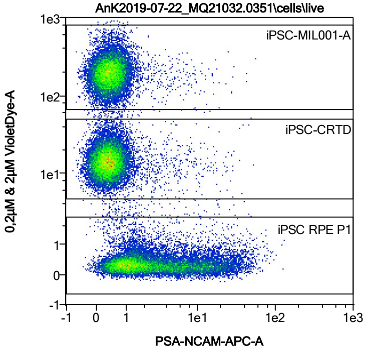


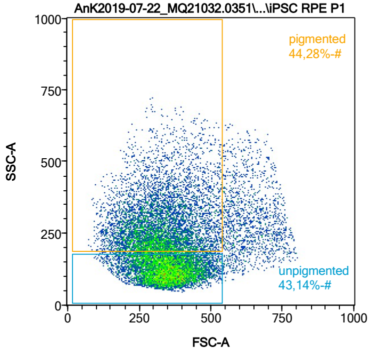


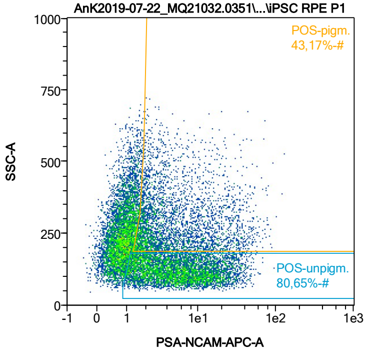


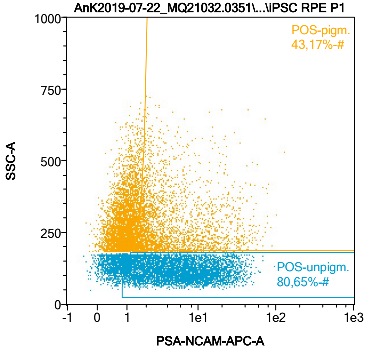


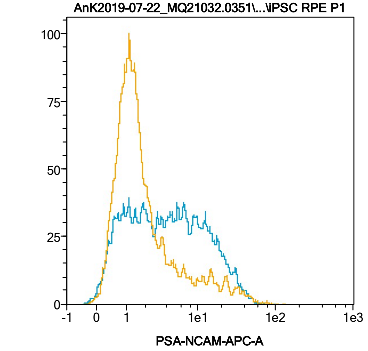


| Name | %-# | PSA-NCAM-APC-A Mean | PSA-NCAM-APC-A Median |
|---|---|---|---|
| POS-unpigm. | 80,65 | 8,35 | 5,26 |
| POS-pigm. | 43,17 | 7,43 | 3,13 |

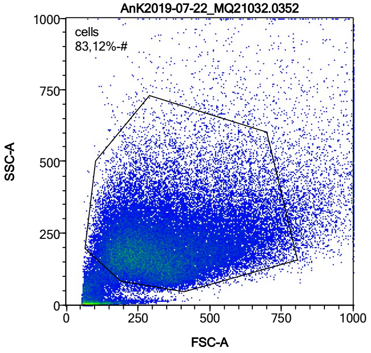


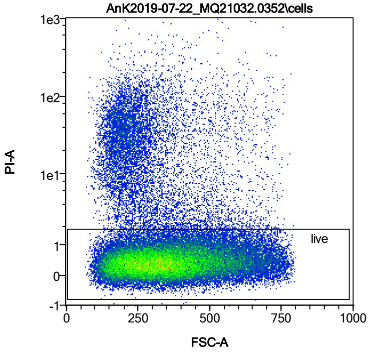


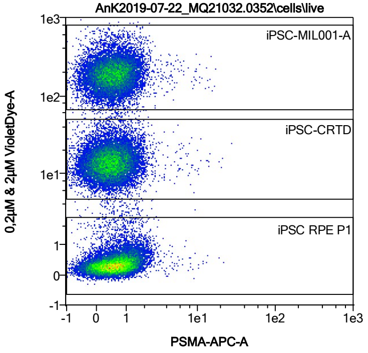


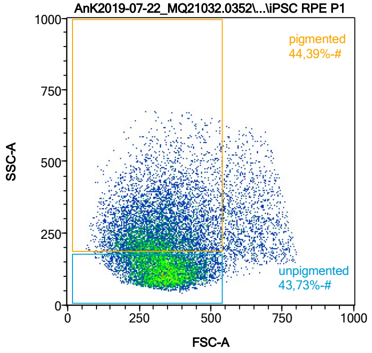


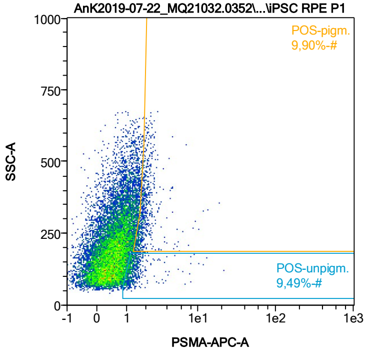


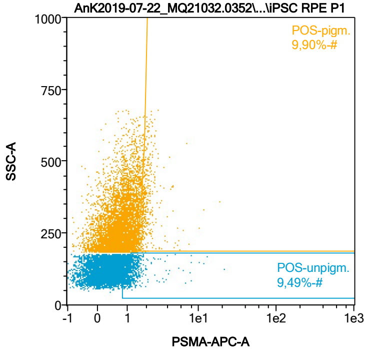


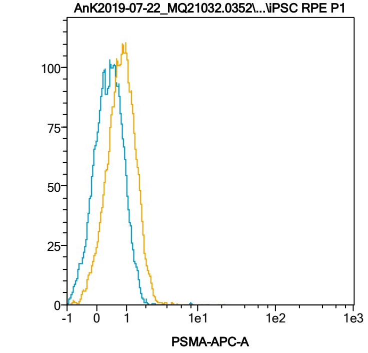


| Name | %-# | PSMA-APC-A Mean | PSMA-APC-A Median |
|---|---|---|---|
| POS-unpigm. | 9,49 | 1,47 | 1,22 |
| POS-pigm. | 9,90 | 2,03 | 1,83 |

Date:2024-06-14

Operator: Sebastian Knoebel

Version:2.13.0+54.g95abf0f4

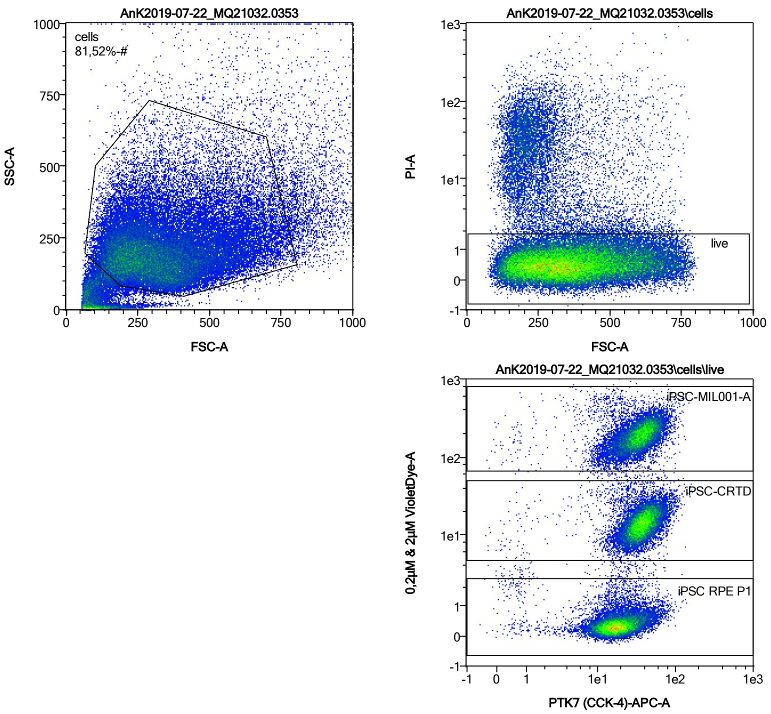


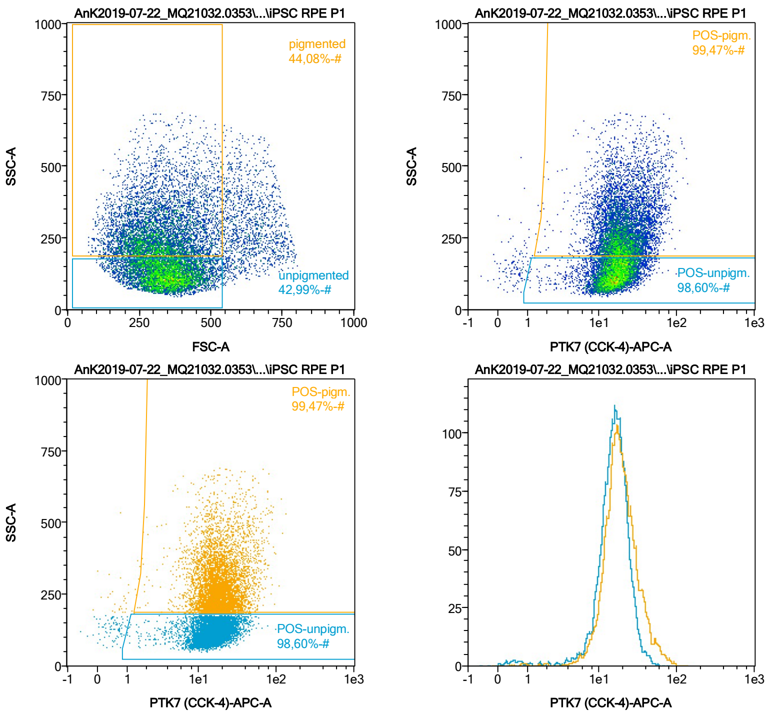


| Name | %-# | PTK7 (CCK-4)-APC-A Mean | PTK7 (CCK-4)-APC-A Median |
|---|---|---|---|
| POS-unpigm. | 98,60 | 16,73 | 15,86 |
| POS-pigm. | 99,47 | 21,27 | 18,25 |

Date:2024-06-14

Operator: Sebastian Knoebel

Version:2.13.0+54.g95abf0f4

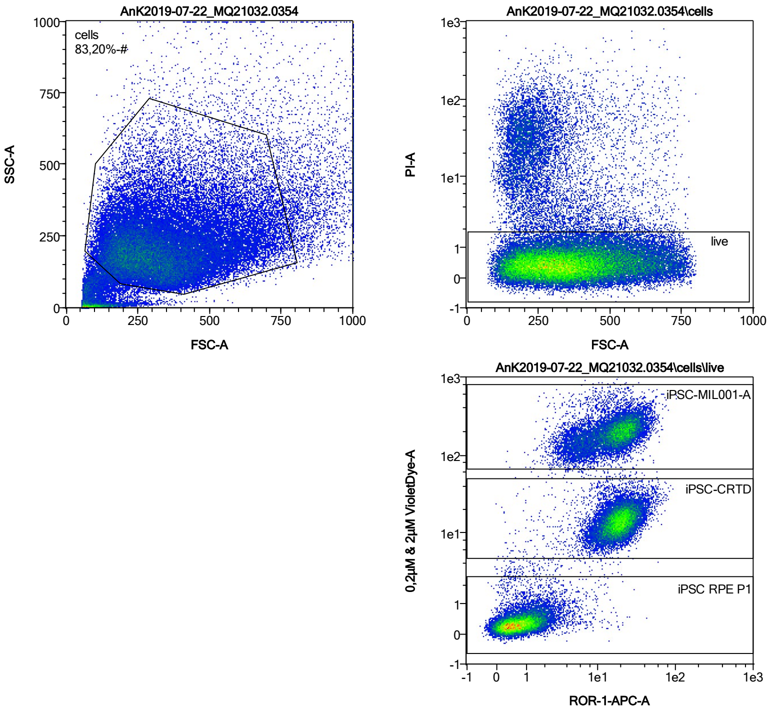


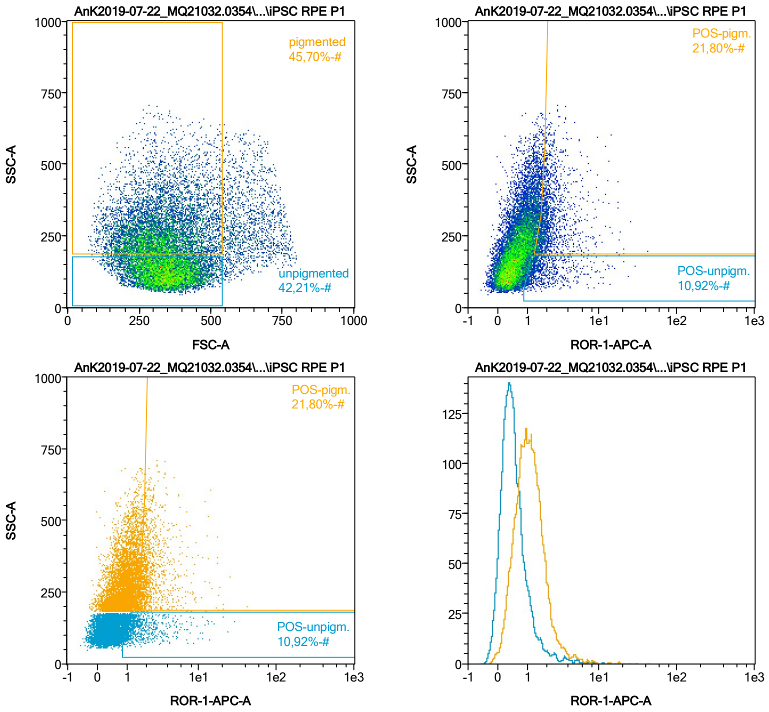


| Name | %-# | ROR-1-APC-A Mean | ROR-1-APC-A Median |
|---|---|---|---|
| POS-unpigm. | 10,92 | 2,20 | 1,45 |
| POS-pigm. | 21,80 | 2,58 | 1,99 |

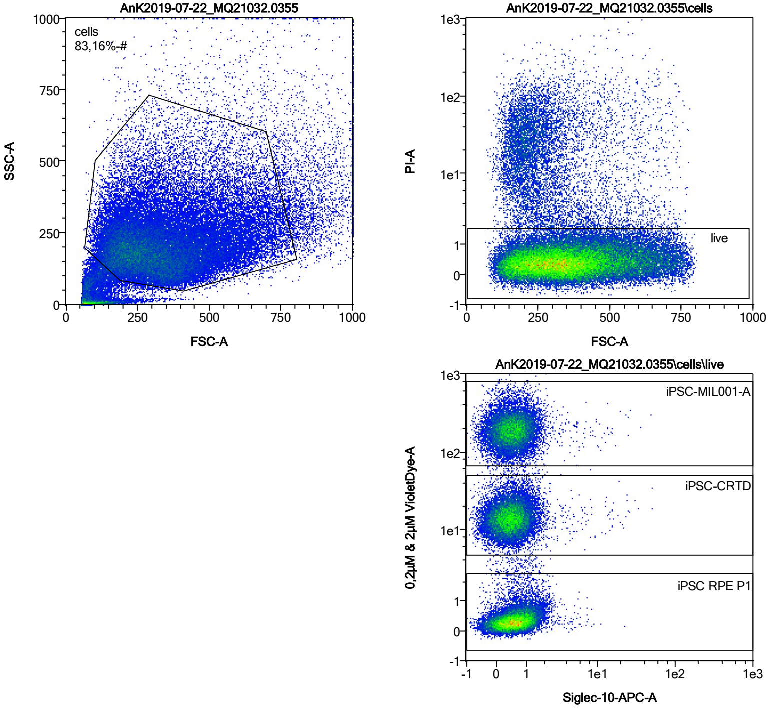


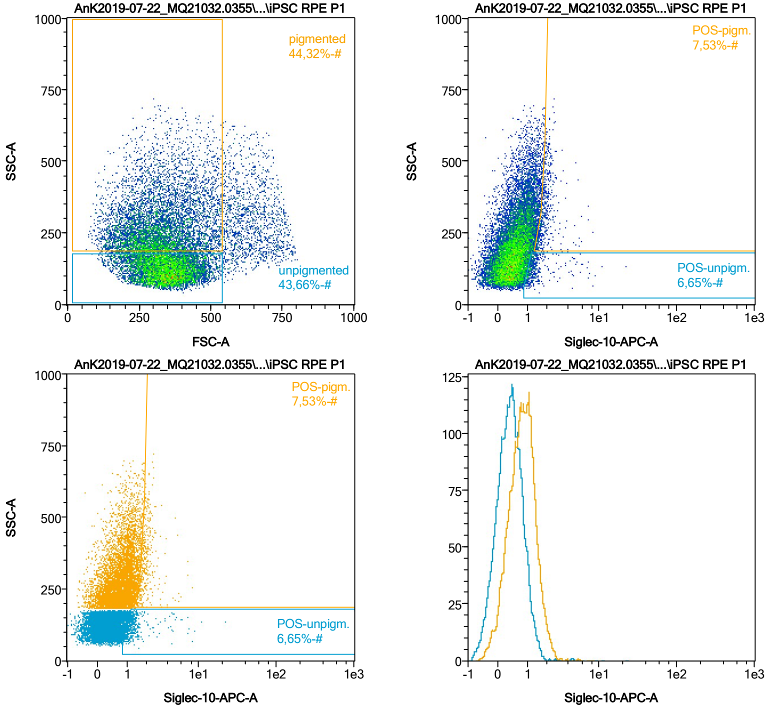


| Name | %-# | Siglec-10-APC-A Mean | Siglec-10-APC-A Median |
|---|---|---|---|
| POS-unpigm. | 6,65 | 1,55 | 1,21 |
| POS-pigm. | 7,53 | 1,99 | 1,84 |

Date:2024-06-14

Operator: Sebastian Knoebel

Version:2.13.0+54.g95abf0f4

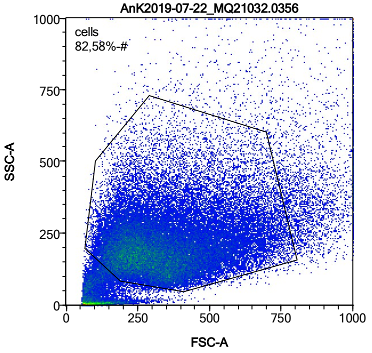


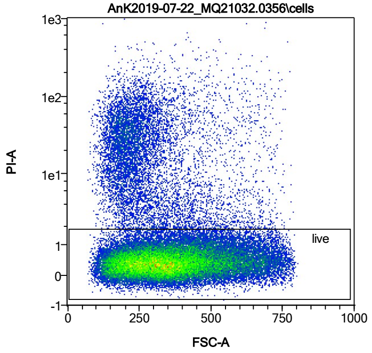


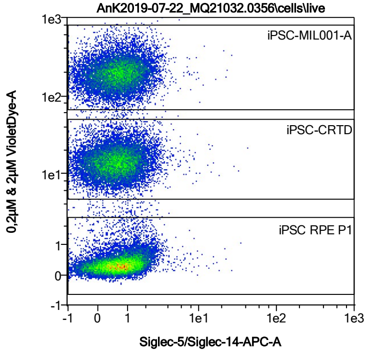


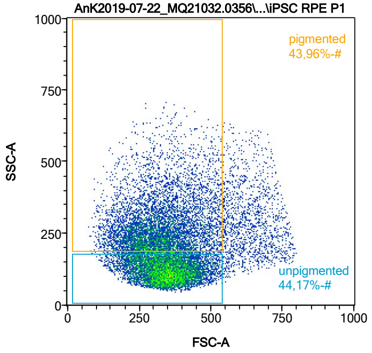


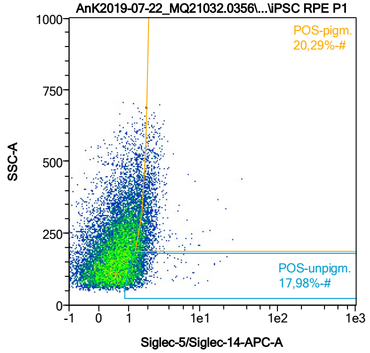


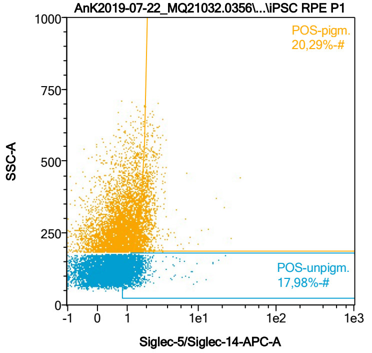


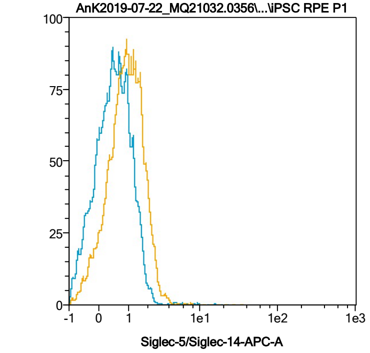


| Name | %-# | Siglec-5/Siglec-14-APC-A Mean | Siglec-5/Siglec-14-APC-A Median |
|---|---|---|---|
| POS-unpigm. | 17,98 | 1,51 | 1,28 |
| POS-pigm. | 20,29 | 2,11 | 1,90 |

Date:2024-06-14
Operator: Sebastian Knoebel
Version:2.13.0+54.g95abf0f4

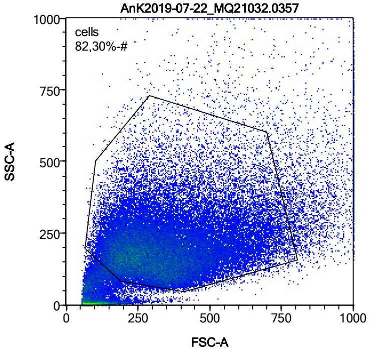


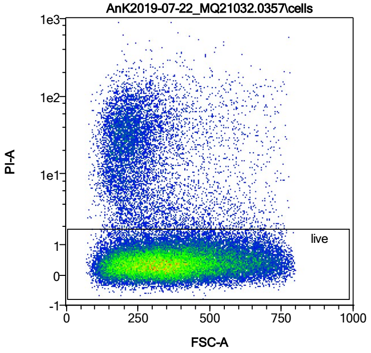


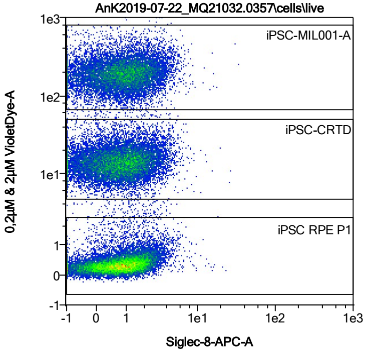


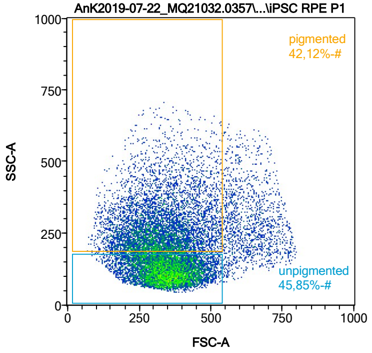


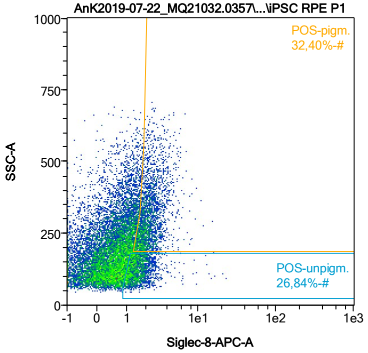


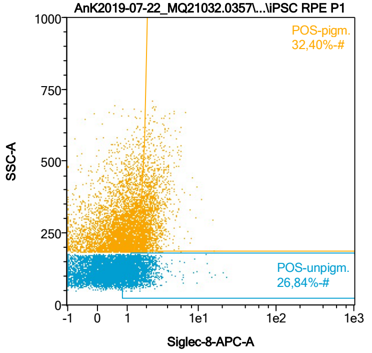


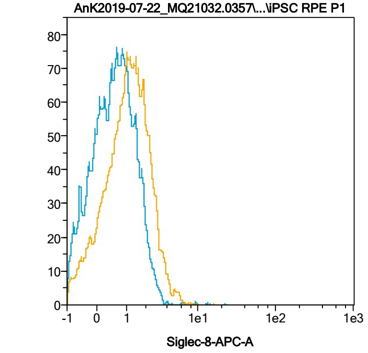


| Name | %-# | Siglec-8-APC-A Mean | Siglec-8-APC-A Median |
|---|---|---|---|
| POS-unpigm. | 26,84 | 1,66 | 1,44 |
| POS-pigm. | 32,40 | 2,35 | 2,10 |

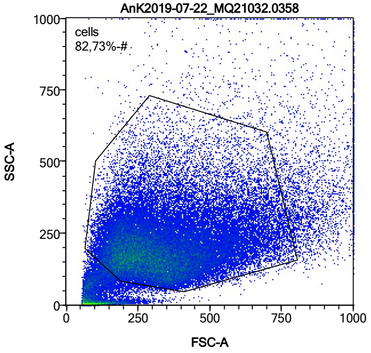


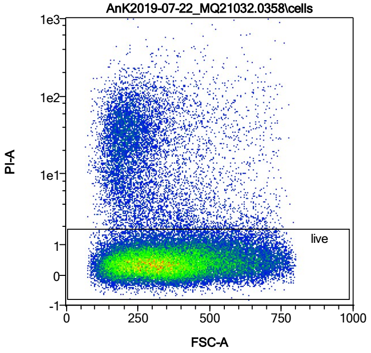


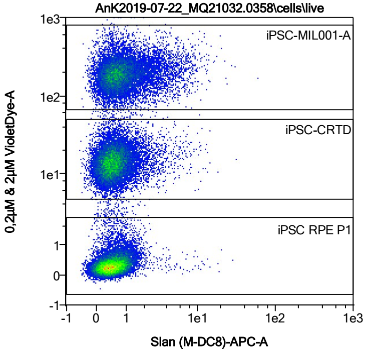


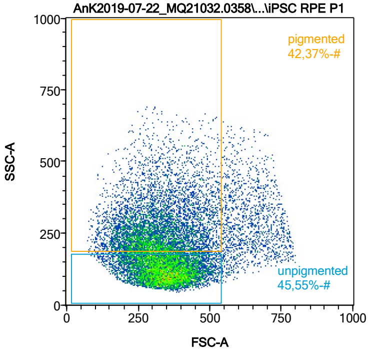


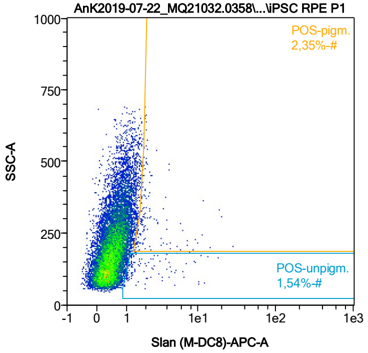


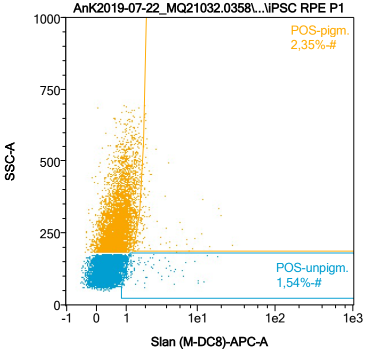


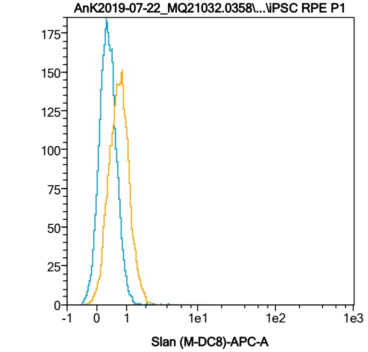


| Name | %-# | Slan (M-DC8)-APC-A Mean | Slan (M-DC8)-APC-A Median |
|---|---|---|---|
| POS-unpigm. | 1,54 | 2,53 | 1,30 |
| POS-pigm. | 2,35 | 2,91 | 1,87 |

Date:2024-06-14
Operator: Sebastian Knoebel
Version:2.13.0+54.g95abf0f4

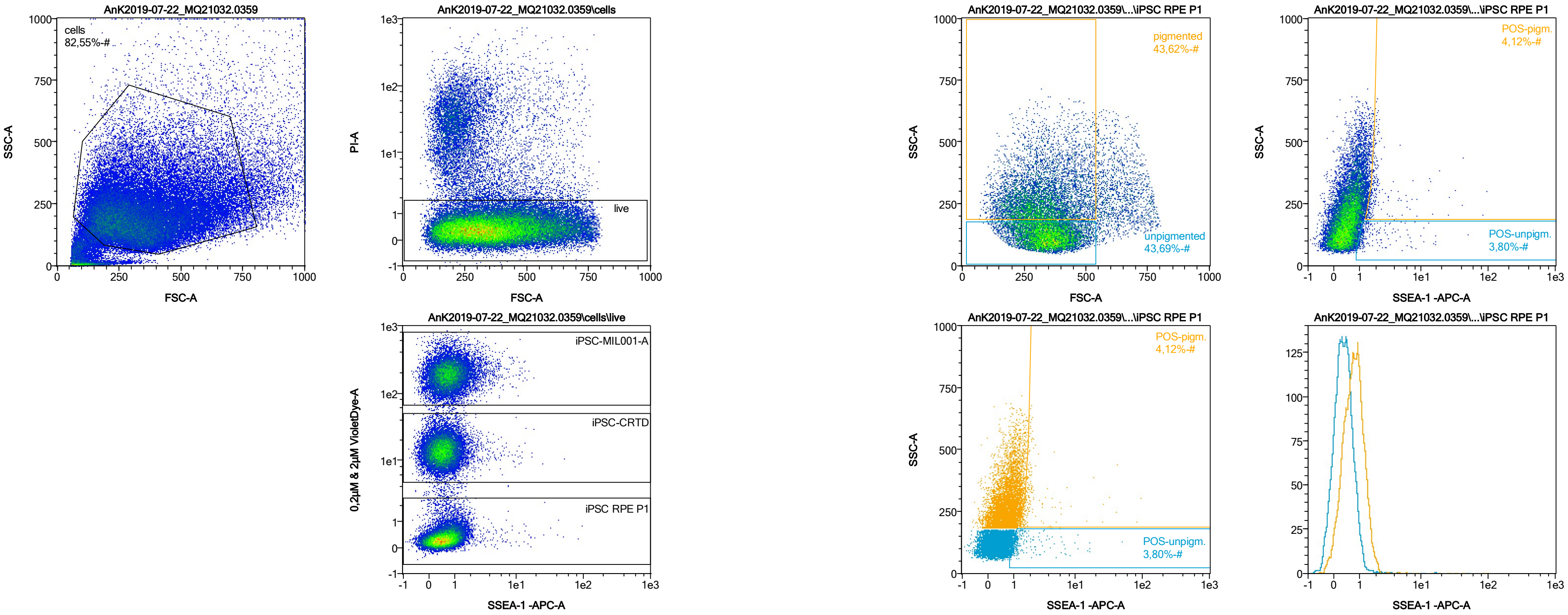


| Name | %-# | SSEA-1 -APC-A Mean | SSEA-1 -APC-A Median |
|---|---|---|---|
| POS-unpigm. | 3,80 | 2,16 | 1,17 |
| POS-pigm. | 4,12 | 3,43 | 1,81 |

Date:2024-06-14

Operator: Sebastian Knoebel

Version:2.13.0+54.g95abf0f4

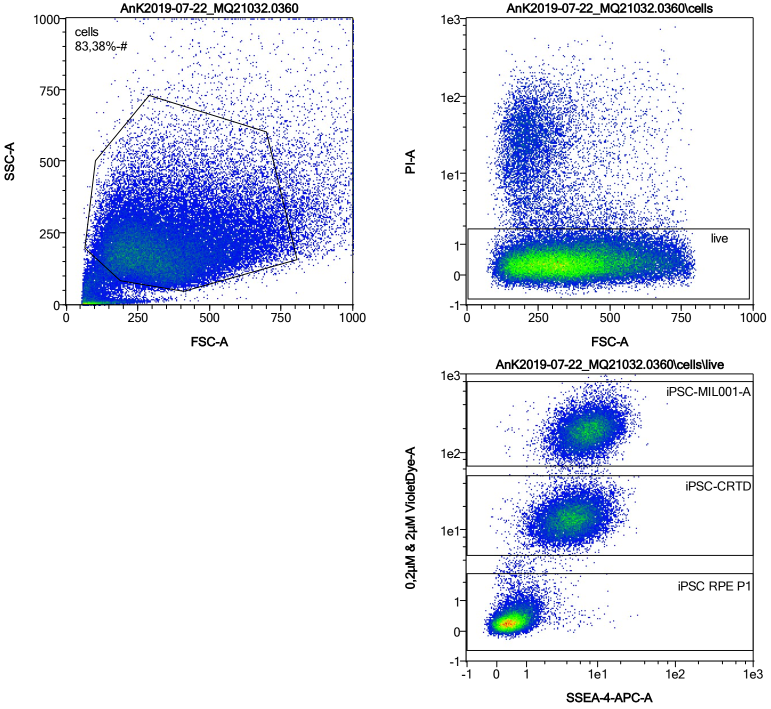


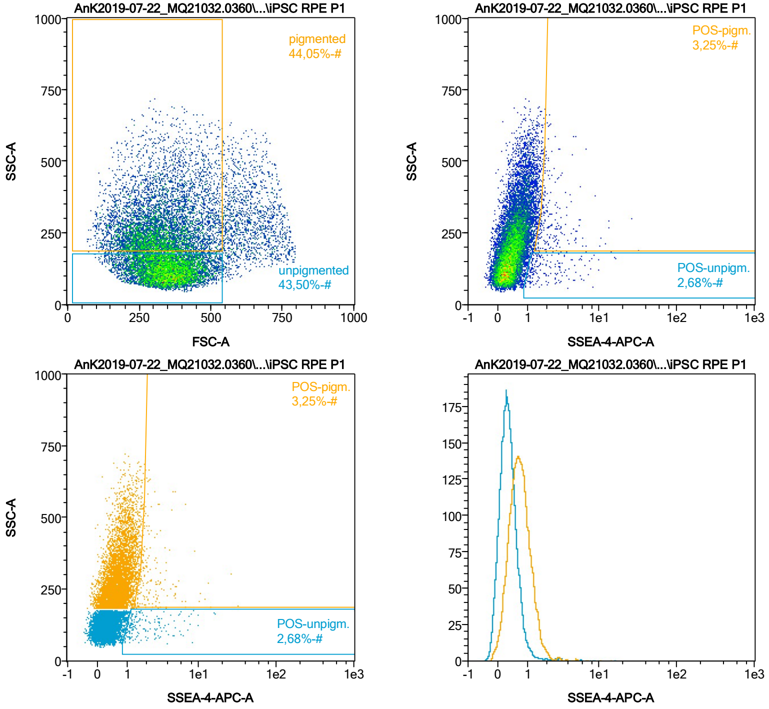


| Name | %-# | SSEA-4-APC-A Mean | SSEA-4-APC-A Median |
|---|---|---|---|
| POS-unpigm. | 2,68 | 2,46 | 1,48 |
| POS-pigm. | 3,25 | 2,89 | 2,01 |

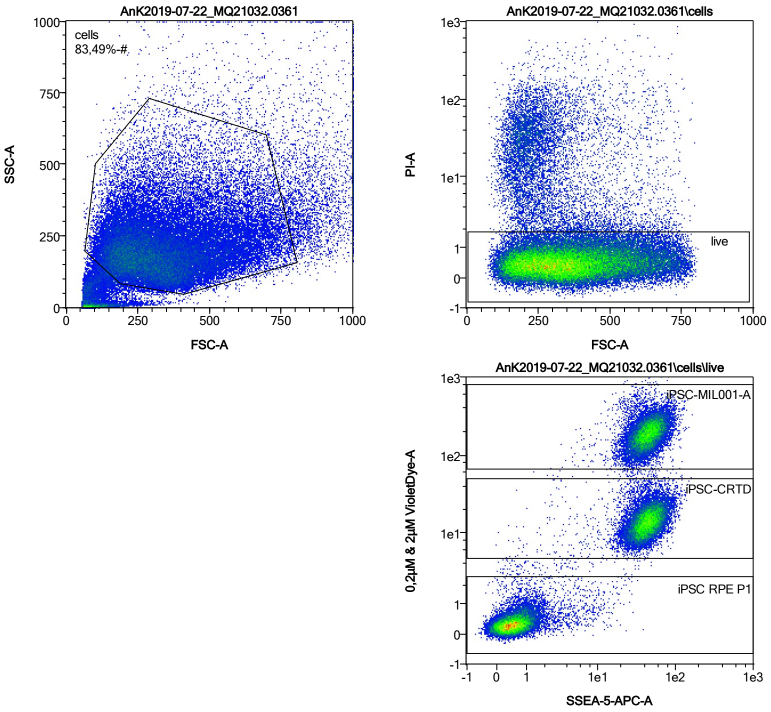


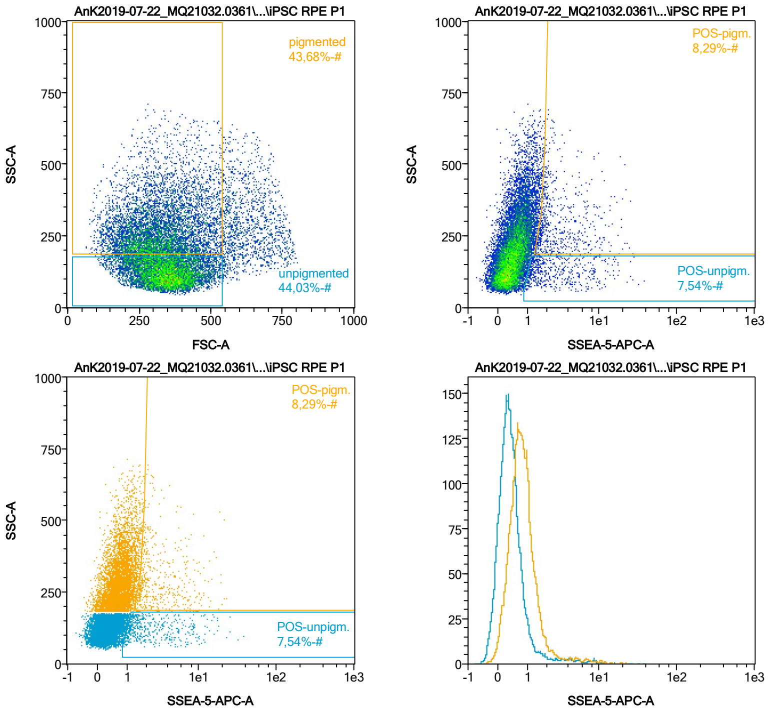


| Name | %-# | SSEA-5-APC-A Mean | SSEA-5-APC-A Median |
|---|---|---|---|
| POS-unpigm. | 7,54 | 3,43 | 1,78 |
| POS-pigm. | 8,29 | 3,75 | 2,29 |

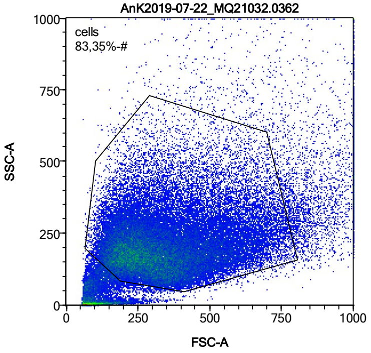


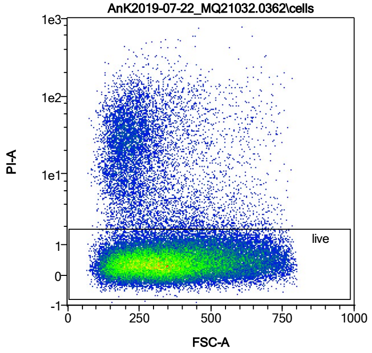


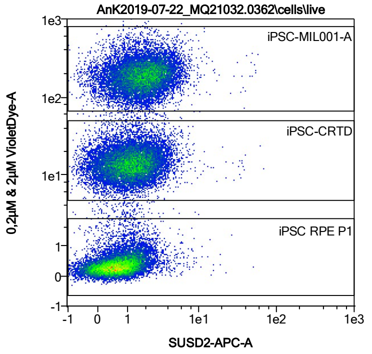


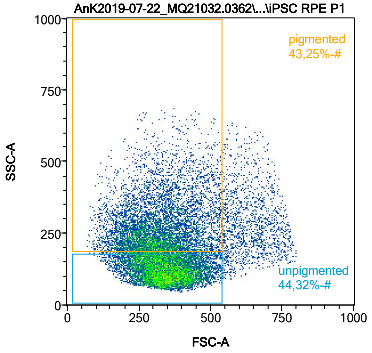


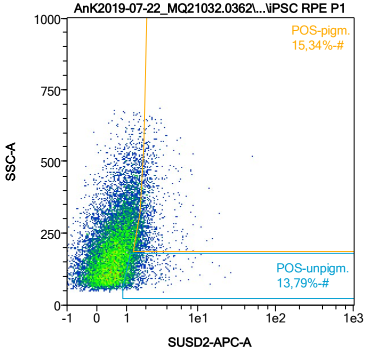


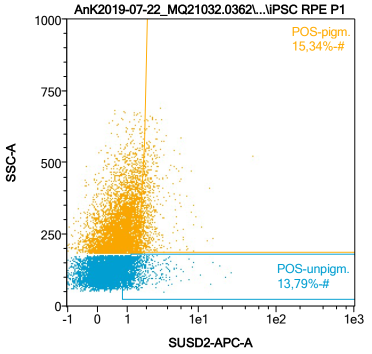


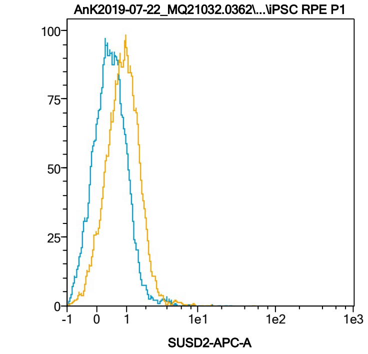


| Name | %-# | SUSD2-APC-A Mean | SUSD2-APC-A Median |
|---|---|---|---|
| POS-unpigm. | 13,79 | 1,65 | 1,28 |
| POS-pigm. | 15,34 | 2,24 | 1,87 |

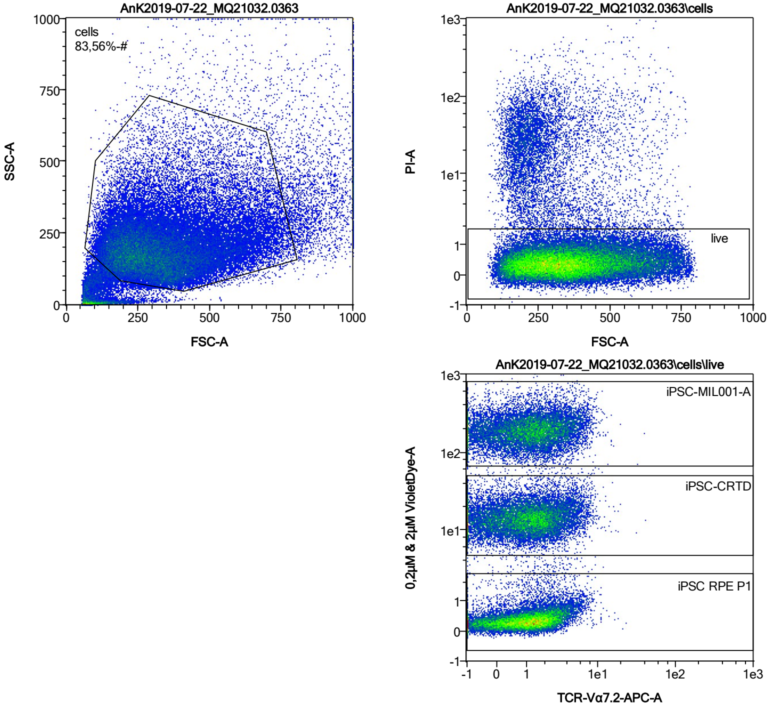


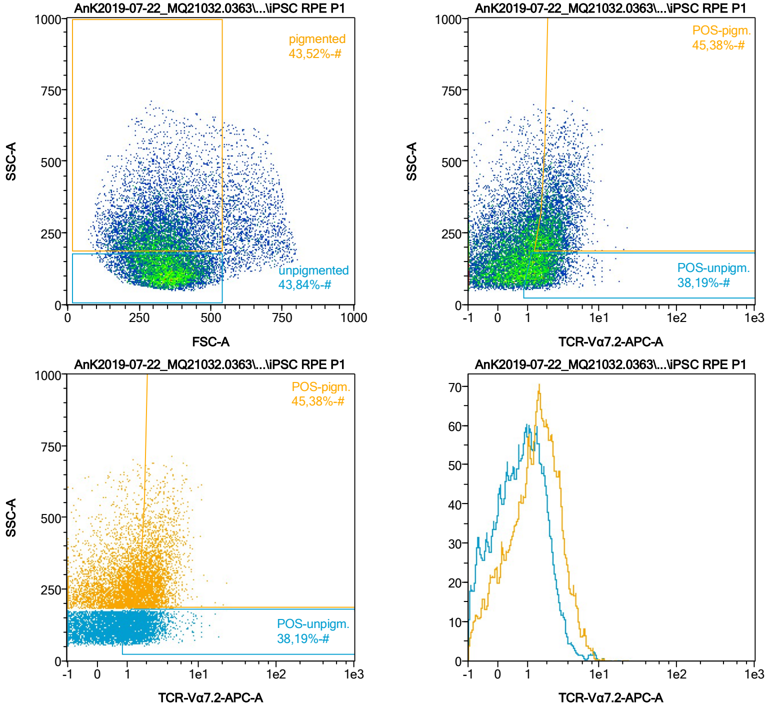


| Name | %-# | TCR-Vα7.2-APC-A Mean | TCR-Vα7.2-APC-A Median |
|---|---|---|---|
| POS-unpigm. | 38,19 | 1,95 | 1,66 |
| POS-pigm. | 45,38 | 2,79 | 2,41 |

Date:2024-06-14

Operator: Sebastian Knoebel

Version:2.13.0+54.g95abf0f4

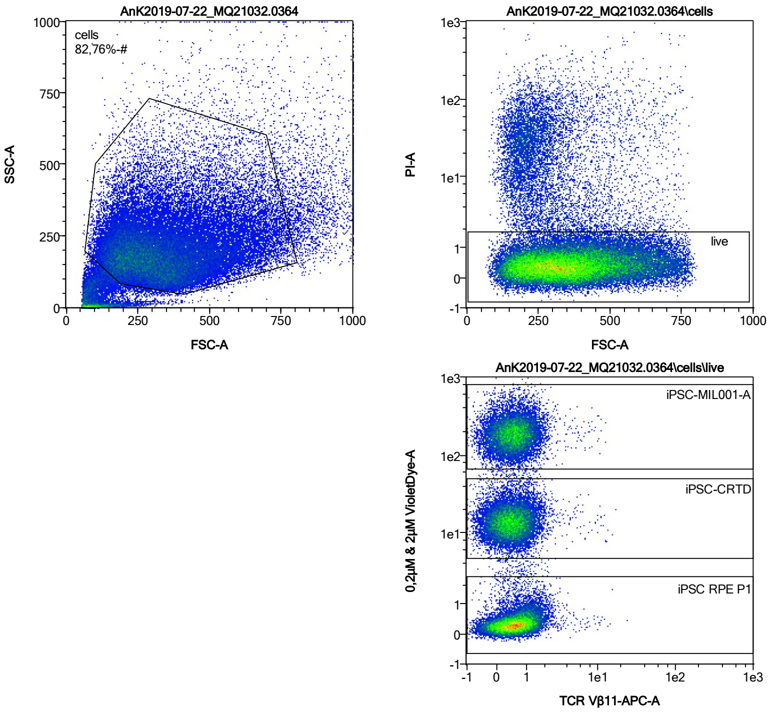


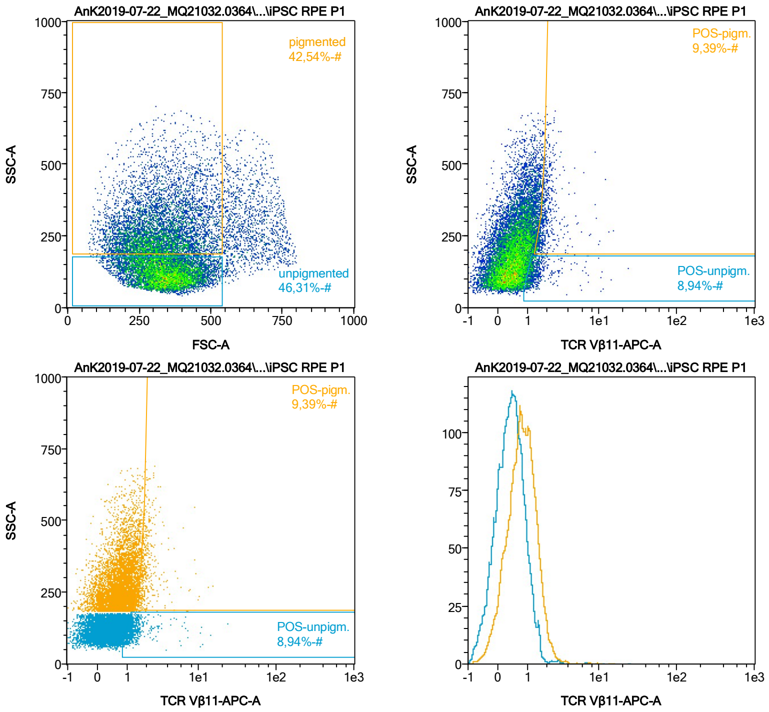


| Name | %-# | TCR Vβ11-APC-A Mean | TCR Vβ11-APC-A Median |
|---|---|---|---|
| POS-unpigm. | 8,94 | 1,53 | 1,23 |
| POS-pigm. | 9,39 | 2,09 | 1,82 |

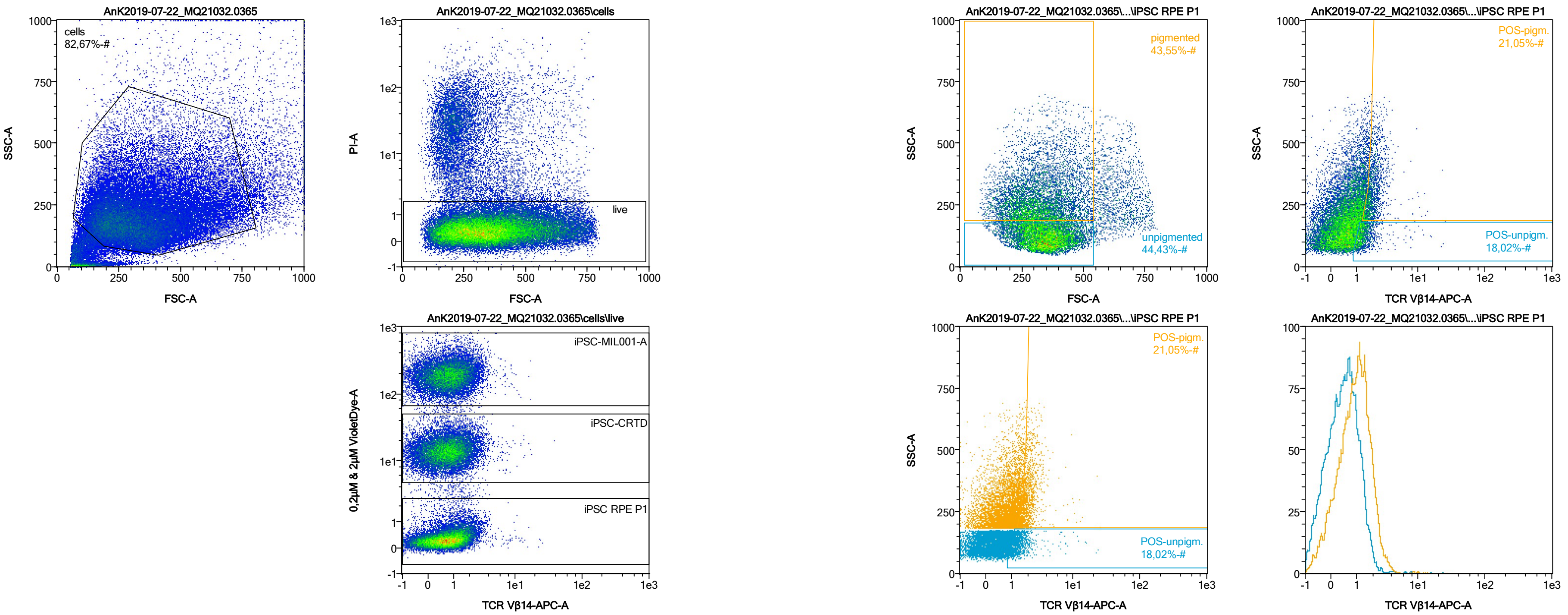


| Name | %-# | TCR Vβ14-APC-A Mean | TCR Vβ14-APC-A Median |
|---|---|---|---|
| POS-unpigm. | 18,02 | 1,54 | 1,29 |
| POS-pigm. | 21,05 | 2,15 | 1,92 |

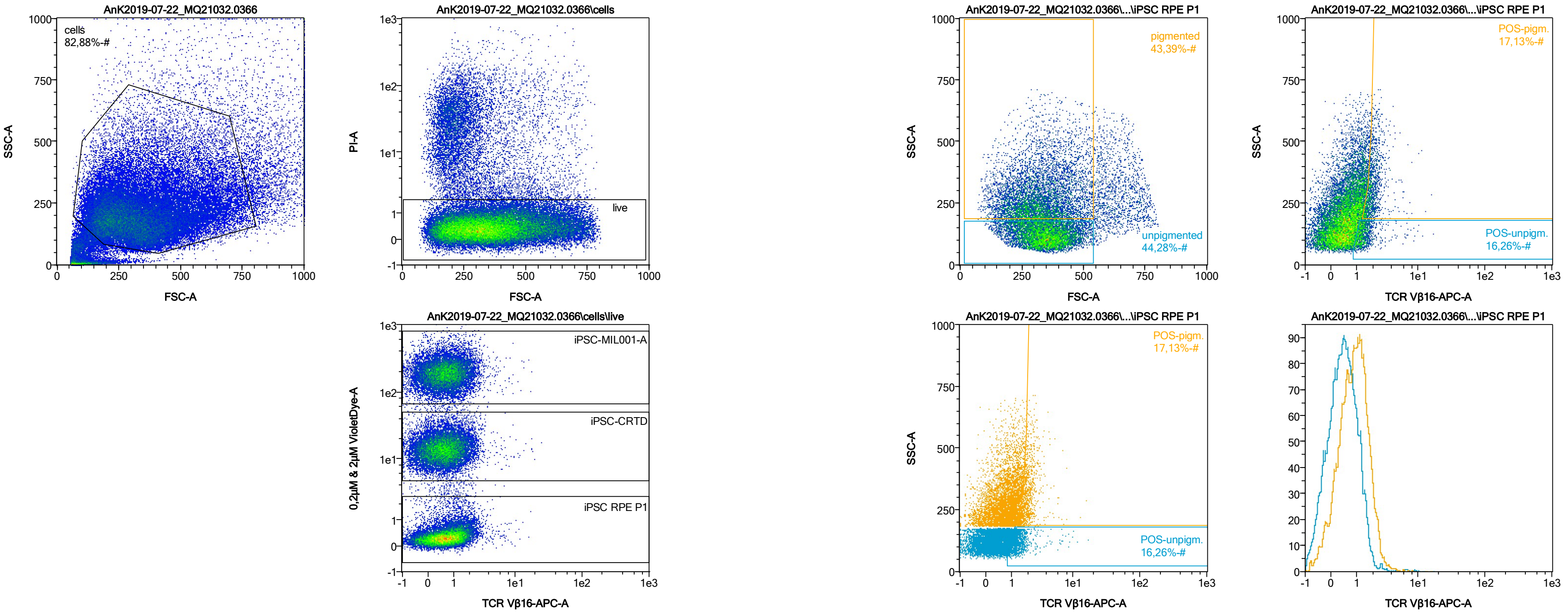


| Name | %-# | TCR Vβ16-APC-A Mean | TCR Vβ16-APC-A Median |
|---|---|---|---|
| POS-unpigm. | 16,26 | 1,50 | 1,27 |
| POS-pigm. | 17,13 | 2,06 | 1,89 |

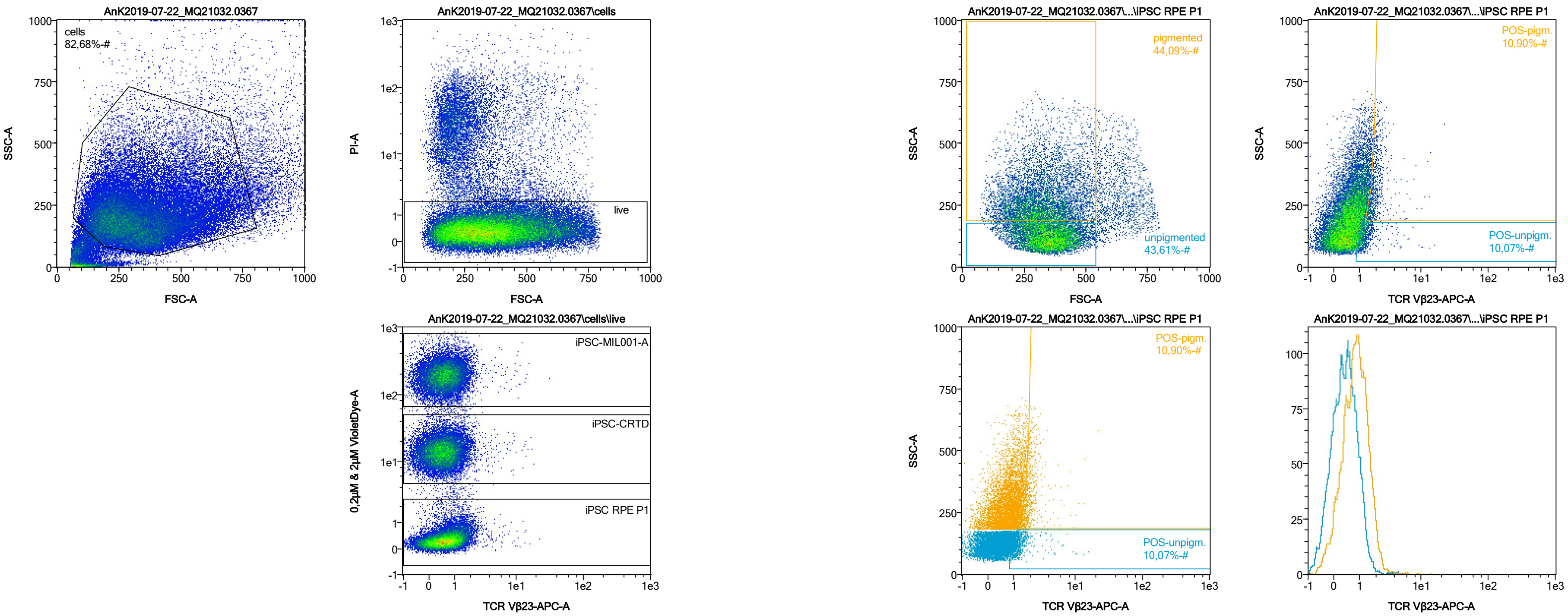


| Name | %-# | TCR Vβ23-APC-A Mean | TCR Vβ23-APC-A Median |
|---|---|---|---|
| POS-unpigm. | 10,07 | 1,47 | 1,22 |
| POS-pigm. | 10,90 | 2,12 | 1,86 |

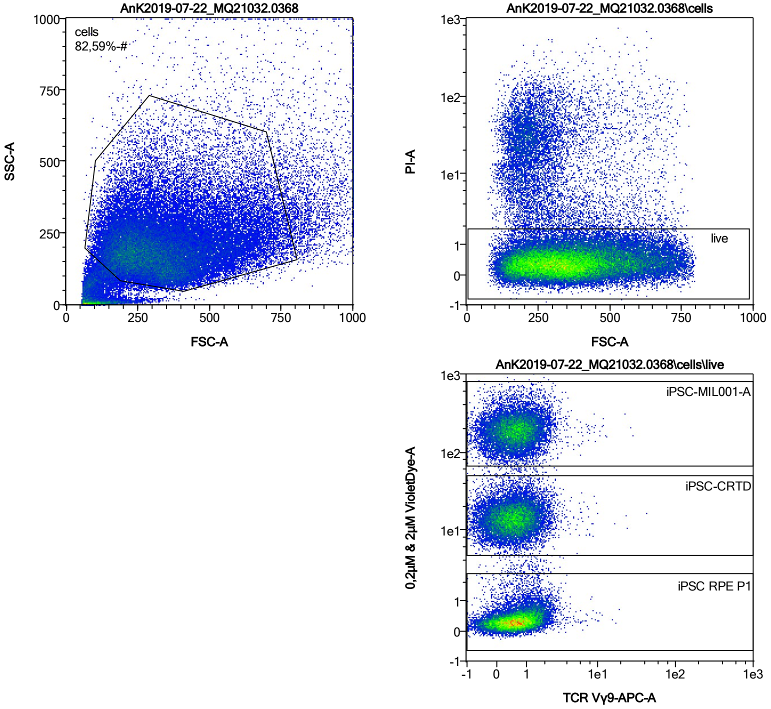


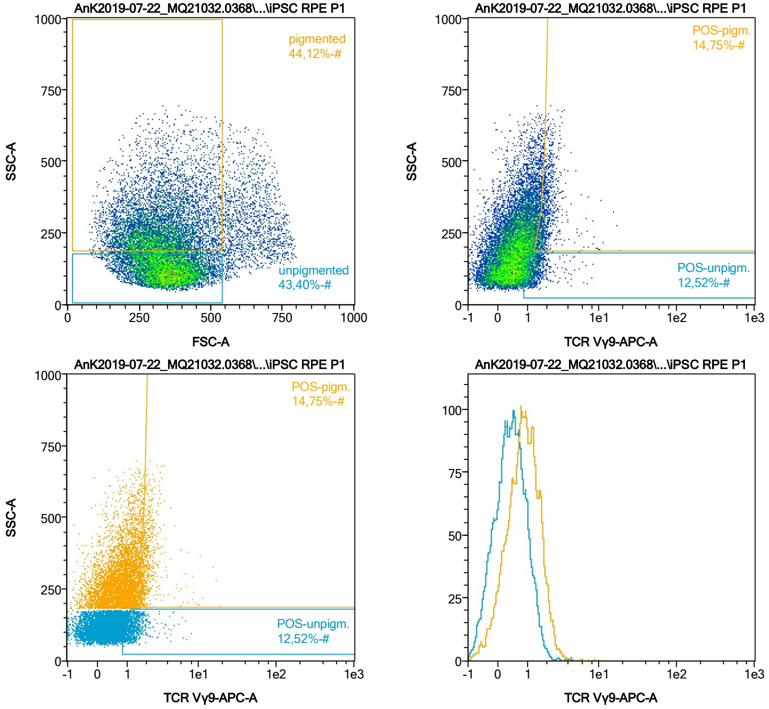


| Name | %-# | TCR Vγ9-APC-A Mean | TCR Vγ9-APC-A Median |
|---|---|---|---|
| POS-unpigm. | 12,52 | 1,42 | 1,27 |
| POS-pigm. | 14,75 | 2,13 | 1,86 |

Date:2024-06-14
Operator: Sebastian Knoebel
Version:2.13.0+54.g95abf0f4

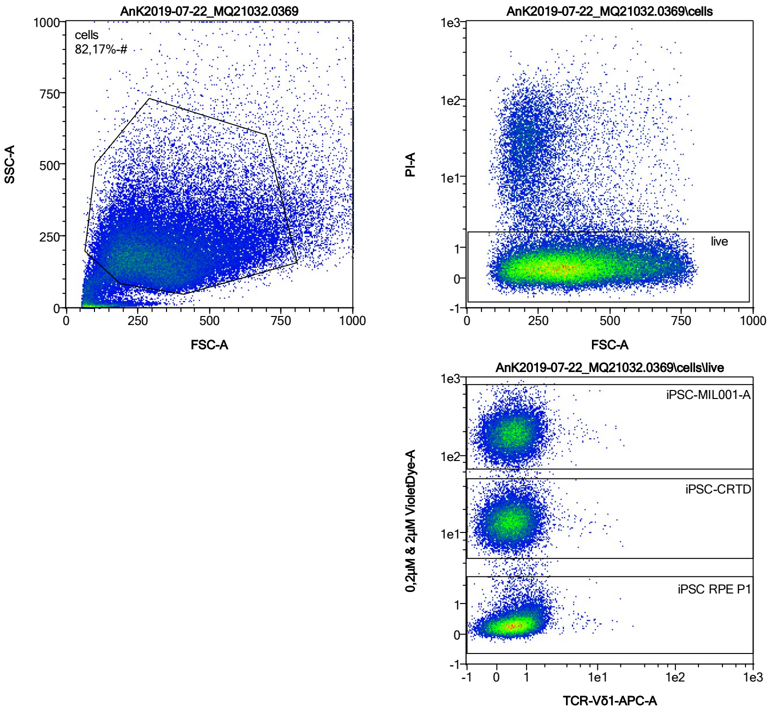


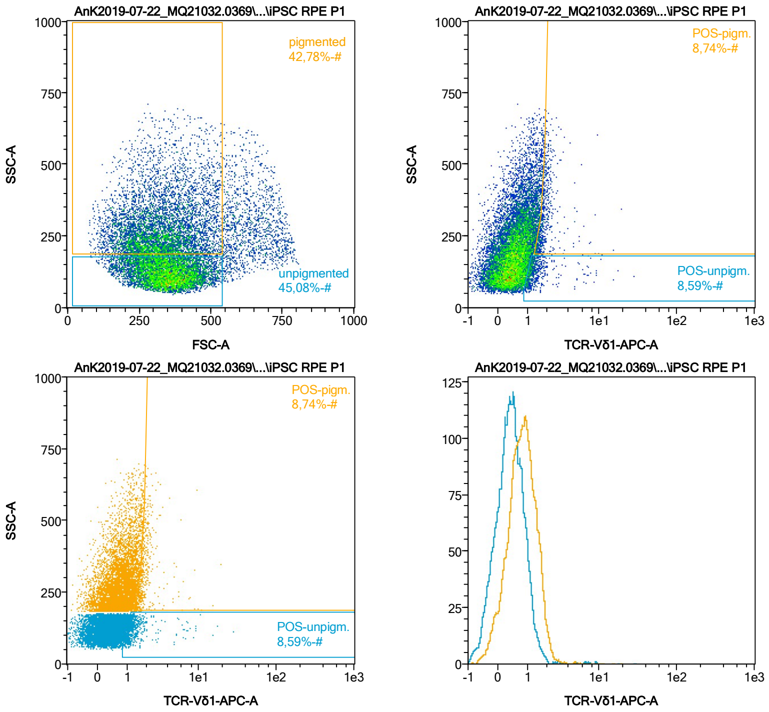


| Name | %-# | TCR-Vδ1-APC-A Mean | TCR-Vδ1-APC-A Median |
|---|---|---|---|
| POS-unpigm. | 8,59 | 1,60 | 1,20 |
| POS-pigm. | 8,74 | 2,06 | 1,77 |

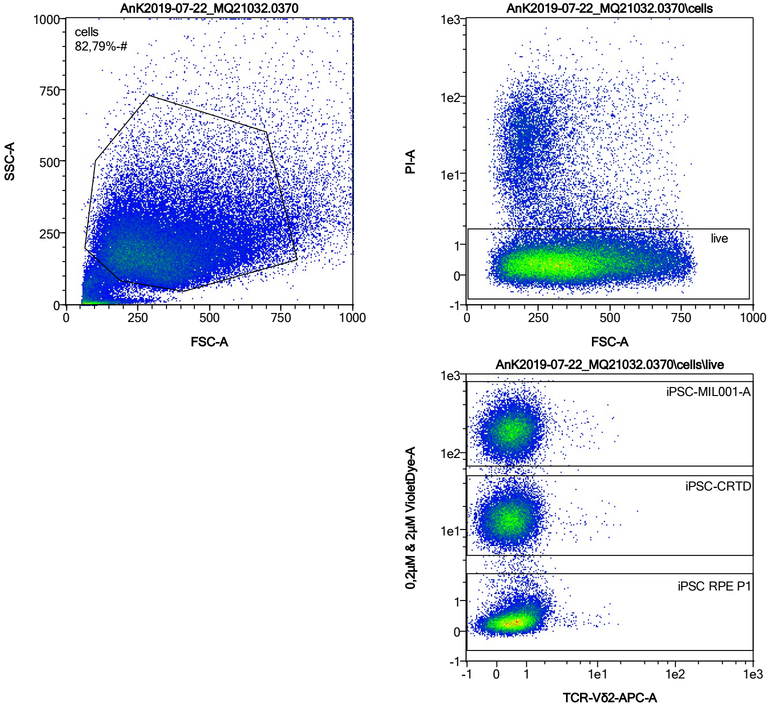


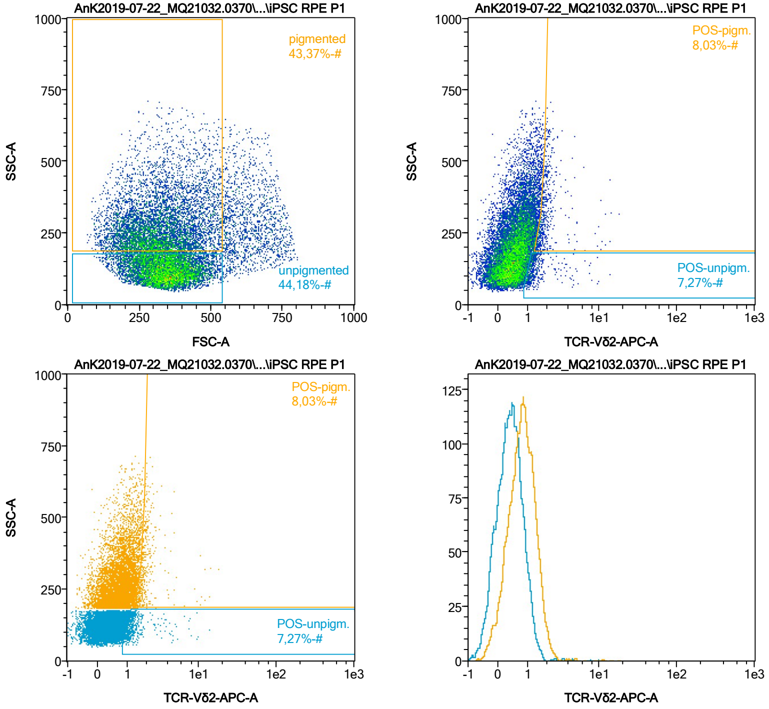


| Name | %-# | TCR-Vδ2-APC-A Mean | TCR-Vδ2-APC-A Median |
|---|---|---|---|
| POS-unpigm. | 7,27 | 1,52 | 1,20 |
| POS-pigm. | 8,03 | 2,10 | 1,78 |

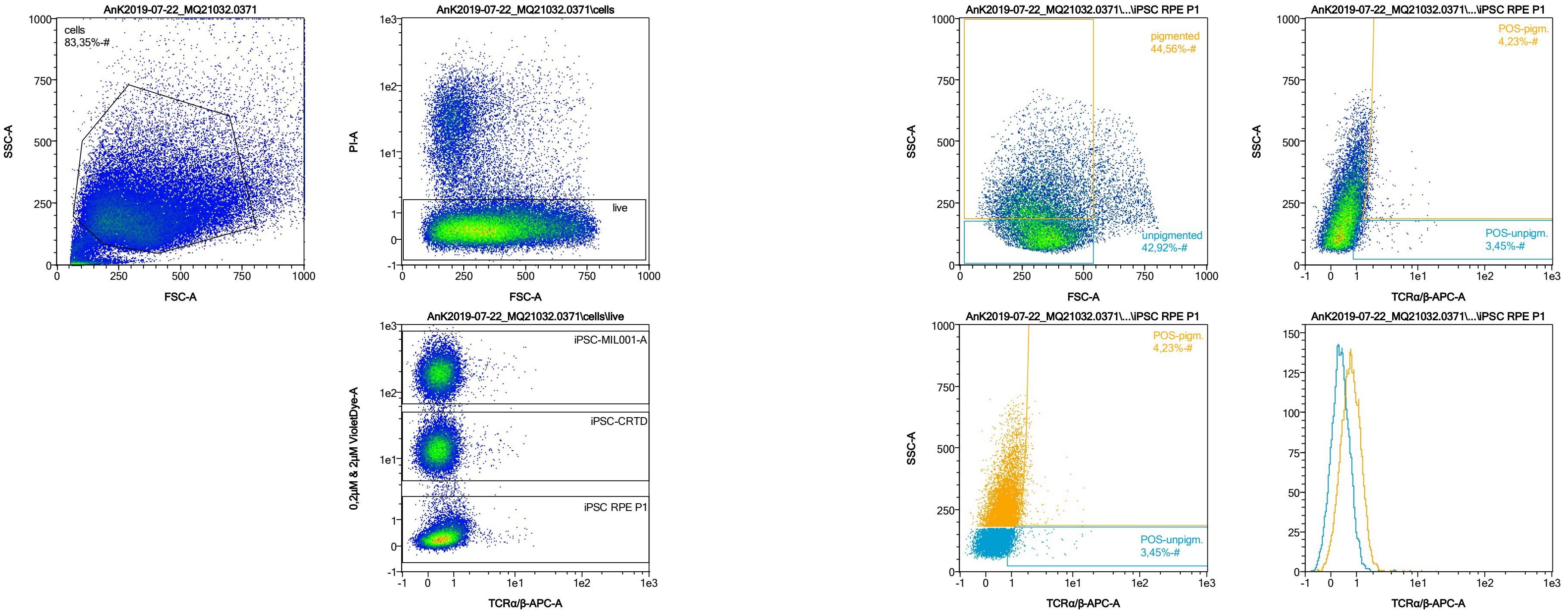


| Name | %-# | TCRα/β-APC-A Mean | TCRα/β-APC-A Median |
|---|---|---|---|
| POS-unpigm. | 3,45 | 1,88 | 1,18 |
| POS-pigm. | 4,23 | 2,29 | 1,80 |

Date:2024-06-14
Operator: Sebastian Knoebel
Version:2.13.0+54.g95abf0f4

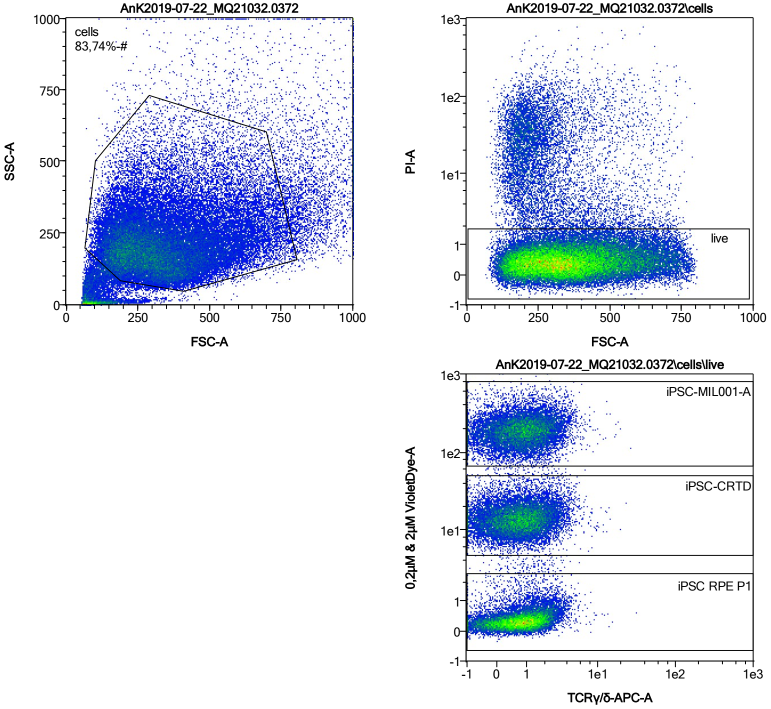


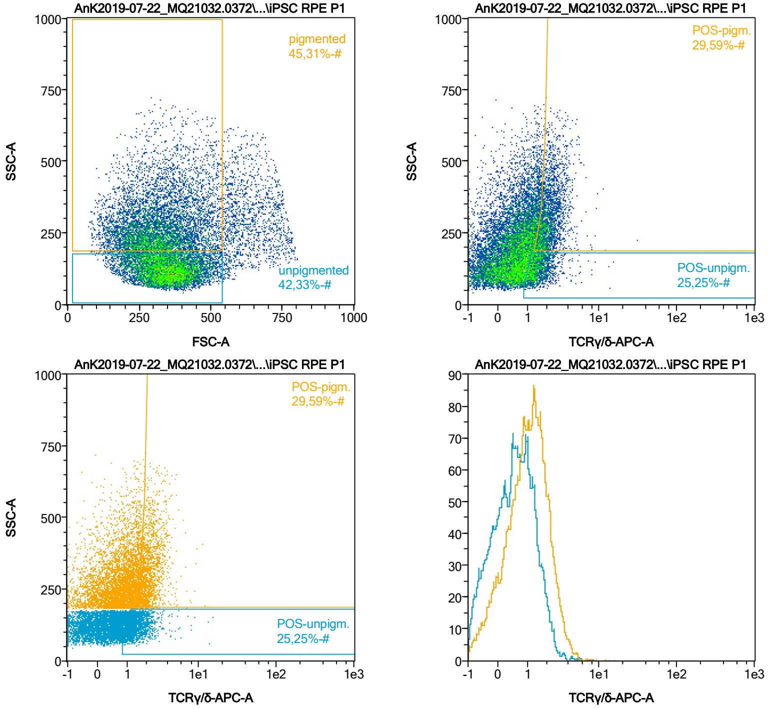


| Name | %-# | TCRγ/δ-APC-A Mean | TCRγ/δ-APC-A Median |
|---|---|---|---|
| POS-unpigm. | 25,25 | 1,59 | 1,39 |
| POS-pigm. | 29,59 | 2,25 | 2,04 |

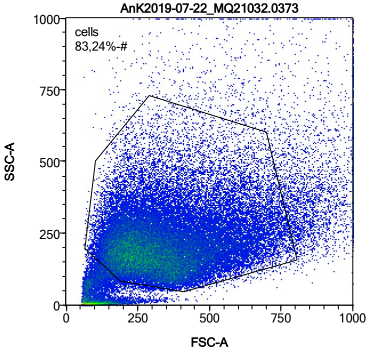


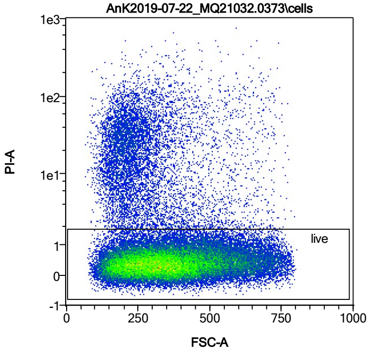


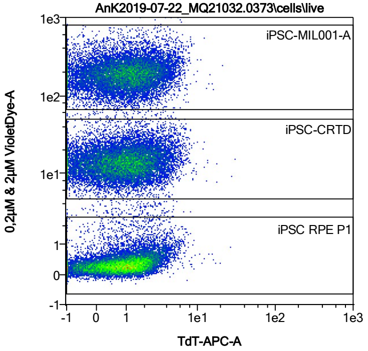


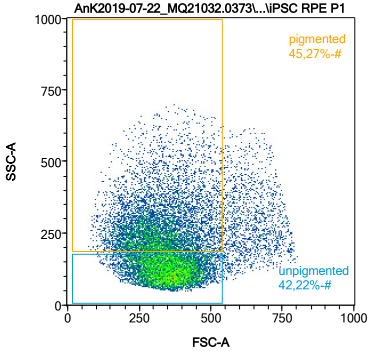


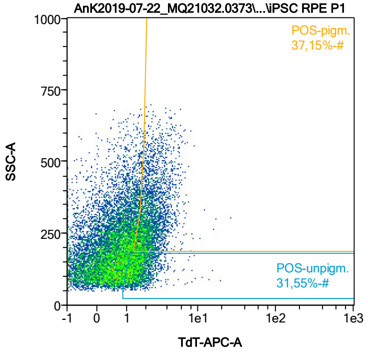


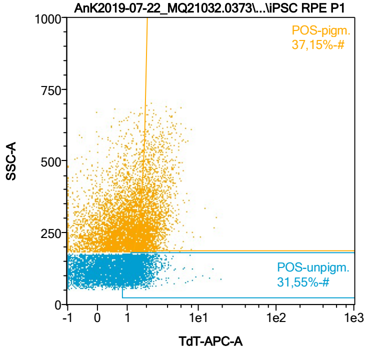


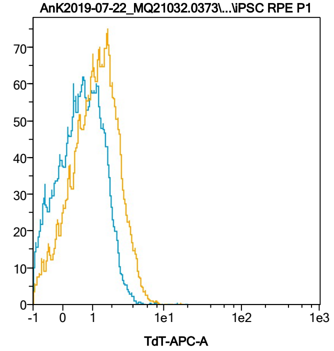


| Name | %-# | TdT-APC-A Mean | TdT-APC-A Median |
|---|---|---|---|
| POS-unpigm. | 31,55 | 1,68 | 1,49 |
| POS-pigm. | 37,15 | 2,49 | 2,20 |

Date:2024-06-14
Operator: Sebastian Knoebel
Version:2.13.0+54.g95abf0f4

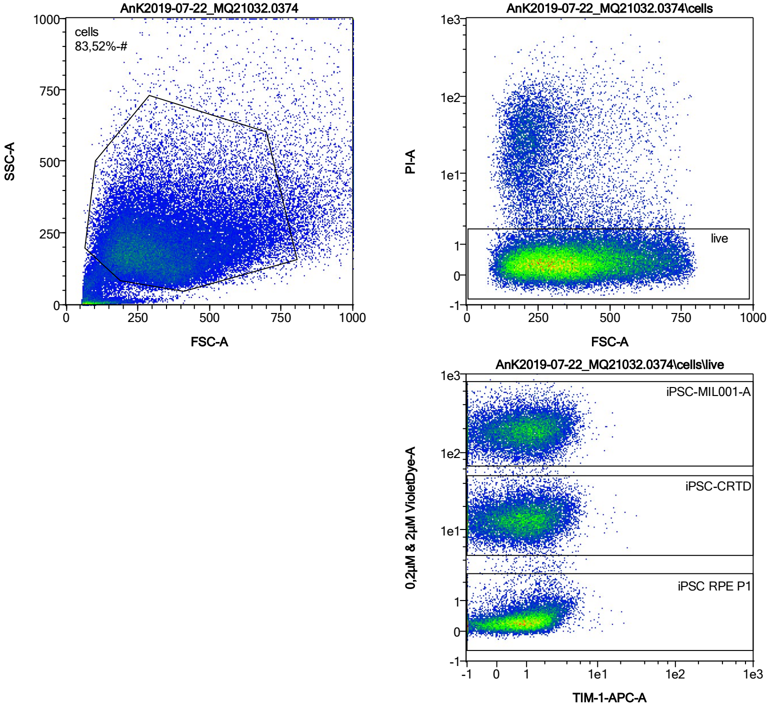


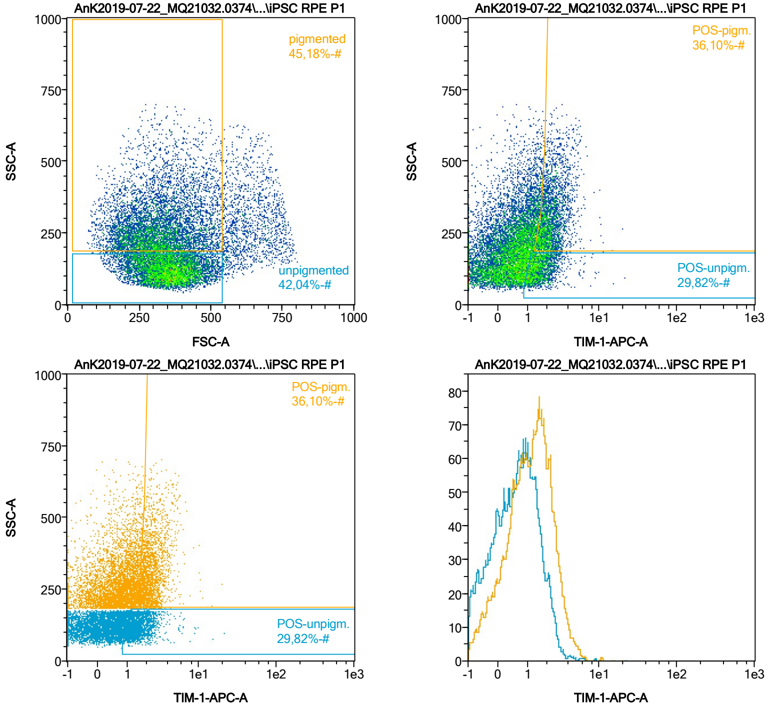


| Name | %-# | TIM-1-APC-A Mean | TIM-1-APC-A Median |
|---|---|---|---|
| POS-unpigm. | 29,82 | 1,68 | 1,47 |
| POS-pigm. | 36,10 | 2,40 | 2,16 |

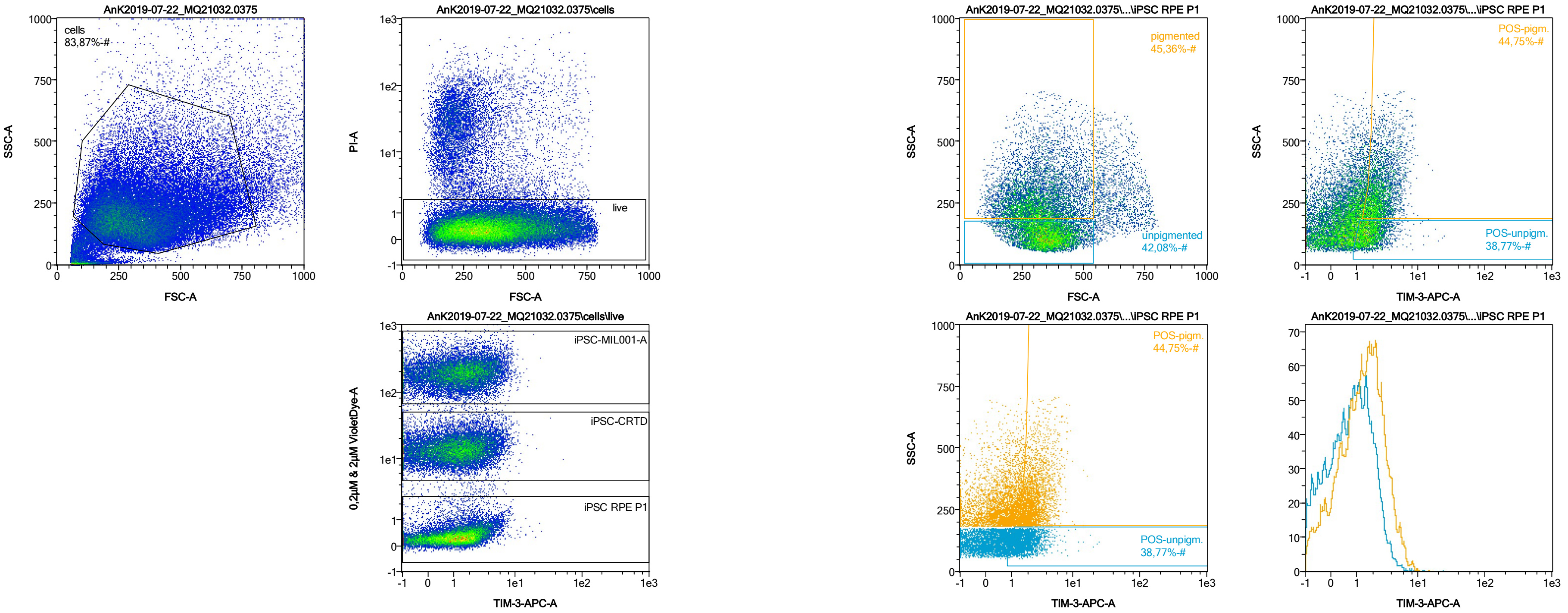


| Name | %-# | TIM-3-APC-A Mean | TIM-3-APC-A Median |
|---|---|---|---|
| POS-unpigm. | 38,77 | 1,93 | 1,67 |
| POS-pigm. | 44,75 | 2,76 | 2,39 |

Date:2024-06-14

Operator: Sebastian Knoebel

Version:2.13.0+54.g95abf0f4

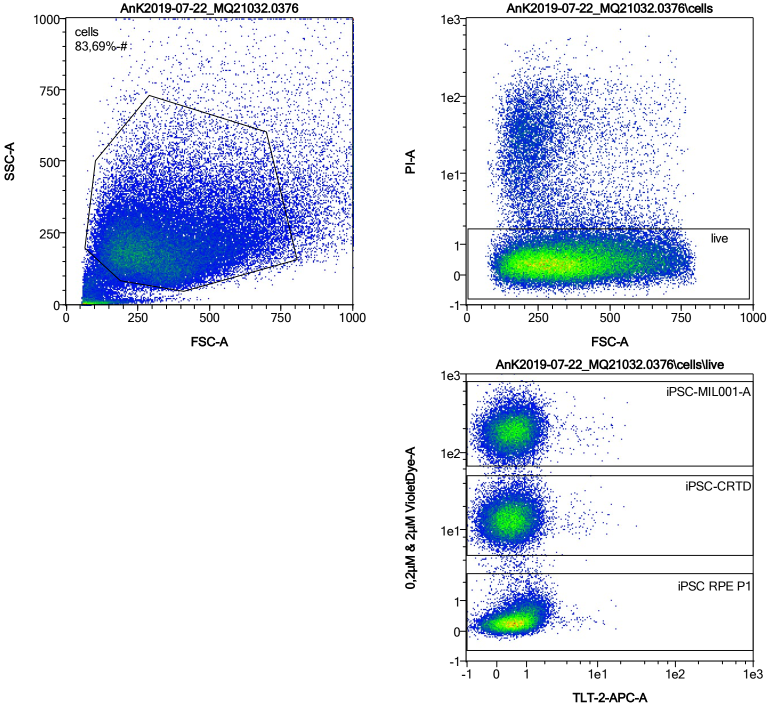


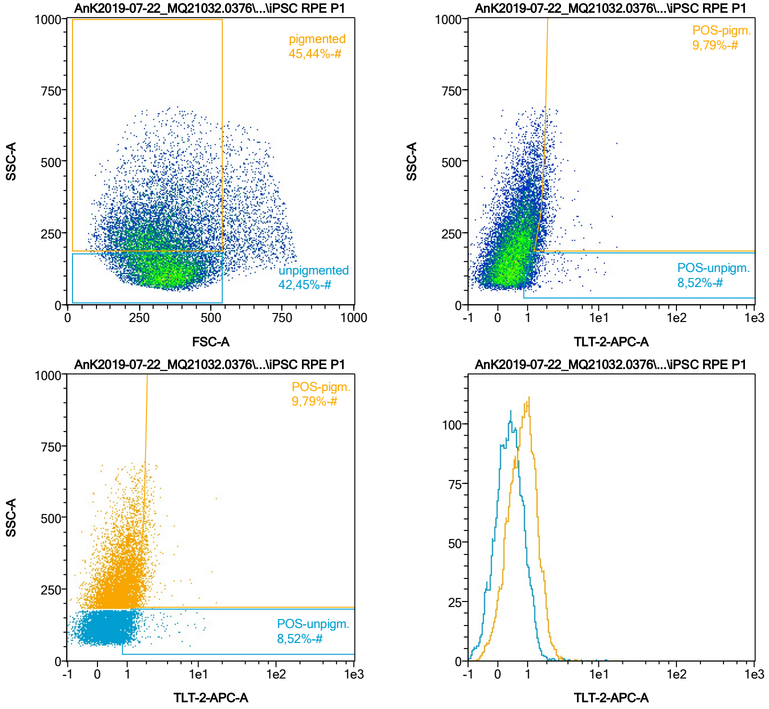


| Name | %-# | TLT-2-APC-A Mean | TLT-2-APC-A Median |
|---|---|---|---|
| POS-unpigm. | 8,52 | 1,53 | 1,21 |
| POS-pigm. | 9,79 | 2,04 | 1,84 |

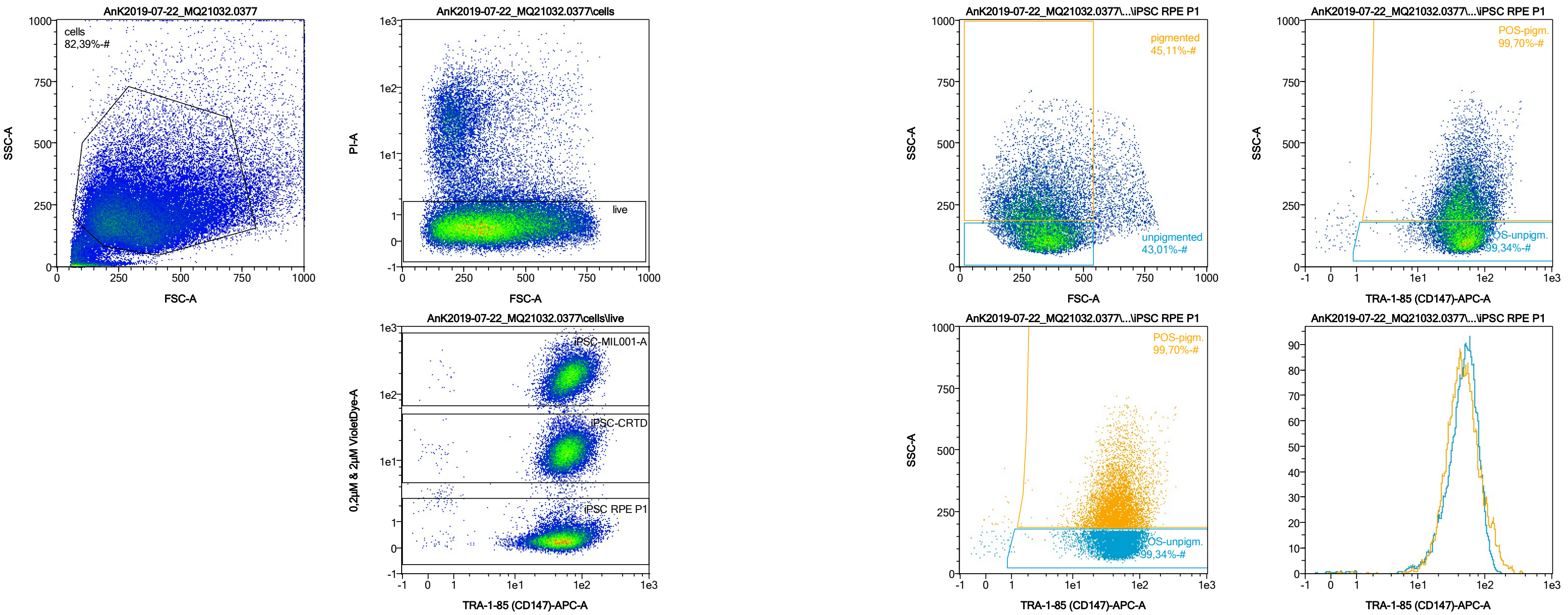


| Name | %-# | TRA-1-85 (CD147)-APC-A Mean | TRA-1-85 (CD147)-APC-A Median |
|---|---|---|---|
| POS-unpigm. | 99,34 | 50,97 | 48,44 |
| POS-pigm. | 99,70 | 54,33 | 45,37 |

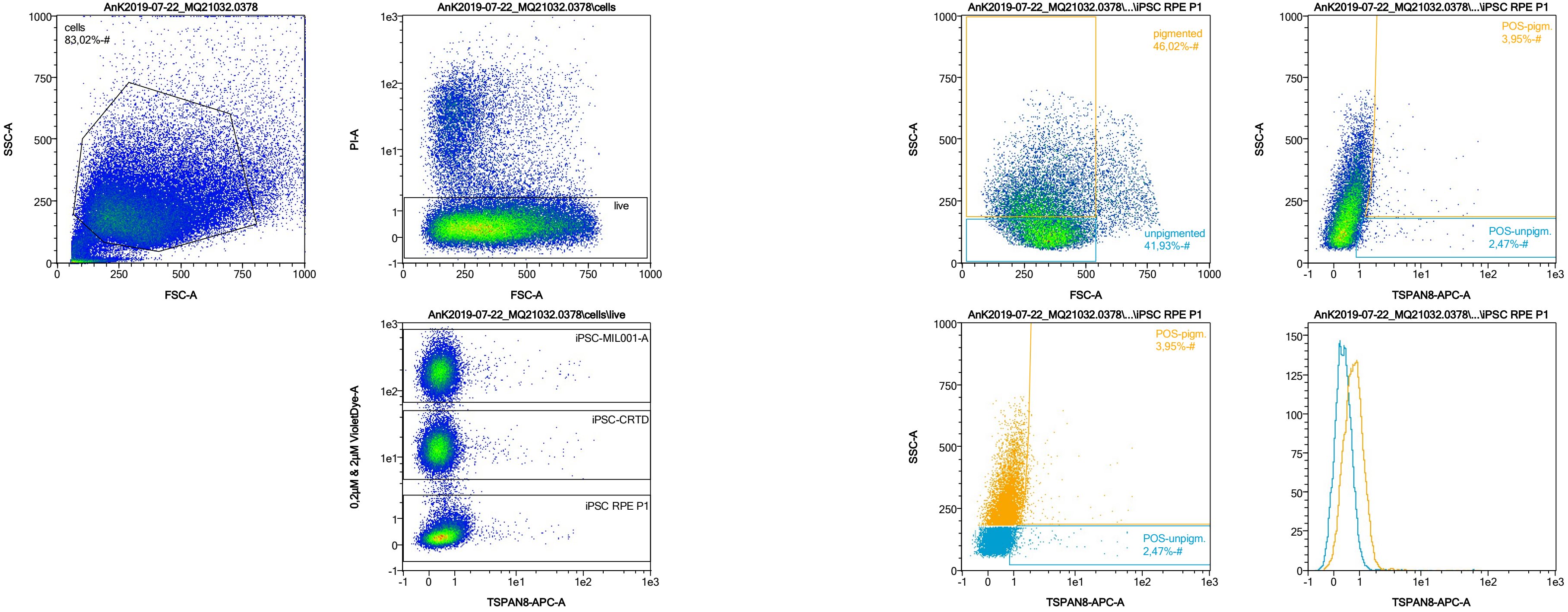


| Name | %-# | TSPAN8-APC-A Mean | TSPAN8-APC-A Median |
|---|---|---|---|
| POS-unpigm. | 2,47 | 4,11 | 1,25 |
| POS-pigm. | 3,95 | 4,02 | 1,83 |

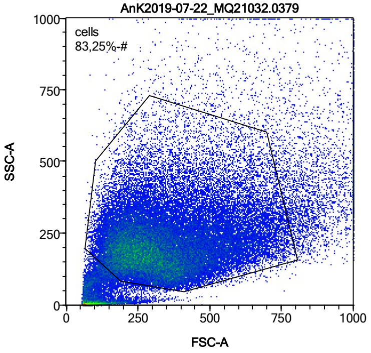


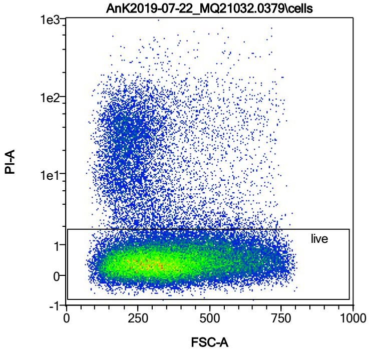


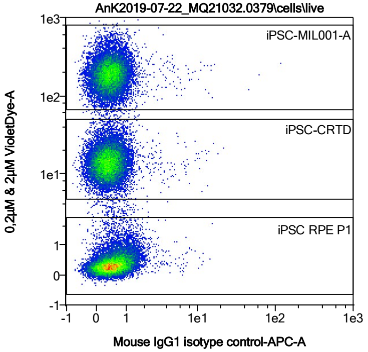


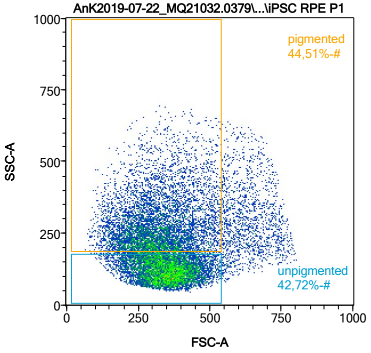


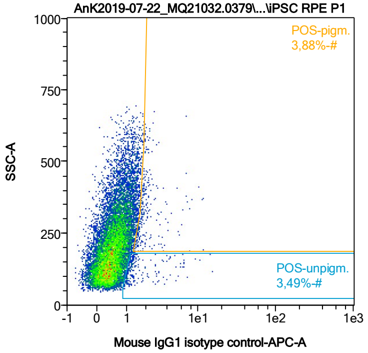


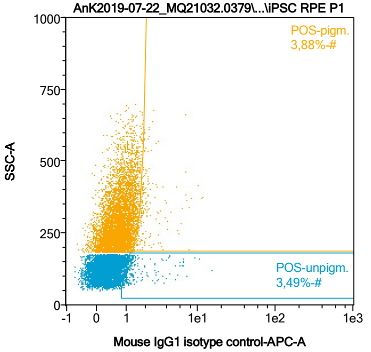


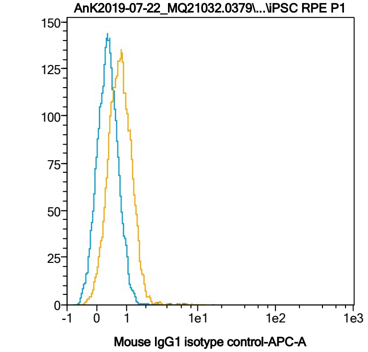


| Name | %-# | Mouse IgG1 isotype control-APC-A Mean | Mouse IgG1 isotype control-APC-A Median |
|---|---|---|---|
| POS-unpigm. | 3,49 | 1,82 | 1,21 |
| POS-pigm. | 3,88 | 2,31 | 1,86 |

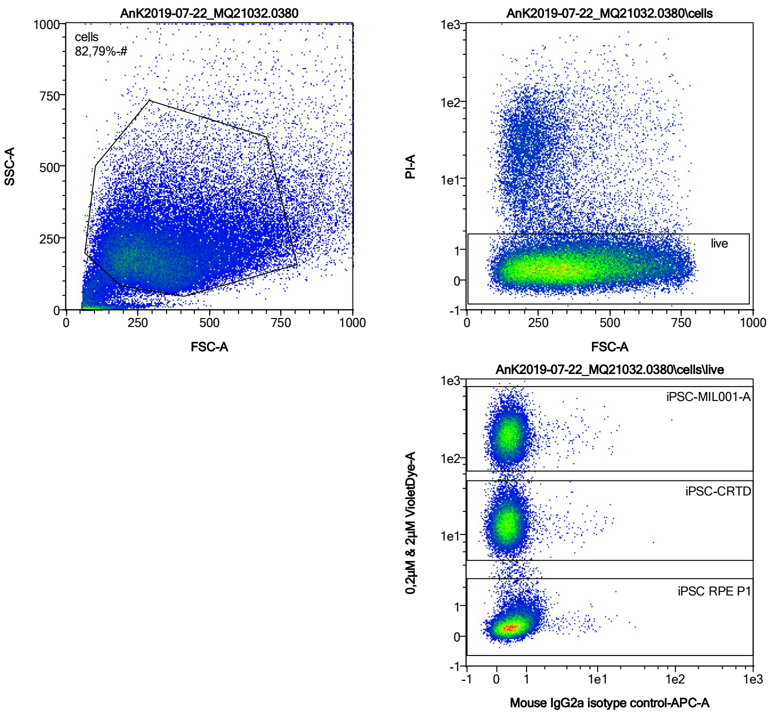


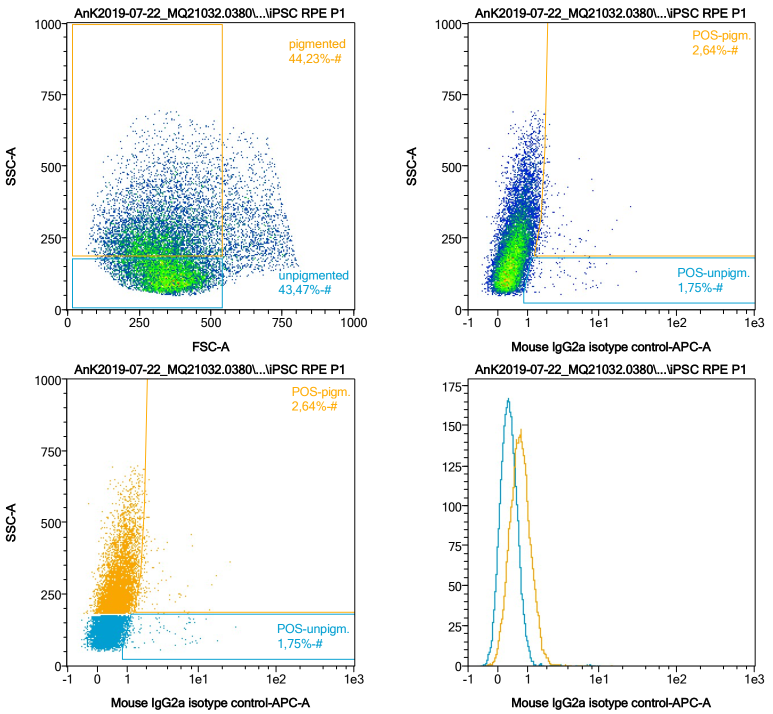


| Name | %-# | Mouse IgG2a isotype control-APC-A Mean | Mouse IgG2a isotype control-APC-A Median |
|---|---|---|---|
| POS-unpigm. | 1,75 | 3,22 | 1,31 |
| POS-pigm. | 2,64 | 2,71 | 1,89 |

Date:2024-06-14
Operator: Sebastian Knoebel
Version:2.13.0+54.g95abf0f4

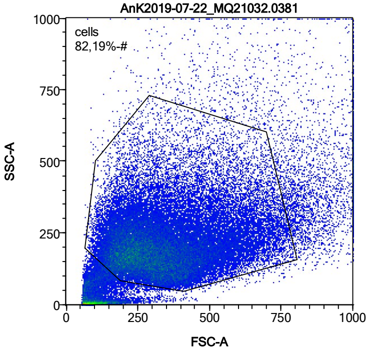


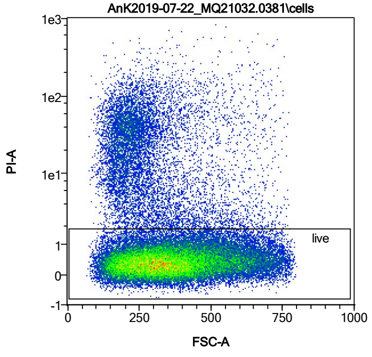


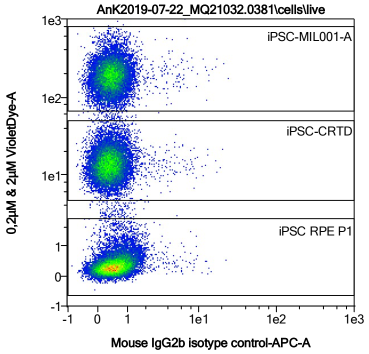


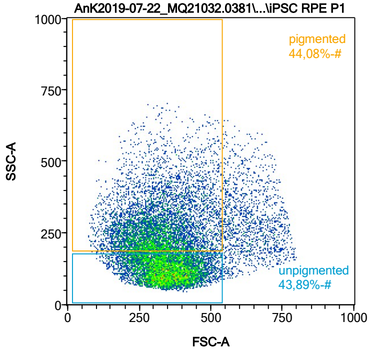


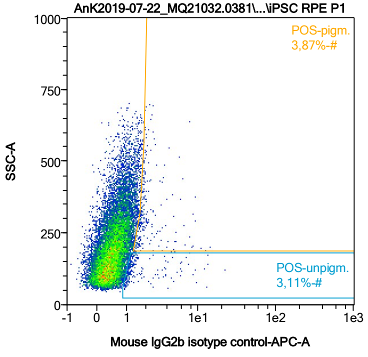


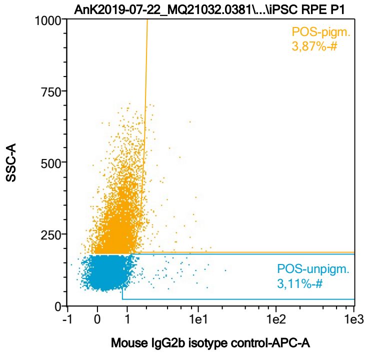


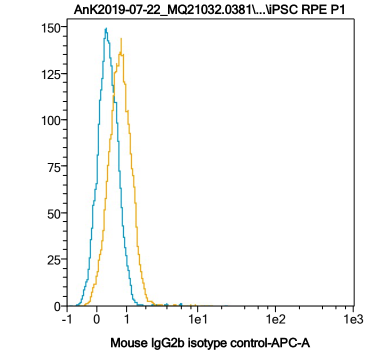


| Name | %-# | Mouse IgG2b isotype control-APC-A Mean | Mouse IgG2b isotype control-APC-A Median |
|---|---|---|---|
| POS-unpigm. | 3,11 | 2,02 | 1,17 |
| POS-pigm. | 3,87 | 2,33 | 1,81 |

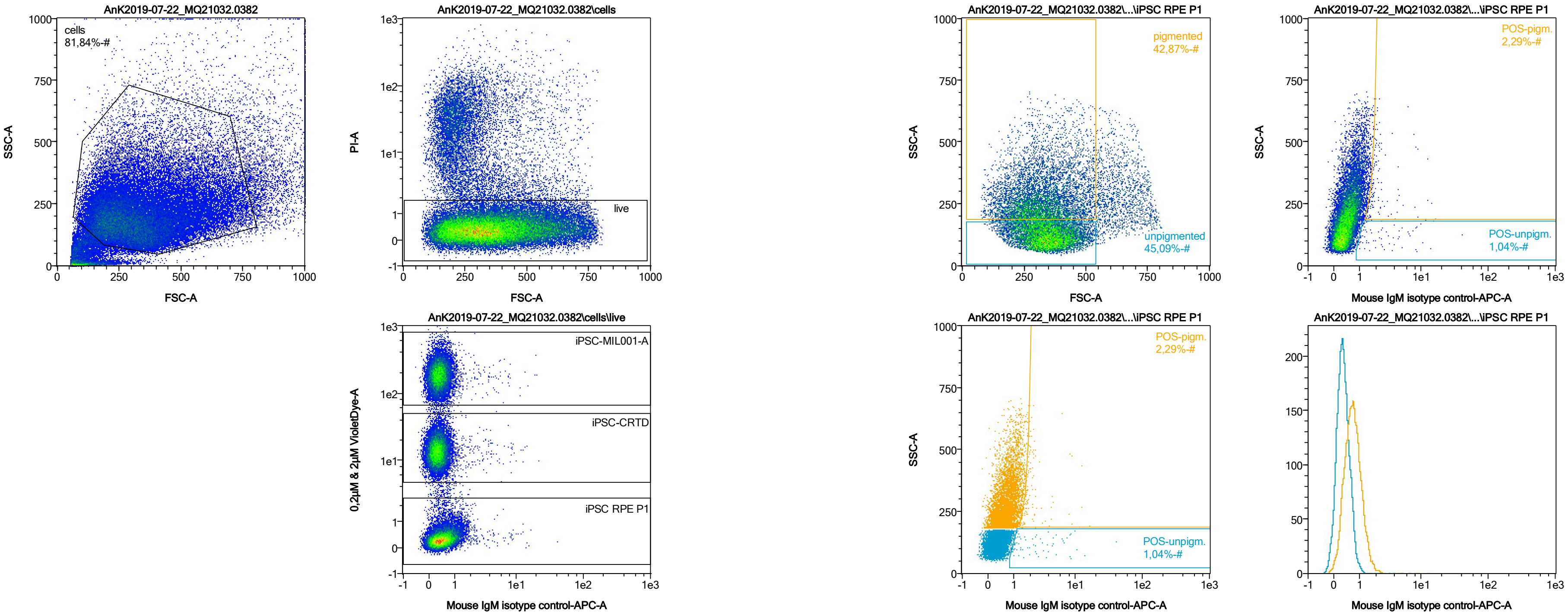


| Name | %-# | Mouse IgM isotype control-APC-A Mean | Mouse IgM isotype control-APC-A Median |
|---|---|---|---|
| POS-unpigm. | 1,04 | 3,67 | 1,94 |
| POS-pigm. | 2,29 | 2,48 | 1,81 |

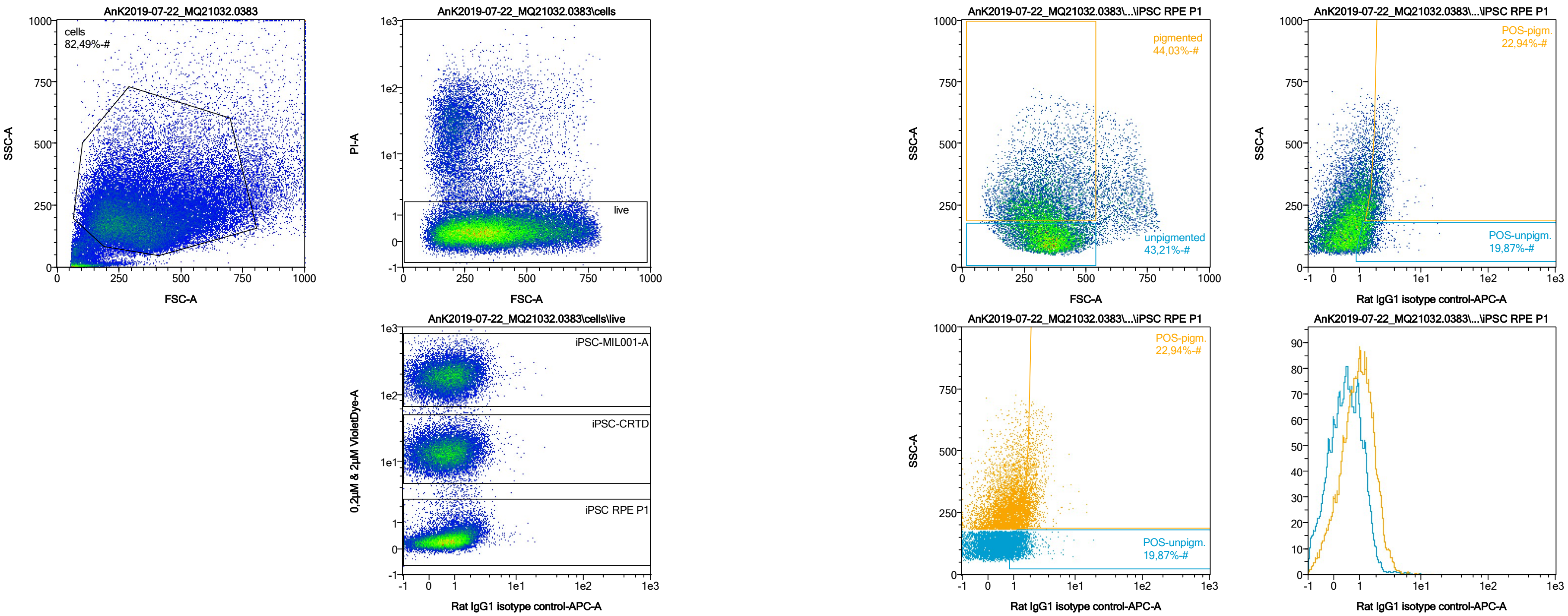


| Name | %-# | Rat IgG1 isotype control-APC-A Mean | Rat IgG1 isotype control-APC-A Median |
|---|---|---|---|
| POS-unpigm. | 19,87 | 1,49 | 1,33 |
| POS-pigm. | 22,94 | 2,15 | 1,95 |

Date:2024-06-14
Operator: Sebastian Knoebel
Version:2.13.0+54.g95abf0f4

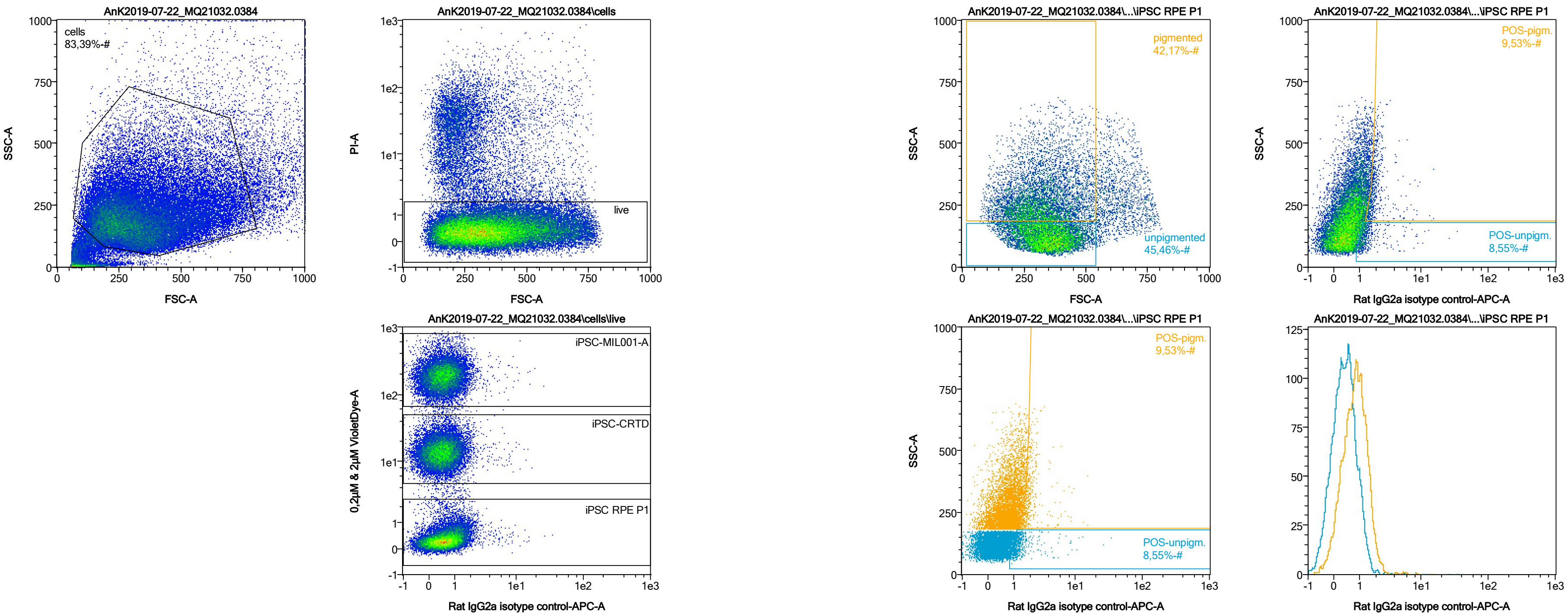


| Name | %-# | Rat IgG2a isotype control-APC-A Mean | Rat IgG2a isotype control-APC-A Median |
|---|---|---|---|
| POS-unpigm. | 8,55 | 1,60 | 1,21 |
| POS-pigm. | 9,53 | 2,11 | 1,80 |

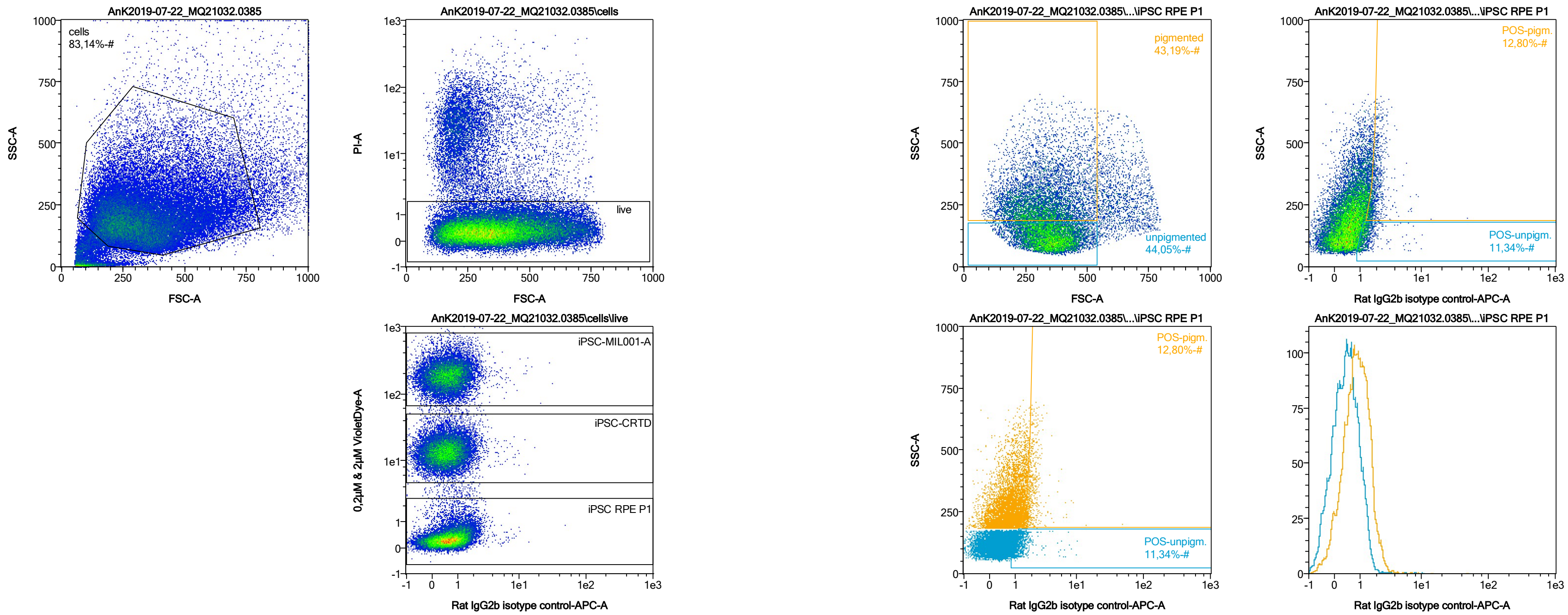


| Name | %-# | Rat IgG2b isotype control-APC-A Mean | Rat IgG2b isotype control-APC-A Median |
|---|---|---|---|
| POS-unpigm. | 11,34 | 1,39 | 1,22 |
| POS-pigm. | 12,80 | 2,03 | 1,81 |

Date:2024-06-14

Operator: Sebastian Knoebel

Version:2.13.0+54.g95abf0f4

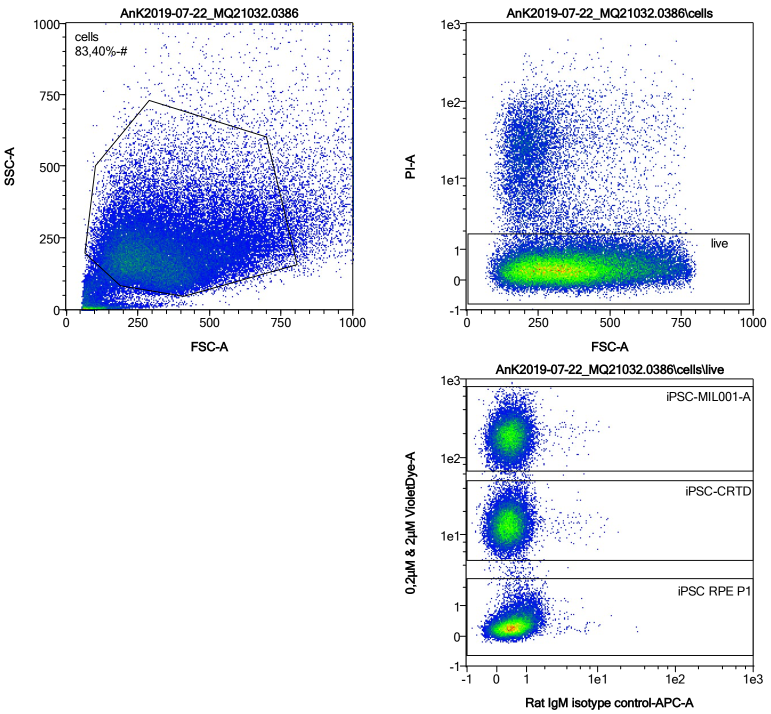


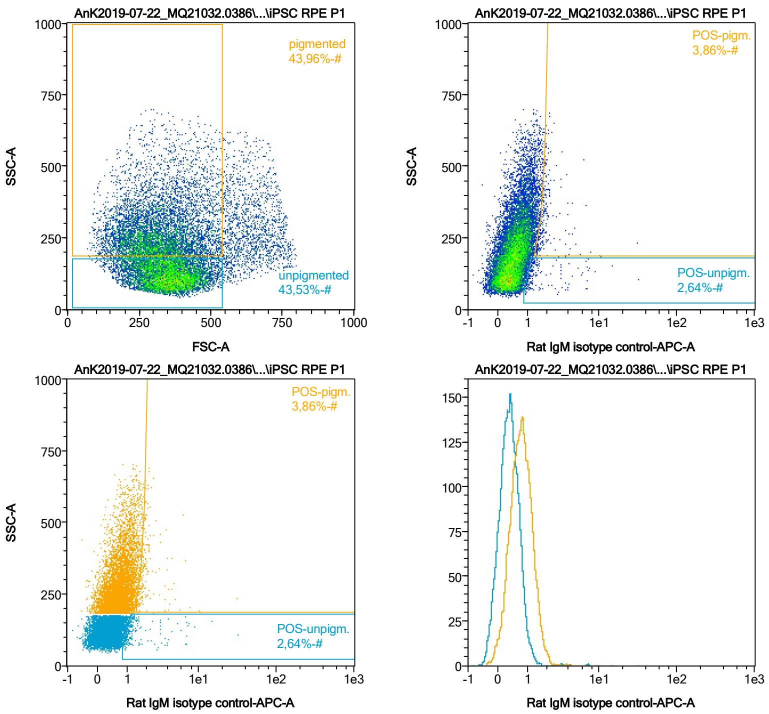


| Name | %-# | Rat IgM isotype control-APC-A Mean | Rat IgM isotype control-APC-A Median |
|---|---|---|---|
| POS-unpigm. | 2,64 | 2,04 | 1,19 |
| POS-pigm. | 3,86 | 2,21 | 1,79 |

Date:2024-06-14

Operator: Sebastian Knoebel

Version:2.13.0+54.g95abf0f4

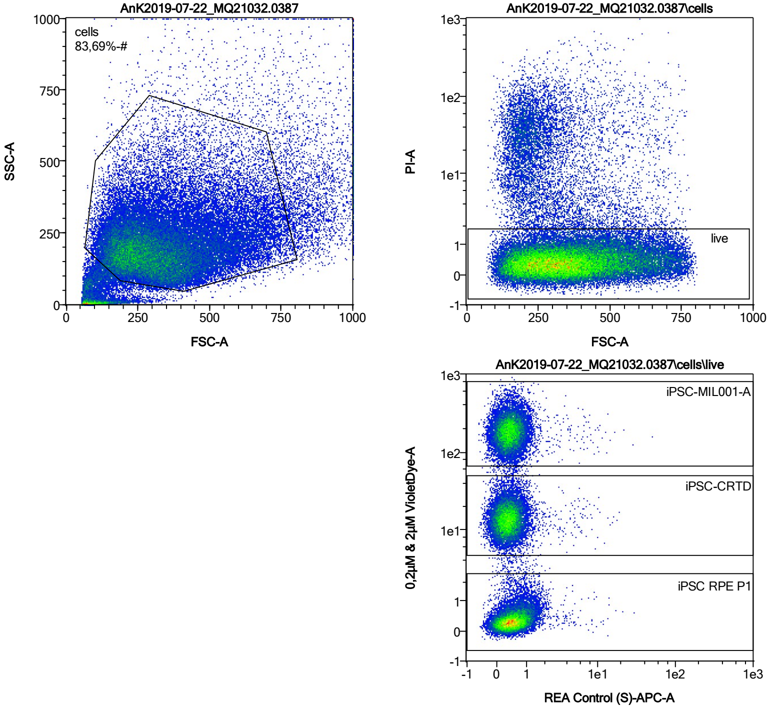


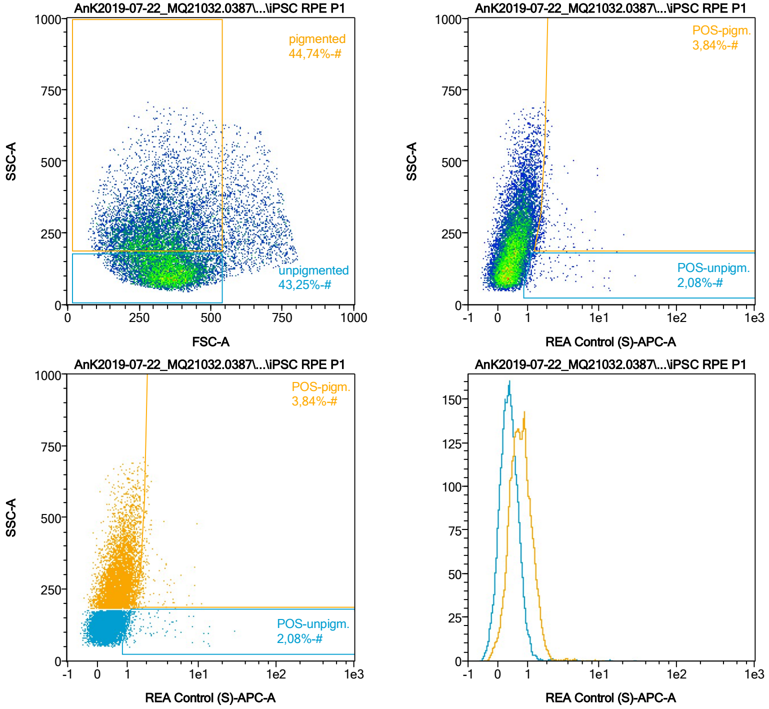


| Name | %-# | REA Control (S)-APC-A Mean | REA Control (S)-APC-A Median |
|---|---|---|---|
| POS-unpigm. | 2,08 | 2,65 | 1,29 |
| POS-pigm. | 3,84 | 2,19 | 1,79 |
