## Supplementary material for "Extensive monolayer formation after transplantation depends on a RPE subpopulation derived from human iPSCs": Document S1

### Document S1. Figures S1-S5.

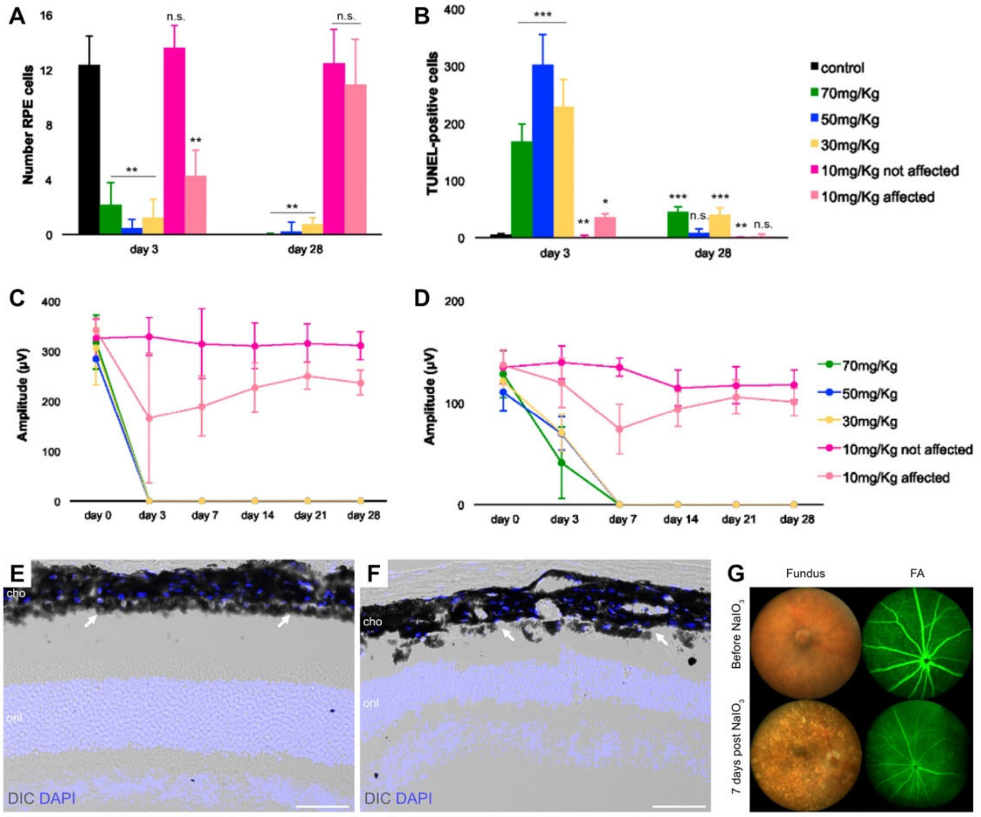


Figure S1: Injection of 30 mg/kg bodyweight $NaIO_3$ suffices to cause RPE degeneration and loss of visual function.

(A) While a low dosage (10 mg/kg bodyweight (bw) $NaIO_3$) had variable effects on RPE cell number, an injection of the intermediate dosage of 30 mg/kg bw $NaIO_3$ was sufficient to cause RPE degeneration to a similar degree as high dosages (50 and 70 mg/kg bw $NaIO_3$) at day 3 and day 28. Control: n=4; 70mg/Kg - day 3: n=14, day 28: n=7; 50mg/Kg - day 3: n=5, day 28: n=5; 30mg/Kg - day 3: n=6, day 28: n=7; 10mg/Kg not affected - day 3: n=4, day 28: n=5; 10mg/Kg affected - day 3: n=1, day 28: n=4. n: individual mice.

(B) In intermediate and high dosage groups, RPE cell death occurred quickly after $NaIO_3$ injection (day 3), and was strongly reduced at 4 weeks after injection (day 28). n: see A.

(C, D) Visual function of both, rods and cones, was compromised after injection of intermediate and high $NaIO_3$ dosages, as shown by the amplitudes of the scotopic (C, 0.003 cd*s*m-2) and photopic (D, 10 cd*s*m-2) b-waves over time. 70mg/Kg - day 0: n=14, day 3: n=14, day 7: n=14, day 14: n=14, day 21: n=11, day 28: n=7; 50mg/Kg - day 0: n=29, day 3: n=5, day 7: n=5, day 14: n=5, day 21: n=5, day 28: n=5; 30mg/Kg - day 0: n=7, day 3: n=11, day 7: n=8, day 14: n=3, day 21: n=8, day 28: n=7; 10mg/Kg - day 0: n=14, day 3: n=14, day 7: n=9, day 14: n=9, day 21: n=9, day 28: n=9. n: individual mice.

(E, F) DAPI staining and DIC imaging at 7 days after injection shows a normal RPE (arrows) and neuroretinal phenotype in the control group (E, saline injection). The RPE in animals injected with 30 mg/kg bw $NaIO_3$ (F) is no longer a continuous monolayer, but patchy and unorganized (arrows). n = 4 biological replicates.

(G) Injection of 30 mg/kg bw $NaIO_3$-injection causes the development of highly reflective speckles in the previously homogenous fundus image. Fluorescein angiography (FA) further shows the change in bloodretina barrier permeability as a result of the $NaIO_3$-injection, with the fluorescein-bright inner plexus vessels clearly standing out from the dark retina initially. At 7 days post $NaIO_3$ injection, the contrast is strongly reduced due to fluorescein leakage through the destroyed RPE layer. n = 4 biological replicates.

Scale bars: 50 µm (E,F). cho, choroid; onl, outer nuclear layer; FA, fluorescein angiography. Data are represented as mean ± SEM.

Statistics: *$p < 0.05$, **$p < 0.01$, ***$p<0.001$ by one-way ANOVA (A-D).

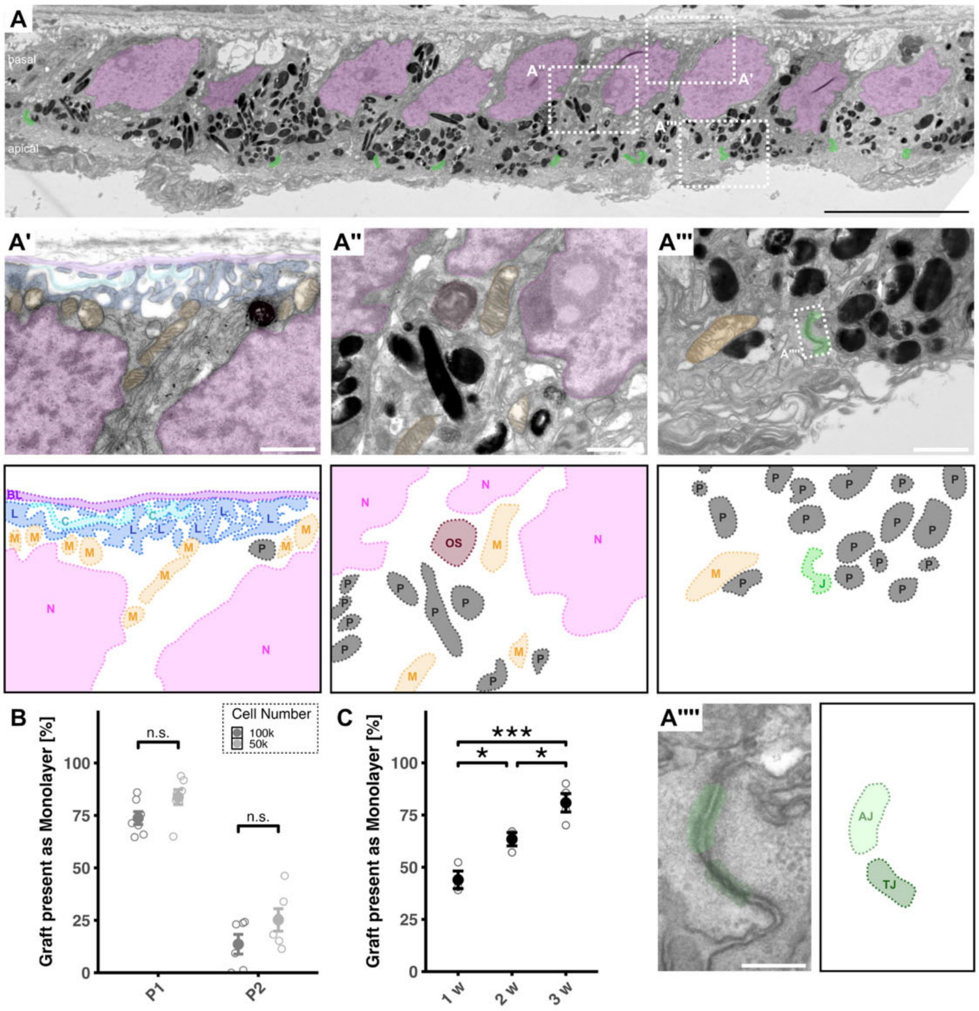


Figure S2: RPE-derived monolayer formation and characterisation

(A) Transmission electron microcopy (TEM) image presented in figure 2E with higher magnifications showing a basal (A’), an intermediate (A’’) and an apical (A’’’) portion of the cells with RPE-characteristic features. Apical junctions are further highlighted per green overlay in A and one junctional complex from A’’’ is magnified in A’’’’. TEM images A’-A’’’’ are depicted with coloured overlays, with the labelled overlay additionally positioned next to the respective image for better visibility. n = 3 biological replicates.

(B) Monolayer percentage data presented in figure 2G, replotted to show differences between eyes having obtained 50,000 or 100,000 cells. While the difference is not statistically significant, transplantation of 50,000 cells tends to generate a higher monolayer percentage than transplantation of 100,000 cells. P1 100k: n=7; P1 50k: n=7; P2 100k: n=6; P2 50k: n=6. All datapoints within one group represent biological replicates.

(C) Monolayer percentage increases over time. Here, empty points represent graft quantification of a single central section per eye, filled points their average. n=3, 3, and 4 biological replicates in groups 1w, 2,w, and 3w, respectively.

Scale bars: 10 µm (A), 1 µm (A’-A’’’), 200 nm (A’’’’). Data are represented as mean ± SEM. AJ, adherens junctions; BL, basal lamina; C, collagen; J, junctional complexes; L, basal labyrinth; M, mitochondrium; N, nucleus; OS, phagocytosed outer segment; P, pigment granule; TJ, tight junctions; w, weeks.

Statistics: $p < 0.05$, **$p < 0.01$, ***$p<0.001$ by two-sided Student’s t-test (B) and one-way ANOVA with Tukey’s post hoc test (C).

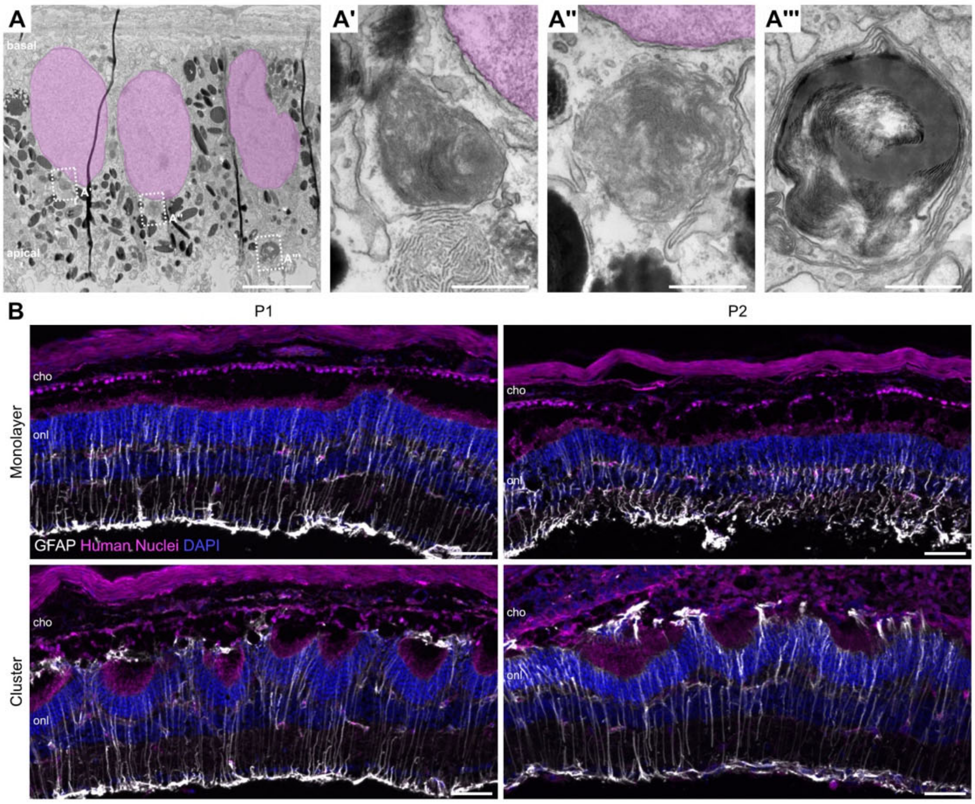


Figure S3: Supplementary data for Figure 3.

(A) Transmission electron microcopy images of RPE-derived monolayers show outer segment particles within transplanted cells. A'-A''' show boxed areas in A magnified. n = 3 biological replicates.

(B) GFAP staining found throughout all neuroretinal layers shows reactive Müller Glia irrespective of RPE passage transplanted or graft cell conformation as monolayer or cluster. n = 5 biological replicates.

Scale bars: 50 µm (B), 5 µm (A) and 500 nm (A'-A'''). cho, choroid; onl, outer nuclear layer.

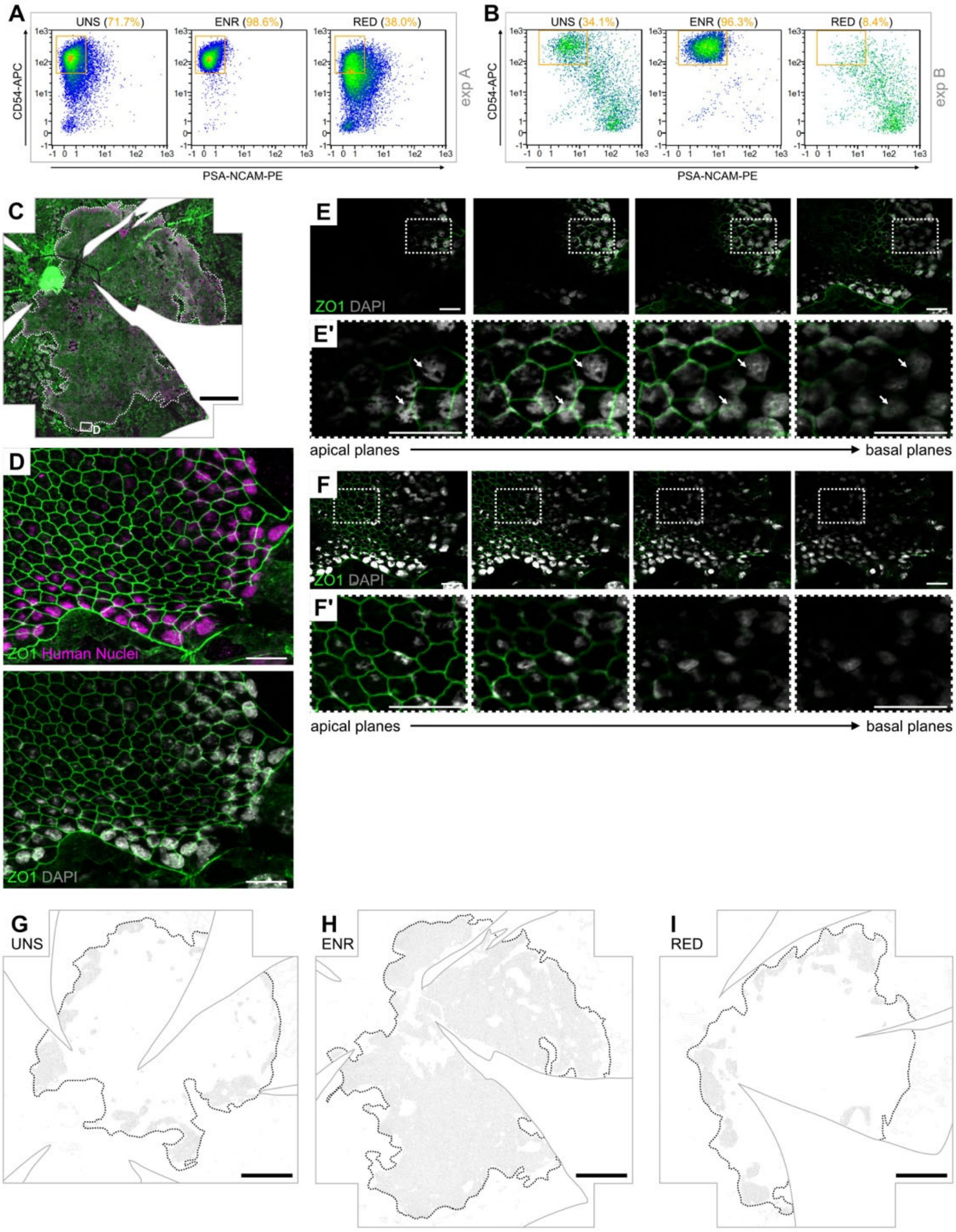


Figure S4: Supplementary data for Figure 5.

(A, B) Flow cytometry re-analysis of unsorted (UNS), CD54$^{+}$/PSA-NCAM$^{-}$-enriched (ENR) and -reduced (RED) fractions before transplantation in experiment A (exp A, see also figure 5I) and experiment B (exp B, see also figure 5J). Orange numbers show purity of the target cell population in each fraction, with a high percentage of CD54$^{+}$/PSA-NCAM$^{-}$ cells in exp A (71.7%) and a lower percentage of CD54$^{+}$/PSA-NCAM$^{-}$ cells in exp B (34.1%).

(C) Flatmount of the ENR graft as presented in figure 5B, here with an additional region framed and magnified in D-F’.

(D) Immunohistochemistry shows higher visibility of the Human Nuclei and DAPI staining towards the edge of the human graft.

(E, F) Single z-planes of the maximum intensity projection in D, with E' and F' showing the boxed regions in E and F enlarged, highlighting the obstruction of nucleus visibility through the apically located pigment (arrows). DAPI brightness is increased in F and F' for better visibility.

(G-I) REShAPE cell outlines of the flatmounted eyecups shown in figures 5A-C, with human graft areas denoted by dotted lines. Eyes received transplantations of cell suspensions that were either unsorted (G), enriched (H), or reduced (I) in CD54$^{+}$/PSA-NCAM$^{-}$ cells.

Scale bars: 500 µm (C, G-I), 20 µm (D-F').

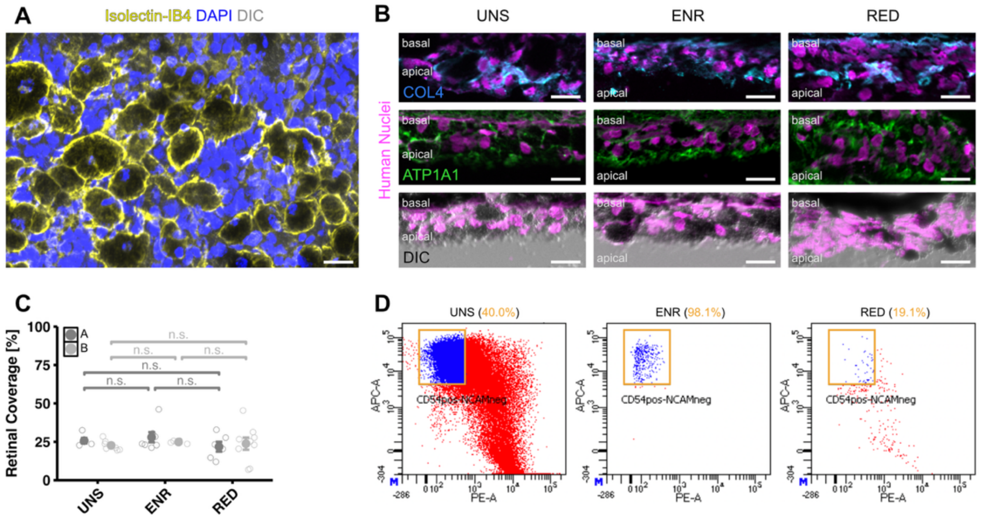


Figure S5: Supplementary data for Figure 5, continued.

(A) Pigmented, spherical clumps likely represent macrophages as they are larger than the average RPE cells, contain a single nucleus and stain positive for isolectin $IB_4$. Here, pigmented clumps are shown stained on top of the outer nuclear layer of a retinal flatmount opposing an unsorted human RPE graft. n = 3 biological replicates.

(B) Clusters formed by all graft fractions lack polarization, as basal marker COL4 and the apical marker ATP1A1 are expressed chaotically throughout the cluster. Similarly, the pigment (dark in DIC images) is not confined to the apical cytosol but mostly situated throughout the cluster. n > 4 biological replicates.

(C) Quantification of retinal coverage after transplantation of RPE cells from experiments A and B (exp A and exp B, representing the experiments depicted in figures 5I and 5J, respectively) shows a similar coverage in all fractions, irrespective of $CD54^+$/PSA-$NCAM^-$ cell purity (exp A: n=5, 7, and 6 biological replicates in groups UNS, ENR and RED, respectively. exp B: n=7, 4, and 9 biological replicates in groups UNS, ENR and RED, respectively).

(D) Reanalysis of the unsorted (UNS), $CD54^+$/PSA-$NCAM^-$-enriched (ENR) and -reduced (RED) fractions of RPE sorted with an Aria III flow cytometer. Orange boxes show the target cell sorting gate and orange numbers show purity of the target cell population per fraction.

Scale bars: 20 µm (A, B).

Statistics: $p < 0.05$, **$p < 0.01$, ***$p<0.001$ by Kruskal-Wallis test (exp. A) and one-way ANOVA (exp. B) (C).
